# Metabolic features of mouse and human retinas: rods vs. cones, macula vs. periphery, retina vs. RPE

**DOI:** 10.1101/2020.07.10.196295

**Authors:** Bo Li, Ting Zhang, Wei Liu, Yekai Wang, Rong Xu, Shaoxue Zeng, Rui Zhang, Siyan Zhu, Mark C Gillies, Ling Zhu, Jianhai Du

**Author notes:** Corresponding Author: Jianhai Du, One Medical Center Dr, PO Box 9193, WVU Eye Institute, Morgantown, WV 26505. These authors contributed equally to this work (Bo Li, Ting Zhang).

## Abstract

Photoreceptors, especially cones, which are enriched in the human macula, have high energy demands, making them vulnerable to metabolic stress. Metabolic dysfunction of photoreceptors and their supporting retinal pigment epithelium (RPE) is an important underlying cause of degenerative retinal diseases. However, how cones and the macula support their exorbitant metabolic demand and communicate with RPE is unclear. By profiling metabolite uptake and release and analyzing metabolic genes, we have found cone-rich retinas and human macula share specific metabolic features with upregulated pathways in pyruvate metabolism, mitochondrial TCA cycle and lipid synthesis. Human neural retina and RPE have distinct but complementary metabolic features. Retinal metabolism centers on NADH production and neurotransmitter biosynthesis. The retina needs aspartate to sustain its aerobic glycolysis and mitochondrial metabolism. RPE metabolism is directed toward NADPH production and biosynthesis of acetyl-rich metabolites, serine and others. RPE consumes multiple nutrients, including proline, to produce metabolites for the retina.

## Introduction

Retinal photoreceptors need an enormous supply of ATP, reducing equivalents and other metabolites to maintain visual transduction and renew their daily-shed outer segments ^1, 2, 3^. Mutations of metabolic genes or defects in metabolic processes have a role in many common degenerative retinal diseases such as inherited retinal degeneration, diabetic retinopathy and age-related macular degeneration (AMD)^3, 4, 5^.

Like tumors, retinas are known for aerobic glycolysis, also called the Warburg effect ^1^. Aerobic glycolysis can produce large amounts of lactate from glucose, even in the presence of O_2_. It is estimated that 80-96% of glucose in isolated retinas is converted into lactate and only a small fraction is oxidized in the mitochondria ^1^. Recent studies confirm that mammalian retinas strongly express enzymes associated with the Warburg effect such as the M2 isoform of pyruvate kinase, hexokinase 2, and lactate dehydrogenase A ^6, 7, 8, 9, 10^. However, retinas are also densely packed with mitochondria which consume oxygen rapidly ^5, 11^. Since glucose is the putative primary nutrient for the retina, it remains unclear how the retina supports its active mitochondrial metabolism while converting most of its glucose to lactate.

Rods and cones are the two major types of photoreceptors in the retina. Rods are responsible for dimly lit vision while cones are for high acuity, central vision and color vision. Cones are highly enriched in the center of the retina, called the macula, in humans and some birds. Cone photoreceptor degeneration is the ultimate cause of clinically significant vision loss in conditions such as AMD ^5, 11^. Cones expend more energy than rods ^12, 13^, but we still know very little about their metabolism compared to rods. The study of cone metabolism is challenging, since cones account for only 2-3% of photoreceptors in mice and rodents do not have the macula. The recent development of all-cone mice by deletion of neural retina leucine zipper (Nrl), a gene required for rod development ^14, 15^, and cone-rich human retinal organoids ^16^ facilitate cone research. To investigate cone photoreceptor metabolism, we used all-cone mice, cone-rich retinal organoids and human macula from post-mortem donors in this study.

Photoreceptors rely on metabolic support from their neighboring retinal pigment epithelium (RPE). Disruption of RPE metabolism contributes to photoreceptor degeneration ^4, 17^. Inhibition of RPE mitochondrial metabolism is sufficient to cause AMD-like retinal degeneration in mice ^18, 19, 20^. Conversely, impairment of photoreceptor metabolism also influences RPE health ^21^. Accumulating evidence supports the concept that photoreceptors and RPE form an inter-dependent metabolic ecosystem ^21, 22, 23, 24, 25^. Photoreceptors shed lipid-rich outer segment discs and export lactate due to the Warburg effect; the RPE converts the outer segment discs into ketone bodies to fuel photoreceptors and it utilizes lactate to preserve glucose for photoreceptors^23, 26^.

We have also reported that reductive carboxylation (reversed TCA cycle) is a very active metabolic pathway in RPE, which it prefers to fuel with proline ^27, 28^. Proline-derived intermediates, including citrate, glutamate and aspartate, can be exported by RPE to fuel retinal mitochondria ^29^. However, most of these studies come from cultured RPE cells and mice. Native human RPE are more dense and multinucleated in the macula than in the periphery ^30, 31^. The macular RPE is where AMD starts, but the metabolic differences between the RPE of the macula and the peripheral retina, and their symbiotic relations with the corresponding neural retinal regions, are still unknown.

We used a targeted metabolomics approach in this study to analyze consumed and released metabolites in the media from cone-rich human retinal organoids, rod-dominant vs. all-cone mouse retinas, human macular vs. peripheral neural retinas and human macular vs. peripheral RPE/choroid. By integrating our results with the analysis of the expression of metabolic genes, we have identified key metabolic features of retinal organoid development, rods vs. cones, macula vs. periphery and neural retina vs. RPE. These features may elucidate the different metabolic needs of retinal mitochondria, cones and human macula and reveal the metabolic communications between human retina and RPE at the macula and peripheral retina.

## Results

### Metabolite consumption and metabolic gene expression in human cone-rich retinal organoids

Cell differentiation in retinal organoids largely recapitulated transcriptomic and electrophysiological properties of retinogenesis *in vivo* ^16^. Retinal organoids at D56 were mainly composed of retinal progenitor cells and early differentiated retinal cells, while retinal organoids at D296 were mostly composed of maturing cones, rods, and Müller glial cells ^16^. To study the nutrient needs during the development of the human retina, we collected spent media 48 hours (h) after the medium change from human retinal organoids on 56 days (D56) and 296 days (D296) (**Fig 1a**). Blank media without cells in the same culture plate for 48h served as baseline control. We targeted 219 metabolites covering major nutrients and pathways (**Table S1**) and detected 89 metabolites from these media. To determine metabolite uptake and release, we divided the ion abundance from organoids by those from the control media (See details in the methods). Retinal organoids at both D56 and D296 consumed glucose, cystine and cystathionine (**Fig 1b-c**). Retinal organoids at D56 used L-Alanyl-Glutamine (GlutaMax), hypoxanthine, lysine, hippurate, glycerol and biotin but D296 did not (**Fig 1b-d**). Surprisingly, retinal organoids at D296 consumed 6-8 fold more aspartate and glutamate from the media, but organoids at D56 did not use them at all (**Fig 1b-c**). These results indicate that the consumption of aspartate and glutamate is specific for mature retinal cells. Additionally, retinal organoids at D296 also consumed histamine. Glucose can be metabolized through glycolysis and the mitochondrial TCA cycle into lactate, pyruvate and other mitochondrial intermediates. GlutaMAX could be hydrolyzed into glutamine and alanine. As expected, the products from glucose and GlutaMAX, including lactate, pyruvate, alanine, glutamine, proline, citrate and other metabolites, were produced significantly by D56 organoids (**Fig 1e**). D296 organoids generated similar amounts of lactate and pyruvate as D56 organoids, but they produced ∼11-fold of hypotaurine and 2-fold of acetyl-carnitine (C2-carnitine) (**Fig 1f-g**).

**Figure 1.**
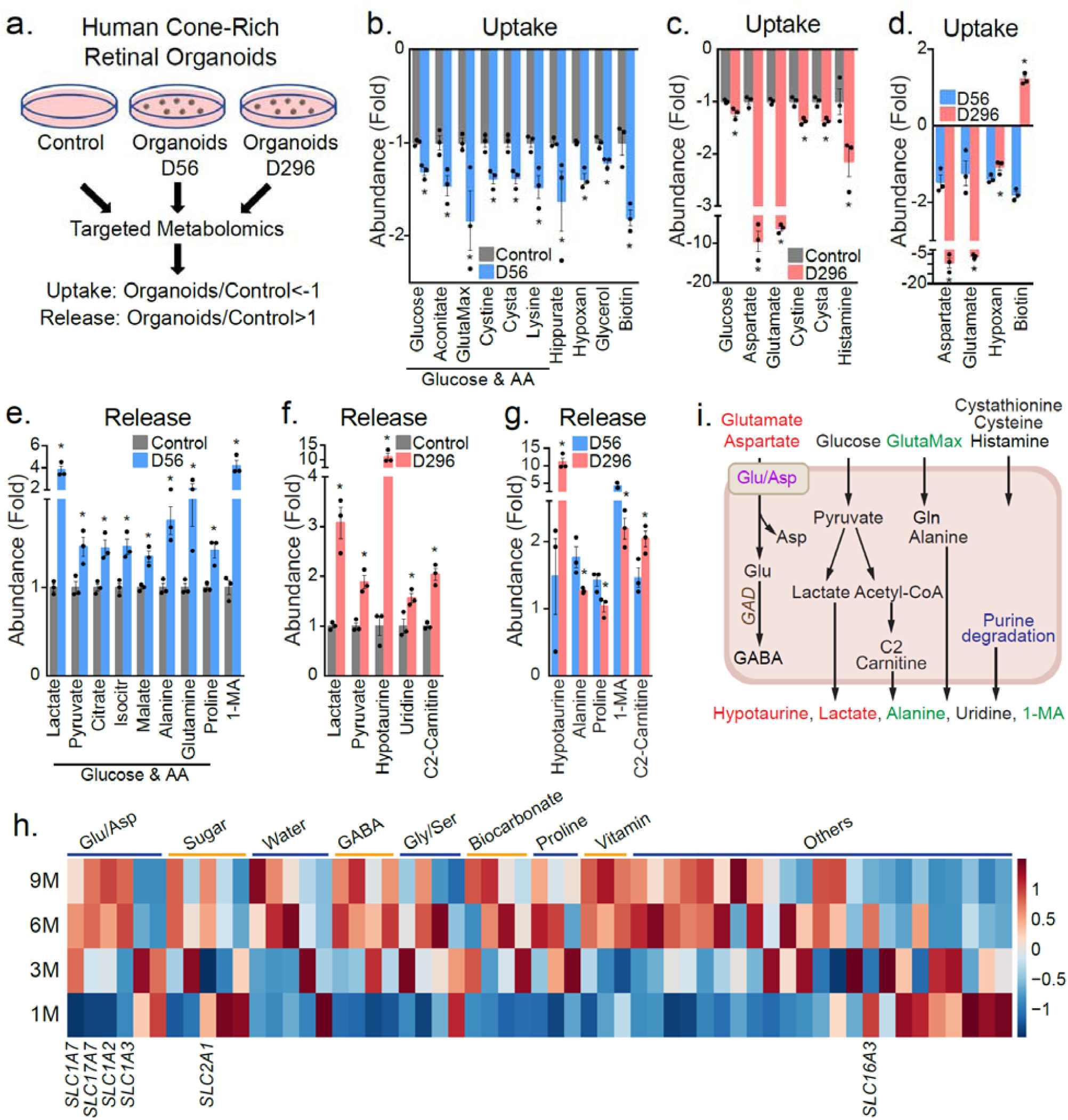
Metabolite consumption and metabolic gene expression in human cone-dominant retinal organoids. **(a)**. A schematic for profiling medium metabolites in retinal organoid culture. Retinal organoids on Day 56 (D56) and D296 were cultured in a medium (See Methods) for 48h. The spent media were analyzed for metabolite consumption or release, and the medium in culture wells without retinal organoids was used as a control. (**b-d**) Metabolite consumption in retinal organoids at D56 and D296. (**e-g**) Metabolite release from organoids at Day 56 and D296. Data were relative ion abundance over the control. N=3, *p<0.05 vs. Control or D56 organoids. (**h)** Heatmap of gene expression of top-ranked small molecule transporters in the human retinal organoids cultured at 1-, 3-, 6- and 9-month stages. FPKM values of bulk RNA sequencing ^16^ was used for plotting. (**i**) A schematic of nutrient uptake and release in retinal organoids. Increased metabolites in mature organoids at D296 are colored in red and increased metabolites in D56 organoids are colored in green.

Small molecule transporters and metabolic enzymes contribute to specific profiles in metabolite uptake and release. To compare the expression of transporters and enzymes, we curated 2,764 classic metabolism genes from KEGG pathways (**Table S2**) and analyzed their expression in the transcriptome of retinal organoids at stages between 1 month and 9 months ^16^. Differentially expressed genes along the time course of human retinal organoids identified previously^16^ were intersected with the genes in metabolic pathways, resulting in 378 metabolic genes that were differentially expressed during the development of retinal organoids. More than half of the differentially expressed genes were ion and small molecule transporters (**Fig S1a, Table S3**). Remarkably, glutamate and aspartate transporters including solute carrier 1A3 (*SLC1A3*), *SLC17A7, SLC1A7* and *SLC1A2* were the most highly enriched of the upregulated small molecule transporters in 9-month organoids (**Fig 1h, Table S3**). These results indicate that mature retinal organoids need more aspartate and glutamate. Other amino acids, bicarbonate, and water transporters were also upregulated in the maturing organoids. Glucose transporter 1 (*SLC2A1*), the major glucose transporter, was expressed similarly during organoid development except for a decrease at 3 months. Monocarboxylate transporter (MCT) is the major lactate/pyruvate transporter in the retina and *MCT4* or *SLC16A3* was expressed similarly between 3-month and 9-month organoids (**Fig 1h**). These results were consistent with our metabolite consumption data on glucose and lactate. Glutamate decarboxylase 1 and 2 (*GAD1/2*) convert glutamate into 4-aminobutyrate (GABA) (**Fig 1i**). They were highly upregulated in D296 organoids (**Fig S1b**), providing further evidence of the need for glutamate. GABA transporters were also increased in the mature organoids (**Table S3**). Genes in cysteine, cystathionine, glutathione and histidine/histamine metabolism were highly enriched in the mature organoids (**Fig S1b**). This may implicate in the changed consumption of cysteine derivatives such as cystine, hypotaurine and histamine. Apart from aspartate and glutamate metabolism, most upregulated genes in mature retinal organoids were involved in glycan, glycolysis, lipids, ATP-binding cassette (ABC) transporter (lipid transporter) and nucleotide, especially cyclic nucleotides and purine degradation (**Fig S1c-e, Table S3**). Increased purine degradation and recycling may be the cause of the reduced consumption of hypoxanthine that we observed (**Fig 1d, 1i**). To sum up, retinal organoids showed distinctive features of nutrient consumption and developmentally regulated expression of metabolic genes. Mature retinal organoids may import and utilize aspartate and glutamate through their upregulated transporters for neurotransmitter synthesis and energy metabolism (**Fig 1i**).

### Metabolite consumption and metabolic gene expression of rod-dominant and all-cone mouse retinas

Rods comprise ∼97% of photoreceptors in mouse retinas. Knockout (KO) of *Nrl* converts all rods into cone-like photoreceptors ^15^ to make all-cone retinas. We cultured retinas from ∼1-month-old *Nrl*-heterozygous (Het) (rod-dominant similar to wild type [WT]) mice and Nrl KO mice in transwells and analyzed metabolites in the media after culturing for 24h (**Fig 2a**). Compared with the control media, 32 metabolites from Nrl Het and 37 metabolites from Nrl KO media were significantly different (**Fig 2b-h**). Most changed metabolites were in a similar pattern in the Het and KO retinas. Interestingly, aspartate and glutamate were among the most consumed metabolites in both Het and KO retinas, with a decrease of 8-11 fold compared with baseline media (**Fig 2b-c**). Additionally, both Het and KO retinas consumed glucose, amino acids, carnitine and intermediates in mitochondrial metabolism including citrate, succinate and fumarate (**Fig 2b-c**). Like retinal organoids, mouse retinas excreted large amounts of lactate, acetylcarnitine and 1-methyladenosine (**Fig 2e-f**). However, mouse retinas also released nucleosides, other acyl-carnitines, amino acids and derivatives from amino acids such as creatine and creatinine (**Fig 2e-f**). Isopentenyl pyrophosphate (IPP), an intermediate the mevalonate pathway, was not detected in the control media but was significantly increased in the spent media of mouse retinas, especially the *Nrl* KO group (**Fig 2g**). These results indicate that mouse retinas have very active mitochondrial, lipid and nucleotide metabolism.

**Figure 2.**
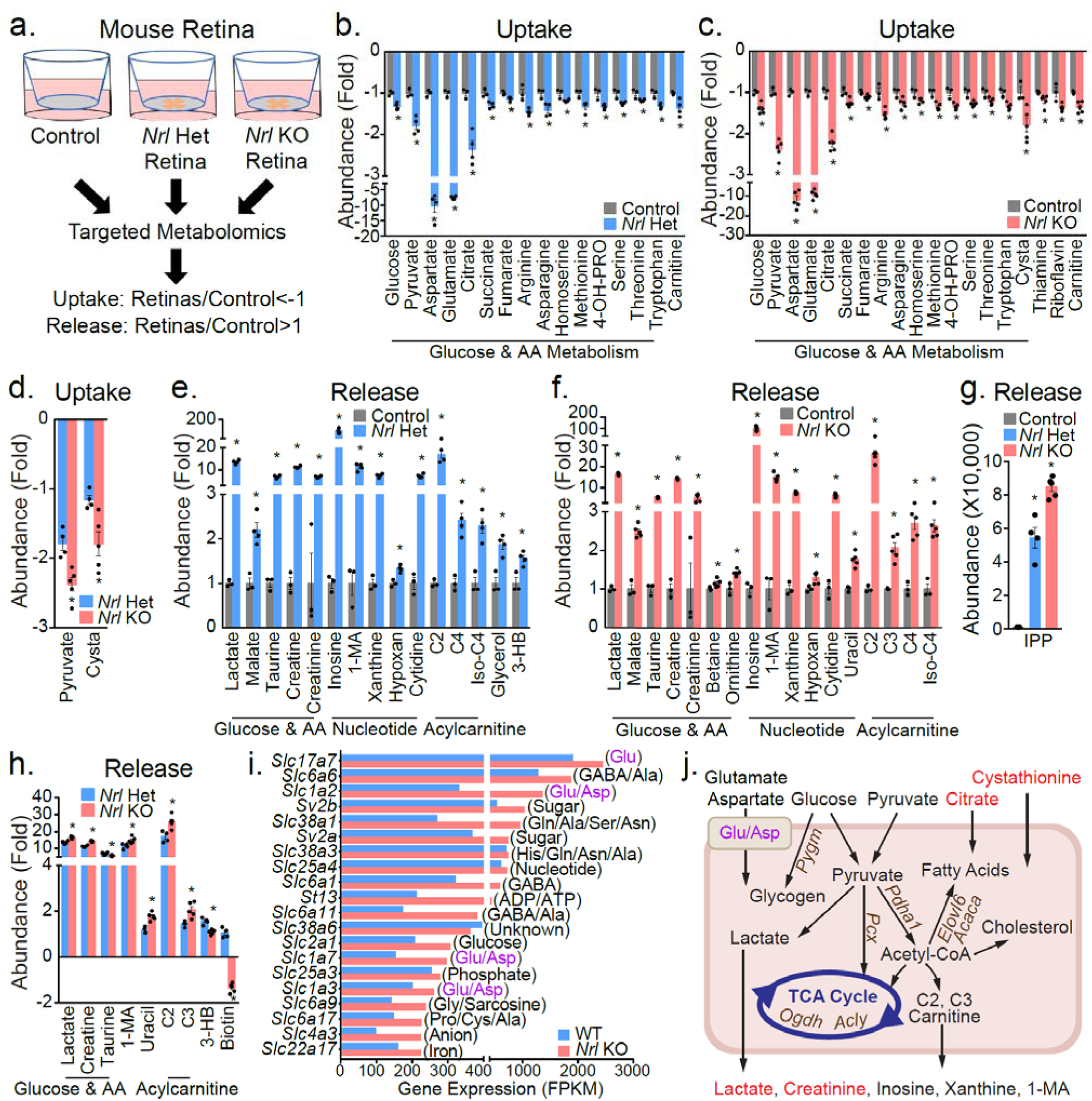
Metabolite consumption and metabolic gene expression in Nrl Het (rod-dominant) and Nrl KO (all-cone) mouse retina. **(a)**. A schematic for profiling medium metabolites in mouse retinal explants. Retinas from Nrl Het and Nrl KO mice were dissected, placed in transwells and maintained in culture for 24h. The culture media without retinas were the baseline control. All media metabolites were analyzed by targeted metabolomics. **(b-d)**. Metabolite consumption from mouse retinal explants of Nrl Het. (**e-h)**. Metabolite release from Nrl Het and KO retinas. Data were relative ion abundance over control or Nrl Het or absolute ion abundance when those metabolites were not detected in the baseline control. N=6, *p<0.05 vs. Control or D56 organoids. (**i)**. Gene expression of the top 20 abundant small molecule transporters in the Nrl KO and Het mouse retinas. (**j**). A schematic of nutrient uptake and release in Nrl KO retinas. Metabolites with enhanced uptake or release in Nrl KO retinas are colored red. Upregulated genes in Nrl KO are colored brown. Glu/Asp represents transporters for glutamate and aspartate.

Twelve metabolites were different between Het and *Nrl* KO retinas (**Fig 2d, 2g-h)**. *Nrl* KO consumed more pyruvate and cystathionine and produced more lactate, acylcarnitines, creatine and 1-methyladenosine (**Fig 2d, 2g-h**). In contrast to retinal organoids, neither *Nrl* Het nor KO retinas excreted hypotaurine; however, mouse retinas exported more taurine, particularly the Het retinas (**Fig 2h**). Ketone bodies are important alternative fuels for neurons. Intriguingly only rod-dominant retinas produced the ketone body, 3-HB (**Fig 2e-h**), suggesting that there might be inter-dependent ketone body utilization between rods and cones.

We then analyzed the expression of metabolic genes from a transcriptome database of WT and *Nrl* KO retinas ^32^. Among the most abundant small molecule transporters ranked by either WT or KO retinas, transporters for glutamate and aspartate were among the most enriched (**Fig 2i, Fig S2**). In particular, Slc17a7 had more abundant transcripts than the other small molecule transporters (**Fig 2i, Fig S2)**. Transporters for glucose, GABA, glutamine, lactate, pyruvate and ketone bodies were also among the top transporters. Compared with WT, 800 metabolic genes were changed, with 401 upregulated in *Nrl* KO retinas (**Fig S3, Table S4**). Ion transport, lipid metabolism, small molecule transport, glycan, amino acid, nucleotide and sugar metabolism were the major changed pathways (**Fig S3**). Key enzymes in glycogen metabolism (*Pygm,Gaa*), pyruvate metabolism (*Pcx, Pdha1,Me1*), TCA cycle (Cs, *Acly, Ogdh, Idh3a*), cysteine/cystathionine metabolism (*Suox, Ado, Cdo1, Gclm*), fatty acid synthesis (*Acaca, Acacb, Oxsm, Elovl6*) were significantly upregulated in *Nrl* KO retinas (**Fig 2j, Table S4**). These data support the nutrient uptake and release results, indicating that cone photoreceptors require more pyruvate and other nutrients for their robust mitochondrial TCA cycle, lipid synthesis, and amino acid metabolism (**Fig 2j**).

### Metabolite consumption and metabolic gene expression of human peripheral and macular retina

To investigate the nutrient utilization of the human macula and peripheral retina, we cultured human neural retinal explants punched from the macular and peripheral regions from the same donor (**Fig 3a**). Human donors had no known retinal diseases and *post mortem* fundus photographs showed no abnormalities (**Fig 3a, Table S5**). After culturing for 24h, media levels of 52 metabolites in the peripheral retina and 51 metabolites in the macula were significantly different (**Fig 3b-h, Fig S4**). Strikingly, aspartate and glutamate were almost completely depleted in spent media from both peripheral retina and macular punches (**Fig 3b-c**). Aspartate and glutamate were supplemented at ∼0.15 mM in all the media of retinal cultures, and they were depleted in mature retinal organoids, mouse retinas and human retinas (**Table S6-8**). These findings demonstrate that retinas need substantial amounts of aspartate and glutamate. As well as glucose, carnitine and amino acids, human retinas also actively consumed vitamins such as pantothenic acid, nicotinamide and biotin (**Fig 3b-d, Fig S4**). β-alanine, which can be metabolized into acetyl-CoA in mitochondria, was also reduced 2-3 fold in the media. Similar to mouse retinas, both macular and peripheral retinas released lactate, nucleosides and acylcarnitines, confirming that human retinas are very active in metabolizing glucose, amino acids and lipids (**Fig 3e-g, Fig S4**). Creatine, creatinine and citrulline increased 3-7 fold in the media with retinal cultures (**Fig 3e-g**). These metabolites, synthesized from arginine, are involved in the synthesis and degradation of the high-energy metabolite phosphocreatine. IPP, guanosine, kynurenine, and N1-methylnicotinamide were not detected in the control media, but significantly increased in spent media from both macular and peripheral neural retinas (**Fig 3g**). These results confirm that the mevalonate pathway, purine catabolism and NAD metabolism are active in the retina.

**Figure 3.**
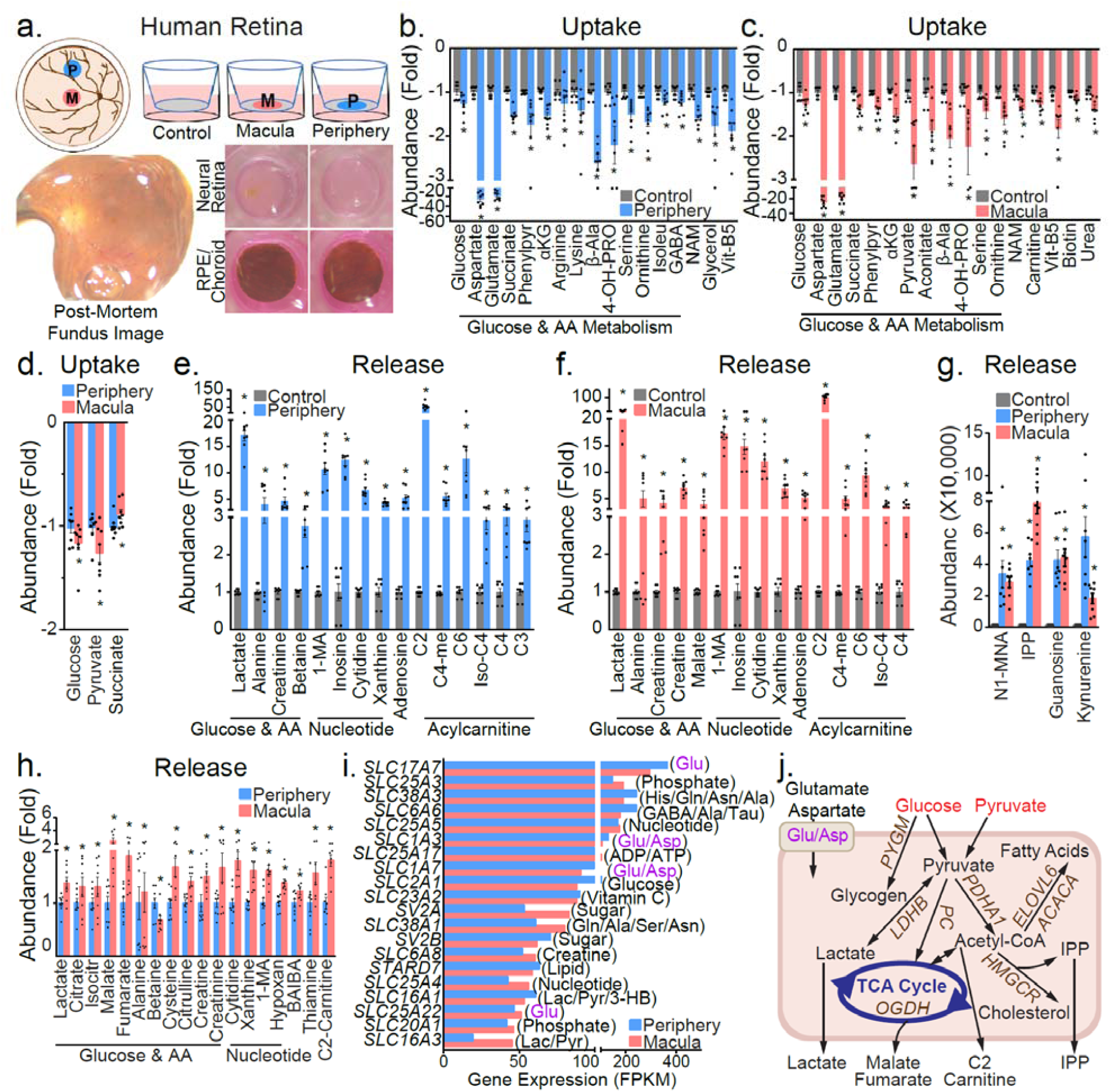
Metabolite consumption and metabolic gene expression in the human macular and peripheral retina. **(a)** Schematic diagram of human macular and peripheral explant culture. Macula and Peripheral punches from neural retinas or their corresponding RPE/choroid punches were cultured on inserts in transwell plates for 24h. Representative fundus images from one donor showed healthy retinas. Representative images of cultured macular and peripheral neural retinas and RPE/choroids in transwell plates. **(b-d)** Metabolite consumption from macular and peripheral neural retinal explant culture. (**e-h**) Metabolite release from the macular and peripheral neural retinal explant culture. Data were relative ion abundance over control or peripheral retinas, or absolute abundance when detected in the baseline control. N=8, *p<0.05 vs. control or peripheral retinas. (**i)** Gene expression of the top 20 abundant small molecule transporters in human peripheral and macular retinas. (**j**) A schematic of nutrient uptake and release in macular retinas. Metabolites with enhanced uptake or release in macular retinas are colored red. Upregulated genes in macular retinas are colored brown. Glu/Asp represents transporters for glutamate and aspartate.

Twenty-five metabolites in the media were significantly different between the macular and the peripheral retina (**Fig 3d, g-h**). In general, the neural macula consumed and released more metabolites than the peripheral retina except for succinate, betaine and kynurenine (**Fig 3d**,**g-h**). Like cone-dominant *Nrl* KO retinas, the neural macula consumed more pyruvate from the media and released more lactate, acetylcarnitine, 1-methyladenosine and creatine into the media. These data suggest that the neural macula has more active energy metabolism than the peripheral retina. In support of this, the neural macula released more nucleosides and several TCA cycle intermediates than the peripheral retina (**Fig 3h**).

To further understand the genetic basis for this differential nutrient consumption, we analyzed the expression of metabolic genes in a human transcriptome database ^33^. As in the mouse retinas, aspartate and glutamate transporters were highly enriched amongst the most abundant small molecular transporters in both the macula and peripheral retina, with SLC17A7 being the most abundant of all (**Fig 3i, Fig S5**). Transporters for glucose, glutamine, taurine, creatine, lactate and pyruvate were also among the most abundant transporters (**Fig 3i, Fig S5**). Metabolic gene analysis found 307 upregulated and 192 downregulated genes in neural macula, compared with peripheral retina (**Table S9**). Similar to mouse *Nrl* KO retinas, the most changed pathways were in ion transport, small molecule transport, lipid, glycan, amino acid, nucleotide and sugar metabolism (**Fig S6**). Changed small molecule transporters were enriched for the transport of sugar, glutathione, glycine, serine, arginine, citrulline and glutamate (**Fig S6**). Glycan, a polysaccharide that conjugates with lipid and protein to make the cell membrane, is synthesized mostly from glucose and fructose. Strikingly, most genes in glycan synthesis were upregulated in the macula, but genes responsible for its degradation were downregulated, indicating the neural macula requires more exogenous glucose to meet the high demand for glycans.

Key enzymes in several major metabolic pathways were significantly upregulated in the macula compared with the peripheral retina such as those responsible for glycogen metabolism (*PYGM, PYGB, GBE1, UGP2*), pyruvate metabolism (*PC, ME2, LDHB, LDHD*), TCA cycle (*OGDH, ACLY, MDH1, ACO1, SDHA*), fatty acid synthesis (*ACACA, ACSS2, FASN, ELOVL6*), mevalonate & cholesterol metabolism (IDI1, *HMGCR, HMGCS1*), ketone body metabolism (*OXCT2, HMGCLL1*) and NAD synthesis (*QPRT, NMNAT2*) (**Table S9, Fig 3j**). These upregulated pathways in human neural macula were similar to all-cone *Nrl* KO retinas. By comparing the changed metabolic genes in *Nrl* KO with the human macula, we found 119 out of 177 genes were changed in the same way, including those in the major metabolic pathways (**Table S10**). These results indicate that all-cone mouse retinas have some metabolic similarities with the cone-enriched human macula. Notably, pyruvate was the only nutrient that was consumed more by both *Nrl* KO retinas and human macula. Pyruvate carboxylase (*PC* in human and *Pcx* in mouse), which utilizes pyruvate for TCA cycle and gluconeogenesis, was also increased in both *Nrl* KO retinas and macular retinas (**Fig 3j**). These data demonstrate that the neural macula meets its higher metabolic requirements by utilizing more glucose and pyruvate for the TCA cycle and synthesis of glycan, glycogen and lipids.

### Metabolite consumption and metabolic gene expression in human peripheral and macular RPE/choroid

The RPE, tightly bound to the choroid, is critical to transport metabolites to and from the retinas. To understand the metabolic communications between retina and RPE, we studied the nutrient uptake and release of punches of human RPE/choroid using the same donor eyes from the neural retinal experiments (**Fig 3a**). The profiles of consumption and release by human RPE/choroid punches from both macula and periphery were similar to those of our previously reported studies of human RPE *in vitro*, ^28^ but quite different from those in human retinas (**Fig 4a-g, Fig S7**). As well as specific nutrients in the media, such as carnitine, nicotinamide, cysteine and proline, human RPE/choroid also consumed nutrients not specified in the medium formula such as N-acetylglycine, erythritol and succinate (**Fig 4a-b**). These metabolites may come from serum and other supplements in the media. In contrast to neural retinas, RPE/choroid did not consume glutamate, and only the macular RPE used a small amount of aspartate (<1.5 folds), confirming that the consumption of glutamate and aspartate is specific to the neural retina. In contrast to human neural retinas, human RPE/choroid consumed, rather than excreted, taurine and betaine (**Fig 4d-g, Fig S7**), suggesting that the neural retina and RPE might exchange these metabolites.

**Figure 4.**
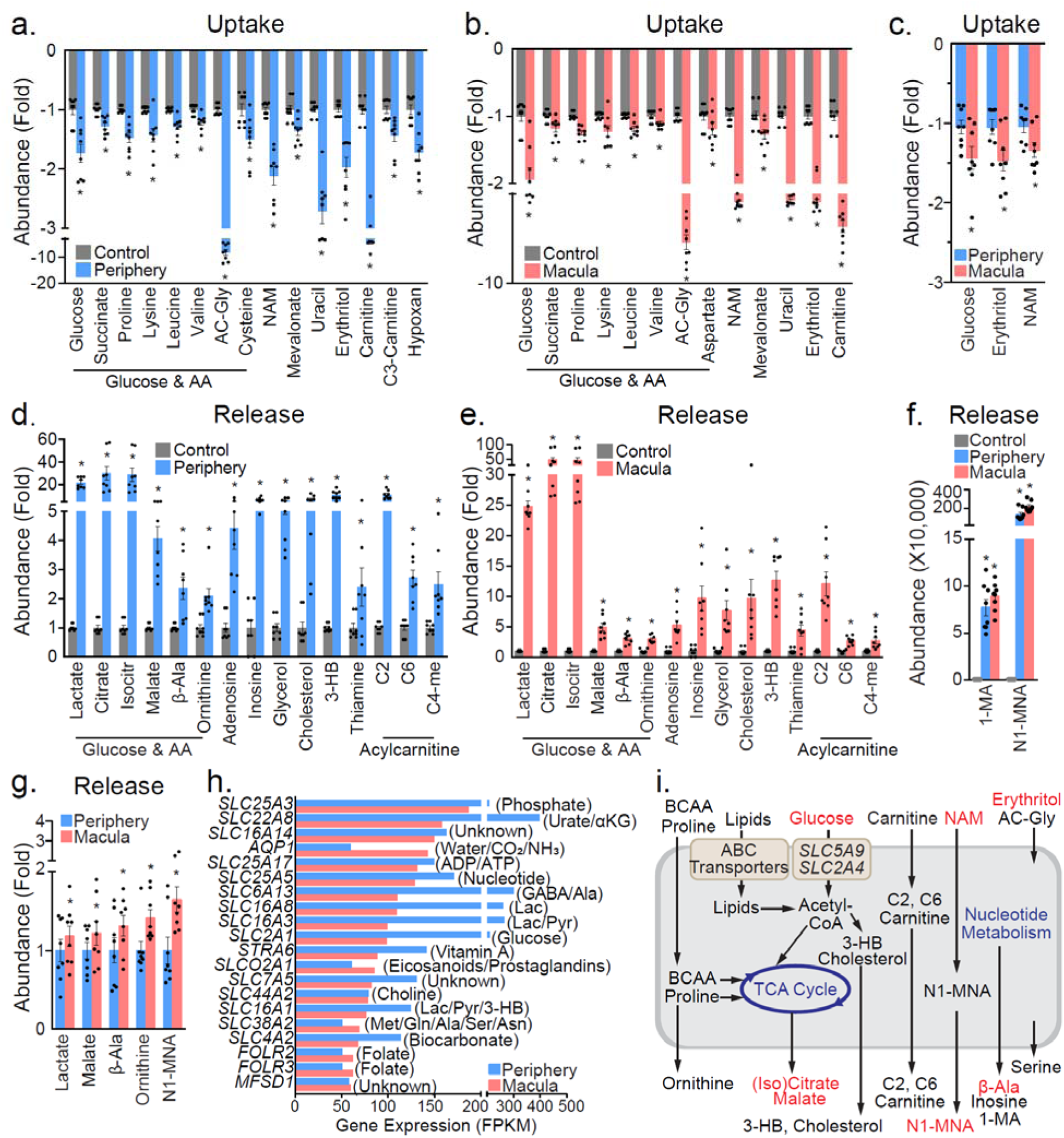
Metabolite consumption and metabolic gene expression in human macular and peripheral RPE/choroids. **(a-c)** Metabolite consumption from macular and peripheral RPE/choroid explant culture. (**d-g**) Metabolite release from the macular and peripheral RPE/choroid explant culture. Data were relative ion abundance over control or peripheral retinas, or absolute abundance when detected in the baseline control. N=8, *p<0.05 vs. Control or peripheral RPE/Choroids. (**h)** Gene expression of the top 20 abundant small molecule transporters in human peripheral and macular RPE/choroids. (**i**) A schematic of nutrient uptake and release in macular RPE/choroid. Red-colored metabolites are those with increased uptake or release in macular RPE/choroid. ABC transporters and glucose transporters (SLC5A9 and SL2A4) are upregulated in macular RPE/choroid.

As well as lactate, nucleosides and acylcarnitines, RPE/choroid from both macula and periphery exported considerable amounts of citrate, isocitrate, 3-HB and cholesterol (**Fig 4d-e**), supporting previous reports ^23, 26^ that RPE actively metabolizes lipids. Compared with their corresponding neural retinas, RPE/choroid exported ∼40 times more N1-methylnicotinamide (methylated from nicotinamide), indicating that RPE/choroid has highly active nicotinamide catabolism (**Fig 4f**). Additionally, RPE/choroid released serine, β-alanine and glycerol, which were consumed by the human neural retinas (**Fig 4d-e, Fig S7**), providing further evidence for metabolic communication between the neural retina and the RPE.

The peripheral RPE/choroid punches consumed and released significantly more metabolites than the macular punches (**Fig 4, Fig S7**). However, the macular punches generally consumed or released more of the changed metabolites that they had in common with the peripheral RPE/choroid punches (**Fig 4c, 4g**). The macular RPE/choroid punches utilized more nicotinamide, erythritol and glucose than their peripheral counterparts (**Fig 4c**). Erythritol comes from either additives or endogenous synthesis from glucose. It can be metabolized into erythronic acid, which is abundant in human aqueous humor ^34^. RPE/choroid punches from the macular region also released more N1-methylnicotinamide, malate, β-alanine and ornithine, which may have been derived from the methylation of nicotinamide and glucose catabolism (**Fig 4f, 4g, 4i**).

Unlike neural retinas, no aspartate/glutamate transporters were found in the top 20 most abundant small molecule transporters in the RPE/choroid (**Fig 4h, Fig S8**). The abundance of small molecule transporters was generally higher in the peripheral RPE than the macula, supporting our data that there were more changed metabolites in the periphery. MCTs for lactate, pyruvate and ketone bodies, and transporters for α-ketoglutarate, sugar, leucine, proline and choline, were abundant in peripheral RPE/choroid (**Fig 4h, Fig S8**), consistent with the altered uptake and release of these metabolites. The analysis of expression of metabolic genes found that there were 127 genes that were differentially expressed between macular and peripheral RPE/choroid (**Table S11**). The major altered pathways included lipid metabolism, transporters and amino acid metabolism (**Fig S9**). Interestingly, genes involved in glycolysis and the TCA cycle were either decreased or unchanged in the macular RPE/choroid, while glucose transporters (*SLC2A4, SLC5A9*) were significantly increased (**Fig S9, Table S11, Fig 4i**). Overall, genes in both lipid synthesis and lipid transporters were upregulated in the macular RPE/choroid punches compared with those from the peripheral RPE/choroid retina **(Fig 4i)**. These data suggest that macular RPE suppresses its glycolysis to preserve more glucose for the neural retina.

### Human retina and RPE/choroid have distinctive but symbiotic metabolic features

To compare the metabolic features of the human retina and RPE/choroid, we studied the uptake and release of metabolites in the media from their cultures (**Fig 3-4, Fig 5a**). We found 25 metabolites were consumed and released in the same patterns between retinas and RPE/choroid such as glucose, nicotinamide, carnitine, lactate and inosine. Twenty-two metabolites were consumed or released only by the neural retina, indicating these metabolites are neural retina-specific. Sixteen metabolites were consumed or released only by RPE/choroid, indicating that they are RPE specific (**Fig 5a**). Interestingly, ten metabolites were consumed or released in opposite patterns, such as serine, ornithine and β-alanine (**Fig 5a**) which were released by RPE/choroid but consumed by the neural retinas, suggesting that retinas and RPE are metabolically inter-dependent. Overall, we found neural retinas and RPE/choroid share some common pathways, but also have their distinctive preferences in nutrient consumption and production.

**Figure 5.**
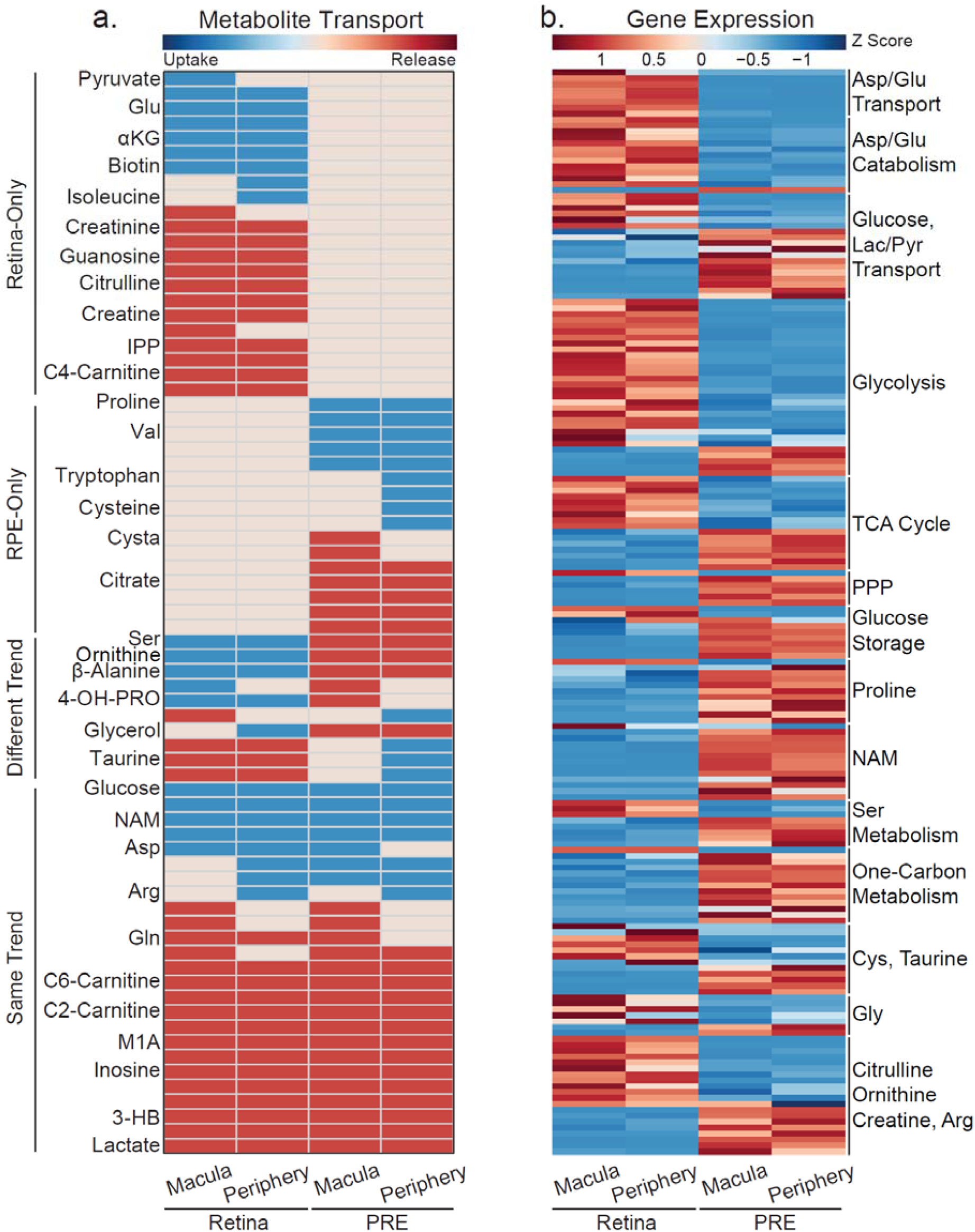
Comparison of metabolite consumption and metabolic genes between human retinas and RPE/choroids. **(a)** Differential metabolite consumption and release between human retinas and RPE/choroids. Metabolites are colored for changes over their baseline controls. Green = metabolite consumption, red = metabolite release and yellow = unchanged. (**b**) Comparing gene expression in major metabolic pathways between human retinas and RPE/choroids. M, Macula; P, Periphery; Pyr, pyruvate.

About half of the 2,764 metabolic genes we studied were expressed differentially in human neural retinas vs. RPE/choroids: 1473 in the macula and 1770 in the periphery (**Table S12-13**). Transporters, lipids, glycan, amino acid, nucleotide and sugar metabolism were again the major altered pathways between retinas and RPE/choroids (**Fig S4**). All genes involved in aspartate/glutamate transport and catabolism were upregulated in the neural retinas except aspartoacylase, which regenerates aspartate from N-acetyl-aspartate. (**Fig 5b, Table S13**). This strongly suggests that retinas have a specific requirement for aspartate and glutamate, as summarized in **Fig 6**. More transporters for glucose, lactate and pyruvate were found in RPE/choroid but they were also enriched in the retinas, indicating that these metabolites are essential nutrients for both tissues. As expected, neural retinas predominantly expressed genes involved in glycolysis to support their robust aerobic glycolysis (**Fig 5b, Fig S10, Table S14**). Interestingly, many mitochondrial TCA cycle genes expressed different isoforms in neural retinas vs. RPE/choroid. For example, NAD^+^-dependent *IDH3A/B*, ADP-dependent *SUCLA2*, ACO2 and L2HGDH were predominantly expressed in the neural retinas, while NADP^+^-dependent IDH1/2, GDP-dependent *SUCLG2, ACO1* and *D2HGDH* were mainly expressed in the RPE/choroid (**Fig 6, Table S14**). Notably, upregulated TCA cycle genes in the neural retinas were mostly NAD^+^-dependent. This differential expression suggests neural retinas and RPE have different roles in mitochondrial bioenergetics and biosynthesis. Strikingly, NADP(H)-associated pathways, including the pentose phosphate pathway (PPP) and one-carbon metabolism, were highly upregulated in the RPE/choroid compared to neural retinas (**Fig 5b**). Genes in serine synthesis were upregulated in RPE/choroid, but serine transporters were upregulated in the neural retinas. This was consistent with our findings that serine was released by RPE/choroid and consumed by retinas (**Fig 5b, Table S14**). Remarkably, genes involved in proline metabolism and nicotinamide/NAD metabolism were significantly upregulated in RPE/choroid, consistent with our findings that RPE/choroid consumed proline and nicotinamide. Many genes in amino acid metabolism pathways, such as those for cysteine, taurine and citrulline, were partly upregulated in neural retinas or RPE/choroids, further indicating that there are symbiotic metabolic relationships between them (**Fig 6**).

**Figure 6.**
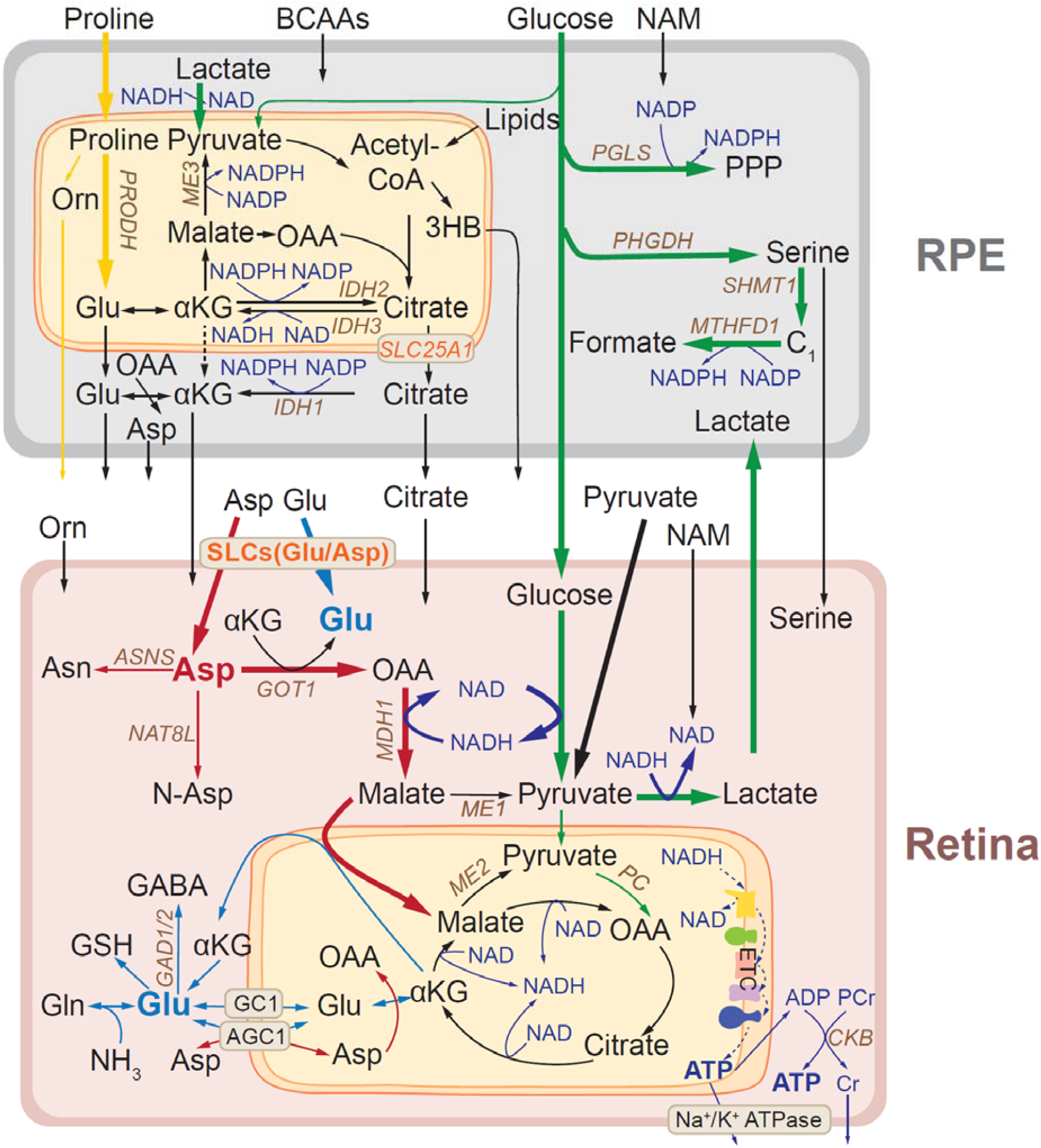
A proposed model of metabolic communications between human retina and RPE/choroid. Glucose transports through RPE from the choroid to the retina (bold green line). RPE strongly expresses genes in PPP, serine de novo synthesis and one-carbon metabolism, which can use glucose to generate NADPH. The retina strongly expresses glycolic genes to produce lactate and NADH. The retina imports aspartate and glutamate through upregulated transporters (bold red line). Aspartate can be converted into OAA and malate to recycle NADH into NAD+ and bring electrons into mitochondria to produce ATP. Aspartate can also be used for the synthesis of glutamate, asparagine (ASN), N-acetylaspartate (Asp). The retina needs a large amounts of glutamate for neurotransmission, the mitochondrial TCA cycle and biosynthesis of GABA, glutathione (GSH) and glutamine (Gln) (bold blue line). RPE mitochondria use different substrates such as proline (yellow line), BCAAs and lipids to synthesize NADPH and other metabolites that can be released to be used by the retina. Cone-rich retinas or human macula consumes more pyruvate (bold black line) for mitochondrial metabolism. Brown colored genes are upregulated genes in the retinas or RPE.

## Discussion

We have compiled an atlas of nutrient consumption and utilization in human retinal organoids, mouse retinas, all cone mouse retinas of *Nrl* KO and human macular and peripheral retinas. By combining nutrient consumption with metabolic gene analysis, we have identified the following key features of neural retinal and RPE metabolism (**Fig 6**).

- Neural retinal metabolism centers on NADH-related metabolism for ATP production and the synthesis of neurotransmitters such as glutamate and GABA. The neural retina consumes glucose, aspartate, glutamate, citrate, nicotinamide and other nutrients to sustain its active glycolysis, mitochondrial oxidative metabolism and neurotransmitter synthesis.
- Cone-rich retinas and the neural macula have more active glycolysis and mitochondrial metabolism, through which they consume more pyruvate, than the rod-rich peripheral neural retina.
- Mature retinal organoids have aspartate/glutamate metabolism that is similar to the human neural retina.
- RPE/choroid metabolism centers on NADPH-related and acetyl-CoA metabolism for biosynthesis. RPE/choroid consumes glucose, proline, nicotinamide, BCAAs and other nutrients for its activate PPP, reductive carboxylation, serine synthesis, one-carbon metabolism, citrate synthesis, ketone body synthesis and cholesterol synthesis.
- Neural retinas and RPE/choroid have specific and complementary profiles in nutrient consumption and production that support their tight metabolic communications.

Uptake of aspartate and glutamate was a common feature in cone-dominant retinal organoids, mouse and human retinas. Transporters and enzymes responsible for aspartate and glutamate metabolism were also highly enriched in the retina. Why do retinas actively consume aspartate and glutamate? Aspartate and glutamate are inter-convertible by transamination with their ketoacids through *GOT1/2* (**Fig 6**). We recently also reported that aspartate transamination is the primary pathway for the catabolism of aspartate and glutamate in mouse neural retinas ^35^. As a major neurotransmitter, glutamate is the most abundant amino acid in the retina ^36, 37^. Retinal cells with the highest glutamate levels also have the highest aspartate levels^37^. We have previously reported that aspartate is utilized robustly to provide both carbons and nitrogen groups for glutamate synthesis ^8, 35, 38^. Notably, aspartate transamination also generates oxaloacetate (OAA) which can be converted to malate by cytosolic malate dehydrogenase (*MDH1*) to carry electrons into mitochondria through the malate-aspartate shuttle (**Fig 6**). This is important because it may help address the paradox of how the retina supports its active mitochondrial metabolism while its metabolism relies on the Warburg effect that allows only a small proportion of pyruvate to be oxidized. *MDH1* and *GOT*1 transcripts are more abundant than any other TCA cycle enzymes in both human and mouse neural retinas (**Fig S11**), supporting that aspartate catabolism is enhanced in the retina. Aspartate catabolism may 1) recycle NADH into NAD^+^ to sustain the highly active glycolysis, 2) provide cytosolic electrons for ATP production, 3) replenish mitochondrial OAA for biosynthesis and 4) produce cytosolic and mitochondrial pyruvate through malic enzymes (**Fig 6**). Interestingly, low glucose or inhibition of glucose oxidation could cause massive accumulation of aspartate in the neural retina with severe depletion of glutamate ^38, 39, 40^. Therefore, aspartate utilization supports active glycolysis, mitochondrial metabolism and glutamate synthesis. Additionally, aspartate is an immediate precursor for asparagine and N-acetylaspartate (NAA), both of which are important for neuronal function ^41, 42^ (**Fig 6**). Deficiency of the mitochondrial aspartate/glutamate carrier 1(*AGC1*) reduces aspartate and NAA, resulting in brain defects and impaired visual function ^42, 43, 44^.

Besides being a neurotransmitter, glutamate is an important substrate for the TCA cycle and the synthesis of glutamine and glutathione (**Fig 6**). Decreased glutamate availability substantially reduces the levels of glutamine and glutathione ^39^. Glutamate also serves as a major nitrogen donor to synthesize non-essential amino acids by transaminases and is a precursor for GABA by decarboxylation (**Fig 6**). Consistently, glutamate alanine transaminase and glutamate decarboxylases are strongly expressed in the neural retina ^33^. Blood levels of aspartate and glutamate are quite low ^45, 46^, it remains unclear how they are transported across retina-blood barriers. RPE may be a source of aspartate and glutamate for the retina. We have reported that RPE can metabolize glucose and proline into glutamate and aspartate, which are exported to be used by the retina ^28, 47^. Notably, glutamate content decreases in cone photoreceptors overlying RPE defects in atrophic AMD ^48^.

Cones are known to have much higher energy needs than rods ^49^. It is postulated that mitochondrial metabolism may support the extra energy demand, as cones have 2-10 times more mitochondria and 3 times more cristae membrane surface than rods ^11, 12, 50^. The generation of higher lactate, acetylcarnitine and IPP (synthesized from acetyl-CoA) in both all-cone mouse retinas and the neural macula, which is cone-rich, provides strong evidence that cones are more metabolically active than rods. We also found all-cone neural retinas and macula consume more pyruvate than the peripheral neural retina, which is rod dominant. This may explain how cones fuel their abundant mitochondria even when they have active aerobic glycolysis. The SLC16A3 (MCT4), which has low affinity for pyruvate in order to preserve intracellular pyruvate levels, ^51^ is strongly expressed in the neural macula (**Table S9**). Inhibition of pyruvate utilization causes visual impairment in mice and fish due to photoreceptor degeneration ^39, 52^. Pyruvate provides metabolic advantages that enhance both glycolysis and mitochondrial metabolism. Pyruvate can regenerate NAD^+^ through lactate dehydrogenase to stimulate glycolysis. Mitochondria need a constant supply of acetyl groups from acetyl-CoA and OAA to synthesize citrate to sustain the TCA cycle for oxidative and biosynthetic metabolism (**Fig 6**). Pyruvate provides both acetyl groups and OAA through pyruvate dehydrogenase and *PC*. The neural retina can also use fatty acids and ketone bodies, but these nutrients only provide acetyl groups ^23, 53^. Consistently, pyruvate carboxylase expression was upregulated in both all-cone retinas and neural macula. Hepatic deletion of *PC* in mice diminishes TCA cycle intermediates, especially aspartate, and depletes NADPH and glutathione ^54^. PPP is the classic pathway to generate NADPH for the regeneration of visual pigment, but its activity is very low in the neural retina and photoreceptors ^55, 56, 57^. The regeneration of visual pigment in cones is at least 100 times faster than rods ^58^. Pyruvate could stimulate PPP activity 15 times in isolated neural retinas^55^ and may contribute to the high demand for NADPH in cones. Pyruvate also increases the availability of glutamate and glutamine *in vitro* ^59, 60^. We have reported that retinal deletion of the mitochondrial pyruvate carrier leads to the accumulation of pyruvate and aspartate but the depletion of glutamate, glutamine and glutathione results in retinal degeneration ^39^. Interestingly, pyruvate supplementation prevents light-induced retinal degeneration ^61, 62, 63^. Further research is warranted to determine whether increasing pyruvate utilization by supplementation or genetic approaches might have a neuroprotective effect on the macula.

The RPE is closely coupled with photoreceptors in nutrient utilization ^20, 22, 23, 35^. In agreement with other studies of RPE cells ^28, 35, 47^, we found that human RPE/choroid explants consumed glucose, proline, nicotinamide and BCAAs, and released plenty of 3-HB, citrate and cholesterol. All these released metabolites are made from acetyl-CoA, indicating that RPE mitochondria function like mitochondria in the liver that oxidize nutrients to support other tissues. Interestingly, retinas could readily utilize the released 3-HB, citrate and cholesterol for bioenergetics and biosynthesis ^23, 28, 39, 55, 64^. The different consumption of metabolites by the neural retina and RPE/choroid suggests an important metabolic coupling between them. For example, RPE released serine, but the retina consumed serine. Serine is an essential precursor for phospholipid synthesis and one-carbon metabolism. Photoreceptors require active phospholipid biosynthesis to renew their daily shedding of outer segments, but enzymes for serine synthesis are highly expressed in RPE (**Fig 5b**) ^65^. The uptake of serine by our neural retinal cultures and upregulated serine transporters sideroflexin 1/3 (*SFXN1/3*) in the neural retina further suggest that retinas need exogenous serine. Mutations of SFXN3 leads to progressive outer retinal degeneration ^66^. Remarkably, low serine from the RPE may account for the synthesis of toxic hydroxysphingophospholipids in the degenerative macular condition, macular telangiectasia type 2 ^67^.

In summary, profiling nutrient uptake and release by tissue explants provides a useful platform to investigate tissue-specific and inter-tissue metabolism and to functionally annotate the ever-increasing tissue-specific metabolic transcriptome. By profiling mouse and human retinal explants, we reveal distinctive metabolic features in progenitor vs. differentiated retinal organoids, mouse rod vs. cone-retinas, and human peripheral neural retina vs. the neural macula. These findings provide a resource of basic information for future studies on the retina and RPE metabolism.

## Methods

### Reagents

All the reagents and resources are detailed in the Key Resources table (**Table S15**)

### Human retinal organoid culture

Cone-rich human retinal organoids were generated from H1 human embryonic stem cells (WiCell) using a method described previously ^16, 68^. For the assay of metabolite consumption, one retinal organoid on day 56 or day 296 was grown in a well of a 24-well plate containing 800 µl of culture medium (DMEM/F12 (3:1), 2% B27, 1x non-essential amino acids (NEAA), 8% fetal bovine serum (FBS), 100 µM Taurine and 2 mM GlutaMAX (see detail information in **Table S6** and **Table S15**). Culture wells containing 800 µl of the medium without retinal organoids were used as baseline controls. After 48 hours of culture, 50 µl of the spent medium was collected from each culture well for metabolite analysis.

### Mouse retinal explant culture

*Nrl* KO mice were generously provided by Anand Swaroop, PhD (National Eye Institute, Bethesda, MD) and were housed and bred at the West Virginia University Animal Research Facility vivarium. The *Nrl* KO mice were bred with C57BL6/J mice purchased from Jackson Laboratories (Bar Harbor, ME) to generate *Nrl* Het mice which were further bred with *Nrl* KO mice. Our studies used littermates at postnatal 36 days with half male and half female. All procedures were in accordance with the National Institutes of Health guidelines and the protocols approved by the Institutional Animal Care and Use Committee of West Virginia University. Mouse explant culture was performed as previously reported ^28^. Briefly, the retina was isolated and made four relief cuts on each side in retinal culture medium containing DMEM/F12 (3:1), 2% B27, 1x NEAA, 1% FBS, and 100 µM Taurine (detailed in **Table S7** and **Table S15**). The retina was then transferred into an insert of 24-well transwell plate with photoreceptor-side facing down. The bottom chamber contained 500 µl and top chamber 100 µl of retinal culture medium. Transwells containing medium without retinas were cultured as the baseline controls. After 24 hours, 50 µl of the spent medium was collected from the bottom chamber for metabolite analysis.

### Human retina and RPE/choroid explant culture

Post-mortem donor eyes without known eye conditions were obtained from Lions NSW eye bank. Ethical approval for this project was obtained from the Human Research Ethics Committee of the University of Sydney. The eyes were kept in CO_2_ independent medium in 4™C until further dissection. All donor eyes within 28 hours post-mortem (**Table S5**) were cut open by removing iris and lens, followed by cutting a circle of the sclera with 0.5cm thickness. Fundus photographs were taken by a digital microscope camera (Jenoptik Optical System) after the anterior segment was removed.

The whole neural retina was gently detached after separated from the ora serrata and optic nerve head. Then it was bluntly separated from vitreous and the neural retina was unfolded in a glass petri dish (150mm×15mm) with neurobasal-A medium. 5mm-diameter circular neural retinas were trephined from the macula and superior mid-peripheral retina using a Biopsy Punch (**Fig. 3a**). The macula punch was on the fovea. Mid-periphery was defined as the mid-point between the macula lutea and the ora serrata. RPE-choroid-sclera complexes (RPE/choroid) were trephined from macula and mid-periphery as the same location in retinal punches. Both retinal and RPE/choroid explants were transferred to the inserts of 24-well transwell plates (Corning, CLS3421) with culture medium. Neural explants were cultured with photoreceptors facing down, and RPE-choroid-sclera complexes were cultured with RPE layer facing up. Retinal culture media was modified from mouse retinal culture (**Table S8 lists detail formulation**) and RPE/choroid culture media was the same as we reported ^28^ (**Table S16**). The volume of medium for neural explants in the insert is 100µl and in well is 600µl. The volume for RPE explant is 150µl in insert and 900µl in well. **Table 15** listed details for the supplies and culture media. Neural retina and RPE explants were cultured for 24h at 37°C in 5% CO_2_ incubator. 10µl spent medium was collected from the lower chamber for metabolite analysis.

### Metabolite analysis

Metabolites were extracted from 10 µl of medium samples, dried with freeze dryer, and analyzed with Liquid chromatography–mass spectrometry (LC MS) and gas chromatography–mass spectrometry (GC MS) as described ^28, 29, 39^. LC MS used a Shimadzu LC Nexera X2 UHPLC coupled with a QTRAP 5500 LC MS (AB Sciex). An ACQUITY UPLC BEH Amide analytic column (2.1 × 50 mm, 1.7 μm, Waters) was used for chromatographic separation. A total of 190 metabolites were measured with optimal parameters for targeted metabolomics (**Table S1**). GC MS used an Agilent 7890B/5977B system with a DB5MS column (30 m × 0.25 mm × 0.25 μm). Mass spectra were collected from 80–600 m/z under selective ion monitoring mode to quantify 28 metabolites (**Table S1**). The ion abundance of each metabolite was divided by those from baseline controls to obtain fold changes in abundance. Metabolite uptake or release defined based on the fold changes <-1 or >1.

### Statistics

The significance of differences between means was determined by unpaired or paired two-tailed t tests using GraphPad Prism 7. The data were presented as mean ± Standard deviation. P < 0.05 was considered to be significant. The differences in metabolic genes were analyzed by Volcano plot using MetaboAnalyst 4.0 (https://www.metaboanalyst.ca/). The changes with fold changes >1.3 and P<0.05 were set as significant.

## Abbreviations

AC-Gly: Acetyl-Glycine;
Ala: Alanine;
Arg: Arginine;
Asn: Asparagine;
Asp: Aspartate;
BAIBA: β-aminoisobutyric acid;
β-Ala: β-alanine;
C2: Acetylcarnitine;
C3: Propionylcarnitine;
C4: Butyrylcarnitine;
C4-me: 2-methylbutyryl-carnitine;
C6: Hexanoylcarnitine;
Cysta: Cystathionine;
GABA: Gamma-aminobutyric acid;
Gln: Glutamine;
Glu: Glutamate;
3-HB: 3-Hydroxybutyrate;
Hypoxan: Hypoxanthine;
Iso-C4: Isobutyrylcarnitine;
Isocitr: Isocitrate;
Isoleu: Isoleucine;
IPP: Isopentyl Pyrophosphate;
α-KG: a-Ketoglutarate;
Leu: Leucine;
Lys: Lysine;
1-MA: 1-Methyladenosine;
Met: Methionine;
NAM: Nicotinamide;
N1-MNA: N1-Methylnicotinamide;
4-OH-PRO: 4-Hydroxyproline;
Phenylpyr: Phenylpyruvate;
Pro: Proline;
Ser: Serine;
Val: Valine;
Vit-B5: Vitamin B5.

## Acknowledgments

We thank Dr. Anand Swaroop and Dr. Zachary Batz from the National Eye Institute for sharing *Nrl* KO animals and their transcriptome datasets. We appreciate the comments from Dr. James Hurley from the University of Washington. We acknowledge the NSW lion tissue bank’s support for this research. This work was supported by NIH Grants EY026030 (J.D.), EY029806 (W.L.), the Retina Research Foundation (J.D.), funds for Core facilities P20 GM103434 (WV INBRE grant), WVCTSI grant GM104942, funds from Australian NHMRC project grant (APP1145121 M.C.G) and the Lowy Medical Research Institute. Professor Mark C. Gillies is a Sydney Medical School Fellow and supported by an Australian NHMRC practitioner fellowship.

## Author Contributions

Conceptualization, J.D; Investigation, B.L., T.Z., W.L., Y.W., R.X., S.Z., R.Z., S.Z., L.Z.; Writing, B.L., T.Z., W.L., L.Z., M.C.G., and J.D.; Funding Acquisition, J.D.; Supervision, L.Z., M.C.G., and J.D.

