## Supplementary Materials for "Metabolic features of mouse and human retinas: rods vs. cones, macula vs. periphery, retina vs. RPE"

**This PDF file includes** Supplementary **Figs. S1 to S11** and **Tables S1 to S16**

**Fig S1.** Metabolic genes are temporally expressed in retinal organoids	2

**Fig S2.** Top small molecule transporters ranked by WT retinas	3

**Fig S3.** Changed metabolic genes in the NRL KO vs WT retinas	4

**Fig S4.** Metabolite consumption and release in human macular and peripheral retinas	5

**Fig S5.** Top small molecule transporters ranked by peripheral retinas	6

**Fig S6.** Changed metabolic genes in the macular and peripheral retinas	7

**Fig S7.** Metabolite consumption and release in human macular and peripheral RPE/choroid	8

**Fig S8.** Top small molecule transporters ranked by peripheral RPE/choroid	9

**Fig S9.** Changed metabolic genes in the macular and peripheral RPE/choroid	10

**Fig S10.** Changed metabolic genes between human retinas and RPE/choroid	11

**Fig S11.** The expression of TCA cycle genes in mouse and human retinas	12

**Table S1.** List of metabolites and parameters by targeted metabolomics	13

**Table S2.** List of 2764 classic metabolism genes and their classifications	20

**Table S3.** Differentially expressed metabolic genes in retinal organoids	100

**Table S4.** Differentially expressed metabolic genes in WT and *Nrl* KO retinas	108

**Table S5.** Donor information for human retina and RPE/choroid explants	126

**Table S6.** Medium formulation for retinal organoid culture	127

**Table S7.** Medium formulation for mouse retinal explant culture	130

**Table S8.** Medium formulation for human retinal explant culture	133

**Table S9.** Differentially expressed metabolic genes in macular and peripheral retinas	135

**Table S10.** Comparing the changed metabolic genes in the retinas between WT vs *Nrl* KO and macula vs periphery	146

**Table S11.** Differentially expressed metabolic genes in macular and peripheral RPE/choroids	151

**Table S12.** Differentially expressed metabolic genes in macula between retinas and RPE/choroid	154

**Table S13.** Differentially expressed metabolic genes in periphery between retinas and RPE/choroid	187

**Table S14.** Differentially expressed major metabolic genes between retinas and RPE/choroid	226

**Table S15.** Key reagent and resources	231

**Table S16.** The medium formulation for human RPE/choroid culture	233

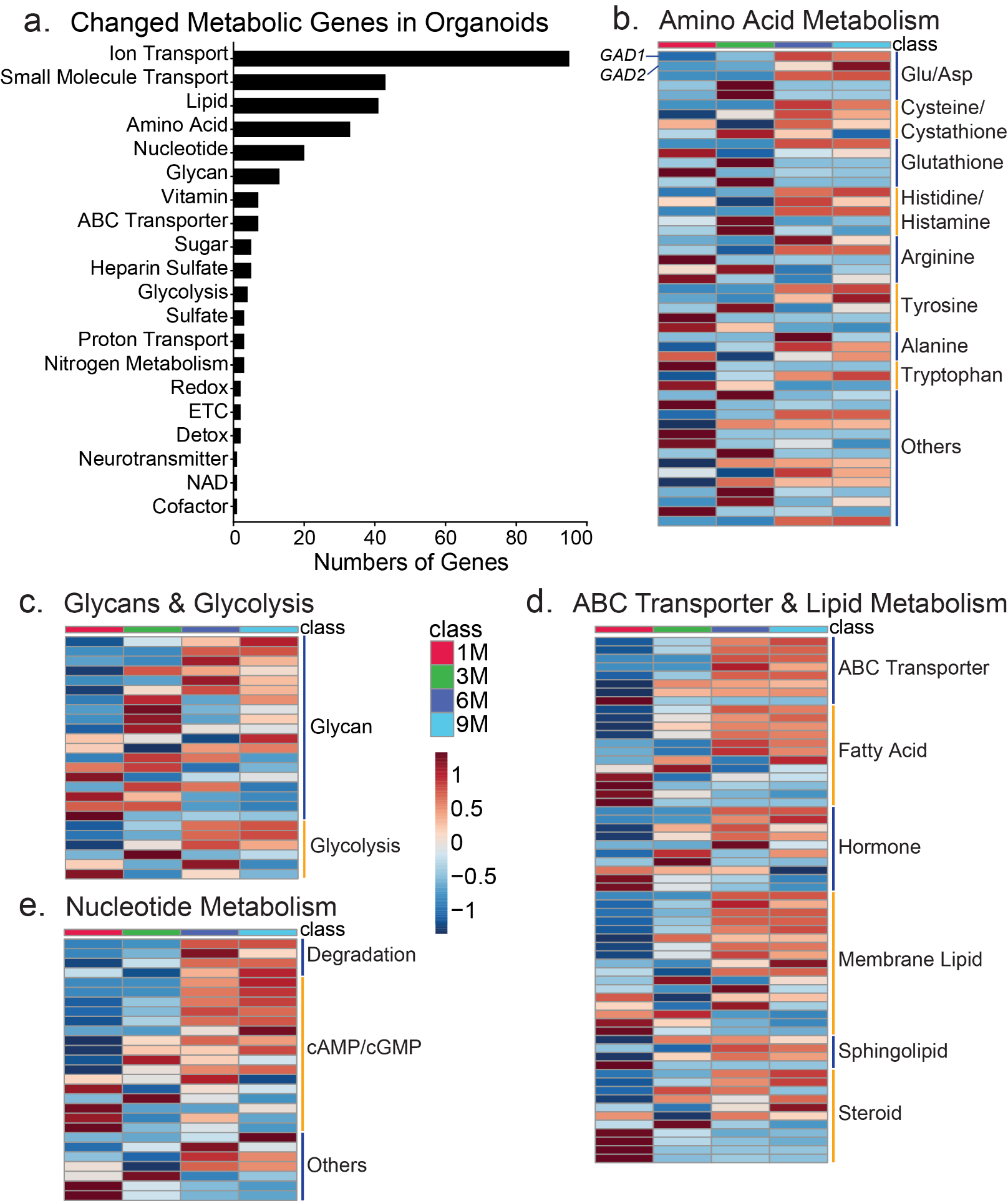

**Fig S1. Metabolic genes are temporally expressed in retinal organoids.** Differentially expressed genes along the time-course of human retinal organoids were intersected with the genes in metabolic pathways, resulting in a list of developmentally regulated metabolic genes. (**a**) Subgroups of developmentally regulated metabolic genes. (**b-e**) Heatmaps showing temporal expression of metabolic genes in (**b**) amino acid, (**c**) glycans & glycolysis, (**d**) ABC transporter & lipid and (**e**) nucleotide in human retinal organoids.

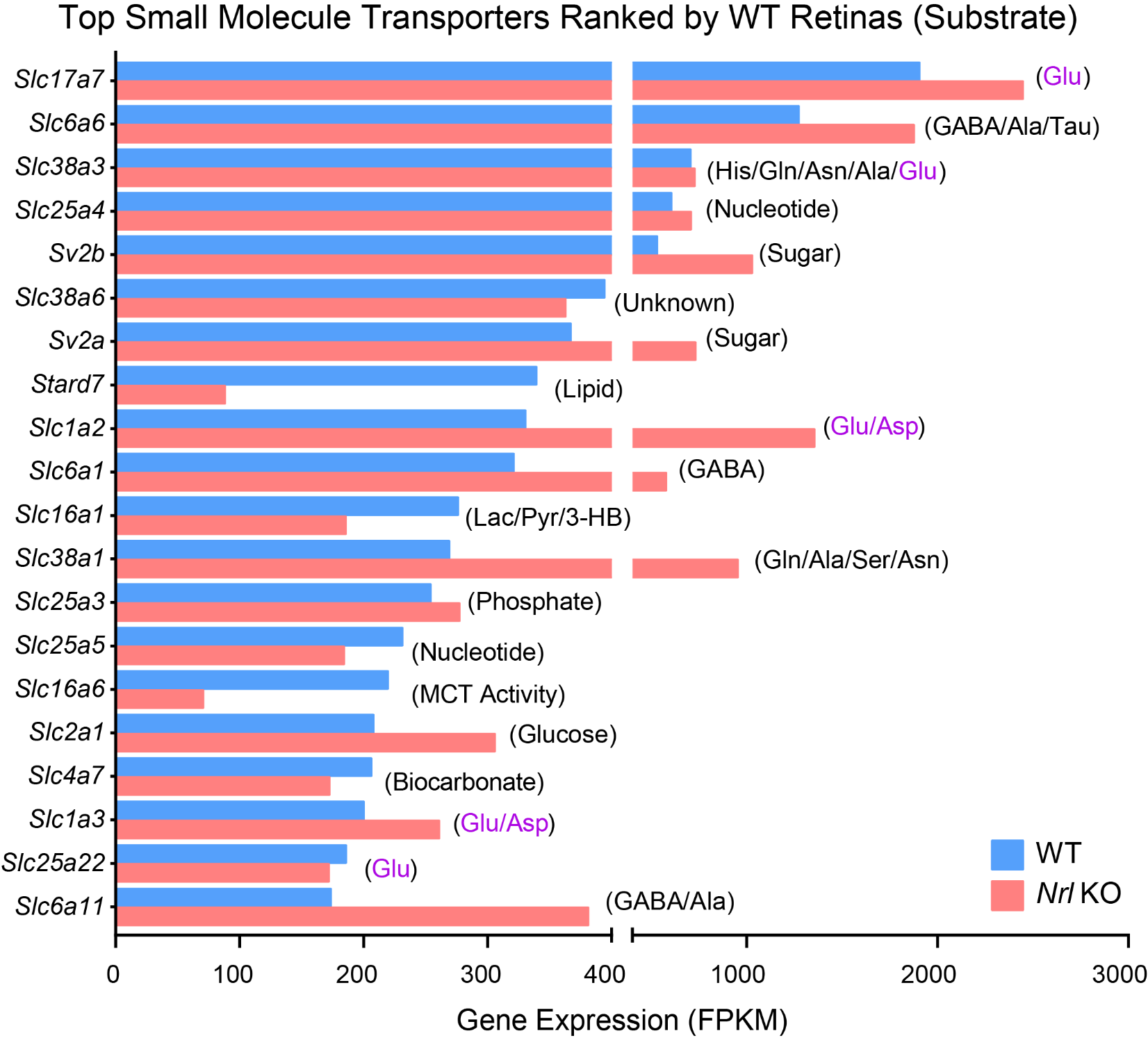

**Fig S2. Top small molecule transporters ranked by WT retinas**. Top 20 most abundant small molecule transporters ranked by the expression in WT retinas.

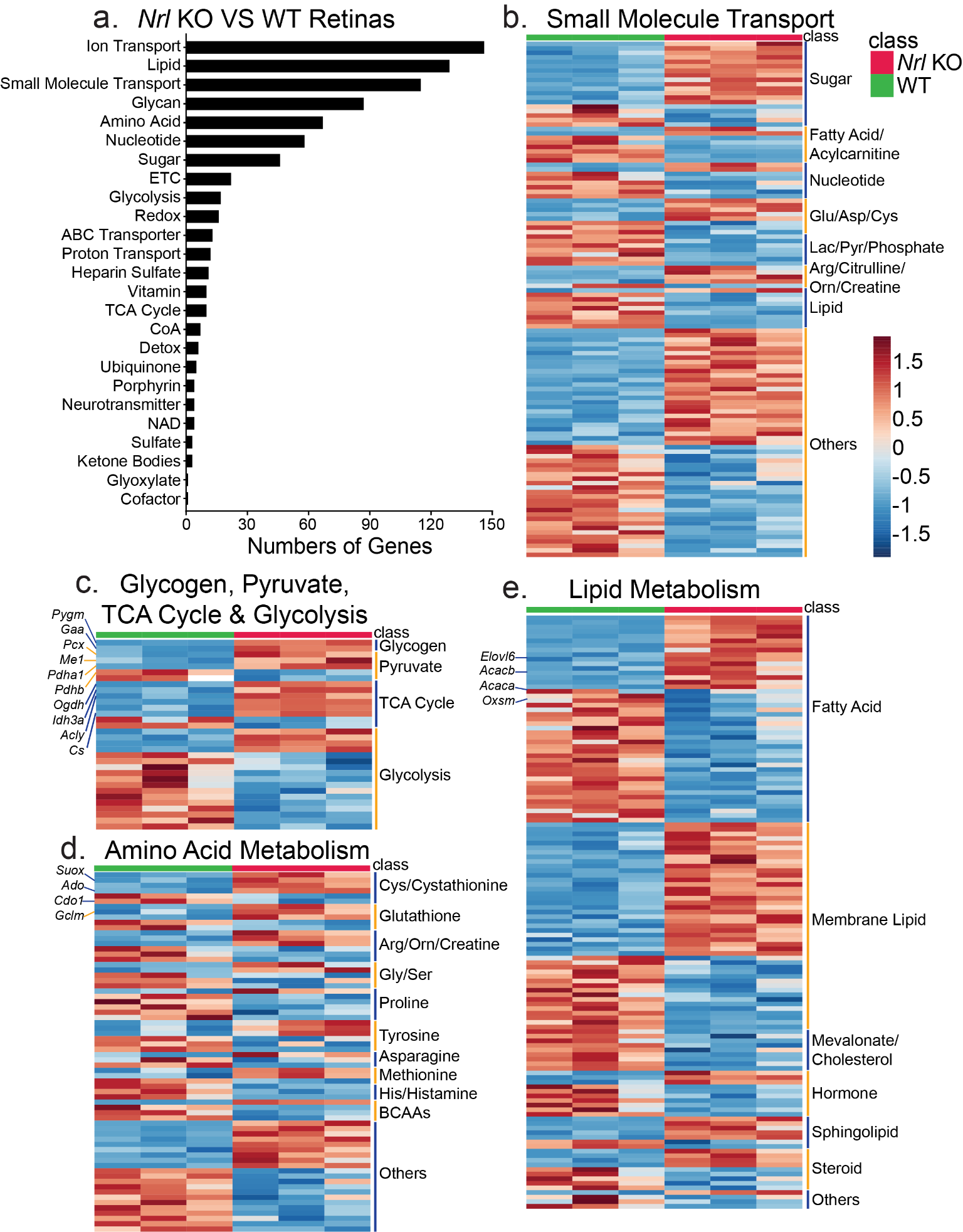

**Fig S3. Changed metabolic genes in the *Nrl* KO vs WT retinas. (a)** Number of changed genes in metabolic pathways in *Nrl* KO retinas. **(b-e)** Heatmaps of differentially expressed genes in **(b)** small molecule transport, **(c)** glycogen, pyruvate, TCA cycle & glycolysis, **(d)** amino acid, **(e)** lipid in *Nrl* KO retinas.

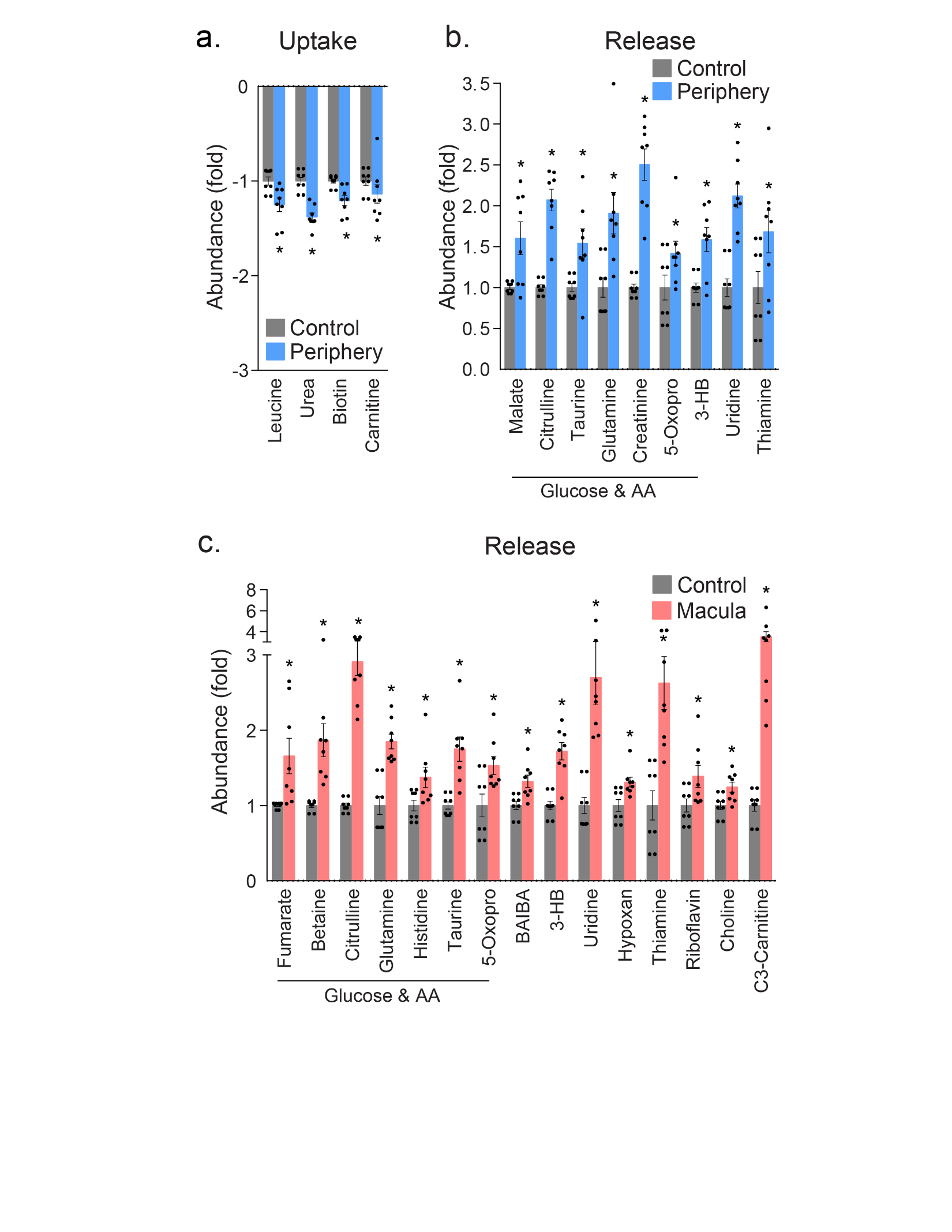

**Fig S4. Metabolite uptake and release in human macular and peripheral retinas. (a)** Uptaked metabolites in the medium from peripheral retina culture. **(b)** Released metabolites in the medium from peripheral retina culture. **(c)** Released metabolites in the medium from macular retina culture. N=8, **P*<0.05 vs baseline control.

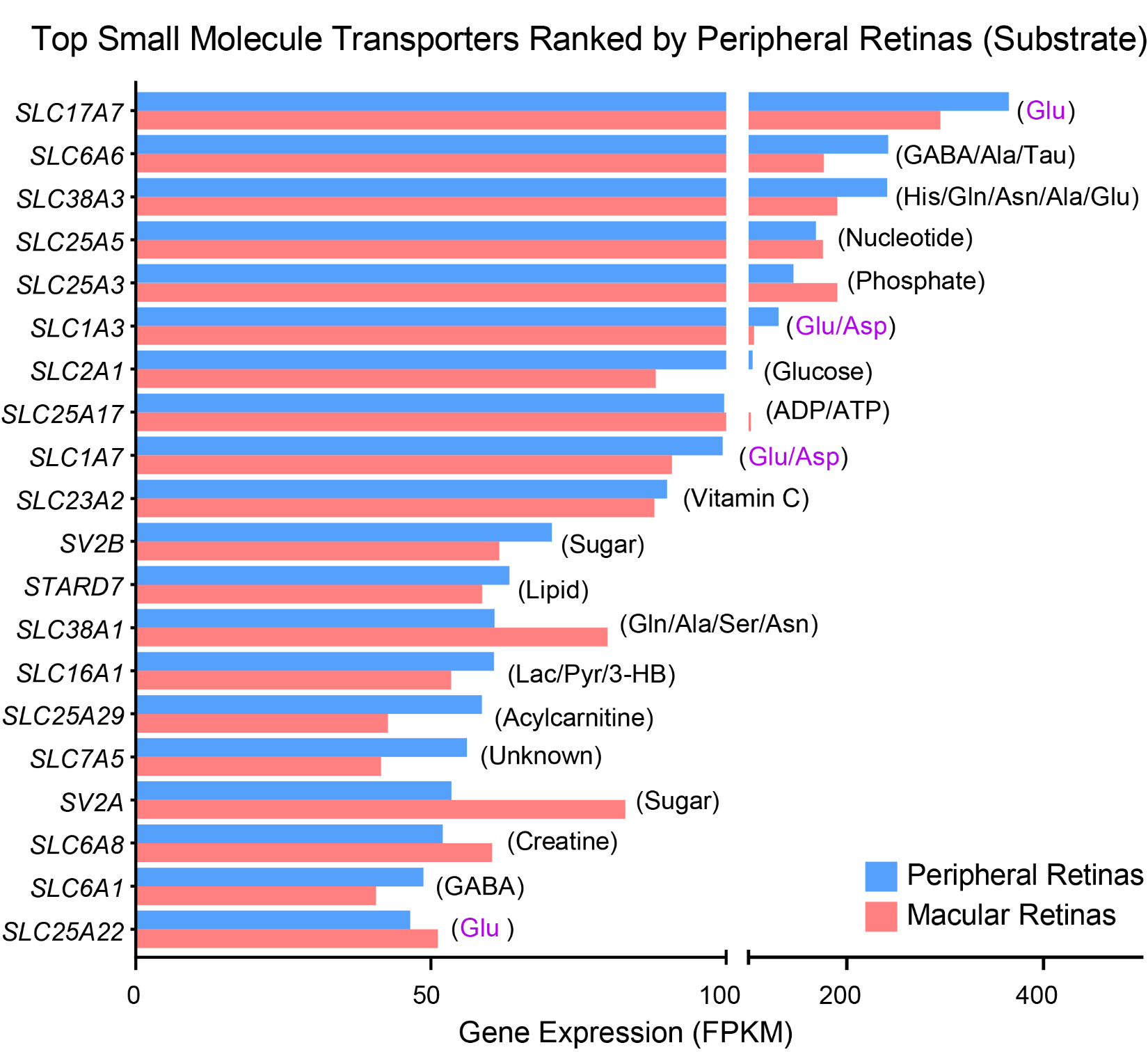

**Fig S5. Top small molecule transporters ranked by peripheral retinas.** Top 20 most abundant small molecule transporters ranked by the expression in human peripheral retinas. Periphery expression was the average of the expression in nasal and temporal regions.

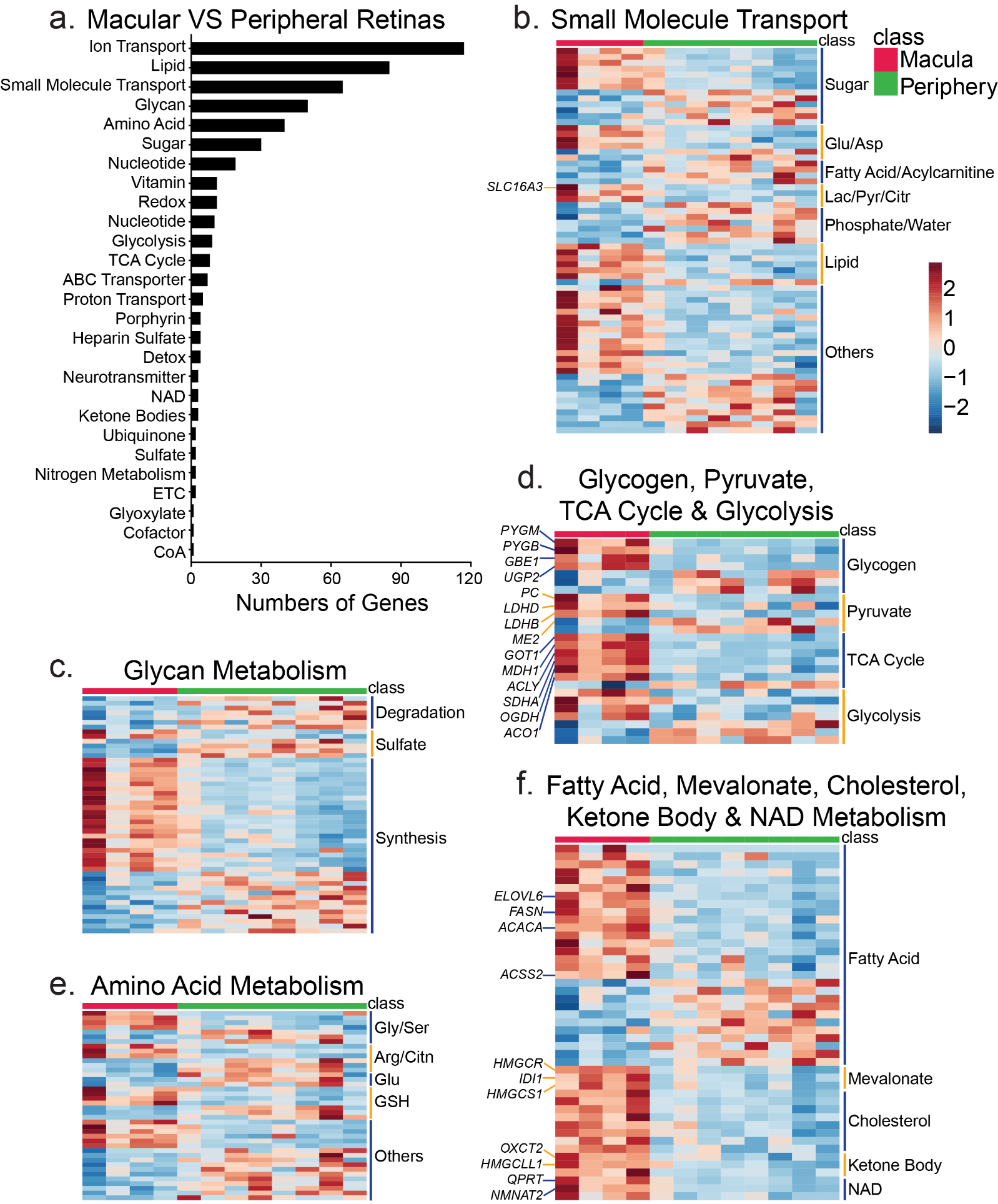

**Fig S6. Changed metabolic genes in the macular and peripheral retinas. (a)** Number of changed genes in metabolic pathways. **(b-e)** Heatmaps of differentially expressed genes in **(b)** small molecule transport, **(c)** glycans, **(d)** glycogen, pyruvate, TCA cycle & glycolysis, **(e)** amino acid, **(f)** fatty acid, mevalonate, cholesterol, ketone body & NAD metabolism.

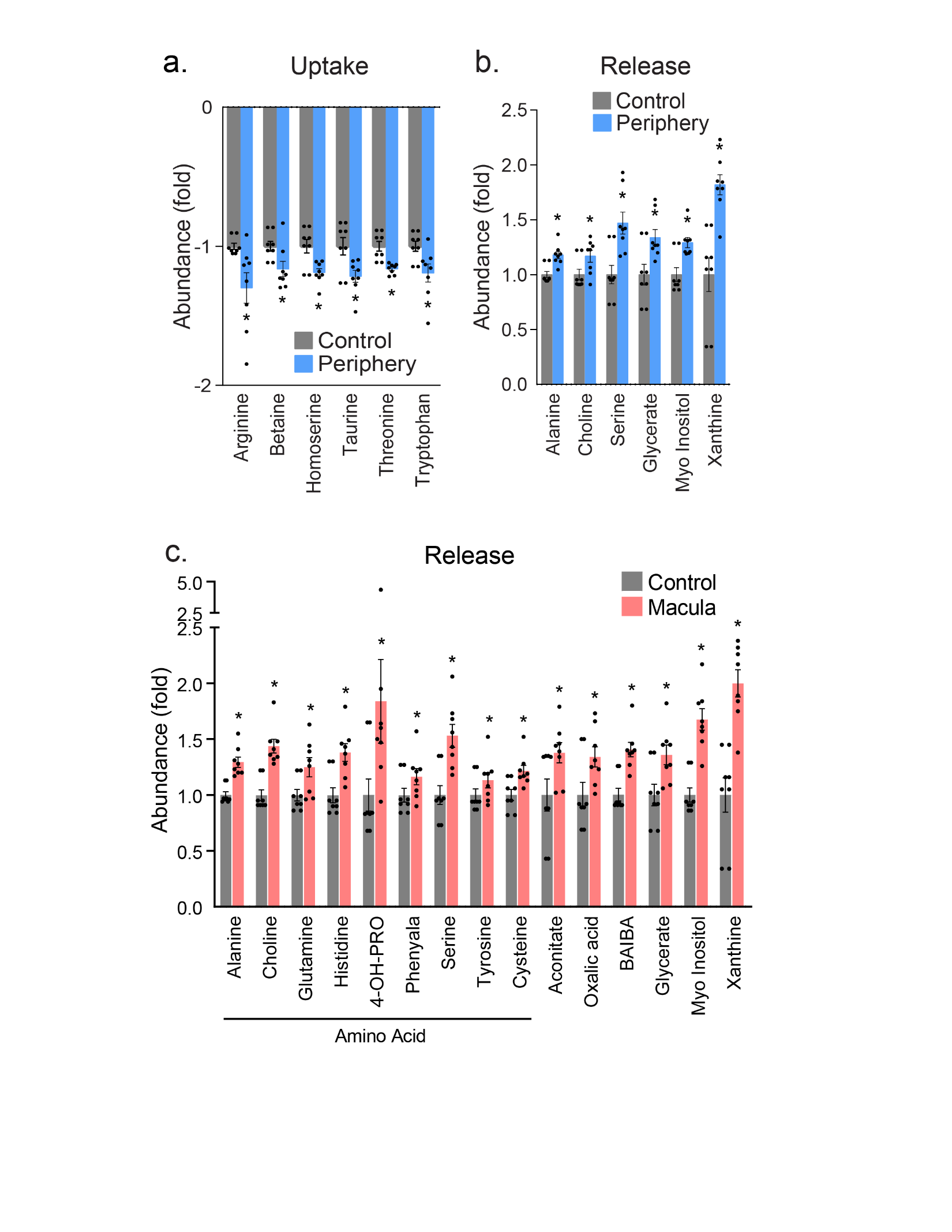

**Fig S7. Metabolite uptake and release in human macular and peripheral RPE/choroid. (a)** Uptaked metabolites in the medium from peripheral RPE/choroid culture. **(b)** Released metabolites in the medium from peripheral RPE/choroid culture. **(c).** Released metabolites in the medium from macular RPE/choroid culture. N=8, **P*<0.05 vs baseline control.

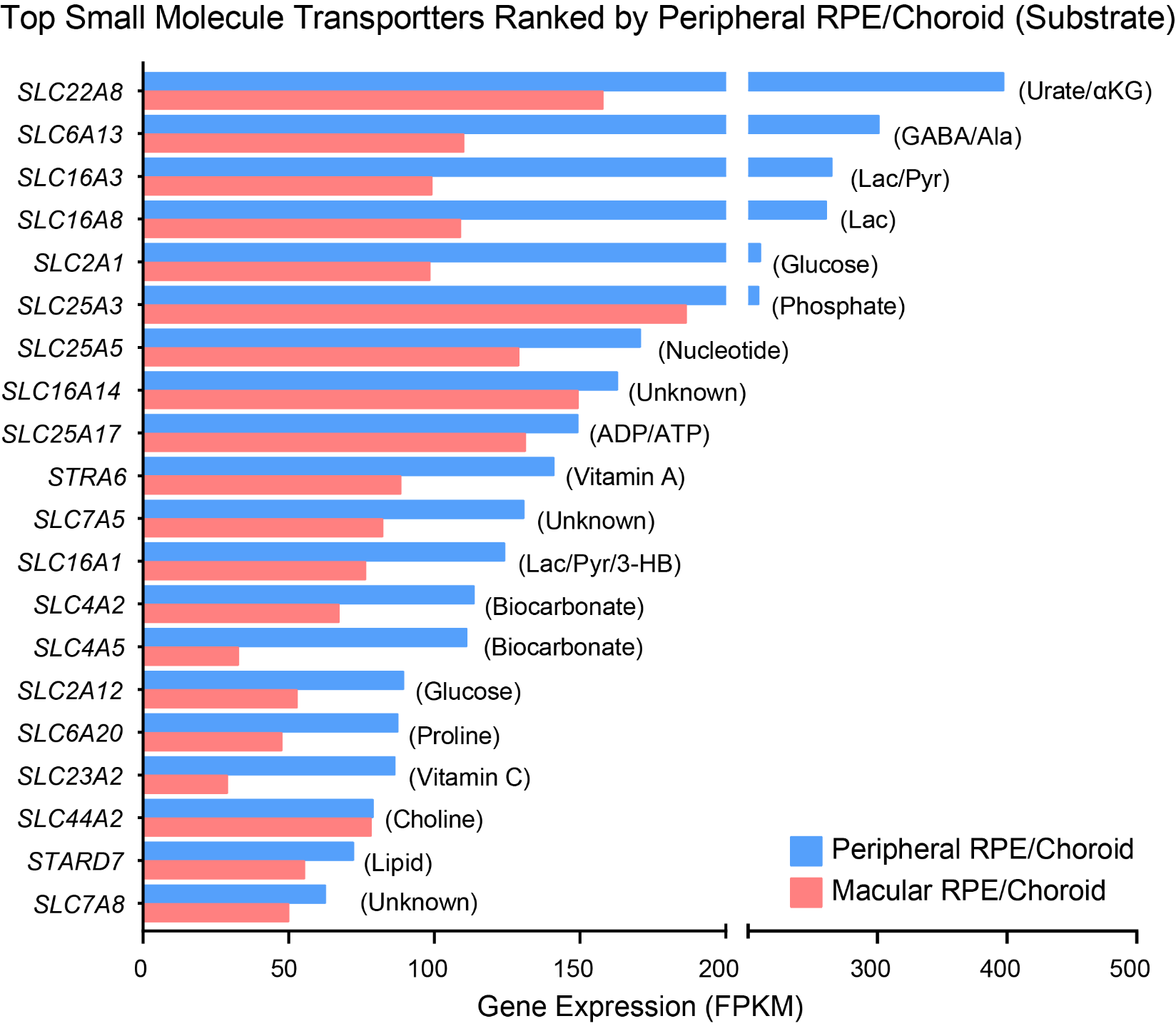

**Fig S8. Top small molecule transporters ranked by peripheral RPE/choroid.** Top 20 most abundant small molecule transporters ranked by the expression in human peripheral RPE/choroid. Periphery expression was the average of the expression in nasal and temporal regions.

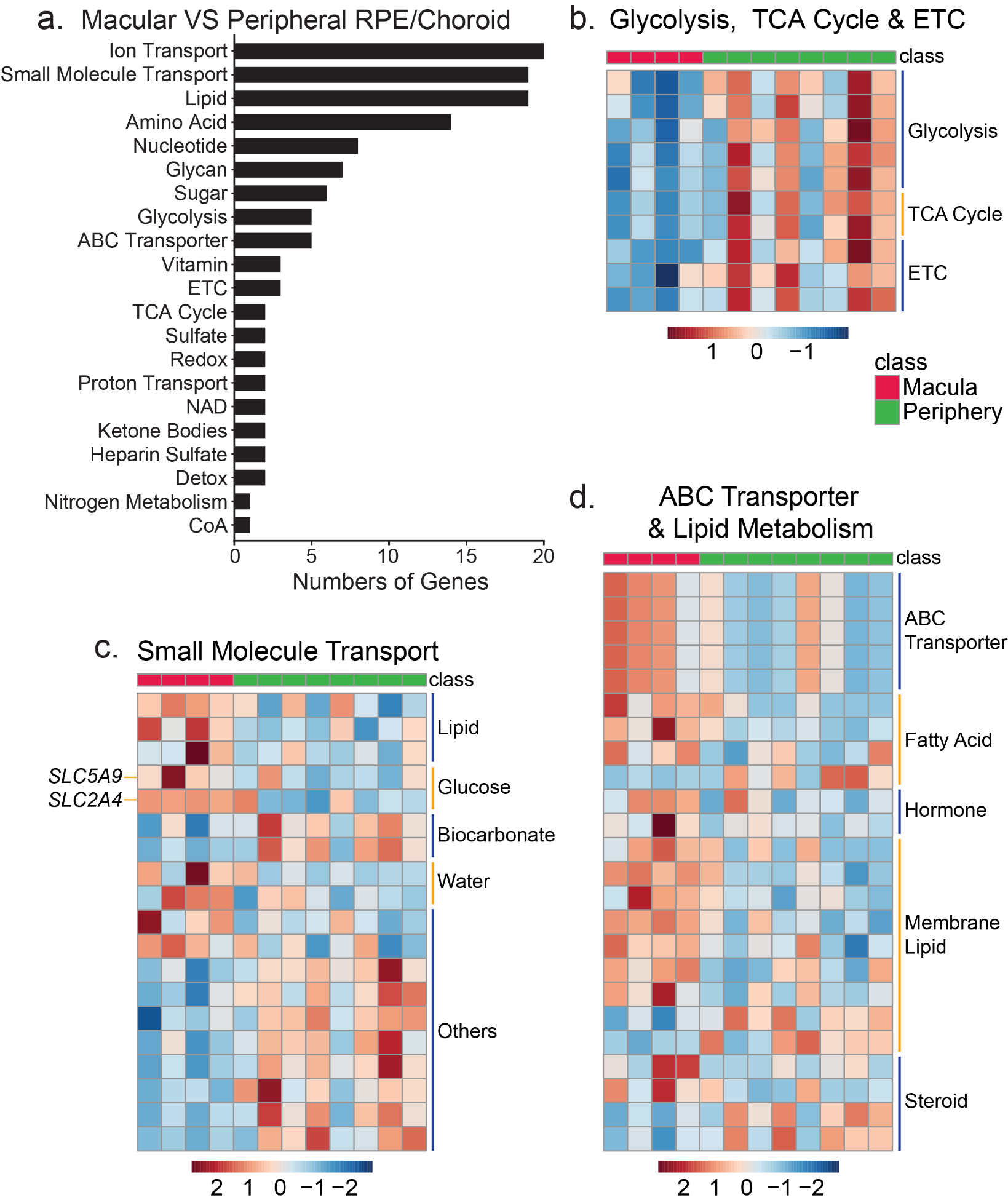

**Fig S9. Changed metabolic genes in the macular and peripheral RPE/choroid. (a)** Number of changed genes in metabolic pathways. **(b-c)** Heatmaps of differentially expressed genes in **(b)** glycolysis, TCA cycle & ETC, **(c)** small molecule transport, **(d)** ABC transporter & lipid.

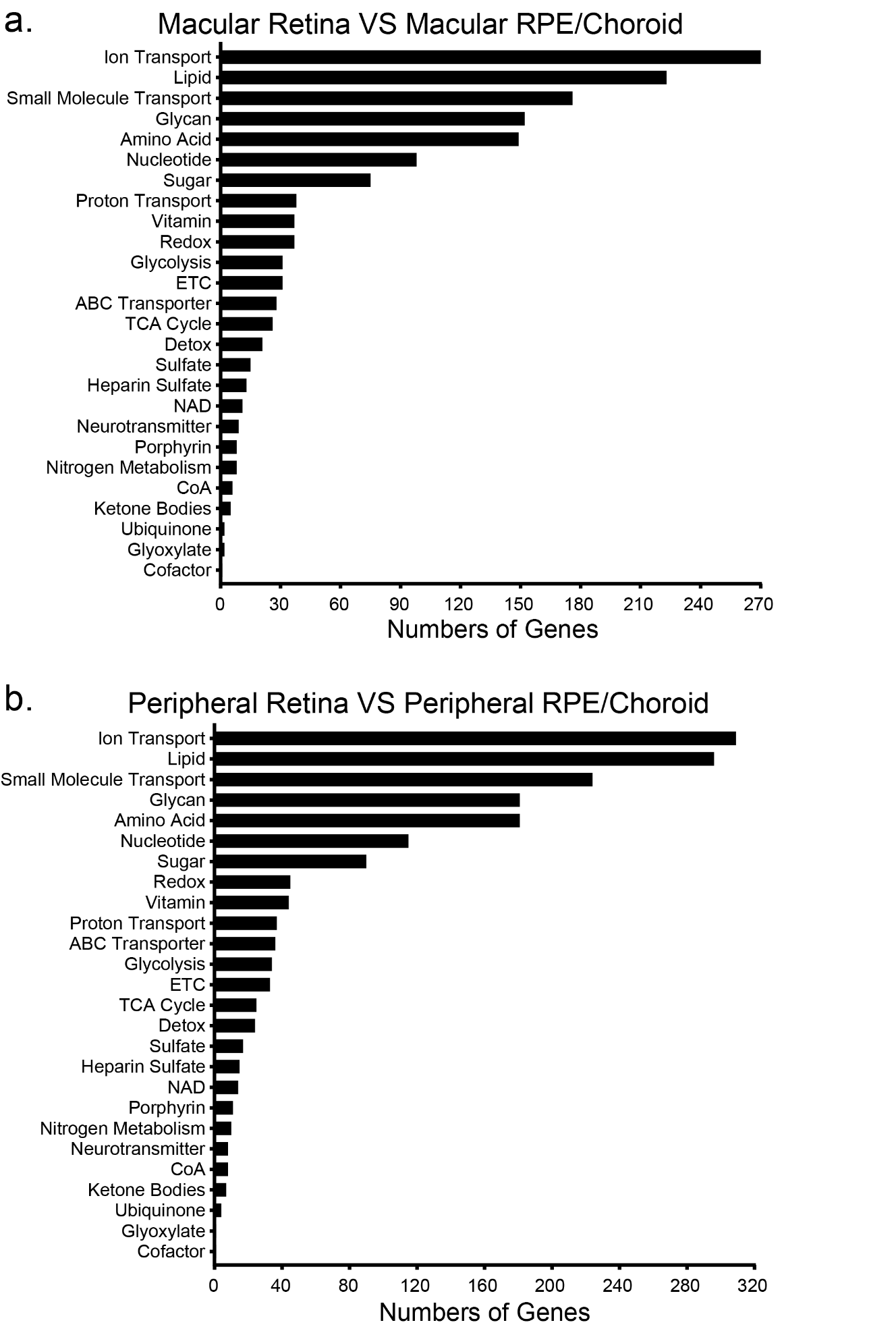

**Fig S10. Changed metabolic genes between human retinas and RPE/choroid. (a)** Number of changed genes in metabolic pathways in the macula. **(b).** Number of changed genes in metabolic pathways in the periphery.

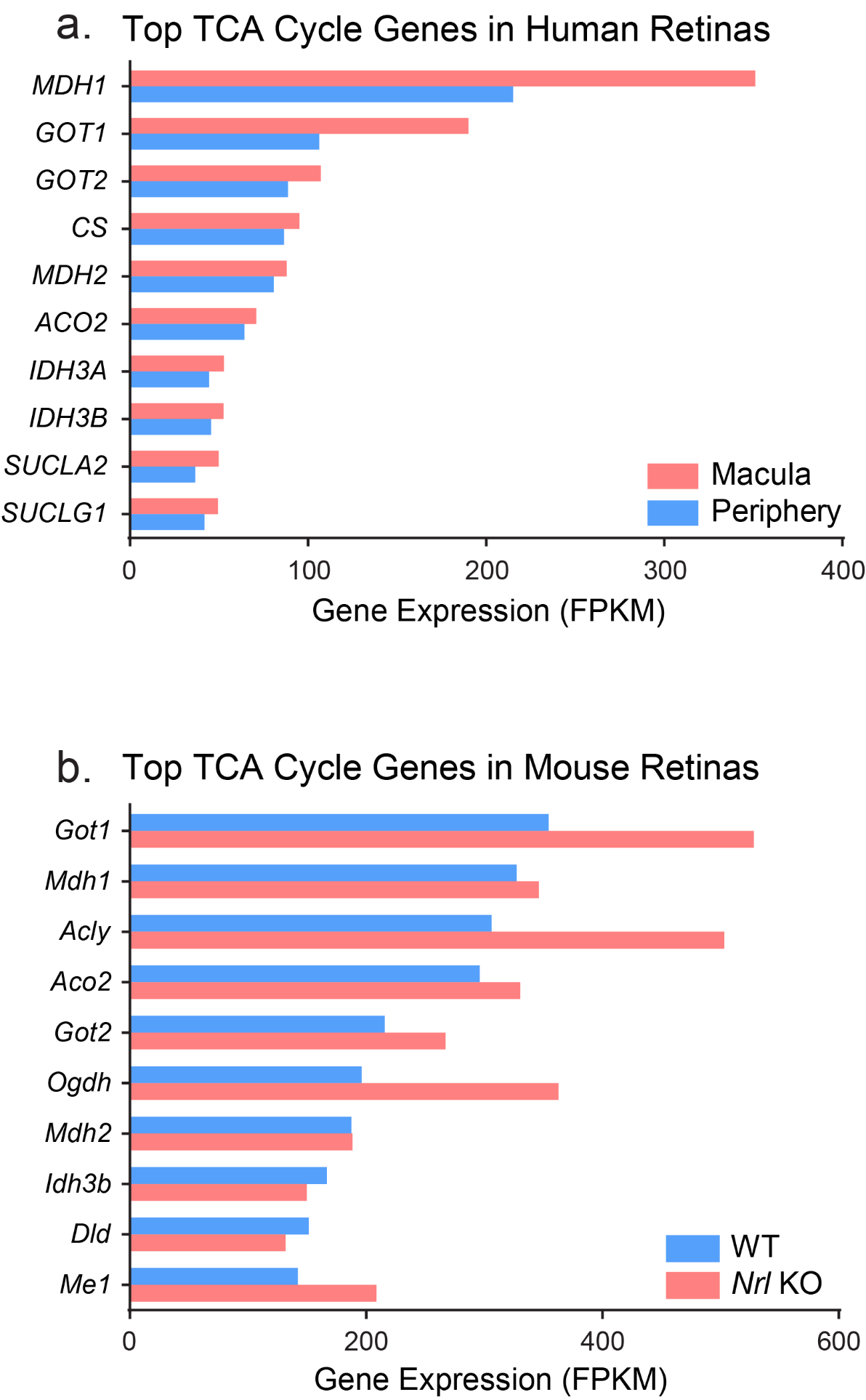

**Fig S11. The expression of TCA cycle genes in mouse and human retinas.**

**Table S1.** **List of metabolites and parameters by targeted metabolomics.**

| **Metabolite** | **CAS** | **MW** | **Positive/ Negative Mode** | **Q1 Mass (Da)** | **Q3 Mass (Da)** | **DP (Volts)** | **CE (Volts)** | **Platform** |
| --- | --- | --- | --- | --- | --- | --- | --- | --- |
| 3-Nitrotyrosine | 621-44-3 | 226.19 | Neg | 225.0 | 163.0 | -45 | -19 | LC MS |
| Hydroxykynurenine | 606-14-4 | 224.08 | Neg | 223.0 | 75.0 | -150 | -55 | LC MS |
| 4-Hydroxyphenylpyruvic acid | 156-39-8 | 180.16 | Neg | 179.1 | 97.1 | -40 | -17 | LC MS |
| 3-Hydroxypicolinic acid | 19621-92-2 | 139.03 | Neg | 138.1 | 94.1 | -83 | -19 | LC MS |
| cis-Aconitic acid | 499-12-7 | 174.02 | Neg | 173.0 | 85.0 | -37 | -18 | LC MS |
| Adenine | 73-24-5 | 135.05 | Neg | 134.0 | 107.0 | -92 | -25 | LC MS |
| Adipic acid | 124-04-9 | 146.06 | Neg | 145.0 | 83.0 | -50 | -17 | LC MS |
| ADP | 58-64-0 | 427.03 | Neg | 426.0 | 79.1 | -65 | -84 | LC MS |
| α-ketoglutarate | 328-50-7 | 146.11 | Neg | 145.0 | 101.0 | -56 | -11 | LC MS |
| Aminoadipic acid | 542-32-5 | 161.07 | Neg | 160.1 | 116.1 | -50 | -20 | LC MS |
| Adenosine monophosphate | 61-19-8 | 347.06 | Neg | 346.0 | 134.0 | -82 | -46 | LC MS |
| Anthranilate | 118-92-3 | 137.05 | Neg | 136.0 | 92.0 | -83 | -21 | LC MS |
| Ascorbic acid | 50-81-7 | 176.03 | Neg | 175.0 | 87.0 | -63 | -30 | LC MS |
| Azelaic acid | 123-99-9 | 188.22 | Neg | 187.0 | 97.1 | -73 | -25 | LC MS |
| Cyclic AMP | 60-92-4 | 329.05 | Neg | 328.0 | 134.0 | -87 | -32 | LC MS |
| Carbamoyl phosphate | 590-55-6 | 140.98 | Neg | 151.0 | 108.0 | -105 | -25 | LC MS |
| Cyclic GMP | 7665-99-8 | 345.05 | Neg | 344.0 | 150.0 | -37 | -30 | LC MS |
| Citraconic acid | 498-23-7 | 130.03 | Neg | 129.0 | 85.0 | -21 | -12 | LC MS |
| Citrulline | 372-75-8 | 175.10 | Neg | 174.1 | 131.1 | -32 | -20 | LC MS |
| L-Cystine | 56-89-3 | 240.02 | Neg | 239.0 | 74.0 | -50 | -25 | LC MS |
| Dihydroxyacetone phosphate | 57-04-5 | 170.00 | Neg | 169.0 | 79.0 | -38 | 37 | LC MS |
| dUMP | 964-26-1 | 308.04 | Neg | 307.0 | 79.0 | -68 | -66 | LC MS |
| D-Erythrose 4-phosphate | 585-18-2 | 200.01 | Neg | 199.0 | 97.0 | -160 | -15 | LC MS |
| FAD | 146-14-5 | 785.16 | Neg | 784.1 | 437.1 | 52 | 43 | LC MS |
| Flavin Mononucleotide | 6184-17-4 | 456.11 | Neg | 455.1 | 243.1 | 14 | 46 | LC MS |
| Folic acid | 59-30-3 | 441.14 | Neg | 440.1 | 311.0 | -132 | -30 | LC MS |
| D-Fructose | 57-48-7 | 180.06 | Neg | 179.1 | 71.0 | -120 | -28 | LC MS |
| Fructose 1,6-bisphosphate | 488-69-7 | 340.00 | Neg | 339.1 | 79.0 | -56 | -75 | LC MS |
| Glyceraldehyde 3-phosphate | 142-10-9 | 170.00 | Neg | 171.0 | 79.0 | -74 | -28 | LC MS |
| Glucose 6-phosphate | 56-73-5 | 260.14 | Neg | 259.0 | 79.0 | -46 | -62 | LC MS |
| Geranyl-PP | 763-10-0 | 314.21 | Neg | 313.1 | 185.1 | -58 | -45 | LC MS |
| Glucoronate | 6556-12-3 | 194.14 | Neg | 193.0 | 73.0 | -56 | -20 | LC MS |
| D-Glucose | 492-62-6 | 180.06 | Neg | 179.0 | 89.0 | -50 | -15 | LC MS |
| Glutaric acid | 110-94-1 | 132.04 | Neg | 131.0 | 87.0 | -45 | -16 | LC MS |
| Guanosine monophosphate | 85-32-5 | 363.06 | Neg | 362.1 | 79.0 | -41 | -61 | LC MS |
| Glutathione | 70-18-8 | 307.08 | Neg | 306.0 | 143.1 | -61 | -26 | LC MS |
| Oxidized Glutathione | 13081-14-6 | 426.09 | Neg | 611.0 | 306.0 | -67 | -34 | LC MS |
| Guanosine triphosphate | 86-01-1 | 522.99 | Neg | 521.9 | 159.0 | -39 | -32 | LC MS |
| Guanosine | 118-00-3 | 283.09 | Neg | 282.1 | 150.0 | -67 | -33 | LC MS |
| Heptadecanoic acid | 506-12-7 | 270.26 | Neg | 269.1 | 135.1 | -77 | -35 | LC MS |
| Hippuric acid | 495-69-2 | 179.06 | Neg | 178.0 | 77.1 | -61 | -23 | LC MS |
| Hypotaurine | 300-84-5 | 109.02 | Neg | 108.1 | 64.0 | -40 | -17 | LC MS |
| Hypoxanthine | 68-94-0 | 136.04 | Neg | 135.0 | 65.0 | -108 | -37 | LC MS |
| Inosinic acid | 131-99-7 | 348.05 | Neg | 347.0 | 79.0 | -118 | -86 | LC MS |
| Inosine | 58-63-9 | 268.08 | Neg | 267.0 | 135.0 | -123 | -30 | LC MS |
| Isopentenyl pyrophosphate | 18687-43-9 | 246.10 | Neg | 245.1 | 79.0 | 60 | 31 | LC MS |
| L-Kynurenine | 343-65-7 | 208.09 | Neg | 207.1 | 144.0 | -58 | -33 | LC MS |
| Lactate | 50-21-5 | 90.03 | Neg | 89.0 | 43.0 | -60 | -17 | LC MS |
| L-Malic acid | 6915-15-7 | 134.02 | Neg | 133.0 | 115.0 | -50 | -14 | LC MS |
| Maleic acid | 110-16-7 | 116.01 | Neg | 115.0 | 71.0 | -34 | -13 | LC MS |
| Malonic acid | 141-82-2 | 104.01 | Neg | 103.0 | 41.0 | -50 | -38 | LC MS |
| Melatonin | 73-31-4 | 232.12 | Neg | 231.1 | 58.0 | -75 | -37 | LC MS |
| Mevalonic acid | 150-97-0 | 148.07 | Neg | 147.1 | 77.0 | -50 | -20 | LC MS |
| myo-Inositol | 87-89-8 | 180.16 | Neg | 179.0 | 87.0 | -105 | -24 | LC MS |
| Acetylglycine | 543-24-8 | 117.10 | Neg | 116.0 | 74.0 | -37 | -14 | LC MS |
| NADH | 58-68-4 | 665.13 | Neg | 664.0 | 397.0 | -7 | -46 | LC MS |
| NADPH | 2646-71-1 | 745.09 | Neg | 744.0 | 408.0 | -35 | -52 | LC MS |
| Nicotinate | 56-67-6 | 122.10 | Neg | 122.0 | 78.0 | -105 | -18 | LC MS |
| Nicotinic acid | 59-67-6 | 123.03 | Neg | 122.0 | 78.0 | -50 | -15 | LC MS |
| Pantothenic acid | 137-08-6 | 219.11 | Neg | 218.1 | 71.0 | -70 | -41 | LC MS |
| Phenylpyruvic acid | 156-06-9 | 164.05 | Neg | 163.0 | 91.0 | -38 | -14 | LC MS |
| Phosphocreatine | 19333-65-4 | 255.08 | Neg | 210.0 | 79.0 | -42 | -49 | LC MS |
| Phosphoenolpyruvic acid | 138-08-9 | 166.98 | Neg | 167.1 | 79.0 | -38 | -20 | LC MS |
| Pyridoxal | 66-72-8 | 167.06 | Neg | 166.0 | 79.0 | -30 | -47 | LC MS |
| Pyridoxal 5'-phosphate | 54-47-7 | 247.14 | Neg | 246.0 | 79.1 | -61 | -61 | LC MS |
| Quinic acid | 77-95-2 | 192.06 | Neg | 191.1 | 85.0 | -60 | -33 | LC MS |
| Quinolinic acid | 89-00-9 | 167.02 | Neg | 166.0 | 78.0 | -53 | -20 | LC MS |
| Retinol | 68-26-8 | 286.23 | Neg | 285.2 | 79.0 | -79 | -18 | LC MS |
| D-Ribulose 5-phosphate | 551-85-9 | 230.02 | Neg | 229.0 | 79.0 | -70 | -45 | LC MS |
| Salicyluric acid | 69-72-7 | 138.02 | Neg | 137.0 | 93.0 | -100 | -38 | LC MS |
| Shikimic acid | 138-59-0 | 174.05 | Neg | 173.0 | 93.0 | -61 | -23 | LC MS |
| Succinic acid semialdehyde | 692-29-5 | 102.03 | Neg | 101.0 | 55.0 | -60 | -45 | LC MS |
| Xanthine | 69-89-6 | 152.03 | Neg | 151.0 | 108.0 | -70 | -23 | LC MS |
| Xanthosine | 146-80-5 | 284.08 | Neg | 283.1 | 151.0 | -92 | -28 | LC MS |
| Xanthurenic acid | 59-00-7 | 205.04 | Neg | 204.0 | 160.0 | -67 | -19 | LC MS |
| 1-Methyladenosine | 15763-06-1 | 281.11 | Pos | 282.1 | 150.1 | 48 | 35 | LC MS |
| N1-Methylnicotinamide | 3106-60-3 | 137.16 | Pos | 138.0 | 78.0 | 110 | 37 | LC MS |
| 2-Methylbutyroylcarnitine | 31023-25-3 | 245.32 | Pos | 246.2 | 85.1 | 100 | 43 | LC MS |
| 3-Aminoisobutanoic acid | 144-90-1 | 103.12 | Pos | 104.1 | 58.0 | 55 | 38 | LC MS |
| (R)-3-Hydroxybutyric acid | 300-85-6 | 104.05 | Pos | 105.1 | 45.0 | 20 | 47 | LC MS |
| 4-Pyridoxic acid | 82-82-6 | 183.05 | Pos | 182.1 | 138.0 | -59 | -20 | LC MS |
| p-Aminobenzoic acid | 99-96-7 | 137.05 | Pos | 138.1 | 120.0 | 63 | 24 | LC MS |
| 4-Hydroxyproline | 51-35-4 | 131.13 | Pos | 132.1 | 86.0 | 60 | 17 | LC MS |
| 5-Aminopentanoic acid | 660-88-8 | 117.08 | Pos | 118.1 | 55.1 | 57 | 24 | LC MS |
| Acetyl coenzyme A | 102029-73-2 | 809.13 | Pos | 810.1 | 303.1 | 116 | 46 | LC MS |
| N-Alpha-acetyllysine | 152473-69-3 | 188.22 | Pos | 189.1 | 84.1 | 50 | 32 | LC MS |
| N-acetyltryptophan | 87-32-1 | 246.26 | Pos | 247.2 | 146.0 | 75 | 35 | LC MS |
| N-Acetyl-L-alanine | 97-69-8 | 131.13 | Pos | 132.1 | 44.0 | 50 | 20 | LC MS |
| N-Acetylasparagine | 4033-40-3 | 174.15 | Pos | 175.1 | 70.1 | 55 | 24 | LC MS |
| N-Acetylglutamic acid | 1188-37-0 | 189.17 | Pos | 190.1 | 84.1 | 51 | 20 | LC MS |
| Acetylglycine | 543-24-8 | 117.10 | Pos | 118.1 | 30.2 | 30 | 16 | LC MS |
| N-Acetyl-L-methionine | 1115-47-5 | 191.25 | Pos | 192.1 | 61.0 | 27 | 43 | LC MS |
| N-Acetyl-L-phenylalanine | 2018-61-3 | 207.23 | Pos | 208.1 | 120.0 | 50 | 18 | LC MS |
| N-Acetylserine | 16354-58-8 | 147.13 | Pos | 148.1 | 60.0 | 40 | 22 | LC MS |
| Adenosine | 58-61-7 | 267.10 | Pos | 268.1 | 136.0 | 95 | 38 | LC MS |
| AICAR | 3031-94-5 | 338.06 | Pos | 259.1 | 111.0 | 85 | 34 | LC MS |
| L-Alanine | 56-41-7 | 89.05 | Pos | 90.0 | 44.0 | 50 | 20 | LC MS |
| Argininosuccinic acid | 2387-71-5 | 290.20 | Pos | 291.1 | 69.9 | 70 | 50 | LC MS |
| L-Aspartic acid | 56-84-8 | 133.04 | Pos | 134.1 | 74.0 | 50 | 20 | LC MS |
| Adenosine triphosphate | 987-65-5 | 507.00 | Pos | 507.9 | 136.1 | 14 | 60 | LC MS |
| Benzoic acid | 65-85-0 | 122.04 | Pos | 123.0 | 77.0 | 220 | 31 | LC MS |
| Betaine | 590-46-5 | 117.08 | Pos | 118.1 | 58.0 | 166 | 56 | LC MS |
| Butyrylcarnitine | 25576-40-3 | 231.15 | Pos | 232.2 | 85.1 | 72 | 30 | LC MS |
| Cadaverine | 462-94-2 | 102.12 | Pos | 103.0 | 77.0 | 190 | 25 | LC MS |
| L-Carnitine | 541-15-1 | 161.11 | Pos | 163.1 | 85.0 | 100 | 15 | LC MS |
| Carnosine | 305-84-0 | 226.11 | Pos | 227.1 | 110.1 | 157 | 32 | LC MS |
| Choline | 62-49-7 | 103.10 | Pos | 104.1 | 60.1 | 95 | 37 | LC MS |
| Coenzyme A | 55672-92-9 | 767.53 | Pos | 768.1 | 261.0 | 34 | 47 | LC MS |
| Creatine | 57-00-1 | 131.07 | Pos | 132.1 | 90.0 | 170 | 17 | LC MS |
| Creatinine | 60-27-5 | 113.06 | Pos | 114.0 | 44.0 | 71 | 33 | LC MS |
| Cyclic adenosine diphosphate-ribose | 119340-53-3 | 541.06 | Pos | 542.0 | 136.0 | 75 | 51 | LC MS |
| Cytidine | 65-46-3 | 243.09 | Pos | 244.0 | 112.1 | 50 | 30 | LC MS |
| Cytosine | 71-30-7 | 111.04 | Pos | 112.0 | 95.0 | 97 | 23 | LC MS |
| D-2-Hydroxyglutaric acid | 103404-90-6 | 148.11 | Pos | 149.1 | 77.0 | 100 | 43 | LC MS |
| Decanoyl-L-carnitine | 3992-45-8 | 315.45 | Pos | 316.3 | 85.1 | 70 | 25 | LC MS |
| Dehydroascorbic acid | 490-83-5 | 174.02 | Pos | 175.0 | 63.1 | 130 | 12 | LC MS |
| Dopamine | 51-61-6 | 153.08 | Pos | 154.0 | 91.0 | 40 | 36 | LC MS |
| Epinephrine | 51-43-4 | 183.09 | Pos | 184.1 | 77.1 | 50 | 52 | LC MS |
| Sphinganine | 764-22-7 | 301.51 | Pos | 302.3 | 81.0 | 160 | 47 | LC MS |
| Erythritol | 149-32-6 | 122.06 | Pos | 123.0 | 80.0 | 40 | 33 | LC MS |
| Glucose 1-phosphate | 59-56-3 | 260.03 | Pos | 259.0 | 79.0 | -85 | -60 | LC MS |
| Gamma-Aminobutyric acid | 56-12-2 | 103.12 | Pos | 104.1 | 87.0 | 57 | 13 | LC MS |
| Guanosine diphosphate | 43139-22-6 | 443.02 | Pos | 444.0 | 152.0 | 65 | 26 | LC MS |
| Glutamax | 39537-23-0 | 217.22 | Pos | 218.1 | 84.0 | 50 | 20 | LC MS |
| L-Glutamic acid | 56-86-0 | 147.05 | Pos | 148.1 | 84.1 | 51 | 20 | LC MS |
| L-Glutamine | 56-85-9 | 146.07 | Pos | 147.1 | 84.1 | 45 | 23 | LC MS |
| Glyceric acid | 14028-62-7 | 250.00 | Pos | 107.0 | 45.0 | 66 | 16 | LC MS |
| Glycerol | 56-81-5 | 92.05 | Pos | 93.0 | 51.0 | 160 | 36 | LC MS |
| Glycine | 56-40-6 | 75.03 | Pos | 76.0 | 30.2 | 30 | 16 | LC MS |
| Guanine | 73-40-5 | 151.05 | Pos | 152.0 | 110.0 | 20 | 28 | LC MS |
| Hexanoylcarnitine | 22671-29-0 | 259.18 | Pos | 260.2 | 85.1 | 80 | 25 | LC MS |
| Histamine | 51-45-6 | 111.08 | Pos | 112.0 | 95.0 | 65 | 18 | LC MS |
| L-Histidine | 332-80-9 | 155.07 | Pos | 156.1 | 110.0 | 50 | 22 | LC MS |
| Homocysteine | 6027-13-0 | 135.04 | Pos | 136.0 | 91.0 | 135 | 24 | LC MS |
| Isobutyryl-L-carnitine | 25518-49-4 | 231.15 | Pos | 232.1 | 85.1 | 70 | 30 | LC MS |
| L-Acetylcarnitine | 14992-62-2 | 203.24 | Pos | 204.1 | 85.0 | 71 | 30 | LC MS |
| L-Arginine | 74-79-3 | 174.11 | Pos | 175.1 | 70.1 | 45 | 33 | LC MS |
| L-Asparagine | 70-47-3 | 132.05 | Pos | 133.1 | 70.1 | 55 | 24 | LC MS |
| L-Leucine | 61-90-5 | 131.10 | Pos | 132.1 | 86.0 | 60 | 16 | LC MS |
| L-Homoserine | 1927-25-9 | 119.06 | Pos | 120.1 | 74.0 | 37 | 16 | LC MS |
| Linoleic acid | 60-33-3 | 280.24 | Pos | 281.1 | 221.0 | 60 | 13 | LC MS |
| L-Lysine | 56-87-1 | 146.11 | Pos | 147.1 | 84.0 | 38 | 22 | LC MS |
| Malonyl coenzyme A | 108347-84-8 | 853.58 | Pos | 854.1 | 347.1 | 75 | 45 | LC MS |
| L-Methionine | 63-68-3 | 149.05 | Pos | 150.1 | 61.0 | 27 | 43 | LC MS |
| Methylmalonic acid | 516-05-2 | 118.03 | Pos | 119.0 | 65.0 | 70 | 37 | LC MS |
| Myristoyl-L-carnitine | 25597-07-3 | 371.30 | Pos | 372.4 | 85.1 | 70 | 25 | LC MS |
| N-Acetylglutamic acid | 1188-37-0 | 189.17 | Pos | 190.0 | 102.0 | 20 | 23 | LC MS |
| N-Acetylputrescine | 5699-41-2 | 130.11 | Pos | 131.1 | 114.0 | 64 | 30 | LC MS |
| NAD | 53-84-9 | 663.11 | Pos | 664.0 | 136.0 | 27 | 48 | LC MS |
| NADP | 53-59-8 | 743.08 | Pos | 744.0 | 136.0 | 9 | 78 | LC MS |
| Nicotinic acid mononucleotide | 321-02-8 | 335.04 | Pos | 336.1 | 124.0 | 75 | 17 | LC MS |
| Niacinamide | 98-92-0 | 122.05 | Pos | 123.0 | 80.0 | 136 | 20 | LC MS |
| β-Nicotinamide adenine dinucleotide | 20111-18-6 | 685.41 | pos | 665.0 | 136.1 | 95 | 63 | LC MS |
| Nicotinic acid N-oxide | 2398-81-4 | 139.11 | Pos | 139.0 | 65.1 | 121 | 36 | LC MS |
| Nicotinamide ribotide | 1094-61-7 | 334.06 | Pos | 335.0 | 123.1 | 40 | 25 | LC MS |
| Nicotinamide riboside | 2181-04-6 | 358.21 | Pos | 256.1 | 123.0 | 68 | 18 | LC MS |
| L-Octanoylcarnitine | 25243-95-2 | 287.21 | Pos | 288.3 | 85.1 | 100 | 25 | LC MS |
| Ophthalmic acid | 495-27-2 | 289.13 | Pos | 290.1 | 58.0 | 139 | 56 | LC MS |
| Palmitoleic acid | 373-49-9 | 254.23 | Pos | 255.1 | 195.0 | 60 | 10 | LC MS |
| L-Palmitoylcarnitine | 1985-18-8 | 399.62 | Pos | 400.4 | 85.1 | 30 | 40 | LC MS |
| Palmityl-CoA | 188174-64-3 | 10004.94 | Pos | 1006.3 | 499.5 | 45 | 53 | LC MS |
| L-Phenylalanine | 63-91-2 | 165.08 | Pos | 166.1 | 120.0 | 50 | 18 | LC MS |
| Phosphoserine | 407-41-0 | 185.07 | Pos | 186.0 | 70.0 | 50 | 26 | LC MS |
| Pipecolic acid | 535-75-1 | 129.08 | Pos | 130.0 | 84.1 | 58 | 32 | LC MS |
| L-Proline | 147-85-3 | 129.08 | Pos | 116.1 | 70.1 | 70 | 44 | LC MS |
| Propionic acid | 79-09-4 | 74.04 | Pos | 75.0 | 57.0 | 29 | 15 | LC MS |
| Propionylcarnitine | 17298-37-2 | 217.13 | Pos | 218.1 | 85.1 | 60 | 24 | LC MS |
| Pyridoxamine | 85-87-0 | 168.09 | Pos | 169.1 | 134.1 | 20 | 36 | LC MS |
| Pyridoxine | 65-23-6 | 169.07 | Pos | 170.1 | 134.0 | 40 | 29 | LC MS |
| Pyroglutamic acid | 98-79-3 | 129.04 | Pos | 130.1 | 84.0 | 100 | 20 | LC MS |
| Riboflavin | 83-88-5 | 376.14 | Pos | 377.1 | 243.1 | 14 | 29 | LC MS |
| S-Adenosylmethionine | 86867-01-8 | 398.44 | Pos | 399.3 | 136.0 | 60 | 30 | LC MS |
| Sarcosine | 107-97-1 | 89.09 | Pos | 90.0 | 44.0 | 38 | 17 | LC MS |
| L-Serine | 56-45-1 | 105.04 | Pos | 106.0 | 60.0 | 40 | 22 | LC MS |
| Spermidine | 124-20-9 | 145.16 | Pos | 146.1 | 72.0 | 60 | 19 | LC MS |
| Spermine | 71-44-3 | 202.22 | Pos | 203.1 | 129.0 | 33 | 26 | LC MS |
| Stearoylcarnitine | 25597-09-5 | 427.37 | Pos | 428.5 | 85.1 | 70 | 25 | LC MS |
| Succinyl-CoA | 108347-97-3 | 867.61 | Pos | 868.1 | 361.2 | 45 | 41 | LC MS |
| Taurine | 107-35-7 | 125.02 | Pos | 126.0 | 108.0 | 200 | 15 | LC MS |
| Thiamine | 59-43-8 | 264.10 | pos | 265.0 | 122.1 | 67 | 40 | LC MS |
| L-Threonine | 72-19-5 | 119.06 | Pos | 120.1 | 102.0 | 50 | 10 | LC MS |
| Trigonelline | 535-83-1 | 137.05 | Pos | 138.0 | 92.0 | 60 | 27 | LC MS |
| Tryptamine | 61-54-1 | 160.10 | Pos | 161.1 | 144.0 | 58 | 15 | LC MS |
| L-Tryptophan | 73-22-3 | 204.09 | Pos | 205.0 | 146.0 | 75 | 35 | LC MS |
| Tyramine | 51-67-2 | 137.08 | Pos | 138.0 | 77.1 | 58 | 41 | LC MS |
| L-Tyrosine | 60-18-4 | 181.07 | Pos | 182.1 | 136.0 | 40 | 17 | LC MS |
| Uridine diphosphate glucose | 133-89-1 | 566.30 | Pos | 611.0 | 499.0 | 40 | 29 | LC MS |
| Uridine | 58-96-8 | 244.07 | Pos | 245.0 | 113.0 | 23 | 51 | LC MS |
| L-Valine | 72-18-4 | 117.08 | Pos | 118.1 | 72.0 | 60 | 14 | LC MS |
| Vitamin A | 68-26-8 | 286.23 | Pos | 287.1 | 247.0 | 60 | 16 | LC MS |
| 2-hydroxyglutarate | ‎2889-31-8 | 148.11 | Pos | 433.0 | NA | NA | NA | GC MS |
| 3-Phosphoglyceric acid | 80731-10-8 | 186.06 | Pos | 585.0 | NA | NA | NA | GC MS |
| Acetoacetic acid | 3483-11-2 | 102.09 | Pos | 188.0 | NA | NA | NA | GC MS |
| Adipic acid | 124-04-9 | 146.14 | Pos | 317.0 | NA | NA | NA | GC MS |
| beta-hydroxybutyrate | 300-85-6 | 104.10 | Pos | 275.0 | NA | NA | NA | GC MS |
| Biotin | 58-85-5 | 244.31 | Pos | 529.4 | NA | NA | NA | GC MS |
| Cholesterol | 57-88-5 | 386.65 | Pos | 443.0 | NA | NA | NA | GC MS |
| Citrate | 77-92-9 | 192.12 | Pos | 591.0 | NA | NA | NA | GC MS |
| Cystathionine | 56-88-2 | 222.00 | Pos | 302.0 | NA | NA | NA | GC MS |
| Cysteine | 56-89-3 | 121.16 | Pos | 406.0 | NA | NA | NA | GC MS |
| Diethylene glycol | 111-46-6 | 106.12 | Pos | 159.1 | NA | NA | NA | GC MS |
| Fumarate | 110-17-8 | 116.07 | Pos | 287.0 | NA | NA | NA | GC MS |
| Glycerate | 6000-40-4 | 106.08 | Pos | 391.0 | NA | NA | NA | GC MS |
| Glycerol | 56-81-5 | 92.09 | Pos | 377.0 | NA | NA | NA | GC MS |
| Isocitrate | 1637-73-6 | 192.12 | Pos | 591.0 | NA | NA | NA | GC MS |
| Isoleucine | 73-32-5 | 131.17 | Pos | 200.0 | NA | NA | NA | GC MS |
| Malate | 6915-15-7 | 134.09 | Pos | 419.0 | NA | NA | NA | GC MS |
| N-Acetylaspartic acid | 997-55-7 | 175.14 | Pos | 460.0 | NA | NA | NA | GC MS |
| Ornithine | 70-26-8 | 132.16 | Pos | 286.0 | NA | NA | NA | GC MS |
| Oxalic acid | 144-62-7 | 90.09 | Pos | 261.0 | NA | NA | NA | GC MS |
| Oxaloacetate | 328-42-7 | 132.07 | Pos | 332.0 | NA | NA | NA | GC MS |
| Phosphoenolpyruvic acid | 5541-93-5 | 168.04 | Pos | 453.0 | NA | NA | NA | GC MS |
| Proline | 147-85-3 | 115.13 | Pos | 184.0 | NA | NA | NA | GC MS |
| Pyruvate | 113-24-6 | 88.06 | Pos | 174.0 | NA | NA | NA | GC MS |
| Quinolinic acid | 89-00-9 | 167.00 | Pos | 338.0 | NA | NA | NA | GC MS |
| Succinate | 110-15-6 | 118.09 | Pos | 289.0 | NA | NA | NA | GC MS |
| Uracil | 66-22-8 | 112.03 | Pos | 283.3 | NA | NA | NA | GC MS |
| Urea | 57-13-6 | 60.06 | Pos | 231.0 | NA | NA | NA | GC MS |
| Beta alanine | 107-95-9 | 89.09 | Pos | 260.2 | NA | NA | NA | GC MS |

**Table S2.** **List of 2764 classic metabolism genes and their classifications.**

| **Number** | **Gene symbol** | **Gene name** | **Entrez Gene ID** | **Classification (Sub-category/substrate)** |
| --- | --- | --- | --- | --- |
| 1 | ABCA1 | ATP-binding cassette, sub-family A (ABC1), member 1 | 19 | ABC Transporter |
| 2 | ABCA10 | ATP-binding cassette, sub-family A (ABC1), member 10 | 10349 | ABC Transporter |
| 3 | ABCA12 | ATP-binding cassette, sub-family A (ABC1), member 12 | 26154 | ABC Transporter |
| 4 | ABCA13 | ATP-binding cassette, sub-family A (ABC1), member 13 | 154664 | ABC Transporter |
| 5 | ABCA2 | ATP-binding cassette, sub-family A (ABC1), member 2 | 20 | ABC Transporter |
| 6 | ABCA3 | ATP-binding cassette, sub-family A (ABC1), member 3 | 21 | ABC Transporter |
| 7 | ABCA4 | ATP-binding cassette, sub-family A (ABC1), member 4 | 24 | ABC Transporter |
| 8 | ABCA5 | ATP-binding cassette, sub-family A (ABC1), member 5 | 23461 | ABC Transporter |
| 9 | ABCA6 | ATP-binding cassette, sub-family A (ABC1), member 6 | 23460 | ABC Transporter |
| 10 | ABCA7 | ATP-binding cassette, sub-family A (ABC1), member 7 | 10347 | ABC Transporter |
| 11 | ABCA8 | ATP-binding cassette, sub-family A (ABC1), member 8 | 10351 | ABC Transporter |
| 12 | ABCA9 | ATP-binding cassette, sub-family A (ABC1), member 9 | 10350 | ABC Transporter |
| 13 | ABCB1 | ATP-binding cassette, sub-family B (MDR/TAP), member 1 | 5243 | ABC Transporter |
| 14 | ABCB10 | ATP-binding cassette, sub-family B (MDR/TAP), member 10 | 23456 | ABC Transporter |
| 15 | ABCB11 | ATP-binding cassette, sub-family B (MDR/TAP), member 11 | 8647 | ABC Transporter |
| 16 | ABCB4 | ATP-binding cassette, sub-family B (MDR/TAP), member 4 | 5244 | ABC Transporter |
| 17 | ABCB5 | ATP-binding cassette, sub-family B (MDR/TAP), member 5 | 340273 | ABC Transporter |
| 18 | ABCB6 | ATP-binding cassette, sub-family B (MDR/TAP), member 6 | 10058 | ABC Transporter |
| 19 | ABCB7 | ATP-binding cassette, sub-family B (MDR/TAP), member 7 | 22 | ABC Transporter |
| 20 | ABCB8 | ATP-binding cassette, sub-family B (MDR/TAP), member 8 | 11194 | ABC Transporter |
| 21 | ABCB9 | ATP-binding cassette, sub-family B (MDR/TAP), member 9 | 23457 | ABC Transporter |
| 22 | ABCC1 | ATP-binding cassette, sub-family C (CFTR/MRP), member 1 | 4363 | ABC Transporter |
| 23 | ABCC10 | ATP-binding cassette, sub-family C (CFTR/MRP), member 10 | 89845 | ABC Transporter |
| 24 | ABCC11 | ATP-binding cassette, sub-family C (CFTR/MRP), member 11 | 85320 | ABC Transporter |
| 25 | ABCC12 | ATP-binding cassette, sub-family C (CFTR/MRP), member 12 | 94160 | ABC Transporter |
| 26 | ABCC13 | ATP-binding cassette, sub-family C (CFTR/MRP), member 13 | 150000 | ABC Transporter |
| 27 | ABCC2 | ATP-binding cassette, sub-family C (CFTR/MRP), member 2 | 1244 | ABC Transporter |
| 28 | ABCC3 | ATP-binding cassette, sub-family C (CFTR/MRP), member 3 | 8714 | ABC Transporter |
| 29 | ABCC4 | ATP-binding cassette, sub-family C (CFTR/MRP), member 4 | 10257 | ABC Transporter |
| 30 | ABCC5 | ATP-binding cassette, sub-family C (CFTR/MRP), member 5 | 10057 | ABC Transporter |
| 31 | ABCC6 | ATP-binding cassette, sub-family C (CFTR/MRP), member 6 | 368 | ABC Transporter |
| 32 | ABCC8 | ATP-binding cassette, sub-family C (CFTR/MRP), member 8 | 6833 | ABC Transporter |
| 33 | ABCC9 | ATP-binding cassette, sub-family C (CFTR/MRP), member 9 | 10060 | ABC Transporter |
| 34 | ABCD1 | ATP-binding cassette, sub-family D (ALD), member 1 | 215 | ABC Transporter |
| 35 | ABCD2 | ATP-binding cassette, sub-family D (ALD), member 2 | 225 | ABC Transporter |
| 36 | ABCD3 | ATP-binding cassette, sub-family D (ALD), member 3 | 5825 | ABC Transporter |
| 37 | ABCD4 | ATP-binding cassette, sub-family D (ALD), member 4 | 5826 | ABC Transporter |
| 38 | ABCE1 | ATP-binding cassette, sub-family E (OABP), member 1 | 6059 | ABC Transporter |
| 39 | ABCF1 | ATP-binding cassette, sub-family F (GCN20), member 1 | 23 | ABC Transporter |
| 40 | ABCF2 | ATP-binding cassette, sub-family F (GCN20), member 2 | 10061 | ABC Transporter |
| 41 | ABCF3 | ATP-binding cassette, sub-family F (GCN20), member 3 | 55324 | ABC Transporter |
| 42 | ABCG1 | ATP-binding cassette, sub-family G (WHITE), member 1 | 9619 | ABC Transporter |
| 43 | ABCG2 | ATP-binding cassette, sub-family G (WHITE), member 2 | 9429 | ABC Transporter |
| 44 | ABCG4 | ATP-binding cassette, sub-family G (WHITE), member 4 | 64137 | ABC Transporter |
| 45 | ABCG5 | ATP-binding cassette, sub-family G (WHITE), member 5 (sterolin 1) | 64240 | ABC Transporter |
| 46 | ABCG8 | ATP-binding cassette, sub-family G (WHITE), member 8 (sterolin 2) | 64241 | ABC Transporter |
| 47 | PAOX | polyamine oxidase (exo-N4-amino) | 196743 | Amino Acid (Polyamine) |
| 48 | SAT1 | spermidine/spermine N1-acetyltransferase 1 | 6303 | Amino Acid (Polyamine) |
| 49 | SAT2 | spermidine/spermine N1-acetyltransferase family member 2 | 112483 | Amino Acid (Polyamine) |
| 50 | SATL1 | spermidine/spermine N1-acetyl transferase-like 1 | 340562 | Amino Acid (Polyamine) |
| 51 | SMOX | spermine oxidase | 54498 | Amino Acid (Polyamine) |
| 52 | SMS | spermine synthase | 6611 | Amino Acid (Polyamine) |
| 53 | SRM | spermidine synthase | 6723 | Amino Acid (Polyamine) |
| 54 | MAK10 | MAK10 homolog, amino-acid N-acetyltransferase subunit (S. cerevisiae) | 60560 | Amino Acid (Acetylation) |
| 55 | NAT1 | N-acetyltransferase 1 (arylamine N-acetyltransferase) | 9 | Amino Acid (Acetylation) |
| 56 | NAT11 | N-acetyltransferase 11 | 79829 | Amino Acid (Acetylation) |
| 57 | NAT12 | N-acetyltransferase 12 | 122830 | Amino Acid (Acetylation) |
| 58 | NAT13 | N-acetyltransferase 13 | 80218 | Amino Acid (Acetylation) |
| 59 | NAT14 | N-acetyltransferase 14 | 57106 | Amino Acid (Acetylation) |
| 60 | NAT15 | N-acetyltransferase 15 | 79903 | Amino Acid (Acetylation) |
| 61 | NAT2 | N-acetyltransferase 2 (arylamine N-acetyltransferase) | 10 | Amino Acid (Acetylation) |
| 62 | NAT5 | N-acetyltransferase 5 | 51126 | Amino Acid (Acetylation) |
| 63 | NAT6 | N-acetyltransferase 6 | 24142 | Amino Acid (Acetylation) |
| 64 | NAT8 | N-acetyltransferase 8 | 9027 | Amino Acid (Acetylation) |
| 65 | NAT8B | N-acetyltransferase 8B | 51471 | Amino Acid (Acetylation) |
| 66 | NAT8L | N-acetyltransferase 8-like | 339983 | Amino Acid (Acetylation) |
| 67 | NAT9 | N-acetyltransferase 9 | 26151 | Amino Acid (Acetylation) |
| 68 | NAT10 | N-acetyltransferase 10 | 55226 | Amino Acid (Acetylation) |
| 69 | AGPS | alkylglycerone phosphate synthase | 8540 | Amino Acid (Alanine) |
| 70 | AGXT | alanine-glyoxylate aminotransferase | 189 | Amino Acid (Alanine) |
| 71 | AGXT2 | alanine-glyoxylate aminotransferase 2 | 64902 | Amino Acid (Alanine) |
| 72 | AGXT2L1 | alanine-glyoxylate aminotransferase 2-like 1 | 64850 | Amino Acid (Alanine) |
| 73 | AGXT2L2 | alanine-glyoxylate aminotransferase 2-like 2 | 85007 | Amino Acid (Alanine) |
| 74 | DAO | D-amino-acid oxidase | 1610 | Amino Acid (Alanine) |
| 75 | ADC | arginine decarboxylase | 113451 | Amino Acid (Arginine) |
| 76 | NAGS | N-acetylglutamate synthase | 162417 | Amino Acid (Arginine) |
| 77 | AOX1 | aldehyde oxidase 1 | 316 | Amino Acid (Arginine) |
| 78 | NOS1 | nitric oxide synthase 1 (neuronal) | 4842 | Amino Acid (Arginine) |
| 79 | NOS2 | Nitric oxide synthase 2 (inducible, hepatocytes) | 4843 | Amino Acid (Arginine) |
| 80 | NOS3 | nitric oxide synthase 3 (endothelial cell) | 4846 | Amino Acid (Arginine) |
| 81 | MOSC1 | MOCO sulphurase C-terminal domain containing 1 | 64757 | Amino Acid (Arginine) |
| 82 | MOSC2 | MOCO sulphurase C-terminal domain containing 2 | 54996 | Amino Acid (Arginine) |
| 83 | DDAH1 | dimethylarginine dimethylaminohydrolase 1 | 23576 | Amino Acid (Arginine, Citrulline) |
| 84 | DDAH2 | dimethylarginine dimethylaminohydrolase 2 | 23564 | Amino Acid (Arginine, Citrulline) |
| 85 | ARG1 | Arginase, liver | 383 | Amino Acid (Arginine, Urea) |
| 86 | ARG2 | arginase, type II | 384 | Amino Acid (Arginine, Urea) |
| 87 | ASL | argininosuccinate lyase | 435 | Amino Acid (Arginine, Urea) |
| 88 | ASS1 | argininosuccinate synthetase 1 | 445 | Amino Acid (Arginine, Urea) |
| 89 | OTC | ornithine carbamoyltransferase | 5009 | Amino Acid (Arginine, Urea) |
| 90 | ASNS | Asparagine synthetase | 440 | Amino Acid (Asparagine) |
| 91 | ASNSD1 | asparagine synthetase domain containing 1 | 54529 | Amino Acid (Asparagine) |
| 92 | ASPG | asparaginase homolog (S. cerevisiae) | 374569 | Amino Acid (Asparagine) |
| 93 | ASRGL1 | asparaginase like 1 | 80150 | Amino Acid (Asparagine) |
| 94 | NIT2 | nitrilase family, member 2 | 56954 | Amino Acid (Asparagine) |
| 95 | ACY3 | aspartoacylase (aminocyclase) 3 | 91703 | Amino Acid (Aspartate) |
| 96 | ASPA | aspartoacylase (Canavan disease) | 443 | Amino Acid (Aspartate) |
| 97 | DDO | D-aspartate oxidase | 8528 | Amino Acid (Aspartate) |
| 98 | BCAT1 | branched chain aminotransferase 1, cytosolic | 586 | Amino Acid (BCAAs) |
| 99 | BCAT2 | branched chain aminotransferase 2, mitochondrial | 587 | Amino Acid (BCAAs) |
| 100 | BCKDHA | branched chain keto acid dehydrogenase E1, alpha polypeptide | 593 | Amino Acid (BCAAs) |
| 101 | BCKDHB | branched chain keto acid dehydrogenase E1, beta polypeptide (maple syrup urine disease) | 594 | Amino Acid (BCAAs) |
| 102 | DBT | dihydrolipoamide branched chain transacylase E2 | 1629 | Amino Acid (BCAAs) |
| 103 | HIBADH | 3-hydroxyisobutyrate dehydrogenase | 11112 | Amino Acid (BCAAs) |
| 104 | HIBCH | 3-hydroxyisobutyryl-Coenzyme A hydrolase | 26275 | Amino Acid (BCAAs) |
| 105 | IVD | isovaleryl Coenzyme A dehydrogenase | 3712 | Amino Acid (BCAAs) |
| 106 | MCCC1 | methylcrotonoyl-Coenzyme A carboxylase 1 (alpha) | 56922 | Amino Acid (BCAAs) |
| 107 | MCCC2 | methylcrotonoyl-Coenzyme A carboxylase 2 (beta) | 64087 | Amino Acid (BCAAs) |
| 108 | ALDH6A1 | aldehyde dehydrogenase 6 family, member A1 | 4329 | Amino Acid (BCAAs) |
| 109 | AUH | AU RNA binding protein/enoyl-Coenzyme A hydratase | 549 | Amino Acid (BCAAs) |
| 110 | PCBD1 | pterin-4 alpha-carbinolamine dehydratase/dimerization cofactor of hepatocyte nuclear factor 1 alpha | 5092 | Amino Acid (BH4) |
| 111 | PCBD2 | pterin-4 alpha-carbinolamine dehydratase/dimerization cofactor of hepatocyte nuclear factor 1 alpha (TCF1) 2 | 84105 | Amino Acid (BH4) |
| 112 | GCH1 | GTP cyclohydrolase 1 (dopa-responsive dystonia) | 2643 | Amino Acid (BH4) |
| 113 | PTS | 6-pyruvoyltetrahydropterin synthase | 5805 | Amino Acid (BH4) |
| 114 | QDPR | quinoid dihydropteridine reductase | 5860 | Amino Acid (BH4) |
| 115 | SPR | sepiapterin reductase (7,8-dihydrobiopterin: NADP+ oxidoreductase) | 6697 | Amino Acid (BH4) |
| 116 | TMLHE | trimethyllysine hydroxylase, epsilon | 55217 | Amino Acid (Carnitine) |
| 117 | AHCY | S-adenosylhomocysteine hydrolase | 191 | Amino Acid (Cofactor) |
| 118 | AHCYL1 | S-adenosylhomocysteine hydrolase-like 1 | 10768 | Amino Acid (Cofactor) |
| 119 | AHCYL2 | S-adenosylhomocysteine hydrolase-like 2 | 23382 | Amino Acid (Cofactor) |
| 120 | BBOX1 | butyrobetaine (gamma), 2-oxoglutarate dioxygenase (gamma-butyrobetaine hydroxylase) 1 | 8424 | Amino Acid (Cofactor) |
| 121 | MOCOS | molybdenum cofactor sulfurase | 55034 | Amino Acid (Cofactor) |
| 122 | MOCS1 | molybdenum cofactor synthesis 1 | 4337 | Amino Acid (Cofactor) |
| 123 | MOCS2 | molybdenum cofactor synthesis 2 | 4338 | Amino Acid (Cofactor) |
| 124 | MOCS3 | molybdenum cofactor synthesis 3 | 27304 | Amino Acid (Cofactor) |
| 125 | CKB | creatine kinase, brain | 1152 | Amino Acid (Creatine) |
| 126 | CKBE | creatine kinase, ectopic expression | 1156 | Amino Acid (Creatine) |
| 127 | CKM | creatine kinase, muscle | 1158 | Amino Acid (Creatine) |
| 128 | CKMT1A | creatine kinase, mitochondrial 1A | 1159 | Amino Acid (Creatine) |
| 129 | CKMT1B | creatine kinase, mitochondrial 1B | 548596 | Amino Acid (Creatine) |
| 130 | CKMT2 | creatine kinase, mitochondrial 2 (sarcomeric) | 1160 | Amino Acid (Creatine) |
| 131 | GAMT | guanidinoacetate N-methyltransferase | 2593 | Amino Acid (Creatine) |
| 132 | GATM | glycine amidinotransferase (L-arginine:glycine amidinotransferase) | 2628 | Amino Acid (Creatine) |
| 133 | CRYM | crystallin, mu | 1428 | Amino Acid (Cystathionine) |
| 134 | CCBL1 | cysteine conjugate-beta lyase; cytoplasmic (glutamine transaminase K, kyneurenine aminotransferase) | 883 | Amino Acid (Cysteine) |
| 135 | CCBL2 | cysteine conjugate-beta lyase 2 | 56267 | Amino Acid (Cysteine) |
| 136 | ADO | 2-aminoethanethiol (cysteamine) dioxygenase | 84890 | Amino Acid (Cysteine) |
| 137 | CBS | cystathionine-beta-synthase | 875 | Amino Acid (Cysteine) |
| 138 | CDO1 | cysteine dioxygenase, type I | 1036 | Amino Acid (Cysteine) |
| 139 | CSAD | cysteine sulfinic acid decarboxylase | 51380 | Amino Acid (Cysteine) |
| 140 | CTH | cystathionase (cystathionine gamma-lyase) | 1491 | Amino Acid (Cysteine) |
| 141 | NFS1 | NFS1 nitrogen fixation 1 homolog (S. cerevisiae) | 9054 | Amino Acid (Cysteine) |
| 142 | PCYOX1 | prenylcysteine oxidase 1 | 51449 | Amino Acid (Cysteine) |
| 143 | PCYOX1L | prenylcysteine oxidase 1 like | 78991 | Amino Acid (Cysteine) |
| 144 | SUOX | sulfite oxidase | 6821 | Amino Acid (Cysteine) |
| 145 | CMBL | carboxymethylenebutenolidase homolog (Pseudomonas) | 134147 | Amino Acid (Cysteine) |
| 146 | ENOPH1 | enolase-phosphatase 1 | 58478 | Amino Acid (Cysteine, Methione) |
| 147 | AOC2 | amine oxidase, copper containing 2 (retina-specific) | 314 | Amino acid (Dopamine, histamine) |
| 148 | AOC3 | amine oxidase, copper containing 3 (vascular adhesion protein 1) | 8639 | Amino acid (Dopamine, histamine) |
| 149 | ALDH9A1 | aldehyde dehydrogenase 9 family, member A1 | 223 | Amino acid (GABA) |
| 150 | ALDH18A1 | aldehyde dehydrogenase 18 family, member A1 | 5832 | Amino Acid (Glutamate) |
| 151 | ALDH5A1 | aldehyde dehydrogenase 5 family, member A1 (succinate-semialdehyde dehydrogenase) | 7915 | Amino Acid (Glutamate) |
| 152 | GAD1 | glutamate decarboxylase 1 (brain, 67kDa) | 2571 | Amino Acid (Glutamate) |
| 153 | GAD2 | Glutamate decarboxylase 2 (pancreatic islets and brain, 65kDa) | 2572 | Amino Acid (Glutamate) |
| 154 | GADL1 | glutamate decarboxylase-like 1 | 339896 | Amino Acid (Glutamate) |
| 155 | GLS | glutaminase | 2744 | Amino Acid (Glutamate) |
| 156 | GLS2 | glutaminase 2 (liver, mitochondrial) | 27165 | Amino Acid (Glutamate) |
| 157 | GLUD1 | glutamate dehydrogenase 1 | 2746 | Amino Acid (Glutamate) |
| 158 | GLUD2 | glutamate dehydrogenase 2 | 2747 | Amino Acid (Glutamate) |
| 159 | GLUL | glutamate-ammonia ligase (glutamine synthetase) | 2752 | Amino Acid (Glutamate) |
| 160 | OAT | ornithine aminotransferase (gyrate atrophy) | 4942 | Amino Acid (Glutamate) |
| 161 | ADHFE1 | alcohol dehydrogenase, iron containing, 1 | 137872 | Amino Acid (Glutamate) |
| 162 | GPT | glutamic-pyruvate transaminase (alanine aminotransferase) | 2875 | Amino Acid (Glutamate, Alaline) |
| 163 | GPT2 | glutamic pyruvate transaminase (alanine aminotransferase) 2 | 84706 | Amino Acid (Glutamate, Alaline) |
| 164 | LGSN | lens protein with glutamine synthetase domain | 51557 | Amino Acid (Glutamine) |
| 165 | GCLC | glutamate-cysteine ligase, catalytic subunit | 2729 | Amino Acid (Glutathione) |
| 166 | GCLM | glutamate-cysteine ligase, modifier subunit | 2730 | Amino Acid (Glutathione) |
| 167 | GPX1 | glutathione peroxidase 1 | 2876 | Amino Acid (Glutathione) |
| 168 | GPX2 | glutathione peroxidase 2 (gastrointestinal) | 2877 | Amino Acid (Glutathione) |
| 169 | GPX3 | glutathione peroxidase 3 (plasma) | 2878 | Amino Acid (Glutathione) |
| 170 | GPX4 | glutathione peroxidase 4 (phospholipid hydroperoxidase) | 2879 | Amino Acid (Glutathione) |
| 171 | GPX5 | glutathione peroxidase 5 (epididymal androgen-related protein) | 2880 | Amino Acid (Glutathione) |
| 172 | GPX6 | glutathione peroxidase 6 (olfactory) | 257202 | Amino Acid (Glutathione) |
| 173 | GPX7 | glutathione peroxidase 7 | 2882 | Amino Acid (Glutathione) |
| 174 | GPX8 | glutathione peroxidase 8 (putative) | 493869 | Amino Acid (Glutathione) |
| 175 | GSR | glutathione reductase | 2936 | Amino Acid (Glutathione) |
| 176 | GSS | glutathione synthetase | 2937 | Amino Acid (Glutathione) |
| 177 | GSTA1 | glutathione S-transferase A1 | 2938 | Amino Acid (Glutathione) |
| 178 | GSTA2 | glutathione S-transferase A2 | 2939 | Amino Acid (Glutathione) |
| 179 | GSTA3 | glutathione S-transferase A3 | 2940 | Amino Acid (Glutathione) |
| 180 | GSTA4 | glutathione S-transferase A4 | 2941 | Amino Acid (Glutathione) |
| 181 | GSTA5 | glutathione S-transferase A5 | 221357 | Amino Acid (Glutathione) |
| 182 | GSTCD | glutathione S-transferase, C-terminal domain containing | 79807 | Amino Acid (Glutathione) |
| 183 | GSTK1 | glutathione S-transferase kappa 1 | 373156 | Amino Acid (Glutathione) |
| 184 | GSTM1 | glutathione S-transferase M1 | 2944 | Amino Acid (Glutathione) |
| 185 | GSTM2 | glutathione S-transferase M2 (muscle) | 2946 | Amino Acid (Glutathione) |
| 186 | GSTM3 | glutathione S-transferase M3 (brain) | 2947 | Amino Acid (Glutathione) |
| 187 | GSTM4 | glutathione S-transferase M4 | 2948 | Amino Acid (Glutathione) |
| 188 | GSTM5 | glutathione S-transferase M5 | 2949 | Amino Acid (Glutathione) |
| 189 | GSTO1 | glutathione S-transferase omega 1 | 9446 | Amino Acid (Glutathione) |
| 190 | GSTO2 | glutathione S-transferase omega 2 | 119391 | Amino Acid (Glutathione) |
| 191 | GSTP1 | glutathione S-transferase pi | 2950 | Amino Acid (Glutathione) |
| 192 | GSTT1 | glutathione S-transferase theta 1 | 2952 | Amino Acid (Glutathione) |
| 193 | GSTT2 | glutathione S-transferase theta 2 | 2953 | Amino Acid (Glutathione) |
| 194 | HAGH | hydroxyacylglutathione hydrolase | 3029 | Amino Acid (Glutathione) |
| 195 | HAGHL | hydroxyacylglutathione hydrolase-like | 84264 | Amino Acid (Glutathione) |
| 196 | MGST1 | microsomal glutathione S-transferase 1 | 4257 | Amino Acid (Glutathione) |
| 197 | MGST2 | microsomal glutathione S-transferase 2 | 4258 | Amino Acid (Glutathione) |
| 198 | MGST3 | microsomal glutathione S-transferase 3 | 4259 | Amino Acid (Glutathione) |
| 199 | GGT1 | gamma-glutamyltransferase 1 | 2678 | Amino Acid (Glutathione, Taurine, Hypotaurine) |
| 200 | GGT2 | gamma-glutamyltransferase 2 | 728441 | Amino Acid (Glutathione, Taurine, Hypotaurine) |
| 201 | GGT5 | gamma-glutamyltransferase 5 | 2687 | Amino Acid (Glutathione, Taurine, Hypotaurine) |
| 202 | GGT6 | gamma-glutamyltransferase 6 homolog (rat) | 124975 | Amino Acid (Glutathione, Taurine, Hypotaurine) |
| 203 | GGT7 | gamma-glutamyltransferase 7 | 2686 | Amino Acid (Glutathione, Taurine, Hypotaurine) |
| 204 | GGTLC1 | gamma-glutamyltransferase-like activity 4 | 92086 | Amino Acid (Glutathione, Taurine, Hypotaurine) |
| 205 | GGTLC2 | gamma-glutamyltransferase light chain 2 | 91227 | Amino Acid (Glutathione, Taurine, Hypotaurine) |
| 206 | GCAT | glycine C-acetyltransferase (2-amino-3-ketobutyrate coenzyme A ligase) | 23464 | Amino Acid (Glycine) |
| 207 | AMT | aminomethyltransferase | 275 | Amino Acid (Glycine) |
| 208 | BHMT | Betaine-homocysteine methyltransferase | 635 | Amino Acid (Glycine) |
| 209 | BHMT2 | betaine-homocysteine methyltransferase 2 | 23743 | Amino Acid (Glycine) |
| 210 | DMGDH | dimethylglycine dehydrogenase | 29958 | Amino Acid (Glycine) |
| 211 | GCSH | glycine cleavage system protein H (aminomethyl carrier) | 2653 | Amino Acid (Glycine) |
| 212 | GLDC | glycine dehydrogenase (decarboxylating) | 2731 | Amino Acid (Glycine) |
| 213 | SARDH | sarcosine dehydrogenase | 1757 | Amino Acid (Glycine) |
| 214 | CHDH | choline dehydrogenase | 55349 | Amino Acid (Glycine) |
| 215 | GLYAT | glycine-N-acyltransferase | 10249 | Amino Acid (Glycine) |
| 216 | GLYATL1 | glycine-N-acyltransferase-like 1 | 92292 | Amino Acid (Glycine) |
| 217 | GLYATL2 | glycine-N-acyltransferase-like 2 | 219970 | Amino Acid (Glycine) |
| 218 | GNMT | glycine N-methyltransferase | 27232 | Amino Acid (Glycine) |
| 219 | PAM | peptidylglycine alpha-amidating monooxygenase | 5066 | Amino Acid (Glycine) |
| 220 | AMDHD1 | amidohydrolase domain containing 1 | 144193 | Amino Acid (histidine) |
| 221 | AMDHD2 | amidohydrolase domain containing 2 | 51005 | Amino Acid (histidine) |
| 222 | FTCD | formiminotransferase cyclodeaminase | 10841 | Amino Acid (histidine) |
| 223 | HAL | histidine ammonia-lyase | 3034 | Amino Acid (histidine) |
| 224 | HDC | Histidine decarboxylase | 3067 | Amino Acid (histidine) |
| 225 | HNMT | histamine N-methyltransferase | 3176 | Amino Acid (histidine) |
| 226 | UROC1 | Urocanase domain containing 1 | 131669 | Amino Acid (histidine) |
| 227 | DPEP1 | dipeptidase 1 (renal) | 1800 | Amino Acid (Hydrolysis) |
| 228 | ACY1 | Aminoacylase 1 | 95 | Amino Acid (Hydrolysis) |
| 229 | ACY1L | Aminoacylase 1-like | 96 | Amino Acid (Hydrolysis) |
| 230 | DPEP2 | dipeptidase 2 | 64174 | Amino Acid (Hydrolysis) |
| 231 | DPEP3 | dipeptidase 3 | 64180 | Amino Acid (Hydrolysis) |
| 232 | APEH | N-acylaminoacyl-peptide hydrolase | 327 | Amino Acid (Hydrolysis) |
| 233 | ESD | esterase D/formylglutathione hydrolase | 2098 | Amino Acid (Hydrolysis) |
| 234 | CNDP1 | carnosine dipeptidase 1 (metallopeptidase M20 family) | 84735 | Amino Acid (Hydroylysis) |
| 235 | CNDP2 | CNDP dipeptidase 2 (metallopeptidase M20 family) | 55748 | Amino Acid (Hydroylysis) |
| 236 | AADAT | aminoadipate aminotransferase | 51166 | Amino Acid (Kyneurenine, glutamate) |
| 237 | AASDH | aminoadipate-semialdehyde dehydrogenase | 132949 | Amino Acid (Lysine) |
| 238 | AASDHPPT | aminoadipate-semialdehyde dehydrogenase-phosphopantetheinyl transferase | 60496 | Amino Acid (Lysine) |
| 239 | AASS | aminoadipate-semialdehyde synthase | 10157 | Amino Acid (Lysine) |
| 240 | PIPOX | pipecolic acid oxidase | 51268 | Amino Acid (Lysine) |
| 241 | GCDH | glutaryl-Coenzyme A dehydrogenase | 2639 | Amino Acid (Lysine, Tryptophan) |
| 242 | GLTP | glycolipid transfer protein | 51228 | Amino Acid (Membrane lipid) |
| 243 | ADI1 | acireductone dioxygenase 1 | 55256 | Amino Acid (Methionine) |
| 244 | MAT1A | methionine adenosyltransferase I, alpha | 4143 | Amino Acid (Methionine) |
| 245 | MAT2A | methionine adenosyltransferase II, alpha | 4144 | Amino Acid (Methionine) |
| 246 | MAT2B | methionine adenosyltransferase II, beta | 27430 | Amino Acid (Methionine) |
| 247 | MSRA | methionine sulfoxide reductase A | 4482 | Amino Acid (Methionine) |
| 248 | MSRB2 | methionine sulfoxide reductase B2 | 22921 | Amino Acid (Methionine) |
| 249 | MSRB3 | methionine sulfoxide reductase B3 | 253827 | Amino Acid (Methionine) |
| 250 | MTR | 5-methyltetrahydrofolate-homocysteine methyltransferase | 4548 | Amino Acid (Methionine) |
| 251 | SEPX1 | selenoprotein X, 1 | 51734 | Amino Acid (Methionine) |
| 252 | AS3MT | arsenic (+3 oxidation state) methyltransferase | 57412 | Amino Acid (Methionion) |
| 253 | AMD1 | adenosylmethionine decarboxylase 1 | 262 | Amino Acid (Methionion, Polyamine) |
| 254 | MTAP | methylthioadenosine phosphorylase | 4507 | Amino Acid (Methionion, Polyamine) |
| 255 | MAOA | monoamine oxidase A | 4128 | Amino Acid (Neurotransmiter) |
| 256 | MAOB | Monoamine oxidase B | 4129 | Amino Acid (Neurotransmiter) |
| 257 | PNMT | phenylethanolamine N-methyltransferase | 5409 | Amino Acid (Neurotransmitter) |
| 258 | SRR | serine racemase | 63826 | Amino Acid (Neurotransmitter) |
| 259 | TH | tyrosine hydroxylase | 7054 | Amino Acid (Neurotransmitter) |
| 260 | TPH1 | tryptophan hydroxylase 1 (tryptophan 5-monooxygenase) | 7166 | Amino Acid (Neurotransmitter) |
| 261 | TPH2 | tryptophan hydroxylase 2 | 121278 | Amino Acid (Neurotransmitter) |
| 262 | ALDH1L1 | aldehyde dehydrogenase 1 family, member L1 | 10840 | Amino Acid (One carbon) |
| 263 | ALDH1L2 | aldehyde dehydrogenase 1 family, member L2 | 160428 | Amino Acid (One carbon) |
| 264 | ODC1 | ornithine decarboxylase 1 | 4953 | Amino Acid (Ornithine, Polyamine) |
| 265 | DDT | D-dopachrome tautomerase | 1652 | Amino Acid (Phenypyruvate, Pigment) |
| 266 | ALDH4A1 | aldehyde dehydrogenase 4 family, member A1 | 8659 | Amino Acid (Proline) |
| 267 | PRODH | proline dehydrogenase (oxidase) 1 | 5625 | Amino Acid (Proline) |
| 268 | PRODH2 | proline dehydrogenase (oxidase) 2 | 58510 | Amino Acid (Proline) |
| 269 | PYCR1 | pyrroline-5-carboxylate reductase 1 | 5831 | Amino Acid (Proline) |
| 270 | PYCR2 | pyrroline-5-carboxylate reductase family, member 2 | 29920 | Amino Acid (Proline) |
| 271 | PYCRL | pyrroline-5-carboxylate reductase-like | 65263 | Amino Acid (Proline) |
| 272 | P4HA1 | prolyl 4-hydroxylase subunit alpha 1 | 5033 | Amino Acid (Proline) |
| 273 | P4HA2 | prolyl 4-hydroxylase subunit alpha 2 | 8974 | Amino Acid (Proline) |
| 274 | P4HA3 | prolyl 4-hydroxylase subunit alpha 3 | 283208 | Amino Acid (Proline) |
| 275 | P4HB | prolyl 4-hydroxylase subunit beta | 5034 | Amino Acid (Proline) |
| 276 | PEPD | peptidase D | 5184 | Amino Acid (Proline) |
| 277 | EIF5A | eukaryotic translation initiation factor 5A | 1984 | Amino Acid (Proline) |
| 278 | PLOD1 | procollagen-lysine,2-oxoglutarate 5-dioxygenase 1 | 5351 | Amino Acid (Proline) |
| 279 | PLOD2 | procollagen-lysine,2-oxoglutarate 5-dioxygenase 2 | 5352 | Amino Acid (Proline) |
| 280 | LEPREL4 | prolyl 3-hydroxylase family member 4 | 10609 | Amino Acid (Proline) |
| 281 | P3H4 | prolyl 3-hydroxylase family member 4 | 10609 | Amino Acid (Proline) |
| 282 | ACYP1 | acylphosphatase 1, erythrocyte (common) type | 97 | Amino Acid (Pyruvate) |
| 283 | ACYP2 | acylphosphatase 2, muscle type | 98 | Amino Acid (Pyruvate) |
| 284 | SDS | serine dehydratase | 10993 | Amino Acid (Serine) |
| 285 | SDSL | serine dehydratase-like | 113675 | Amino Acid (Serine) |
| 286 | BPHL | biphenyl hydrolase-like (serine hydrolase; breast epithelial mucin-associated antigen) | 670 | Amino Acid (Serine) |
| 287 | PHGDH | phosphoglycerate dehydrogenase | 26227 | Amino Acid (Serine) |
| 288 | PSAT | phosphoserine transaminase | 29968 | Amino Acid (Serine) |
| 289 | PSPH | phosphoserine phosphatase | 5723 | Amino Acid (Serine) |
| 290 | SCLY | selenocysteine lyase | 51540 | Amino Acid (Slenenocysteine) |
| 291 | SEPHS1 | selenophosphate synthetase 1 | 22929 | Amino Acid (Slenocysteine) |
| 292 | SEPHS2 | selenophosphate synthetase 2 | 22928 | Amino Acid (Slenocysteine) |
| 293 | THNSL1 | threonine synthase-like 1 (S. cerevisiae) | 79896 | Amino Acid (Threonine) |
| 294 | THNSL2 | threonine synthase-like 2 (S. cerevisiae) | 55258 | Amino Acid (Threonine) |
| 295 | DCT | dopachrome tautomerase (dopachrome delta-isomerase, tyrosine-related protein 2) | 1638 | Amino Acid (Tryosine, Pigment) |
| 296 | TYR | tyrosinase (oculocutaneous albinism IA) | 7299 | Amino Acid (Tryosine, Pigment) |
| 297 | TYRP1 | tyrosinase-related protein 1 | 7306 | Amino Acid (Tryosine, Pigment) |
| 298 | ACMSD | aminocarboxymuconate semialdehyde decarboxylase | 130013 | Amino Acid (Tryptophan) |
| 299 | AFMID | arylformamidase | 125061 | Amino Acid (Tryptophan) |
| 300 | INMT | indolethylamine N-methyltransferase | 11185 | Amino Acid (Tryptophan) |
| 301 | KMO | kynurenine 3-monooxygenase (kynurenine 3-hydroxylase) | 8564 | Amino Acid (Tryptophan) |
| 302 | KYNU | kynureninase (L-kynurenine hydrolase) | 8942 | Amino Acid (Tryptophan) |
| 303 | TDO2 | tryptophan 2,3-dioxygenase | 6999 | Amino Acid (Tryptophan) |
| 304 | HAAO | 3-hydroxyanthranilate 3,4-dioxygenase | 23498 | Amino Acid (Tryptophan, NAD)) |
| 305 | IDO1 | indoleamine 2,3-dioxygenase 1 | 3620 | Amino Acid (Tryptophan, NAD)) |
| 306 | IDO2 | indoleamine-pyrrole 2,3 dioxygenase 2 | 169355 | Amino Acid (Tryptophan, NAD)) |
| 307 | GSTZ1 | glutathione transferase zeta 1 (maleylacetoacetate isomerase) | 2954 | Amino Acid (Tryrosine) |
| 308 | FAH | fumarylacetoacetate hydrolase (fumarylacetoacetase) | 2184 | Amino Acid (Tyrosine) |
| 309 | FAHD1 | fumarylacetoacetate hydrolase domain containing 1 | 81889 | Amino Acid (Tyrosine) |
| 310 | FAHD2A | fumarylacetoacetate hydrolase domain containing 2A | 51011 | Amino Acid (Tyrosine) |
| 311 | FAHD2B | fumarylacetoacetate hydrolase domain containing 2B | 151313 | Amino Acid (Tyrosine) |
| 312 | HGD | homogentisate 1,2-dioxygenase (homogentisate oxidase) | 3081 | Amino Acid (Tyrosine) |
| 313 | HPD | 4-hydroxyphenylpyruvate dioxygenase | 3242 | Amino Acid (Tyrosine) |
| 314 | HPDL | 4-hydroxyphenylpyruvate dioxygenase-like | 84842 | Amino Acid (Tyrosine) |
| 315 | PAH | phenylalanine hydroxylase | 5053 | Amino Acid (Tyrosine) |
| 316 | TAT | tyrosine aminotransferase | 6898 | Amino Acid (Tyrosine) |
| 317 | AGMAT | agmatine ureohydrolase (agmatinase) | 79814 | Amino Acid (Urea) |
| 318 | CPS1 | carbamoyl-phosphate synthetase 1, mitochondrial | 1373 | Amino Acid (Urea) |
| 319 | ACOT1 | acyl-CoA thioesterase 1 | 641371 | CoA |
| 320 | ACOT11 | acyl-CoA thioesterase 11 | 26027 | CoA |
| 321 | ACOT12 | acyl-CoA thioesterase 12 | 134526 | CoA |
| 322 | ACOT2 | acyl-CoA thioesterase 2 | 10965 | CoA |
| 323 | ACOT4 | acyl-CoA thioesterase 4 | 122970 | CoA |
| 324 | ACOT6 | acyl-CoA thioesterase 6 | 641372 | CoA |
| 325 | ACOT7 | acyl-CoA thioesterase 7 | 11332 | CoA |
| 326 | ACOT8 | Acyl-CoA thioesterase 8 | 10005 | CoA |
| 327 | ACOT9 | acyl-CoA thioesterase 9 | 23597 | CoA |
| 328 | COASY | Coenzyme A synthase | 80347 | CoA |
| 329 | PANK1 | pantothenate kinase 1 | 53354 | CoA |
| 330 | PANK2 | pantothenate kinase 2 (Hallervorden-Spatz syndrome) | 80025 | CoA |
| 331 | PANK3 | pantothenate kinase 3 | 79646 | CoA |
| 332 | PANK4 | pantothenate kinase 4 | 55229 | CoA |
| 333 | PPCDC | phosphopantothenoylcysteine decarboxylase | 60490 | CoA |
| 334 | PPCS | phosphopantothenoylcysteine synthetase | 79717 | CoA |
| 335 | ARSF | arylsulfatase F | 416 | Cofactor (Calcium) |
| 336 | FLAD1 | FAD1 flavin adenine dinucleotide synthetase homolog (S. cerevisiae) | 80308 | Cofactor (FAD) |
| 337 | HLCS | holocarboxylase synthetase (biotin-(proprionyl-Coenzyme A-carboxylase (ATP-hydrolysing)) ligase) | 3141 | Cofactor(molybdenum) |
| 338 | ALDH1B1 | aldehyde dehydrogenase 1 family, member B1 | 219 | Detox |
| 339 | ALDH7A1 | aldehyde dehydrogenase 7 family, member A1 | 501 | Detox |
| 340 | EPX | eosinophil peroxidase | 8288 | Detox |
| 341 | CAT | Catalase | 847 | Detox |
| 342 | DHDH | dihydrodiol dehydrogenase (dimeric) | 27294 | Detox |
| 343 | TPMT | thiopurine S-methyltransferase | 7172 | Detox |
| 344 | UGT1A1 | UDP glucuronosyltransferase 1 family, polypeptide A1 | 54658 | Detox |
| 345 | UGT1A10 | UDP glucuronosyltransferase 1 family, polypeptide A10 | 54575 | Detox |
| 346 | UGT1A3 | UDP glucuronosyltransferase 1 family, polypeptide A3 | 54659 | Detox |
| 347 | UGT1A4 | UDP glucuronosyltransferase 1 family, polypeptide A4 | 54657 | Detox |
| 348 | UGT1A5 | UDP glucuronosyltransferase 1 family, polypeptide A5 | 54579 | Detox |
| 349 | UGT1A6 | UDP glucuronosyltransferase 1 family, polypeptide A6 | 54578 | Detox |
| 350 | UGT1A7 | UDP glucuronosyltransferase 1 family, polypeptide A7 | 54577 | Detox |
| 351 | UGT1A8 | UDP glucuronosyltransferase 1 family, polypeptide A8 | 54576 | Detox |
| 352 | UGT1A9 | UDP glucuronosyltransferase 1 family, polypeptide A9 | 54600 | Detox |
| 353 | UGT2A1 | UDP glucuronosyltransferase 2 family, polypeptide A1 | 10941 | Detox |
| 354 | UGT2A2 | UDP glucuronosyltransferase 2 family, polypeptide A2 | 574537 | Detox |
| 355 | UGT2A3 | UDP glucuronosyltransferase 2 family, polypeptide A3 | 79799 | Detox |
| 356 | UGT2B10 | UDP glucuronosyltransferase 2 family, polypeptide B10 | 7365 | Detox |
| 357 | UGT2B11 | UDP glucuronosyltransferase 2 family, polypeptide B11 | 10720 | Detox |
| 358 | UGT2B15 | UDP glucuronosyltransferase 2 family, polypeptide B15 | 7366 | Detox |
| 359 | UGT2B17 | UDP glucuronosyltransferase 2 family, polypeptide B17 | 7367 | Detox |
| 360 | UGT2B28 | UDP glucuronosyltransferase 2 family, polypeptide B28 | 54490 | Detox |
| 361 | UGT2B4 | UDP glucuronosyltransferase 2 family, polypeptide B4 | 7363 | Detox |
| 362 | UGT2B7 | UDP glucuronosyltransferase 2 family, polypeptide B7 | 7364 | Detox |
| 363 | UGT3A1 | UDP glycosyltransferase 3 family, polypeptide A1 | 133688 | Detox |
| 364 | UGT3A2 | UDP glycosyltransferase 3 family, polypeptide A2 | 167127 | Detox |
| 365 | MPO | myeloperoxidase | 4353 | Detox |
| 366 | AADAC | arylacetamide deacetylase (esterase) | 13 | Detox |
| 367 | AADACL1 | arylacetamide deacetylase-like 1 | 57552 | Detox |
| 368 | AADACL2 | arylacetamide deacetylase-like 2 | 344752 | Detox |
| 369 | AADACL3 | arylacetamide deacetylase-like 3 | 126767 | Detox |
| 370 | AADACL4 | arylacetamide deacetylase-like 4 | 343066 | Detox |
| 371 | AKR7L | aldo-keto reductase family 7-like | 246181 | Detox |
| 372 | BCHE | butyrylcholinesterase | 590 | Detox |
| 373 | CES1 | carboxylesterase 1 (monocyte/macrophage serine esterase 1) | 1066 | Detox |
| 374 | CES2 | carboxylesterase 2 (intestine, liver) | 8824 | Detox |
| 375 | CES3 | carboxylesterase 3 (brain) | 23491 | Detox |
| 376 | CES7 | carboxylesterase 7 | 221223 | Detox |
| 377 | EPHX1 | epoxide hydrolase 1, microsomal (xenobiotic) | 2052 | Detox |
| 378 | EPHX2 | epoxide hydrolase 2, cytoplasmic | 2053 | Detox |
| 379 | CBR1 | carbonyl reductase 1 | 873 | Detox |
| 380 | CBR3 | carbonyl reductase 3 | 874 | Detox |
| 381 | PON1 | paraoxonase 1 | 5444 | Detox |
| 382 | PON2 | paraoxonase 2 | 5445 | Detox |
| 383 | PON3 | paraoxonase 3 | 5446 | Detox |
| 384 | FMO1 | flavin containing monooxygenase 1 | 2326 | Detox |
| 385 | FMO2 | flavin containing monooxygenase 2 | 2327 | Detox |
| 386 | FMO3 | flavin containing monooxygenase 3 | 2328 | Detox |
| 387 | FMO4 | flavin containing monooxygenase 4 | 2329 | Detox |
| 388 | FMO5 | flavin containing monooxygenase 5 | 2330 | Detox |
| 389 | SOD1 | superoxide dismutase 1, soluble (amyotrophic lateral sclerosis 1 (adult)) | 6647 | Detox |
| 390 | SOD2 | Superoxide dismutase 2, mitochondrial | 6648 | Detox |
| 391 | SOD3 | superoxide dismutase 3, extracellular | 6649 | Detox |
| 392 | ALDH3A1 | aldehyde dehydrogenase 3 family, memberA1 | 218 | Detox |
| 393 | ALDH2 | aldehyde dehydrogenase 2 family (mitochondrial) | 217 | Detox (Ethanoll) |
| 394 | MT-ND1 | michondrially encoded NADH dehydrogenase 1 | 4535 | ETC (Complex I) |
| 395 | MT-ND2 | michondrially encoded NADH dehydrogenase 2 | 4536 | ETC (Complex I) |
| 396 | MT-ND3 | michondrially encoded NADH dehydrogenase 3 | 4537 | ETC (Complex I) |
| 397 | MT-ND4 | michondrially encoded NADH dehydrogenase 4 | 4538 | ETC (Complex I) |
| 398 | MT-ND4L | michondrially encoded NADH dehydrogenase 4 like | 4539 | ETC (Complex I) |
| 399 | MT-ND5 | michondrially encoded NADH dehydrogenase 5 | 4540 | ETC (Complex I) |
| 400 | MT-ND6 | michondrially encoded NADH dehydrogenase 6 | 4541 | ETC (Complex I) |
| 401 | NDUFA1 | NADH dehydrogenase (ubiquinone) 1 alpha subcomplex, 1, 7.5kDa | 4694 | ETC (Complex I) |
| 402 | NDUFA10 | NADH dehydrogenase (ubiquinone) 1 alpha subcomplex, 10, 42kDa | 4705 | ETC (Complex I) |
| 403 | NDUFA11 | NADH dehydrogenase (ubiquinone) 1 alpha subcomplex, 11, 14.7kDa | 126328 | ETC (Complex I) |
| 404 | NDUFA12 | NADH dehydrogenase (ubiquinone) 1 alpha subcomplex, 12 | 55967 | ETC (Complex I) |
| 405 | NDUFA13 | NADH dehydrogenase (ubiquinone) 1 alpha subcomplex, 13 | 51079 | ETC (Complex I) |
| 406 | NDUFA2 | NADH dehydrogenase (ubiquinone) 1 alpha subcomplex, 2, 8kDa | 4695 | ETC (Complex I) |
| 407 | NDUFA3 | NADH dehydrogenase (ubiquinone) 1 alpha subcomplex, 3, 9kDa | 4696 | ETC (Complex I) |
| 408 | NDUFA4 | NADH dehydrogenase (ubiquinone) 1 alpha subcomplex, 4, 9kDa | 4697 | ETC (Complex I) |
| 409 | NDUFA4L | NADH dehydrogenase (ubiquinone) 1 alpha subcomplex, 4-like | 404215 | ETC (Complex I) |
| 410 | NDUFA4L2 | NADH dehydrogenase (ubiquinone) 1 alpha subcomplex, 4-like 2 | 56901 | ETC (Complex I) |
| 411 | NDUFA5 | NADH dehydrogenase (ubiquinone) 1 alpha subcomplex, 5, 13kDa | 4698 | ETC (Complex I) |
| 412 | NDUFA6 | NADH dehydrogenase (ubiquinone) 1 alpha subcomplex, 6, 14kDa | 4700 | ETC (Complex I) |
| 413 | NDUFA7 | NADH dehydrogenase (ubiquinone) 1 alpha subcomplex, 7, 14.5kDa | 4701 | ETC (Complex I) |
| 414 | NDUFA8 | NADH dehydrogenase (ubiquinone) 1 alpha subcomplex, 8, 19kDa | 4702 | ETC (Complex I) |
| 415 | NDUFA9 | NADH dehydrogenase (ubiquinone) 1 alpha subcomplex, 9, 39kDa | 4704 | ETC (Complex I) |
| 416 | NDUFAB1 | NADH dehydrogenase (ubiquinone) 1, alpha/beta subcomplex, 1, 8kDa | 4706 | ETC (Complex I) |
| 417 | NDUFAF1 | NADH dehydrogenase (ubiquinone) 1 alpha subcomplex, assembly factor 1 | 51103 | ETC (Complex I) |
| 418 | NDUFAF2 | NADH dehydrogenase (ubiquinone) 1 alpha subcomplex, assembly factor 2 | 91942 | ETC (Complex I) |
| 419 | NDUFB1 | NADH dehydrogenase (ubiquinone) 1 beta subcomplex, 1, 7kDa | 4707 | ETC (Complex I) |
| 420 | NDUFB10 | NADH dehydrogenase (ubiquinone) 1 beta subcomplex, 10, 22kDa | 4716 | ETC (Complex I) |
| 421 | NDUFB11 | NADH dehydrogenase (ubiquinone) 1 beta subcomplex, 11, 17.3kDa | 54539 | ETC (Complex I) |
| 422 | NDUFB2 | NADH dehydrogenase (ubiquinone) 1 beta subcomplex, 2, 8kDa | 4708 | ETC (Complex I) |
| 423 | NDUFB3 | NADH dehydrogenase (ubiquinone) 1 beta subcomplex, 3, 12kDa | 4709 | ETC (Complex I) |
| 424 | NDUFB4 | NADH dehydrogenase (ubiquinone) 1 beta subcomplex, 4, 12kDa | 4710 | ETC (Complex I) |
| 425 | NDUFB5 | NADH dehydrogenase (ubiquinone) 1 beta subcomplex, 5, 16kDa | 4711 | ETC (Complex I) |
| 426 | NDUFB6 | NADH dehydrogenase (ubiquinone) 1 beta subcomplex, 6, 17kDa | 4712 | ETC (Complex I) |
| 427 | NDUFB7 | NADH dehydrogenase (ubiquinone) 1 beta subcomplex, 7, 18kDa | 4713 | ETC (Complex I) |
| 428 | NDUFB8 | NADH dehydrogenase (ubiquinone) 1 beta subcomplex, 8, 19kDa | 4714 | ETC (Complex I) |
| 429 | NDUFB9 | NADH dehydrogenase (ubiquinone) 1 beta subcomplex, 9, 22kDa | 4715 | ETC (Complex I) |
| 430 | NDUFC1 | NADH dehydrogenase (ubiquinone) 1, subcomplex unknown, 1, 6kDa | 4717 | ETC (Complex I) |
| 431 | NDUFC2 | NADH dehydrogenase (ubiquinone) 1, subcomplex unknown, 2, 14.5kDa | 4718 | ETC (Complex I) |
| 432 | NDUFS1 | NADH dehydrogenase (ubiquinone) Fe-S protein 1, 75kDa (NADH-coenzyme Q reductase) | 4719 | ETC (Complex I) |
| 433 | NDUFS2 | NADH dehydrogenase (ubiquinone) Fe-S protein 2, 49kDa (NADH-coenzyme Q reductase) | 4720 | ETC (Complex I) |
| 434 | NDUFS3 | NADH dehydrogenase (ubiquinone) Fe-S protein 3, 30kDa (NADH-coenzyme Q reductase) | 4722 | ETC (Complex I) |
| 435 | NDUFS4 | NADH dehydrogenase (ubiquinone) Fe-S protein 4, 18kDa (NADH-coenzyme Q reductase) | 4724 | ETC (Complex I) |
| 436 | NDUFS5 | NADH dehydrogenase (ubiquinone) Fe-S protein 5, 15kDa (NADH-coenzyme Q reductase) | 4725 | ETC (Complex I) |
| 437 | NDUFS6 | NADH dehydrogenase (ubiquinone) Fe-S protein 6, 13kDa (NADH-coenzyme Q reductase) | 4726 | ETC (Complex I) |
| 438 | NDUFS7 | NADH dehydrogenase (ubiquinone) Fe-S protein 7, 20kDa (NADH-coenzyme Q reductase) | 374291 | ETC (Complex I) |
| 439 | NDUFS8 | NADH dehydrogenase (ubiquinone) Fe-S protein 8, 23kDa (NADH-coenzyme Q reductase) | 4728 | ETC (Complex I) |
| 440 | NDUFV1 | NADH dehydrogenase (ubiquinone) flavoprotein 1, 51kDa | 4723 | ETC (Complex I) |
| 441 | NDUFV2 | NADH dehydrogenase (ubiquinone) flavoprotein 2, 24kDa | 4729 | ETC (Complex I) |
| 442 | NDUFV3 | NADH dehydrogenase (ubiquinone) flavoprotein 3, 10kDa | 4731 | ETC (Complex I) |
| 443 | SDHA | succinate dehydrogenase complex, subunit A, flavoprotein (Fp) | 6389 | ETC (Complex II) |
| 444 | SDHB | succinate dehydrogenase complex, subunit B, iron sulfur (Ip) | 6390 | ETC (Complex II) |
| 445 | SDHC | succinate dehydrogenase complex, subunit C, integral membrane protein, 15kDa | 6391 | ETC (Complex II) |
| 446 | SDHD | succinate dehydrogenase complex, subunit D, integral membrane protein | 6392 | ETC (Complex II) |
| 447 | ACN9 | ACN9 homolog (S. cerevisiae) | 57001 | ETC (Complex II) |
| 448 | CYC1 | cytochrome c-1 | 1537 | ETC (Complex III) |
| 449 | CYCS | cytochrome c, somatic | 54205 | ETC (Complex III) |
| 450 | MT-CYB | Cytochrome b | 4519 | ETC (Complex III) |
| 451 | UCRC | ubiquinol-cytochrome c reductase complex (7.2 kD) | 29796 | ETC (Complex III) |
| 452 | UQCR | ubiquinol-cytochrome c reductase, 6.4kDa subunit | 10975 | ETC (Complex III) |
| 453 | UQCRB | ubiquinol-cytochrome c reductase binding protein | 7381 | ETC (Complex III) |
| 454 | UQCRC1 | ubiquinol-cytochrome c reductase core protein I | 7384 | ETC (Complex III) |
| 455 | UQCRC2 | ubiquinol-cytochrome c reductase core protein II | 7385 | ETC (Complex III) |
| 456 | UQCRFS1 | ubiquinol-cytochrome c reductase, Rieske iron-sulfur polypeptide 1 | 7386 | ETC (Complex III) |
| 457 | UQCRH | ubiquinol-cytochrome c reductase hinge protein | 7388 | ETC (Complex III) |
| 458 | UQCRQ | ubiquinol-cytochrome c reductase, complex III subunit VII, 9.5kDa | 27089 | ETC (Complex III) |
| 459 | COX1 | mitochondrially encoded cytochrome c oxidase I | 4512 | ETC (Complex IV) |
| 460 | COX10 | COX10 homolog, cytochrome c oxidase assembly protein, heme A: farnesyltransferase (yeast) | 1352 | ETC (Complex IV) |
| 461 | COX11 | COX11 homolog, cytochrome c oxidase assembly protein (yeast) | 1353 | ETC (Complex IV) |
| 462 | COX15 | COX15 homolog, cytochrome c oxidase assembly protein (yeast) | 1355 | ETC (Complex IV) |
| 463 | COX16 | COX16 cytochrome c oxidase assembly homolog (S. cerevisiae) | 51241 | ETC (Complex IV) |
| 464 | COX17 | COX17 cytochrome c oxidase assembly homolog (S. cerevisiae) | 10063 | ETC (Complex IV) |
| 465 | COX18 | COX18 cytochrome c oxidase assembly homolog (S. cerevisiae) | 285521 | ETC (Complex IV) |
| 466 | COX19 | COX19 cytochrome c oxidase assembly homolog (S. cerevisiae) | 90639 | ETC (Complex IV) |
| 467 | COX2 | mitochondrially encoded cytochrome c oxidase II | 4513 | ETC (Complex IV) |
| 468 | COX3 | mitochondrially encoded cytochrome c oxidase III | 4514 | ETC (Complex IV) |
| 469 | COX4I1 | cytochrome c oxidase subunit IV isoform 1 | 1327 | ETC (Complex IV) |
| 470 | COX4I2 | cytochrome c oxidase subunit IV isoform 2 (lung) | 84701 | ETC (Complex IV) |
| 471 | COX5A | cytochrome c oxidase subunit Va | 9377 | ETC (Complex IV) |
| 472 | COX5B | Cytochrome c oxidase subunit Vb | 1329 | ETC (Complex IV) |
| 473 | COX6A1 | cytochrome c oxidase subunit VIa polypeptide 1 | 1337 | ETC (Complex IV) |
| 474 | COX6A2 | cytochrome c oxidase subunit VIa polypeptide 2 | 1339 | ETC (Complex IV) |
| 475 | COX6B1 | cytochrome c oxidase subunit Vib polypeptide 1 (ubiquitous) | 1340 | ETC (Complex IV) |
| 476 | COX6B2 | cytochrome c oxidase subunit VIb polypeptide 2 (testis) | 125965 | ETC (Complex IV) |
| 477 | COX6C | cytochrome c oxidase subunit VIc | 1345 | ETC (Complex IV) |
| 478 | COX7A1 | cytochrome c oxidase subunit VIIa polypeptide 1 (muscle) | 1346 | ETC (Complex IV) |
| 479 | COX7A2 | cytochrome c oxidase subunit VIIa polypeptide 2 (liver) | 1347 | ETC (Complex IV) |
| 480 | COX7A2L | cytochrome c oxidase subunit VIIa polypeptide 2 like | 9167 | ETC (Complex IV) |
| 481 | COX7B | cytochrome c oxidase subunit VIIb | 1349 | ETC (Complex IV) |
| 482 | COX7B2 | cytochrome c oxidase subunit VIIb2 | 170712 | ETC (Complex IV) |
| 483 | COX7C | cytochrome c oxidase subunit VIIc | 1350 | ETC (Complex IV) |
| 484 | COX8A | cytochrome c oxidase subunit 8A (ubiquitous) | 1351 | ETC (Complex IV) |
| 485 | COX8C | cytochrome c oxidase subunit 8C | 341947 | ETC (Complex IV) |
| 486 | A4GALT | alpha 1,4-galactosyltransferase (globotriaosylceramide synthase) | 53947 | Glycan |
| 487 | A4GNT | alpha-1,4-N-acetylglucosaminyltransferase | 51146 | Glycan |
| 488 | ABO | ABO blood group (transferase A, alpha 1-3-N-acetylgalactosaminyltransferase; transferase B, alpha 1-3-galactosyltransferase) | 28 | Glycan |
| 489 | ALG1 | asparagine-linked glycosylation 1 homolog (S. cerevisiae, beta-1,4-mannosyltransferase) | 56052 | Glycan |
| 490 | ALG10 | asparagine-linked glycosylation 10 homolog (yeast, alpha-1,2-glucosyltransferase) | 84920 | Glycan |
| 491 | ALG10B | asparagine-linked glycosylation 10 homolog B (yeast, alpha-1,2-glucosyltransferase) | 144245 | Glycan |
| 492 | ALG11 | asparagine-linked glycosylation 11 homolog | 440138 | Glycan |
| 493 | ALG12 | asparagine-linked glycosylation 12 homolog (S. cerevisiae, alpha-1,6-mannosyltransferase) | 79087 | Glycan |
| 494 | ALG13 | asparagine-linked glycosylation 13 homolog (S. cerevisiae) | 55849 | Glycan |
| 495 | ALG14 | asparagine-linked glycosylation 14 homolog (S. cerevisiae) | 199857 | Glycan |
| 496 | ALG2 | asparagine-linked glycosylation 2 homolog (S. cerevisiae, alpha-1,3-mannosyltransferase) | 85365 | Glycan |
| 497 | ALG3 | asparagine-linked glycosylation 3 homolog (S. cerevisiae, alpha-1,3-mannosyltransferase) | 10195 | Glycan |
| 498 | ALG5 | Asparagine-linked glycosylation 5 homolog (S. cerevisiae, dolichyl-phosphate beta-glucosyltransferase) | 29880 | Glycan |
| 499 | ALG6 | asparagine-linked glycosylation 6 homolog (S. cerevisiae, alpha-1,3-glucosyltransferase) | 29929 | Glycan |
| 500 | ALG8 | asparagine-linked glycosylation 8 homolog (S. cerevisiae, alpha-1,3-glucosyltransferase) | 79053 | Glycan |
| 501 | ALG9 | asparagine-linked glycosylation 9 homolog (S. cerevisiae, alpha- 1,2-mannosyltransferase) | 79796 | Glycan |
| 502 | B3GALNT1 | beta-1,3-N-acetylgalactosaminyltransferase 1 (globoside blood group) | 8706 | Glycan |
| 503 | B3GALNT2 | beta-1,3-N-acetylgalactosaminyltransferase 2 | 148789 | Glycan |
| 504 | B3GALT1 | UDP-Gal: betaGlcNAc beta 1,3-galactosyltransferase, polypeptide 1 | 8708 | Glycan |
| 505 | B3GALT2 | UDP-Gal: betaGlcNAc beta 1,3-galactosyltransferase, polypeptide 2 | 8707 | Glycan |
| 506 | B3GALT4 | UDP-Gal: betaGlcNAc beta 1,3-galactosyltransferase, polypeptide 4 | 8705 | Glycan |
| 507 | B3GALT5 | UDP-Gal: betaGlcNAc beta 1,3-galactosyltransferase, polypeptide 5 | 10317 | Glycan |
| 508 | B3GALT6 | UDP-Gal: betaGal beta 1,3-galactosyltransferase polypeptide 6 | 126792 | Glycan |
| 509 | B3GALTL | beta 1,3-galactosyltransferase-like | 145173 | Glycan |
| 510 | B3GAT1 | beta-1,3-glucuronyltransferase 1 (glucuronosyltransferase P) | 27087 | Glycan |
| 511 | B3GAT2 | beta-1,3-glucuronyltransferase 2 (glucuronosyltransferase P) | 135152 | Glycan |
| 512 | B3GAT3 | beta-1,3-glucuronyltransferase 3 (glucuronosyltransferase I) | 26229 | Glycan |
| 513 | B3GNT1 | UDP-GlcNAc: betaGal beta-1,3-N-acetylglucosaminyltransferase 1 | 11041 | Glycan |
| 514 | B3GNT2 | UDP-GlcNAc: betaGal beta-1,3-N-acetylglucosaminyltransferase 2 | 10678 | Glycan |
| 515 | B3GNT3 | UDP-GlcNAc: betaGal beta-1,3-N-acetylglucosaminyltransferase 3 | 10331 | Glycan |
| 516 | B3GNT4 | UDP-GlcNAc: betaGal beta-1,3-N-acetylglucosaminyltransferase 4 | 79369 | Glycan |
| 517 | B3GNT5 | UDP-GlcNAc: betaGal beta-1,3-N-acetylglucosaminyltransferase 5 | 84002 | Glycan |
| 518 | B3GNT6 | UDP-GlcNAc: betaGal beta-1,3-N-acetylglucosaminyltransferase 6 (core 3 synthase) | 192134 | Glycan |
| 519 | B3GNT7 | UDP-GlcNAc: betaGal beta-1,3-N-acetylglucosaminyltransferase 7 | 93010 | Glycan |
| 520 | B3GNT8 | UDP-GlcNAc: betaGal beta-1,3-N-acetylglucosaminyltransferase 8 | 374907 | Glycan |
| 521 | B3GNT9 | UDP-GlcNAc: betaGal beta-1,3-N-acetylglucosaminyltransferase 9 | 84752 | Glycan |
| 522 | B3GNTL1 | UDP-GlcNAc: betaGal beta-1,3-N-acetylglucosaminyltransferase-like 1 | 146712 | Glycan |
| 523 | B4GALNT1 | beta-1,4-N-acetyl-galactosaminyl transferase 1 | 2583 | Glycan |
| 524 | B4GALNT2 | beta-1,4-N-acetyl-galactosaminyl transferase 2 | 124872 | Glycan |
| 525 | B4GALNT3 | beta-1,4-N-acetyl-galactosaminyl transferase 3 | 283358 | Glycan |
| 526 | B4GALNT4 | beta-1,4-N-acetyl-galactosaminyl transferase 4 | 338707 | Glycan |
| 527 | B4GALT1 | UDP-Gal: betaGlcNAc beta 1,4- galactosyltransferase, polypeptide 1 | 2683 | Glycan |
| 528 | B4GALT2 | UDP-Gal: betaGlcNAc beta 1,4- galactosyltransferase, polypeptide 2 | 8704 | Glycan |
| 529 | B4GALT3 | UDP-Gal: betaGlcNAc beta 1,4- galactosyltransferase, polypeptide 3 | 8703 | Glycan |
| 530 | B4GALT4 | UDP-Gal: betaGlcNAc beta 1,4- galactosyltransferase, polypeptide 4 | 8702 | Glycan |
| 531 | B4GALT5 | UDP-Gal: betaGlcNAc beta 1,4- galactosyltransferase, polypeptide 5 | 9334 | Glycan |
| 532 | B4GALT6 | UDP-Gal: betaGlcNAc beta 1,4- galactosyltransferase, polypeptide 6 | 9331 | Glycan |
| 533 | B4GALT7 | xylosylprotein beta 1,4-galactosyltransferase, polypeptide 7 (galactosyltransferase I) | 11285 | Glycan |
| 534 | C1GALT1 | core 1 synthase, glycoprotein-N-acetylgalactosamine 3-beta-galactosyltransferase, 1 | 56913 | Glycan |
| 535 | CTBS | chitobiase, di-N-acetyl- | 1486 | Glycan |
| 536 | DAD1 | defender against cell death | 1603 | Glycan |
| 537 | DDOST | dolichyl-diphosphooligosaccharide-protein glycosyltransferase | 1650 | Glycan |
| 538 | DHDDS | dehydrodolichyl diphosphate synthase | 79947 | Glycan |
| 539 | DOLK | dolichol kinase | 22845 | Glycan |
| 540 | DOLPP1 | dolichyl pyrophosphate phosphatase 1 | 57171 | Glycan |
| 541 | DPAGT1 | dolichyl-phosphate (UDP-N-acetylglucosamine) N-acetylglucosaminephosphotransferase 1 (GlcNAc-1-P transferase) | 1798 | Glycan |
| 542 | FN3K | fructosamine 3 kinase | 64122 | Glycan |
| 543 | FN3KRP | fructosamine-3-kinase-related protein | 79672 | Glycan |
| 544 | FUT1 | fucosyltransferase 1 (galactoside 2-alpha-L-fucosyltransferase, H blood group) | 2523 | Glycan |
| 545 | FUT10 | fucosyltransferase 10 (alpha (1,3) fucosyltransferase) | 84750 | Glycan |
| 546 | FUT11 | fucosyltransferase 11 (alpha (1,3) fucosyltransferase) | 170384 | Glycan |
| 547 | FUT2 | fucosyltransferase 2 (secretor status included) | 2524 | Glycan |
| 548 | FUT3 | fucosyltransferase 3 (galactoside 3(4)-L-fucosyltransferase, Lewis blood group) | 2525 | Glycan |
| 549 | FUT4 | fucosyltransferase 4 (alpha (1,3) fucosyltransferase, myeloid-specific) | 2526 | Glycan |
| 550 | FUT5 | fucosyltransferase 5 (alpha (1,3) fucosyltransferase) | 2527 | Glycan |
| 551 | FUT6 | Fucosyltransferase 6 (alpha (1,3) fucosyltransferase) | 2528 | Glycan |
| 552 | FUT7 | fucosyltransferase 7 (alpha (1,3) fucosyltransferase) | 2529 | Glycan |
| 553 | FUT8 | fucosyltransferase 8 (alpha (1,6) fucosyltransferase) | 2530 | Glycan |
| 554 | FUT9 | fucosyltransferase 9 (alpha (1,3) fucosyltransferase) | 10690 | Glycan |
| 555 | GALE | UDP-galactose-4-epimerase | 2582 | Glycan |
| 556 | GALNT1 | UDP-N-acetyl-alpha-D-galactosamine: polypeptide N-acetylgalactosaminyltransferase 1 (GalNAc-T1) | 2589 | Glycan |
| 557 | GALNT10 | UDP-N-acetyl-alpha-D-galactosamine: polypeptide N-acetylgalactosaminyltransferase 10 (GalNAc-T10) | 55568 | Glycan |
| 558 | GALNT11 | UDP-N-acetyl-alpha-D-galactosamine: polypeptide N-acetylgalactosaminyltransferase 11 (GalNAc-T11) | 63917 | Glycan |
| 559 | GALNT12 | UDP-N-acetyl-alpha-D-galactosamine: polypeptide N-acetylgalactosaminyltransferase 12 (GalNAc-T12) | 79695 | Glycan |
| 560 | GALNT13 | UDP-N-acetyl-alpha-D-galactosamine: polypeptide N-acetylgalactosaminyltransferase 13 (GalNAc-T13) | 114805 | Glycan |
| 561 | GALNT14 | UDP-N-acetyl-alpha-D-galactosamine: polypeptide N-acetylgalactosaminyltransferase 14 (GalNAc-T14) | 79623 | Glycan |
| 562 | GALNT2 | UDP-N-acetyl-alpha-D-galactosamine: polypeptide N-acetylgalactosaminyltransferase 2 (GalNAc-T2) | 2590 | Glycan |
| 563 | GALNT3 | UDP-N-acetyl-alpha-D-galactosamine: polypeptide N-acetylgalactosaminyltransferase 3 (GalNAc-T3) | 2591 | Glycan |
| 564 | GALNT4 | UDP-N-acetyl-alpha-D-galactosamine: polypeptide N-acetylgalactosaminyltransferase 4 (GalNAc-T4) | 8693 | Glycan |
| 565 | GALNT5 | UDP-N-acetyl-alpha-D-galactosamine: polypeptide N-acetylgalactosaminyltransferase 5 (GalNAc-T5) | 11227 | Glycan |
| 566 | GALNT6 | UDP-N-acetyl-alpha-D-galactosamine: polypeptide N-acetylgalactosaminyltransferase 6 (GalNAc-T6) | 11226 | Glycan |
| 567 | GALNT7 | UDP-N-acetyl-alpha-D-galactosamine: polypeptide N-acetylgalactosaminyltransferase 7 (GalNAc-T7) | 51809 | Glycan |
| 568 | GALNT8 | UDP-N-acetyl-alpha-D-galactosamine: polypeptide N-acetylgalactosaminyltransferase 8 (GalNAc-T8) | 26290 | Glycan |
| 569 | GALNT9 | UDP-N-acetyl-alpha-D-galactosamine: polypeptide N-acetylgalactosaminyltransferase 9 (GalNAc-T9) | 50614 | Glycan |
| 570 | GALNTL1 | UDP-N-acetyl-alpha-D-galactosamine: polypeptide N-acetylgalactosaminyltransferase-like 1 | 57452 | Glycan |
| 571 | GALNTL2 | UDP-N-acetyl-alpha-D-galactosamine: polypeptide N-acetylgalactosaminyltransferase-like 2 | 117248 | Glycan |
| 572 | GALNTL4 | UDP-N-acetyl-alpha-D-galactosamine: polypeptide N-acetylgalactosaminyltransferase-like 4 | 374378 | Glycan |
| 573 | GALNTL5 | UDP-N-acetyl-alpha-D-galactosamine: polypeptide N-acetylgalactosaminyltransferase-like 5 | 168391 | Glycan |
| 574 | GALNTL6 | UDP-N-acetyl-alpha-D-galactosamine: polypeptide N-acetylgalactosaminyltransferase-like 6 | 442117 | Glycan |
| 575 | GANAB | glucosidase, alpha; neutral AB | 23193 | Glycan |
| 576 | GANC | glucosidase, alpha; neutral C | 2595 | Glycan |
| 577 | GBA | glucosidase, beta; acid (includes glucosylceramidase) | 2629 | Glycan |
| 578 | GBA2 | glucosidase, beta (bile acid) 2 | 57704 | Glycan |
| 579 | GBA3 | glucosidase, beta, acid 3 (cytosolic) | 57733 | Glycan |
| 580 | GBGT1 | globoside alpha-1,3-N-acetylgalactosaminyltransferase 1 | 26301 | Glycan |
| 581 | GCNT1 | glucosaminyl (N-acetyl) transferase 1, core 2 (beta-1,6-N-acetylglucosaminyltransferase) | 2650 | Glycan |
| 582 | GCNT2 | glucosaminyl (N-acetyl) transferase 2, I-branching enzyme (I blood group) | 2651 | Glycan |
| 583 | GCNT3 | glucosaminyl (N-acetyl) transferase 3, mucin type | 9245 | Glycan |
| 584 | GCNT4 | glucosaminyl (N-acetyl) transferase 4, core 2 (beta-1,6-N-acetylglucosaminyltransferase) | 51301 | Glycan |
| 585 | GCNT6 | glucosaminyl (N-acetyl) transferase family member 6 | 644378 | Glycan |
| 586 | GCNT7 | glucosaminyl (N-acetyl) transferase family member 7 | 140687 | Glycan |
| 587 | GCS1 | glucosidase I | 7841 | Glycan |
| 588 | GLT1D1 | glycosyltransferase 1 domain containing 1 | 144423 | Glycan |
| 589 | GLT25D1 | glycosyltransferase 25 domain containing 1 | 79709 | Glycan |
| 590 | GLT25D2 | glycosyltransferase 25 domain containing 2 | 23127 | Glycan |
| 591 | GLT6D1 | glycosyltransferase 6 domain containing 1 | 360203 | Glycan |
| 592 | GLT8D1 | glycosyltransferase 8 domain containing 1 | 55830 | Glycan |
| 593 | GLT8D2 | glycosyltransferase 8 domain containing 2 | 83468 | Glycan |
| 594 | GLT8D3 | glycosyltransferase 8 domain containing 3 | 283464 | Glycan |
| 595 | GLT8D4 | glycosyltransferase 8 domain containing 4 | 727936 | Glycan |
| 596 | GTDC1 | glycosyltransferase-like domain containing 1 | 79712 | Glycan |
| 597 | GYLTL1B | glycosyltransferase-like 1B | 120071 | Glycan |
| 598 | HAS1 | hyaluronan synthase 1 | 3036 | Glycan |
| 599 | HAS2 | hyaluronan synthase 2 | 3037 | Glycan |
| 600 | HAS3 | hyaluronan synthase 3 | 3038 | Glycan |
| 601 | LARGE | like-glycosyltransferase | 9215 | Glycan |
| 602 | LYCAT | lysocardiolipin acyltransferase | 253558 | Glycan |
| 603 | MFNG | MFNG O-fucosylpeptide 3-beta-N-acetylglucosaminyltransferase | 4242 | Glycan |
| 604 | MGAT1 | mannosyl (alpha-1,3-)-glycoprotein beta-1,2-N-acetylglucosaminyltransferase | 4245 | Glycan |
| 605 | MGAT2 | mannosyl (alpha-1,6-)-glycoprotein beta-1,2-N-acetylglucosaminyltransferase | 4247 | Glycan |
| 606 | MGAT3 | mannosyl (beta-1,4-)-glycoprotein beta-1,4-N-acetylglucosaminyltransferase | 4248 | Glycan |
| 607 | MGAT4A | Mannosyl (alpha-1,3-)-glycoprotein beta-1,4-N-acetylglucosaminyltransferase, isozyme A | 11320 | Glycan |
| 608 | MGAT4B | mannosyl (alpha-1,3-)-glycoprotein beta-1,4-N-acetylglucosaminyltransferase, isozyme B | 11282 | Glycan |
| 609 | MGAT4C | mannosyl (alpha-1,3-)-glycoprotein beta-1,4-N-acetylglucosaminyltransferase, isozyme C (putative) | 25834 | Glycan |
| 610 | MGAT5 | mannosyl (alpha-1,6-)-glycoprotein beta-1,6-N-acetyl-glucosaminyltransferase | 4249 | Glycan |
| 611 | MGAT5B | mannosyl (alpha-1,6-)-glycoprotein beta-1,6-N-acetyl-glucosaminyltransferase, isozyme B | 146664 | Glycan |
| 612 | NAGA | N-acetylgalactosaminidase, alpha- | 4668 | Glycan |
| 613 | NAGPA | N-acetylglucosamine-1-phosphodiester alpha-N-acetylglucosaminidase | 51172 | Glycan |
| 614 | POFUT1 | protein O-fucosyltransferase 1 | 23509 | Glycan |
| 615 | POFUT2 | protein O-fucosyltransferase 2 | 23275 | Glycan |
| 616 | POMGNT1 | protein O-linked mannose beta1,2-N-acetylglucosaminyltransferase | 55624 | Glycan |
| 617 | POMT1 | protein-O-mannosyltransferase 1 | 10585 | Glycan |
| 618 | POMT2 | protein-O-mannosyltransferase 2 | 29954 | Glycan |
| 619 | ST3GAL1 | ST3 beta-galactoside alpha-2,3-sialyltransferase 1 | 6482 | Glycan |
| 620 | ST3GAL2 | ST3 beta-galactoside alpha-2,3-sialyltransferase 2 | 6483 | Glycan |
| 621 | ST3GAL3 | ST3 beta-galactoside alpha-2,3-sialyltransferase 3 | 6487 | Glycan |
| 622 | ST3GAL4 | ST3 beta-galactoside alpha-2,3-sialyltransferase 4 | 6484 | Glycan |
| 623 | ST3GAL5 | ST3 beta-galactoside alpha-2,3-sialyltransferase 5 | 8869 | Glycan |
| 624 | ST3GAL6 | ST3 beta-galactoside alpha-2,3-sialyltransferase 6 | 10402 | Glycan |
| 625 | ST6GAL1 | ST6 beta-galactosamide alpha-2,6-sialyltranferase 1 | 6480 | Glycan |
| 626 | ST6GAL2 | ST6 beta-galactosamide alpha-2,6-sialyltranferase 2 | 84620 | Glycan |
| 627 | ST6GALNAC1 | ST6 (alpha-N-acetyl-neuraminyl-2,3-beta-galactosyl-1,3)-N-acetylgalactosaminide alpha-2,6-sialyltransferase 1 | 55808 | Glycan |
| 628 | ST6GALNAC2 | ST6 (alpha-N-acetyl-neuraminyl-2,3-beta-galactosyl-1,3)-N-acetylgalactosaminide alpha-2,6-sialyltransferase 2 | 10610 | Glycan |
| 629 | ST6GALNAC3 | ST6 (alpha-N-acetyl-neuraminyl-2,3-beta-galactosyl-1,3)-N-acetylgalactosaminide alpha-2,6-sialyltransferase 3 | 256435 | Glycan |
| 630 | ST6GALNAC4 | ST6 (alpha-N-acetyl-neuraminyl-2,3-beta-galactosyl-1,3)-N-acetylgalactosaminide alpha-2,6-sialyltransferase 4 | 27090 | Glycan |
| 631 | ST6GALNAC5 | ST6 (alpha-N-acetyl-neuraminyl-2,3-beta-galactosyl-1,3)-N-acetylgalactosaminide alpha-2,6-sialyltransferase 5 | 81849 | Glycan |
| 632 | ST6GALNAC6 | ST6 (alpha-N-acetyl-neuraminyl-2,3-beta-galactosyl-1,3)-N-acetylgalactosaminide alpha-2,6-sialyltransferase 6 | 30815 | Glycan |
| 633 | ST8SIA1 | ST8 alpha-N-acetyl-neuraminide alpha-2,8-sialyltransferase 1 | 6489 | Glycan |
| 634 | ST8SIA2 | ST8 alpha-N-acetyl-neuraminide alpha-2,8-sialyltransferase 2 | 8128 | Glycan |
| 635 | ST8SIA3 | ST8 alpha-N-acetyl-neuraminide alpha-2,8-sialyltransferase 3 | 51046 | Glycan |
| 636 | ST8SIA4 | ST8 alpha-N-acetyl-neuraminide alpha-2,8-sialyltransferase 4 | 7903 | Glycan |
| 637 | ST8SIA5 | ST8 alpha-N-acetyl-neuraminide alpha-2,8-sialyltransferase 5 | 29906 | Glycan |
| 638 | ST8SIA6 | ST8 alpha-N-acetyl-neuraminide alpha-2,8-sialyltransferase 6 | 338596 | Glycan |
| 639 | WBSCR17 | Williams-Beuren syndrome chromosome region 17 | 64409 | Glycan |
| 640 | XYLT1 | xylosyltransferase I | 64131 | Glycan |
| 641 | XYLT2 | xylosyltransferase II | 64132 | Glycan |
| 642 | DPM1 | dolichyl-phosphate mannosyltransferase polypeptide 1, catalytic subunit | 8813 | Glycan (Anchor) |
| 643 | DPM2 | dolichyl-phosphate mannosyltransferase polypeptide 2, regulatory subunit | 8818 | Glycan (Anchor) |
| 644 | DPM3 | dolichyl-phosphate mannosyltransferase polypeptide 3 | 54344 | Glycan (Anchor) |
| 645 | GPLD1 | glycosylphosphatidylinositol specific phospholipase D1 | 2822 | Glycan (Anchor) |
| 646 | PIGA | phosphatidylinositol glycan anchor biosynthesis, class A (paroxysmal nocturnal hemoglobinuria) | 5277 | Glycan (Anchor) |
| 647 | PIGB | phosphatidylinositol glycan anchor biosynthesis, class B | 9488 | Glycan (Anchor) |
| 648 | PIGC | phosphatidylinositol glycan anchor biosynthesis, class C | 5279 | Glycan (Anchor) |
| 649 | PIGF | phosphatidylinositol glycan anchor biosynthesis, class F | 5281 | Glycan (Anchor) |
| 650 | PIGG | phosphatidylinositol glycan anchor biosynthesis, class G | 54872 | Glycan (Anchor) |
| 651 | PIGH | phosphatidylinositol glycan anchor biosynthesis, class H | 5283 | Glycan (Anchor) |
| 652 | PIGK | phosphatidylinositol glycan anchor biosynthesis, class K | 10026 | Glycan (Anchor) |
| 653 | PIGL | phosphatidylinositol glycan anchor biosynthesis, class L | 9487 | Glycan (Anchor) |
| 654 | PIGM | phosphatidylinositol glycan anchor biosynthesis, class M | 93183 | Glycan (Anchor) |
| 655 | PIGN | phosphatidylinositol glycan anchor biosynthesis, class N | 23556 | Glycan (Anchor) |
| 656 | PIGO | phosphatidylinositol glycan anchor biosynthesis, class O | 84720 | Glycan (Anchor) |
| 657 | PIGP | phosphatidylinositol glycan anchor biosynthesis, class P | 51227 | Glycan (Anchor) |
| 658 | PIGQ | phosphatidylinositol glycan anchor biosynthesis, class Q | 9091 | Glycan (Anchor) |
| 659 | PIGS | phosphatidylinositol glycan anchor biosynthesis, class S | 94005 | Glycan (Anchor) |
| 660 | PIGT | phosphatidylinositol glycan anchor biosynthesis, class T | 51604 | Glycan (Anchor) |
| 661 | PIGU | phosphatidylinositol glycan anchor biosynthesis, class U | 128869 | Glycan (Anchor) |
| 662 | PIGV | phosphatidylinositol glycan anchor biosynthesis, class V | 55650 | Glycan (Anchor) |
| 663 | PIGW | phosphatidylinositol glycan anchor biosynthesis, class W | 284098 | Glycan (Anchor) |
| 664 | PIGX | phosphatidylinositol glycan anchor biosynthesis, class X | 54965 | Glycan (Anchor) |
| 665 | PIGY | phosphatidylinositol glycan anchor biosynthesis, class Y | 84992 | Glycan (Anchor) |
| 666 | PIGZ | phosphatidylinositol glycan anchor biosynthesis, class Z | 80235 | Glycan (Anchor) |
| 667 | AGA | aspartylglucosaminidase | 175 | Glycan (Degradation) |
| 668 | EDEM1 | ER degradation enhancer, mannosidase alpha-like 1 | 9695 | Glycan (Degradation) |
| 669 | EDEM2 | ER degradation enhancer, mannosidase alpha-like 2 | 55741 | Glycan (Degradation) |
| 670 | EDEM3 | ER degradation enhancer, mannosidase alpha-like 3 | 80267 | Glycan (Degradation) |
| 671 | FLJ21865 | endo-beta-N-acetylglucosaminidase | 64772 | Glycan (Degradation) |
| 672 | FUCA1 | fucosidase, alpha-L- 1, tissue | 2517 | Glycan (Degradation) |
| 673 | FUCA2 | fucosidase, alpha-L- 2, plasma | 2519 | Glycan (Degradation) |
| 674 | GALNS | galactosamine (N-acetyl)-6-sulfate sulfatase (Morquio syndrome, mucopolysaccharidosis type IVA) | 2588 | Glycan (Degradation) |
| 675 | GLA | galactosidase, alpha | 2717 | Glycan (Degradation) |
| 676 | GLB1 | galactosidase, beta 1 | 2720 | Glycan (Degradation) |
| 677 | GLB1L | galactosidase, beta 1-like | 79411 | Glycan (Degradation) |
| 678 | GLB1L2 | galactosidase, beta 2-like | 89944 | Glycan (Degradation) |
| 679 | GLB1L3 | galactosidase, beta 1 like 3 | 112937 | Glycan (Degradation) |
| 680 | GNS | glucosamine (N-acetyl)-6-sulfatase (Sanfilippo disease IIID) | 2799 | Glycan (Degradation) |
| 681 | GUSB | glucuronidase, beta | 2990 | Glycan (Degradation) |
| 682 | HEXA | hexosaminidase A (alpha polypeptide) | 3073 | Glycan (Degradation) |
| 683 | HEXB | hexosaminidase B (beta polypeptide) | 3074 | Glycan (Degradation) |
| 684 | HEXDC | hexosaminidase (glycosyl hydrolase family 20, catalytic domain) containing | 284004 | Glycan (Degradation) |
| 685 | HYAL1 | hyaluronoglucosaminidase 1 | 3373 | Glycan (Degradation) |
| 686 | HYAL2 | hyaluronoglucosaminidase 2 | 8692 | Glycan (Degradation) |
| 687 | HYAL3 | hyaluronoglucosaminidase 3 | 8372 | Glycan (Degradation) |
| 688 | HYAL4 | hyaluronoglucosaminidase 4 | 23553 | Glycan (Degradation) |
| 689 | IDS | iduronate 2-sulfatase (Hunter syndrome) | 3423 | Glycan (Degradation) |
| 690 | IDUA | iduronidase, alpha-L- | 3425 | Glycan (Degradation) |
| 691 | LYG1 | lysozyme G-like 1 | 129530 | Glycan (Degradation) |
| 692 | LYZ | lysozyme (renal amyloidosis) | 4069 | Glycan (Degradation) |
| 693 | LYZL1 | lysozyme-like 1 | 119180 | Glycan (Degradation) |
| 694 | LYZL2 | lysozyme-like 2 | 84569 | Glycan (Degradation) |
| 695 | LYZL4 | lysozyme-like 4 | 131375 | Glycan (Degradation) |
| 696 | LYZL6 | lysozyme-like 6 | 57151 | Glycan (Degradation) |
| 697 | MAN1A1 | mannosidase, alpha, class 1A, member 1 | 4121 | Glycan (Degradation) |
| 698 | MAN1A2 | mannosidase, alpha, class 1A, member 2 | 10905 | Glycan (Degradation) |
| 699 | MAN1B1 | mannosidase, alpha, class 1B, member 1 | 11253 | Glycan (Degradation) |
| 700 | MAN1C1 | mannosidase, alpha, class 1C, member 1 | 57134 | Glycan (Degradation) |
| 701 | MAN2A1 | mannosidase, alpha, class 2A, member 1 | 4124 | Glycan (Degradation) |
| 702 | MAN2A2 | mannosidase, alpha, class 2A, member 2 | 4122 | Glycan (Degradation) |
| 703 | MAN2B1 | mannosidase, alpha, class 2B, member 1 | 4125 | Glycan (Degradation) |
| 704 | MAN2B2 | mannosidase, alpha, class 2B, member 2 | 23324 | Glycan (Degradation) |
| 705 | MAN2C1 | mannosidase, alpha, class 2C, member 1 | 4123 | Glycan (Degradation) |
| 706 | MANBA | mannosidase, beta A, lysosomal | 4126 | Glycan (Degradation) |
| 707 | MANBAL | mannosidase, beta A, lysosomal-like | 63905 | Glycan (Degradation) |
| 708 | MANEA | mannosidase, endo-alpha | 79694 | Glycan (Degradation) |
| 709 | MANEAL | mannosidase, endo-alpha-like | 149175 | Glycan (Degradation) |
| 710 | NAGLU | N-acetylglucosaminidase, alpha- (Sanfilippo disease IIIB) | 4669 | Glycan (Degradation) |
| 711 | NEU1 | sialidase 1 (lysosomal sialidase) | 4758 | Glycan (Degradation) |
| 712 | NEU2 | sialidase 2 (cytosolic sialidase) | 4759 | Glycan (Degradation) |
| 713 | NEU3 | sialidase 3 (membrane sialidase) | 10825 | Glycan (Degradation) |
| 714 | NEU4 | sialidase 4 | 129807 | Glycan (Degradation) |
| 715 | CHPF | chondroitin polymerizing factor | 79586 | Glycan (Sulfate) |
| 716 | CHPF2 | chondroitin sulfate glucuronyltransferase | 54480 | Glycan (Sulfate) |
| 717 | CHST1 | carbohydrate (keratan sulfate Gal-6) sulfotransferase 1 | 8534 | Glycan (Sulfate) |
| 718 | CHST10 | carbohydrate sulfotransferase 10 | 9486 | Glycan (Sulfate) |
| 719 | CHST11 | carbohydrate (chondroitin 4) sulfotransferase 11 | 50515 | Glycan (Sulfate) |
| 720 | CHST12 | carbohydrate (chondroitin 4) sulfotransferase 12 | 55501 | Glycan (Sulfate) |
| 721 | CHST13 | carbohydrate (chondroitin 4) sulfotransferase 13 | 166012 | Glycan (Sulfate) |
| 722 | CHST14 | carbohydrate (N-acetylgalactosamine 4-0) sulfotransferase 14 | 113189 | Glycan (Sulfate) |
| 723 | CHST2 | carbohydrate (N-acetylglucosamine-6-O) sulfotransferase 2 | 9435 | Glycan (Sulfate) |
| 724 | CHST3 | carbohydrate (chondroitin 6) sulfotransferase 3 | 9469 | Glycan (Sulfate) |
| 725 | CHST4 | carbohydrate (N-acetylglucosamine 6-O) sulfotransferase 4 | 10164 | Glycan (Sulfate) |
| 726 | CHST5 | carbohydrate (N-acetylglucosamine 6-O) sulfotransferase 5 | 23563 | Glycan (Sulfate) |
| 727 | CHST6 | carbohydrate (N-acetylglucosamine 6-O) sulfotransferase 6 | 4166 | Glycan (Sulfate) |
| 728 | CHST7 | carbohydrate (N-acetylglucosamine 6-O) sulfotransferase 7 | 56548 | Glycan (Sulfate) |
| 729 | CHST8 | carbohydrate (N-acetylgalactosamine 4-0) sulfotransferase 8 | 64377 | Glycan (Sulfate) |
| 730 | CHST9 | carbohydrate (N-acetylgalactosamine 4-0) sulfotransferase 9 | 83539 | Glycan (Sulfate) |
| 731 | CHSY1 | carbohydrate (chondroitin) synthase 1 | 22856 | Glycan (Sulfate) |
| 732 | CHSY3 | chondroitin synthase-2 | 337876 | Glycan (Sulfate) |
| 733 | CSGALNACT1 | chondroitin sulfate N-acetylgalactosaminyltransferase 1 | 55790 | Glycan (Sulfate) |
| 734 | CSGALNACT2 | chondroitin sulfate N-acetylgalactosaminyltransferase 2 | 55454 | Glycan (Sulfate) |
| 735 | GAL3ST1 | galactose-3-O-sulfotransferase 1 | 9514 | Glycan (Sulfate) |
| 736 | GAL3ST2 | galactose-3-O-sulfotransferase 2 | 64090 | Glycan (Sulfate) |
| 737 | GAL3ST3 | galactose-3-O-sulfotransferase 3 | 89792 | Glycan (Sulfate) |
| 738 | GAL3ST4 | galactose-3-O-sulfotransferase 4 | 79690 | Glycan (Sulfate) |
| 739 | SGSH | N-sulfoglucosamine sulfohydrolase (sulfamidase) | 6448 | Glycan (Sulfate) |
| 740 | SULF1 | sulfatase 1 | 23213 | Glycan (Sulfate) |
| 741 | SULF2 | sulfatase 2 | 55959 | Glycan (Sulfate) |
| 742 | UST | uronyl-2-sulfotransferase | 10090 | Glycan (Sulfate) |
| 743 | LDHA | lactate dehydrogenase A | 3939 | Glycolysis |
| 744 | LDHAL6A | lactate dehydrogenase A-like 6A | 160287 | Glycolysis |
| 745 | LDHAL6B | lactate dehydrogenase A-like 6B | 92483 | Glycolysis |
| 746 | LDHB | lactate dehydrogenase B | 3945 | Glycolysis |
| 747 | LDHC | lactate dehydrogenase C | 3948 | Glycolysis |
| 748 | LDHD | lactate dehydrogenase D | 197257 | Glycolysis |
| 749 | ADPGK | ADP-dependent glucokinase | 83440 | Glycolysis |
| 750 | ALDOA | aldolase A, fructose-bisphosphate | 226 | Glycolysis |
| 751 | ALDOB | aldolase B, fructose-bisphosphate | 229 | Glycolysis |
| 752 | ALDOC | aldolase C, fructose-bisphosphate | 230 | Glycolysis |
| 753 | BPGM | 2,3-bisphosphoglycerate mutase | 669 | Glycolysis |
| 754 | DLAT | dihydrolipoamide S-acetyltransferase (E2 component of pyruvate dehydrogenase complex) | 1737 | Glycolysis |
| 755 | ENO1 | enolase 1, (alpha) | 2023 | Glycolysis |
| 756 | ENO2 | enolase 2 (gamma, neuronal) | 2026 | Glycolysis |
| 757 | ENO3 | enolase 3 (beta, muscle) | 2027 | Glycolysis |
| 758 | GAPDH | glyceraldehyde-3-phosphate dehydrogenase | 2597 | Glycolysis |
| 759 | GAPDHS | glyceraldehyde-3-phosphate dehydrogenase, spermatogenic | 26330 | Glycolysis |
| 760 | GCK | glucokinase (hexokinase 4, maturity onset diabetes of the young 2) | 2645 | Glycolysis |
| 761 | GPI | glucose phosphate isomerase | 2821 | Glycolysis |
| 762 | HK1 | hexokinase 1 | 3098 | Glycolysis |
| 763 | HK2 | hexokinase 2 | 3099 | Glycolysis |
| 764 | HK3 | hexokinase 3 (white cell) | 3101 | Glycolysis |
| 765 | HKDC1 | hexokinase domain containing 1 | 80201 | Glycolysis |
| 766 | PFKFB1 | 6-phosphofructo-2-kinase/fructose-2,6-biphosphatase 1 | 5207 | Glycolysis |
| 767 | PFKFB2 | 6-phosphofructo-2-kinase/fructose-2,6-biphosphatase 2 | 5208 | Glycolysis |
| 768 | PFKFB3 | 6-phosphofructo-2-kinase/fructose-2,6-biphosphatase 3 | 5209 | Glycolysis |
| 769 | PFKFB4 | 6-phosphofructo-2-kinase/fructose-2,6-biphosphatase 4 | 5210 | Glycolysis |
| 770 | PFKL | phosphofructokinase, liver | 5211 | Glycolysis |
| 771 | PFKM | phosphofructokinase, muscle | 5213 | Glycolysis |
| 772 | PFKP | phosphofructokinase, platelet | 5214 | Glycolysis |
| 773 | PGAM1 | phosphoglycerate mutase 1 (brain) | 5223 | Glycolysis |
| 774 | PGAM2 | phosphoglycerate mutase 2 (muscle) | 5224 | Glycolysis |
| 775 | PGAM4 | phosphoglycerate mutase family member 4 | 441531 | Glycolysis |
| 776 | PGAM5 | phosphoglycerate mutase family member 5 | 192111 | Glycolysis |
| 777 | PGK1 | phosphoglycerate kinase 1 | 5230 | Glycolysis |
| 778 | PGK2 | phosphoglycerate kinase 2 | 5232 | Glycolysis |
| 779 | PGM1 | phosphoglucomutase 1 | 5236 | Glycolysis |
| 780 | PGM2 | phosphoglucomutase 2 | 55276 | Glycolysis |
| 781 | PGM2L1 | Phosphoglucomutase 2-like 1 | 283209 | Glycolysis |
| 782 | PGM5 | phosphoglucomutase 5 | 5239 | Glycolysis |
| 783 | PKLR | pyruvate kinase, liver and RBC | 5313 | Glycolysis |
| 784 | PKM2 | pyruvate kinase, muscle | 5315 | Glycolysis |
| 785 | TPI1 | triosephosphate isomerase 1 | 7167 | Glycolysis |
| 786 | UEVLD | UEV and lactate/malate dehyrogenase domains | 55293 | Glycolysis |
| 787 | GLO1 | glyoxalase I | 2739 | Glyoxylate |
| 788 | GLOD4 | glyoxalase domain containing 4 | 51031 | Glyoxylate |
| 789 | GLOD5 | glyoxalase domain containing 5 | 392465 | Glyoxylate |
| 790 | HAO2 | hydroxyacid oxidase 2 (long chain) | 51179 | Glyoxylate |
| 791 | HAO1 | hydroxyacid oxidase (glycolate oxidase) 1 | 54363 | Glyoxylate |
| 792 | EXT1 | exostoses (multiple) 1 | 2131 | Heparin Sulfate |
| 793 | EXT2 | exostoses (multiple) 2 | 2132 | Heparin Sulfate |
| 794 | EXT3 | exostoses (multiple) 3 | 2133 | Heparin Sulfate |
| 795 | EXTL1 | exostoses (multiple)-like 1 | 2134 | Heparin Sulfate |
| 796 | EXTL2 | exostoses (multiple)-like 2 | 2135 | Heparin Sulfate |
| 797 | EXTL3 | exostoses (multiple)-like 3 | 2137 | Heparin Sulfate |
| 798 | GLCE | glucuronic acid epimerase | 26035 | Heparin Sulfate |
| 799 | HGSNAT | heparan-alpha-glucosaminide N-acetyltransferase | 138050 | Heparin Sulfate |
| 800 | HPSE | heparanase | 10855 | Heparin Sulfate |
| 801 | HPSE2 | heparanase 2 | 60495 | Heparin Sulfate |
| 802 | HS2ST1 | heparan sulfate 2-O-sulfotransferase 1 | 9653 | Heparin Sulfate |
| 803 | HS3ST1 | heparan sulfate (glucosamine) 3-O-sulfotransferase 1 | 9957 | Heparin Sulfate |
| 804 | HS3ST2 | heparan sulfate (glucosamine) 3-O-sulfotransferase 2 | 9956 | Heparin Sulfate |
| 805 | HS3ST3A1 | heparan sulfate (glucosamine) 3-O-sulfotransferase 3A1 | 9955 | Heparin Sulfate |
| 806 | HS3ST3B1 | Heparan sulfate (glucosamine) 3-O-sulfotransferase 3B1 | 9953 | Heparin Sulfate |
| 807 | HS3ST4 | heparan sulfate (glucosamine) 3-O-sulfotransferase 4 | 9951 | Heparin Sulfate |
| 808 | HS3ST5 | heparan sulfate (glucosamine) 3-O-sulfotransferase 5 | 222537 | Heparin Sulfate |
| 809 | HS3ST6 | heparan sulfate (glucosamine) 3-O-sulfotransferase 6 | 64711 | Heparin Sulfate |
| 810 | HS6ST1 | Heparan sulfate 6-O-sulfotransferase 1 | 9394 | Heparin Sulfate |
| 811 | HS6ST2 | heparan sulfate 6-O-sulfotransferase 2 | 90161 | Heparin Sulfate |
| 812 | HS6ST3 | heparan sulfate 6-O-sulfotransferase 3 | 266722 | Heparin Sulfate |
| 813 | NDST1 | N-deacetylase/N-sulfotransferase (heparan glucosaminyl) 1 | 3340 | Heparin Sulfate |
| 814 | NDST2 | N-deacetylase/N-sulfotransferase (heparan glucosaminyl) 2 | 8509 | Heparin Sulfate |
| 815 | NDST3 | N-deacetylase/N-sulfotransferase (heparan glucosaminyl) 3 | 9348 | Heparin Sulfate |
| 816 | NDST4 | N-deacetylase/N-sulfotransferase (heparan glucosaminyl) 4 | 64579 | Heparin Sulfate |
| 817 | ATP11A | ATPase, class VI, type 11A | 23250 | Ion Transport |
| 818 | ATP11B | ATPase, class VI, type 11B | 23200 | Ion Transport |
| 819 | ATP11C | ATPase, class VI, type 11C | 286410 | Ion Transport |
| 820 | ATP12A | ATPase, H+/K+ transporting, nongastric, alpha polypeptide | 479 | Ion Transport |
| 821 | ATP1A1 | ATPase, Na+/K+ transporting, alpha 1 polypeptide | 476 | Ion Transport |
| 822 | ATP1A2 | ATPase, Na+/K+ transporting, alpha 2 polypeptide | 477 | Ion Transport |
| 823 | ATP1A3 | ATPase, Na+/K+ transporting, alpha 3 polypeptide | 478 | Ion Transport |
| 824 | ATP1A4 | ATPase, Na+/K+ transporting, alpha 4 polypeptide | 480 | Ion Transport |
| 825 | ATP1B1 | ATPase, Na+/K+ transporting, beta 1 polypeptide | 481 | Ion Transport |
| 826 | ATP1B2 | ATPase, Na+/K+ transporting, beta 2 polypeptide | 482 | Ion Transport |
| 827 | ATP1B3 | ATPase, Na+/K+ transporting, beta 3 polypeptide | 483 | Ion Transport |
| 828 | ATP1B4 | ATPase, (Na+)/K+ transporting, beta 4 polypeptide | 23439 | Ion Transport |
| 829 | ATP2A1 | ATPase, Ca++ transporting, cardiac muscle, fast twitch 1 | 487 | Ion Transport |
| 830 | ATP2A2 | ATPase, Ca++ transporting, cardiac muscle, slow twitch 2 | 488 | Ion Transport |
| 831 | ATP2A3 | ATPase, Ca++ transporting, ubiquitous | 489 | Ion Transport |
| 832 | ATP2B1 | ATPase, Ca++ transporting, plasma membrane 1 | 490 | Ion Transport |
| 833 | ATP2B2 | ATPase, Ca++ transporting, plasma membrane 2 | 491 | Ion Transport |
| 834 | ATP2B3 | ATPase, Ca++ transporting, plasma membrane 3 | 492 | Ion Transport |
| 835 | ATP2B4 | ATPase, Ca++ transporting, plasma membrane 4 | 493 | Ion Transport |
| 836 | ATP2C1 | ATPase, Ca++-sequestering | 27032 | Ion Transport |
| 837 | ATP2C2 | ATPase, Ca++ transporting, type 2C, member 2 | 9914 | Ion Transport |
| 838 | ATP3 | ATPase, Mg++ transporting | 494 | Ion Transport |
| 839 | ATP4A | ATPase, H+/K+ exchanging, gastric, alpha polypeptide | 495 | Ion Transport |
| 840 | ATP4B | ATPase, H+/K+ exchanging, beta polypeptide | 496 | Ion Transport |
| 841 | ATP7A | ATPase, Cu++ transporting, alpha polypeptide | 538 | Ion Transport |
| 842 | ATP7B | ATPase, Cu++ transporting, beta polypeptide | 540 | Ion Transport |
| 843 | BEST1 | Bestrophin 1 | 7439 | Ion Transport |
| 844 | BEST2 | Bestrophin 2 | 54831 | Ion Transport |
| 845 | BEST3 | Bestrophin 3 | 144453 | Ion Transport |
| 846 | BEST4 | Bestrophin 4 | 266675 | Ion Transport |
| 847 | BSND | Bartter syndrome, infantile, with sensorineural deafness (Barttin) | 7809 | Ion Transport |
| 848 | C14orf68 | Chromosome 14 open reading frame 68 | 283600 | Ion Transport |
| 849 | C20orf59 | Chromosome 20 open reading frame 59 | 63910 | Ion Transport |
| 850 | C6orf192 | Chromosome 6 open reading frame 192 | 116843 | Ion Transport |
| 851 | CACNA1A | calcium channel, voltage-dependent, P/Q type, alpha 1A subunit | 773 | Ion Transport |
| 852 | CACNA1B | calcium channel, voltage-dependent, N type, alpha 1B subunit | 774 | Ion Transport |
| 853 | CACNA1C | calcium channel, voltage-dependent, L type, alpha 1C subunit | 775 | Ion Transport |
| 854 | CACNA1D | calcium channel, voltage-dependent, L type, alpha 1D subunit | 776 | Ion Transport |
| 855 | CACNA1E | calcium channel, voltage-dependent, R type, alpha 1E subunit | 777 | Ion Transport |
| 856 | CACNA1F | calcium channel, voltage-dependent, alpha 1F subunit | 778 | Ion Transport |
| 857 | CACNA1G | calcium channel, voltage-dependent, T type, alpha 1G subunit | 8913 | Ion Transport |
| 858 | CACNA1H | calcium channel, voltage-dependent, T type, alpha 1H subunit | 8912 | Ion Transport |
| 859 | CACNA1I | calcium channel, voltage-dependent, T type, alpha 1I subunit | 8911 | Ion Transport |
| 860 | CACNA1S | calcium channel, voltage-dependent, L type, alpha 1S subunit | 779 | Ion Transport |
| 861 | CACNA2D1 | calcium channel, voltage-dependent, alpha2/delta subunit 1 | 781 | Ion Transport |
| 862 | CACNA2D2 | calcium channel, voltage-dependent, alpha 2/delta subunit 2 | 9254 | Ion Transport |
| 863 | CACNA2D3 | calcium channel, voltage-dependent, alpha2/delta subunit 3 | 55799 | Ion Transport |
| 864 | CACNA2D4 | calcium channel, voltage-dependent, alpha 2/delta subunit 4 | 93589 | Ion Transport |
| 865 | CACNB1 | calcium channel, voltage-dependent, beta 1 subunit | 782 | Ion Transport |
| 866 | CACNB2 | calcium channel, voltage-dependent, beta 2 subunit | 783 | Ion Transport |
| 867 | CACNB3 | calcium channel, voltage-dependent, beta 3 subunit | 784 | Ion Transport |
| 868 | CACNB4 | calcium channel, voltage-dependent, beta 4 subunit | 785 | Ion Transport |
| 869 | CACNG1 | calcium channel, voltage-dependent, gamma subunit 1 | 786 | Ion Transport |
| 870 | CACNG2 | calcium channel, voltage-dependent, gamma subunit 2 | 10369 | Ion Transport |
| 871 | CACNG3 | calcium channel, voltage-dependent, gamma subunit 3 | 10368 | Ion Transport |
| 872 | CACNG4 | calcium channel, voltage-dependent, gamma subunit 4 | 27092 | Ion Transport |
| 873 | CACNG5 | calcium channel, voltage-dependent, gamma subunit 5 | 27091 | Ion Transport |
| 874 | CACNG6 | calcium channel, voltage-dependent, gamma subunit 6 | 59285 | Ion Transport |
| 875 | CACNG7 | calcium channel, voltage-dependent, gamma subunit 7 | 59284 | Ion Transport |
| 876 | CACNG8 | calcium channel, voltage-dependent, gamma subunit 8 | 59283 | Ion Transport |
| 877 | CATSPER1 | cation channel, sperm associated 1 | 117144 | Ion Transport |
| 878 | CATSPER2 | cation channel, sperm associated 2 | 117155 | Ion Transport |
| 879 | CATSPER3 | cation channel, sperm associated 3 | 347732 | Ion Transport |
| 880 | CATSPERB | cation channel, sperm-associated, beta | 79820 | Ion Transport |
| 881 | CFTR | Cystic fibrosis transmembrane conductance regulator (ATP-binding cassette sub-family C, member 7) | 1080 | Ion Transport |
| 882 | CHRNA1 | Cholinergic receptor, nicotinic, alpha 1 (muscle) | 1134 | Ion Transport |
| 883 | CHRNA10 | Cholinergic receptor, nicotinic, alpha 10 | 57053 | Ion Transport |
| 884 | CHRNA2 | Cholinergic receptor, nicotinic, alpha 2 (neuronal) | 1135 | Ion Transport |
| 885 | CHRNA3 | Cholinergic receptor, nicotinic, alpha 3 | 1136 | Ion Transport |
| 886 | CHRNA4 | Cholinergic receptor, nicotinic, alpha 4 | 1137 | Ion Transport |
| 887 | CHRNA5 | Cholinergic receptor, nicotinic, alpha 5 | 1138 | Ion Transport |
| 888 | CHRNA6 | Cholinergic receptor, nicotinic, alpha 6 | 8973 | Ion Transport |
| 889 | CHRNA7 | Cholinergic receptor, nicotinic, alpha 7 | 1139 | Ion Transport |
| 890 | CHRNA9 | Cholinergic receptor, nicotinic, alpha 9 | 55584 | Ion Transport |
| 891 | CHRNB1 | Cholinergic receptor, nicotinic, beta 1 (muscle) | 1140 | Ion Transport |
| 892 | CHRNB2 | Cholinergic receptor, nicotinic, beta 2 (neuronal) | 1141 | Ion Transport |
| 893 | CHRNB3 | Cholinergic receptor, nicotinic, beta 3 | 1142 | Ion Transport |
| 894 | CHRNB4 | Cholinergic receptor, nicotinic, beta 4 | 1143 | Ion Transport |
| 895 | CHRND | Cholinergic receptor, nicotinic, delta | 1144 | Ion Transport |
| 896 | CHRNE | Cholinergic receptor, nicotinic, epsilon | 1145 | Ion Transport |
| 897 | CHRNG | Cholinergic receptor, nicotinic, gamma | 1146 | Ion Transport |
| 898 | CLCA1 | chloride channel calcium activated 1 | 1179 | Ion Transport |
| 899 | CLCA2 | chloride channel calcium activated 2 | 9635 | Ion Transport |
| 900 | CLCA3 | chloride channel calcium activated 3 | 9629 | Ion Transport |
| 901 | CLCA4 | chloride channel calcium activated 4 | 22802 | Ion Transport |
| 902 | CLCC1 | chloride channel CLIC-like 1 | 23155 | Ion Transport |
| 903 | CLCN1 | chloride channel 1 | 1180 | Ion Transport |
| 904 | CLCN2 | chloride channel 2 | 1181 | Ion Transport |
| 905 | CLCN3 | chloride channel 3 | 1182 | Ion Transport |
| 906 | CLCN4 | chloride channel 4-2 | 1183 | Ion Transport |
| 907 | CLCN5 | chloride channel 5 | 1184 | Ion Transport |
| 908 | CLCN6 | chloride channel 6 | 1185 | Ion Transport |
| 909 | CLCN7 | chloride channel 7 | 1186 | Ion Transport |
| 910 | CLCNKA | Chloride channel Ka | 1187 | Ion Transport |
| 911 | CLCNKB | chloride channel Kb | 1188 | Ion Transport |
| 912 | CLIC1 | chloride intracellular channel 1 | 1192 | Ion Transport |
| 913 | CLIC2 | chloride intracellular channel 2 | 1193 | Ion Transport |
| 914 | CLIC3 | chloride intracellular channel 3 | 9022 | Ion Transport |
| 915 | CLIC4 | chloride intracellular channel 4 (mitochondrial) | 25932 | Ion Transport |
| 916 | CLIC5 | chloride intracellular channel 5 | 53405 | Ion Transport |
| 917 | CLIC6 | chloride intracellular channel 6 | 54102 | Ion Transport |
| 918 | CNGA1 | cyclic nucleotide gated channel alpha 1 | 1259 | Ion Transport |
| 919 | CNGA2 | Cyclic nucleotide gated channel alpha 2 | 1260 | Ion Transport |
| 920 | CNGA3 | cyclic nucleotide gated channel alpha 3 | 1261 | Ion Transport |
| 921 | CNGA4 | cyclic nucleotide gated channel alpha 4 | 1262 | Ion Transport |
| 922 | CNGB1 | cyclic nucleotide gated channel beta 1 | 1258 | Ion Transport |
| 923 | CNGB3 | cyclic nucleotide gated channel beta 3 | 54714 | Ion Transport |
| 924 | GABRA1 | Gamma-aminobutyric acid (GABA) A receptor, alpha 1 | 2554 | Ion Transport |
| 925 | GABRA2 | Gamma-aminobutyric acid (GABA) A receptor, alpha 2 | 2555 | Ion Transport |
| 926 | GABRA3 | Gamma-aminobutyric acid (GABA) A receptor, alpha 3 | 2556 | Ion Transport |
| 927 | GABRA4 | Gamma-aminobutyric acid (GABA) A receptor, alpha 4 | 2557 | Ion Transport |
| 928 | GABRA5 | Gamma-aminobutyric acid (GABA) A receptor, alpha 5 | 2558 | Ion Transport |
| 929 | GABRA6 | Gamma-aminobutyric acid (GABA) A receptor, alpha 6 | 2559 | Ion Transport |
| 930 | GABRB1 | Gamma-aminobutyric acid (GABA) A receptor, beta 1 | 2560 | Ion Transport |
| 931 | GABRB2 | Gamma-aminobutyric acid (GABA) A receptor, beta 2 | 2561 | Ion Transport |
| 932 | GABRB3 | Gamma-aminobutyric acid (GABA) A receptor, beta 3 | 2562 | Ion Transport |
| 933 | GABRD | Gamma-aminobutyric acid (GABA) A receptor, delta | 2563 | Ion Transport |
| 934 | GABRE | Gamma-aminobutyric acid (GABA) A receptor, epsilon | 2564 | Ion Transport |
| 935 | GABRG1 | Gamma-aminobutyric acid (GABA) A receptor, gamma 1 | 2565 | Ion Transport |
| 936 | GABRG2 | Gamma-aminobutyric acid (GABA) A receptor, gamma 2 | 2566 | Ion Transport |
| 937 | GABRG3 | Gamma-aminobutyric acid (GABA) A receptor, gamma 3 | 2567 | Ion Transport |
| 938 | GABRP | Gamma-aminobutyric acid (GABA) A receptor, pi | 2568 | Ion Transport |
| 939 | GABRQ | Gamma-aminobutyric acid (GABA) receptor, theta | 55879 | Ion Transport |
| 940 | GABRR1 | Gamma-aminobutyric acid (GABA) receptor, rho 1 | 2569 | Ion Transport |
| 941 | GABRR2 | Gamma-aminobutyric acid (GABA) receptor, rho 2 | 2570 | Ion Transport |
| 942 | GABRR3 | Gamma-aminobutyric acid (GABA) receptor, rho 3 | 200959 | Ion Transport |
| 943 | GLRA1 | Glycine receptor, alpha 1 (startle disease/hyperekplexia) | 2741 | Ion Transport |
| 944 | GLRA2 | Glycine receptor, alpha 2 | 2742 | Ion Transport |
| 945 | GLRA3 | Glycine receptor, alpha 3 | 8001 | Ion Transport |
| 946 | GLRA4 | Glycine receptor, alpha 4 | 441509 | Ion Transport |
| 947 | GLRB | Glycine receptor, beta | 2743 | Ion Transport |
| 948 | GRIA1 | Glutamate receptor, ionotropic, AMPA 1 | 2890 | Ion Transport |
| 949 | GRIA2 | Glutamate receptor, ionotropic, AMPA 2 | 2891 | Ion Transport |
| 950 | GRIA3 | Glutamate receptor, ionotrophic, AMPA 3 | 2892 | Ion Transport |
| 951 | GRIA4 | Glutamate receptor, ionotrophic, AMPA 4 | 2893 | Ion Transport |
| 952 | GRID1 | Glutamate receptor, ionotropic, delta 1 | 2894 | Ion Transport |
| 953 | GRID2 | Glutamate receptor, ionotropic, delta 2 | 2895 | Ion Transport |
| 954 | GRIK1 | Glutamate receptor, ionotropic, kainate 1 | 2897 | Ion Transport |
| 955 | GRIK2 | Glutamate receptor, ionotropic, kainate 2 | 2898 | Ion Transport |
| 956 | GRIK3 | Glutamate receptor, ionotropic, kainate 3 | 2899 | Ion Transport |
| 957 | GRIK4 | Glutamate receptor, ionotropic, kainate 4 | 2900 | Ion Transport |
| 958 | GRIK5 | Glutamate receptor, ionotropic, kainate 5 | 2901 | Ion Transport |
| 959 | GRIN1 | Glutamate receptor, ionotropic, N-methyl D-aspartate 1 | 2902 | Ion Transport |
| 960 | GRIN2A | Glutamate receptor, ionotropic, N-methyl D-aspartate 2A | 2903 | Ion Transport |
| 961 | GRIN2B | Glutamate receptor, ionotropic, N-methyl D-aspartate 2B | 2904 | Ion Transport |
| 962 | GRIN2C | Glutamate receptor, ionotropic, N-methyl D-aspartate 2C | 2905 | Ion Transport |
| 963 | GRIN2D | Glutamate receptor, ionotropic, N-methyl D-aspartate 2D | 2906 | Ion Transport |
| 964 | GRIN3A | Glutamate receptor, ionotropic, N-methyl-D-aspartate 3A | 116443 | Ion Transport |
| 965 | GRIN3B | Glutamate receptor, ionotropic, N-methyl-D-aspartate 3B | 116444 | Ion Transport |
| 966 | HCN1 | hyperpolarization activated cyclic nucleotide-gated potassium channel 1 | 348980 | Ion Transport |
| 967 | HCN2 | hyperpolarization activated cyclic nucleotide-gated potassium channel 2 | 610 | Ion Transport |
| 968 | HCN3 | hyperpolarization activated cyclic nucleotide-gated potassium channel 3 | 57657 | Ion Transport |
| 969 | HCN4 | hyperpolarization activated cyclic nucleotide-gated potassium channel 4 | 10021 | Ion Transport |
| 970 | HEPH | hephaestin | 9843 | Ion Transport |
| 971 | HTR3A | 5-hydroxytryptamine (serotonin) receptor 3A | 3359 | Ion Transport |
| 972 | HTR3B | 5-hydroxytryptamine (serotonin) receptor 3B | 9177 | Ion Transport |
| 973 | HTR3C | 5-hydroxytryptamine (serotonin) receptor 3, family member C | 170572 | Ion Transport |
| 974 | HTR3D | 5-hydroxytryptamine (serotonin) receptor 3 family member D | 200909 | Ion Transport |
| 975 | HTR3E | 5-hydroxytryptamine (serotonin) receptor 3, family member E | 285242 | Ion Transport |
| 976 | ITPR1 | Inositol 1,4,5-triphosphate receptor, type 1 | 3708 | Ion Transport |
| 977 | ITPR2 | inositol 1,4,5-triphosphate receptor, type 2 | 3709 | Ion Transport |
| 978 | ITPR3 | inositol 1,4,5-triphosphate receptor, type 3 | 3710 | Ion Transport |
| 979 | KCNA1 | potassium voltage-gated channel, shaker-related subfamily, member 1 | 3736 | Ion Transport |
| 980 | KCNA10 | potassium voltage-gated channel, shaker-related subfamily, member 10 | 3744 | Ion Transport |
| 981 | KCNA2 | potassium voltage-gated channel, shaker-related subfamily, member 2 | 3737 | Ion Transport |
| 982 | KCNA3 | potassium voltage-gated channel, shaker-related subfamily, member 3 | 3738 | Ion Transport |
| 983 | KCNA4 | potassium voltage-gated channel, shaker-related subfamily, member 4 | 3739 | Ion Transport |
| 984 | KCNA5 | potassium voltage-gated channel, shaker-related subfamily, member 5 | 3741 | Ion Transport |
| 985 | KCNA6 | potassium voltage-gated channel, shaker-related, subfamily, member 6 | 3742 | Ion Transport |
| 986 | KCNA7 | potassium voltage-gated channel, shaker-related subfamily, member 7 | 3743 | Ion Transport |
| 987 | KCNAB1 | potassium voltage-gated channel, shaker-related subfamily, beta member 1 | 7881 | Ion Transport |
| 988 | KCNAB2 | potassium voltage-gated channel, shaker-related subfamily, beta member 2 | 8514 | Ion Transport |
| 989 | KCNAB3 | potassium voltage-gated channel, shaker-related subfamily, beta member 3 | 9196 | Ion Transport |
| 990 | KCNB1 | potassium voltage gated channel, Shab-related subfamily, member 1 | 3745 | Ion Transport |
| 991 | KCNB2 | potassium voltage gated channel, Shab-related subfamily, member 2 | 9312 | Ion Transport |
| 992 | KCNC1 | potassium voltage gated channel, Shaw-related subfamily, member 1 | 3746 | Ion Transport |
| 993 | KCNC2 | potassium voltage gated channel, Shaw-related subfamily, member 2 | 3747 | Ion Transport |
| 994 | KCNC3 | potassium voltage gated channel, Shaw-related subfamily, member 3 | 3748 | Ion Transport |
| 995 | KCNC4 | potassium voltage gated channel, Shaw-related subfamily, member 4 | 3749 | Ion Transport |
| 996 | KCND1 | potassium voltage-gated channel, Shal-related family, member 1 | 3750 | Ion Transport |
| 997 | KCND2 | potassium voltage-gated channel, Shal-related family, member 2 | 3751 | Ion Transport |
| 998 | KCND3 | potassium voltage-gated channel, Shal-related family, member 3 | 3752 | Ion Transport |
| 999 | KCNE1 | potassium voltage-gated channel, Isk-related subfamily, member 1 | 3753 | Ion Transport |
| 1000 | KCNE1L | potassium voltage-gated channel, Isk-related family, member 1-like | 23630 | Ion Transport |
| 1001 | KCNE2 | potassium voltage-gated channel, Isk-related subfamily, gene 2 | 9992 | Ion Transport |
| 1002 | KCNE3 | potassium voltage-gated channel, Isk-related subfamily, gene 3 | 10008 | Ion Transport |
| 1003 | KCNE4 | potassium voltage-gated channel, Isk-related subfamily, gene 4 | 23704 | Ion Transport |
| 1004 | KCNF1 | potassium voltage-gated channel, subfamily F, member 1 | 3754 | Ion Transport |
| 1005 | KCNG1 | potassium voltage-gated channel, subfamily G, member 1 | 3755 | Ion Transport |
| 1006 | KCNG2 | potassium voltage-gated channel, subfamily G, member 2 | 26251 | Ion Transport |
| 1007 | KCNG3 | potassium voltage-gated channel, subfamily G, member 3 | 170850 | Ion Transport |
| 1008 | KCNG4 | potassium voltage-gated channel, subfamily G, member 4 | 93107 | Ion Transport |
| 1009 | KCNH1 | potassium voltage-gated channel, subfamily H (eag-related), member 1 | 3756 | Ion Transport |
| 1010 | KCNH2 | potassium voltage-gated channel, subfamily H (eag-related), member 2 | 3757 | Ion Transport |
| 1011 | KCNH3 | potassium voltage-gated channel, subfamily H (eag-related), member 3 | 23416 | Ion Transport |
| 1012 | KCNH4 | potassium voltage-gated channel, subfamily H (eag-related), member 4 | 23415 | Ion Transport |
| 1013 | KCNH5 | potassium voltage-gated channel, subfamily H (eag-related), member 5 | 27133 | Ion Transport |
| 1014 | KCNH6 | potassium voltage-gated channel, subfamily H (eag-related), member 6 | 81033 | Ion Transport |
| 1015 | KCNH7 | potassium voltage-gated channel, subfamily H (eag-related), member 7 | 90134 | Ion Transport |
| 1016 | KCNH8 | potassium voltage-gated channel, subfamily H (eag-related), member 8 | 131096 | Ion Transport |
| 1017 | KCNJ1 | potassium inwardly-rectifying channel, subfamily J, member 1 | 3758 | Ion Transport |
| 1018 | KCNJ10 | potassium inwardly-rectifying channel, subfamily J, member 10 | 3766 | Ion Transport |
| 1019 | KCNJ11 | potassium inwardly rectifying channel, subfamily J, member 11 | 3767 | Ion Transport |
| 1020 | KCNJ12 | potassium inwardly-rectifying channel, subfamily J, member 12 | 3768 | Ion Transport |
| 1021 | KCNJ13 | potassium inwardly-rectifying channel, subfamily J, member 13 | 3769 | Ion Transport |
| 1022 | KCNJ14 | potassium inwardly-rectifying channel, subfamily J, member 14 | 3770 | Ion Transport |
| 1023 | KCNJ15 | potassium inwardly-rectifying channel, subfamily J, member 15 | 3772 | Ion Transport |
| 1024 | KCNJ16 | potassium inwardly-rectifying channel, subfamily J, member 16 | 3773 | Ion Transport |
| 1025 | KCNJ2 | potassium inwardly-rectifying channel, subfamily J, member 2 | 3759 | Ion Transport |
| 1026 | KCNJ3 | potassium inwardly-rectifying channel, subfamily J, member 3 | 3760 | Ion Transport |
| 1027 | KCNJ4 | potassium inwardly-rectifying channel, subfamily J, member 4 | 3761 | Ion Transport |
| 1028 | KCNJ5 | potassium inwardly-rectifying channel, subfamily J, member 5 | 3762 | Ion Transport |
| 1029 | KCNJ6 | potassium inwardly-rectifying channel, subfamily J, member 6 | 3763 | Ion Transport |
| 1030 | KCNJ8 | potassium inwardly-rectifying channel, subfamily J, member 8 | 3764 | Ion Transport |
| 1031 | KCNJ9 | potassium inwardly-rectifying channel, subfamily J, member 9 | 3765 | Ion Transport |
| 1032 | KCNK1 | potassium channel, subfamily K, member 1 | 3775 | Ion Transport |
| 1033 | KCNK10 | potassium channel, subfamily K, member 10 | 54207 | Ion Transport |
| 1034 | KCNK12 | potassium channel, subfamily K, member 12 | 56660 | Ion Transport |
| 1035 | KCNK13 | potassium channel, subfamily K, member 13 | 56659 | Ion Transport |
| 1036 | KCNK15 | potassium channel, subfamily K, member 15 | 60598 | Ion Transport |
| 1037 | KCNK16 | potassium channel, subfamily K, member 16 | 83795 | Ion Transport |
| 1038 | KCNK17 | potassium channel, subfamily K, member 17 | 89822 | Ion Transport |
| 1039 | KCNK18 | Potassium channel, subfamily K, member 18 | 338567 | Ion Transport |
| 1040 | KCNK2 | potassium channel, subfamily K, member 2 | 3776 | Ion Transport |
| 1041 | KCNK3 | potassium channel, subfamily K, member 3 | 3777 | Ion Transport |
| 1042 | KCNK4 | potassium channel, subfamily K, member 4 | 50801 | Ion Transport |
| 1043 | KCNK5 | potassium channel, subfamily K, member 5 | 8645 | Ion Transport |
| 1044 | KCNK6 | potassium inwardly-rectifying channel, subfamily K, member 6 | 9424 | Ion Transport |
| 1045 | KCNK7 | potassium channel, subfamily K, member 7 | 10089 | Ion Transport |
| 1046 | KCNK9 | potassium channel, subfamily K, member 9 | 51305 | Ion Transport |
| 1047 | KCNMA1 | potassium large conductance calcium-activated channel, subfamily M, alpha member 1 | 3778 | Ion Transport |
| 1048 | KCNMB1 | potassium large conductance calcium-activated channel, subfamily M, beta member 1 | 3779 | Ion Transport |
| 1049 | KCNMB2 | potassium large conductance calcium-activated channel, subfamily M, beta member 2 | 10242 | Ion Transport |
| 1050 | KCNMB3 | potassium large conductance calcium-activated channel, subfamily M beta member 3 | 27094 | Ion Transport |
| 1051 | KCNMB4 | potassium large conductance calcium-activated channel, subfamily M, beta member 4 | 27345 | Ion Transport |
| 1052 | KCNN1 | potassium intermediate/small conductance calcium-activated channel, subfamily N, member 1 | 3780 | Ion Transport |
| 1053 | KCNN2 | potassium intermediate/small conductance calcium-activated channel, subfamily N, member 2 | 3781 | Ion Transport |
| 1054 | KCNN3 | potassium intermediate/small conductance calcium-activated channel, subfamily N, member 3 | 3782 | Ion Transport |
| 1055 | KCNN4 | potassium intermediate/small conductance calcium-activated channel, subfamily N, member 4 | 3783 | Ion Transport |
| 1056 | KCNQ1 | potassium voltage-gated channel, subfamily Q, member 1 | 3784 | Ion Transport |
| 1057 | KCNQ2 | potassium voltage-gated channel, subfamily Q, member 2 | 3785 | Ion Transport |
| 1058 | KCNQ3 | potassium voltage-gated channel, subfamily Q, member 3 | 3786 | Ion Transport |
| 1059 | KCNQ4 | potassium voltage-gated channel, subfamily Q, member 4 | 9132 | Ion Transport |
| 1060 | KCNQ5 | potassium voltage-gated channel, subfamily Q, member 5 | 56479 | Ion Transport |
| 1061 | KCNS1 | K+ voltage-gated channel, subfamily S, 1 | 3787 | Ion Transport |
| 1062 | KCNS2 | K+ voltage-gated channel, subfamily S, 2 | 3788 | Ion Transport |
| 1063 | KCNS3 | potassium voltage-gated channel, delayed-rectifier, subfamily S, member 3 | 3790 | Ion Transport |
| 1064 | KCNT1 | potassium channel, subfamily T, member 1 | 57582 | Ion Transport |
| 1065 | KCNT2 | potassium channel, subfamily T, member 2 | 343450 | Ion Transport |
| 1066 | KCNU1 | potassium channel, subfamily U, member 1 | 157855 | Ion Transport |
| 1067 | KCNV1 | potassium channel, subfamily V, member 1 | 27012 | Ion Transport |
| 1068 | KCNV2 | potassium channel, subfamily V, member 2 | 169522 | Ion Transport |
| 1069 | KCTD1 | potassium channel tetramerisation domain containing 1 | 284252 | Ion Transport |
| 1070 | KCTD10 | potassium channel tetramerisation domain containing 10 | 83892 | Ion Transport |
| 1071 | KCTD11 | potassium channel tetramerisation domain containing 11 | 147040 | Ion Transport |
| 1072 | KCTD12 | potassium channel tetramerisation domain containing 12 | 115207 | Ion Transport |
| 1073 | KCTD13 | potassium channel tetramerisation domain containing 13 | 253980 | Ion Transport |
| 1074 | KCTD14 | potassium channel tetramerisation domain containing 14 | 65987 | Ion Transport |
| 1075 | KCTD15 | potassium channel tetramerisation domain containing 15 | 79047 | Ion Transport |
| 1076 | KCTD16 | potassium channel tetramerisation domain containing 16 | 57528 | Ion Transport |
| 1077 | KCTD17 | potassium channel tetramerisation domain containing 17 | 79734 | Ion Transport |
| 1078 | KCTD18 | potassium channel tetramerisation domain containing 18 | 130535 | Ion Transport |
| 1079 | KCTD19 | potassium channel tetramerisation domain containing 19 | 146212 | Ion Transport |
| 1080 | KCTD2 | potassium channel tetramerisation domain containing 2 | 23510 | Ion Transport |
| 1081 | KCTD20 | potassium channel tetramerisation domain containing 20 | 222658 | Ion Transport |
| 1082 | KCTD21 | potassium channel tetramerisation domain containing 21 | 283219 | Ion Transport |
| 1083 | KCTD3 | potassium channel tetramerisation domain containing 3 | 51133 | Ion Transport |
| 1084 | KCTD4 | potassium channel tetramerisation domain containing 4 | 386618 | Ion Transport |
| 1085 | KCTD5 | potassium channel tetramerisation domain containing 5 | 54442 | Ion Transport |
| 1086 | KCTD6 | potassium channel tetramerisation domain containing 6 | 200845 | Ion Transport |
| 1087 | KCTD7 | potassium channel tetramerisation domain containing 7 | 154881 | Ion Transport |
| 1088 | KCTD8 | potassium channel tetramerisation domain containing 8 | 386617 | Ion Transport |
| 1089 | KCTD9 | potassium channel tetramerisation domain containing 9 | 54793 | Ion Transport |
| 1090 | LGICZ1 | ligand-gated ion channel, zinc activated 1 | 353174 | Ion Transport |
| 1091 | LST-3TM12 | organic anion transporter LST-3b | 338821 | Ion Transport |
| 1092 | MAGT1 | magnesium transporter 1 | 84061 | Ion Transport |
| 1093 | MCOLN1 | Mucolipin 1 | 57192 | Ion Transport |
| 1094 | MCOLN2 | Mucolipin 2 | 255231 | Ion Transport |
| 1095 | MCOLN3 | Mucolipin 3 | 55283 | Ion Transport |
| 1096 | MMAA | methylmalonic aciduria (cobalamin deficiency) cblA type | 166785 | Ion Transport |
| 1097 | NALCN | sodium leak channel, non-selective | 259232 | Ion Transport |
| 1098 | NIPA1 | non imprinted in Prader-Willi/Angelman syndrome 1 | 123606 | Ion Transport |
| 1099 | OCA2 | Oculocutaneous albinism II (pink-eye dilution homolog, mouse) | 4948 | Ion Transport |
| 1100 | ORAI1 | ORAI calcium release-activated calcium modulator 1 | 84876 | Ion Transport |
| 1101 | ORAI2 | ORAI calcium release-activated calcium modulator 2 | 80228 | Ion Transport |
| 1102 | ORAI3 | ORAI calcium release-activated calcium modulator 3 | 93129 | Ion Transport |
| 1103 | P2RX1 | purinergic receptor P2X, ligand-gated ion channel, 1 | 5023 | Ion Transport |
| 1104 | P2RX2 | purinergic receptor P2X, ligand-gated ion channel, 2 | 22953 | Ion Transport |
| 1105 | P2RX3 | purinergic receptor P2X, ligand-gated ion channel, 3 | 5024 | Ion Transport |
| 1106 | P2RX4 | purinergic receptor P2X, ligand-gated ion channel, 4 | 5025 | Ion Transport |
| 1107 | P2RX5 | purinergic receptor P2X, ligand-gated ion channel, 5 | 5026 | Ion Transport |
| 1108 | P2RX6 | purinergic receptor P2X, ligand-gated ion channel, 6 | 9127 | Ion Transport |
| 1109 | P2RX7 | purinergic receptor P2X, ligand-gated ion channel, 7 | 5027 | Ion Transport |
| 1110 | PKD1 | Polycystic kidney disease 1 (autosomal dominant) | 5310 | Ion Transport |
| 1111 | PKD2 | Polycystic kidney disease 2 (autosomal dominant) | 5311 | Ion Transport |
| 1112 | PKD2L1 | Polycystic kidney disease 2-like 1 | 9033 | Ion Transport |
| 1113 | RYR1 | ryanodine receptor 1 (skeletal) | 6261 | Ion Transport |
| 1114 | RYR2 | ryanodine receptor 2 (cardiac) | 6262 | Ion Transport |
| 1115 | RYR3 | ryanodine receptor 3 | 6263 | Ion Transport |
| 1116 | SCN10A | sodium channel, voltage-gated, type X, alpha | 6336 | Ion Transport |
| 1117 | SCN11A | sodium channel, voltage-gated, type XI, alpha | 11280 | Ion Transport |
| 1118 | SCN1A | sodium channel, voltage-gated, type I, alpha | 6323 | Ion Transport |
| 1119 | SCN1B | sodium channel, voltage-gated, type I, beta | 6324 | Ion Transport |
| 1120 | SCN2A | sodium channel, voltage-gated, type II, alpha 1 | 6326 | Ion Transport |
| 1121 | SCN2B | sodium channel, voltage-gated, type II, beta | 6327 | Ion Transport |
| 1122 | SCN3A | sodium channel, voltage-gated, type III, alpha | 6328 | Ion Transport |
| 1123 | SCN3B | sodium channel, voltage-gated, type III, beta | 55800 | Ion Transport |
| 1124 | SCN4A | sodium channel, voltage-gated, type IV, alpha | 6329 | Ion Transport |
| 1125 | SCN4B | sodium channel, voltage-gated, type IV, beta | 6330 | Ion Transport |
| 1126 | SCN5A | sodium channel, voltage-gated, type V, alpha | 6331 | Ion Transport |
| 1127 | SCN7A | sodium channel, voltage-gated, type VII, alpha | 6332 | Ion Transport |
| 1128 | SCN8A | sodium channel, voltage-gated, type VIII, alpha | 6334 | Ion Transport |
| 1129 | SCN9A | sodium channel, voltage-gated, type IX, alpha | 6335 | Ion Transport |
| 1130 | SCNN1A | sodium channel, nonvoltage-gated, type I, alpha | 6337 | Ion Transport |
| 1131 | SCNN1B | sodium channel, nonvoltage-gated 1 beta | 6338 | Ion Transport |
| 1132 | SCNN1D | sodium channel, nonvoltage-gated 1, delta | 6339 | Ion Transport |
| 1133 | SCNN1G | sodium channel, nonvoltage-gated 1 gamma | 6340 | Ion Transport |
| 1134 | SLC11A1 | solute carrier family 11 (proton-coupled divalent metal ion transporters), member 1 | 6556 | Ion Transport |
| 1135 | SLC11A2 | solute carrier family 11 (proton-coupled divalent metal ion transporters), member 2 | 4891 | Ion Transport |
| 1136 | SLC12A1 | solute carrier family 12 (sodium/potassium/chloride transporters), member 1 | 6557 | Ion Transport |
| 1137 | SLC12A2 | solute carrier family 12, member 2 | 6558 | Ion Transport |
| 1138 | SLC12A3 | solute carrier family 12, member 3 | 6559 | Ion Transport |
| 1139 | SLC12A4 | solute carrier family 12, member 4 | 6560 | Ion Transport |
| 1140 | SLC12A5 | solute carrier family 12, member 5 | 57468 | Ion Transport |
| 1141 | SLC12A6 | solute carrier family 12, member 6 | 9990 | Ion Transport |
| 1142 | SLC12A7 | solute carrier family 12, member 7 | 10723 | Ion Transport |
| 1143 | SLC12A8 | solute carrier family 12 (potassium/chloride transporters), member 8 | 84561 | Ion Transport |
| 1144 | SLC12A9 | solute carrier family 12 (potassium/chloride transporters), member 9 | 56996 | Ion Transport |
| 1145 | SLC13A1 | solute carrier family 13 (sodium/sulphate symporters), member 1 | 6561 | Ion Transport |
| 1146 | SLC13A4 | solute carrier family 13 (sodium/sulfate symporters), member 4 | 26266 | Ion Transport |
| 1147 | SLC22A24 | Solute carrier family 22, member 24 | 283238 | Ion Transport |
| 1148 | SLC22A25 | Solute carrier family 22, member 25 | 387601 | Ion Transport |
| 1149 | SLC24A1 | solute carrier family 24 (sodium/potassium/calcium exchanger), member 1 | 9187 | Ion Transport |
| 1150 | SLC24A2 | solute carrier family 24 (sodium/potassium/calcium exchanger), member 2 | 25769 | Ion Transport |
| 1151 | SLC24A3 | solute carrier family 24 (sodium/potassium/calcium exchanger), member 3 | 57419 | Ion Transport |
| 1152 | SLC24A4 | solute carrier family 24 (sodium/potassium/calcium exchanger), member 4 | 123041 | Ion Transport |
| 1153 | SLC24A5 | solute carrier family 24, member 5 | 283652 | Ion Transport |
| 1154 | SLC24A6 | solute carrier family 24 (sodium/potassium/calcium exchanger), member 6 | 80024 | Ion Transport |
| 1155 | SLC30A1 | solute carrier family 30 (zinc transporter), member 1 | 7779 | Ion Transport |
| 1156 | SLC30A10 | solute carrier family 30, member 10 | 55532 | Ion Transport |
| 1157 | SLC30A2 | solute carrier family 30 (zinc transporter), member 2 | 7780 | Ion Transport |
| 1158 | SLC30A3 | solute carrier family 30 (zinc transporter), member 3 | 7781 | Ion Transport |
| 1159 | SLC30A4 | solute carrier family 30 (zinc transporter), member 4 | 7782 | Ion Transport |
| 1160 | SLC30A5 | solute carrier family 30 (zinc transporter), member 5 | 64924 | Ion Transport |
| 1161 | SLC30A6 | solute carrier family 30 (zinc transporter), member 6 | 55676 | Ion Transport |
| 1162 | SLC30A7 | solute carrier family 30 (zinc transporter), member 7 | 148867 | Ion Transport |
| 1163 | SLC30A8 | solute carrier family 30 (zinc transporter), member 8 | 169026 | Ion Transport |
| 1164 | SLC30A9 | solute carrier family 30 (zinc transporter), member 9 | 10463 | Ion Transport |
| 1165 | SLC31A1 | solute carrier family 31, member 1 | 1317 | Ion Transport |
| 1166 | SLC31A2 | solute carrier family 31, member 2 | 1318 | Ion Transport |
| 1167 | SLC39A1 | solute carrier family 39 (zinc transporter), member 1 | 27173 | Ion Transport |
| 1168 | SLC39A10 | solute carrier family 39 (zinc transporter), member 10 | 57181 | Ion Transport |
| 1169 | SLC39A11 | solute carrier family 39 (metal ion transporter), member 11 | 201266 | Ion Transport |
| 1170 | SLC39A12 | solute carrier family 39 (metal ion transporter), member 12 | 221074 | Ion Transport |
| 1171 | SLC39A13 | solute carrier family 39 (metal ion transporter), member 13 | 91252 | Ion Transport |
| 1172 | SLC39A14 | solute carrier family 39 (zinc transporter), member 14 | 23516 | Ion Transport |
| 1173 | SLC39A2 | solute carrier family 39 (zinc transporter), member 2 | 29986 | Ion Transport |
| 1174 | SLC39A3 | solute carrier family 39 (zinc transporter), member 3 | 29985 | Ion Transport |
| 1175 | SLC39A4 | solute carrier family 39 (zinc transporter), member 4 | 55630 | Ion Transport |
| 1176 | SLC39A5 | solute carrier family 39 (metal ion transporter), member 5 | 283375 | Ion Transport |
| 1177 | SLC39A6 | solute carrier family 39 (metal ion transporter), member 6 | 25800 | Ion Transport |
| 1178 | SLC39A7 | solute carrier family 39, member 7 | 7922 | Ion Transport |
| 1179 | SLC39A8 | solute carrier family 39 (metal ion transporter), member 8 | 64116 | Ion Transport |
| 1180 | SLC39A9 | solute carrier family 39 (zinc transporter), member 9 | 55334 | Ion Transport |
| 1181 | SLC3A1 | solute carrier family 3, member 1 | 6519 | Ion Transport |
| 1182 | SLC3A2 | solute carrier family 3 (activators of dibasic and neutral amino acid transport), member 2 | 6520 | Ion Transport |
| 1183 | SLC40A1 | solute carrier family 40 (iron-regulated transporter), member 1 | 30061 | Ion Transport |
| 1184 | SLC41A1 | solute carrier family 41, member 1 | 254428 | Ion Transport |
| 1185 | SLC41A2 | solute carrier family 41, member 2 | 84102 | Ion Transport |
| 1186 | SLC41A3 | solute carrier family 41, member 3 | 54946 | Ion Transport |
| 1187 | SLN | sarcolipin | 6588 | Ion Transport |
| 1188 | TCN1 | transcobalamin I (vitamin B12 binding protein, R binder family) | 6947 | Ion Transport |
| 1189 | TCN2 | transcobalamin II; macrocytic anemia | 6948 | Ion Transport |
| 1190 | TF | transferrin | 7018 | Ion Transport |
| 1191 | TRPA1 | transient receptor potential cation channel, subfamily A, member 1 | 8989 | Ion Transport |
| 1192 | TRPC1 | transient receptor potential cation channel, subfamily C, member 1 | 7220 | Ion Transport |
| 1193 | TRPC3 | transient receptor potential cation channel, subfamily C, member 3 | 7222 | Ion Transport |
| 1194 | TRPC4 | transient receptor potential cation channel, subfamily C, member 4 | 7223 | Ion Transport |
| 1195 | TRPC5 | transient receptor potential cation channel, subfamily C, member 5 | 7224 | Ion Transport |
| 1196 | TRPC6 | transient receptor potential cation channel, subfamily C, member 6 | 7225 | Ion Transport |
| 1197 | TRPC7 | transient receptor potential cation channel, subfamily C, member 7 | 57113 | Ion Transport |
| 1198 | TRPM1 | transient receptor potential cation channel, subfamily M, member 1 | 4308 | Ion Transport |
| 1199 | TRPM2 | transient receptor potential cation channel, subfamily M, member 2 | 7226 | Ion Transport |
| 1200 | TRPM3 | transient receptor potential cation channel, subfamily M, member 3 | 80036 | Ion Transport |
| 1201 | TRPM4 | transient receptor potential cation channel, subfamily M, member 4 | 54795 | Ion Transport |
| 1202 | TRPM5 | transient receptor potential cation channel, subfamily M, member 5 | 29850 | Ion Transport |
| 1203 | TRPM6 | transient receptor potential cation channel, subfamily M, member 6 | 140803 | Ion Transport |
| 1204 | TRPM7 | transient receptor potential cation channel, subfamily M, member 7 | 54822 | Ion Transport |
| 1205 | TRPM8 | transient receptor potential cation channel, subfamily M, member 8 | 79054 | Ion Transport |
| 1206 | TRPV1 | transient receptor potential cation channel, subfamily V, member 1 | 7442 | Ion Transport |
| 1207 | TRPV2 | transient receptor potential cation channel, subfamily V, member 2 | 51393 | Ion Transport |
| 1208 | TRPV3 | transient receptor potential cation channel, subfamily V, member 3 | 162514 | Ion Transport |
| 1209 | TRPV4 | transient receptor potential cation channel, subfamily V, member 4 | 59341 | Ion Transport |
| 1210 | TRPV5 | transient receptor potential cation channel, subfamily V, member 5 | 56302 | Ion Transport |
| 1211 | TRPV6 | transient receptor potential cation channel, subfamily V, member 6 | 55503 | Ion Transport |
| 1212 | TTYH1 | Tweety homolog 1 (Drosophila) | 57348 | Ion Transport |
| 1213 | TTYH2 | Tweety homolog 2 (Drosophila) | 94015 | Ion Transport |
| 1214 | TTYH3 | Tweety homolog 3 (Drosophila) | 80727 | Ion Transport |
| 1215 | VDAC1 | voltage-dependent anion channel 1 | 7416 | Ion Transport |
| 1216 | VDAC2 | voltage-dependent anion channel 2 | 7417 | Ion Transport |
| 1217 | VDAC3 | voltage-dependent anion channel 3 | 7419 | Ion Transport |
| 1218 | ZACN | zinc activated ligand-gated ion channel | 353174 | Ion Transport |
| 1219 | ARSD | arylsulfatase D | 414 | Ion Transport |
| 1220 | ARSK | Arylsulfatase family, member K | 153642 | Ion Transport |
| 1221 | TMCO3 | Transmembrane and coiled-coil domains 3 | 55002 | Ion Transport |
| 1222 | TPCN1 | two pore segment channel 1 | 53373 | Ion Transport |
| 1223 | TPCN2 | Two pore segment channel 2 | 219931 | Ion Transport |
| 1224 | SLC8A1 | solute carrier family 8 (sodium/calcium exchanger), member 1 | 6546 | Ion Transport |
| 1225 | SLC8A2 | solute carrier family 8 (sodium/calcium exchanger), member 2 | 6543 | Ion Transport |
| 1226 | SLC8A3 | solute carrier family 8 (sodium/calcium exchanger), member 3 | 6547 | Ion Transport |
| 1227 | SLC9A1 | solute carrier family 9 (sodium/hydrogen exchanger), member 1 | 6548 | Ion Transport |
| 1228 | SLC9A10 | solute carrier family 9 (sodium/hydrogen exchanger), isoform 10 | 285335 | Ion Transport |
| 1229 | SLC9A11 | solute carrier family 9, member 11 | 284525 | Ion Transport |
| 1230 | SLC9A2 | solute carrier family 9 (sodium/hydrogen exchanger), member 2 | 6549 | Ion Transport |
| 1231 | SLC9A3 | solute carrier family 9 (sodium/hydrogen exchanger), member 3 | 6550 | Ion Transport |
| 1232 | SLC9A4 | solute carrier family 9 (sodium/hydrogen exchanger), member 4 | 389015 | Ion Transport |
| 1233 | SLC9A5 | solute carrier family 9 (sodium/hydrogen exchanger), member 5 | 6553 | Ion Transport |
| 1234 | SLC9A6 | solute carrier family 9 (sodium/hydrogen exchanger), isoform 6 | 10479 | Ion Transport |
| 1235 | SLC9A7 | solute carrier family 9 (sodium/hydrogen exchanger), isoform 7 | 84679 | Ion Transport |
| 1236 | SLC9A8 | solute carrier family 9 (sodium/hydrogen exchanger), member 8 | 23315 | Ion Transport |
| 1237 | SLC9A9 | solute carrier family 9 (sodium/hydrogen exchanger), isoform 9 | 285195 | Ion Transport |
| 1238 | SLC22A23 | solute carrier family 22, member 23 | 63027 | Ion Transport |
| 1239 | MFSD5 | Major Facilitator Superfamily Domain Containing 5 | 84975 | Ion Transport |
| 1240 | MFSD8 | Major Facilitator Superfamily Domain Containing 8 | 256471 | Ion Transport |
| 1241 | ACCN1 | amiloride-sensitive cation channel 1, neuronal (degenerin) | 40 | Ion Transport |
| 1242 | ACCN2 | amiloride-sensitive cation channel 2, neuronal | 41 | Ion Transport |
| 1243 | ACCN3 | amiloride-sensitive cation channel 3 | 9311 | IonTransport |
| 1244 | ACCN4 | amiloride-sensitive cation channel 4, pituitary | 55515 | Ion Transport |
| 1245 | ACCN5 | amiloride-sensitive cation channel 5, intestinal | 51802 | Ion Transport |
| 1246 | AACS | acetoacetyl-CoA synthetase | 65985 | Ketone Bodies |
| 1247 | BDH1 | 3-hydroxybutyrate dehydrogenase, type 1 | 622 | Ketone Bodies |
| 1248 | BDH2 | 3-hydroxybutyrate dehydrogenase, type 2 | 56898 | Ketone Bodies |
| 1249 | OXCT1 | 3-oxoacid CoA transferase 1 | 5019 | Ketone Bodies |
| 1250 | OXCT2 | 3-oxoacid CoA transferase 2 | 64064 | Ketone Bodies |
| 1251 | HMGCS2 | 3-hydroxy-3-methylglutaryl-Coenzyme A synthase 2 (mitochondrial) | 3158 | Ketone Bodies |
| 1252 | HMGCL | 3-hydroxymethyl-3-methylglutaryl-Coenzyme A lyase (hydroxymethylglutaricaciduria) | 3155 | Ketone Bodies |
| 1253 | HMGCLL1 | 3-hydroxymethyl-3-methylglutaryl-Coenzyme A lyase-like 1 | 54511 | Ketone Bodies |
| 1254 | PTER | phosphotriesterase related | 9317 | Lipid |
| 1255 | LIAS | lipoic acid synthetase | 11019 | Lipid |
| 1256 | LIPT1 | lipoyltransferase 1 | 51601 | Lipid |
| 1257 | IAH1 | isoamyl acetate-hydrolyzing esterase 1 homolog (S. cerevisiae) | 285148 | Lipid |
| 1258 | BAAT | bile acid Coenzyme A: amino acid N-acyltransferase (glycine N-choloyltransferase) | 570 | Lipid (Bile Acid) |
| 1259 | CYP27A1 | cytochrome P450, family 27, subfamily A, polypeptide 1 | 1593 | Lipid (Bile Acid) |
| 1260 | CYP7A1 | cytochrome P450, family 7, subfamily A, polypeptide 1 | 1581 | Lipid (Bile Acid) |
| 1261 | CYP7B1 | cytochrome P450, family 7, subfamily B, polypeptide 1 | 9420 | Lipid (Bile Acid) |
| 1262 | CYP8B1 | cytochrome P450, family 8, subfamily B, polypeptide 1 | 1582 | Lipid (Bile Acid) |
| 1263 | HSD3B7 | hydroxy-delta-5-steroid dehydrogenase, 3 beta- and steroid delta-isomerase 7 | 80270 | Lipid (Bile Acid) |
| 1264 | CYP51A1 | cytochrome P450, family 51, subfamily A, polypeptide 1 | 1595 | Lipid (Cholesterol) |
| 1265 | DHCR24 | 24-dehydrocholesterol reductase | 1718 | Lipid (Cholesterol) |
| 1266 | DHCR7 | 7-dehydrocholesterol reductase | 1717 | Lipid (Cholesterol) |
| 1267 | EBP | emopamil binding protein (sterol isomerase) | 10682 | Lipid (Cholesterol) |
| 1268 | EBPL | emopamil binding protein-like | 84650 | Lipid (Cholesterol) |
| 1269 | FDFT1 | farnesyl-diphosphate farnesyltransferase 1 | 2222 | Lipid (Cholesterol) |
| 1270 | FDPS | farnesyl diphosphate synthase (farnesyl pyrophosphate synthetase, dimethylallyltranstransferase, geranyltranstransferase) | 2224 | Lipid (Cholesterol) |
| 1271 | HSD17B7 | hydroxysteroid (17-beta) dehydrogenase 7 | 51478 | Lipid (Cholesterol) |
| 1272 | LBR | Lamin B receptor | 3930 | Lipid (Cholesterol) |
| 1273 | LCAT | lecithin-cholesterol acyltransferase | 3931 | Lipid (Cholesterol) |
| 1274 | LSS | lanosterol synthase (2,3-oxidosqualene-lanosterol cyclase) | 4047 | Lipid (Cholesterol) |
| 1275 | NSDHL | NAD(P) dependent steroid dehydrogenase-like | 50814 | Lipid (Cholesterol) |
| 1276 | SC4MOL | sterol-C4-methyl oxidase-like | 6307 | Lipid (Cholesterol) |
| 1277 | SC5DL | sterol-C5-desaturase (ERG3 delta-5-desaturase homolog, S. cerevisiae)-like | 6309 | Lipid (Cholesterol) |
| 1278 | SQLE | Squalene epoxidase | 6713 | Lipid (Cholesterol) |
| 1279 | TM7SF2 | transmembrane 7 superfamily member 2 | 7108 | Lipid (Cholesterol) |
| 1280 | CEL | carboxyl ester lipase (bile salt-stimulated lipase) | 1056 | Lipid (Cofactor) |
| 1281 | CRAT | carnitine acetyltransferase | 1384 | Lipid (Cofactor) |
| 1282 | CUBN | cubilin (intrinsic factor-cobalamin receptor) | 8029 | Lipid (Cofactor) |
| 1283 | CYP24A1 | cytochrome P450, family 24, subfamily A, polypeptide 1 | 1591 | Lipid (Cofactor) |
| 1284 | CYP27B1 | cytochrome P450, family 27, subfamily B, polypeptide 1 | 1594 | Lipid (Cofactor) |
| 1285 | DAK | dihydroxyacetone kinase 2 homolog (S. cerevisiae) | 26007 | Lipid (Cofactor) |
| 1286 | ILVBL | ilvB (bacterial acetolactate synthase)-like | 10994 | Lipid (Fatty Acid) |
| 1287 | ALDH3A2 | aldehyde dehydrogenase 3 family, member A2 | 224 | Lipid (Fatty Acid) |
| 1288 | ALDH3B1 | aldehyde dehydrogenase 3 family, member B1 | 221 | Lipid (Fatty Acid) |
| 1289 | ALDH3B2 | aldehyde dehydrogenase 3 family, member B2 | 222 | Lipid (Fatty Acid) |
| 1290 | AMACR | alpha-methylacyl-CoA racemase | 23600 | Lipid (Fatty acid) |
| 1291 | ACAA1 | acetyl-Coenzyme A acyltransferase 1 (peroxisomal 3-oxoacyl-Coenzyme A thiolase) | 30 | Lipid (Fatty Acid) |
| 1292 | ACAA2 | acetyl-Coenzyme A acyltransferase 2 (mitochondrial 3-oxoacyl-Coenzyme A thiolase) | 10449 | Lipid (Fatty Acid) |
| 1293 | ACACA | acetyl-Coenzyme A carboxylase alpha | 31 | Lipid (Fatty Acid) |
| 1294 | ACACB | acetyl-Coenzyme A carboxylase beta | 32 | Lipid (Fatty Acid) |
| 1295 | ACAD10 | acyl-Coenzyme A dehydrogenase family, member 10 | 80724 | Lipid (Fatty Acid) |
| 1296 | ACAD11 | acyl-Coenzyme A dehydrogenase family, member 11 | 84129 | Lipid (Fatty Acid) |
| 1297 | ACAD8 | acyl-Coenzyme A dehydrogenase family, member 8 | 27034 | Lipid (Fatty Acid) |
| 1298 | ACAD9 | acyl-Coenzyme A dehydrogenase family, member 9 | 28976 | Lipid (Fatty Acid) |
| 1299 | ACADL | acyl-Coenzyme A dehydrogenase, long chain | 33 | Lipid (Fatty Acid) |
| 1300 | ACADM | acyl-Coenzyme A dehydrogenase, C-4 to C-12 straight chain | 34 | Lipid (Fatty Acid) |
| 1301 | ACADS | acyl-Coenzyme A dehydrogenase, C-2 to C-3 short chain | 35 | Lipid (Fatty Acid) |
| 1302 | ACADSB | acyl-Coenzyme A dehydrogenase, short/branched chain | 36 | Lipid (Fatty Acid) |
| 1303 | ACADVL | acyl-Coenzyme A dehydrogenase, very long chain | 37 | Lipid (Fatty Acid) |
| 1304 | ACAT1 | acetyl-Coenzyme A acetyltransferase 1 (acetoacetyl Coenzyme A thiolase) | 38 | Lipid (Fatty Acid) |
| 1305 | ACAT2 | acetyl-Coenzyme A acetyltransferase 2 (acetoacetyl Coenzyme A thiolase) | 39 | Lipid (Fatty Acid) |
| 1306 | ACBD3 | acyl-Coenzyme A binding domain containing 3 | 64746 | Lipid (Fatty Acid) |
| 1307 | ACBD4 | acyl-Coenzyme A binding domain containing 4 | 79777 | Lipid (Fatty Acid) |
| 1308 | ACBD5 | acyl-Coenzyme A binding domain containing 5 | 91452 | Lipid (Fatty Acid) |
| 1309 | ACBD6 | acyl-Coenzyme A binding domain containing 6 | 84320 | Lipid (Fatty Acid) |
| 1310 | ACBD7 | acyl-Coenzyme A binding domain containing 7 | 414149 | Lipid (Fatty Acid) |
| 1311 | ACOX1 | acyl-Coenzyme A oxidase 1, palmitoyl | 51 | Lipid (Fatty Acid) |
| 1312 | ACOX2 | acyl-Coenzyme A oxidase 2, branched chain | 8309 | Lipid (Fatty Acid) |
| 1313 | ACOX3 | acyl-Coenzyme A oxidase 3, pristanoyl | 8310 | Lipid (Fatty Acid) |
| 1314 | ACOXL | acyl-Coenzyme A oxidase-like | 55289 | Lipid (Fatty Acid) |
| 1315 | ACSBG1 | acyl-CoA synthetase bubblegum family member 1 | 23205 | Lipid (Fatty Acid) |
| 1316 | ACSBG2 | acyl-CoA synthetase bubblegum family member 2 | 81616 | Lipid (Fatty Acid) |
| 1317 | ACSF2 | acyl-CoA synthetase family member 2 | 80221 | Lipid (Fatty Acid) |
| 1318 | ACSF3 | acyl-CoA synthetase family member 3 | 197322 | Lipid (Fatty Acid) |
| 1319 | ACSL1 | acyl-CoA synthetase long-chain family member 1 | 2180 | Lipid (Fatty Acid) |
| 1320 | ACSL3 | acyl-CoA synthetase long-chain family member 3 | 2181 | Lipid (Fatty Acid) |
| 1321 | ACSL4 | Acyl-CoA synthetase long-chain family member 4 | 2182 | Lipid (Fatty Acid) |
| 1322 | ACSL5 | acyl-CoA synthetase long-chain family member 5 | 51703 | Lipid (Fatty Acid) |
| 1323 | ACSL6 | acyl-CoA synthetase long-chain family member 6 | 23305 | Lipid (Fatty Acid) |
| 1324 | ACSM1 | acyl-CoA synthetase medium-chain family member 1 | 116285 | Lipid (Fatty Acid) |
| 1325 | ACSM2A | acyl-CoA synthetase medium-chain family member 2A | 123876 | Lipid (Fatty Acid) |
| 1326 | ACSM2B | acyl-CoA synthetase medium-chain family member 2B | 348158 | Lipid (Fatty Acid) |
| 1327 | ACSM3 | acyl-CoA synthetase medium-chain family member 3 | 6296 | Lipid (Fatty Acid) |
| 1328 | ACSM4 | acyl-CoA synthetase medium-chain family member 4 | 341392 | Lipid (Fatty Acid) |
| 1329 | ACSM5 | acyl-CoA synthetase medium-chain family member 5 | 54988 | Lipid (Fatty Acid) |
| 1330 | ACSS1 | Acyl-CoA synthetase short-chain family member 1 | 84532 | Lipid (Fatty Acid) |
| 1331 | ACSS2 | acyl-CoA synthetase short-chain family member 2 | 55902 | Lipid (Fatty Acid) |
| 1332 | ACSS3 | acyl-CoA synthetase short-chain family member 3 | 79611 | Lipid (Fatty Acid) |
| 1333 | CLYBL | citrate lyase beta like | 171425 | Lipid (Fatty Acid) |
| 1334 | CPT1A | carnitine palmitoyltransferase 1A (liver) | 1374 | Lipid (Fatty Acid) |
| 1335 | CPT1B | carnitine palmitoyltransferase 1B (muscle) | 1375 | Lipid (Fatty Acid) |
| 1336 | CPT1C | carnitine palmitoyltransferase 1C | 126129 | Lipid (Fatty Acid) |
| 1337 | CPT2 | carnitine palmitoyltransferase II | 1376 | Lipid (Fatty Acid) |
| 1338 | CYP4A11 | cytochrome P450, family 4, subfamily A, polypeptide 11 | 1579 | Lipid (Fatty Acid) |
| 1339 | CYP4A22 | cytochrome P450, family 4, subfamily A, polypeptide 22 | 284541 | Lipid (Fatty Acid) |
| 1340 | CYP4B1 | cytochrome P450, family 4, subfamily B, polypeptide 1 | 1580 | Lipid (Fatty Acid) |
| 1341 | CYP4F11 | cytochrome P450, family 4, subfamily F, polypeptide 11 | 57834 | Lipid (Fatty Acid) |
| 1342 | CYP4F12 | cytochrome P450, family 4, subfamily F, polypeptide 12 | 66002 | Lipid (Fatty Acid) |
| 1343 | CYP4F2 | cytochrome P450, family 4, subfamily F, polypeptide 2 | 8529 | Lipid (Fatty Acid) |
| 1344 | CYP4F22 | cytochrome P450, family 4, subfamily F, polypeptide 22 | 126410 | Lipid (Fatty Acid) |
| 1345 | CYP4F3 | cytochrome P450, family 4, subfamily F, polypeptide 3 | 4051 | Lipid (Fatty Acid) |
| 1346 | CYP4F8 | cytochrome P450, family 4, subfamily F, polypeptide 8 | 11283 | Lipid (Fatty Acid) |
| 1347 | CYP4V2 | cytochrome P450, family 4, subfamily V, polypeptide 2 | 285440 | Lipid (Fatty Acid) |
| 1348 | CYP4X1 | cytochrome P450, family 4, subfamily X, polypeptide 1 | 260293 | Lipid (Fatty Acid) |
| 1349 | CYP4Z1 | cytochrome P450, family 4, subfamily Z, polypeptide 1 | 199974 | Lipid (Fatty Acid) |
| 1350 | DCI | dodecenoyl-Coenzyme A delta isomerase (3,2 trans-enoyl-Coenzyme A isomerase) | 1632 | Lipid (Fatty Acid) |
| 1351 | DECR1 | 2,4-dienoyl CoA reductase 1, mitochondrial | 1666 | Lipid (Fatty Acid) |
| 1352 | DECR2 | 2,4-dienoyl CoA reductase 2, peroxisomal | 26063 | Lipid (Fatty Acid) |
| 1353 | DGAT1 | diacylglycerol O-acyltransferase homolog 1 (mouse) | 8694 | Lipid (Fatty Acid) |
| 1354 | DGAT2 | diacylglycerol O-acyltransferase homolog 2 (mouse) | 84649 | Lipid (Fatty Acid) |
| 1355 | DGAT2L3 | diacylglycerol O-acyltransferase 2-like 3 | 158833 | Lipid (Fatty Acid) |
| 1356 | DGAT2L4 | diacylglycerol O-acyltransferase 2-like 4 | 158835 | Lipid (Fatty Acid) |
| 1357 | DGAT2L6 | diacylglycerol O-acyltransferase 2-like 6 | 347516 | Lipid (Fatty Acid) |
| 1358 | DGAT2L7 | diacylglycerol O-acyltransferase 2-like 7 | 646409 | Lipid (Fatty Acid) |
| 1359 | ECH1 | enoyl Coenzyme A hydratase 1, peroxisomal | 1891 | Lipid (Fatty Acid) |
| 1360 | ECHDC1 | enoyl Coenzyme A hydratase domain containing 1 | 55862 | Lipid (Fatty Acid) |
| 1361 | ECHDC2 | enoyl Coenzyme A hydratase domain containing 2 | 55268 | Lipid (Fatty Acid) |
| 1362 | ECHDC3 | enoyl Coenzyme A hydratase domain containing 3 | 79746 | Lipid (Fatty Acid) |
| 1363 | ECHS1 | enoyl Coenzyme A hydratase, short chain, 1, mitochondrial | 1892 | Lipid (Fatty Acid) |
| 1364 | EHHADH | enoyl-Coenzyme A, hydratase/3-hydroxyacyl Coenzyme A dehydrogenase | 1962 | Lipid (Fatty Acid) |
| 1365 | ELOVL1 | elongation of very long chain fatty acids (FEN1/Elo2, SUR4/Elo3, yeast)-like 1 | 64834 | Lipid (Fatty Acid) |
| 1366 | ELOVL2 | elongation of very long chain fatty acids (FEN1/Elo2, SUR4/Elo3, yeast)-like 2 | 54898 | Lipid (Fatty Acid) |
| 1367 | ELOVL3 | elongation of very long chain fatty acids (FEN1/Elo2, SUR4/Elo3, yeast)-like 3 | 83401 | Lipid (Fatty Acid) |
| 1368 | ELOVL4 | elongation of very long chain fatty acids (FEN1/Elo2, SUR4/Elo3, yeast)-like 4 | 6785 | Lipid (Fatty Acid) |
| 1369 | ELOVL5 | ELOVL family member 5, elongation of long chain fatty acids (FEN1/Elo2, SUR4/Elo3-like, yeast) | 60481 | Lipid (Fatty Acid) |
| 1370 | ELOVL6 | ELOVL family member 6, elongation of long chain fatty acids (FEN1/Elo2, SUR4/Elo3-like, yeast) | 79071 | Lipid (Fatty Acid) |
| 1371 | ELOVL7 | ELOVL family member 7, elongation of long chain fatty acids (yeast) | 79993 | Lipid (Fatty Acid) |
| 1372 | ETFA | electron-transfer-flavoprotein, alpha polypeptide (glutaric aciduria II) | 2108 | Lipid (Fatty Acid) |
| 1373 | ETFB | electron-transfer-flavoprotein, beta polypeptide | 2109 | Lipid (Fatty Acid) |
| 1374 | ETFDH | electron-transferring-flavoprotein dehydrogenase | 2110 | Lipid (Fatty Acid) |
| 1375 | FA2H | fatty acid 2-hydroxylase | 79152 | Lipid (Fatty Acid) |
| 1376 | FAAH | fatty acid amide hydrolase | 2166 | Lipid (Fatty Acid) |
| 1377 | FAAH2 | fatty acid amide hydrolase 2 | 158584 | Lipid (Fatty Acid) |
| 1378 | FADS1 | fatty acid desaturase 1 | 3992 | Lipid (Fatty Acid) |
| 1379 | FADS2 | fatty acid desaturase 2 | 9415 | Lipid (Fatty Acid) |
| 1380 | FADS3 | fatty acid desaturase 3 | 3995 | Lipid (Fatty Acid) |
| 1381 | FAR1 | fatty acyl CoA reductase 1 | 84188 | Lipid (Fatty Acid) |
| 1382 | FAR2 | fatty acyl CoA reductase 2 | 55711 | Lipid (Fatty Acid) |
| 1383 | FASN | fatty acid synthase | 2194 | Lipid (Fatty Acid) |
| 1384 | GPSN2 | glycoprotein, synaptic 2 | 9524 | Lipid (Fatty Acid) |
| 1385 | HACL1 | 2-hydroxyacyl-CoA lyase 1 | 26061 | Lipid (Fatty Acid) |
| 1386 | HADH | hydroxyacyl-Coenzyme A dehydrogenase | 3033 | Lipid (Fatty Acid) |
| 1387 | HADHA | hydroxyacyl-Coenzyme A dehydrogenase/3-ketoacyl-Coenzyme A thiolase/enoyl-Coenzyme A hydratase (trifunctional protein), alpha subunit | 3030 | Lipid (Fatty Acid) |
| 1388 | HADHB | hydroxyacyl-Coenzyme A dehydrogenase/3-ketoacyl-Coenzyme A thiolase/enoyl-Coenzyme A hydratase (trifunctional protein), beta subunit | 3032 | Lipid (Fatty Acid) |
| 1389 | LIPC | lipase, hepatic | 3990 | Lipid (Fatty Acid) |
| 1390 | LIPF | lipase, gastric | 8513 | Lipid (Fatty Acid) |
| 1391 | LIPG | lipase, endothelial | 9388 | Lipid (Fatty Acid) |
| 1392 | LIPH | lipase, member H | 200879 | Lipid (Fatty Acid) |
| 1393 | LIPI | lipase, member I | 149998 | Lipid (Fatty Acid) |
| 1394 | LIPJ | lipase, family member J | 142910 | Lipid (Fatty Acid) |
| 1395 | LIPK | lipase, mamber K | 643414 | Lipid (Fatty Acid) |
| 1396 | LIPM | lipase, member M | 340654 | Lipid (Fatty Acid) |
| 1397 | LIPN | lipase, member N | 643418 | Lipid (Fatty Acid) |
| 1398 | LPL | lipoprotein lipase | 4023 | Lipid (Fatty Acid) |
| 1399 | LPPR2 | lipid phosphate phosphatase-related protein type 2 | 64748 | Lipid (Fatty Acid) |
| 1400 | MCAT | malonyl CoA:ACP acyltransferase (mitochondrial) | 27349 | Lipid (Fatty Acid) |
| 1401 | MECR | mitochondrial trans-2-enoyl-CoA reductase | 51102 | Lipid (Fatty Acid) |
| 1402 | MGLL | monoglyceride lipase | 11343 | Lipid (Fatty Acid) |
| 1403 | MLYCD | malonyl-CoA decarboxylase | 23417 | Lipid (Fatty Acid) |
| 1404 | MTTP | microsomal triglyceride transfer protein | 4547 | Lipid (Fatty Acid) |
| 1405 | OLAH | oleoyl-ACP hydrolase | 55301 | Lipid (Fatty Acid) |
| 1406 | OXSM | 3-oxoacyl-ACP synthase, mitochondrial | 54995 | Lipid (Fatty Acid) |
| 1407 | PECI | peroxisomal D3,D2-enoyl-CoA isomerase | 10455 | Lipid (Fatty Acid) |
| 1408 | PECR | peroxisomal trans-2-enoyl-CoA reductase | 55825 | Lipid (Fatty Acid) |
| 1409 | PHYH | phytanoyl-CoA 2-hydroxylase | 5264 | Lipid (Fatty Acid) |
| 1410 | PHYHD1 | phytanoyl-CoA dioxygenase domain containing 1 | 254295 | Lipid (Fatty Acid) |
| 1411 | PNLIP | pancreatic lipase | 5406 | Lipid (Fatty Acid) |
| 1412 | PNLIPRP1 | pancreatic lipase-related protein 1 | 5407 | Lipid (Fatty Acid) |
| 1413 | PNLIPRP2 | pancreatic lipase-related protein 2 | 5408 | Lipid (Fatty Acid) |
| 1414 | PNLIPRP3 | pancreatic lipase-related protein 3 | 119548 | Lipid (Fatty Acid) |
| 1415 | PNPLA1 | patatin-like phospholipase domain containing 1 | 285848 | Lipid (Fatty Acid) |
| 1416 | PNPLA2 | patatin-like phospholipase domain containing 2 | 57104 | Lipid (Fatty Acid) |
| 1417 | PNPLA3 | patatin-like phospholipase domain containing 3 | 80339 | Lipid (Fatty Acid) |
| 1418 | PNPLA4 | patatin-like phospholipase domain containing 4 | 8228 | Lipid (Fatty Acid) |
| 1419 | PNPLA5 | patatin-like phospholipase domain containing 5 | 150379 | Lipid (Fatty Acid) |
| 1420 | PNPLA6 | patatin-like phospholipase domain containing 6 | 10908 | Lipid (Fatty Acid) |
| 1421 | PNPLA7 | patatin-like phospholipase domain containing 7 | 375775 | Lipid (Fatty Acid) |
| 1422 | PNPLA8 | patatin-like phospholipase domain containing 8 | 50640 | Lipid (Fatty Acid) |
| 1423 | SCD | stearoyl-CoA desaturase (delta-9-desaturase) | 6319 | Lipid (Fatty Acid) |
| 1424 | SCD5 | stearoyl-CoA desaturase 5 | 79966 | Lipid (Fatty Acid) |
| 1425 | SCP2 | sterol carrier protein 2 | 6342 | Lipid (Fatty Acid) |
| 1426 | SDR16C6 | short chain dehydrogenase/reductase family 16C, member 6 | 442388 | Lipid (Fatty Acid) |
| 1427 | SDR39U1 | short chain dehydrogenase/reductase family 39U, member 1 | 56948 | Lipid (Fatty Acid) |
| 1428 | SDR42E1 | short chain dehydrogenase/reductase family 42E, member 1 | 93517 | Lipid (Fatty Acid) |
| 1429 | SDR9C7 | short chain dehydrogenase/reductase family 9C, member 7 | 121214 | Lipid (Fatty Acid) |
| 1430 | CBR4 | carbonyl reductase 4 | 84869 | Lipid (Fatty Acid) |
| 1431 | ABHD5 | Abhydrolase domain containing 5 | 51099 | Lipid (Fatty Acid) |
| 1432 | ABHD1 | abhydrolase domain containing 1 | 84696 | Lipid (Fatty Acid) |
| 1433 | ABHD11 | abhydrolase domain containing 11 | 83451 | Lipid (Fatty Acid) |
| 1434 | ABHD12 | abhydrolase domain containing 12 | 26090 | Lipid (Fatty Acid) |
| 1435 | ABHD12B | abhydrolase domain containing 12B | 145447 | Lipid (Fatty Acid) |
| 1436 | ABHD13 | abhydrolase domain containing 13 | 84945 | Lipid (Fatty Acid) |
| 1437 | ABHD14A | abhydrolase domain containing 14A | 25864 | Lipid (Fatty Acid) |
| 1438 | ABHD14B | abhydrolase domain containing 14B | 84836 | Lipid (Fatty Acid) |
| 1439 | ABHD2 | abhydrolase domain containing 2 | 11057 | Lipid (Fatty Acid) |
| 1440 | ABHD3 | abhydrolase domain containing 3 | 171586 | Lipid (Fatty Acid) |
| 1441 | ABHD4 | abhydrolase domain containing 4 | 63874 | Lipid (Fatty Acid) |
| 1442 | ABHD6 | abhydrolase domain containing 6 | 57406 | Lipid (Fatty Acid) |
| 1443 | ABHD7 | abhydrolase domain containing 7 | 253152 | Lipid (Fatty Acid) |
| 1444 | ABHD8 | abhydrolase domain containing 8 | 79575 | Lipid (Fatty Acid) |
| 1445 | ABHD9 | abhydrolase domain containing 9 | 79852 | Lipid (Fatty Acid) |
| 1446 | AOAH | acyloxyacyl hydrolase (neutrophil) | 313 | Lipid (Fatty Acid) |
| 1447 | THEM4 | thioesterase superfamily member 4 | 117145 | Lipid (Fatty Acid) |
| 1448 | DDHD2 | DDHD domain containing 2 | 23259 | Lipid (Fatty Acid) |
| 1449 | THEM2 | thioesterase superfamily member 2 | 55856 | Lipid (Fatty Acid) |
| 1450 | THEM5 | thioesterase superfamily member 5 | 284486 | Lipid (Fatty Acid) |
| 1451 | AANAT | arylalkylamine N-acetyltransferase | 15 | Lipid (Hormone) |
| 1452 | ACE | angiotensin I converting enzyme (peptidyl-dipeptidase A) 1 | 1636 | Lipid (Hormone) |
| 1453 | ACE2 | angiotensin I converting enzyme (peptidyl-dipeptidase A) 2 | 59272 | Lipid (Hormone) |
| 1454 | ALOX12 | arachidonate 12-lipoxygenase | 239 | Lipid (Hormone) |
| 1455 | ALOX12B | arachidonate 12-lipoxygenase, 12R type | 242 | Lipid (Hormone) |
| 1456 | ALOX15 | arachidonate 15-lipoxygenase | 246 | Lipid (Hormone) |
| 1457 | ALOX15B | arachidonate 15-lipoxygenase, type B | 247 | Lipid (Hormone) |
| 1458 | ALOX5 | arachidonate 5-lipoxygenase | 240 | Lipid (Hormone) |
| 1459 | ALOX5AP | arachidonate 5-lipoxygenase-activating protein | 241 | Lipid (Hormone) |
| 1460 | ALOXE3 | arachidonate lipoxygenase 3 | 59344 | Lipid (Hormone) |
| 1461 | AOF1 | amine oxidase (flavin containing) domain 1 | 221656 | Lipid (Hormone) |
| 1462 | AOF2 | amine oxidase (flavin containing) domain 2 | 23028 | Lipid (Hormone) |
| 1463 | ASMT | acetylserotonin O-methyltransferase | 438 | Lipid (Hormone) |
| 1464 | ASMTL | acetylserotonin O-methyltransferase-like | 8623 | Lipid (Hormone) |
| 1465 | CYP20A1 | cytochrome P450, family 20, subfamily A, polypeptide 1 | 57404 | Lipid (Hormone) |
| 1466 | DIO1 | deiodinase, iodothyronine, type I | 1733 | Lipid (Hormone) |
| 1467 | DIO2 | deiodinase, iodothyronine, type II | 1734 | Lipid (Hormone) |
| 1468 | DIO3 | deiodinase, iodothyronine, type III | 1735 | Lipid (Hormone) |
| 1469 | HPGD | hydroxyprostaglandin dehydrogenase 15-(NAD) | 3248 | Lipid (Hormone) |
| 1470 | IYD | iodotyrosine deiodinase | 389434 | Lipid (Hormone) |
| 1471 | LNPEP | leucyl/cystinyl aminopeptidase | 4012 | Lipid (Hormone) |
| 1472 | LTA4H | leukotriene A4 hydrolase | 4048 | Lipid (Hormone) |
| 1473 | LTC4S | leukotriene C4 synthase | 4056 | Lipid (Hormone) |
| 1474 | MME | membrane metallo-endopeptidase | 4311 | Lipid (Hormone) |
| 1475 | MMEL1 | membrane metallo-endopeptidase-like 1 | 79258 | Lipid (Hormone) |
| 1476 | PGDS | prostaglandin D2 synthase, hematopoietic | 27306 | Lipid (Hormone) |
| 1477 | PTGDS | prostaglandin D2 synthase 21kDa (brain) | 5730 | Lipid (Hormone) |
| 1478 | PTGES | prostaglandin E synthase | 9536 | Lipid (Hormone) |
| 1479 | PTGES2 | prostaglandin E synthase 2 | 80142 | Lipid (Hormone) |
| 1480 | PTGES3 | prostaglandin E synthase 3 (cytosolic) | 10728 | Lipid (Hormone) |
| 1481 | PTGIS | prostaglandin I2 (prostacyclin) synthase | 5740 | Lipid (Hormone) |
| 1482 | PTGR1 | prostaglandin reductase 1 | 22949 | Lipid (Hormone) |
| 1483 | PTGR2 | prostaglandin reductase 2 | 145482 | Lipid (Hormone) |
| 1484 | PTGS1 | prostaglandin-endoperoxide synthase 1 (prostaglandin G/H synthase and cyclooxygenase) | 5742 | Lipid (Hormone) |
| 1485 | PTGS2 | prostaglandin-endoperoxide synthase 2 (prostaglandin G/H synthase and cyclooxygenase) | 5743 | Lipid (Hormone) |
| 1486 | TBXAS1 | thromboxane A synthase 1 (platelet, cytochrome P450, family 5, subfamily A) | 6916 | Lipid (Hormone) |
| 1487 | TPO | thyroid peroxidase | 7173 | Lipid (Hormone) |
| 1488 | SCCPDH | saccharopine dehydrogenase (putative) | 51097 | Lipid (Membrane lipd) |
| 1489 | AGPAT1 | 1-acylglycerol-3-phosphate O-acyltransferase 1 (lysophosphatidic acid acyltransferase, alpha) | 10554 | Lipid (Membrane Lipid) |
| 1490 | AGPAT2 | 1-acylglycerol-3-phosphate O-acyltransferase 2 (lysophosphatidic acid acyltransferase, beta) | 10555 | Lipid (Membrane Lipid) |
| 1491 | AGPAT3 | 1-acylglycerol-3-phosphate O-acyltransferase 3 | 56894 | Lipid (Membrane Lipid) |
| 1492 | AGPAT4 | 1-acylglycerol-3-phosphate O-acyltransferase 4 (lysophosphatidic acid acyltransferase, delta) | 56895 | Lipid (Membrane Lipid) |
| 1493 | AGPAT5 | 1-acylglycerol-3-phosphate O-acyltransferase 5 (lysophosphatidic acid acyltransferase, epsilon) | 55326 | Lipid (Membrane Lipid) |
| 1494 | AGPAT6 | 1-acylglycerol-3-phosphate O-acyltransferase 6 (lysophosphatidic acid acyltransferase, zeta) | 137964 | Lipid (Membrane Lipid) |
| 1495 | AGPAT9 | 1-acylglycerol-3-phosphate O-acyltransferase 9 | 84803 | Lipid (Membrane Lipid) |
| 1496 | CDS1 | CDP-diacylglycerol synthase (phosphatidate cytidylyltransferase) 1 | 1040 | Lipid (Membrane Lipid) |
| 1497 | CDS2 | CDP-diacylglycerol synthase (phosphatidate cytidylyltransferase) 2 | 8760 | Lipid (Membrane Lipid) |
| 1498 | CEPT1 | choline/ethanolamine phosphotransferase 1 | 10390 | Lipid (Membrane Lipid) |
| 1499 | CHKA | choline kinase alpha | 1119 | Lipid (Membrane Lipid) |
| 1500 | CHKB | choline kinase beta | 1120 | Lipid (Membrane Lipid) |
| 1501 | CHPT1 | choline phosphotransferase 1 | 56994 | Lipid (Membrane Lipid) |
| 1502 | CLC | Charcot-Leyden crystal protein | 1178 | Lipid (Membrane Lipid) |
| 1503 | CRLS1 | cardiolipin synthase 1 | 54675 | Lipid (Membrane Lipid) |
| 1504 | CROT | carnitine O-octanoyltransferase | 54677 | Lipid (Membrane Lipid) |
| 1505 | DAGLA | diacylglycerol lipase, alpha | 747 | Lipid (Membrane Lipid) |
| 1506 | DAGLB | diacylglycerol lipase, beta | 221955 | Lipid (Membrane Lipid) |
| 1507 | DGKA | diacylglycerol kinase, alpha 80kDa | 1606 | Lipid (Membrane Lipid) |
| 1508 | DGKB | diacylglycerol kinase, beta 90kDa | 1607 | Lipid (Membrane Lipid) |
| 1509 | DGKD | diacylglycerol kinase, delta 130kDa | 8527 | Lipid (Membrane Lipid) |
| 1510 | DGKE | diacylglycerol kinase, epsilon 64kDa | 8526 | Lipid (Membrane Lipid) |
| 1511 | DGKG | diacylglycerol kinase, gamma 90kDa | 1608 | Lipid (Membrane Lipid) |
| 1512 | DGKH | diacylglycerol kinase, eta | 160851 | Lipid (Membrane Lipid) |
| 1513 | DGKI | diacylglycerol kinase, iota | 9162 | Lipid (Membrane Lipid) |
| 1514 | DGKK | diacylglycerol kinase, kappa | 139189 | Lipid (Membrane Lipid) |
| 1515 | DGKQ | diacylglycerol kinase, theta 110kDa | 1609 | Lipid (Membrane Lipid) |
| 1516 | DGKZ | diacylglycerol kinase, zeta 104kDa | 8525 | Lipid (Membrane Lipid) |
| 1517 | ETNK1 | ethanolamine kinase 1 | 55500 | Lipid (Membrane Lipid) |
| 1518 | ETNK2 | ethanolamine kinase 2 | 55224 | Lipid (Membrane Lipid) |
| 1519 | GDE1 | glycerophosphodiester phosphodiesterase 1 | 51573 | Lipid (Membrane Lipid) |
| 1520 | GK | glycerol kinase | 2710 | Lipid (Membrane Lipid) |
| 1521 | GK2 | glycerol kinase 2 | 2712 | Lipid (Membrane Lipid) |
| 1522 | GK5 | glycerol kinase 5 (putative) | 256356 | Lipid (Membrane Lipid) |
| 1523 | GNPAT | glyceronephosphate O-acyltransferase | 8443 | Lipid (Membrane Lipid) |
| 1524 | GPAM | glycerol-3-phosphate acyltransferase, mitochondrial | 57678 | Lipid (Membrane Lipid) |
| 1525 | GPD1 | glycerol-3-phosphate dehydrogenase 1 (soluble) | 2819 | Lipid (Membrane Lipid) |
| 1526 | GPD1L | glycerol-3-phosphate dehydrogenase 1-like | 23171 | Lipid (Membrane Lipid) |
| 1527 | GPD2 | glycerol-3-phosphate dehydrogenase 2 (mitochondrial) | 2820 | Lipid (Membrane Lipid) |
| 1528 | LPCAT1 | lysophosphatidylcholine acyltransferase 1 | 79888 | Lipid (Membrane Lipid) |
| 1529 | LPCAT2 | lysophosphatidylcholine acyltransferase 2 | 54947 | Lipid (Membrane Lipid) |
| 1530 | LPCAT3 | membrane bound O-acyltransferase domain containing 5 | 10162 | Lipid (Membrane Lipid) |
| 1531 | LPCAT4 | lysophosphatidylcholine acyltransferase 4 | 254531 | Lipid (Membrane Lipid) |
| 1532 | LPGAT1 | lysophosphatidylglycerol acyltransferase 1 | 9926 | Lipid (Membrane Lipid) |
| 1533 | LPIN1 | lipin 1 | 23175 | Lipid (Membrane Lipid) |
| 1534 | LPIN2 | lipin 2 | 9663 | Lipid (Membrane Lipid) |
| 1535 | LPIN3 | lipin 3 | 64900 | Lipid (Membrane Lipid) |
| 1536 | LYPLA1 | lysophospholipase I | 10434 | Lipid (Membrane Lipid) |
| 1537 | LYPLA2 | lysophospholipase II | 11313 | Lipid (Membrane Lipid) |
| 1538 | LYPLAL1 | lysophospholipase-like 1 | 127018 | Lipid (Membrane Lipid) |
| 1539 | MBOAT1 | membrane bound O-acyltransferase domain containing 1 | 154141 | Lipid (Membrane Lipid) |
| 1540 | MBOAT2 | membrane bound O-acyltransferase domain containing 2 | 129642 | Lipid (Membrane Lipid) |
| 1541 | MBOAT4 | membrane bound O-acyltransferase domain containing 4 | 619373 | Lipid (Membrane Lipid) |
| 1542 | MBOAT7 | membrane bound O-acyltransferase domain containing 7 | 79143 | Lipid (Membrane Lipid) |
| 1543 | MOGAT1 | monoacylglycerol O-acyltransferase 1 | 116255 | Lipid (Membrane Lipid) |
| 1544 | MOGAT2 | monoacylglycerol O-acyltransferase 2 | 80168 | Lipid (Membrane Lipid) |
| 1545 | MOGAT3 | monoacylglycerol O-acyltransferase 3 | 346606 | Lipid (Membrane Lipid) |
| 1546 | NAPE-PLD | N-acyl-phosphatidylethanolamine-hydrolyzing phospholipase D | 222236 | Lipid (Membrane Lipid) |
| 1547 | PAP2D | phosphatidic acid phosphatase type 2 | 163404 | Lipid (Membrane Lipid) |
| 1548 | PCYT1A | phosphate cytidylyltransferase 1, choline, alpha | 5130 | Lipid (Membrane Lipid) |
| 1549 | PCYT1B | phosphate cytidylyltransferase 1, choline, beta | 9468 | Lipid (Membrane Lipid) |
| 1550 | PCYT2 | phosphate cytidylyltransferase 2, ethanolamine | 5833 | Lipid (Membrane Lipid) |
| 1551 | PEMT | phosphatidylethanolamine N-methyltransferase | 10400 | Lipid (Membrane Lipid) |
| 1552 | PGS1 | phosphatidylglycerophosphate synthase 1 | 9489 | Lipid (Membrane Lipid) |
| 1553 | PHOSPHO1 | phosphatase orphan 1 | 162466 | Lipid (Membrane Lipid) |
| 1554 | PISD | phosphatidylserine decarboxylase | 23761 | Lipid (Membrane Lipid) |
| 1555 | PLA1A | phospholipase A1 member A | 51365 | Lipid (Membrane Lipid) |
| 1556 | PLA2G10 | phospholipase A2, group X | 8399 | Lipid (Membrane Lipid) |
| 1557 | PLA2G12A | phospholipase A2, group XIIA | 81579 | Lipid (Membrane Lipid) |
| 1558 | PLA2G12B | phospholipase A2, group XIIB | 84647 | Lipid (Membrane Lipid) |
| 1559 | PLA2G15 | phospholipase A2, group XV | 23659 | Lipid (Membrane Lipid) |
| 1560 | PLA2G16 | phospholipase A2, group XVI | 11145 | Lipid (Membrane Lipid) |
| 1561 | PLA2G1B | phospholipase A2, group IB (pancreas) | 5319 | Lipid (Membrane Lipid) |
| 1562 | PLA2G2A | phospholipase A2, group IIA (platelets, synovial fluid) | 5320 | Lipid (Membrane Lipid) |
| 1563 | PLA2G2C | phospholipase A2, group IIC | 391013 | Lipid (Membrane Lipid) |
| 1564 | PLA2G2D | phospholipase A2, group IID | 26279 | Lipid (Membrane Lipid) |
| 1565 | PLA2G2E | phospholipase A2, group IIE | 30814 | Lipid (Membrane Lipid) |
| 1566 | PLA2G2F | phospholipase A2, group IIF | 64600 | Lipid (Membrane Lipid) |
| 1567 | PLA2G3 | phospholipase A2, group III | 50487 | Lipid (Membrane Lipid) |
| 1568 | PLA2G4A | phospholipase A2, group IVA (cytosolic, calcium-dependent) | 5321 | Lipid (Membrane Lipid) |
| 1569 | PLA2G4B | phospholipase A2, group IVB (cytosolic) | ######## | Lipid (Membrane Lipid) |
| 1570 | PLA2G4C | phospholipase A2, group IVC (cytosolic, calcium-independent) | 8605 | Lipid (Membrane Lipid) |
| 1571 | PLA2G4D | phospholipase A2, group IVD (cytosolic) | 283748 | Lipid (Membrane Lipid) |
| 1572 | PLA2G4E | phospholipase A2, group IVE | 123745 | Lipid (Membrane Lipid) |
| 1573 | PLA2G4F | Phospholipase A2, group IVF | 255189 | Lipid (Membrane Lipid) |
| 1574 | PLA2G5 | phospholipase A2, group V | 5322 | Lipid (Membrane Lipid) |
| 1575 | PLA2G6 | phospholipase A2, group VI (cytosolic, calcium-independent) | 8398 | Lipid (Membrane Lipid) |
| 1576 | PLA2G7 | phospholipase A2, group VII (platelet-activating factor acetylhydrolase, plasma) | 7941 | Lipid (Membrane Lipid) |
| 1577 | PLB1 | phospholipase B1 | 151056 | Lipid (Membrane Lipid) |
| 1578 | PLCB1 | phospholipase C, beta 1 (phosphoinositide-specific) | 23236 | Lipid (Membrane Lipid) |
| 1579 | PLCB2 | phospholipase C, beta 2 | 5330 | Lipid (Membrane Lipid) |
| 1580 | PLCB3 | phospholipase C, beta 3 (phosphatidylinositol-specific) | 5331 | Lipid (Membrane Lipid) |
| 1581 | PLCB4 | phospholipase C, beta 4 | 5332 | Lipid (Membrane Lipid) |
| 1582 | PLCD1 | phospholipase C, delta 1 | 5333 | Lipid (Membrane Lipid) |
| 1583 | PLCD3 | phospholipase C, delta 3 | 113026 | Lipid (Membrane Lipid) |
| 1584 | PLCD4 | phospholipase C, delta 4 | 84812 | Lipid (Membrane Lipid) |
| 1585 | PLCE1 | Phospholipase C, epsilon 1 | 51196 | Lipid (Membrane Lipid) |
| 1586 | PLCG1 | phospholipase C, gamma 1 | 5335 | Lipid (Membrane Lipid) |
| 1587 | PLCG2 | Phospholipase C, gamma 2 (phosphatidylinositol-specific) | 5336 | Lipid (Membrane Lipid) |
| 1588 | PLCH1 | phospholipase C, eta 1 | 23007 | Lipid (Membrane Lipid) |
| 1589 | PLCH2 | phospholipase C, eta 2 | 9651 | Lipid (Membrane Lipid) |
| 1590 | PLCL1 | phospholipase C-like 1 | 5334 | Lipid (Membrane Lipid) |
| 1591 | PLCL2 | phospholipase C-like 2 | 23228 | Lipid (Membrane Lipid) |
| 1592 | PLCXD1 | phosphatidylinositol-specific phospholipase C, X domain containing 1 | 55344 | Lipid (Membrane Lipid) |
| 1593 | PLCXD2 | phosphatidylinositol-specific phospholipase C, X domain containing 2 | 257068 | Lipid (Membrane Lipid) |
| 1594 | PLCXD3 | phosphatidylinositol-specific phospholipase C, X domain containing 3 | 345557 | Lipid (Membrane Lipid) |
| 1595 | PLCZ1 | phospholipase C, zeta 1 | 89869 | Lipid (Membrane Lipid) |
| 1596 | PLD1 | phospholipase D1, phosphatidylcholine-specific | 5337 | Lipid (Membrane Lipid) |
| 1597 | PLD2 | phospholipase D2 | 5338 | Lipid (Membrane Lipid) |
| 1598 | PLD3 | phospholipase D family, member 3 | 23646 | Lipid (Membrane Lipid) |
| 1599 | PLD4 | phospholipase D family, member 4 | 122618 | Lipid (Membrane Lipid) |
| 1600 | PLD5 | phospholipase D family, member 5 | 200150 | Lipid (Membrane Lipid) |
| 1601 | PLD6 | phospholipase D family, member 6 | 201164 | Lipid (Membrane Lipid) |
| 1602 | PPAP2A | phosphatidic acid phosphatase type 2A | 8611 | Lipid (Membrane Lipid) |
| 1603 | PPAP2B | phosphatidic acid phosphatase type 2B | 8613 | Lipid (Membrane Lipid) |
| 1604 | PPAP2C | phosphatidic acid phosphatase type 2C | 8612 | Lipid (Membrane Lipid) |
| 1605 | PPAPDC1A | phosphatidic acid phosphatase type 2 domain containing 1A | 196051 | Lipid (Membrane Lipid) |
| 1606 | PPAPDC1B | phosphatidic acid phosphatase type 2 domain containing 1B | 84513 | Lipid (Membrane Lipid) |
| 1607 | PPAPDC2 | phosphatidic acid phosphatase type 2 domain containing 2 | 403313 | Lipid (Membrane Lipid) |
| 1608 | PPAPDC3 | phosphatidic acid phosphatase type 2 domain containing 3 | 84814 | Lipid (Membrane Lipid) |
| 1609 | PRG2 | plasticity-related gene 2 | 79948 | Lipid (Membrane Lipid) |
| 1610 | PTDSS1 | phosphatidylserine synthase 1 | 9791 | Lipid (Membrane Lipid) |
| 1611 | PTDSS2 | phosphatidylserine synthase 2 | 81490 | Lipid (Membrane Lipid) |
| 1612 | ARSE | arylsulfatase E (chondrodysplasia punctata 1) | 415 | Lipid (Membrane Lipid) |
| 1613 | ARSJ | arylsulfatase family, member J | 79642 | Lipid (Membrane Lipid) |
| 1614 | LPPR4 | plasticity related gene 1 | 9890 | Lipid (Membrane Lipid) |
| 1615 | DDHD1 | DDHD domain containing 1 | 80821 | Lipid (Membrane Lipid) |
| 1616 | GDPD1 | glycerophosphodiester phosphodiesterase domain containing 1 | 284161 | Lipid (Membrane Lipid) |
| 1617 | GDPD2 | glycerophosphodiester phosphodiesterase domain containing 2 | 54857 | Lipid (Membrane Lipid) |
| 1618 | GDPD3 | glycerophosphodiester phosphodiesterase domain containing 3 | 79153 | Lipid (Membrane Lipid) |
| 1619 | GDPD4 | glycerophosphodiester phosphodiesterase domain containing 4 | 220032 | Lipid (Membrane Lipid) |
| 1620 | GDPD5 | glycerophosphodiester phosphodiesterase domain containing 5 | 81544 | Lipid (Membrane Lipid) |
| 1621 | HMGCR | 3-hydroxy-3-methylglutaryl-Coenzyme A reductase | 3156 | Lipid (Mevalonate) |
| 1622 | HMGCS1 | 3-hydroxy-3-methylglutaryl-Coenzyme A synthase 1 (soluble) | 3157 | Lipid (Mevalonate) |
| 1623 | IDI1 | isopentenyl-diphosphate delta isomerase 1 | 3422 | Lipid (Mevalonate) |
| 1624 | IDI2 | isopentenyl-diphosphate delta isomerase 2 | 91734 | Lipid (Mevalonate) |
| 1625 | MVD | mevalonate (diphospho) decarboxylase | 4597 | Lipid (Mevalonate) |
| 1626 | MVK | mevalonate kinase (mevalonic aciduria) | 4598 | Lipid (Mevalonate) |
| 1627 | PMVK | phosphomevalonate kinase | 10654 | Lipid (Mevalonate) |
| 1628 | MCEE | methylmalonyl CoA epimerase | 84693 | Lipid (Propanoate) |
| 1629 | MUT | methylmalonyl Coenzyme A mutase | 4594 | Lipid (Propanoate) |
| 1630 | PCCA | propionyl Coenzyme A carboxylase, alpha polypeptide | 5095 | Lipid (Propanoate) |
| 1631 | PCCB | Propionyl Coenzyme A carboxylase, beta polypeptide | 5096 | Lipid (Propanoate) |
| 1632 | CERK | ceramide kinase | 64781 | Lipid (Sphingolipid) |
| 1633 | CERKL | ceramide kinase-like | 375298 | Lipid (Sphingolipid) |
| 1634 | DEGS1 | degenerative spermatocyte homolog 1, lipid desaturase (Drosophila) | 8560 | Lipid (Sphingolipid) |
| 1635 | DEGS2 | degenerative spermatocyte homolog 2, lipid desaturase (Drosophila) | 123099 | Lipid (Sphingolipid) |
| 1636 | GALC | galactosylceramidase | 2581 | Lipid (Sphingolipid) |
| 1637 | LASS1 | LAG1 homolog, ceramide synthase 1 | 10715 | Lipid (Sphingolipid) |
| 1638 | LASS2 | LAG1 homolog, ceramide synthase 2 | 29956 | Lipid (Sphingolipid) |
| 1639 | LASS3 | LAG1 homolog, ceramide synthase 3 | 204219 | Lipid (Sphingolipid) |
| 1640 | LASS4 | LAG1 homolog, ceramide synthase 4 | 79603 | Lipid (Sphingolipid) |
| 1641 | LASS5 | LAG1 homolog, ceramide synthase 5 | 91012 | Lipid (Sphingolipid) |
| 1642 | LASS6 | LAG1 homolog, ceramide synthase 6 | 253782 | Lipid (Sphingolipid) |
| 1643 | SGMS1 | sphingomyelin synthase 1 | 259230 | Lipid (Sphingolipid) |
| 1644 | SGMS2 | sphingomyelin synthase 2 | 166929 | Lipid (Sphingolipid) |
| 1645 | SGPL1 | sphingosine-1-phosphate lyase 1 | 8879 | Lipid (Sphingolipid) |
| 1646 | SGPP1 | sphingosine-1-phosphate phosphatase 1 | 81537 | Lipid (Sphingolipid) |
| 1647 | SGPP2 | sphingosine-1-phosphate phosphotase 2 | 130367 | Lipid (Sphingolipid) |
| 1648 | SMPD1 | sphingomyelin phosphodiesterase 1, acid lysosomal (acid sphingomyelinase) | 6609 | Lipid (Sphingolipid) |
| 1649 | SMPD2 | sphingomyelin phosphodiesterase 2, neutral membrane (neutral sphingomyelinase) | 6610 | Lipid (Sphingolipid) |
| 1650 | SMPD3 | sphingomyelin phosphodiesterase 3, neutral membrane (neutral sphingomyelinase II) | 55512 | Lipid (Sphingolipid) |
| 1651 | SMPD4 | sphingomyelin phosphodiesterase 4, neutral membrane (neutral sphingomyelinase-3) | 55627 | Lipid (Sphingolipid) |
| 1652 | SMPDL3A | Sphingomyelin phosphodiesterase, acid-like 3A | 10924 | Lipid (Sphingolipid) |
| 1653 | SMPDL3B | sphingomyelin phosphodiesterase, acid-like 3B | 27293 | Lipid (Sphingolipid) |
| 1654 | SPHK1 | sphingosine kinase 1 | 8877 | Lipid (Sphingolipid) |
| 1655 | SPHK2 | sphingosine kinase 2 | 56848 | Lipid (Sphingolipid) |
| 1656 | SPTLC1 | serine palmitoyltransferase, long chain base subunit 1 | 10558 | Lipid (Sphingolipid) |
| 1657 | SPTLC2 | serine palmitoyltransferase, long chain base subunit 2 | 9517 | Lipid (Sphingolipid) |
| 1658 | SPTLC3 | serine palmitoyltransferase, long chain base subunit 3 | 55304 | Lipid (Sphingolipid) |
| 1659 | UGCG | UDP-glucose ceramide glucosyltransferase | 7357 | Lipid (Sphingolipid) |
| 1660 | UGCGL1 | UDP-glucose ceramide glucosyltransferase-like 1 | 56886 | Lipid (Sphingolipid) |
| 1661 | UGCGL2 | UDP-glucose ceramide glucosyltransferase-like 2 | 55757 | Lipid (Sphingolipid) |
| 1662 | UGT8 | UDP glycosyltransferase 8 (UDP-galactose ceramide galactosyltransferase) | 7368 | Lipid (Sphingolipid) |
| 1663 | ACER1 | alkaline ceramidase 1 | 125981 | Lipid (Sphingosine) |
| 1664 | ACER2 | alkaline ceramidase 2 | 340485 | Lipid (Sphingosine) |
| 1665 | ACER3 | phytoceramidase, alkaline | 55331 | Lipid (Sphingosine) |
| 1666 | ASAH1 | N-acylsphingosine amidohydrolase (acid ceramidase) 1 | 427 | Lipid (Sphingosine) |
| 1667 | ASAH2 | N-acylsphingosine amidohydrolase (non-lysosomal ceramidase) 2 | 56624 | Lipid (Sphingosine) |
| 1668 | ASAH2B | N-acylsphingosine amidohydrolase (non-lysosomal ceramidase) 2B | 653308 | Lipid (Sphingosine) |
| 1669 | ASAH2C | N-acylsphingosine amidohydrolase (non-lysosomal ceramidase) 2C | 653365 | Lipid (Sphingosine) |
| 1670 | ASAHL | N-acylsphingosine amidohydrolase (acid ceramidase)-like | 27163 | Lipid (Sphingosine) |
| 1671 | KDSR | 3-ketodihydrosphingosine reductase | 2531 | Lipid (Sphingosine) |
| 1672 | AKR1C4 | aldo-keto reductase family 1, member C4 (chlordecone reductase; 3-alpha hydroxysteroid dehydrogenase, type I; dihydrodiol dehydrogenase 4) | 1109 | Lipid (Steroid) |
| 1673 | AKR1D1 | aldo-keto reductase family 1, member D1 (delta 4-3-ketosteroid-5-beta-reductase) | 6718 | Lipid (Steroid) |
| 1674 | CH25H | cholesterol 25-hydroxylase | 9023 | Lipid (Steroid) |
| 1675 | CYP11A1 | cytochrome P450, family 11, subfamily A, polypeptide 1 | 1583 | Lipid (Steroid) |
| 1676 | CYP11B1 | cytochrome P450, family 11, subfamily B, polypeptide 1 | 1584 | Lipid (Steroid) |
| 1677 | CYP11B2 | cytochrome P450, family 11, subfamily B, polypeptide 2 | 1585 | Lipid (Steroid) |
| 1678 | CYP17A1 | cytochrome P450, family 17, subfamily A, polypeptide 1 | 1586 | Lipid (Steroid) |
| 1679 | CYP19A1 | cytochrome P450, family 19, subfamily A, polypeptide 1 | 1588 | Lipid (Steroid) |
| 1680 | CYP1A1 | cytochrome P450, family 1, subfamily A, polypeptide 1 | 1543 | Lipid (Steroid) |
| 1681 | CYP1A2 | cytochrome P450, family 1, subfamily A, polypeptide 2 | 1544 | Lipid (Steroid) |
| 1682 | CYP1B1 | cytochrome P450, family 1, subfamily B, polypeptide 1 | 1545 | Lipid (Steroid) |
| 1683 | CYP21A2 | cytochrome P450, family 21, subfamily A, polypeptide 2 | 1589 | Lipid (Steroid) |
| 1684 | CYP27C1 | cytochrome P450, family 27, subfamily C, polypeptide 1 | 339761 | Lipid (Steroid) |
| 1685 | CYP2A13 | cytochrome P450, family 2, subfamily A, polypeptide 13 | 1553 | Lipid (Steroid) |
| 1686 | CYP2A6 | cytochrome P450, family 2, subfamily A, polypeptide 6 | 1548 | Lipid (Steroid) |
| 1687 | CYP2A7 | cytochrome P450, family 2, subfamily A, polypeptide 7 | 1549 | Lipid (Steroid) |
| 1688 | CYP2B6 | cytochrome P450, family 2, subfamily B, polypeptide 6 | 1555 | Lipid (Steroid) |
| 1689 | CYP2C18 | cytochrome P450, family 2, subfamily C, polypeptide 18 | 1562 | Lipid (Steroid) |
| 1690 | CYP2C19 | cytochrome P450, family 2, subfamily C, polypeptide 19 | 1557 | Lipid (Steroid) |
| 1691 | CYP2C8 | cytochrome P450, family 2, subfamily C, polypeptide 8 | 1558 | Lipid (Steroid) |
| 1692 | CYP2C9 | Cytochrome P450, family 2, subfamily C, polypeptide 9 | 1559 | Lipid (Steroid) |
| 1693 | CYP2D6 | Cytochrome P450, family 2, subfamily D, polypeptide 6 | 1565 | Lipid (Steroid) |
| 1694 | CYP2E1 | cytochrome P450, family 2, subfamily E, polypeptide 1 | 1571 | Lipid (Steroid) |
| 1695 | CYP2F1 | cytochrome P450, family 2, subfamily F, polypeptide 1 | 1572 | Lipid (Steroid) |
| 1696 | CYP2J2 | cytochrome P450, family 2, subfamily J, polypeptide 2 | 1573 | Lipid (Steroid) |
| 1697 | CYP2R1 | cytochrome P450, family 2, subfamily R, polypeptide 1 | 120227 | Lipid (Steroid) |
| 1698 | CYP2S1 | cytochrome P450, family 2, subfamily S, polypeptide 1 | 29785 | Lipid (Steroid) |
| 1699 | CYP2U1 | cytochrome P450, family 2, subfamily U, polypeptide 1 | 113612 | Lipid (Steroid) |
| 1700 | CYP2W1 | cytochrome P450, family 2, subfamily W, polypeptide 1 | 54905 | Lipid (Steroid) |
| 1701 | CYP39A1 | cytochrome P450, family 39, subfamily A, polypeptide 1 | 51302 | Lipid (Steroid) |
| 1702 | CYP3A4 | Cytochrome P450, family 3, subfamily A, polypeptide 4 | 1576 | Lipid (Steroid) |
| 1703 | CYP3A43 | cytochrome P450, family 3, subfamily A, polypeptide 43 | 64816 | Lipid (Steroid) |
| 1704 | CYP3A5 | cytochrome P450, family 3, subfamily A, polypeptide 5 | 1577 | Lipid (Steroid) |
| 1705 | CYP3A7 | cytochrome P450, family 3, subfamily A, polypeptide 7 | 1551 | Lipid (Steroid) |
| 1706 | CYP46A1 | cytochrome P450, family 46, subfamily A, polypeptide 1 | 10858 | Lipid (Steroid) |
| 1707 | GGPS1 | geranylgeranyl diphosphate synthase 1 | 9453 | Lipid (Steroid) |
| 1708 | HSD11B1 | hydroxysteroid (11-beta) dehydrogenase 1 | 3290 | Lipid (Steroid) |
| 1709 | HSD11B1L | hydroxysteroid (11-beta) dehydrogenase 1-like | 374875 | Lipid (Steroid) |
| 1710 | HSD11B2 | hydroxysteroid (11-beta) dehydrogenase 2 | 3291 | Lipid (Steroid) |
| 1711 | HSD17B1 | hydroxysteroid (17-beta) dehydrogenase 1 | 3292 | Lipid (Steroid) |
| 1712 | HSD17B10 | hydroxysteroid (17-beta) dehydrogenase 10 | 3028 | Lipid (Steroid) |
| 1713 | HSD17B11 | hydroxysteroid (17-beta) dehydrogenase 11 | 51170 | Lipid (Steroid) |
| 1714 | HSD17B12 | hydroxysteroid (17-beta) dehydrogenase 12 | 51144 | Lipid (Steroid) |
| 1715 | HSD17B13 | short-chain dehydrogenase/reductase 9 | 345275 | Lipid (Steroid) |
| 1716 | HSD17B14 | hydroxysteroid (17-beta) dehydrogenase 14 | 51171 | Lipid (Steroid) |
| 1717 | HSD17B2 | hydroxysteroid (17-beta) dehydrogenase 2 | 3294 | Lipid (Steroid) |
| 1718 | HSD17B3 | hydroxysteroid (17-beta) dehydrogenase 3 | 3293 | Lipid (Steroid) |
| 1719 | HSD17B4 | Hydroxysteroid (17-beta) dehydrogenase 4 | 3295 | Lipid (Steroid) |
| 1720 | HSD17B6 | hydroxysteroid (17-beta) dehydrogenase 6 homolog (mouse) | 8630 | Lipid (Steroid) |
| 1721 | HSD17B8 | hydroxysteroid (17-beta) dehydrogenase 8 | 7923 | Lipid (Steroid) |
| 1722 | HSD3B1 | hydroxy-delta-5-steroid dehydrogenase, 3 beta- and steroid delta-isomerase 1 | 3283 | Lipid (Steroid) |
| 1723 | HSD3B2 | hydroxy-delta-5-steroid dehydrogenase, 3 beta- and steroid delta-isomerase 2 | 3284 | Lipid (Steroid) |
| 1724 | HSDL1 | hydroxysteroid dehydrogenase like 1 | 83693 | Lipid (Steroid) |
| 1725 | HSDL2 | hydroxysteroid dehydrogenase like 2 | 84263 | Lipid (Steroid) |
| 1726 | KL | klotho | 9365 | Lipid (Steroid) |
| 1727 | KLB | klotho beta | 152831 | Lipid (Steroid) |
| 1728 | LIPA | Lipase A, lysosomal acid, cholesterol esterase (Wolman disease) | 3988 | Lipid (Steroid) |
| 1729 | LIPE | lipase, hormone-sensitive | 3991 | Lipid (Steroid) |
| 1730 | SOAT1 | sterol O-acyltransferase (acyl-Coenzyme A: cholesterol acyltransferase) 1 | 6646 | Lipid (Steroid) |
| 1731 | SOAT2 | sterol O-acyltransferase 2 | 8435 | Lipid (Steroid) |
| 1732 | SRD5A1 | steroid-5-alpha-reductase, alpha polypeptide 1 (3-oxo-5 alpha-steroid delta 4-dehydrogenase alpha 1) | 6715 | Lipid (Steroid) |
| 1733 | SRD5A2 | steroid-5-alpha-reductase, alpha polypeptide 2 (3-oxo-5 alpha-steroid delta 4-dehydrogenase alpha 2) | 6716 | Lipid (Steroid) |
| 1734 | SRD5A2L2 | steroid 5 alpha-reductase 2-like 2 | 253017 | Lipid (Steroid) |
| 1735 | SRD5A3 | steroid 5 alpha-reductase 3 | 79644 | Lipid (Steroid) |
| 1736 | BST1 | bone marrow stromal cell antigen 1 | 683 | NAD |
| 1737 | C9ORF95 | Chromosome 9 ORF 95 | 54981 | NAD |
| 1738 | DUOX1 | dual oxidase 1 | 53905 | NAD |
| 1739 | DUOX2 | dual oxidase 2 | 50506 | NAD |
| 1740 | NADK | NAD kinase | 65220 | NAD |
| 1741 | NADSYN1 | NAD synthetase 1 | 55191 | NAD |
| 1742 | NAPRT1 | nicotinate phosphoribosyltransferase domain containing 1 | 93100 | NAD |
| 1743 | NMNAT1 | nicotinamide nucleotide adenylyltransferase 1 | 64802 | NAD |
| 1744 | NMNAT2 | nicotinamide nucleotide adenylyltransferase 2 | 23057 | NAD |
| 1745 | NMNAT3 | nicotinamide nucleotide adenylyltransferase 3 | 349565 | NAD |
| 1746 | NNMT | nicotinamide N-methyltransferase | 4837 | NAD |
| 1747 | NNT | nicotinamide nucleotide transhydrogenase | 23530 | NAD |
| 1748 | NOX1 | NADPH oxidase 1 | 27035 | NAD |
| 1749 | NOX3 | NADPH oxidase 3 | 50508 | NAD |
| 1750 | NOX4 | NADPH oxidase 4 | 50507 | NAD |
| 1751 | NOX5 | NADPH oxidase, EF-hand calcium binding domain 5 | 79400 | NAD |
| 1752 | NQO1 | NAD(P)H dehydrogenase, quinone 1 | 1728 | NAD |
| 1753 | NQO2 | NAD(P)H dehydrogenase, quinone 2 | 4835 | NAD |
| 1754 | QPRT | quinolinate phosphoribosyltransferase (nicotinate-nucleotide pyrophosphorylase (carboxylating)) | 23475 | NAD |
| 1755 | CD38 | CD38 | 952 | NAD |
| 1756 | NAMPT | nicotinamide phosphoribosyltransferase | 10135 | NAD |
| 1757 | ABAT | 4-aminobutyrate aminotransferase | 18 | Neurotransmitter |
| 1758 | ACHE | acetylcholinesterase (Yt blood group) | 43 | Neurotransmitter |
| 1759 | CHAT | choline acetyltransferase | 1103 | Neurotransmitter |
| 1760 | COMT | catechol-O-methyltransferase | 1312 | Neurotransmitter |
| 1761 | COMTD1 | catechol-O-methyltransferase domain containing 1 | 118881 | Neurotransmitter |
| 1762 | DBH | dopamine beta-hydroxylase (dopamine beta-monooxygenase) | 1621 | Neurotransmitter |
| 1763 | DDC | dopa decarboxylase (aromatic L-amino acid decarboxylase) | 1644 | Neurotransmitter |
| 1764 | NAALAD2 | N-acetylated alpha-linked acidic dipeptidase 2 | 10003 | Neurotransmitter |
| 1765 | NAALADL1 | N-acetylated alpha-linked acidic dipeptidase-like 1 | 10004 | Neurotransmitter |
| 1766 | NAALADL2 | N-acetylated alpha-linked acidic dipeptidase-like 2 | 254827 | Neurotransmitter |
| 1767 | CA1 | carbonic anhydrase I | 759 | Nitrogen Metabolism |
| 1768 | CA10 | carbonic anhydrase X | 56934 | Nitrogen Metabolism |
| 1769 | CA11 | carbonic anhydrase XI | 770 | Nitrogen Metabolism |
| 1770 | CA12 | carbonic anhydrase XII | 771 | Nitrogen Metabolism |
| 1771 | CA13 | carbonic anhydrase XIII | 377677 | Nitrogen Metabolism |
| 1772 | CA14 | carbonic anhydrase XIV | 23632 | Nitrogen Metabolism |
| 1773 | CA2 | carbonic anhydrase II | 760 | Nitrogen Metabolism |
| 1774 | CA3 | carbonic anhydrase III, muscle specific | 761 | Nitrogen Metabolism |
| 1775 | CA4 | carbonic anhydrase IV | 762 | Nitrogen Metabolism |
| 1776 | CA5A | carbonic anhydrase VA, mitochondrial | 763 | Nitrogen Metabolism |
| 1777 | CA5B | carbonic anhydrase VB, mitochondrial | 11238 | Nitrogen Metabolism |
| 1778 | CA6 | carbonic anhydrase VI | 765 | Nitrogen Metabolism |
| 1779 | CA7 | carbonic anhydrase VII | 766 | Nitrogen Metabolism |
| 1780 | CA8 | carbonic anhydrase VIII | 767 | Nitrogen Metabolism |
| 1781 | CA9 | carbonic anhydrase IX | 768 | Nitrogen Metabolism |
| 1782 | NIT1 | nitrilase 1 | 4817 | Nitrogen Metabolism |
| 1783 | ABP1 | amiloride binding protein 1 (amine oxidase (copper-containing)) | 26 | Nucleotide |
| 1784 | BPNT1 | 3'(2'), 5'-bisphosphate nucleotidase 1 | 10380 | Nucleotide |
| 1785 | CANT1 | calcium activated nucleotidase 1 | 124583 | Nucleotide |
| 1786 | DCK | deoxycytidine kinase | 1633 | Nucleotide |
| 1787 | ENPP1 | ectonucleotide pyrophosphatase/phosphodiesterase 1 | 5167 | Nucleotide |
| 1788 | ENPP2 | ectonucleotide pyrophosphatase/phosphodiesterase 2 (autotaxin) | 5168 | Nucleotide |
| 1789 | ENPP3 | ectonucleotide pyrophosphatase/phosphodiesterase 3 | 5169 | Nucleotide |
| 1790 | ENPP4 | ectonucleotide pyrophosphatase/phosphodiesterase 4 (putative function) | 22875 | Nucleotide |
| 1791 | ENPP5 | ectonucleotide pyrophosphatase/phosphodiesterase 5 (putative function) | 59084 | Nucleotide |
| 1792 | ENPP6 | ectonucleotide pyrophosphatase/phosphodiesterase 6 | 133121 | Nucleotide |
| 1793 | ENPP7 | ectonucleotide pyrophosphatase/phosphodiesterase 7 | 339221 | Nucleotide |
| 1794 | ENTPD1 | ectonucleoside triphosphate diphosphohydrolase 1 | 953 | Nucleotide |
| 1795 | ENTPD2 | ectonucleoside triphosphate diphosphohydrolase 2 | 954 | Nucleotide |
| 1796 | ENTPD3 | ectonucleoside triphosphate diphosphohydrolase 3 | 956 | Nucleotide |
| 1797 | ENTPD4 | ectonucleoside triphosphate diphosphohydrolase 4 | 9583 | Nucleotide |
| 1798 | ENTPD5 | ectonucleoside triphosphate diphosphohydrolase 5 | 957 | Nucleotide |
| 1799 | ENTPD6 | ectonucleoside triphosphate diphosphohydrolase 6 (putative function) | 955 | Nucleotide |
| 1800 | ENTPD7 | ectonucleoside triphosphate diphosphohydrolase 7 | 57089 | Nucleotide |
| 1801 | ENTPD8 | ectonucleoside triphosphate diphosphohydrolase 8 | 377841 | Nucleotide |
| 1802 | ITPA | inosine triphosphatase (nucleoside triphosphate pyrophosphatase) | 3704 | Nucleotide |
| 1803 | NME1 | non-metastatic cells 1, protein (NM23A) expressed in | 4830 | Nucleotide |
| 1804 | NME2 | non-metastatic cells 2, protein (NM23A) expressed in | 4831 | Nucleotide |
| 1805 | NME3 | non-metastatic cells 3, protein expressed in | 4832 | Nucleotide |
| 1806 | NME4 | non-metastatic cells 4, protein expressed in | 4833 | Nucleotide |
| 1807 | NME5 | non-metastatic cells 5, protein expressed in (nucleoside-diphosphate kinase) | 8382 | Nucleotide |
| 1808 | NME6 | non-metastatic cells 6, protein expressed in (nucleoside-diphosphate kinase) | 10201 | Nucleotide |
| 1809 | NME7 | non-metastatic cells 7, protein expressed in (nucleoside-diphosphate kinase) | 29922 | Nucleotide |
| 1810 | NT5C | 5', 3'-nucleotidase, cytosolic | 30833 | Nucleotide |
| 1811 | NT5C1A | 5'-nucleotidase, cytosolic IA | 84618 | Nucleotide |
| 1812 | NT5C1B | 5'-nucleotidase, cytosolic IB | 93034 | Nucleotide |
| 1813 | NT5C2 | 5'-nucleotidase, cytosolic II | 22978 | Nucleotide |
| 1814 | NT5C3 | 5'-nucleotidase, cytosolic III | 51251 | Nucleotide |
| 1815 | NT5C3L | 5'-nucleotidase, cytosolic III-like | 115024 | Nucleotide |
| 1816 | NT5DC1 | 5'-nucleotidase domain containing 1 | 221294 | Nucleotide |
| 1817 | NT5DC2 | 5'-nucleotidase domain containing 2 | 64943 | Nucleotide |
| 1818 | NT5DC3 | 5'-nucleotidase domain containing 3 | 51559 | Nucleotide |
| 1819 | NT5DC4 | 5'-nucleotidase domain containing 4 | 284958 | Nucleotide |
| 1820 | NT5E | 5'-nucleotidase, ecto (CD73) | 4907 | Nucleotide |
| 1821 | NT5M | 5',3'-nucleotidase, mitochondrial | 56953 | Nucleotide |
| 1822 | NUDT1 | nudix (nucleoside diphosphate linked moiety X)-type motif 1 | 4521 | Nucleotide |
| 1823 | NUDT10 | nudix (nucleoside diphosphate linked moiety X)-type motif 10 | 170685 | Nucleotide |
| 1824 | NUDT11 | nudix (nucleoside diphosphate linked moiety X)-type motif 11 | 55190 | Nucleotide |
| 1825 | NUDT12 | nudix (nucleoside diphosphate linked moiety X)-type motif 12 | 83594 | Nucleotide |
| 1826 | NUDT13 | nudix (nucleoside diphosphate linked moiety X)-type motif 13 | 25961 | Nucleotide |
| 1827 | NUDT14 | Nudix (nucleoside diphosphate linked moiety X)-type motif 14 | 256281 | Nucleotide |
| 1828 | NUDT15 | nudix (nucleoside diphosphate linked moiety X)-type motif 15 | 55270 | Nucleotide |
| 1829 | NUDT16 | nudix (nucleoside diphosphate linked moiety X)-type motif 16 | 131870 | Nucleotide |
| 1830 | NUDT16L1 | nudix (nucleoside diphosphate linked moiety X)-type motif 16-like 1 | 84309 | Nucleotide |
| 1831 | NUDT17 | nudix (nucleoside diphosphate linked moiety X)-type motif 17 | 200035 | Nucleotide |
| 1832 | NUDT18 | nudix (nucleoside diphosphate linked moiety X)-type motif 18 | 79873 | Nucleotide |
| 1833 | NUDT19 | nudix (nucleoside diphosphate linked moiety X)-type motif 19 | 390916 | Nucleotide |
| 1834 | NUDT2 | nudix (nucleoside diphosphate linked moiety X)-type motif 2 | 318 | Nucleotide |
| 1835 | NUDT21 | nudix (nucleoside diphosphate linked moiety X)-type motif 21 | 11051 | Nucleotide |
| 1836 | NUDT22 | Nudix (nucleoside diphosphate linked moiety X)-type motif 22 | 84304 | Nucleotide |
| 1837 | NUDT3 | nudix (nucleoside diphosphate linked moiety X)-type motif 3 | 11165 | Nucleotide |
| 1838 | NUDT4 | nudix (nucleoside diphosphate linked moiety X)-type motif 4 | 11163 | Nucleotide |
| 1839 | NUDT5 | nudix (nucleoside diphosphate linked moiety X)-type motif 5 | 11164 | Nucleotide |
| 1840 | NUDT6 | nudix (nucleoside diphosphate linked moiety X)-type motif 6 | 11162 | Nucleotide |
| 1841 | NUDT7 | nudix (nucleoside diphosphate linked moiety X)-type motif 7 | 283927 | Nucleotide |
| 1842 | NUDT8 | nudix (nucleoside diphosphate linked moiety X)-type motif 8 | 254552 | Nucleotide |
| 1843 | NUDT9 | nudix (nucleoside diphosphate linked moiety X)-type motif 9 | 53343 | Nucleotide |
| 1844 | RRM1 | ribonucleotide reductase M1 polypeptide | 6240 | Nucleotide |
| 1845 | RRM2 | ribonucleotide reductase M2 polypeptide | 6241 | Nucleotide |
| 1846 | RRM2B | ribonucleotide reductase M2 B (TP53 inducible) | 50484 | Nucleotide |
| 1847 | A1CF | APOBEC1 complementation factor | 29974 | Nucleotide |
| 1848 | ADCY1 | adenylate cyclase 1 (brain) | 107 | Nucleotide (cAMP) |
| 1849 | ADCY10 | testicular soluble adenylyl cyclase | 55811 | Nucleotide (cAMP) |
| 1850 | ADCY2 | adenylate cyclase 2 (brain) | 108 | Nucleotide (cAMP) |
| 1851 | ADCY3 | adenylate cyclase 3 | 109 | Nucleotide (cAMP) |
| 1852 | ADCY4 | adenylate cyclase 4 | 196883 | Nucleotide (cAMP) |
| 1853 | ADCY5 | adenylate cyclase 5 | 111 | Nucleotide (cAMP) |
| 1854 | ADCY6 | adenylate cyclase 6 | 112 | Nucleotide (cAMP) |
| 1855 | ADCY7 | adenylate cyclase 7 | 113 | Nucleotide (cAMP) |
| 1856 | ADCY8 | adenylate cyclase 8 (brain) | 114 | Nucleotide (cAMP) |
| 1857 | ADCY9 | Adenylate cyclase 9 | 115 | Nucleotide (cAMP) |
| 1858 | GUCY1A2 | guanylate cyclase 1, soluble, alpha 2 | 2977 | Nucleotide (cGMP) |
| 1859 | GUCY1A3 | guanylate cyclase 1, soluble, alpha 3 | 2982 | Nucleotide (cGMP) |
| 1860 | GUCY1B3 | guanylate cyclase 1, soluble, beta 3 | 2983 | Nucleotide (cGMP) |
| 1861 | GUCY2C | guanylate cyclase 2C (heat stable enterotoxin receptor) | 2984 | Nucleotide (cGMP) |
| 1862 | GUCY2D | guanylate cyclase 2D, membrane (retina-specific) | 3000 | Nucleotide (cGMP) |
| 1863 | GUCY2F | guanylate cyclase 2F, retinal | 2986 | Nucleotide (cGMP) |
| 1864 | NPR1 | natriuretic peptide receptor A/guanylate cyclase A (atrionatriuretic peptide receptor A) | 4881 | Nucleotide (cGMP) |
| 1865 | NPR2 | natriuretic peptide receptor B/guanylate cyclase B (atrionatriuretic peptide receptor B) | 4882 | Nucleotide (cGMP) |
| 1866 | NPR3 | natriuretic peptide receptor C/guanylate cyclase C (atrionatriuretic peptide receptor C) | 4883 | Nucleotide (cGMP) |
| 1867 | PDE1C | phosphodiesterase 1C, calmodulin-dependent 70kDa | 5137 | Nucleotide (cGMP) |
| 1868 | PDE2A | phosphodiesterase 2A, cGMP-stimulated | 5138 | Nucleotide (cGMP) |
| 1869 | PDE3A | phosphodiesterase 3A, cGMP-inhibited | 5139 | Nucleotide (cGMP) |
| 1870 | PDE3B | phosphodiesterase 3B, cGMP-inhibited | 5140 | Nucleotide (cGMP) |
| 1871 | PDE4A | phosphodiesterase 4A, cAMP-specific (phosphodiesterase E2 dunce homolog, Drosophila) | 5141 | Nucleotide (cGMP) |
| 1872 | PDE4B | phosphodiesterase 4B, cAMP-specific (phosphodiesterase E4 dunce homolog, Drosophila) | 5142 | Nucleotide (cGMP) |
| 1873 | PDE4C | phosphodiesterase 4C, cAMP-specific (phosphodiesterase E1 dunce homolog, Drosophila) | 5143 | Nucleotide (cGMP) |
| 1874 | PDE4D | phosphodiesterase 4D, cAMP-specific (phosphodiesterase E3 dunce homolog, Drosophila) | 5144 | Nucleotide (cGMP) |
| 1875 | PDE5A | phosphodiesterase 5A, cGMP-specific | 8654 | Nucleotide (cGMP) |
| 1876 | PDE6A | phosphodiesterase 6A, cGMP-specific, rod, alpha | 5145 | Nucleotide (cGMP) |
| 1877 | PDE6B | phosphodiesterase 6B, cGMP-specific, rod, beta (congenital stationary night blindness 3, autosomal dominant) | 5158 | Nucleotide (cGMP) |
| 1878 | PDE6C | phosphodiesterase 6C, cGMP-specific, cone, alpha prime | 5146 | Nucleotide (cGMP) |
| 1879 | PDE6D | phosphodiesterase 6D, cGMP-specific, rod, delta | 5147 | Nucleotide (cGMP) |
| 1880 | PDE6G | phosphodiesterase 6G, cGMP-specific, rod, gamma | 5148 | Nucleotide (cGMP) |
| 1881 | PDE6H | phosphodiesterase 6H, cGMP-specific, cone, gamma | 5149 | Nucleotide (cGMP) |
| 1882 | PDE9A | phosphodiesterase 9A | 5152 | Nucleotide (cGMP) |
| 1883 | C9ORF98 | Chromosome 9 ORF 98 | 158067 | Nucleotide (cyclic nucleotide) |
| 1884 | CNP | 2',3'-cyclic nucleotide 3' phosphodiesterase | 1267 | Nucleotide (cyclic nucleotide) |
| 1885 | PDE10A | phosphodiesterase 10A | 10846 | Nucleotide (cyclic nucleotide) |
| 1886 | PDE11A | phosphodiesterase 11A | 50940 | Nucleotide (cyclic nucleotide) |
| 1887 | PDE12 | 2'-phosphodiesterase | 201626 | Nucleotide (cyclic nucleotide) |
| 1888 | PDE1A | phosphodiesterase 1A, calmodulin-dependent | 5136 | Nucleotide (cyclic nucleotide) |
| 1889 | PDE1B | phosphodiesterase 1B, calmodulin-dependent | 5153 | Nucleotide (cyclic nucleotide) |
| 1890 | PDE7A | phosphodiesterase 7A | 5150 | Nucleotide (cyclic nucleotide) |
| 1891 | PDE7B | phosphodiesterase 7B | 27115 | Nucleotide (cyclic nucleotide) |
| 1892 | PDE8A | phosphodiesterase 8A | 5151 | Nucleotide (cyclic nucleotide) |
| 1893 | PDE8B | phosphodiesterase 8B | 8622 | Nucleotide (cyclic nucleotide) |
| 1894 | ADA | adenosine deaminase | 100 | Nucleotide (Purine) |
| 1895 | ADAD1 | Adenosine deaminase domain containing 1 (testis-specific) | 132612 | Nucleotide (Purine) |
| 1896 | ADAD2 | adenosine deaminase domain containing 2 | 161931 | Nucleotide (Purine) |
| 1897 | ADAL | adenosine deaminase-like | 161823 | Nucleotide (Purine) |
| 1898 | ADK | adenosine kinase | 132 | Nucleotide (Purine) |
| 1899 | ADSL | adenylosuccinate lyase | 158 | Nucleotide (Purine) |
| 1900 | ADSS | Adenylosuccinate synthase | 159 | Nucleotide (Purine) |
| 1901 | ADSSL1 | adenylosuccinate synthase like 1 | 122622 | Nucleotide (Purine) |
| 1902 | AK1 | adenylate kinase 1 | 203 | Nucleotide (Purine) |
| 1903 | AK2 | adenylate kinase 2 | 204 | Nucleotide (Purine) |
| 1904 | AK3 | adenylate kinase 3 | 50808 | Nucleotide (Purine) |
| 1905 | AK3L1 | adenylate kinase 3-like 1 | 205 | Nucleotide (Purine) |
| 1906 | AK5 | adenylate kinase 5 | 26289 | Nucleotide (Purine) |
| 1907 | AK7 | adenylate kinase 7 | 122481 | Nucleotide (Purine) |
| 1908 | AMPD1 | adenosine monophosphate deaminase 1 (isoform M) | 270 | Nucleotide (Purine) |
| 1909 | AMPD2 | adenosine monophosphate deaminase 2 (isoform L) | 271 | Nucleotide (Purine) |
| 1910 | AMPD3 | adenosine monophosphate deaminase (isoform E) | 272 | Nucleotide (Purine) |
| 1911 | APRT | adenine phosphoribosyltransferase | 353 | Nucleotide (Purine) |
| 1912 | ATIC | 5-aminoimidazole-4-carboxamide ribonucleotide formyltransferase/IMP cyclohydrolase | 471 | Nucleotide (Purine) |
| 1913 | CECR1 | cat eye syndrome chromosome region, candidate 1 | 51816 | Nucleotide (Purine) |
| 1914 | DGUOK | deoxyguanosine kinase | 1716 | Nucleotide (Purine) |
| 1915 | FHIT | Fragile histidine triad gene | 2272 | Nucleotide (Purine) |
| 1916 | GART | phosphoribosylglycinamide formyltransferase, phosphoribosylglycinamide synthetase, phosphoribosylaminoimidazole synthetase | 2618 | Nucleotide (Purine) |
| 1917 | GDA | guanine deaminase | 9615 | Nucleotide (Purine) |
| 1918 | GMPR | guanosine monophosphate reductase | 2766 | Nucleotide (Purine) |
| 1919 | GMPR2 | guanosine monophosphate reductase 2 | 51292 | Nucleotide (Purine) |
| 1920 | GMPS | guanine monphosphate synthetase | 8833 | Nucleotide (Purine) |
| 1921 | GUK1 | guanylate kinase 1 | 2987 | Nucleotide (Purine) |
| 1922 | HPRT1 | Hypoxanthine phosphoribosyltransferase 1 (Lesch-Nyhan syndrome) | 3251 | Nucleotide (Purine) |
| 1923 | IMPDH1 | IMP (inosine monophosphate) dehydrogenase 1 | 3614 | Nucleotide (Purine) |
| 1924 | IMPDH2 | IMP (inosine monophosphate) dehydrogenase 2 | 3615 | Nucleotide (Purine) |
| 1925 | NP | nucleoside phosphorylase | 4860 | Nucleotide (Purine) |
| 1926 | PAICS | phosphoribosylaminoimidazole carboxylase, phosphoribosylaminoimidazole succinocarboxamide synthetase | 10606 | Nucleotide (Purine) |
| 1927 | PFAS | phosphoribosylformylglycinamidine synthase (FGAR amidotransferase) | 5198 | Nucleotide (Purine) |
| 1928 | PPAT | phosphoribosyl pyrophosphate amidotransferase | 5471 | Nucleotide (Purine) |
| 1929 | PRPS1 | phosphoribosyl pyrophosphate synthetase 1 | 5631 | Nucleotide (Purine) |
| 1930 | PRPS1L1 | phosphoribosyl pyrophosphate synthetase 1-like 1 | 221823 | Nucleotide (Purine) |
| 1931 | PRPS2 | phosphoribosyl pyrophosphate synthetase 2 | 5634 | Nucleotide (Purine) |
| 1932 | PRPSAP1 | Phosphoribosyl pyrophosphate synthetase-associated protein 1 | 5635 | Nucleotide (Purine) |
| 1933 | PRPSAP2 | phosphoribosyl pyrophosphate synthetase-associated protein 2 | 5636 | Nucleotide (Purine) |
| 1934 | XDH | xanthine dehydrogenase | 7498 | Nucleotide (Purine) |
| 1935 | CAD | Carbamoyl-phosphate synthetase 2, aspartate transcarbamylase, and dihydroorotase | 790 | Nucleotide (Pyrimidine) |
| 1936 | CDA | cytidine deaminase | 978 | Nucleotide (Pyrimidine) |
| 1937 | CDADC1 | cytidine and dCMP deaminase domain containing 1 | 81602 | Nucleotide (Pyrimidine) |
| 1938 | CMPK | cytidylate kinase | 51727 | Nucleotide (Pyrimidine) |
| 1939 | CTPS | CTP synthase | 1503 | Nucleotide (Pyrimidine) |
| 1940 | CTPS2 | CTP synthase II | 56474 | Nucleotide (Pyrimidine) |
| 1941 | DCTD | dCMP deaminase | 1635 | Nucleotide (Pyrimidine) |
| 1942 | DHODH | dihydroorotate dehydrogenase | 1723 | Nucleotide (Pyrimidine) |
| 1943 | DPYD | dihydropyrimidine dehydrogenase | 1806 | Nucleotide (Pyrimidine) |
| 1944 | DPYS | dihydropyrimidinase | 1807 | Nucleotide (Pyrimidine) |
| 1945 | DPYSL2 | dihydropyrimidinase-like 2 | 1808 | Nucleotide (Pyrimidine) |
| 1946 | DPYSL3 | dihydropyrimidinase-like 3 | 1809 | Nucleotide (Pyrimidine) |
| 1947 | DPYSL4 | dihydropyrimidinase-like 4 | 10570 | Nucleotide (Pyrimidine) |
| 1948 | DPYSL5 | dihydropyrimidinase-like 5 | 56896 | Nucleotide (Pyrimidine) |
| 1949 | DTYMK | deoxythymidylate kinase (thymidylate kinase) | 1841 | Nucleotide (Pyrimidine) |
| 1950 | DUT | deoxyuridine triphosphatase | 1854 | Nucleotide (Pyrimidine) |
| 1951 | TK1 | thymidine kinase 1, soluble | 7083 | Nucleotide (Pyrimidine) |
| 1952 | TK2 | thymidine kinase 2, mitochondrial | 7084 | Nucleotide (Pyrimidine) |
| 1953 | TYMP | Thymidine phosphorylase | 1890 | Nucleotide (Pyrimidine) |
| 1954 | TYMS | thymidylate synthetase | 7298 | Nucleotide (Pyrimidine) |
| 1955 | UCK1 | uridine-cytidine kinase 1 | 83549 | Nucleotide (Pyrimidine) |
| 1956 | UCK2 | uridine-cytidine kinase 2 | 7371 | Nucleotide (Pyrimidine) |
| 1957 | UCKL1 | uridine-cytidine kinase 1-like 1 | 54963 | Nucleotide (Pyrimidine) |
| 1958 | UMPS | uridine monophosphate synthetase (orotate phosphoribosyl transferase and orotidine-5'-decarboxylase) | 7372 | Nucleotide (Pyrimidine) |
| 1959 | UPB1 | ureidopropionase, beta | 51733 | Nucleotide (Pyrimidine) |
| 1960 | UPP1 | uridine phosphorylase 1 | 7378 | Nucleotide (Pyrimidine) |
| 1961 | UPP2 | uridine phosphorylase 2 | 151531 | Nucleotide (Pyrimidine) |
| 1962 | UPRT | uracil phosphoribosyltransferase (FUR1) homolog (S. cerevisiae) | 139596 | Nucleotide (Pyrimidine) |
| 1963 | OAS1 | 2',5'-oligoadenylate synthetase 1, 40/46kDa | 4938 | Nucleotide (Ribonuclease) |
| 1964 | OAS2 | 2'-5'-oligoadenylate synthetase 2, 69/71kDa | 4939 | Nucleotide (Ribonuclease) |
| 1965 | OAS3 | 2'-5'-oligoadenylate synthetase 3, 100kDa | 4940 | Nucleotide (Ribonuclease) |
| 1966 | OASL | 2'-5'-oligoadenylate synthetase-like | 8638 | Nucleotide (Ribonuclease) |
| 1967 | FECH | ferrochelatase (protoporphyria) | 2235 | Porphyrin |
| 1968 | FXN | frataxin | 2395 | Porphyrin |
| 1969 | ALAS1 | aminolevulinate, delta-, synthase 1 | 211 | Porphyrin |
| 1970 | ALAS2 | aminolevulinate, delta-, synthase 2 (sideroblastic/hypochromic anemia) | 212 | Porphyrin |
| 1971 | ALAD | aminolevulinate, delta-, dehydratase | 210 | Porphyrin |
| 1972 | CPOX | coproporphyrinogen oxidase | 1371 | Porphyrin |
| 1973 | HMBS | hydroxymethylbilane synthase | 3145 | Porphyrin |
| 1974 | PPOX | protoporphyrinogen oxidase | 5498 | Porphyrin |
| 1975 | UROD | uroporphyrinogen decarboxylase | 7389 | Porphyrin |
| 1976 | UROS | uroporphyrinogen III synthase (congenital erythropoietic porphyria) | 7390 | Porphyrin |
| 1977 | BLVRA | Biliverdin reductase A | 644 | Porphyrin |
| 1978 | BLVRB | biliverdin reductase B (flavin reductase (NADPH)) | 645 | Porphyrin |
| 1979 | HMOX1 | heme oxygenase (decycling) 1 | 3162 | Porphyrin |
| 1980 | HMOX2 | heme oxygenase (decycling) 2 | 3163 | Porphyrin |
| 1981 | ATP13A1 | ATPase type 13A1 | 57130 | Proton Transport |
| 1982 | ATP13A2 | ATPase type 13A2 | 23400 | Proton Transport |
| 1983 | ATP13A3 | ATPase type 13A3 | 79572 | Proton Transport |
| 1984 | ATP13A4 | ATPase type 13A4 | 84239 | Proton Transport |
| 1985 | ATP13A5 | ATPase type 13A5 | 344905 | Proton Transport |
| 1986 | PPA2 | pyrophosphatase (inorganic) 2 | 27068 | Proton Transport |
| 1987 | MFSD3 | Major Facilitator Superfamily Domain Containing 3 | 113655 | Proton Transport |
| 1988 | TAP1 | transporter 1, ATP-binding cassette, sub-family B (MDR/TAP) | 6890 | Proton Transport |
| 1989 | TAP2 | transporter 2, ATP-binding cassette, sub-family B (MDR/TAP) | 6891 | Proton Transport |
| 1990 | ATP5A1 | ATP synthase, H+ transporting, mitochondrial F1 complex, alpha subunit, isoform 1 | 498 | Proton Transport |
| 1991 | ATP5B | ATP synthase, H+ transporting mitochondrial F1 complex, beta subunit | 506 | Proton Transport |
| 1992 | ATP5C1 | ATP synthase, H+ transporting, mitochondrial F1 complex, gamma polypeptide 1 | 509 | Proton Transport |
| 1993 | ATP5C2 | ATP synthase, H+ transporting, mitochondrial F1 complex, gamma polypeptide 2 | 56689 | Proton Transport |
| 1994 | ATP5D | ATP synthase, H+ transporting, mitochondrial F1 complex, delta subunit | 513 | Proton Transport |
| 1995 | ATP5E | ATP synthase, H+ transporting, mitochondrial F1 complex, epsilon subunit | 514 | Proton Transport |
| 1996 | ATP5F1 | ATP synthase, H+ transporting, mitochondrial F0 complex, subunit b, isoform 1 | 515 | Proton Transport |
| 1997 | ATP5G1 | ATP synthase, H+ transporting, mitochondrial F0 complex, subunit c (subunit 9), isoform 1 | 516 | Proton Transport |
| 1998 | ATP5G2 | ATP synthase, H+ transporting, mitochondrial F0 complex, subunit c (subunit 9), isoform 2 | 517 | Proton Transport |
| 1999 | ATP5G3 | ATP synthase, H+ transporting, mitochondrial F0 complex, subunit c (subunit 9), isoform 3 | 518 | Proton Transport |
| 2000 | ATP5H | ATP synthase, H+ transporting, mitochondrial F0 complex, subunit d | 10476 | Proton Transport |
| 2001 | ATP5I | ATP synthase, H+ transporting, mitochondrial F0 complex, subunit E | 521 | Proton Transport |
| 2002 | ATP5J | ATP synthase, H+ transporting, mitochondrial F0 complex, subunit F | 522 | Proton Transport |
| 2003 | ATP5J2 | ATP synthase, H+ transporting, mitochondrial F0 complex, subunit f, isoform 2 | 9551 | Proton Transport |
| 2004 | ATP5L | ATP synthase, H+ transporting, mitochondrial F0 complex, subunit g | 10632 | Proton Transport |
| 2005 | ATP5L2 | ATP synthase, H+ transporting, mitochondrial F0 complex, subunit G2 | 267020 | Proton Transport |
| 2006 | ATP5O | ATP synthase, H+ transporting, mitochondrial F1 complex, O subunit | 539 | Proton Transport |
| 2007 | ATP5S | ATP synthase, H+ transporting, mitochondrial F0 complex, subunit s | 27109 | Proton Transport |
| 2008 | ATP5SL | ATP5S-like | 55101 | Proton Transport |
| 2009 | ATP6 | mitochondrially encoded ATP synthase 6 | 4508 | Proton Transport |
| 2010 | ATP6AP1 | ATPase, H+ transporting, lysosomal accessory protein 1 | 537 | Proton Transport |
| 2011 | ATP6AP2 | ATPase, H+ transporting, lysosomal accessory protein 2 | 10159 | Proton Transport |
| 2012 | ATP6V0A1 | ATPase, H+ transporting, lysosomal V0 subunit A1 | 535 | Proton Transport |
| 2013 | ATP6V0A2 | ATPase, H+ transporting, lysosomal V0 subunit A2 | 23545 | Proton Transport |
| 2014 | ATP6V0A4 | ATPase, H+ transporting, lysosomal V0 subunit A4 | 50617 | Proton Transport |
| 2015 | ATP6V0B | ATPase, H+ transporting, lysosomal V0 subunit B | 533 | Proton Transport |
| 2016 | ATP6V0C | ATPase, H+ transporting, lysosomal V0 subunit C | 527 | Proton Transport |
| 2017 | ATP6V0D1 | ATPase, H+ transporting, lysosomal V0 subunit D1 | 9114 | Proton Transport |
| 2018 | ATP6V0D2 | ATPase, H+ transporting, lysosomal V0 subunit D2 | 245972 | Proton Transport |
| 2019 | ATP6V0E1 | ATPase, H+ transporting, lysosomal V0 subunit E | 8992 | Proton Transport |
| 2020 | ATP6V0E2 | ATPase, H+ transporting, lysosomal V0 subunit E2 | 155066 | Proton Transport |
| 2021 | ATP6V1A | ATPase, H+ transporting, lysosomal V1 subunit A | 523 | Proton Transport |
| 2022 | ATP6V1B1 | ATPase, H+ transporting, lysosomal V1 subunit B1 | 525 | Proton Transport |
| 2023 | ATP6V1B2 | ATPase, H+ transporting, lysosomal V1 subunit B2 | 526 | Proton Transport |
| 2024 | ATP6V1C1 | ATPase, H+ transporting, lysosomal V1 subunit C1 | 528 | Proton Transport |
| 2025 | ATP6V1C2 | ATPase, H+ transporting, lysosomal V1 subunit C2 | 245973 | Proton Transport |
| 2026 | ATP6V1D | ATPase, H+ transporting, lysosomal V1 subunit D | 51382 | Proton Transport |
| 2027 | ATP6V1E1 | ATPase, H+ transporting, lysosomal V1 subunit E1 | 529 | Proton Transport |
| 2028 | ATP6V1E2 | ATPase, H+ transporting, lysosomal V1 subunit E2 | 90423 | Proton Transport |
| 2029 | ATP6V1F | ATPase, H+ transporting, lysosomal V1 subunit F | 9296 | Proton Transport |
| 2030 | ATP6V1G1 | ATPase, H+ transporting, lysosomal V1 subunit G1 | 9550 | Proton Transport |
| 2031 | ATP6V1G2 | ATPase, H+ transporting, lysosomal V1 subunit G2 | 534 | Proton Transport |
| 2032 | ATP6V1G3 | ATPase, H+ transporting, lysosomal V1 subunit G3 | 127124 | Proton Transport |
| 2033 | ATP6V1H | ATPase, H+ transporting, lysosomal V1 subunit H | 51606 | Proton Transport |
| 2034 | HVCN1 | hydrogen voltage-gated channel 1 | 84329 | Proton Transport |
| 2035 | TCIRG1 | T-cell, immune regulator 1, ATPase, H+ transporting, lysosomal V0 subunit A3 | 10312 | Proton Transport |
| 2036 | UCP1 | uncoupling protein 1 (mitochondrial, proton carrier) | 7350 | Proton Transport |
| 2037 | UCP2 | uncoupling protein 2 (mitochondrial, proton carrier) | 7351 | Proton Transport |
| 2038 | UCP3 | uncoupling protein 3 (mitochondrial, proton carrier) | 7352 | Proton Transport |
| 2039 | ASNA1 | ArsA arsenite transporter, ATP-binding, homolog 1 (bacterial) | 439 | Proton Transport |
| 2040 | ATP10A | ATPase, class V, type 10A | 57194 | Proton Transport |
| 2041 | ATP10B | ATPase, class V, type 10B | 23120 | Proton Transport |
| 2042 | ATP10D | ATPase, class V, type 10D | 57205 | Proton Transport |
| 2043 | ATP8 | mitochondrially encoded ATP synthase 8 | 4509 | Proton Transport |
| 2044 | ATP8A1 | ATPase, aminophospholipid transporter (APLT), class I, type 8A, member 1 | 10396 | Proton Transport |
| 2045 | ATP8A2 | ATPase, aminophospholipid transporter-like, class I, type 8A, member 2 | 51761 | Proton Transport |
| 2046 | ATP8B1 | ATPase, class I, type 8B, member 1 | 5205 | Proton Transport |
| 2047 | ATP8B2 | ATPase, class I, type 8B, member 2 | 57198 | Proton Transport |
| 2048 | ATP8B3 | ATPase, class I, type 8B, member 3 | 148229 | Proton Transport |
| 2049 | ATP8B4 | ATPase, class I, type 8B, member 4 | 79895 | Proton Transport |
| 2050 | ATP9A | ATPase, class II, type 9A | 10079 | Proton Transport |
| 2051 | ATP9B | ATPase, class II, type 9B | 374868 | Proton Transport |
| 2052 | SLC25A27 | solute carrier family 25, member 27 | 9481 | Proton Transport |
| 2053 | POR | P450 (cytochrome) oxidoreductase | 5447 | Redox |
| 2054 | ZADH2 | zinc binding alcohol dehydrogenase, domain containing 2 | 284273 | Redox |
| 2055 | CP | ceruloplasmin (ferroxidase) | 1356 | Redox |
| 2056 | CRYZ | crystallin, zeta (quinone reductase) | 1429 | Redox |
| 2057 | CRYZL1 | crystallin, zeta (quinone reductase)-like 1 | 9946 | Redox |
| 2058 | CYB561 | cytochrome b-561 | 1534 | Redox |
| 2059 | CYB5A | cytochrome b5 type A (microsomal) | 1528 | Redox |
| 2060 | CYB5B | cytochrome b5 type B (outer mitochondrial membrane) | 80777 | Redox |
| 2061 | CYB5D1 | cytochrome b5 domain containing 1 | 124637 | Redox |
| 2062 | CYB5D2 | cytochrome b5 domain containing 2 | 124936 | Redox |
| 2063 | CYB5R1 | cytochrome b5 reductase 1 | 51706 | Redox |
| 2064 | CYB5R2 | cytochrome b5 reductase 2 | 51700 | Redox |
| 2065 | CYB5R3 | cytochrome b5 reductase 3 | 1727 | Redox |
| 2066 | CYB5R4 | cytochrome b5 reductase 4 | 51167 | Redox |
| 2067 | CYB5RL | cytochrome b5 reductase like | 606495 | Redox |
| 2068 | CYBA | cytochrome b-245, alpha polypeptide | 1535 | Redox |
| 2069 | CYBASC3 | cytochrome b, ascorbate dependent 3 | 220002 | Redox |
| 2070 | CYBB | cytochrome b-245, beta polypeptide (chronic granulomatous disease) | 1536 | Redox |
| 2071 | CYBRD1 | cytochrome b reductase 1 | 79901 | Redox |
| 2072 | DHRS1 | dehydrogenase/reductase (SDR family) member 1 | 115817 | Redox |
| 2073 | DHRS11 | dehydrogenase/reductase (SDR family) member 11 | 79154 | Redox |
| 2074 | DHRS12 | dehydrogenase/reductase (SDR family) member 12 | 79758 | Redox |
| 2075 | DHRS13 | dehydrogenase/reductase (SDR family) member 13 | 147015 | Redox |
| 2076 | DHRS2 | dehydrogenase/reductase (SDR family) member 2 | 10202 | Redox |
| 2077 | DHRS3 | dehydrogenase/reductase (SDR family) member 3 | 9249 | Redox |
| 2078 | DHRS4 | dehydrogenase/reductase (SDR family) member 4 | 10901 | Redox |
| 2079 | DHRS4L1 | dehydrogenase/reductase (SDR family) member 4 like 1 | 728635 | Redox |
| 2080 | DHRS4L2 | dehydrogenase/reductase (SDR family) member 4 like 2 | 317749 | Redox |
| 2081 | DHRS7 | dehydrogenase/reductase (SDR family) member 7 | 51635 | Redox |
| 2082 | DHRS7B | dehydrogenase/reductase (SDR family) member 7B | 25979 | Redox |
| 2083 | DHRS7C | dehydrogenase/reductase (SDR family) member 7C | 201140 | Redox |
| 2084 | DHRS9 | dehydrogenase/reductase (SDR family) member 9 | 10170 | Redox |
| 2085 | DHRSX | dehydrogenase/reductase (SDR family) X-linked | 207063 | Redox |
| 2086 | DHTKD1 | dehydrogenase E1 and transketolase domain containing 1 | 55526 | Redox |
| 2087 | ENOX1 | ecto-NOX disulfide-thiol exchanger 1 | 55068 | Redox |
| 2088 | ENOX2 | ecto-NOX disulfide-thiol exchanger 2 | 10495 | Redox |
| 2089 | FDX1 | ferredoxin 1 | 2230 | Redox |
| 2090 | FDX1L | ferredoxin 1-like | 112812 | Redox |
| 2091 | FDXR | ferredoxin reductase | 2232 | Redox |
| 2092 | FOXRED1 | FAD-dependent oxidoreductase domain containing 1 | 55572 | Redox |
| 2093 | FOXRED2 | FAD-dependent oxidoreductase domain containing 2 | 80020 | Redox |
| 2094 | FRRS1 | ferric-chelate reductase 1 | 391059 | Redox |
| 2095 | GLRX | glutaredoxin (thioltransferase) | 2745 | Redox |
| 2096 | GLRX2 | glutaredoxin 2 | 51022 | Redox |
| 2097 | GLRX3 | glutaredoxin 3 | 10539 | Redox |
| 2098 | GLRX5 | glutaredoxin 5 | 51218 | Redox |
| 2099 | GRXCR1 | glutaredoxin, cysteine rich 1 | 389207 | Redox |
| 2100 | GRXCR2 | glutaredoxin, cysteine rich 2 | 643226 | Redox |
| 2101 | ISCA1 | iron-sulfur cluster assembly 1 homolog (S. cerevisiae) | 81689 | Redox |
| 2102 | ISCA1L | iron-sulfur cluster assembly 1 homolog (S. cerevisiae)-l | 389293 | Redox |
| 2103 | ISCA2 | iron-sulfur cluster assembly 2 homolog (S. cerevisiae) | 122961 | Redox |
| 2104 | LPO | lactoperoxidase | 4025 | Redox |
| 2105 | MICAL1 | microtubule associated monoxygenase, calponin and LIM domain containing 1 | 64780 | Redox |
| 2106 | MICAL2 | microtubule associated monoxygenase, calponin and LIM domain containing 2 | 9645 | Redox |
| 2107 | MICAL3 | microtubule associated monoxygenase, calponin and LIM domain containing 3 | 57553 | Redox |
| 2108 | NDOR1 | NADPH dependent diflavin oxidoreductase 1 | 27158 | Redox |
| 2109 | NMRAL1 | NmrA-like family domain containing 1 | 57407 | Redox |
| 2110 | OGFOD1 | 2-oxoglutarate and iron-dependent oxygenase domain containing 1 | 55239 | Redox |
| 2111 | OGFOD2 | 2-oxoglutarate and iron-dependent oxygenase domain containing 2 | 79676 | Redox |
| 2112 | OXA1L | oxidase (cytochrome c) assembly 1-like | 5018 | Redox |
| 2113 | PRDX1 | peroxiredoxin 1 | 5052 | Redox |
| 2114 | PRDX2 | peroxiredoxin 2 | 7001 | Redox |
| 2115 | PRDX3 | peroxiredoxin 3 | 10935 | Redox |
| 2116 | PRDX4 | peroxiredoxin 4 | 10549 | Redox |
| 2117 | PRDX5 | peroxiredoxin 5 | 25824 | Redox |
| 2118 | PRDX6 | peroxiredoxin 6 | 9588 | Redox |
| 2119 | PYROXD1 | pyridine nucleotide-disulphide oxidoreductase domain 1 | 79912 | Redox |
| 2120 | TXNRD1 | Thioredoxin reductase 1 | 7296 | Redox |
| 2121 | TXNRD2 | Thioredoxin reductase 2 | 10587 | Redox |
| 2122 | TXNRD3 | Thioredoxin reductase 3 | 114112 | Redox |
| 2123 | DIRC2 | Disrupted in renal carcinoma 2 | 84925 | Small Molecule Transport |
| 2124 | SFXN4 | Sideroflexin 4 | 119559 | Small Molecule Transport |
| 2125 | SVOP | SV2 related protein homolog (rat) | 55530 | Small Molecule Transport |
| 2126 | MFSD11 | Major Facilitator Superfamily Domain Containing 11 | 79157 | Small Molecule Transport |
| 2127 | MFSD7 | Major Facilitator Superfamily Domain Containing 7 | 84179 | Small Molecule Transport |
| 2128 | MFSD9 | Major Facilitator Superfamily Domain Containing 9 | 84804 | Small Molecule Transport |
| 2129 | MFSD10 | Major Facilitator Superfamily Domain Containing 10 | 10227 | Small Molecule Transport (Anion) |
| 2130 | SLC25A34 | solute carrier family 25, member 34 | 284723 | Small Molecule Transport ( Di-Tri-carboxylic acid) |
| 2131 | SLC25A35 | solute carrier family 25, member 35 | 399512 | Small Molecule Transport ( Di-Tri-carboxylic acid) |
| 2132 | SLC25A31 | solute carrier family 25 (mitochondrial carrier; adenine nucleotide translocator), member 31 | 83447 | Small Molecule Transport ( Nucleotide) |
| 2133 | SLC6A14 | solute carrier family 6 (neurotransmitter transporter), member 14 | 11254 | Small Molecule Transport ( β-Alanine) |
| 2134 | SLC33A1 | solute carrier family 33 (acetyl-CoA transporter), member 1 | 9197 | Small Molecule Transport (Acetyl CoA) |
| 2135 | SLC18A3 | solute carrier family 18 (vesicular monoamine), member 3 | 6572 | Small Molecule Transport (Acetylcholine, Choline) |
| 2136 | SLC25A29 | solute carrier family 25 (mitochondrial carrier, palmitoylcarnitine transporter), member 29 | 123096 | Small Molecule Transport (Acylcarnitine) |
| 2137 | SLC25A17 | solute carrier family 25 (mitochondrial carrier, peroxisomal membrane protein), member 17 | 10478 | Small Molecule Transport (ADP/ATP) |
| 2138 | SLC25A21 | solute carrier family 25 (mitochondrial oxodicarboxylate carrier), member 21 | 89874 | Small Molecule Transport (aKG) |
| 2139 | SLC6A18 | solute carrier family 6 (neurotransmitter transporter), member 18 | 348932 | Small Molecule Transport (Ala, Gly, Meth, Phe) |
| 2140 | SLC1A4 | solute carrier family 1 (glutamate/neutral amino acid transporter), member 4 | 6509 | Small Molecule Transport (Alanine, Cysteine, Serine) |
| 2141 | SLC1A5 | solute carrier family 1 (neutral amino acid transporter), member 5 | 6510 | Small Molecule Transport (Alanine, Cysteine, Serine) |
| 2142 | TMEM104 | Transmembrane protein 104 | 54868 | Small Molecule Transport (Amino Acid) |
| 2143 | SLC25A38 | solute carrier family 25, member 38 | 54977 | Small Molecule Transport (Amino acid) |
| 2144 | SLC25A39 | solute carrier family 25, member 39 | 51629 | Small Molecule Transport (Amino acid) |
| 2145 | SLC25A40 | solute carrier family 25, member 40 | 55972 | Small Molecule Transport (Amino acid) |
| 2146 | SLC25A44 | solute carrier family 25, member 44 | 9673 | Small Molecule Transport (Amino acid) |
| 2147 | SLC25A45 | solute carrier family 25, member 45 | 283130 | Small Molecule Transport (Amino acid) |
| 2148 | SLC38A6 | solute carrier family 38, member 6 | 145389 | Small Molecule Transport (Amino acid) |
| 2149 | SLC38A7 | solute carrier family 38, member 7 | 55238 | Small Molecule Transport (Amino acid) |
| 2150 | SLC43A3 | solute carrier family 43, member 3 | 29015 | Small Molecule Transport (Amino acid) |
| 2151 | SLC6A16 | solute carrier family 6 (neurotransmitter transporter), member 16 | 28968 | Small Molecule Transport (Amino acid) |
| 2152 | SLC7A10 | solute carrier family 7 (cationic amino acid transporter, y+ system), member 10 | 56301 | Small Molecule Transport (Amino acid) |
| 2153 | SLC7A14 | solute carrier family 7 (cationic amino acid transporter, y+ system), member 14 | 57709 | Small Molecule Transport (Amino acid) |
| 2154 | SLC7A4 | solute carrier family 7 (cationic amino acid transporter, y+ system), member 4 | 6545 | Small Molecule Transport (Amino acid) |
| 2155 | SLC7A6 | solute carrier family 7 (cationic amino acid transporter, y+ system), member 6 | 9057 | Small Molecule Transport (Amino acid) |
| 2156 | SLC7A7 | solute carrier family 7 (cationic amino acid transporter, y+ system), member 7 | 9056 | Small Molecule Transport (Amino acid) |
| 2157 | SLC7A8 | solute carrier family 7 (cationic amino acid transporter, y+ system), member 8 | 23428 | Small Molecule Transport (Amino acid) |
| 2158 | SLC7A9 | solute carrier family 7 (cationic amino acid transporter, y+ system), member 9 | 11136 | Small Molecule Transport (Amino acid) |
| 2159 | RHAG | Rh-associated glycoprotein | 6005 | Small Molecule Transport (Ammonium) |
| 2160 | RHBG | Rh family, B glycoprotein | 57127 | Small Molecule Transport (Ammonium) |
| 2161 | RHCG | Rh family, C glycoprotein | 51458 | Small Molecule Transport (Ammonium) |
| 2162 | SLC4A1AP | solute carrier family 4 (anion exchanger), member 1, adaptor protein | 22950 | Small Molecule Transport (Anion) |
| 2163 | SLC22A6 | solute carrier family 22 (organic anion transporter), member 6 | 9356 | Small Molecule Transport (Anion) |
| 2164 | SLC22A7 | solute carrier family 22 (organic anion transporter), member 7 | 10864 | Small Molecule Transport (Anion) |
| 2165 | SLC22A9 | solute carrier family 22 (organic anion transporter), member 9 | 114571 | Small Molecule Transport (Anion) |
| 2166 | SLC4A1 | solute carrier family 4 (anion exchanger), member 1 | 6521 | Small Molecule Transport (Anion) |
| 2167 | SLC4A3 | solute carrier family 4 (anion exchanger), member 3 | 6508 | Small Molecule Transport (Anion) |
| 2168 | SLC4A9 | solute carrier family 4, sodium bicarbonate cotransporter, member 9 | 83697 | Small Molecule Transport (Anion) |
| 2169 | SLC7A1 | solute carrier family 7 (cationic amino acid transporter, y+ system), member 1 | 6541 | Small Molecule Transport (Ar, Lys, Orn, His) |
| 2170 | SLC25A2 | solute carrier family 25 (mitochondrial carrier, ornithine transporter) member 2 | 83884 | Small Molecule Transport (Arg, Citrulline,lys) |
| 2171 | SLC25A15 | solute carrier family 25 (mitochondrial carrier ornithine transporter), member 15 | 10166 | Small Molecule Transport (Arg, Citrulline,lys, Orn) |
| 2172 | SLC7A3 | solute carrier family 7 (cationic amino acid transporter, y+ system), member 3 | 84889 | Small Molecule Transport (Arg, Lys, Orn) |
| 2173 | SLC7A2 | solute carrier family 7 (cationic amino acid transporter, y+ system), member 2 | 6542 | Small Molecule Transport (Arg, Lys, Orn, His) |
| 2174 | SLC38A5 | solute carrier family 38, member 5 | 92745 | Small Molecule Transport (Asn, Se, His, Gln) |
| 2175 | SLC7A13 | solute carrier family 7, (cationic amino acid transporter, y+ system) member 13 | 157724 | Small Molecule Transport (Aspartate, Glutamate) |
| 2176 | SLC43A1 | solute carrier family 43, member 1 | 8501 | Small Molecule Transport (BCAA) |
| 2177 | SLC43A2 | solute carrier family 43, member 2 | 124935 | Small Molecule Transport (BCAA) |
| 2178 | SLC6A19 | solute carrier family 6 (neurotransmitter transporter), member 19 | 340024 | Small Molecule Transport (BCAA) |
| 2179 | OSTalpha | organic solute transporter alpha | 200931 | Small Molecule Transport (Bile Acid) |
| 2180 | OSTbeta | organic solute transporter beta | 123264 | Small Molecule Transport (Bile Acid) |
| 2181 | SLC10A1 | solute carrier family 10 (sodium/bile acid cotransporter family), member 1 | 6554 | Small Molecule Transport (Bile acid) |
| 2182 | SLC10A2 | solute carrier family 10, member 2 | 6555 | Small Molecule Transport (Bile acid) |
| 2183 | SLC10A3 | solute carrier family 10 (sodium/bile acid cotransporter family), member 3 | 8273 | Small Molecule Transport (Bile acid) |
| 2184 | SLC10A4 | solute carrier family 10 (sodium/bile acid cotransporter family), member 4 | 201780 | Small Molecule Transport (Bile acid) |
| 2185 | SLC10A5 | solute carrier family 10 (sodium/bile acid cotransporter family), member 5 | 347051 | Small Molecule Transport (Bile acid) |
| 2186 | SLC10A6 | solute carrier family 10 (sodium/bile acid cotransporter family), member 6 | 345274 | Small Molecule Transport (Bile acid) |
| 2187 | SLC10A7 | solute carrier family 10 (sodium/bile acid cotransporter family), member 7 | 84068 | Small Molecule Transport (Bile acid) |
| 2188 | SLCO1A2 | solute carrier organic anion transporter family, member 1a2 | 6579 | Small Molecule Transport (Bile acid) |
| 2189 | SLCO1B1 | solute carrier organic anion transporter family, member 1b1 | 10599 | Small Molecule Transport (Bile acid) |
| 2190 | SLCO1B3 | solute carrier organic anion transporter family, member 1b3 | 28234 | Small Molecule Transport (Bile acid) |
| 2191 | SLCO4A1 | solute carrier organic anion transporter family, member 4a1 | 28231 | Small Molecule Transport (bile acids,prostaglandins,steroid conjugates,thyroid hormone) |
| 2192 | SLC26A10 | solute carrier family 26, member 10 | 65012 | Small Molecule Transport (Biocarbonate) |
| 2193 | SLC26A3 | solute carrier family 26, member 3 | 1811 | Small Molecule Transport (Biocarbonate) |
| 2194 | SLC26A4 | solute carrier family 26, member 4 | 5172 | Small Molecule Transport (Biocarbonate) |
| 2195 | SLC26A5 | solute carrier family 26, member 5 | 375611 | Small Molecule Transport (Biocarbonate) |
| 2196 | SLC26A6 | solute carrier family 26, member 6 | 65010 | Small Molecule Transport (Biocarbonate) |
| 2197 | SLC26A7 | solute carrier family 26, member 7 | 115111 | Small Molecule Transport (Biocarbonate) |
| 2198 | SLC26A8 | solute carrier family 26, member 8 | 116369 | Small Molecule Transport (Biocarbonate) |
| 2199 | SLC26A9 | solute carrier family 26, member 9 | 115019 | Small Molecule Transport (Biocarbonate) |
| 2200 | SLC4A10 | solute carrier family 4, sodium bicarbonate cotransporter-like, member 10 | 57282 | Small Molecule Transport (Biocarbonate) |
| 2201 | SLC4A11 | solute carrier family 4, member 11 | 83959 | Small Molecule Transport (Biocarbonate) |
| 2202 | SLC4A2 | solute carrier family 4 (anion exchanger), member 2 | 6522 | Small Molecule Transport (Biocarbonate) |
| 2203 | SLC4A4 | solute carrier family 4 (anion exchanger), member 4 | 8671 | Small Molecule Transport (Biocarbonate) |
| 2204 | SLC4A5 | solute carrier family 4 (anion exchanger), member 5 | 57835 | Small Molecule Transport (Biocarbonate) |
| 2205 | SLC4A7 | solute carrier family 4 (anion exchanger), member 7 | 9497 | Small Molecule Transport (Biocarbonate) |
| 2206 | SLC4A8 | solute carrier family 4 (anion exchanger), member 8 | 9498 | Small Molecule Transport (Biocarbonate) |
| 2207 | SLCO4C1 | solute carrier organic anion transporter family, member 4C1 | 353189 | Small Molecule Transport (cAMP,steroid conjugates,thyroid hormones) |
| 2208 | SLC25A20 | solute carrier family 25 (mitochondrial carnitine/acylcarnitine translocase), member 20 | 788 | Small Molecule Transport (Carnitine) |
| 2209 | SLC22A16 | solute carrier family 22 (organic cation transporter), member 16 | 85413 | Small Molecule Transport (Carnitine) |
| 2210 | SLC22A4 | solute carrier family 22 (organic cation transporter), member 4 | 6583 | Small Molecule Transport (Carnitine) |
| 2211 | SLC22A5 | solute carrier family 22 (organic cation transporter), member 5 | 6584 | Small Molecule Transport (Carnitine) |
| 2212 | NPC1 | Niemann-Pick disease, type C1 | 4864 | Small Molecule Transport (Cholesterol) |
| 2213 | NPC1L1 | NPC1 (Niemann-Pick disease, type C1, gene)-like 1 | 29881 | Small Molecule Transport (Cholesterol) |
| 2214 | NPC2 | Niemann-Pick disease, type C2 | 10577 | Small Molecule Transport (Cholesterol) |
| 2215 | STAR | steroidogenic acute regulatory protein | 6770 | Small Molecule Transport (Cholesterol) |
| 2216 | STARD3 | StAR-related lipid transfer (START) domain containing 3 | 10948 | Small Molecule Transport (Cholesterol) |
| 2217 | STARD3NL | STARD3 N-terminal like | 83930 | Small Molecule Transport (Cholesterol) |
| 2218 | STARD4 | StAR-related lipid transfer (START) domain containing 4 | 134429 | Small Molecule Transport (Cholesterol) |
| 2219 | STARD5 | StAR-related lipid transfer (START) domain containing 5 | 80765 | Small Molecule Transport (Cholesterol) |
| 2220 | STARD6 | StAR-related lipid transfer (START) domain containing 6 | 147323 | Small Molecule Transport (Cholesterol) |
| 2221 | SLC44A1 | solute carrier family 44, member 1 | 23446 | Small Molecule Transport (Choline) |
| 2222 | SLC44A2 | solute carrier family 44, member 2 | 57153 | Small Molecule Transport (Choline) |
| 2223 | SLC44A3 | solute carrier family 44, member 3 | 126969 | Small Molecule Transport (Choline) |
| 2224 | SLC44A4 | solute carrier family 44, member 4 | 80736 | Small Molecule Transport (Choline) |
| 2225 | SLC44A5 | solute carrier family 44, member 5 | 204962 | Small Molecule Transport (Choline) |
| 2226 | SLC5A7 | solute carrier family 5 (choline transporter), member 7 | 60482 | Small Molecule Transport (Choline) |
| 2227 | SFXN5 | Sideroflexin 5 | 94097 | Small Molecule Transport (Citrate) |
| 2228 | SLC23A3 | solute carrier family 23 (nucleobase transporters), member 3 | 151295 | Small Molecule Transport (Citrate, Malate) |
| 2229 | SLC25A1 | solute carrier family 25 (mitochondrial carrier, citrate transporter), member 1 | 6576 | Small Molecule Transport (Citrate, Malate) |
| 2230 | SLC13A5 | solute carrier family 13 (sodium/sulfate symporters), member 5 | 284111 | Small Molecule Transport (Citrate, Pyruvate) |
| 2231 | SLC13A2 | solute carrier family 13 (sodium-dependent dicarboxylate transporter), member 2 | 9058 | Small Molecule Transport (Citrate, Succinate) |
| 2232 | SLC13A3 | solute carrier family 13 (sodium-dependent dicarboxylate transporter), member 3 | 64849 | Small Molecule Transport (Citrate, Succinate) |
| 2233 | SLC25A16 | solute carrier family 25 (mitochondrial carrier, Graves disease autoantigen), member 16 | 8034 | Small Molecule Transport (CoA) |
| 2234 | SLC16A12 | solute carrier family 16 (monocarboxylic acid transporters), member 12 | 387700 | Small Molecule Transport (Creatine) |
| 2235 | SLC6A8 | solute carrier family 6 (neurotransmitter transporter, creatine), member 8 | 6535 | Small Molecule Transport (Creatine) |
| 2236 | CTNS | Cystinosis, nephropathic | 1497 | Small Molecule Transport (Cystine) |
| 2237 | SLC7A11 | solute carrier family 7 (cationic amino acid transporter, y+ system), member 11 | 23657 | Small Molecule Transport (Cystine, Glutamate) |
| 2238 | SLC25A19 | solute carrier family 25 (mitochondrial deoxynucleotide carrier), member 19 | 60386 | Small Molecule Transport (dNTP, dNDPs) |
| 2239 | SLC22A2 | solute carrier family 22 (organic cation transporter), member 2 | 6582 | Small Molecule Transport (dopamine  histamine,PGE2) |
| 2240 | SLC29A4 | solute carrier family 29 (nucleoside transporters), member 4 | 222962 | Small Molecule Transport (Dopamine) |
| 2241 | SLCO2A1 | solute carrier organic anion transporter family, member 2a1 | 6578 | Small Molecule Transport (eicosanoids,prostaglandins) |
| 2242 | MFSD2 | Major Facilitator Superfamily Domain Containing 2 | 84879 | Small Molecule Transport (Fatty acid) |
| 2243 | SLC27A1 | solute carrier family 27 (fatty acid transporter), member 1 | 376497 | Small Molecule Transport (Fatty acid) |
| 2244 | SLC27A2 | solute carrier family 27 (fatty acid transporter), member 2 | 11001 | Small Molecule Transport (Fatty acid) |
| 2245 | SLC27A3 | solute carrier family 27 (fatty acid transporter), member 3 | 11000 | Small Molecule Transport (Fatty acid) |
| 2246 | SLC27A4 | solute carrier family 27 (fatty acid transporter), member 4 | 10999 | Small Molecule Transport (Fatty acid) |
| 2247 | SLC27A5 | solute carrier family 27 (fatty acid transporter), member 5 | 10998 | Small Molecule Transport (Fatty acid) |
| 2248 | SLC27A6 | solute carrier family 27 (fatty acid transporter), member 6 | 28965 | Small Molecule Transport (Fatty acid) |
| 2249 | FOLR1 | folate receptor 1 (adult) | 2348 | Small Molecule Transport (Folate) |
| 2250 | FOLR2 | folate receptor 2 (fetal) | 2350 | Small Molecule Transport (Folate) |
| 2251 | FOLR3 | folate receptor 3 (gamma) | 2352 | Small Molecule Transport (Folate) |
| 2252 | SLC46A1 | Solute carrier family 46 (folate transporter), member 1 | 113235 | Small Molecule Transport (Folate) |
| 2253 | SLC46A2 | solute carrier family 46, member 2 | 57864 | Small Molecule Transport (Folate) |
| 2254 | SLC46A3 | solute carrier family 46, member 3 | 283537 | Small Molecule Transport (Folate) |
| 2255 | SLC37A2 | solute carrier family 37 (glycerol-3-phosphate transporter), member 2 | 219855 | Small Molecule Transport (G6P) |
| 2256 | SLC37A4 | solute carrier family 37 (glycerol-6-phosphate transporter), member 4 | 2542 | Small Molecule Transport (G6P) |
| 2257 | SLC37A1 | solute carrier family 37 (glycerol-3-phosphate transporter), member 1 | 54020 | Small Molecule Transport (G6P, G3P) |
| 2258 | SLC6A1 | solute carrier family 6 (neurotransmitter transporter, GABA), member 1 | 6529 | Small Molecule Transport (GABA) |
| 2259 | SLC6A6 | solute carrier family 6 (neurotransmitter transporter, taurine), member 6 | 6533 | Small Molecule Transport (GABA,  β-alanine,taurine) |
| 2260 | SLC6A12 | solute carrier family 6 (neurotransmitter transporter, betaine/GABA), member 12 | 6539 | Small Molecule Transport (GABA, Betaine |
| 2261 | SLC32A1 | solute carrier family 32 (GABA vesicular transporter), member 1 | 140679 | Small Molecule Transport (GABA, Gly) |
| 2262 | SLC6A11 | solute carrier family 6 (neurotransmitter transporter, GABA), member 11 | 6538 | Small Molecule Transport (GABA, β-Alanine) |
| 2263 | SLC6A13 | solute carrier family 6 (neurotransmitter transporter, GABA), member 13 | 6540 | Small Molecule Transport (GABA, β-Alanine) |
| 2264 | SLC35C1 | solute carrier family 35, member C1 | 55343 | Small Molecule Transport (GDP-Fructose) |
| 2265 | SLC38A1 | solute carrier family 38, member 1 | 81539 | Small Molecule Transport (Gln, Ala, Ser, Asn) |
| 2266 | SLC2A10 | solute carrier family 2 (facilitated glucose transporter), member 10 | 81031 | Small Molecule Transport (Glucose) |
| 2267 | SLC2A12 | solute carrier family 2 (facilitated glucose transporter), member 12 | 154091 | Small Molecule Transport (Glucose) |
| 2268 | SLC2A14 | Solute carrier family 2 (facilitated glucose transporter), member 14 | 144195 | Small Molecule Transport (Glucose) |
| 2269 | SLC2A3 | solute carrier family 2 (facilitated glucose transporter), member 3 | 6515 | Small Molecule Transport (Glucose) |
| 2270 | SLC2A6 | solute carrier family 2 (facilitated glucose transporter), member 6 | 11182 | Small Molecule Transport (Glucose) |
| 2271 | SLC2A8 | solute carrier family 2, (facilitated glucose transporter), member 8 | 29988 | Small Molecule Transport (Glucose) |
| 2272 | SLC45A1 | solute carrier family 45, member 1 | 50651 | Small Molecule Transport (Glucose) |
| 2273 | SLC5A1 | solute carrier family 5 (sodium/glucose cotransporter), member 1 | 6523 | Small Molecule Transport (Glucose) |
| 2274 | SLC5A10 | solute carrier family 5 (sodium/glucose cotransporter), member 10 | 125206 | Small Molecule Transport (Glucose) |
| 2275 | SLC5A2 | solute carrier family 5 (sodium/glucose cotransporter), member 2 | 6524 | Small Molecule Transport (Glucose) |
| 2276 | SLC5A4 | solute carrier family 5, member 4a | 6527 | Small Molecule Transport (Glucose) |
| 2277 | SLC5A9 | solute carrier family 5 (sodium/glucose cotransporter), member 9 | 200010 | Small Molecule Transport (Glucose) |
| 2278 | MFSD4 | Major Facilitator Superfamily Domain Containing 4 | 148808 | Small Molecule Transport (Glucose) |
| 2279 | SLC2A11 | solute carrier family 2 (facilitated glucose transporter), member 11 | 66035 | Small Molecule Transport (Glucose, Fructose) |
| 2280 | SLC2A5 | solute carrier family 2 (facilitated glucose transporter), member 5 | 6518 | Small Molecule Transport (Glucose, Fructose) |
| 2281 | SLC2A7 | solute carrier family 2 (facilitated glucose transporter), member 7 | 155184 | Small Molecule Transport (Glucose, Fructose) |
| 2282 | SLC2A1 | solute carrier family 2 (facilitated glucose transporter), member 1 | 6513 | Small Molecule Transport (Glucose, Glucosamine) |
| 2283 | SLC2A2 | solute carrier family 2 (facilitated glucose transporter), member 2 | 6514 | Small Molecule Transport (Glucose, Glucosamine) |
| 2284 | SLC2A4 | solute carrier family 2 (facilitated glucose transporter), member 4 | 6517 | Small Molecule Transport (Glucose, Glucosamine) |
| 2285 | SLC25A18 | solute carrier family 25 (mitochondrial carrier), member 18 | 83733 | Small Molecule Transport (Glutamate) |
| 2286 | SLC25A22 | solute carrier family 25 (mitochondrial carrier, glutamate), member 22 | 79751 | Small Molecule Transport (Glutamate) |
| 2287 | SLC17A6 | solute carrier family 17 (sodium-dependent inorganic phosphate cotransporter), member 6 | 57084 | Small Molecule Transport (Glutamate) |
| 2288 | SLC17A7 | solute carrier family 17 (sodium-dependent inorganic phosphate cotransporter), member 7 | 57030 | Small Molecule Transport (Glutamate) |
| 2289 | SLC17A8 | solute carrier family 17 (sodium-dependent inorganic phosphate cotransporter), member 8 | 246213 | Small Molecule Transport (Glutamate) |
| 2290 | SLC1A6 | solute carrier family 1 (high affinity aspartate/glutamate transporter), member 6 | 6511 | Small Molecule Transport (Glutamate, Asparate) |
| 2291 | SLC1A7 | solute carrier family 1 (glutamate transporter), member 7 | 6512 | Small Molecule Transport (Glutamate, Asparate) |
| 2292 | SLC1A1 | solute carrier family 1 (neuronal/epithelial high affinity glutamate transporter, system Xag), member 1 | 6505 | Small Molecule Transport (Glutamate, Asparate) |
| 2293 | SLC1A2 | solute carrier family 1 (glial high affinity glutamate transporter), member 2 | 6506 | Small Molecule Transport (Glutamate, Asparate) |
| 2294 | SLC1A3 | solute carrier family 1 (glial high affinity glutamate transporter), member 3 | 6507 | Small Molecule Transport (Glutamate, Asparate) |
| 2295 | SLC17A5 | solute carrier family 17 (anion/sugar transporter), member 5 | 26503 | Small Molecule Transport (Glutamate, Asparate, Lactate) |
| 2296 | SLC25A12 | solute carrier family 25 (mitochondrial carrier, Aralar), member 12 | 8604 | Small Molecule Transport (Glutamate, Aspartate) |
| 2297 | SLC25A13 | solute carrier family 25 (mitochondrial carrier, adenine nucleotide translocator), member 13 | 10165 | Small Molecule Transport (Glutamate, Aspartate) |
| 2298 | SLC36A1 | solute carrier family 36 (proton/amino acid symporter), member 1 | 206358 | Small Molecule Transport (Glycine) |
| 2299 | SLC36A2 | solute carrier family 36 (proton/amino acid symporter), member 2 | 153201 | Small Molecule Transport (Glycine) |
| 2300 | SLC36A3 | solute carrier family 36 (proton/amino acid symporter), member 3 | 285641 | Small Molecule Transport (Glycine) |
| 2301 | SLC6A5 | solute carrier family 6 (neurotransmitter transporter, glycine), member 5 | 9152 | Small Molecule Transport (Glycine) |
| 2302 | SLC6A9 | solute carrier family 6 (neurotransmitter transporter, glycine), member 9 | 6536 | Small Molecule Transport (Glycine, Sarcosine) |
| 2303 | FLVCR1 | Feline leukemia virus subgroup C cellular receptor 1 | 28982 | Small Molecule Transport (Hem) |
| 2304 | FLVCR2 | Feline leukemia virus subgroup C cellular receptor family, member 2 | 55640 | Small Molecule Transport (Hem) |
| 2305 | SLC38A4 | solute carrier family 38, member 4 | 55089 | Small Molecule Transport (His, Ar, Alan, Asn) |
| 2306 | SLC38A3 | solute carrier family 38, member 3 | 10991 | Small Molecule Transport (His, Gln, Asn, Ala, Glutamate) |
| 2307 | SLC28A2 | solute carrier family 28 (sodium-coupled nucleoside transporter), member 2 | 9153 | Small Molecule Transport (inosine) |
| 2308 | SLC28A3 | solute carrier family 28 (sodium-coupled nucleoside transporter), member 3 | 64078 | Small Molecule Transport (inosine) |
| 2309 | SLC29A1 | solute carrier family 29 (nucleoside transporters), member 1 | 2030 | Small Molecule Transport (inosine) |
| 2310 | SLC29A2 | solute carrier family 29 (nucleoside transporters), member 2 | 3177 | Small Molecule Transport (inosine) |
| 2311 | SLC29A3 | solute carrier family 29 (nucleoside transporters), member 3 | 55315 | Small Molecule Transport (inosine) |
| 2312 | SLC5A5 | solute carrier family 5 (sodium iodide symporter), member 5 | 6528 | Small Molecule Transport (iodide) |
| 2313 | SLC5A8 | solute carrier family 5 (iodide transporter), member 8 | 160728 | Small Molecule Transport (iodide) |
| 2314 | SLC16A8 | solute carrier family 16 (monocarboxylic acid transporters), member 8 | 23539 | Small Molecule Transport (Lactate) |
| 2315 | SLC16A3 | solute carrier family 16 (monocarboxylic acid transporters), member 3 | 9123 | Small Molecule Transport (Lactate/Pyruvate) |
| 2316 | SLC16A7 | solute carrier family 16 (monocarboxylic acid transporters), member 7 | 9194 | Small Molecule Transport (Lactate/Pyruvate) |
| 2317 | SLC16A1 | solute carrier family 16 (monocarboxylic acid transporters), member 1 | 6566 | Small Molecule Transport (Lactate/Pyruvate/3-HB) |
| 2318 | STARD13 | StAR-related lipid transfer (START) domain containing 13 | 90627 | Small Molecule Transport (Lipid) |
| 2319 | STARD7 | StAR-related lipid transfer (START) domain containing 7 | 56910 | Small Molecule Transport (Lipid) |
| 2320 | STARD8 | StAR-related lipid transfer (START) domain containing 8 | 9754 | Small Molecule Transport (Lipid) |
| 2321 | STARD9 | StAR-related lipid transfer (START) domain containing 9 | 57519 | Small Molecule Transport (Lipid) |
| 2322 | SLC25A11 | solute carrier family 25 (mitochondrial carrier oxoglutarate carrier), member 11 | 8402 | Small Molecule Transport (Malate, aKG) |
| 2323 | SLC25A10 | solute carrier family 25 (mitochondrial carrier, dicarboxylate transporter), member 10 | 1468 | Small Molecule Transport (Malate, Succinate) |
| 2324 | SLC16A13 | solute carrier family 16 (monocarboxylic acid transporters), member 13 | 201232 | Small Molecule Transport (MCT activity) |
| 2325 | SLC16A4 | solute carrier family 16 (monocarboxylic acid transporters), member 4 | 9122 | Small Molecule Transport (MCT activity) |
| 2326 | SLC16A5 | solute carrier family 16 (monocarboxylic acid transporters), member 5 | 9121 | Small Molecule Transport (MCT activity) |
| 2327 | SLC16A6 | solute carrier family 16 (monocarboxylic acid transporters), member 6 | 9120 | Small Molecule Transport (MCT activity) |
| 2328 | SLC16A9 | solute carrier family 16 (monocarboxylic acid transporters), member 9 | 220963 | Small Molecule Transport (MCT activity) |
| 2329 | SLC25A14 | solute carrier family 25 (mitochondrial carrier, brain), member 14 | 9016 | Small Molecule Transport (MCT activity) |
| 2330 | SLC6A15 | solute carrier family 6 (neurotransmitter transporter), member 15 | 55117 | Small Molecule Transport (Met, BCAA) |
| 2331 | SLC38A2 | solute carrier family 38, member 2 | 54407 | Small Molecule Transport (Met, Gln, Ala, Ser, Asn) |
| 2332 | MCART1 | Mitochondrial carrier triple repeat 1 | 92014 | Small Molecule Transport (Mitochondrial carrier) |
| 2333 | MCART2 | Mitochondrial carrier triple repeat 2 | 147407 | Small Molecule Transport (Mitochondrial carrier) |
| 2334 | MCART6 | Mitochondrial carrier triple repeat 6 | 401612 | Small Molecule Transport (Mitochondrial carrier) |
| 2335 | SLC25A23 | solute carrier family 25 (mitochondrial carrier; phosphate carrier), member 23 | 79085 | Small Molecule Transport (Mitochondrial Phosphate) |
| 2336 | SLC25A24 | solute carrier family 25 (mitochondrial carrier, phosphate carrier), member 24 | 29957 | Small Molecule Transport (Mitochondrial Phosphate) |
| 2337 | SLC25A25 | solute carrier family 25 (mitochondrial carrier, phosphate carrier), member 25 | 114789 | Small Molecule Transport (Mitochondrial Phosphate) |
| 2338 | SLC25A3 | solute carrier family 25 (mitochondrial carrier, phosphate carrier), member 3 | 5250 | Small Molecule Transport (Mitochondrial Phosphate) |
| 2339 | SLC34A1 | solute carrier family 34 (sodium phosphate), member 1 | 6569 | Small Molecule Transport (Mitochondrial Phosphate) |
| 2340 | SLC34A2 | solute carrier family 34 (sodium phosphate), member 2 | 10568 | Small Molecule Transport (Mitochondrial Phosphate) |
| 2341 | SLC34A3 | solute carrier family 34 (sodium phosphate), member 3 | 142680 | Small Molecule Transport (Mitochondrial Phosphate) |
| 2342 | SLC25A30 | solute carrier family 25, member 30 | 253512 | Small Molecule Transport (mitochondrial transport) |
| 2343 | SLC25A37 | solute carrier family 25, member 37 | 51312 | Small Molecule Transport (mitochondrial transport) |
| 2344 | SLC25A46 | solute carrier family 25, member 46 | 91137 | Small Molecule Transport (mitochondrial transport) |
| 2345 | SLC25A28 | solute carrier family 25, member 28 | 81894 | Small Molecule Transport (Mitohondrial Iron) |
| 2346 | SLC18A1 | solute carrier family 18 (vesicular monoamine), member 1 | 6570 | Small Molecule Transport (Monoamine) |
| 2347 | SLC18A2 | solute carrier family 18 (vesicular monoamine), member 2 | 6571 | Small Molecule Transport (Monoamine) |
| 2348 | SLC22A3 | solute carrier family 22 (organic cation transporter), member 3 | 6581 | Small Molecule Transport (Monoamine) |
| 2349 | SLC6A2 | solute carrier family 6 (neurotransmitter transporter, noradrenalin), member 2 | 6530 | Small Molecule Transport (Monoamine) |
| 2350 | SLC6A3 | solute carrier family 6 (neurotransmitter transporter, dopamine), member 3 | 6531 | Small Molecule Transport (Monoamine) |
| 2351 | SLC6A4 | solute carrier family 6 (neurotransmitter transporter, serotonin), member 4 | 6532 | Small Molecule Transport (Monoamine) |
| 2352 | SLC2A13 | solute carrier family 2 (facilitated glucose transporter), member 13 | 114134 | Small Molecule Transport (Myo-Inoisitol) |
| 2353 | SLC22A13 | solute carrier family 22 (organic cation transporter), member 13 | 9390 | Small Molecule Transport (Nicotinic acid) |
| 2354 | SLC5A12 | solute carrier family 5 (sodium/glucose cotransporter), member 12 | 159963 | Small Molecule Transport (Nicotinic acid, Lactate, Pyruvate) |
| 2355 | SLC35A1 | solute carrier family 35 (CMP-sialic acid transporter), member 1 | 10559 | Small Molecule Transport (Nucleotide sugar) |
| 2356 | SLC35A2 | solute carrier family 35 (UDP-galactose transporter), member 2 | 7355 | Small Molecule Transport (Nucleotide sugar) |
| 2357 | SLC35A5 | solute carrier family 35, member A5 | 55032 | Small Molecule Transport (Nucleotide sugar) |
| 2358 | SLC35B1 | solute carrier family 35, member B1 | 10237 | Small Molecule Transport (Nucleotide sugar) |
| 2359 | SLC35B2 | solute carrier family 35, member B2 | 347734 | Small Molecule Transport (Nucleotide sugar) |
| 2360 | SLC35B3 | solute carrier family 35, member B3 | 51000 | Small Molecule Transport (Nucleotide sugar) |
| 2361 | SLC35C2 | solute carrier family 35, member C2 | 51006 | Small Molecule Transport (Nucleotide sugar) |
| 2362 | SLC35D2 | solute carrier family 35 (UDP-glucuronic acid/UDP-N-acetylgalactosamine dual transporter), member D2 | 11046 | Small Molecule Transport (Nucleotide sugar) |
| 2363 | SLC35D3 | solute carrier family 35, member D3 | 340146 | Small Molecule Transport (Nucleotide sugar) |
| 2364 | SLC35E1 | solute carrier family 35, member E1 | 79939 | Small Molecule Transport (Nucleotide sugar) |
| 2365 | SLC35E2 | solute carrier family 35, member E2 | 9906 | Small Molecule Transport (Nucleotide sugar) |
| 2366 | SLC35E3 | solute carrier family 35, member E3 | 55508 | Small Molecule Transport (Nucleotide sugar) |
| 2367 | SLC35E4 | solute carrier family 35, member E4 | 339665 | Small Molecule Transport (Nucleotide sugar) |
| 2368 | SLC35F1 | solute carrier family 35, member F1 | 222553 | Small Molecule Transport (Nucleotide sugar) |
| 2369 | SLC35F2 | solute carrier family 35, member F2 | 54733 | Small Molecule Transport (Nucleotide sugar) |
| 2370 | SLC35F3 | solute carrier family 35, member F3 | 148641 | Small Molecule Transport (Nucleotide sugar) |
| 2371 | SLC35F4 | solute carrier family 35, member F4 | 341880 | Small Molecule Transport (Nucleotide sugar) |
| 2372 | SLC35F5 | solute carrier family 35, member F5 | 80255 | Small Molecule Transport (Nucleotide sugar) |
| 2373 | SLC25A32 | solute carrier family 25, member 32 | 81034 | Small Molecule Transport (Nucleotide) |
| 2374 | SLC25A33 | solute carrier family 25, member 33 | 84275 | Small Molecule Transport (Nucleotide) |
| 2375 | SLC25A36 | solute carrier family 25, member 36 | 55186 | Small Molecule Transport (Nucleotide) |
| 2376 | SLC25A4 | solute carrier family 25 (mitochondrial carrier, adenine nucleotide translocator), member 4 | 291 | Small Molecule Transport (Nucleotide) |
| 2377 | SLC25A41 | solute carrier family 25, member 41 | 284427 | Small Molecule Transport (Nucleotide) |
| 2378 | SLC25A42 | solute carrier family 25, member 42 | 284439 | Small Molecule Transport (Nucleotide) |
| 2379 | SLC25A43 | solute carrier family 25, member 43 | 203427 | Small Molecule Transport (Nucleotide) |
| 2380 | SLC25A5 | solute carrier family 25 (mitochondrial carrier, adenine nucleotide translocator), member 5 | 292 | Small Molecule Transport (Nucleotide) |
| 2381 | SLC25A6 | solute carrier family 25 (mitochondrial carrier; adenine nucleotide translocator), member 6 | 293 | Small Molecule Transport (Nucleotide) |
| 2382 | SLC22A14 | solute carrier family 22 (organic cation transporter), member 14 | 9389 | Small Molecule Transport (Organic Cation) |
| 2383 | SLC22A15 | solute carrier family 22 (organic cation transporter), member 15 | 55356 | Small Molecule Transport (Organic Cation) |
| 2384 | SLC22A18 | solute carrier family 22 (organic cation transporter), member 18 | 5002 | Small Molecule Transport (Organic Cation) |
| 2385 | SLC22A18AS | solute carrier family 22 (organic cation transporter), member 18 antisense | 5003 | Small Molecule Transport (Organic Cation) |
| 2386 | SLC22A20 | solute carrier family 22 (organic anion transporter), member 20 | 440044 | Small Molecule Transport (Organic Cation) |
| 2387 | SLC22A17 | solute carrier family 22 (organic cation transporter), member 17 | 51310 | Small Molecule Transport (Organic Cation, Iron) |
| 2388 | PCTP | Phosphatidylcholine transfer protein | 58488 | Small Molecule Transport (PC) |
| 2389 | STARD10 | StAR-related lipid transfer (START) domain containing 10 | 10809 | Small Molecule Transport (PC, PE) |
| 2390 | SLC15A1 | solute carrier family 15 (oligopeptide transporter), member 1 | 6564 | Small Molecule Transport (peptide) |
| 2391 | SLC15A2 | solute carrier family 15 (H+/peptide transporter), member 2 | 6565 | Small Molecule Transport (peptide) |
| 2392 | SLC15A3 | solute carrier family 15, member 3 | 51296 | Small Molecule Transport (peptide) |
| 2393 | SLC15A4 | solute carrier family 15, member 4 | 121260 | Small Molecule Transport (peptide) |
| 2394 | SLCO5A1 | solute carrier organic anion transporter family, member 5A1 | 81796 | Small Molecule Transport (peptide) |
| 2395 | SLCO6A1 | solute carrier organic anion transporter family, member 6A1 | 133482 | Small Molecule Transport (peptide) |
| 2396 | SLC22A1 | solute carrier family 22 (organic cation transporter), member 1 | 6580 | Small Molecule Transport (PGE2,PGF2α) |
| 2397 | SLC17A2 | solute carrier family 17 (sodium phosphate), member 2 | 10246 | Small Molecule Transport (Phosphate) |
| 2398 | SLC17A3 | solute carrier family 17 (sodium phosphate), member 3 | 10786 | Small Molecule Transport (Phosphate) |
| 2399 | SLC17A4 | solute carrier family 17 (sodium phosphate), member 4 | 10050 | Small Molecule Transport (Phosphate) |
| 2400 | SLC20A1 | solute carrier family 20, member 1 | 6574 | Small Molecule Transport (Phosphate) |
| 2401 | SLC20A2 | solute carrier family 20, member 2 | 6575 | Small Molecule Transport (Phosphate) |
| 2402 | SLC37A3 | solute carrier family 37 (glycerol-3-phosphate transporter), member 3 | 84255 | Small Molecule Transport (Phosphosugar) |
| 2403 | ANKH | Ankylosis, progressive homolog (mouse) | 56172 | Small Molecule Transport (Ppi) |
| 2404 | SLC6A17 | solute carrier family 6 (neurotransmitter transporter), member 17 | 388662 | Small Molecule Transport (Pro, Cys, Ala) |
| 2405 | SLC36A4 | solute carrier family 36 (proton/amino acid symporter), member 4 | 120103 | Small Molecule Transport (Proline) |
| 2406 | SLC6A20 | solute carrier family 6 (neurotransmitter transporter), member 20 | 54716 | Small Molecule Transport (Proline) |
| 2407 | SLC6A7 | solute carrier family 6 (neurotransmitter transporter, L-proline), member 7 | 6534 | Small Molecule Transport (Proline) |
| 2408 | SLC16A11 | solute carrier family 16 (monocarboxylic acid transporters), member 11 | 162515 | Small Molecule Transport (Pyryuvate) |
| 2409 | SLC25A26 | solute carrier family 25 (mitochondrial carrier, phosphate carrier), member 26 | 115286 | Small Molecule Transport (S-adenosyl methionine) |
| 2410 | SFXN2 | Sideroflexin 2 | 118980 | Small Molecule Transport (Serine) |
| 2411 | SFXN1 | Sideroflexin 1 | 94081 | Small Molecule Transport (Serine) |
| 2412 | SFXN3 | Sideroflexin 3 | 81855 | Small Molecule Transport (Serine) |
| 2413 | SPNS1 | Spinster homolog 1 (Drosophila) | 83985 | Small Molecule Transport (Sphingolipid) |
| 2414 | SLC45A2 | solute carrier family 45, member 2 | 51151 | Small Molecule Transport (Sugar) |
| 2415 | SLC45A3 | solute carrier family 45, member 3 | 85414 | Small Molecule Transport (Sugar) |
| 2416 | SLC45A4 | solute carrier family 45, member 4 | 57210 | Small Molecule Transport (Sugar) |
| 2417 | SLC5A11 | solute carrier family 5 (sodium/glucose cotransporter), member 11 | 115584 | Small Molecule Transport (Sugar) |
| 2418 | SLC5A3 | solute carrier family 5 (inositol transporters), member 3 | 6526 | Small Molecule Transport (Sugar) |
| 2419 | SV2A | Synaptic vesicle glycoprotein 2A | 9900 | Small Molecule Transport (Sugar) |
| 2420 | SV2B | Synaptic vesicle glycoprotein 2B | 9899 | Small Molecule Transport (Sugar) |
| 2421 | SV2C | Synaptic vesicle glycoprotein 2C | 22987 | Small Molecule Transport (Sugar) |
| 2422 | SLC26A1 | solute carrier family 26 (sulfate transporter), member 1 | 10861 | Small Molecule Transport (Sulfate) |
| 2423 | SLC26A11 | solute carrier family 26, member 11 | 284129 | Small Molecule Transport (Sulfate) |
| 2424 | SLC26A2 | solute carrier family 26 (sulfate transporter), member 2 | 1836 | Small Molecule Transport (Sulfate) |
| 2425 | SLC16A2 | solute carrier family 16 (monocarboxylic acid transporters), member 2 | 6567 | Small Molecule Transport (T4) |
| 2426 | SLCO2B1 | solute carrier organic anion transporter family, member 2b1 | 11309 | Small Molecule Transport (T4) |
| 2427 | SLC19A2 | solute carrier family 19 (thiamine transporter), member 2 | 10560 | Small Molecule Transport (Thiamine) |
| 2428 | SLC19A3 | solute carrier family 19 (sodium/hydrogen exchanger), member 3 | 80704 | Small Molecule Transport (Thiamine) |
| 2429 | SLC47A1 | solute carrier family 47, member 1 | 55244 | Small Molecule Transport (Thiamine) |
| 2430 | SLC47A2 | solute carrier family 47, member 2 | 146802 | Small Molecule Transport (Thiamine) |
| 2431 | SLC19A1 | solute carrier family 19 (folate transporter), member 1 | 6573 | Small Molecule Transport (Thiamine, Folate) |
| 2432 | SLCO1C1 | solute carrier organic anion transporter family, member 1c1 | 53919 | Small Molecule Transport (Thyroid) |
| 2433 | SLC16A10 | solute carrier family 16 (monocarboxylic acid transporters), member 10 | 117247 | Small Molecule Transport (tryptophan, phenylalanine, levodopa, tyrosine) |
| 2434 | SLC35A3 | solute carrier family 35 (UDP-N-acetylglucosamine (UDP-GlcNAc) transporter), member 3 | 23443 | Small Molecule Transport (UDP-N-Acetyl-Glucosomine) |
| 2435 | SLC35A4 | solute carrier family 35, member A4 | 113829 | Small Molecule Transport (UDP-N-Acetyl-Glucosomine) |
| 2436 | SLC35B4 | solute carrier family 35, member B4 | 84912 | Small Molecule Transport (UDP-N-Acetyl-Glucosomine) |
| 2437 | SLC35D1 | solute carrier family 35 (UDP-glucuronic acid/UDP-N-acetylgalactosamine dual transporter), member D1 | 23169 | Small Molecule Transport (UDP-N-Acetyl-Glucosomine) |
| 2438 | MFSD1 | Major Facilitator Superfamily Domain Containing 1 | 64747 | Small Molecule Transport (Unknown) |
| 2439 | SLC16A14 | solute carrier family 16 (monocarboxylic acid transporters), member 14 | 151473 | Small Molecule Transport (Unknown) |
| 2440 | SLC7A5 | solute carrier family 7 (cationic amino acid transporter, y+ system), member 5 | 8140 | Small Molecule Transport (Unkown) |
| 2441 | SLC17A1 | solute carrier family 17 (sodium phosphate), member 1 | 6568 | Small Molecule Transport (Urate) |
| 2442 | SLC22A12 | solute carrier family 22 (organic anion/cation transporter), member 12 | 116085 | Small Molecule Transport (Urate) |
| 2443 | SLC2A9 | solute carrier family 2 (facilitated glucose transporter), member 9 | 56606 | Small Molecule Transport (Urate) |
| 2444 | SLC22A8 | solute carrier family 22 (organic anion transporter), member 8 | 9376 | Small Molecule Transport (Urate, aKG) |
| 2445 | SLC14A1 | solute carrier family 14 (urea transporter), member 1 | 6563 | Small Molecule Transport (Urea) |
| 2446 | SLC14A2 | solute carrier family 14 (urea transporter), member 2 | 8170 | Small Molecule Transport (Urea) |
| 2447 | SLC22A11 | solute carrier family 22 (organic anion/cation transporter), member 11 | 55867 | Small Molecule Transport (Uric acid |
| 2448 | SLC28A1 | solute carrier family 28 (sodium-coupled nucleoside transporter), member 1 | 9154 | Small Molecule Transport (Uridine) |
| 2449 | SLCO3A1 | solute carrier organic anion transporter family, member 3a1 | 28232 | Small Molecule Transport (vasopressin, prostaglandins, thyroid hormones) |
| 2450 | STRA6 | stimulated by retinoic acid gene 6 homolog (mouse) | 64220 | Small Molecule Transport (Vitamine A) |
| 2451 | SLC23A1 | solute carrier family 23 (nucleobase transporters), member 1 | 9963 | Small Molecule Transport (Vitamine C) |
| 2452 | SLC23A2 | solute carrier family 23 (nucleobase transporters), member 2 | 9962 | Small Molecule Transport (Vitamine C) |
| 2453 | SLC5A6 | solute carrier family 5 (sodium-dependent vitamin transporter), member 6 | 8884 | Small Molecule Transport (Vitamine) |
| 2454 | AQP12A | aquaporin 12A | 375318 | Small Molecule Transport (Water) |
| 2455 | AQP12B | aquaporin 12B | 653437 | Small Molecule Transport (Water) |
| 2456 | AQP2 | aquaporin 2 (collecting duct) | 359 | Small Molecule Transport (Water) |
| 2457 | AQP3 | aquaporin 3 (Gill blood group) | 360 | Small Molecule Transport (Water) |
| 2458 | AQP4 | aquaporin 4 | 361 | Small Molecule Transport (Water) |
| 2459 | AQP5 | aquaporin 5 | 362 | Small Molecule Transport (Water) |
| 2460 | AQP6 | aquaporin 6, kidney specific | 363 | Small Molecule Transport (Water) |
| 2461 | AQP7 | aquaporin 7 | 364 | Small Molecule Transport (Water) |
| 2462 | AQP8 | aquaporin 8 | 343 | Small Molecule Transport (Water) |
| 2463 | MIP | Major intrinsic protein of lens fiber | 4284 | Small Molecule Transport (Water) |
| 2464 | AQP10 | aquaporin 10 | 89872 | Small Molecule Transport (Water, Glycerol) |
| 2465 | AQP11 | aquaporin 11 | 282679 | Small Molecule Transport (Water, Glycerol) |
| 2466 | AQP9 | aquaporin 9 | 366 | Small Molecule Transport (Water, Glycerol) |
| 2467 | AQP1 | aquaporin 1 (Colton blood group) | 358 | Small Molecule Transport (Water, CO2, Ammonium) |
| 2468 | ABHD10 | abhydrolase domain containing 10 | 55347 | Sugar |
| 2469 | WWOX | WW domain containing oxidoreductase | 51741 | Sugar |
| 2470 | C9ORF103 | Chromosome 9 ORF 103 | 414328 | Sugar |
| 2471 | CRYL1 | crystallin, lambda 1 | 51084 | Sugar |
| 2472 | GFOD1 | glucose-fructose oxidoreductase domain containing 1 | 54438 | Sugar |
| 2473 | GFOD2 | glucose-fructose oxidoreductase domain containing 2 | 81577 | Sugar |
| 2474 | GRHPR | glyoxylate reductase/hydroxypyruvate reductase | 9380 | Sugar |
| 2475 | H6PD | Hexose-6-phosphate dehydrogenase (glucose 1-dehydrogenase) | 9563 | Sugar |
| 2476 | HYI | hydroxypyruvate isomerase homolog (E. coli) | 81888 | Sugar |
| 2477 | LCT | lactase | 3938 | Sugar |
| 2478 | RBKS | ribokinase | 64080 | Sugar |
| 2479 | SI | sucrase-isomaltase (alpha-glucosidase) | 6476 | Sugar |
| 2480 | SIAE | sialic acid acetylesterase | 54414 | Sugar |
| 2481 | SORD | sorbitol dehydrogenase | 6652 | Sugar |
| 2482 | TREH | trehalase (brush-border membrane glycoprotein) | 11181 | Sugar |
| 2483 | TSTA3 | tissue specific transplantation antigen P35B | 7264 | Sugar |
| 2484 | LCTL | lactase-like | 197021 | Sugar |
| 2485 | CHI3L1 | chitinase 3-like 1 (cartilage glycoprotein-39) | 1116 | Sugar (Aminosugar) |
| 2486 | CHI3L2 | chitinase 3-like 2 | 1117 | Sugar (Aminosugar) |
| 2487 | CHIA | chitinase, acidic | 27159 | Sugar (Aminosugar) |
| 2488 | CHID1 | chitinase domain containing 1 | 66005 | Sugar (Aminosugar) |
| 2489 | CHIT1 | chitinase 1 | 1118 | Sugar (Aminosugar) |
| 2490 | CMAS | Cytidine monophosphate N-acetylneuraminic acid synthetase | 55907 | Sugar (Aminosugar) |
| 2491 | GFPT1 | glutamine-fructose-6-phosphate transaminase 1 | 2673 | Sugar (Aminosugar) |
| 2492 | GFPT2 | glutamine-fructose-6-phosphate transaminase 2 | 9945 | Sugar (Aminosugar) |
| 2493 | GNE | glucosamine (UDP-N-acetyl)-2-epimerase/N-acetylmannosamine kinase | 10020 | Sugar (Aminosugar) |
| 2494 | GNPDA1 | glucosamine-6-phosphate deaminase 1 | 10007 | Sugar (Aminosugar) |
| 2495 | GNPDA2 | glucosamine-6-phosphate deaminase 2 | 132789 | Sugar (Aminosugar) |
| 2496 | GNPNAT1 | glucosamine-phosphate N-acetyltransferase 1 | 64841 | Sugar (Aminosugar) |
| 2497 | GNPTAB | N-acetylglucosamine-1-phosphate transferase, alpha and beta subunits | 79158 | Sugar (Aminosugar) |
| 2498 | GNPTG | N-acetylglucosamine-1-phosphate transferase, gamma subunit | 84572 | Sugar (Aminosugar) |
| 2499 | NAGK | N-acetylglucosamine kinase | 55577 | Sugar (Aminosugar) |
| 2500 | NANP | N-acetylneuraminic acid phosphatase | 140838 | Sugar (Aminosugar) |
| 2501 | NANS | N-acetylneuraminic acid synthase (sialic acid synthase) | 54187 | Sugar (Aminosugar) |
| 2502 | NPL | N-acetylneuraminate pyruvate lyase (dihydrodipicolinate synthase) | 80896 | Sugar (Aminosugar) |
| 2503 | PGM3 | phosphoglucomutase 3 | 5238 | Sugar (Aminosugar) |
| 2504 | RENBP | renin binding protein | 5973 | Sugar (Aminosugar) |
| 2505 | UAP1 | UDP-N-acteylglucosamine pyrophosphorylase 1 | 6675 | Sugar (Aminosugar) |
| 2506 | UAP1L1 | UDP-N-acteylglucosamine pyrophosphorylase 1-like 1 | 91373 | Sugar (Aminosugar) |
| 2507 | AMY1A | amylase, alpha 1A (salivary) | 276 | Sugar (Amylase) |
| 2508 | AMY1B | amylase, alpha 1B (salivary) | 277 | Sugar (Amylase) |
| 2509 | AMY1C | amylase, alpha 1C (salivary) | 278 | Sugar (Amylase) |
| 2510 | AMY2A | amylase, alpha 2A (pancreatic) | 279 | Sugar (Amylase) |
| 2511 | AMY2B | amylase, alpha 2B (pancreatic) | 280 | Sugar (Amylase) |
| 2512 | FPGT | fucose-1-phosphate guanylyltransferase | 8790 | Sugar (Fructose) |
| 2513 | FUK | fucokinase | 197258 | Sugar (Fructose) |
| 2514 | KHK | ketohexokinase (fructokinase) | 3795 | Sugar (Fructose) |
| 2515 | ENOSF1 | enolase superfamily member 1 | 55556 | Sugar (Fructose, mannose) |
| 2516 | GALK1 | galactokinase 1 | 2584 | Sugar (Galactose) |
| 2517 | GALK2 | galactokinase 2 | 2585 | Sugar (Galactose) |
| 2518 | GALM | Galactose mutarotase (aldose 1-epimerase) | 130589 | Sugar (Galactose) |
| 2519 | GALT | galactose-1-phosphate uridylyltransferase | 2592 | Sugar (Galactose) |
| 2520 | FBP1 | fructose-1,6-bisphosphatase 1 | 2203 | Sugar (gluconeogenesi) |
| 2521 | FBP2 | fructose-1,6-bisphosphatase 2 | 8789 | Sugar (gluconeogenesi) |
| 2522 | G6PC | glucose-6-phosphatase, catalytic subunit | 2538 | Sugar (gluconeogenesi) |
| 2523 | G6PC2 | glucose-6-phosphatase, catalytic, 2 | 57818 | Sugar (gluconeogenesi) |
| 2524 | G6PC3 | glucose 6 phosphatase, catalytic, 3 | 92579 | Sugar (gluconeogenesi) |
| 2525 | PCK1 | phosphoenolpyruvate carboxykinase 1 (soluble) | 5105 | Sugar (gluconeogenesi) |
| 2526 | PCK2 | phosphoenolpyruvate carboxykinase 2 (mitochondrial) | 5106 | Sugar (gluconeogenesi) |
| 2527 | AGL | amylo-1, 6-glucosidase, 4-alpha-glucanotransferase (glycogen debranching enzyme, glycogen storage disease type III) | 178 | Sugar (Glycogen) |
| 2528 | GAA | glucosidase, alpha; acid (Pompe disease, glycogen storage disease type II) | 2548 | Sugar (Glycogen) |
| 2529 | PYGB | phosphorylase, glycogen; brain | 5834 | Sugar (Glycogen) |
| 2530 | PYGL | phosphorylase, glycogen; liver (Hers disease, glycogen storage disease type VI) | 5836 | Sugar (Glycogen) |
| 2531 | PYGM | phosphorylase, glycogen; muscle (McArdle syndrome, glycogen storage disease type V) | 5837 | Sugar (Glycogen) |
| 2532 | GBE1 | glucan (1,4-alpha-), branching enzyme 1 (glycogen branching enzyme, Andersen disease, glycogen storage disease type IV) | 2632 | Sugar (Glycogen) |
| 2533 | GYG1 | glycogenin 1 | 2992 | Sugar (Glycogen) |
| 2534 | GYG2 | glycogenin 2 | 8908 | Sugar (Glycogen) |
| 2535 | GYS1 | glycogen synthase 1 (muscle) | 2997 | Sugar (Glycogen) |
| 2536 | GYS2 | glycogen synthase 2 (liver) | 2998 | Sugar (Glycogen) |
| 2537 | UGP2 | UDP-glucose pyrophosphorylase 2 | 7360 | Sugar (Glycogen) |
| 2538 | HISPPD1 | histidine acid phosphatase domain containing 1 | 23262 | Sugar (Inositol Phosphate) |
| 2539 | HISPPD2A | histidine acid phosphatase domain containing 2A | 9677 | Sugar (Inositol Phosphate) |
| 2540 | CDIPT | CDP-diacylglycerol--inositol 3-phosphatidyltransferase (phosphatidylinositol synthase) | 10423 | Sugar (Inositol Phosphate) |
| 2541 | FIG4 | FIG4 homolog (S. cerevisiae) | 9896 | Sugar (Inositol Phosphate) |
| 2542 | IMPA1 | inositol(myo)-1(or 4)-monophosphatase 1 | 3612 | Sugar (Inositol Phosphate) |
| 2543 | IMPA2 | inositol(myo)-1(or 4)-monophosphatase 2 | 3613 | Sugar (Inositol Phosphate) |
| 2544 | IMPAD1 | inositol monophosphatase domain containing 1 | 54928 | Sugar (Inositol Phosphate) |
| 2545 | INPP1 | inositol polyphosphate-1-phosphatase | 3628 | Sugar (Inositol Phosphate) |
| 2546 | INPP4A | inositol polyphosphate-4-phosphatase, type I, 107kDa | 3631 | Sugar (Inositol Phosphate) |
| 2547 | INPP4B | inositol polyphosphate-4-phosphatase, type II, 105kDa | 8821 | Sugar (Inositol Phosphate) |
| 2548 | INPP5A | inositol polyphosphate-5-phosphatase, 40kDa | 3632 | Sugar (Inositol Phosphate) |
| 2549 | INPP5B | inositol polyphosphate-5-phosphatase, 75kDa | 3633 | Sugar (Inositol Phosphate) |
| 2550 | INPP5D | inositol polyphosphate-5-phosphatase, 145kDa | 3635 | Sugar (Inositol Phosphate) |
| 2551 | INPP5E | inositol polyphosphate-5-phosphatase, 72 kDa | 56623 | Sugar (Inositol Phosphate) |
| 2552 | INPP5F | inositol polyphosphate-5-phosphatase F | 22876 | Sugar (Inositol Phosphate) |
| 2553 | INPP5J | inositol polyphosphate-5-phosphatase J | 27124 | Sugar (Inositol Phosphate) |
| 2554 | INPP5K | inositol polyphosphate-5-phosphatase K | 51763 | Sugar (Inositol Phosphate) |
| 2555 | INPPL1 | inositol polyphosphate phosphatase-like 1 | 3636 | Sugar (Inositol Phosphate) |
| 2556 | IP6K1 | inositol hexaphosphate kinase 1 | 9807 | Sugar (Inositol Phosphate) |
| 2557 | IP6K2 | inositol hexaphosphate kinase 2 | 51447 | Sugar (Inositol Phosphate) |
| 2558 | IP6K3 | inositol hexaphosphate kinase 3 | 117283 | Sugar (Inositol Phosphate) |
| 2559 | IPMK | inositol polyphosphate multikinase | 253430 | Sugar (Inositol Phosphate) |
| 2560 | IPPK | inositol 1,3,4,5,6-pentakisphosphate 2-kinase | 64768 | Sugar (Inositol Phosphate) |
| 2561 | ISYNA1 | myo-inositol 1-phosphate synthase A1 | 51477 | Sugar (Inositol Phosphate) |
| 2562 | ITPK1 | inositol 1,3,4-triphosphate 5/6 kinase | 3705 | Sugar (Inositol Phosphate) |
| 2563 | ITPKA | inositol 1,4,5-trisphosphate 3-kinase A | 3706 | Sugar (Inositol Phosphate) |
| 2564 | ITPKB | inositol 1,4,5-trisphosphate 3-kinase B | 3707 | Sugar (Inositol Phosphate) |
| 2565 | ITPKC | inositol 1,4,5-trisphosphate 3-kinase C | 80271 | Sugar (Inositol Phosphate) |
| 2566 | MINPP1 | multiple inositol polyphosphate histidine phosphatase, 1 | 9562 | Sugar (Inositol Phosphate) |
| 2567 | MIOX | myo-inositol oxygenase | 55586 | Sugar (Inositol Phosphate) |
| 2568 | PGAP1 | GPI deacylase | 80055 | Sugar (Inositol Phosphate) |
| 2569 | PI4K2A | phosphatidylinositol 4-kinase type 2 alpha | 55361 | Sugar (Inositol Phosphate) |
| 2570 | PI4K2B | Phosphatidylinositol 4-kinase type 2 beta | 55300 | Sugar (Inositol Phosphate) |
| 2571 | PI4KA | phosphatidylinositol 4-kinase, catalytic, alpha | 5297 | Sugar (Inositol Phosphate) |
| 2572 | PI4KB | phosphatidylinositol 4-kinase, catalytic, beta | 5298 | Sugar (Inositol Phosphate) |
| 2573 | PIK3C2A | phosphoinositide-3-kinase, class 2, alpha polypeptide | 5286 | Sugar (Inositol Phosphate) |
| 2574 | PIK3C2B | phosphoinositide-3-kinase, class 2, beta polypeptide | 5287 | Sugar (Inositol Phosphate) |
| 2575 | PIK3C2G | phosphoinositide-3-kinase, class 2, gamma polypeptide | 5288 | Sugar (Inositol Phosphate) |
| 2576 | PIK3C3 | phosphoinositide-3-kinase, class 3 | 5289 | Sugar (Inositol Phosphate) |
| 2577 | PIK3CA | phosphoinositide-3-kinase, catalytic, alpha polypeptide | 5290 | Sugar (Inositol Phosphate) |
| 2578 | PIK3CB | phosphoinositide-3-kinase, catalytic, beta polypeptide | 5291 | Sugar (Inositol Phosphate) |
| 2579 | PIK3CD | phosphoinositide-3-kinase, catalytic, delta polypeptide | 5293 | Sugar (Inositol Phosphate) |
| 2580 | PIK3CG | phosphoinositide-3-kinase, catalytic, gamma polypeptide | 5294 | Sugar (Inositol Phosphate) |
| 2581 | PIK3R1 | phosphoinositide-3-kinase, regulatory subunit 1 (p85 alpha) | 5295 | Sugar (Inositol Phosphate) |
| 2582 | PIK3R2 | phosphoinositide-3-kinase, regulatory subunit 2 (p85 beta) | 5296 | Sugar (Inositol Phosphate) |
| 2583 | PIK3R3 | phosphoinositide-3-kinase, regulatory subunit 3 (p55, gamma) | 8503 | Sugar (Inositol Phosphate) |
| 2584 | PIK3R4 | phosphoinositide-3-kinase, regulatory subunit 4, p150 | 30849 | Sugar (Inositol Phosphate) |
| 2585 | PIK3R5 | phosphoinositide-3-kinase, regulatory subunit 5, p101 | 23533 | Sugar (Inositol Phosphate) |
| 2586 | PIK3R6 | phosphoinositide-3-kinase, regulatory subunit 6 | 146850 | Sugar (Inositol Phosphate) |
| 2587 | PIP4K2A | phosphatidylinositol-5-phosphate 4-kinase, type II, alpha | 5305 | Sugar (Inositol Phosphate) |
| 2588 | PIP4K2B | phosphatidylinositol-5-phosphate 4-kinase, type II, beta | 8396 | Sugar (Inositol Phosphate) |
| 2589 | PIP4K2C | phosphatidylinositol-5-phosphate 4-kinase, type II, gamma | 79837 | Sugar (Inositol Phosphate) |
| 2590 | PIP5K1A | phosphatidylinositol-4-phosphate 5-kinase, type I, alpha | 8394 | Sugar (Inositol Phosphate) |
| 2591 | PIP5K1B | phosphatidylinositol-4-phosphate 5-kinase, type I, beta | 8395 | Sugar (Inositol Phosphate) |
| 2592 | PIP5K1C | phosphatidylinositol-4-phosphate 5-kinase, type I, gamma | 23396 | Sugar (Inositol Phosphate) |
| 2593 | PIP5K3 | phosphatidylinositol-3-phosphate/phosphatidylinositol 5-kinase, type III | 200576 | Sugar (Inositol Phosphate) |
| 2594 | PIP5KL1 | phosphatidylinositol-4-phosphate 5-kinase-like 1 | 138429 | Sugar (Inositol Phosphate) |
| 2595 | PTEN | phosphatase and tensin homolog | 5728 | Sugar (Inositol Phosphate) |
| 2596 | SYNJ1 | synaptojanin 1 | 8867 | Sugar (Inositol Phosphate) |
| 2597 | SYNJ2 | synaptojanin 2 | 8871 | Sugar (Inositol Phosphate) |
| 2598 | LALBA | lactalbumin, alpha- | 3906 | Sugar (Lactose) |
| 2599 | GLYCTK | glycerate kinase | 132158 | Sugar (Mannose) |
| 2600 | GMDS | GDP-mannose 4,6-dehydratase | 2762 | Sugar (Mannose) |
| 2601 | GMPPA | GDP-mannose pyrophosphorylase A | 29926 | Sugar (Mannose) |
| 2602 | GMPPB | GDP-mannose pyrophosphorylase B | 29925 | Sugar (Mannose) |
| 2603 | MGAM | maltase-glucoamylase (alpha-glucosidase) | 8972 | Sugar (Mannose) |
| 2604 | MPI | mannose phosphate isomerase | 4351 | Sugar (Mannose) |
| 2605 | PMM1 | phosphomannomutase 1 | 5372 | Sugar (Mannose) |
| 2606 | PMM2 | phosphomannomutase 2 | 5373 | Sugar (Mannose) |
| 2607 | TGDS | TDP-glucose 4,6-dehydratase | 23483 | Sugar (Nucleotide Sugar) |
| 2608 | UGDH | UDP-glucose dehydrogenase | 7358 | Sugar (Nucleotide Sugar) |
| 2609 | UXS1 | UDP-glucuronate decarboxylase 1 | 80146 | Sugar (Nucleotide Sugar) |
| 2610 | PPA1 | pyrophosphatase (inorganic) 1 | 5464 | Sugar (PPi hydrolysis) |
| 2611 | DERA | 2-deoxyribose-5-phosphate aldolase homolog (C. elegans) | 51071 | Sugar (PPP) |
| 2612 | G6PD | glucose-6-phosphate dehydrogenase | 2539 | Sugar (PPP) |
| 2613 | PGD | phosphogluconate dehydrogenase | 5226 | Sugar (PPP) |
| 2614 | PGLS | 6-phosphogluconolactonase | 25796 | Sugar (PPP) |
| 2615 | RPE | ribulose-5-phosphate-3-epimerase | 6120 | Sugar (PPP) |
| 2616 | RPIA | ribose 5-phosphate isomerase A (ribose 5-phosphate epimerase) | 22934 | Sugar (PPP) |
| 2617 | TALDO1 | transaldolase 1 | 6888 | Sugar (PPP) |
| 2618 | TKT | transketolase (Wernicke-Korsakoff syndrome) | 7086 | Sugar (PPP) |
| 2619 | TKTL1 | transketolase-like 1 | 8277 | Sugar (PPP) |
| 2620 | TKTL2 | transketolase-like 2 | 84076 | Sugar (PPP) |
| 2621 | AGK | acylglycerol kinase | 55750 | Sugar (PPP) |
| 2622 | DCXR | dicarbonyl/L-xylulose reductase | 51181 | Sugar (Xyulose) |
| 2623 | XYLB | xylulokinase homolog (H. influenzae) | 9942 | Sugar (Xyulose) |
| 2624 | STS | steroid sulfatase (microsomal), isozyme S | 412 | Sulfate |
| 2625 | SULT2A1 | sulfotransferase family, cytosolic, 2A, dehydroepiandrosterone (DHEA)-preferring, member 1 | 6822 | Sulfate |
| 2626 | SULT4A1 | sulfotransferase family 4A, member 1 | 25830 | Sulfate |
| 2627 | ARSA | Arylsulfatase A | 410 | Sulfate |
| 2628 | ARSB | arylsulfatase B | 411 | Sulfate |
| 2629 | ARSH | arylsulfatase H | 347527 | Sulfate |
| 2630 | ARSI | arylsulfatase family, member I | 340075 | Sulfate |
| 2631 | ARSG | arylsulfatase G | 22901 | Sulfate |
| 2632 | PAPSS1 | 3'-phosphoadenosine 5'-phosphosulfate synthase 1 | 9061 | Sulfate |
| 2633 | PAPSS2 | 3'-phosphoadenosine 5'-phosphosulfate synthase 2 | 9060 | Sulfate |
| 2634 | SULT1B1 | sulfotransferase family, cytosolic, 1B, member 1 | 27284 | Sulfate |
| 2635 | SULT1E1 | sulfotransferase family 1E, estrogen-preferring, member 1 | 6783 | Sulfate |
| 2636 | SULT2B1 | sulfotransferase family, cytosolic, 2B, member 1 | 6820 | Sulfate |
| 2637 | SULT1A1 | sulfotransferase family, cytosolic, 1A, phenol-preferring, member 1 | 6817 | Sulfate |
| 2638 | DSE | dermatan sulfate epimerase | 29940 | Sulfate |
| 2639 | DSEL | dermatan sulfate epimerase-like | 92126 | Sulfate |
| 2640 | MPST | mercaptopyruvate sulfurtransferase | 4357 | Sulfate |
| 2641 | GALNAC4S-6ST | B cell RAG associated protein | 51363 | Sulfate |
| 2642 | SULT1A2 | sulfotransferase family, cytosolic, 1A, phenol-preferring, member 2 | 6799 | Sulfate |
| 2643 | SULT1A3 | sulfotransferase family, cytosolic, 1A, phenol-preferring, member 3 | 6818 | Sulfate |
| 2644 | SULT1A4 | sulfotransferase family, cytosolic, 1A, phenol-preferring, member 4 | 445329 | Sulfate |
| 2645 | SULT1C3 | sulfotransferase family, cytosolic, 1C, member 3 | 442038 | Sulfate |
| 2646 | SULT1C4 | sulfotransferase family, cytosolic, 1C, member 4 | 27233 | Sulfate |
| 2647 | SULT6B1 | sulfotransferase family, cytosolic, 6B, member 1 | 391365 | Sulfate |
| 2648 | SRXN1 | sulfiredoxin 1 homolog (S. cerevisiae) | 140809 | Sulfate |
| 2649 | SULT1C2 | sulfotransferase family, cytosolic, 1C, member 2 | 6819 | Sulfate |
| 2650 | TST | thiosulfate sulfurtransferase (rhodanese) | 7263 | Sulfate |
| 2651 | ACLY | ATP citrate lyase | 47 | TCA Cycle |
| 2652 | ACO1 | aconitase 1, soluble | 48 | TCA Cycle |
| 2653 | ACO2 | aconitase 2, mitochondrial | 50 | TCA Cycle |
| 2654 | CS | citrate synthase | 1431 | TCA Cycle |
| 2655 | D2HGDH | D-2-hydroxyglutarate dehydrogenase | 728294 | TCA Cycle |
| 2656 | DLST | dihydrolipoamide S-succinyltransferase (E2 component of 2-oxo-glutarate complex) | 1743 | TCA Cycle |
| 2657 | FH | fumarate hydratase | 2271 | TCA Cycle |
| 2658 | HCCS | holocytochrome c synthase (cytochrome c heme-lyase) | 3052 | TCA Cycle |
| 2659 | IDH1 | isocitrate dehydrogenase 1 (NADP+), soluble | 3417 | TCA Cycle |
| 2660 | IDH2 | isocitrate dehydrogenase 2 (NADP+), mitochondrial | 3418 | TCA Cycle |
| 2661 | IDH3A | isocitrate dehydrogenase 3 (NAD+) alpha | 3419 | TCA Cycle |
| 2662 | IDH3B | isocitrate dehydrogenase 3 (NAD+) beta | 3420 | TCA Cycle |
| 2663 | IDH3G | isocitrate dehydrogenase 3 (NAD+) gamma | 3421 | TCA Cycle |
| 2664 | L2HGDH | L-2-hydroxyglutarate dehydrogenase | 79944 | TCA Cycle |
| 2665 | MDH1 | malate dehydrogenase 1, NAD (soluble) | 4190 | TCA Cycle |
| 2666 | MDH1B | malate dehydrogenase 1B, NAD (soluble) | 130752 | TCA Cycle |
| 2667 | MDH2 | malate dehydrogenase 2, NAD (mitochondrial) | 4191 | TCA Cycle |
| 2668 | ME1 | malic enzyme 1, NADP (+)-dependent, cytosolic | 4199 | TCA Cycle |
| 2669 | ME2 | malic enzyme 2, NAD (+)-dependent, mitochondrial | 4200 | TCA Cycle |
| 2670 | ME3 | malic enzyme 3, NADP (+)-dependent, mitochondrial | 10873 | TCA Cycle |
| 2671 | OGDH | oxoglutarate (alpha-ketoglutarate) dehydrogenase (lipoamide) | 4967 | TCA Cycle |
| 2672 | OGDHL | oxoglutarate dehydrogenase-like | 55753 | TCA Cycle |
| 2673 | PC | pyruvate carboxylase | 5091 | TCA Cycle |
| 2674 | SUCLA2 | succinate-CoA ligase, ADP-forming, beta subunit | 8803 | TCA Cycle |
| 2675 | SUCLG1 | succinate-CoA ligase, alpha subunit | 8802 | TCA Cycle |
| 2676 | SUCLG2 | succinate-CoA ligase, GDP-forming, beta subunit | 8801 | TCA Cycle |
| 2677 | GOT1 | glutamic-oxaloacetic transaminase 1, soluble (aspartate aminotransferase 1) | 2805 | TCA Cycle |
| 2678 | GOT1L1 | glutamic-oxaloacetic transaminase 1-like 1 | 137362 | TCA Cycle |
| 2679 | GOT2 | glutamic-oxaloacetic transaminase 2, mitochondrial (aspartate aminotransferase 2) | 2806 | TCA Cycle |
| 2680 | PDHA1 | pyruvate dehydrogenase (lipoamide) alpha 1 | 5160 | TCA Cycle |
| 2681 | PDHA2 | pyruvate dehydrogenase (lipoamide) alpha 2 | 5161 | TCA Cycle |
| 2682 | PDHB | pyruvate dehydrogenase (lipoamide) beta | 5162 | TCA Cycle |
| 2683 | PDHX | pyruvate dehydrogenase complex, component X | 8050 | TCA Cycle |
| 2684 | DLD | dihydrolipoamide dehydrogenase | 1738 | TCA Cycle |
| 2685 | COQ7 | coenzyme Q7 homolog, ubiquinone (yeast) | 10229 | Ubiquinone |
| 2686 | COQ9 | coenzyme Q9 homolog (S. cerevisiae) | 57017 | Ubiquinone |
| 2687 | COQ2 | coenzyme Q2 homolog, prenyltransferase (yeast) | 27235 | Ubiquinone |
| 2688 | COQ3 | coenzyme Q3 homolog, methyltransferase (S. cerevisiae) | 51805 | Ubiquinone |
| 2689 | COQ10A | coenzyme Q10 homolog A (S. cerevisiae) | 93058 | Ubiquinone |
| 2690 | COQ10B | Coenzyme Q10 homolog B (S. cerevisiae) | 80219 | Ubiquinone |
| 2691 | PDSS1 | prenyl (decaprenyl) diphosphate synthase, subunit 1 | 23590 | Ubiquinone |
| 2692 | PDSS2 | prenyl (decaprenyl) diphosphate synthase, subunit 2 | 57107 | Ubiquinone |
| 2693 | COQ5 | coenzyme Q5 homolog, methyltransferase (S. cerevisiae) | 84274 | Ubiquinone |
| 2694 | CABC1 | chaperone, ABC1 activity of bc1 complex homolog (S. pombe) | 56997 | Ubiquinone |
| 2695 | SQRDL | sulfide quinone reductase-like (yeast) | 58472 | Ubiquinone |
| 2696 | COQ6 | coenzyme Q6 homolog, monooxygenase (S. cerevisiae) | 51004 | Ubiquinone |
| 2697 | COQ4 | coenzyme Q4 homolog (S. cerevisiae) | 51117 | Ubiquinone |
| 2698 | BCMO1 | beta carotene 15, 15 monooxygenase 1 | 53630 | Vitamin (A) |
| 2699 | BCO2 | beta-carotene dioxygenase 2 | 83875 | Vitamin (A) |
| 2700 | CYP26A1 | cytochrome P450, family 26, subfamily A, polypeptide 1 | 1592 | Vitamin (A) |
| 2701 | CYP26B1 | cytochrome P450, family 26, subfamily B, polypeptide 1 | 56603 | Vitamin (A) |
| 2702 | CYP26C1 | cytochrome P450, family 26, subfamily C, polypeptide 1 | 340665 | Vitamin (A) |
| 2703 | LRAT | lecithin retinol acyltransferase (phosphatidylcholine--retinol O-acyltransferase) | 9227 | Vitamin (A) |
| 2704 | RDH10 | retinol dehydrogenase 10 (all-trans) | 157506 | Vitamin (A) |
| 2705 | RDH11 | retinol dehydrogenase 11 (all-trans/9-cis/11-cis) | 51109 | Vitamin (A) |
| 2706 | RDH12 | retinol dehydrogenase 12 (all-trans/9-cis/11-cis) | 145226 | Vitamin (A) |
| 2707 | RDH13 | retinol dehydrogenase 13 (all-trans/9-cis) | 112724 | Vitamin (A) |
| 2708 | RDH14 | retinol dehydrogenase 14 (all-trans/9-cis/11-cis) | 57665 | Vitamin (A) |
| 2709 | RDH16 | retinol dehydrogenase 16 (all-trans) | 8608 | Vitamin (A) |
| 2710 | RDH5 | retinol dehydrogenase 5 (11-cis/9-cis) | 5959 | Vitamin (A) |
| 2711 | RDH8 | retinol dehydrogenase 8 (all-trans) | 50700 | Vitamin (A) |
| 2712 | RETSAT | retinol saturase (all-trans-retinol 13,14-reductase) | 54884 | Vitamin (A) |
| 2713 | RPE65 | retinal pigment epithelium-specific protein 65kDa | 6121 | Vitamin (A) |
| 2714 | SDR16C5 | short chain dehydrogenase/reductase family 16C, member 5 | 195814 | Vitamin (A) |
| 2715 | MMAB | methylmalonic aciduria (cobalamin deficiency) cblB type | 326625 | Vitamin (B12) |
| 2716 | MMACHC | methylmalonic aciduria (cobalamin deficiency) cblC type, with homocystinuria | 25974 | Vitamin (B12) |
| 2717 | MMADHC | methylmalonic aciduria (cobalamin deficiency) cblD type, with homocystinuria | 27249 | Vitamin (B12) |
| 2718 | MTRR | 5-methyltetrahydrofolate-homocysteine methyltransferase reductase | 4552 | Vitamin (B12) |
| 2719 | PDXK | pyridoxal (pyridoxine, vitamin B6) kinase | 8566 | Vitamin (B6) |
| 2720 | PDXP | pyridoxal (pyridoxine, vitamin B6) phosphatase | 57026 | Vitamin (B6) |
| 2721 | PHOSPHO2 | phosphatase orphan 2 | 493911 | Vitamin (B6) |
| 2722 | PNPO | pyridoxamine 5'-phosphate oxidase | 55163 | Vitamin (B6) |
| 2723 | BTD | biotinidase | 686 | Vitamin (Biotin) |
| 2724 | FOLH1 | folate hydrolase (prostate-specific membrane antigen) 1 | 2346 | Vitamin (Folate) |
| 2725 | GPHN | gephyrin | 10243 | Vitamin (Folate) |
| 2726 | DHFR | dihydrofolate reductase | 1719 | Vitamin (Folate) |
| 2727 | DHFRL1 | dihydrofolate reductase-like 1 | 200895 | Vitamin (Folate) |
| 2728 | FPGS | folylpolyglutamate synthase | 2356 | Vitamin (Folate) |
| 2729 | MTHFD1 | methylenetetrahydrofolate dehydrogenase (NADP+ dependent) 1, methenyltetrahydrofolate cyclohydrolase, formyltetrahydrofolate synthetase | 4522 | Vitamin (Folate) |
| 2730 | MTHFD1L | Methylenetetrahydrofolate dehydrogenase (NADP+ dependent) 1-like | 25902 | Vitamin (Folate) |
| 2731 | MTHFD2 | methylenetetrahydrofolate dehydrogenase (NADP+ dependent) 2, methenyltetrahydrofolate cyclohydrolase | 10797 | Vitamin (Folate) |
| 2732 | MTHFD2L | methylenetetrahydrofolate dehydrogenase (NADP+ dependent) 2-like | 441024 | Vitamin (Folate) |
| 2733 | MTHFR | 5,10-methylenetetrahydrofolate reductase (NADPH) | 4524 | Vitamin (Folate) |
| 2734 | MTHFS | 5,10-methenyltetrahydrofolate synthetase (5-formyltetrahydrofolate cyclo-ligase) | 10588 | Vitamin (Folate) |
| 2735 | MTHFSD | methenyltetrahydrofolate synthetase domain containing | 64779 | Vitamin (Folate) |
| 2736 | SHMT1 | serine hydroxymethyltransferase 1 (soluble) | 6470 | Vitamin (Folate) |
| 2737 | SHMT2 | serine hydroxymethyltransferase 2 (mitochondrial) | 6472 | Vitamin (Folate) |
| 2738 | GGH | gamma-glutamyl hydrolase (conjugase, folylpolygammaglutamyl hydrolase) | 8836 | Vitamin (Folate) |
| 2739 | VKORC1 | vitamin K epoxide reductase complex, subunit 1 | 79001 | Vitamin (K) |
| 2740 | VKORC1L1 | vitamin K epoxide reductase complex, subunit 1-like 1 | 154807 | Vitamin (K) |
| 2741 | RFK | riboflavin kinase | 55312 | Vitamin (Riboflavin) |
| 2742 | THTPA | Thiamine triphosphatase | 79178 | Vitamin (Thiamine) |
| 2743 | TPK1 | thiamin pyrophosphokinase 1 | 27010 | Vitamin (Thiamine) |
| 2744 | ALDH1A3 | aldehyde dehydrogenase 1 family, member A3 | 220 | Vitamine (Retinoic acidl, Aldehyde) |
| 2745 | ADH1A | alcohol dehydrogenase 1A (class I), alpha polypeptide | 124 | Vitamine (Retinoic acid, Alcohol) |
| 2746 | ADH1B | Alcohol dehydrogenase 1B (class I), beta polypeptide | 125 | Vitamine (Retinoic acid, Alcohol) |
| 2747 | ADH1C | alcohol dehydrogenase 1C (class I), gamma polypeptide | 126 | Vitamine (Retinoic acid, Alcohol) |
| 2748 | ADH4 | alcohol dehydrogenase 4 (class II), pi polypeptide | 127 | Vitamine (Retinoic acid, Alcohol) |
| 2749 | ADH5 | alcohol dehydrogenase 5 (class III), chi polypeptide | 128 | Vitamine (Retinoic acid, Alcohol) |
| 2750 | ADH6 | alcohol dehydrogenase 6 (class V) | 130 | Vitamine (Retinoic acid, Alcohol) |
| 2751 | ADH7 | alcohol dehydrogenase 7 (class IV), mu or sigma polypeptide | 131 | Vitamine (Retinoic acid, Alcohol) |
| 2752 | AKR1A1 | Aldo-keto reductase family 1, member A1 (aldehyde reductase) | 10327 | Vitamine (Retinoic acid, Alcohol) |
| 2753 | AKR1B1 | aldo-keto reductase family 1, member B1 (aldose reductase) | 231 | Vitamine (Retinoic acid, Alcohol) |
| 2754 | AKR1B10 | aldo-keto reductase family 1, member B10 (aldose reductase) | 57016 | Vitamine (Retinoic acid, Alcohol) |
| 2755 | AKR1C1 | aldo-keto reductase family 1, member C1 (dihydrodiol dehydrogenase 1; 20-alpha (3-alpha)-hydroxysteroid dehydrogenase) | 1645 | Vitamine (Retinoic acid, Alcohol) |
| 2756 | AKR1C2 | Aldo-keto reductase family 1, member C2 (dihydrodiol dehydrogenase 2; bile acid binding protein; 3-alpha hydroxysteroid dehydrogenase, type III) | 1646 | Vitamine (Retinoic acid, Alcohol) |
| 2757 | AKR1C3 | aldo-keto reductase family 1, member C3 (3-alpha hydroxysteroid dehydrogenase, type II) | 8644 | Vitamine (Retinoic acid, Alcohol) |
| 2758 | AKR1CL2 | aldo-keto reductase family 1, member C-like 2 | 83592 | Vitamine (Retinoic acid, Alcohol) |
| 2759 | AKR7A2 | aldo-keto reductase family 7, member A2 (aflatoxin aldehyde reductase) | 8574 | Vitamine (Retinoic acid, Alcohol) |
| 2760 | AKR7A3 | aldo-keto reductase family 7, member A3 (aflatoxin aldehyde reductase) | 22977 | Vitamine (Retinoic acid, Alcohol) |
| 2761 | ALDH16A1 | aldehyde dehydrogenase 16 family, member A1 | 126133 | Vitamine (Retinoic acid, Alcohol) |
| 2762 | ALDH1A1 | aldehyde dehydrogenase 1 family, member A1 | 216 | Vitamine (Retinoic acid, Alcohol) |
| 2763 | ALDH1A2 | aldehyde dehydrogenase 1 family, member A2 | 8854 | Vitamine (Retinoic acid, Alcohol) |
| 2764 | ALDH8A1 | aldehyde dehydrogenase 8 family, member A1 | 64577 | Vitamine (Retinoic acid, Alcohol) |

**Table S3.** **Differentially expressed metabolic genes in retinal organoids.**

| **Gene symbol** | **1M_1** | **1M_2** | **1M_3** | **3M_1** | **3M_2** | **3M_3** | **6M_1** | **6M_2** | **6M_3** | **9M_1** | **9M_2** | **9M_3** | **Classification (Sub-category/substrate)** | **In figures** |
| --- | --- | --- | --- | --- | --- | --- | --- | --- | --- | --- | --- | --- | --- | --- |
| SLC1A7 | 0.41 | 0.52 | 0.69 | 36.0 | 42.6 | 40.0 | 34.7 | 34.6 | 39.3 | 27.4 | 28.6 | 27.4 | Small Molecule Transport (Glutamate, Asparate) | Fig 1i |
| SLC17A7 | 2.31 | 2.77 | 2.93 | 48.2 | 52.0 | 52.2 | 83.3 | 86.4 | 94.2 | 79.9 | 99.9 | 78.5 | Small Molecule Transport (Glutamate) | Fig 1i |
| SLC1A2 | 0.81 | 1.12 | 0.69 | 7.60 | 9.81 | 8.28 | 12.9 | 11.3 | 12.6 | 15.2 | 13.6 | 16.1 | Small Molecule Transport (Glutamate, Asparate) | Fig 1i |
| SLC1A3 | 8.98 | 6.83 | 7.33 | 11.7 | 17.2 | 11.7 | 108 | 159 | 128 | 161 | 91.8 | 112 | Small Molecule Transport (Glutamate, Asparate) | Fig 1i |
| SLC17A8 | 0.15 | 0.21 | 0.35 | 0.44 | 0.20 | 0.17 | 0.21 | 0.24 | 0.17 | 0.31 | 0.18 | 0.11 | Small Molecule Transport (Glutamate) | Fig 1i |
| SLC17A6 | 2.06 | 3.15 | 3.84 | 4.63 | 1.76 | 1.66 | 0.27 | 0.13 | 0.13 | 0.17 | 0.08 | 0.12 | Small Molecule Transport (Glutamate) | Fig 1i |
| SV2B | 0.42 | 0.62 | 0.63 | 13.9 | 17.3 | 18.9 | 36.2 | 41.1 | 41.5 | 41.2 | 44.9 | 33.0 | Small Molecule Transport (Sugar) | Fig 1i |
| SLC45A3 | 0.32 | 0.28 | 0.28 | 0.36 | 0.53 | 0.66 | 0.60 | 0.34 | 0.25 | 0.29 | 0.33 | 0.45 | Small Molecule Transport (Sugar) | Fig 1i |
| SLC2A1 | 155 | 153 | 131 | 111 | 67.0 | 66.6 | 207 | 116 | 123 | 229 | 98.5 | 132 | Small Molecule Transport (Glucose, Glucosamine) | Fig 1i |
| SLC2A4 | 1.26 | 1.29 | 1.15 | 0.19 | 0.30 | 0.10 | 0.58 | 0.50 | 0.56 | 0.65 | 0.26 | 0.59 | Small Molecule Transport (Glucose, Glucosamine) | Fig 1i |
| SLC35F3 | 2.60 | 2.32 | 2.41 | 1.80 | 1.13 | 1.30 | 0.58 | 0.49 | 0.64 | 0.37 | 0.42 | 0.41 | Small Molecule Transport (Nucleotide sugar) | Fig 1i |
| AQP4 | 0.26 | 0.24 | 0.29 | 0.22 | 1.36 | 0.23 | 4.69 | 6.08 | 13.2 | 25.6 | 5.19 | 24.3 | Small Molecule Transport (Water) | Fig 1i |
| AQP1 | 1.24 | 1.06 | 1.32 | 1.14 | 3.28 | 1.42 | 28.4 | 12.4 | 15.7 | 27.5 | 5.26 | 11.7 | Small Molecule Transport(Water, CO2, Ammonium) | Fig 1i |
| AQP9 | 0.01 | 0.00 | 0.01 | 0.00 | 0.00 | 0.00 | 0.01 | 0.11 | 0.06 | 0.04 | 0.01 | 0.03 | Small Molecule Transport (Water, Glycerol) | Fig 1i |
| AQP3 | 1.21 | 1.25 | 1.21 | 6.25 | 7.20 | 8.16 | 3.63 | 3.13 | 2.85 | 2.82 | 3.35 | 2.19 | Small Molecule Transport (Water) | Fig 1i |
| MIP | 4.05 | 16.5 | 13.1 | 1.58 | 1.13 | 0.24 | 0.10 | 0.10 | 0.19 | 0.08 | 0.07 | 0.06 | Small Molecule Transport (Water) | Fig 1i |
| SLC6A11 | 0.21 | 0.07 | 0.03 | 0.72 | 3.79 | 1.93 | 7.25 | 9.46 | 9.01 | 10.1 | 5.19 | 5.94 | Small Molecule Transport (GABA, β-Alanine) | Fig 1i |
| SLC6A6 | 9.23 | 8.95 | 9.61 | 23.1 | 32.6 | 29.6 | 45.3 | 63.3 | 74.3 | 68.1 | 98.9 | 61.4 | Small Molecule Transport (GABA, | Fig 1i |
| SLC32A1 | 0.21 | 0.31 | 0.06 | 1.45 | 3.34 | 1.97 | 2.08 | 1.54 | 1.32 | 1.11 | 1.64 | 1.09 | Small Molecule Transport (GABA, Gly) | Fig 1i |
| SLC6A1 | 1.43 | 2.33 | 2.54 | 1.94 | 7.44 | 2.92 | 5.45 | 7.58 | 7.85 | 7.01 | 5.46 | 5.65 | Small Molecule Transport (GABA) | Fig 1i |
| SLC38A5 | 0.86 | 0.98 | 1.16 | 40.4 | 35.2 | 36.9 | 14.9 | 14.8 | 14.3 | 7.95 | 8.61 | 8.00 | Small Molecule Transport (Asn, Se, His, Gln) | Fig 1i |
| SFXN3 | 2.23 | 3.07 | 3.51 | 13.9 | 13.4 | 13.1 | 22.7 | 21.9 | 19.8 | 20.7 | 20.4 | 16.2 | Small Molecule Transport (Serine) | Fig 1i |
| SLC6A5 | 0.24 | 0.08 | 0.15 | 0.44 | 0.24 | 0.13 | 1.21 | 0.18 | 0.09 | 0.36 | 0.00 | 0.02 | Small Molecule Transport (Glycine) | Fig 1i |
| SLC6A18 | 0.08 | 0.11 | 0.09 | 0.09 | 0.05 | 0.09 | 0.05 | 0.00 | 0.08 | 0.02 | 0.02 | 0.04 | Small Molecule Transport (Ala, Gly, Meth, Phe) | Fig 1i |
| SLC26A3 | 0.00 | 0.00 | 0.00 | 0.00 | 0.02 | 0.00 | 0.00 | 0.00 | 0.00 | 0.00 | 0.00 | 0.02 | Small Molecule Transport (Biocarbonate) | Fig 1i |
| SLC4A10 | 0.05 | 0.13 | 0.11 | 0.29 | 1.43 | 0.31 | 4.33 | 6.54 | 5.28 | 7.18 | 7.14 | 6.09 | Small Molecule Transport (Biocarbonate) | Fig 1i |
| SLC26A7 | 0.13 | 0.15 | 0.10 | 0.92 | 0.32 | 0.65 | 1.02 | 2.12 | 0.80 | 0.56 | 0.85 | 0.69 | Small Molecule Transport (Biocarbonate) | Fig 1i |
| SLC26A10 | 0.57 | 0.59 | 0.55 | 2.86 | 3.07 | 2.89 | 1.58 | 1.61 | 1.84 | 1.38 | 1.27 | 1.50 | Small Molecule Transport (Biocarbonate) | Fig 1i |
| SLC6A20 | 0.03 | 0.02 | 0.04 | 0.64 | 0.16 | 0.17 | 1.09 | 1.78 | 0.45 | 1.33 | 0.37 | 0.66 | Small Molecule Transport (Proline) | Fig 1i |
| SLC6A17 | 0.19 | 0.23 | 0.34 | 5.82 | 7.27 | 11.1 | 13.2 | 5.76 | 11.4 | 2.86 | 2.35 | 11.2 | Small Molecule Transport (Pro, Cys, Ala) | Fig 1i |
| SLC6A7 | 0.01 | 0.00 | 0.00 | 0.03 | 0.07 | 0.03 | 0.01 | 0.00 | 0.00 | 0.01 | 0.03 | 0.02 | Small Molecule Transport (Proline) | Fig 1i |
| SLC46A3 | 0.50 | 0.56 | 0.69 | 4.29 | 4.46 | 3.94 | 7.33 | 9.21 | 8.62 | 8.13 | 9.30 | 8.77 | Small Molecule Transport (Folate) | Fig 1i |
| FOLR1 | 0.09 | 0.33 | 0.31 | 0.12 | 0.07 | 0.08 | 1.67 | 1.14 | 0.73 | 2.61 | 1.58 | 1.15 | Small Molecule Transport (Folate) | Fig 1i |
| SLC47A1 | 0.40 | 0.38 | 0.39 | 0.29 | 0.17 | 0.00 | 0.63 | 0.66 | 0.66 | 0.79 | 0.37 | 0.70 | Small Molecule Transport (Thiamine) | Fig 1i |
| SLC13A2 | 0.00 | 0.00 | 0.01 | 0.04 | 0.03 | 0.03 | 0.50 | 1.95 | 1.53 | 1.15 | 1.05 | 0.79 | Small Molecule Transport (Citrate, Succinate) | Fig 1i |
| SLC22A8 | 0.01 | 0.00 | 0.00 | 0.05 | 0.08 | 0.18 | 1.25 | 1.58 | 0.98 | 0.71 | 0.53 | 0.36 | Small Molecule Transport (Urate, aKG) | Fig 1i |
| SVOP | 0.57 | 0.82 | 0.85 | 10.1 | 11.1 | 11.8 | 17.1 | 17.0 | 19.2 | 13.4 | 16.7 | 17.1 | Small Molecule Transport | Fig 1i |
| SLC7A14 | 0.13 | 0.12 | 0.13 | 0.47 | 2.24 | 0.92 | 1.77 | 2.39 | 1.69 | 1.87 | 2.25 | 2.14 | Small Molecule Transport (Amino acid) | Fig 1i |
| SLC16A12 | 0.03 | 0.09 | 0.09 | 0.19 | 0.12 | 0.12 | 0.61 | 1.53 | 0.56 | 1.15 | 1.11 | 0.80 | Small Molecule Transport (Creatine) | Fig 1i |
| SLC25A45 | 0.03 | 0.01 | 0.01 | 0.04 | 0.08 | 0.11 | 0.17 | 0.35 | 0.12 | 0.13 | 0.10 | 0.12 | Small Molecule Transport (Amino acid) | Fig 1i |
| SLC5A5 | 0.02 | 0.04 | 0.01 | 0.06 | 0.05 | 0.05 | 0.05 | 0.00 | 0.02 | 0.28 | 0.05 | 0.01 | Small Molecule Transport (iodide) | Fig 1i |
| SLC7A5 | 7.34 | 6.97 | 5.90 | 7.05 | 10.1 | 9.45 | 25.2 | 15.1 | 13.8 | 19.2 | 11.3 | 11.5 | Small Molecule Transport (Unkown) | Fig 1i |
| SLC4A1 | 0.01 | 0.01 | 0.02 | 0.07 | 0.06 | 0.01 | 0.03 | 0.02 | 0.03 | 0.03 | 4.8E-03 | 0.05 | Small Molecule Transport (Anion) | Fig 1i |
| RHBG | 0.00 | 0.00 | 0.02 | 0.02 | 0.03 | 0.00 | 0.03 | 0.05 | 0.00 | 0.00 | 0.03 | 0.00 | Small Molecule Transport (Ammonium) | Fig 1i |
| SLC6A19 | 0.02 | 0.01 | 0.01 | 0.16 | 0.15 | 0.14 | 0.10 | 0.03 | 0.13 | 0.02 | 0.02 | 0.02 | Small Molecule Transport (BCAA) | Fig 1i |
| SLC44A3 | 0.94 | 0.98 | 1.09 | 1.03 | 0.79 | 0.77 | 1.68 | 1.71 | 1.28 | 2.29 | 1.00 | 1.46 | Small Molecule Transport (Choline) | Fig 1i |
| SLC7A2 | 2.67 | 2.27 | 2.43 | 0.98 | 0.58 | 0.69 | 2.25 | 2.92 | 2.77 | 4.01 | 3.11 | 2.71 | Small Molecule Transport (Arg, Lys, Orn, His) | Fig 1i |
| SLC22A15 | 0.57 | 0.75 | 0.95 | 1.43 | 0.86 | 0.72 | 0.78 | 0.91 | 0.62 | 0.86 | 0.73 | 0.81 | Small Molecule Transport (Organic Cation) | Fig 1i |
| SLC16A3 | 22.2 | 18.4 | 16.4 | 18.5 | 8.70 | 8.04 | 31.0 | 12.2 | 11.1 | 19.8 | 9.57 | 10.2 | Small Molecule Transport (Lactate/Pyruvate) | Fig 1i |
| SLCO5A1 | 1.62 | 1.88 | 1.92 | 6.49 | 3.51 | 3.65 | 2.15 | 1.11 | 0.88 | 0.97 | 1.06 | 1.23 | Small Molecule Transport (peptide) | Fig 1i |
| SLC18A3 | 0.54 | 0.76 | 0.48 | 0.11 | 1.16 | 0.18 | 0.64 | 0.18 | 0.28 | 0.53 | 0.11 | 0.41 | Small Molecule Transport (Acetylcholine, Choline) | Fig 1i |
| SLC25A21 | 0.71 | 0.52 | 0.54 | 0.23 | 0.00 | 0.11 | 0.30 | 0.59 | 0.50 | 0.29 | 0.20 | 0.24 | Small Molecule Transport (aKG) | Fig 1i |
| SLC7A10 | 0.32 | 0.49 | 0.47 | 0.51 | 0.51 | 0.50 | 0.22 | 0.18 | 0.13 | 0.24 | 0.13 | 0.14 | Small Molecule Transport (Amino acid) | Fig 1i |
| SLC6A2 | 0.07 | 0.09 | 0.22 | 0.49 | 0.07 | 0.02 | 0.02 | 0.00 | 0.05 | 0.01 | 0.03 | 0.04 | Small Molecule Transport (Monoamine) | Fig 1i |
| SLC34A2 | 0.44 | 0.37 | 0.42 | 0.05 | 0.08 | 0.01 | 0.18 | 0.10 | 0.08 | 0.12 | 0.03 | 0.07 | Small Molecule Transport (Mitochondrial Phosphate) | Fig 1i |
| SLC6A3 | 0.39 | 0.33 | 0.32 | 0.27 | 0.15 | 0.19 | 0.09 | 0.08 | 0.08 | 0.06 | 0.04 | 0.01 | Small Molecule Transport (Monoamine) | Fig 1i |
| SLC38A4 | 3.82 | 3.02 | 3.12 | 2.13 | 0.48 | 0.67 | 0.07 | 0.14 | 0.07 | 0.03 | 0.05 | 0.04 | Small Molecule Transport (His, Ar, Alan, Asn) | Fig 1i |
| GAD1 | 2.22 | 1.88 | 2.01 | 3.96 | 6.16 | 4.60 | 10.7 | 13.3 | 14.0 | 9.42 | 11.5 | 13.8 | Amino Acid (Glutamate) | Fig S1b |
| GAD2 | 2.04 | 2.52 | 2.71 | 2.28 | 3.37 | 2.09 | 5.22 | 3.35 | 2.74 | 5.00 | 5.95 | 6.00 | Amino Acid (Glutamate) | Fig S1b |
| DDO | 0.02 | 0.02 | 0.00 | 0.00 | 0.00 | 0.00 | 0.35 | 0.47 | 0.52 | 0.54 | 0.26 | 0.57 | Amino Acid (Aspartate) | Fig S1b |
| GADL1 | 0.01 | 0.01 | 0.01 | 0.03 | 0.08 | 0.00 | 0.00 | 0.02 | 0.00 | 0.01 | 0.01 | 0.01 | Amino Acid (Glutamate) | Fig S1b |
| ACY3 | 0.00 | 0.00 | 0.00 | 0.00 | 0.14 | 0.05 | 0.02 | 0.00 | 0.00 | 0.00 | 0.02 | 0.00 | Amino Acid (Aspartate) | Fig S1b |
| CRYM | 8.84 | 11.1 | 10.8 | 9.70 | 9.77 | 7.75 | 36.2 | 56.8 | 45.8 | 49.8 | 28.6 | 42.5 | Amino Acid (Cystathionine) | Fig S1b |
| CSAD | 3.92 | 4.04 | 3.87 | 4.82 | 4.67 | 4.59 | 5.21 | 4.93 | 5.45 | 5.18 | 4.65 | 5.02 | Amino Acid (Cysteine) | Fig S1b |
| CDO1 | 20.3 | 18.9 | 19.4 | 14.2 | 17.3 | 10.4 | 22.0 | 23.4 | 16.7 | 27.5 | 12.8 | 16.5 | Amino Acid (Cysteine) | Fig S1b |
| ADO | 9.47 | 8.66 | 9.15 | 9.54 | 9.73 | 10.4 | 9.62 | 9.40 | 9.17 | 9.70 | 8.36 | 7.92 | Amino Acid (Cysteine) | Fig S1b |
| GPX3 | 8.79 | 15.5 | 14.9 | 7.33 | 10.4 | 8.26 | 205 | 412 | 299 | 336 | 324 | 217 | Amino Acid (Glutathione) | Fig S1b |
| MGST1 | 7.93 | 8.96 | 8.50 | 1.64 | 3.89 | 1.60 | 5.63 | 3.48 | 5.61 | 10.3 | 2.00 | 4.52 | Amino Acid (Glutathione) | Fig S1b |
| GPX6 | 0.02 | 0.02 | 0.04 | 0.14 | 0.21 | 0.43 | 0.03 | 0.00 | 0.00 | 0.03 | 0.02 | 0.00 | Amino Acid (Glutathione) | Fig S1b |
| GPX8 | 12.2 | 11.7 | 11.1 | 3.36 | 1.61 | 1.23 | 3.77 | 3.05 | 2.90 | 3.46 | 1.99 | 2.54 | Amino Acid (Glutathione) | Fig S1b |
| GPX5 | 0.04 | 0.05 | 0.10 | 0.41 | 0.60 | 0.78 | 0.05 | 0.00 | 0.00 | 0.00 | 0.02 | 0.02 | Amino Acid (Glutathione) | Fig S1b |
| HNMT | 0.08 | 0.04 | 0.06 | 0.11 | 0.46 | 0.04 | 0.68 | 1.00 | 0.67 | 1.60 | 0.16 | 0.84 | Amino Acid (histidine) | Fig S1b |
| HDC | 0.09 | 0.10 | 0.07 | 0.01 | 0.00 | 0.02 | 0.04 | 0.19 | 0.13 | 0.08 | 0.05 | 0.14 | Amino Acid (histidine) | Fig S1b |
| AOC3 | 1.08 | 0.91 | 1.08 | 1.04 | 1.50 | 1.57 | 3.61 | 10.0 | 5.30 | 2.78 | 11.4 | 4.76 | Amino acid (Dopamine, histamine) | Fig S1b |
| FTCD | 0.36 | 0.25 | 0.42 | 0.45 | 0.90 | 0.59 | 0.22 | 0.23 | 0.23 | 0.29 | 0.26 | 0.26 | Amino Acid (histidine) | Fig S1b |
| UROC1 | 0.05 | 0.04 | 0.03 | 0.02 | 0.12 | 0.15 | 0.04 | 0.03 | 0.05 | 0.01 | 0.03 | 0.03 | Amino Acid (histidine) | Fig S1b |
| NOS1 | 0.03 | 0.04 | 0.02 | 0.01 | 0.01 | 0.01 | 0.14 | 0.27 | 0.28 | 0.09 | 0.10 | 0.13 | Amino Acid (Arginine) | Fig S1b |
| MOSC2 | 1.65 | 1.39 | 2.01 | 1.62 | 1.38 | 1.37 | 1.72 | 1.88 | 2.52 | 2.59 | 1.62 | 1.91 | Amino Acid (Arginine) | Fig S1b |
| CPS1 | 11.3 | 10.5 | 9.99 | 1.77 | 1.09 | 0.88 | 1.84 | 1.21 | 1.68 | 1.42 | 0.73 | 1.07 | Amino Acid (Urea) | Fig S1b |
| ARG1 | 0.13 | 0.15 | 0.14 | 0.09 | 0.21 | 0.42 | 0.06 | 0.00 | 0.06 | 0.09 | 0.11 | 0.08 | Amino Acid (Arginine,Urea) | Fig S1b |
| NOS2 | 0.15 | 0.12 | 0.08 | 0.06 | 0.09 | 0.04 | 0.06 | 0.08 | 0.02 | 0.08 | 0.09 | 0.07 | Amino Acid (Arginine) | Fig S1b |
| TYRP1 | 0.96 | 0.80 | 1.76 | 4.34 | 1.45 | 1.20 | 25.7 | 23.8 | 11.5 | 47.0 | 7.54 | 13.0 | Amino Acid (Tryosine, Pigment) | Fig S1b |
| TYR | 0.66 | 0.78 | 0.93 | 1.54 | 0.08 | 0.19 | 2.56 | 1.80 | 0.76 | 5.50 | 0.77 | 1.23 | Amino Acid (Tryosine, Pigment) | Fig S1b |
| DCT | 15.2 | 14.3 | 17.3 | 56.5 | 38.6 | 54.1 | 9.46 | 7.84 | 4.03 | 63.8 | 2.93 | 6.39 | Amino Acid (Tryosine, Pigment) | Fig S1b |
| HPD | 2.45 | 3.71 | 3.34 | 0.00 | 0.04 | 0.05 | 0.09 | 0.09 | 0.13 | 0.19 | 0.11 | 0.04 | Amino Acid (Tyrosine) | Fig S1b |
| TH | 0.85 | 0.71 | 0.87 | 0.49 | 0.29 | 0.62 | 0.23 | 0.10 | 0.00 | 0.07 | 0.01 | 0.08 | Amino Acid (Neurotransmitter) | Fig S1b |
| AGXT | 0.00 | 0.04 | 0.04 | 0.00 | 0.00 | 0.00 | 5.48 | 0.22 | 0.39 | 0.60 | 0.00 | 0.00 | Amino Acid (Alanine) | Fig S1b |
| DAO | 0.02 | 0.00 | 0.02 | 0.00 | 0.04 | 0.08 | 0.09 | 0.06 | 0.11 | 0.08 | 0.12 | 0.02 | Amino Acid (Alanine) | Fig S1b |
| AGXT2L1 | 0.87 | 0.76 | 0.87 | 0.26 | 0.11 | 0.09 | 0.46 | 0.69 | 0.57 | 0.66 | 0.76 | 0.86 | Amino Acid (Alanine) | Fig S1b |
| KMO | 0.13 | 0.31 | 0.29 | 0.04 | 0.03 | 0.02 | 0.02 | 0.02 | 0.02 | 0.03 | 0.01 | 0.02 | Amino Acid (Tryptophan) | Fig S1b |
| KYNU | 0.00 | 0.00 | 0.00 | 0.00 | 0.05 | 0.00 | 0.02 | 0.00 | 0.07 | 0.05 | 0.05 | 0.02 | Amino Acid (Tryptophan) | Fig S1b |
| TPH2 | 0.08 | 0.06 | 0.06 | 0.05 | 0.02 | 0.03 | 0.00 | 0.00 | 0.00 | 0.00 | 0.01 | 0.00 | Amino Acid (Neurotransmitter) | Fig S1b |
| ASPG | 0.00 | 0.00 | 0.00 | 0.00 | 0.18 | 0.10 | 0.00 | 0.00 | 0.00 | 0.02 | 0.01 | 0.01 | Amino Acid (Asparagine) | Fig S1b |
| BCAT1 | 24.6 | 22.1 | 19.7 | 3.19 | 3.40 | 2.27 | 7.39 | 3.06 | 3.52 | 3.99 | 2.01 | 2.97 | Amino Acid (BCAAs) | Fig S1b |
| CKMT1B | 0.23 | 0.42 | 0.57 | 3.58 | 5.01 | 5.09 | 9.80 | 17.2 | 17.3 | 12.5 | 18.8 | 12.4 | Amino Acid (Creatine) | Fig S1b |
| CKM | 0.11 | 0.08 | 0.14 | 0.15 | 0.47 | 0.85 | 0.52 | 0.48 | 0.38 | 0.47 | 0.62 | 0.24 | Amino Acid (Creatine) | Fig S1b |
| BHMT | 7.09 | 47.4 | 32.2 | 0.09 | 0.09 | 0.03 | 0.03 | 0.03 | 0.10 | 0.07 | 0.06 | 0.10 | Amino Acid (Glycine) | Fig S1b |
| GLDC | 8.88 | 8.49 | 8.36 | 5.27 | 4.45 | 4.46 | 6.55 | 5.07 | 5.10 | 5.29 | 3.29 | 3.39 | Amino Acid (Glycine) | Fig S1b |
| LGSN | 0.04 | 0.00 | 0.02 | 0.39 | 0.38 | 0.01 | 0.02 | 0.05 | 0.00 | 0.01 | 0.03 | 0.01 | Amino Acid (Glutamine) | Fig S1b |
| PIPOX | 1.45 | 1.33 | 1.27 | 3.09 | 3.47 | 3.04 | 3.25 | 2.33 | 3.75 | 3.81 | 2.26 | 2.82 | Amino Acid (Lysine) | Fig S1b |
| MSRB3 | 2.29 | 2.86 | 2.90 | 1.14 | 1.17 | 1.23 | 3.77 | 4.80 | 3.71 | 4.18 | 3.14 | 2.92 | Amino Acid (Methionine) | Fig S1b |
| MAOA | 1.32 | 1.27 | 1.59 | 20.1 | 18.5 | 19.7 | 16.1 | 16.4 | 15.8 | 19.9 | 17.3 | 11.2 | Amino Acid (Neurotransmiter) | Fig S1b |
| PNMT | 0.44 | 0.48 | 0.65 | 1.12 | 1.38 | 1.60 | 0.78 | 0.49 | 0.66 | 0.60 | 0.61 | 0.47 | Amino Acid (Neurotransmitter) | Fig S1b |
| PRODH2 | 0.06 | 0.04 | 0.10 | 0.10 | 0.07 | 0.16 | 0.11 | 0.05 | 0.05 | 0.03 | 0.09 | 0.13 | Amino Acid (Proline) | Fig S1b |
| PHGDH | 27.8 | 23.5 | 25.2 | 15.5 | 10.7 | 8.00 | 20.9 | 8.04 | 7.51 | 14.8 | 5.52 | 8.29 | Amino Acid (Serine) | Fig S1b |
| NAT2 | 0.02 | 0.00 | 0.00 | 0.00 | 0.00 | 0.00 | 0.07 | 0.00 | 0.14 | 0.14 | 0.02 | 0.07 | Amino Acid (Acetylation) | Fig S1b |
| FUT3 | 1.00 | 0.80 | 0.88 | 0.97 | 2.34 | 2.40 | 1.63 | 2.73 | 2.69 | 3.30 | 2.65 | 3.38 | Glycan | Fig S1c |
| B3GALT5 | 0.21 | 0.10 | 0.24 | 0.15 | 0.20 | 0.17 | 0.69 | 0.55 | 0.58 | 0.66 | 0.37 | 0.81 | Glycan | Fig S1c |
| NEU4 | 0.18 | 0.12 | 0.12 | 0.06 | 0.09 | 0.13 | 0.48 | 0.82 | 0.74 | 0.63 | 0.37 | 0.35 | Glycan (Degradation) | Fig S1c |
| B3GALT2 | 0.94 | 0.96 | 0.88 | 4.05 | 3.65 | 3.16 | 3.14 | 3.86 | 2.47 | 2.76 | 3.02 | 2.32 | Glycan | Fig S1c |
| CSGALNACT1 | 0.41 | 0.55 | 0.62 | 0.70 | 0.76 | 0.41 | 2.50 | 1.98 | 1.69 | 2.41 | 0.67 | 1.07 | Glycan (Sulfate) | Fig S1c |
| B3GAT2 | 1.95 | 1.70 | 1.76 | 4.55 | 3.51 | 3.77 | 5.19 | 4.84 | 5.22 | 5.33 | 3.63 | 4.01 | Glycan | Fig S1c |
| FUT6 | 0.09 | 0.12 | 0.18 | 0.27 | 0.39 | 0.38 | 0.15 | 0.15 | 0.18 | 0.36 | 0.23 | 0.30 | Glycan | Fig S1c |
| FUT7 | 0.07 | 0.03 | 0.08 | 0.15 | 0.28 | 0.40 | 0.12 | 0.07 | 0.04 | 0.10 | 0.16 | 0.13 | Glycan | Fig S1c |
| B3GNT4 | 0.29 | 0.30 | 0.30 | 0.49 | 0.61 | 0.72 | 0.46 | 0.18 | 0.30 | 0.44 | 0.50 | 0.42 | Glycan | Fig S1c |
| ST8SIA3 | 0.94 | 0.86 | 0.75 | 1.44 | 2.42 | 1.33 | 1.55 | 1.42 | 0.88 | 1.60 | 1.02 | 1.16 | Glycan | Fig S1c |
| MGAT4C | 0.32 | 0.65 | 0.52 | 0.56 | 0.58 | 0.24 | 0.33 | 0.32 | 0.27 | 0.86 | 0.53 | 0.37 | Glycan | Fig S1c |
| CHST6 | 0.40 | 0.42 | 0.41 | 0.26 | 0.17 | 0.18 | 0.75 | 0.22 | 0.37 | 1.03 | 0.14 | 0.22 | Glycan (Sulfate) | Fig S1c |
| ST8SIA4 | 1.65 | 1.32 | 0.98 | 2.88 | 1.95 | 1.45 | 2.77 | 1.49 | 1.68 | 2.39 | 0.87 | 0.91 | Glycan | Fig S1c |
| FUT5 | 0.86 | 0.87 | 1.02 | 0.59 | 1.34 | 1.09 | 0.34 | 0.28 | 0.44 | 0.47 | 0.47 | 0.57 | Glycan | Fig S1c |
| GALNT5 | 0.29 | 0.17 | 0.26 | 0.06 | 0.09 | 0.00 | 0.10 | 0.16 | 0.08 | 0.19 | 0.09 | 0.11 | Glycan | Fig S1c |
| GLB1L3 | 0.03 | 0.04 | 0.04 | 0.10 | 0.08 | 0.20 | 0.06 | 0.16 | 0.12 | 4.6E-03 | 0.04 | 0.01 | Glycan (Degradation) | Fig S1c |
| HAS1 | 0.87 | 0.75 | 1.24 | 0.25 | 0.95 | 0.87 | 0.55 | 0.29 | 0.25 | 0.28 | 0.31 | 0.34 | Glycan | Fig S1c |
| MFNG | 3.04 | 2.56 | 3.55 | 4.78 | 2.13 | 2.44 | 1.53 | 0.20 | 0.32 | 0.36 | 0.43 | 0.40 | Glycan | Fig S1c |
| HAS2 | 27.4 | 22.3 | 23.3 | 0.80 | 0.41 | 0.51 | 2.81 | 3.44 | 2.13 | 1.77 | 0.80 | 2.48 | Glycan | Fig S1c |
| ENO2 | 83.6 | 74.5 | 76.1 | 203 | 188 | 159 | 369 | 318 | 320 | 368 | 422 | 264 | Glycolysis | Fig S1c |
| ALDOC | 33.2 | 25.7 | 26.0 | 50.7 | 50.6 | 38.6 | 153 | 90.1 | 105 | 139 | 136 | 92.6 | Glycolysis | Fig S1c |
| PFKP | 28.6 | 24.1 | 24.8 | 53.3 | 56.5 | 49.7 | 99.4 | 55.1 | 57.1 | 64.1 | 68.2 | 53.0 | Glycolysis | Fig S1c |
| PKLR | 0.36 | 0.57 | 0.34 | 0.95 | 0.87 | 0.66 | 0.43 | 0.44 | 0.39 | 0.54 | 0.37 | 0.55 | Glycolysis | Fig S1c |
| GCK | 0.74 | 0.93 | 0.94 | 0.96 | 0.58 | 0.62 | 1.93 | 0.80 | 0.43 | 1.48 | 0.20 | 0.36 | Glycolysis | Fig S1c |
| PFKFB4 | 30.9 | 22.7 | 17.5 | 19.3 | 8.99 | 7.74 | 26.9 | 13.5 | 12.1 | 13.4 | 13.1 | 14.6 | Glycolysis | Fig S1c |
| ABCA4 | 0.43 | 0.37 | 0.56 | 18.6 | 30.9 | 27.1 | 45.6 | 55.2 | 65.9 | 56.5 | 85.3 | 48.0 | ABC Transporter | Fig S1d |
| ABCG4 | 0.25 | 0.25 | 0.25 | 8.20 | 7.97 | 9.14 | 15.9 | 18.6 | 20.3 | 16.9 | 23.5 | 15.0 | ABC Transporter | Fig S1d |
| ABCA6 | 4.6E-03 | 0.00 | 0.01 | 0.01 | 0.01 | 0.01 | 0.14 | 0.21 | 0.20 | 0.12 | 0.21 | 0.20 | ABC Transporter | Fig S1d |
| ABCA8 | 0.05 | 0.07 | 0.08 | 0.08 | 0.10 | 0.15 | 1.70 | 4.74 | 2.41 | 1.67 | 1.75 | 2.42 | ABC Transporter | Fig S1d |
| ABCA7 | 2.58 | 3.03 | 2.99 | 18.4 | 14.9 | 16.8 | 36.9 | 41.2 | 44.7 | 37.7 | 45.3 | 39.2 | ABC Transporter | Fig S1d |
| ABCG1 | 1.59 | 2.25 | 2.31 | 9.50 | 14.5 | 15.8 | 12.7 | 12.2 | 13.1 | 12.6 | 12.7 | 10.8 | ABC Transporter | Fig S1d |
| ABCC8 | 0.13 | 0.14 | 0.25 | 0.36 | 0.52 | 0.27 | 0.53 | 0.33 | 0.34 | 0.32 | 0.58 | 0.32 | ABC Transporter | Fig S1d |
| ABCB11 | 5.13 | 4.41 | 4.21 | 0.41 | 0.26 | 0.23 | 0.06 | 0.00 | 0.09 | 0.03 | 0.05 | 0.04 | ABC Transporter | Fig S1d |
| ACSL6 | 0.22 | 0.25 | 0.23 | 12.1 | 17.2 | 17.6 | 24.3 | 31.5 | 36.1 | 26.5 | 30.4 | 27.5 | Lipid (Fatty Acid) | Fig S1d |
| ECHDC2 | 0.29 | 0.14 | 0.15 | 5.74 | 7.10 | 8.63 | 6.89 | 11.2 | 12.3 | 13.6 | 11.7 | 8.41 | Lipid (Fatty Acid) | Fig S1d |
| MGLL | 1.00 | 0.95 | 0.99 | 3.27 | 4.54 | 4.27 | 5.72 | 4.16 | 4.48 | 5.36 | 4.29 | 4.38 | Lipid (Fatty Acid) | Fig S1d |
| ACSL1 | 2.30 | 2.82 | 2.89 | 8.10 | 9.80 | 9.68 | 12.3 | 12.9 | 13.2 | 12.0 | 13.5 | 11.9 | Lipid (Fatty Acid) | Fig S1d |
| ACOX2 | 0.25 | 0.44 | 0.49 | 0.12 | 0.10 | 0.25 | 1.47 | 2.00 | 1.67 | 1.79 | 1.22 | 1.49 | Lipid (Fatty Acid) | Fig S1d |
| ELOVL7 | 0.25 | 0.23 | 0.19 | 0.15 | 0.12 | 0.13 | 0.55 | 0.79 | 0.52 | 0.59 | 0.38 | 0.60 | Lipid (Fatty Acid) | Fig S1d |
| ABHD1 | 0.15 | 0.10 | 0.09 | 0.09 | 0.04 | 0.38 | 0.10 | 0.13 | 0.06 | 0.25 | 0.15 | 0.18 | Lipid (Fatty Acid) | Fig S1d |
| SDR9C7 | 0.15 | 0.04 | 0.09 | 0.16 | 0.13 | 0.11 | 0.07 | 0.00 | 0.10 | 0.02 | 0.13 | 0.11 | Lipid (Fatty Acid) | Fig S1d |
| CPT1A | 7.78 | 9.98 | 9.68 | 6.13 | 7.77 | 7.87 | 7.25 | 8.59 | 8.39 | 7.32 | 9.31 | 7.16 | Lipid (Fatty Acid) | Fig S1d |
| FAR2 | 9.10 | 7.03 | 7.17 | 3.65 | 1.08 | 1.36 | 3.29 | 2.06 | 2.31 | 2.56 | 1.53 | 2.05 | Lipid (Fatty Acid) | Fig S1d |
| ACAT2 | 20.8 | 18.5 | 20.0 | 15.6 | 8.38 | 7.34 | 7.90 | 6.19 | 5.17 | 7.04 | 4.11 | 3.91 | Lipid (Fatty Acid) | Fig S1d |
| LIPC | 0.28 | 0.48 | 0.34 | 0.08 | 0.00 | 0.00 | 0.04 | 0.00 | 0.00 | 0.00 | 0.00 | 0.05 | Lipid (Fatty Acid) | Fig S1d |
| PTGDS | 0.65 | 1.62 | 1.12 | 8.16 | 7.88 | 3.19 | 170 | 151 | 96.9 | 179 | 122 | 125 | Lipid (Hormone) | Fig S1d |
| DIO2 | 0.35 | 0.21 | 0.24 | 0.45 | 0.65 | 0.67 | 3.79 | 6.41 | 4.59 | 8.08 | 4.37 | 6.40 | Lipid (Hormone) | Fig S1d |
| AANAT | 0.32 | 0.27 | 0.35 | 5.00 | 6.27 | 6.95 | 10.4 | 6.76 | 5.45 | 6.22 | 4.91 | 3.93 | Lipid (Hormone) | Fig S1d |
| ACE | 0.09 | 0.13 | 0.23 | 0.38 | 0.67 | 0.57 | 0.77 | 0.59 | 0.80 | 0.71 | 0.71 | 0.55 | Lipid (Hormone) | Fig S1d |
| ALOX15 | 0.28 | 0.09 | 0.15 | 0.16 | 0.08 | 0.12 | 4.97 | 1.16 | 0.85 | 1.72 | 0.19 | 0.07 | Lipid (Hormone) | Fig S1d |
| ALOXE3 | 0.08 | 0.03 | 0.05 | 0.13 | 0.13 | 0.08 | 0.11 | 0.07 | 0.03 | 0.08 | 0.11 | 0.11 | Lipid (Hormone) | Fig S1d |
| PTGS1 | 0.21 | 0.26 | 0.19 | 0.64 | 1.30 | 1.33 | 0.32 | 0.18 | 0.16 | 0.20 | 0.11 | 0.25 | Lipid (Hormone) | Fig S1d |
| PTGS2 | 3.35 | 0.76 | 0.46 | 2.06 | 1.00 | 1.24 | 2.37 | 1.05 | 0.78 | 1.23 | 0.59 | 0.71 | Lipid (Hormone) | Fig S1d |
| MME | 0.72 | 1.05 | 0.99 | 0.24 | 0.47 | 0.65 | 0.39 | 0.40 | 0.35 | 0.16 | 0.10 | 0.34 | Lipid (Hormone) | Fig S1d |
| TPO | 0.17 | 0.16 | 0.23 | 0.21 | 0.00 | 0.04 | 0.08 | 0.03 | 0.06 | 0.00 | 0.05 | 0.05 | Lipid (Hormone) | Fig S1d |
| PLA2G5 | 0.01 | 0.00 | 0.00 | 0.06 | 0.25 | 0.14 | 2.77 | 5.07 | 5.97 | 4.48 | 6.13 | 3.32 | Lipid (Membrane Lipid) | Fig S1d |
| PLD4 | 0.00 | 0.00 | 0.02 | 0.58 | 0.82 | 0.68 | 2.69 | 1.44 | 2.44 | 2.31 | 0.79 | 1.49 | Lipid (Membrane Lipid) | Fig S1d |
| PHOSPHO1 | 0.05 | 0.02 | 0.06 | 0.62 | 1.03 | 0.80 | 3.32 | 3.21 | 3.28 | 3.61 | 4.01 | 2.33 | Lipid (Membrane Lipid) | Fig S1d |
| PLCXD3 | 0.04 | 0.04 | 0.02 | 0.39 | 0.60 | 0.31 | 1.06 | 1.37 | 1.17 | 1.96 | 0.66 | 0.96 | Lipid (Membrane Lipid) | Fig S1d |
| PLA2G4C | 0.40 | 0.55 | 0.47 | 2.86 | 6.73 | 5.53 | 11.9 | 10.3 | 10.1 | 11.4 | 12.2 | 12.3 | Lipid (Membrane Lipid) | Fig S1d |
| LPPR4 | 0.38 | 0.49 | 0.46 | 9.78 | 7.42 | 8.42 | 5.78 | 8.96 | 6.00 | 8.65 | 6.15 | 6.26 | Lipid (Membrane Lipid) | Fig S1d |
| GDPD5 | 1.48 | 1.91 | 2.20 | 14.5 | 17.1 | 15.2 | 24.3 | 24.8 | 27.4 | 22.8 | 28.5 | 22.6 | Lipid (Membrane Lipid) | Fig S1d |
| PLCH2 | 3.92 | 4.04 | 4.67 | 22.7 | 27.1 | 28.7 | 42.7 | 42.6 | 49.0 | 32.9 | 40.4 | 36.4 | Lipid (Membrane Lipid) | Fig S1d |
| DGKB | 0.21 | 0.18 | 0.24 | 0.11 | 0.22 | 0.15 | 0.28 | 0.26 | 0.44 | 0.99 | 0.20 | 0.40 | Lipid (Membrane Lipid) | Fig S1d |
| PLD1 | 0.37 | 0.45 | 0.56 | 0.33 | 0.16 | 0.21 | 0.78 | 0.86 | 0.45 | 0.84 | 0.63 | 0.65 | Lipid (Membrane Lipid) | Fig S1d |
| PLA2G1B | 0.00 | 0.26 | 0.08 | 0.30 | 0.15 | 0.33 | 0.00 | 0.00 | 0.22 | 0.00 | 0.20 | 0.28 | Lipid (Membrane Lipid) | Fig S1d |
| PLA2G2A | 0.00 | 0.24 | 0.08 | 0.00 | 0.15 | 0.00 | 0.23 | 0.31 | 0.35 | 0.10 | 0.00 | 0.23 | Lipid (Membrane Lipid) | Fig S1d |
| DGKG | 0.46 | 1.19 | 0.95 | 0.07 | 0.10 | 0.02 | 0.96 | 0.68 | 0.38 | 1.12 | 0.29 | 0.63 | Lipid (Membrane Lipid) | Fig S1d |
| PLCG2 | 0.47 | 0.44 | 0.48 | 0.11 | 0.03 | 0.04 | 1.04 | 0.52 | 0.85 | 0.64 | 0.09 | 0.10 | Lipid (Membrane Lipid) | Fig S1d |
| PLCL1 | 1.20 | 1.72 | 1.57 | 1.54 | 2.14 | 1.61 | 0.65 | 0.82 | 0.60 | 0.59 | 0.66 | 0.58 | Lipid (Membrane Lipid) | Fig S1d |
| DGKK | 0.14 | 0.15 | 0.21 | 0.09 | 0.11 | 0.08 | 0.05 | 0.06 | 0.00 | 0.02 | 0.02 | 0.02 | Lipid (Membrane Lipid) | Fig S1d |
| PLA2G3 | 1.23 | 0.81 | 0.73 | 0.33 | 0.22 | 0.07 | 0.07 | 0.03 | 0.00 | 0.03 | 0.01 | 0.03 | Lipid (Membrane Lipid) | Fig S1d |
| SMPD3 | 1.49 | 1.82 | 1.84 | 5.86 | 7.13 | 5.32 | 5.63 | 6.33 | 5.85 | 5.18 | 5.27 | 4.47 | Lipid (Sphingolipid) | Fig S1d |
| ASAH2 | 3.88 | 2.57 | 2.06 | 2.21 | 0.95 | 0.70 | 5.38 | 7.40 | 5.50 | 6.68 | 4.68 | 5.67 | Lipid (Sphingosine) | Fig S1d |
| CERKL | 8.22 | 6.79 | 7.88 | 15.8 | 10.4 | 15.9 | 13.6 | 17.2 | 17.4 | 15.6 | 17.1 | 12.9 | Lipid (Sphingolipid) | Fig S1d |
| SPHK1 | 44.1 | 45.1 | 48.3 | 1.23 | 2.27 | 3.27 | 1.93 | 2.21 | 1.51 | 1.92 | 1.98 | 2.48 | Lipid (Sphingolipid) | Fig S1d |
| CYP27A1 | 1.15 | 0.94 | 1.00 | 3.58 | 2.64 | 1.96 | 11.2 | 7.45 | 7.44 | 12.3 | 8.52 | 8.95 | Lipid (Bile Acid) | Fig S1d |
| HSD17B3 | 0.05 | 0.07 | 0.21 | 0.31 | 0.45 | 0.69 | 0.94 | 0.84 | 0.84 | 1.02 | 0.52 | 1.54 | Lipid (Steroid) | Fig S1d |
| KL | 0.02 | 0.03 | 0.03 | 0.20 | 0.32 | 0.29 | 0.34 | 0.26 | 0.15 | 0.14 | 0.06 | 0.11 | Lipid (Steroid) | Fig S1d |
| CYP2E1 | 0.75 | 1.08 | 1.19 | 2.37 | 3.09 | 3.64 | 2.16 | 2.24 | 2.73 | 3.36 | 3.51 | 2.71 | Lipid (Steroid) | Fig S1d |
| CYP19A1 | 0.00 | 0.04 | 0.01 | 0.00 | 0.00 | 0.01 | 0.01 | 0.06 | 0.00 | 0.05 | 0.06 | 0.03 | Lipid (Steroid) | Fig S1d |
| HSD17B2 | 6.91 | 7.15 | 7.43 | 1.66 | 2.52 | 0.24 | 6.58 | 11.5 | 4.33 | 7.93 | 4.58 | 5.73 | Lipid (Steroid) | Fig S1d |
| CYP2W1 | 0.07 | 0.13 | 0.20 | 0.09 | 0.42 | 0.31 | 0.22 | 0.07 | 0.07 | 0.09 | 0.11 | 0.07 | Lipid (Steroid) | Fig S1d |
| CYP3A43 | 0.19 | 0.08 | 0.15 | 0.04 | 0.09 | 0.13 | 0.05 | 0.04 | 0.13 | 0.06 | 0.06 | 0.11 | Lipid (Steroid) | Fig S1d |
| DHCR24 | 16.8 | 12.1 | 14.5 | 8.50 | 2.75 | 1.87 | 3.76 | 3.56 | 2.21 | 3.95 | 1.21 | 2.32 | Lipid (Cholesterol) | Fig S1d |
| CYP1B1 | 179 | 241 | 199 | 12.0 | 6.41 | 7.57 | 11.9 | 10.0 | 5.20 | 13.8 | 10.4 | 7.15 | Lipid (Steroid) | Fig S1d |
| SRD5A2 | 0.22 | 0.85 | 1.15 | 0.03 | 0.00 | 0.03 | 0.01 | 0.03 | 0.14 | 0.02 | 0.02 | 0.04 | Lipid (Steroid) | Fig S1d |
| NT5E | 0.01 | 0.02 | 0.01 | 0.09 | 0.48 | 0.51 | 1.31 | 7.85 | 6.66 | 4.64 | 5.09 | 6.32 | Nucleotide | Fig S1e |
| ENTPD8 | 0.08 | 0.14 | 0.15 | 0.16 | 0.30 | 0.36 | 1.45 | 1.41 | 0.96 | 0.75 | 0.37 | 0.88 | Nucleotide | Fig S1e |
| AMPD3 | 1.05 | 0.71 | 0.83 | 2.23 | 2.15 | 2.40 | 4.26 | 3.47 | 2.76 | 4.03 | 3.61 | 2.94 | Nucleotide (Purine) | Fig S1e |
| GMPR | 0.53 | 1.39 | 1.37 | 0.22 | 0.44 | 0.27 | 1.50 | 2.08 | 1.14 | 3.14 | 0.79 | 2.31 | Nucleotide (Purine) | Fig S1e |
| PDE6A | 4.3E-03 | 0.02 | 0.02 | 0.32 | 0.70 | 0.53 | 11.9 | 20.6 | 27.4 | 28.6 | 36.9 | 17.8 | Nucleotide (cGMP) | Fig S1e |
| PDE6G | 0.08 | 0.10 | 0.14 | 1.91 | 4.27 | 2.95 | 44.3 | 81.8 | 126 | 71.6 | 165 | 66.4 | Nucleotide (cGMP) | Fig S1e |
| PDE6H | 1.38 | 1.82 | 2.31 | 209 | 241 | 213 | 452 | 654 | 618 | 631 | 856 | 477 | Nucleotide (cGMP) | Fig S1e |
| PDE6C | 0.13 | 0.15 | 0.21 | 5.43 | 8.92 | 6.60 | 13.6 | 30.5 | 28.6 | 22.8 | 28.8 | 15.5 | Nucleotide (cGMP) | Fig S1e |
| GUCY2D | 0.24 | 0.15 | 0.19 | 6.14 | 7.18 | 6.94 | 14.5 | 14.1 | 15.2 | 14.0 | 18.2 | 13.8 | Nucleotide (cGMP) | Fig S1e |
| GUCY2F | 0.02 | 0.02 | 0.02 | 0.02 | 0.02 | 0.00 | 0.03 | 0.09 | 0.07 | 0.03 | 0.17 | 0.24 | Nucleotide (cGMP) | Fig S1e |
| ADCY1 | 3.40 | 4.41 | 5.11 | 23.9 | 24.7 | 23.4 | 35.3 | 27.6 | 32.5 | 26.0 | 33.7 | 25.9 | Nucleotide (cAMP) | Fig S1e |
| ADCY8 | 0.60 | 0.38 | 0.17 | 1.46 | 2.05 | 1.36 | 2.48 | 1.18 | 1.16 | 1.84 | 1.40 | 3.03 | Nucleotide (cAMP) | Fig S1e |
| PDE1C | 1.74 | 1.74 | 1.61 | 17.2 | 14.0 | 16.4 | 12.2 | 9.24 | 9.64 | 7.93 | 8.74 | 7.72 | Nucleotide (cGMP) | Fig S1e |
| ADCY2 | 1.22 | 1.11 | 1.38 | 2.38 | 4.28 | 2.86 | 3.86 | 3.88 | 4.18 | 5.80 | 3.36 | 3.44 | Nucleotide (cAMP) | Fig S1e |
| PDE4C | 1.89 | 1.49 | 1.33 | 1.95 | 1.20 | 1.40 | 3.55 | 0.94 | 0.93 | 1.62 | 1.08 | 0.93 | Nucleotide (cGMP) | Fig S1e |
| PDE7B | 2.21 | 1.66 | 1.81 | 0.80 | 0.78 | 0.81 | 0.99 | 1.77 | 1.32 | 1.32 | 1.20 | 1.26 | Nucleotide (cyclic nucleotide) | Fig S1e |
| GUCY2C | 0.01 | 0.02 | 0.01 | 0.02 | 0.04 | 0.02 | 0.01 | 0.02 | 0.02 | 0.01 | 0.02 | 0.00 | Nucleotide (cGMP) | Fig S1e |
| PDE11A | 1.33 | 0.96 | 1.60 | 0.43 | 0.37 | 0.48 | 0.34 | 0.50 | 0.44 | 0.68 | 0.85 | 0.66 | Nucleotide (cyclic nucleotide) | Fig S1e |
| ADCY4 | 0.11 | 0.09 | 0.09 | 0.01 | 0.00 | 0.00 | 0.06 | 0.05 | 0.07 | 0.02 | 0.01 | 0.02 | Nucleotide (cAMP) | Fig S1e |
| NPR1 | 0.03 | 0.25 | 0.25 | 0.04 | 0.07 | 0.00 | 0.11 | 0.04 | 0.06 | 0.04 | 0.01 | 0.02 | Nucleotide (cGMP) | Fig S1e |
| OAS1 | 0.12 | 0.14 | 0.19 | 0.07 | 0.00 | 0.09 | 1.11 | 0.48 | 0.28 | 1.99 | 0.45 | 5.34 | Nucleotide (Ribonuclease) | Fig S1e |
| AK5 | 0.48 | 0.68 | 0.63 | 1.99 | 3.95 | 3.77 | 5.71 | 10.7 | 5.93 | 3.71 | 3.67 | 2.83 | Nucleotide (Purine) | Fig S1e |
| ENPP2 | 6.17 | 6.51 | 5.66 | 3.98 | 2.62 | 1.84 | 15.6 | 19.3 | 7.75 | 16.3 | 10.8 | 9.01 | Nucleotide | Fig S1e |
| NUDT4 | 22.1 | 22.3 | 21.7 | 9.88 | 11.2 | 10.1 | 22.8 | 34.0 | 26.9 | 32.4 | 20.2 | 26.3 | Nucleotide | Fig S1e |
| UPB1 | 0.22 | 0.21 | 0.36 | 0.54 | 0.45 | 0.60 | 0.13 | 0.00 | 0.13 | 0.16 | 0.19 | 0.17 | Nucleotide (Pyrimidine) | Fig S1e |
| RRM2 | 20.6 | 20.5 | 22.1 | 12.4 | 3.63 | 3.56 | 3.97 | 2.21 | 2.11 | 1.22 | 0.43 | 0.66 | Nucleotide | Fig S1e |
| TK1 | 8.22 | 7.89 | 8.30 | 4.55 | 0.68 | 1.22 | 1.01 | 0.34 | 0.56 | 0.49 | 0.13 | 0.21 | Nucleotide (Pyrimidine) | Fig S1e |
| ARSF | 0.16 | 0.18 | 0.27 | 0.00 | 0.03 | 0.00 | 0.00 | 0.17 | 0.00 | 0.12 | 0.02 | 0.04 | Cofactor (Calcium) |  |
| BCHE | 18.2 | 18.5 | 19.6 | 17.9 | 12.3 | 13.0 | 13.1 | 17.9 | 15.0 | 15.3 | 9.17 | 9.93 | Detox |  |
| FMO2 | 0.00 | 0.00 | 0.00 | 0.11 | 0.21 | 0.16 | 0.15 | 0.34 | 0.20 | 0.28 | 0.58 | 0.47 | Detox |  |
| SOD3 | 0.44 | 0.43 | 0.49 | 0.19 | 0.04 | 0.18 | 0.78 | 0.77 | 0.18 | 1.61 | 0.17 | 0.69 | Detox |  |
| NDUFA4L2 | 0.55 | 0.93 | 0.49 | 1.85 | 3.90 | 2.68 | 29.4 | 6.51 | 6.70 | 27.6 | 7.28 | 12.3 | ETC (Complex I) |  |
| COX4I2 | 0.20 | 0.32 | 0.30 | 0.74 | 0.99 | 0.49 | 0.89 | 0.00 | 0.00 | 0.40 | 0.29 | 0.21 | ETC (Complex IV) |  |
| HPSE | 0.23 | 0.88 | 0.66 | 0.03 | 0.03 | 0.01 | 0.06 | 0.02 | 0.00 | 0.01 | 0.05 | 0.04 | Heparin Sulfate |  |
| HS3ST2 | 0.23 | 0.48 | 0.50 | 0.12 | 0.27 | 0.33 | 0.09 | 0.04 | 0.09 | 0.15 | 0.09 | 0.13 | Heparin Sulfate |  |
| HS6ST2 | 4.16 | 4.42 | 3.84 | 0.48 | 2.82 | 0.54 | 1.11 | 0.79 | 1.21 | 2.01 | 0.76 | 1.74 | Heparin Sulfate |  |
| HPSE2 | 0.43 | 0.27 | 0.32 | 0.02 | 0.00 | 0.02 | 0.01 | 0.00 | 0.00 | 0.10 | 0.10 | 0.05 | Heparin Sulfate |  |
| HS3ST3B1 | 0.73 | 0.81 | 0.79 | 4.81 | 5.82 | 5.33 | 7.82 | 7.55 | 8.19 | 7.67 | 7.15 | 6.05 | Heparin Sulfate |  |
| GLRA4 | 0.08 | 0.08 | 0.11 | 0.05 | 0.07 | 0.08 | 0.02 | 0.00 | 0.00 | 0.03 | 0.02 | 0.03 | Ion Transport |  |
| CHRNA4 | 1.97 | 2.37 | 2.33 | 2.06 | 1.70 | 1.36 | 1.16 | 1.37 | 0.99 | 0.98 | 0.99 | 0.75 | Ion Transport |  |
| P2RX2 | 0.56 | 0.64 | 0.45 | 0.17 | 0.31 | 0.17 | 0.27 | 0.23 | 0.12 | 0.11 | 0.03 | 0.07 | Ion Transport |  |
| SCN4A | 0.09 | 0.09 | 0.10 | 0.01 | 0.00 | 0.00 | 0.00 | 0.00 | 0.00 | 0.01 | 3.E-03 | 0.00 | Ion Transport |  |
| KCNJ13 | 4.93 | 5.25 | 5.35 | 0.75 | 0.35 | 0.40 | 1.65 | 3.13 | 1.25 | 1.45 | 2.05 | 1.25 | Ion Transport |  |
| P2RX1 | 0.22 | 0.12 | 0.15 | 0.05 | 0.04 | 0.09 | 0.04 | 0.03 | 0.03 | 0.06 | 0.07 | 0.06 | Ion Transport |  |
| KCNMB2 | 1.13 | 1.21 | 0.98 | 0.52 | 0.29 | 0.28 | 0.56 | 0.78 | 0.97 | 0.44 | 0.19 | 0.17 | Ion Transport |  |
| CLCN1 | 0.02 | 0.03 | 0.03 | 0.04 | 0.02 | 0.02 | 0.00 | 0.00 | 0.03 | 0.01 | 0.02 | 0.01 | Ion Transport |  |
| RYR1 | 1.84 | 2.07 | 2.13 | 2.25 | 1.41 | 1.30 | 0.52 | 0.44 | 0.37 | 0.36 | 0.48 | 0.56 | Ion Transport |  |
| CHRNA1 | 1.72 | 1.52 | 1.73 | 2.93 | 1.39 | 1.71 | 0.42 | 0.53 | 0.13 | 0.30 | 0.24 | 0.08 | Ion Transport |  |
| SCN7A | 0.26 | 0.29 | 0.35 | 0.28 | 0.05 | 0.11 | 0.02 | 0.06 | 0.02 | 0.04 | 0.04 | 0.05 | Ion Transport |  |
| CHRNB3 | 1.81 | 3.01 | 4.21 | 2.56 | 0.69 | 0.80 | 0.18 | 0.13 | 0.04 | 0.04 | 0.12 | 0.18 | Ion Transport |  |
| P2RX3 | 3.37 | 7.70 | 8.97 | 6.22 | 1.38 | 1.35 | 0.12 | 0.00 | 0.19 | 0.12 | 0.11 | 0.02 | Ion Transport |  |
| KCNH4 | 0.07 | 0.11 | 0.12 | 0.18 | 0.34 | 0.20 | 0.03 | 0.02 | 0.04 | 0.01 | 0.02 | 0.01 | Ion Transport |  |
| KCNK6 | 0.09 | 0.12 | 0.05 | 0.18 | 0.51 | 0.52 | 0.16 | 0.03 | 0.06 | 0.09 | 0.05 | 0.03 | Ion Transport |  |
| CACNG6 | 3.20 | 2.15 | 2.85 | 4.05 | 5.04 | 5.38 | 1.40 | 0.76 | 1.02 | 0.95 | 0.79 | 1.14 | Ion Transport |  |
| TRPC5 | 0.26 | 0.23 | 0.27 | 0.24 | 0.24 | 0.30 | 0.06 | 0.08 | 0.11 | 0.05 | 0.09 | 0.05 | Ion Transport |  |
| CFTR | 0.05 | 0.01 | 0.03 | 0.10 | 0.07 | 0.04 | 0.01 | 0.00 | 0.00 | 0.05 | 4.E-03 | 0.00 | Ion Transport |  |
| GABRA2 | 1.29 | 1.58 | 1.46 | 2.31 | 2.16 | 1.62 | 0.94 | 0.74 | 0.64 | 1.30 | 0.42 | 0.64 | Ion Transport |  |
| GABRP | 0.01 | 0.01 | 0.00 | 0.00 | 0.00 | 0.00 | 0.00 | 0.03 | 0.03 | 0.00 | 0.00 | 0.00 | Ion Transport |  |
| CACNA2D3 | 1.10 | 1.07 | 1.12 | 0.55 | 1.44 | 0.60 | 0.98 | 1.43 | 1.19 | 1.14 | 1.13 | 1.11 | Ion Transport |  |
| KCNK7 | 0.28 | 0.29 | 0.33 | 0.21 | 0.35 | 0.15 | 0.23 | 0.36 | 0.22 | 0.23 | 0.27 | 0.34 | Ion Transport |  |
| KCNK1 | 7.74 | 6.51 | 8.14 | 1.53 | 2.20 | 1.76 | 4.24 | 7.44 | 4.67 | 7.88 | 4.63 | 6.63 | Ion Transport |  |
| TRPC6 | 0.05 | 0.04 | 0.08 | 0.02 | 0.01 | 0.00 | 0.01 | 0.12 | 0.02 | 0.04 | 0.02 | 0.02 | Ion Transport |  |
| CLIC6 | 1.62 | 1.43 | 1.38 | 2.20 | 1.27 | 1.56 | 3.95 | 5.51 | 0.67 | 4.46 | 1.98 | 2.37 | Ion Transport |  |
| TRPM1 | 0.38 | 0.40 | 0.52 | 1.49 | 0.23 | 0.28 | 4.03 | 5.10 | 1.90 | 3.72 | 2.98 | 2.67 | Ion Transport |  |
| TF | 0.97 | 0.90 | 1.03 | 8.79 | 11.2 | 14.0 | 343 | 422 | 412 | 697 | 465 | 555 | Ion Transport |  |
| GABRB1 | 0.16 | 0.12 | 0.17 | 0.05 | 0.50 | 0.06 | 0.99 | 1.98 | 1.25 | 2.03 | 1.57 | 1.54 | Ion Transport |  |
| KCNH5 | 0.00 | 0.03 | 0.06 | 0.19 | 0.16 | 0.18 | 0.55 | 0.68 | 0.50 | 0.78 | 0.67 | 0.68 | Ion Transport |  |
| KCNJ10 | 0.27 | 0.41 | 0.33 | 2.03 | 2.87 | 2.38 | 4.97 | 9.67 | 7.93 | 9.45 | 6.75 | 8.75 | Ion Transport |  |
| NALCN | 0.28 | 0.24 | 0.26 | 0.34 | 0.29 | 0.13 | 0.81 | 0.67 | 0.68 | 1.33 | 0.67 | 0.58 | Ion Transport |  |
| KCND3 | 0.15 | 0.10 | 0.25 | 0.52 | 1.23 | 0.84 | 2.42 | 2.39 | 2.30 | 2.11 | 1.67 | 2.28 | Ion Transport |  |
| ATP2B2 | 0.19 | 0.19 | 0.25 | 0.88 | 1.95 | 1.21 | 3.13 | 4.34 | 3.06 | 4.12 | 3.07 | 2.59 | Ion Transport |  |
| GABRG2 | 0.12 | 0.13 | 0.13 | 2.61 | 3.26 | 2.77 | 5.37 | 7.23 | 5.31 | 6.43 | 5.29 | 4.14 | Ion Transport |  |
| GABRR1 | 0.07 | 0.07 | 0.16 | 0.04 | 0.35 | 0.04 | 2.71 | 3.00 | 2.64 | 2.91 | 3.00 | 2.54 | Ion Transport |  |
| GABRR3 | 0.00 | 0.00 | 0.02 | 0.38 | 0.84 | 0.40 | 13.5 | 13.4 | 15.5 | 17.3 | 14.3 | 13.5 | Ion Transport |  |
| ATP1A2 | 8.70 | 7.45 | 4.25 | 5.76 | 11.5 | 5.05 | 48.4 | 44.1 | 32.1 | 54.3 | 35.6 | 26.5 | Ion Transport |  |
| GABRA1 | 0.04 | 0.04 | 0.04 | 0.45 | 0.83 | 0.61 | 2.50 | 1.77 | 1.50 | 1.83 | 2.27 | 1.71 | Ion Transport |  |
| SLC30A8 | 0.01 | 0.01 | 0.00 | 0.00 | 0.00 | 0.00 | 0.03 | 0.12 | 0.03 | 0.04 | 0.01 | 0.05 | Ion Transport |  |
| SLC39A5 | 0.08 | 0.02 | 0.13 | 1.10 | 1.28 | 1.26 | 2.35 | 4.02 | 4.39 | 3.09 | 3.03 | 4.02 | Ion Transport |  |
| CACNA1S | 0.00 | 0.00 | 0.02 | 0.46 | 0.92 | 0.58 | 1.90 | 1.91 | 2.70 | 1.88 | 2.85 | 2.68 | Ion Transport |  |
| CACNA2D4 | 0.42 | 0.52 | 0.44 | 8.01 | 11.3 | 11.1 | 19.7 | 25.7 | 28.9 | 23.0 | 31.9 | 26.2 | Ion Transport |  |
| KCNB2 | 1.87 | 1.56 | 1.86 | 5.56 | 7.25 | 6.63 | 11.1 | 12.1 | 12.4 | 9.80 | 14.4 | 12.5 | Ion Transport |  |
| KCNB1 | 0.59 | 0.71 | 0.81 | 9.85 | 12.1 | 12.9 | 24.0 | 29.2 | 34.6 | 27.5 | 35.3 | 23.8 | Ion Transport |  |
| KCNV2 | 0.34 | 0.31 | 0.54 | 45.7 | 51.0 | 50.5 | 99.3 | 145 | 155 | 116 | 159 | 95.3 | Ion Transport |  |
| SLC24A4 | 0.03 | 0.03 | 0.03 | 4.90 | 5.72 | 5.71 | 7.76 | 11.2 | 12.7 | 10.4 | 14.6 | 9.96 | Ion Transport |  |
| CACNA1F | 0.13 | 0.17 | 0.21 | 7.71 | 8.76 | 8.32 | 15.0 | 19.0 | 20.0 | 18.0 | 25.7 | 15.7 | Ion Transport |  |
| KCNC2 | 0.09 | 0.15 | 0.10 | 0.14 | 0.67 | 0.13 | 1.15 | 1.16 | 1.02 | 0.54 | 0.98 | 0.89 | Ion Transport |  |
| GABRG1 | 0.01 | 4.6E-03 | 0.01 | 0.31 | 0.46 | 0.32 | 0.58 | 1.31 | 1.20 | 0.59 | 1.23 | 0.78 | Ion Transport |  |
| TRPC7 | 0.00 | 0.01 | 0.02 | 0.10 | 0.29 | 0.05 | 0.40 | 0.82 | 0.97 | 0.47 | 0.69 | 0.27 | Ion Transport |  |
| HCN1 | 0.27 | 0.26 | 0.23 | 4.47 | 5.74 | 6.12 | 9.68 | 17.2 | 17.2 | 15.0 | 18.1 | 12.7 | Ion Transport |  |
| CNGB1 | 0.15 | 0.24 | 0.23 | 10.5 | 30.4 | 24.4 | 120 | 187 | 230 | 138 | 186 | 149 | Ion Transport |  |
| SLC24A2 | 0.06 | 0.07 | 0.12 | 0.98 | 2.58 | 1.53 | 6.04 | 12.9 | 13.1 | 10.1 | 15.7 | 10.5 | Ion Transport |  |
| CNGB3 | 0.02 | 0.01 | 0.03 | 0.91 | 3.53 | 1.28 | 9.00 | 22.2 | 23.5 | 14.7 | 23.4 | 11.9 | Ion Transport |  |
| KCTD16 | 4.13 | 4.55 | 3.34 | 3.66 | 1.84 | 1.95 | 4.28 | 3.63 | 2.39 | 3.86 | 3.11 | 3.44 | Ion Transport |  |
| CLCNKA | 0.21 | 0.25 | 0.40 | 0.08 | 0.07 | 0.24 | 1.22 | 0.68 | 0.37 | 0.46 | 0.05 | 0.11 | Ion Transport |  |
| KCNK5 | 0.86 | 0.78 | 0.55 | 0.14 | 0.12 | 0.07 | 1.73 | 1.13 | 0.68 | 1.02 | 0.04 | 0.00 | Ion Transport |  |
| CHRND | 0.12 | 0.12 | 0.07 | 0.37 | 0.82 | 0.27 | 0.14 | 0.05 | 0.41 | 1.29 | 0.07 | 0.53 | Ion Transport |  |
| KCNA1 | 0.31 | 0.29 | 0.26 | 0.08 | 0.06 | 0.04 | 1.18 | 1.18 | 0.81 | 1.77 | 0.21 | 0.15 | Ion Transport |  |
| ATP1A4 | 0.60 | 0.27 | 0.11 | 0.10 | 0.82 | 0.43 | 1.22 | 0.33 | 1.28 | 0.56 | 1.25 | 0.87 | Ion Transport |  |
| GRIN2A | 0.08 | 0.11 | 0.06 | 0.34 | 0.55 | 0.43 | 0.46 | 0.46 | 0.35 | 0.89 | 0.30 | 0.29 | Ion Transport |  |
| SLC8A3 | 0.63 | 0.55 | 0.44 | 0.81 | 1.77 | 0.97 | 1.77 | 1.33 | 0.93 | 1.24 | 1.18 | 0.87 | Ion Transport |  |
| SCNN1B | 0.00 | 0.03 | 0.01 | 0.01 | 0.02 | 0.00 | 0.93 | 0.20 | 0.26 | 1.08 | 0.03 | 0.12 | Ion Transport |  |
| TRPM8 | 0.02 | 0.07 | 0.03 | 0.05 | 0.09 | 0.05 | 0.17 | 0.03 | 0.04 | 0.18 | 0.04 | 0.03 | Ion Transport |  |
| KCNE4 | 0.18 | 0.14 | 0.17 | 0.06 | 0.34 | 0.11 | 0.49 | 0.28 | 0.28 | 0.61 | 0.18 | 0.44 | Ion Transport |  |
| SCN1A | 0.11 | 0.13 | 0.09 | 0.57 | 1.45 | 0.66 | 1.62 | 0.90 | 0.91 | 2.11 | 0.49 | 1.53 | Ion Transport |  |
| TCN1 | 0.00 | 0.00 | 0.00 | 0.03 | 0.00 | 0.04 | 0.08 | 0.00 | 0.00 | 0.03 | 0.03 | 0.02 | Ion Transport |  |
| KCNK9 | 0.13 | 0.06 | 0.08 | 0.29 | 0.76 | 0.59 | 2.37 | 0.71 | 0.29 | 0.69 | 0.34 | 0.25 | Ion Transport |  |
| SCN9A | 0.36 | 0.63 | 0.60 | 1.02 | 0.53 | 0.64 | 1.29 | 1.02 | 1.07 | 1.08 | 0.36 | 0.67 | Ion Transport |  |
| SCN3A | 1.43 | 2.27 | 1.65 | 5.49 | 5.18 | 3.72 | 4.81 | 4.57 | 3.74 | 3.72 | 3.23 | 3.94 | Ion Transport |  |
| CACNG3 | 0.03 | 0.18 | 0.15 | 0.90 | 1.09 | 0.98 | 1.83 | 1.46 | 1.34 | 1.81 | 0.96 | 0.93 | Ion Transport |  |
| GRID2 | 0.43 | 0.45 | 0.30 | 1.71 | 1.99 | 1.29 | 2.75 | 2.60 | 2.03 | 2.55 | 1.44 | 1.51 | Ion Transport |  |
| GRIK4 | 0.24 | 0.42 | 0.37 | 2.17 | 2.03 | 2.08 | 3.90 | 3.21 | 2.17 | 3.07 | 4.09 | 2.71 | Ion Transport |  |
| ATP1B2 | 17.6 | 16.3 | 18.3 | 64.1 | 89.4 | 86.3 | 129 | 127 | 146 | 127 | 111 | 126 | Ion Transport |  |
| SLC8A1 | 2.60 | 2.27 | 1.80 | 14.1 | 14.0 | 15.1 | 17.3 | 20.3 | 17.7 | 15.4 | 15.6 | 16.0 | Ion Transport |  |
| KCNG4 | 0.12 | 0.12 | 0.13 | 3.82 | 6.35 | 6.93 | 9.50 | 7.02 | 7.32 | 7.52 | 4.62 | 7.46 | Ion Transport |  |
| KCNK4 | 0.00 | 0.08 | 0.07 | 0.47 | 0.29 | 0.51 | 0.49 | 0.39 | 0.39 | 0.33 | 0.55 | 0.32 | Ion Transport |  |
| SLC40A1 | 1.47 | 1.42 | 1.41 | 23.8 | 12.9 | 20.6 | 22.5 | 21.8 | 23.9 | 14.8 | 18.7 | 17.8 | Ion Transport |  |
| SCN2A | 0.20 | 0.39 | 0.37 | 3.25 | 4.20 | 2.65 | 2.94 | 3.88 | 3.48 | 3.28 | 3.81 | 3.79 | Ion Transport |  |
| CNGA3 | 0.60 | 0.57 | 0.62 | 7.53 | 10.0 | 11.1 | 6.92 | 9.71 | 10.4 | 8.49 | 11.0 | 7.04 | Ion Transport |  |
| GRIN1 | 0.15 | 0.14 | 0.22 | 0.60 | 1.00 | 0.97 | 0.97 | 0.93 | 0.84 | 0.77 | 1.03 | 0.75 | Ion Transport |  |
| CHRNA3 | 9.36 | 10.7 | 13.0 | 66.4 | 64.5 | 79.0 | 73.3 | 74.5 | 73.4 | 56.7 | 53.5 | 62.2 | Ion Transport |  |
| ATP1A3 | 15.3 | 17.0 | 19.4 | 109 | 110 | 102 | 112 | 107 | 112 | 98.6 | 115 | 83.6 | Ion Transport |  |
| KCNH6 | 0.65 | 0.80 | 1.14 | 9.85 | 11.0 | 11.4 | 11.9 | 12.4 | 10.9 | 10.0 | 11.2 | 7.79 | Ion Transport |  |
| CACNA1G | 1.05 | 1.04 | 1.12 | 2.40 | 2.86 | 3.08 | 2.67 | 3.74 | 3.18 | 2.26 | 1.93 | 2.73 | Ion Transport |  |
| GRIK3 | 3.24 | 3.51 | 3.43 | 6.04 | 6.48 | 6.57 | 7.64 | 10.5 | 7.66 | 6.76 | 4.50 | 5.49 | Ion Transport |  |
| KCNA2 | 0.58 | 0.79 | 0.66 | 8.68 | 10.2 | 9.57 | 9.77 | 7.42 | 6.35 | 8.68 | 5.94 | 6.01 | Ion Transport |  |
| KCTD8 | 0.39 | 0.34 | 0.46 | 1.77 | 1.41 | 1.54 | 1.74 | 1.43 | 1.31 | 0.92 | 0.70 | 0.95 | Ion Transport |  |
| KCNV1 | 0.09 | 0.13 | 0.09 | 0.13 | 0.12 | 0.11 | 0.11 | 0.20 | 0.17 | 0.10 | 0.01 | 0.03 | Ion Transport |  |
| KCNT1 | 0.03 | 0.07 | 0.08 | 0.13 | 0.38 | 0.32 | 0.14 | 0.29 | 0.05 | 0.22 | 0.14 | 0.24 | Ion Transport |  |
| KCND1 | 0.90 | 0.87 | 0.96 | 1.43 | 2.59 | 2.30 | 1.62 | 1.84 | 2.02 | 1.72 | 2.34 | 2.36 | Ion Transport |  |
| CACNG2 | 0.04 | 0.05 | 0.05 | 0.70 | 1.55 | 0.81 | 0.72 | 1.12 | 0.65 | 0.70 | 0.76 | 0.57 | Ion Transport |  |
| GLRA2 | 0.35 | 0.50 | 0.26 | 2.84 | 6.44 | 2.75 | 1.43 | 2.11 | 1.28 | 0.96 | 1.31 | 0.87 | Ion Transport |  |
| CACNA1E | 0.33 | 0.35 | 0.38 | 0.33 | 0.97 | 0.36 | 0.66 | 0.45 | 0.37 | 0.33 | 0.48 | 0.31 | Ion Transport |  |
| BEST2 | 0.03 | 0.07 | 0.08 | 0.19 | 0.15 | 0.03 | 0.09 | 0.09 | 0.14 | 0.09 | 0.18 | 0.09 | Ion Transport |  |
| SCN3B | 2.03 | 2.66 | 2.76 | 19.1 | 22.0 | 25.0 | 13.4 | 14.9 | 17.1 | 10.2 | 13.5 | 14.1 | Ion Transport |  |
| CHRNB4 | 6.22 | 6.75 | 8.41 | 36.8 | 37.9 | 46.2 | 26.5 | 21.6 | 25.9 | 13.9 | 14.0 | 18.6 | Ion Transport |  |
| CACNG5 | 2.43 | 2.88 | 3.34 | 13.1 | 18.2 | 13.8 | 11.4 | 9.66 | 7.86 | 5.92 | 7.39 | 5.39 | Ion Transport |  |
| KCNJ9 | 0.15 | 0.27 | 0.31 | 1.65 | 2.19 | 1.55 | 1.30 | 1.43 | 1.35 | 1.12 | 1.06 | 0.93 | Ion Transport |  |
| SLC8A2 | 0.69 | 0.79 | 0.78 | 5.11 | 4.98 | 4.48 | 3.92 | 3.93 | 3.54 | 2.57 | 2.91 | 2.52 | Ion Transport |  |
| BEST3 | 0.01 | 0.03 | 0.02 | 0.52 | 0.39 | 0.55 | 0.17 | 0.12 | 0.16 | 0.27 | 0.15 | 0.17 | Ion Transport |  |
| KCNA5 | 0.26 | 0.28 | 0.19 | 1.31 | 1.97 | 2.29 | 1.01 | 0.53 | 0.67 | 0.96 | 0.53 | 0.48 | Ion Transport |  |
| KCNK3 | 1.26 | 1.46 | 1.69 | 14.9 | 14.9 | 18.4 | 9.35 | 6.49 | 5.49 | 4.72 | 5.16 | 5.33 | Ion Transport |  |
| KCNS2 | 0.12 | 0.22 | 0.20 | 1.39 | 1.68 | 2.48 | 0.71 | 0.52 | 0.40 | 0.35 | 0.55 | 0.50 | Ion Transport |  |
| TRPV2 | 0.03 | 0.08 | 0.08 | 1.41 | 1.49 | 2.38 | 0.58 | 0.81 | 0.72 | 0.31 | 0.22 | 0.30 | Ion Transport |  |
| CACNA1B | 0.84 | 1.12 | 1.30 | 3.40 | 3.24 | 2.73 | 2.04 | 1.55 | 1.09 | 1.29 | 1.53 | 1.32 | Ion Transport |  |
| GABRR2 | 0.13 | 0.62 | 0.31 | 16.1 | 12.9 | 14.3 | 5.39 | 4.48 | 2.69 | 2.56 | 3.31 | 3.55 | Ion Transport |  |
| GRIK2 | 1.06 | 1.07 | 1.17 | 5.15 | 3.52 | 4.10 | 2.12 | 2.05 | 1.90 | 1.28 | 1.09 | 1.31 | Ion Transport |  |
| KCNK15 | 0.01 | 0.00 | 0.00 | 1.08 | 0.68 | 1.00 | 0.34 | 0.33 | 0.17 | 0.16 | 0.08 | 0.19 | Ion Transport |  |
| ATP2B3 | 0.39 | 0.48 | 0.53 | 0.86 | 1.32 | 0.84 | 0.69 | 0.46 | 0.44 | 0.92 | 0.56 | 0.64 | Ion Transport |  |
| GRIA2 | 2.20 | 3.29 | 3.40 | 8.52 | 11.1 | 6.92 | 5.26 | 4.99 | 5.05 | 6.52 | 5.38 | 4.83 | Ion Transport |  |
| CHRNA6 | 0.07 | 0.13 | 0.31 | 1.67 | 0.71 | 1.02 | 0.30 | 0.26 | 0.33 | 0.12 | 0.14 | 0.16 | Ion Transport |  |
| ACCN1 | 0.53 | 0.63 | 0.57 | 3.66 | 3.74 | 3.47 | 6.81 | 7.97 | 6.03 | 6.33 | 6.90 | 5.36 | Ion Transport |  |
| ACCN4 | 1.47 | 1.53 | 1.62 | 9.56 | 8.93 | 9.56 | 7.09 | 5.08 | 5.82 | 4.78 | 5.06 | 4.92 | Ion Transport |  |
| DUOX2 | 0.06 | 0.05 | 0.05 | 0.08 | 0.19 | 0.16 | 0.10 | 0.28 | 0.09 | 0.46 | 0.18 | 0.07 | NAD |  |
| NMNAT2 | 1.38 | 2.16 | 2.42 | 6.86 | 8.45 | 4.52 | 2.17 | 1.39 | 1.35 | 1.68 | 1.37 | 1.17 | NAD |  |
| CHAT | 0.16 | 0.22 | 0.14 | 0.19 | 0.30 | 0.18 | 0.16 | 0.04 | 0.08 | 0.16 | 0.04 | 0.15 | Neurotransmitter |  |
| DDC | 0.61 | 0.75 | 0.68 | 1.29 | 1.09 | 0.84 | 1.29 | 1.77 | 1.45 | 3.40 | 5.01 | 1.85 | Neurotransmitter |  |
| CA8 | 7.70 | 7.48 | 7.35 | 1.74 | 0.86 | 0.51 | 1.17 | 0.30 | 0.38 | 0.61 | 0.44 | 0.59 | Nitrogen Metabolism |  |
| CA10 | 0.21 | 0.42 | 0.73 | 1.65 | 3.11 | 1.94 | 10.9 | 9.35 | 7.40 | 8.80 | 7.67 | 7.21 | Nitrogen Metabolism |  |
| CA9 | 0.29 | 0.83 | 0.58 | 0.34 | 0.35 | 0.30 | 16.0 | 2.95 | 3.83 | 7.96 | 2.74 | 4.20 | Nitrogen Metabolism |  |
| CA12 | 0.37 | 0.16 | 0.32 | 0.51 | 0.92 | 0.33 | 11.6 | 4.80 | 4.11 | 8.87 | 0.79 | 1.69 | Nitrogen Metabolism |  |
| HMOX1 | 0.69 | 0.79 | 0.69 | 0.64 | 1.10 | 0.42 | 11.6 | 3.71 | 4.86 | 7.58 | 1.07 | 2.18 | Porphyrin |  |
| ATP6V0A4 | 0.05 | 0.05 | 0.04 | 0.04 | 0.17 | 0.02 | 0.15 | 0.67 | 0.74 | 0.22 | 0.31 | 0.23 | Proton Transport |  |
| ATP6V0D2 | 0.01 | 0.03 | 0.00 | 1.70 | 3.25 | 3.64 | 5.12 | 7.98 | 9.59 | 4.81 | 7.35 | 7.11 | Proton Transport |  |
| ATP8A1 | 4.74 | 5.46 | 5.46 | 14.0 | 11.5 | 12.6 | 7.79 | 10.3 | 9.32 | 7.37 | 11.1 | 8.97 | Proton Transport |  |
| TXNRD2 | 0.13 | 0.24 | 0.27 | 0.06 | 0.00 | 0.07 | 0.04 | 0.00 | 0.00 | 0.00 | 0.00 | 0.01 | Redox |  |
| CP | 15.7 | 34.4 | 29.7 | 11.1 | 6.88 | 12.4 | 65.1 | 102 | 65.7 | 95.8 | 44.0 | 81.0 | Redox |  |
| CRYL1 | 13.4 | 14.2 | 14.3 | 10.1 | 8.42 | 7.80 | 14.1 | 11.3 | 10.5 | 17.6 | 11.3 | 14.1 | Sugar |  |
| LCTL | 1.17 | 2.03 | 1.89 | 0.63 | 0.25 | 0.52 | 1.33 | 1.13 | 0.65 | 0.75 | 0.36 | 0.27 | Sugar |  |
| CHI3L1 | 0.07 | 0.02 | 0.02 | 0.04 | 0.03 | 0.04 | 0.86 | 0.57 | 0.29 | 3.82 | 0.21 | 0.84 | Sugar (Aminosugar) |  |
| G6PC2 | 0.04 | 0.03 | 0.02 | 0.09 | 0.24 | 0.14 | 2.90 | 7.05 | 5.88 | 5.90 | 6.96 | 8.53 | Sugar (gluconeogenesi) |  |
| SYNJ2 | 0.65 | 0.67 | 0.69 | 0.35 | 0.57 | 0.77 | 0.90 | 0.84 | 0.54 | 1.27 | 0.37 | 0.57 | Sugar (Inositol Phosphate) |  |
| TKTL1 | 1.11 | 1.46 | 1.12 | 0.45 | 0.95 | 0.44 | 2.84 | 2.49 | 3.91 | 4.23 | 1.74 | 6.62 | Sugar (PPP) |  |
| TKTL2 | 0.01 | 0.01 | 0.01 | 0.03 | 0.06 | 0.07 | 0.00 | 0.09 | 0.00 | 0.04 | 0.04 | 0.06 | Sugar (PPP) |  |
| SULT1E1 | 1.70 | 0.08 | 0.11 | 0.27 | 0.33 | 0.41 | 0.05 | 0.05 | 0.00 | 0.00 | 0.03 | 0.02 | Sulfate |  |
| ARSI | 0.33 | 0.33 | 0.33 | 0.21 | 0.09 | 0.20 | 0.45 | 0.42 | 0.21 | 0.30 | 0.22 | 0.21 | Sulfate |  |
| SULT1C2 | 0.24 | 0.21 | 0.24 | 0.10 | 0.04 | 0.07 | 0.61 | 0.48 | 0.39 | 0.48 | 0.22 | 0.52 | Sulfate |  |
| CYP26A1 | 9.25 | 15.4 | 12.8 | 1.40 | 17.7 | 14.2 | 5.57 | 6.44 | 5.92 | 4.84 | 4.69 | 3.99 | Vitamin (A) |  |
| RDH10 | 27.7 | 27.3 | 28.0 | 20.4 | 29.7 | 29.9 | 53.3 | 59.7 | 53.5 | 80.8 | 49.4 | 79.2 | Vitamin (A) |  |
| RPE65 | 0.10 | 0.04 | 0.13 | 0.19 | 0.87 | 0.24 | 1.27 | 1.55 | 0.68 | 2.66 | 1.33 | 1.02 | Vitamin (A) |  |
| LRAT | 0.13 | 0.08 | 0.10 | 0.02 | 0.15 | 0.11 | 0.95 | 1.86 | 0.77 | 1.13 | 0.58 | 0.93 | Vitamin (A) |  |
| CYP26B1 | 3.15 | 2.48 | 1.14 | 1.48 | 1.96 | 1.54 | 4.26 | 3.32 | 2.99 | 3.86 | 1.44 | 4.09 | Vitamin (A) |  |
| ALDH1A3 | 33.8 | 32.0 | 11.7 | 8.15 | 8.11 | 8.54 | 14.8 | 11.6 | 4.67 | 11.0 | 6.11 | 5.78 | Vitamine (Retinoic acidl, Aldehyde) |  |
| ALDH1A1 | 213 | 244 | 260 | 21.5 | 1.77 | 3.18 | 2.70 | 1.62 | 2.20 | 1.93 | 2.82 | 2.38 | Vitamine (Retinoic acid, Alcohol) |  |
| AKR1C1 | 3.08 | 3.76 | 4.02 | 5.66 | 3.31 | 3.23 | 4.19 | 5.99 | 4.59 | 7.61 | 3.71 | 3.58 | Vitamine (Retinoic acid, Alcohol) |  |
| AKR1C2 | 1.19 | 1.40 | 1.66 | 4.47 | 2.73 | 1.83 | 2.70 | 4.63 | 3.00 | 3.24 | 3.13 | 2.52 | Vitamine (Retinoic acid, Alcohol) |  |

**Table S4.** **Differentially expressed metabolic genes in WT and *Nrl* KO retinas.**

| **Gene symbol** | **FC** | **P value** | **Classification (Sub-category/substrate)** | **In figures** |
| --- | --- | --- | --- | --- |
| *Mfsd4b3-ps* | 8.57 | 8.33E-03 | Small Molecule Transport (Glucose) | Fig S3b |
| *Slc2a4* | 3.82 | 2.33E-03 | Small Molecule Transport (Glucose, Glucosamine) | Fig S3b |
| *Slc45a3* | 2.71 | 1.25E-02 | Small Molecule Transport (Sugar) | Fig S3b |
| *Mfsd4a* | 2.02 | 4.62E-03 | Small Molecule Transport (Glucose) | Fig S3b |
| *Sv2a* | 1.75 | 1.12E-03 | Small Molecule Transport (Sugar) | Fig S3b |
| *Sv2b* | 1.73 | 2.85E-04 | Small Molecule Transport (Sugar) | Fig S3b |
| *Slc2a13* | 1.57 | 5.80E-03 | Small Molecule Transport (Myo-Inoisitol) | Fig S3b |
| *Slc35f1* | 1.52 | 3.91E-03 | Small Molecule Transport (Nucleotide sugar) | Fig S3b |
| *Slc35f3* | 1.52 | 4.15E-02 | Small Molecule Transport (Nucleotide sugar) | Fig S3b |
| *Sv2c* | 1.52 | 1.08E-02 | Small Molecule Transport (Sugar) | Fig S3b |
| *Slc35c2* | 1.44 | 6.92E-03 | Small Molecule Transport (Nucleotide sugar) | Fig S3b |
| *Slc45a4* | 1.43 | 1.90E-02 | Small Molecule Transport (Sugar) | Fig S3b |
| *Slc35b4* | 1.42 | 1.94E-02 | Small Molecule Transport (UDP-N-Acetyl-Glucosomine) | Fig S3b |
| *Slc2a1* | 1.31 | 3.56E-02 | Small Molecule Transport (Glucose, Glucosamine) | Fig S3b |
| *Slc35b1* | 0.74 | 3.20E-02 | Small Molecule Transport (Nucleotide sugar) | Fig S3b |
| *Slc35a1* | 0.74 | 1.16E-02 | Small Molecule Transport (Nucleotide sugar) | Fig S3b |
| *Slc35d1* | 0.71 | 1.46E-02 | Small Molecule Transport (UDP-N-Acetyl-Glucosomine) | Fig S3b |
| *Slc35b3* | 0.62 | 2.55E-02 | Small Molecule Transport (Nucleotide sugar) | Fig S3b |
| *Slc35f4* | 0.53 | 5.72E-03 | Small Molecule Transport (Nucleotide sugar) | Fig S3b |
| *Mfsd2a* | 3.39 | 3.66E-02 | Small Molecule Transport (Fatty acid) | Fig S3b |
| *Slc27a4* | 1.54 | 1.46E-04 | Small Molecule Transport (Fatty acid) | Fig S3b |
| *Slc27a1* | 0.64 | 5.71E-04 | Small Molecule Transport (Fatty acid) | Fig S3b |
| *Slc27a3* | 0.64 | 4.82E-02 | Small Molecule Transport (Fatty acid) | Fig S3b |
| *Slc22a5* | 0.54 | 2.13E-03 | Small Molecule Transport (Carnitine) | Fig S3b |
| *Slc25a29* | 0.36 | 1.90E-05 | Small Molecule Transport (Acylcarnitine) | Fig S3b |
| *Mfsd2b* | 0.17 | 7.17E-04 | Small Molecule Transport (Fatty acid) | Fig S3b |
| *Slc27a5* | 0.04 | 3.12E-03 | Small Molecule Transport (Fatty acid) | Fig S3b |
| *Slc29a3* | 2.01 | 4.28E-03 | Small Molecule Transport (inosine) | Fig S3b |
| *Slc25a32* | 1.70 | 2.33E-03 | Small Molecule Transport (Nucleotide) | Fig S3b |
| *Slc25a5* | 0.71 | 4.85E-03 | Small Molecule Transport (Nucleotide) | Fig S3b |
| *Slc29a1* | 0.64 | 4.91E-03 | Small Molecule Transport (inosine) | Fig S3b |
| *Slc25a19* | 0.51 | 3.14E-02 | Small Molecule Transport (dNTP, dNDPs) | Fig S3b |
| *Slc29a2* | 0.44 | 7.03E-03 | Small Molecule Transport (inosine) | Fig S3b |
| *Slc25a43* | 0.44 | 1.53E-02 | Small Molecule Transport (Nucleotide) | Fig S3b |
| *Slc25a42* | 0.40 | 3.47E-04 | Small Molecule Transport (Nucleotide) | Fig S3b |
| *Slc1a2* | 6.55 | 4.28E-02 | Small Molecule Transport (Creatine) | Fig S3b |
| *Slc1a7* | 2.29 | 3.47E-02 | Small Molecule Transport (Arg, Lys, Orn, His) | Fig S3b |
| *Slc1a1* | 2.12 | 1.35E-02 | Small Molecule Transport (Creatine) | Fig S3b |
| *Slc1a4* | 1.36 | 4.37E-03 | Small Molecule Transport (Arg, Citrulline,lys, Orn) | Fig S3b |
| *Slc6a17* | 0.30 | 3.01E-02 | Small Molecule Transport (Arg, Citrulline,lys) | Fig S3b |
| *Slc1a5* | 3.61 | 2.14E-03 | Small Molecule Transport (Glutamate, Asparate) | Fig S3b |
| *Ctns* | 1.71 | 1.82E-03 | Small Molecule Transport (Glutamate, Asparate) | Fig S3b |
| *Slc25a18* | 1.56 | 9.18E-03 | Small Molecule Transport (Glutamate, Asparate) | Fig S3b |
| *Slc16a1* | 0.46 | 1.72E-02 | Small Molecule Transport (Glutamate) | Fig S3b |
| *Slc16a3* | 1.44 | 4.51E-02 | Small Molecule Transport (Alanine, Cysteine, Serine) | Fig S3b |
| *Slc16a11* | 1.35 | 1.26E-02 | Small Molecule Transport (Pro, Cys, Ala) | Fig S3b |
| *Slc25a25* | 0.63 | 4.40E-03 | Small Molecule Transport (Alanine, Cysteine, Serine) | Fig S3b |
| *Slc17a4* | 0.61 | 4.95E-03 | Small Molecule Transport (Cystine) | Fig S3b |
| *Slc17a2* | 0.38 | 2.35E-03 | Small Molecule Transport (Mitochondrial Phosphate) | Fig S3b |
| *Slc17a3* | 0.19 | 4.85E-02 | Small Molecule Transport (Phosphate) | Fig S3b |
| *Slc16a12* | 0.19 | 1.44E-03 | Small Molecule Transport (Phosphate) | Fig S3b |
| *Slc7a2* | 0.12 | 9.57E-04 | Small Molecule Transport (Phosphate) | Fig S3b |
| *Slc6a8* | 0.60 | 2.28E-03 | Small Molecule Transport (Lactate/Pyruvate/3-HB) | Fig S3b |
| *Slc25a15* | 0.57 | 2.34E-03 | Small Molecule Transport (Lactate/Pyruvate) | Fig S3b |
| *Slc25a2* | 0.46 | 4.39E-03 | Small Molecule Transport (Pyryuvate) | Fig S3b |
| *Slc10a4* | 1.48 | 2.21E-02 | Small Molecule Transport (Bile acid) | Fig S3b |
| *Stard3* | 0.71 | 2.88E-03 | Small Molecule Transport (Cholesterol) | Fig S3b |
| *Stard3nl* | 0.63 | 1.41E-02 | Small Molecule Transport (Cholesterol) | Fig S3b |
| *Star* | 0.60 | 2.18E-03 | Small Molecule Transport (Cholesterol) | Fig S3b |
| *Stard10* | 0.48 | 6.74E-03 | Small Molecule Transport (PC, PE) | Fig S3b |
| *Npc1l1* | 0.25 | 1.13E-02 | Small Molecule Transport (Cholesterol) | Fig S3b |
| *Stard9* | 0.25 | 3.97E-03 | Small Molecule Transport (Lipid) | Fig S3b |
| *Stard7* | 0.23 | 1.22E-04 | Small Molecule Transport (Lipid) | Fig S3b |
| *Slco4a1* | 0.19 | 2.86E-04 | Small Molecule Transport (bile acids, prostaglandins, steroid conjugates, thyroid hormone) | Fig S3b |
| *Rhbg* | 35.76 | 6.17E-03 | Small Molecule Transport (Ammonium) | Fig S3b |
| *Slc38a1* | 3.12 | 6.12E-04 | Small Molecule Transport (Gln, Ala, Ser, Asn) | Fig S3b |
| *Slc16a5* | 2.92 | 2.33E-02 | Small Molecule Transport (MCT activity) | Fig S3b |
| *Slc15a3* | 2.09 | 1.87E-02 | Small Molecule Transport (peptide) | Fig S3b |
| *Slc4a3* | 2.02 | 1.01E-03 | Small Molecule Transport (Anion) | Fig S3b |
| *Mfsd9* | 1.98 | 2.32E-02 | Small Molecule Transport | Fig S3b |
| *Slc6a11* | 1.95 | 1.77E-03 | Small Molecule Transport (GABA, β-Alanine) | Fig S3b |
| *Slc7a4* | 1.89 | 1.86E-03 | Small Molecule Transport (Amino acid) | Fig S3b |
| *Slc7a5* | 1.81 | 1.45E-02 | Small Molecule Transport (Unkown) | Fig S3b |
| *Slc16a2* | 1.78 | 1.02E-02 | Small Molecule Transport (T4) | Fig S3b |
| *Slc25a45* | 1.78 | 4.62E-03 | Small Molecule Transport (Amino acid) | Fig S3b |
| *Slc7a14* | 1.63 | 1.06E-02 | Small Molecule Transport (Amino acid) | Fig S3b |
| *Slc14a1* | 1.62 | 4.11E-04 | Small Molecule Transport (Urea) | Fig S3b |
| *Slc4a4* | 1.61 | 3.58E-02 | Small Molecule Transport (Biocarbonate) | Fig S3b |
| *Slc6a1* | 1.58 | 8.41E-03 | Small Molecule Transport (GABA) | Fig S3b |
| *Svop* | 1.52 | 1.74E-04 | Small Molecule Transport | Fig S3b |
| *Slc36a4* | 1.50 | 1.21E-02 | Small Molecule Transport (Proline) | Fig S3b |
| *Slc6a9* | 1.48 | 1.44E-02 | Small Molecule Transport (Glycine, Sarcosine) | Fig S3b |
| *Slc7a8* | 1.47 | 2.42E-02 | Small Molecule Transport (Amino acid) | Fig S3b |
| *Slc38a2* | 1.44 | 2.12E-02 | Small Molecule Transport (Met, Gln, Ala, Ser, Asn) | Fig S3b |
| *Slc43a2* | 1.41 | 2.98E-02 | Small Molecule Transport (BCAA) | Fig S3b |
| *Slc32a1* | 1.38 | 2.26E-02 | Small Molecule Transport (GABA, Gly) | Fig S3b |
| *Slc18a3* | 1.36 | 1.12E-02 | Small Molecule Transport (Acetylcholine, Choline) | Fig S3b |
| *Slc18a2* | 1.35 | 4.72E-02 | Small Molecule Transport (Monoamine) | Fig S3b |
| *Slc6a5* | 1.34 | 4.31E-02 | Small Molecule Transport (Glycine) | Fig S3b |
| *Slc19a1* | 1.32 | 3.68E-02 | Small Molecule Transport (Thiamine, Folate) | Fig S3b |
| *Slc25a17* | 0.75 | 9.55E-03 | Small Molecule Transport (ADP/ATP) | Fig S3b |
| *Slc5a5* | 0.73 | 1.42E-02 | Small Molecule Transport (iodide) | Fig S3b |
| *Slc25a37* | 0.67 | 1.39E-02 | Small Molecule Transport (mitochondrial transport) | Fig S3b |
| *Sfxn4* | 0.65 | 3.47E-02 | Small Molecule Transport | Fig S3b |
| *Slco1c1* | 0.60 | 2.00E-02 | Small Molecule Transport (Thyroid) | Fig S3b |
| *Slc25a28* | 0.59 | 1.20E-02 | Small Molecule Transport (Mitohondrial Iron) | Fig S3b |
| *Sfxn2* | 0.57 | 4.20E-02 | Small Molecule Transport (Serine) | Fig S3b |
| *Mfsd10* | 0.57 | 1.97E-02 | Small Molecule Transport (Anion) | Fig S3b |
| *Slc25a34* | 0.56 | 3.69E-02 | Small Molecule Transport (Di-Tri-carboxylic acid) | Fig S3b |
| *Slco4c1* | 0.52 | 3.10E-02 | Small Molecule Transport (cAMP, steroid conjugates, thyroid hormones) | Fig S3b |
| *Slc46a1* | 0.49 | 1.13E-03 | Small Molecule Transport (Folate) | Fig S3b |
| *Slc26a2* | 0.48 | 4.17E-04 | Small Molecule Transport (Sulfate) | Fig S3b |
| *Slc25a35* | 0.47 | 1.54E-03 | Small Molecule Transport (Di-Tri-carboxylic acid) | Fig S3b |
| *Flvcr1* | 0.46 | 3.27E-03 | Small Molecule Transport (Hem) | Fig S3b |
| *Slc5a8* | 0.38 | 4.04E-02 | Small Molecule Transport (iodide) | Fig S3b |
| *Slc26a11* | 0.33 | 2.92E-05 | Small Molecule Transport (Sulfate) | Fig S3b |
| *Slc15a1* | 0.32 | 5.04E-03 | Small Molecule Transport (peptide) | Fig S3b |
| *Slc16a4* | 0.30 | 1.47E-03 | Small Molecule Transport (MCT activity) | Fig S3b |
| *Aqp7* | 0.30 | 2.34E-02 | Small Molecule Transport (Water) | Fig S3b |
| *Slc23a1* | 0.29 | 2.18E-02 | Small Molecule Transport (Vitamine C) | Fig S3b |
| *Slc16a6* | 0.29 | 1.74E-05 | Small Molecule Transport (MCT activity) | Fig S3b |
| *Slc17a1* | 0.27 | 6.55E-03 | Small Molecule Transport (Urate) | Fig S3b |
| *Folr2* | 0.16 | 3.01E-02 | Small Molecule Transport (Folate) | Fig S3b |
| *Slc22a13* | 0.14 | 2.14E-02 | Small Molecule Transport (Nicotinic acid) | Fig S3b |
| *Aqp1* | 0.01 | 4.35E-04 | Small Molecule Transport (Water, CO2, Ammonium) | Fig S3b |
| *Pygm* | 5.45 | 3.80E-05 | Sugar (Glycogen) | Fig S3c |
| *Gaa* | 1.46 | 1.06E-03 | Sugar (Glycogen) | Fig S3c |
| *Pcx* | 1.59 | 8.21E-03 | TCA Cycle | Fig S3c |
| *Me1* | 1.34 | 3.72E-02 | TCA Cycle | Fig S3c |
| *Pdha1* | 1.31 | 3.65E-03 | TCA Cycle | Fig S3c |
| *Pdhb* | 0.69 | 6.58E-04 | TCA Cycle | Fig S3c |
| *Gpt* | 0.57 | 4.91E-04 | Amino Acid (Glutamate, Alaline) | Fig S3c |
| *Ogdh* | 1.64 | 1.23E-03 | TCA Cycle | Fig S3c |
| *Idh3a* | 1.51 | 8.60E-03 | TCA Cycle | Fig S3c |
| *Acly* | 1.46 | 3.87E-03 | TCA Cycle | Fig S3c |
| *Ogdhl* | 1.44 | 5.66E-05 | TCA Cycle | Fig S3c |
| *Got1* | 1.33 | 2.35E-03 | TCA Cycle | Fig S3c |
| *Cs* | 1.31 | 8.40E-04 | TCA Cycle | Fig S3c |
| *Aco1* | 0.76 | 2.41E-02 | TCA Cycle | Fig S3c |
| *Idh1* | 0.68 | 1.07E-03 | TCA Cycle | Fig S3c |
| *Hkdc1* | 1.66 | 9.82E-03 | Glycolysis | Fig S3c |
| *Pgm2l1* | 1.55 | 1.03E-02 | Glycolysis | Fig S3c |
| *Pfkm* | 1.48 | 3.71E-04 | Glycolysis | Fig S3c |
| *Eno2* | 1.33 | 7.37E-04 | Glycolysis | Fig S3c |
| *Aldoa* | 0.73 | 1.95E-03 | Glycolysis | Fig S3c |
| *Pfkl* | 0.72 | 6.40E-03 | Glycolysis | Fig S3c |
| *Eno1* | 0.72 | 3.01E-02 | Glycolysis | Fig S3c |
| *Ldha* | 0.69 | 1.78E-03 | Glycolysis | Fig S3c |
| *Pgk1* | 0.67 | 8.90E-03 | Glycolysis | Fig S3c |
| *Gapdh* | 0.65 | 8.63E-03 | Glycolysis | Fig S3c |
| *Pfkfb4* | 0.54 | 1.66E-02 | Glycolysis | Fig S3c |
| *Pgm2* | 0.52 | 1.43E-02 | Glycolysis | Fig S3c |
| *Pfkfb1* | 0.42 | 2.52E-03 | Glycolysis | Fig S3c |
| *Gapdhs* | 0.38 | 2.23E-02 | Glycolysis | Fig S3c |
| *Pfkp* | 0.34 | 2.87E-05 | Glycolysis | Fig S3c |
| *Pfkfb2* | 0.25 | 1.21E-02 | Glycolysis | Fig S3c |
| *Aldob* | 0.16 | 3.22E-03 | Glycolysis | Fig S3c |
| *Crym* | 2.24 | 4.91E-03 | Amino Acid (Cystathionine) | Fig S3d |
| *Suox* | 1.39 | 3.99E-03 | Amino Acid (Cysteine) | Fig S3d |
| *Pcyox1* | 1.35 | 3.31E-03 | Amino Acid (Cysteine) | Fig S3d |
| *Ado* | 1.33 | 8.10E-04 | Amino Acid (Cysteine) | Fig S3d |
| *Nfs1* | 0.77 | 3.38E-03 | Amino Acid (Cysteine) | Fig S3d |
| *Cdo1* | 0.53 | 3.59E-03 | Amino Acid (Cysteine) | Fig S3d |
| *Gstm1* | 1.49 | 8.01E-03 | Amino Acid (Glutathione) | Fig S3d |
| *Gclm* | 1.49 | 2.84E-02 | Amino Acid (Glutathione) | Fig S3d |
| *Gsr* | 1.45 | 1.23E-02 | Amino Acid (Glutathione) | Fig S3d |
| *Gss* | 0.59 | 3.23E-03 | Amino Acid (Glutathione) | Fig S3d |
| *Gpx4* | 0.56 | 2.22E-02 | Amino Acid (Glutathione) | Fig S3d |
| *Nos1* | 1.73 | 2.40E-02 | Amino Acid (Arginine) | Fig S3d |
| *Ckb* | 1.59 | 3.10E-03 | Amino Acid (Creatine) | Fig S3d |
| *Ddah1* | 1.53 | 7.66E-03 | Amino Acid (Arginine, Citrulline) | Fig S3d |
| *Asl* | 0.72 | 2.67E-02 | Amino Acid (Arginine, Urea) | Fig S3d |
| *Odc1* | 0.56 | 8.40E-03 | Amino Acid (Ornithine, Polyamine) | Fig S3d |
| *Arg2* | 0.28 | 1.87E-03 | Amino Acid (Arginine, Urea) | Fig S3d |
| *Gldc* | 2.28 | 4.92E-02 | Amino Acid (Glycine) | Fig S3d |
| *Sdsl* | 2.00 | 1.77E-02 | Amino Acid (Serine) | Fig S3d |
| *Psph* | 0.76 | 2.71E-02 | Amino Acid (Serine) | Fig S3d |
| *Amt* | 0.67 | 1.71E-02 | Amino Acid (Glycine) | Fig S3d |
| *Gnmt* | 0.33 | 3.56E-03 | Amino Acid (Glycine) | Fig S3d |
| *P3h4* | 1.60 | 4.90E-02 | Amino Acid (Proline) | Fig S3d |
| *P4ha2* | 0.76 | 1.32E-03 | Amino Acid (Proline) | Fig S3d |
| *P4ha1* | 0.71 | 1.46E-02 | Amino Acid (Proline) | Fig S3d |
| *Aldh4a1* | 0.69 | 3.29E-03 | Amino Acid (Proline) | Fig S3d |
| *P4ha3* | 0.45 | 1.26E-02 | Amino Acid (Proline) | Fig S3d |
| *Prodh2* | 0.44 | 3.43E-02 | Amino Acid (Proline) | Fig S3d |
| *Dct* | 3.17 | 3.56E-02 | Amino Acid (Tryosine, Pigment) | Fig S3d |
| *Tyr* | 2.92 | 2.42E-02 | Amino Acid (Tryosine, Pigment) | Fig S3d |
| *Tyrp1* | 2.86 | 4.07E-02 | Amino Acid (Tryosine, Pigment) | Fig S3d |
| *Fah* | 0.58 | 1.11E-02 | Amino Acid (Tyrosine) | Fig S3d |
| *Hgd* | 0.27 | 4.34E-02 | Amino Acid (Tyrosine) | Fig S3d |
| *Hpdl* | 0.13 | 3.18E-04 | Amino Acid (Tyrosine) | Fig S3d |
| *Aspg* | 3.12 | 2.94E-02 | Amino Acid (Asparagine) | Fig S3d |
| *Nit2* | 0.74 | 4.52E-02 | Amino Acid (Asparagine) | Fig S3d |
| *Asrgl1* | 0.28 | 3.35E-03 | Amino Acid (Asparagine) | Fig S3d |
| *Adi1* | 1.32 | 2.22E-02 | Amino Acid (Methionine) | Fig S3d |
| *Mtap* | 1.34 | 2.81E-02 | Amino Acid (Methionion, Polyamine) | Fig S3d |
| *Amd1* | 0.61 | 1.94E-03 | Amino Acid (Methionion, Polyamine) | Fig S3d |
| *Amdhd2* | 0.61 | 6.11E-03 | Amino Acid (histidine) | Fig S3d |
| *Aoc2* | 0.44 | 1.33E-03 | Amino acid (Dopamine, histamine) | Fig S3d |
| *Hdc* | 1.23E-04 | 1.55E-02 | Amino Acid (histidine) | Fig S3d |
| *Bcat1* | 2.32 | 1.25E-04 | Amino Acid (BCAAs) | Fig S3d |
| *Aldh6a1* | 0.72 | 8.77E-03 | Amino Acid (BCAAs) | Fig S3d |
| *Mccc2* | 0.67 | 3.60E-03 | Amino Acid (BCAAs) | Fig S3d |
| *Ivd* | 0.44 | 7.72E-03 | Amino Acid (BCAAs) | Fig S3d |
| *Tph1* | 9.61 | 3.54E-03 | Amino Acid (Neurotransmitter) | Fig S3d |
| *Ahcyl2* | 2.90 | 2.48E-03 | Amino Acid (Cofactor) | Fig S3d |
| *Aldh1l1* | 2.53 | 1.29E-02 | Amino Acid (One carbon) | Fig S3d |
| *Thnsl2* | 1.87 | 2.95E-03 | Amino Acid (Threonine) | Fig S3d |
| *Aldh5a1* | 1.64 | 1.60E-04 | Amino Acid (Glutamate) | Fig S3d |
| *Tph2* | 1.54 | 2.25E-02 | Amino Acid (Neurotransmitter) | Fig S3d |
| *Gad2* | 1.50 | 1.08E-02 | Amino Acid (Glutamate) | Fig S3d |
| *Smox* | 1.49 | 3.96E-02 | Amino Acid (Polyamine) | Fig S3d |
| *Gad1* | 1.33 | 5.72E-03 | Amino Acid (Glutamate) | Fig S3d |
| *Aldh9a1* | 0.73 | 2.58E-03 | Amino acid (GABA) | Fig S3d |
| *Ahcyl1* | 0.66 | 2.64E-03 | Amino Acid (Cofactor) | Fig S3d |
| *Nat10* | 0.66 | 2.44E-02 | Amino Acid (Acetylation) | Fig S3d |
| *Acy1* | 0.65 | 5.53E-03 | Amino Acid (Hydrolysis) | Fig S3d |
| *Gpt* | 0.57 | 4.91E-04 | Amino Acid (Glutamate, Alaline) | Fig S3d |
| *Acmsd* | 0.52 | 3.06E-02 | Amino Acid (Tryptophan) | Fig S3d |
| *Apeh* | 0.51 | 3.15E-03 | Amino Acid (Hydrolysis) | Fig S3d |
| *Pipox* | 0.47 | 6.91E-03 | Amino Acid (Lysine) | Fig S3d |
| *Aasdh* | 0.39 | 2.47E-03 | Amino Acid (Lysine) | Fig S3d |
| *Dpep2* | 0.32 | 1.57E-02 | Amino Acid (Hydrolysis) | Fig S3d |
| *Tmlhe* | 0.26 | 6.15E-04 | Amino Acid (Carnitine) | Fig S3d |
| *Pnmt* | 0.24 | 7.28E-03 | Amino Acid (Neurotransmitter) | Fig S3d |
| *Them5* | 6.72 | 8.91E-06 | Lipid (Fatty Acid) | Fig S3e |
| *Acbd6* | 4.85 | 2.53E-04 | Lipid (Fatty Acid) | Fig S3e |
| *Elovl7* | 3.96 | 3.63E-03 | Lipid (Fatty Acid) | Fig S3e |
| *Elovl2* | 3.95 | 7.09E-04 | Lipid (Fatty Acid) | Fig S3e |
| *Elovl6* | 2.61 | 7.20E-03 | Lipid (Fatty Acid) | Fig S3e |
| *Acads* | 2.57 | 4.11E-03 | Lipid (Fatty Acid) | Fig S3e |
| *Elovl5* | 2.16 | 4.80E-06 | Lipid (Fatty Acid) | Fig S3e |
| *Acacb* | 2.15 | 2.71E-02 | Lipid (Fatty Acid) | Fig S3e |
| *Acsf2* | 1.82 | 4.41E-03 | Lipid (Fatty Acid) | Fig S3e |
| *Abhd2* | 1.74 | 3.64E-03 | Lipid (Fatty Acid) | Fig S3e |
| *Ehhadh* | 1.59 | 4.42E-02 | Lipid (Fatty Acid) | Fig S3e |
| *Acaca* | 1.53 | 2.82E-02 | Lipid (Fatty Acid) | Fig S3e |
| *Acsl3* | 1.51 | 2.47E-02 | Lipid (Fatty Acid) | Fig S3e |
| *Oxsm* | 1.51 | 1.24E-02 | Lipid (Fatty Acid) | Fig S3e |
| *Fads2* | 1.45 | 3.94E-02 | Lipid (Fatty Acid) | Fig S3e |
| *Aldh3a2* | 1.41 | 1.43E-02 | Lipid (Fatty Acid) | Fig S3e |
| *Hadhb* | 0.77 | 4.34E-02 | Lipid (Fatty Acid) | Fig S3e |
| *Acss2* | 0.75 | 1.89E-02 | Lipid (Fatty Acid) | Fig S3e |
| *Abhd4* | 0.74 | 1.26E-02 | Lipid (Fatty Acid) | Fig S3e |
| *Acad9* | 0.72 | 2.97E-03 | Lipid (Fatty Acid) | Fig S3e |
| *Acaa1a* | 0.71 | 4.15E-02 | Lipid (Fatty Acid) | Fig S3e |
| *Etfdh* | 0.68 | 8.49E-05 | Lipid (Fatty Acid) | Fig S3e |
| *Acox3* | 0.68 | 1.07E-02 | Lipid (Fatty Acid) | Fig S3e |
| *Far2* | 0.68 | 2.92E-02 | Lipid (Fatty Acid) | Fig S3e |
| *Acat1* | 0.65 | 4.65E-02 | Lipid (Fatty Acid) | Fig S3e |
| *Acat2* | 0.65 | 1.48E-03 | Lipid (Fatty Acid) | Fig S3e |
| *Echdc2* | 0.63 | 4.86E-03 | Lipid (Fatty Acid) | Fig S3e |
| *Hadha* | 0.60 | 2.17E-02 | Lipid (Fatty Acid) | Fig S3e |
| *Acad8* | 0.60 | 9.46E-05 | Lipid (Fatty Acid) | Fig S3e |
| *Abhd11* | 0.58 | 6.95E-04 | Lipid (Fatty Acid) | Fig S3e |
| *Elovl4* | 0.56 | 2.54E-03 | Lipid (Fatty Acid) | Fig S3e |
| *Faah* | 0.55 | 7.89E-03 | Lipid (Fatty Acid) | Fig S3e |
| *Cpt1b* | 0.51 | 1.69E-02 | Lipid (Fatty Acid) | Fig S3e |
| *Amacr* | 0.49 | 8.77E-04 | Lipid (Fatty acid) | Fig S3e |
| *Elovl1* | 0.43 | 1.90E-04 | Lipid (Fatty Acid) | Fig S3e |
| *Phyhd1* | 0.40 | 6.17E-03 | Lipid (Fatty Acid) | Fig S3e |
| *Acadvl* | 0.33 | 5.83E-04 | Lipid (Fatty Acid) | Fig S3e |
| *Dgat1* | 0.33 | 1.74E-03 | Lipid (Fatty Acid) | Fig S3e |
| *Pnpla1* | 0.32 | 8.15E-04 | Lipid (Fatty Acid) | Fig S3e |
| *Abhd14a* | 0.28 | 9.54E-04 | Lipid (Fatty Acid) | Fig S3e |
| *Pnliprp2* | 0.27 | 2.28E-04 | Lipid (Fatty Acid) | Fig S3e |
| *Mecr* | 0.26 | 2.22E-05 | Lipid (Fatty Acid) | Fig S3e |
| *Acox2* | 0.25 | 3.87E-02 | Lipid (Fatty Acid) | Fig S3e |
| *Dgat2l6* | 0.13 | 2.55E-02 | Lipid (Fatty Acid) | Fig S3e |
| *Acoxl* | 0.03 | 3.78E-03 | Lipid (Fatty Acid) | Fig S3e |
| *Plcl1* | 4.12 | 9.07E-04 | Lipid (Membrane Lipid) | Fig S3e |
| *Crot* | 2.84 | 1.95E-04 | Lipid (Membrane Lipid) | Fig S3e |
| *Plcxd3* | 2.77 | 2.27E-02 | Lipid (Membrane Lipid) | Fig S3e |
| *Plce1* | 2.76 | 2.08E-02 | Lipid (Membrane Lipid) | Fig S3e |
| *Dgkk* | 2.63 | 2.13E-02 | Lipid (Membrane Lipid) | Fig S3e |
| *Dgki* | 2.61 | 2.93E-02 | Lipid (Membrane Lipid) | Fig S3e |
| *Mogat1* | 2.42 | 2.09E-03 | Lipid (Membrane Lipid) | Fig S3e |
| *Phospho1* | 2.37 | 1.11E-02 | Lipid (Membrane Lipid) | Fig S3e |
| *Plcg2* | 2.03 | 3.66E-02 | Lipid (Membrane Lipid) | Fig S3e |
| *Cds2* | 1.83 | 4.68E-03 | Lipid (Membrane Lipid) | Fig S3e |
| *Pgs1* | 1.74 | 1.62E-04 | Lipid (Membrane Lipid) | Fig S3e |
| *Mboat7* | 1.70 | 3.95E-03 | Lipid (Membrane Lipid) | Fig S3e |
| *Dagla* | 1.64 | 7.62E-03 | Lipid (Membrane Lipid) | Fig S3e |
| *Lpin1* | 1.59 | 1.64E-02 | Lipid (Membrane Lipid) | Fig S3e |
| *Plcb1* | 1.57 | 1.96E-02 | Lipid (Membrane Lipid) | Fig S3e |
| *Plcl2* | 1.54 | 1.02E-02 | Lipid (Membrane Lipid) | Fig S3e |
| *Gpd2* | 1.54 | 6.72E-03 | Lipid (Membrane Lipid) | Fig S3e |
| *Plcb4* | 1.49 | 4.56E-03 | Lipid (Membrane Lipid) | Fig S3e |
| *Mboat2* | 1.48 | 3.13E-02 | Lipid (Membrane Lipid) | Fig S3e |
| *Gdpd5* | 1.46 | 1.79E-03 | Lipid (Membrane Lipid) | Fig S3e |
| *Pla2g2c* | 1.45 | 4.58E-02 | Lipid (Membrane Lipid) | Fig S3e |
| *Gpd1* | 1.45 | 8.21E-03 | Lipid (Membrane Lipid) | Fig S3e |
| *Pla2g15* | 1.41 | 3.07E-03 | Lipid (Membrane Lipid) | Fig S3e |
| *Dgkg* | 1.40 | 2.26E-02 | Lipid (Membrane Lipid) | Fig S3e |
| *Plcb3* | 1.36 | 4.67E-02 | Lipid (Membrane Lipid) | Fig S3e |
| *Mboat1* | 1.35 | 3.09E-02 | Lipid (Membrane Lipid) | Fig S3e |
| *Lypla1* | 1.34 | 6.01E-03 | Lipid (Membrane Lipid) | Fig S3e |
| *Dgkd* | 1.33 | 4.25E-02 | Lipid (Membrane Lipid) | Fig S3e |
| *Gpd1l* | 1.32 | 8.88E-05 | Lipid (Membrane Lipid) | Fig S3e |
| *Plcd1* | 0.75 | 2.51E-02 | Lipid (Membrane Lipid) | Fig S3e |
| *Gk* | 0.65 | 2.86E-02 | Lipid (Membrane Lipid) | Fig S3e |
| *Gdpd1* | 0.64 | 7.88E-04 | Lipid (Membrane Lipid) | Fig S3e |
| *Chpt1* | 0.62 | 6.76E-03 | Lipid (Membrane Lipid) | Fig S3e |
| *Dgka* | 0.61 | 3.91E-04 | Lipid (Membrane Lipid) | Fig S3e |
| *Ddhd1* | 0.61 | 1.78E-02 | Lipid (Membrane Lipid) | Fig S3e |
| *Pcyt2* | 0.58 | 8.23E-03 | Lipid (Membrane Lipid) | Fig S3e |
| *Plcd4* | 0.57 | 1.52E-02 | Lipid (Membrane Lipid) | Fig S3e |
| *Etnk2* | 0.55 | 2.50E-02 | Lipid (Membrane Lipid) | Fig S3e |
| *Pemt* | 0.50 | 1.15E-02 | Lipid (Membrane Lipid) | Fig S3e |
| *Pla2g7* | 0.43 | 3.83E-04 | Lipid (Membrane Lipid) | Fig S3e |
| *Gdpd3* | 0.39 | 1.10E-02 | Lipid (Membrane Lipid) | Fig S3e |
| *Agpat3* | 0.33 | 2.73E-04 | Lipid (Membrane Lipid) | Fig S3e |
| *Plcd3* | 0.32 | 2.28E-04 | Lipid (Membrane Lipid) | Fig S3e |
| *Pla2g4b* | 0.31 | 9.98E-03 | Lipid (Membrane Lipid) | Fig S3e |
| *Pla2g3* | 0.26 | 4.11E-05 | Lipid (Membrane Lipid) | Fig S3e |
| *Idi1* | 0.57 | 5.60E-03 | Lipid (Mevalonate) | Fig S3e |
| *Hsd3b7* | 0.46 | 1.20E-02 | Lipid (Bile Acid) | Fig S3e |
| *Baat* | 0.20 | 2.67E-02 | Lipid (Bile Acid) | Fig S3e |
| *Fdft1* | 0.74 | 1.18E-03 | Lipid (Cholesterol) | Fig S3e |
| *Dhcr7* | 0.69 | 1.29E-02 | Lipid (Cholesterol) | Fig S3e |
| *Nsdhl* | 0.64 | 4.62E-04 | Lipid (Cholesterol) | Fig S3e |
| *Fdps* | 0.61 | 1.43E-03 | Lipid (Cholesterol) | Fig S3e |
| *Tm7sf2* | 0.48 | 5.08E-03 | Lipid (Cholesterol) | Fig S3e |
| *Ebpl* | 0.09 | 1.02E-05 | Lipid (Cholesterol) | Fig S3e |
| *Ptgs1* | 2.42 | 2.42E-02 | Lipid (Hormone) | Fig S3e |
| *Ace* | 1.47 | 5.70E-04 | Lipid (Hormone) | Fig S3e |
| *Dio2* | 1.45 | 3.75E-02 | Lipid (Hormone) | Fig S3e |
| *Hpgd* | 0.72 | 1.47E-02 | Lipid (Hormone) | Fig S3e |
| *Lta4h* | 0.65 | 6.74E-03 | Lipid (Hormone) | Fig S3e |
| *Ptgis* | 0.59 | 3.34E-02 | Lipid (Hormone) | Fig S3e |
| *Ptges2* | 0.48 | 2.93E-04 | Lipid (Hormone) | Fig S3e |
| *Alox5ap* | 0.38 | 1.81E-02 | Lipid (Hormone) | Fig S3e |
| *Aloxe3* | 0.32 | 4.93E-02 | Lipid (Hormone) | Fig S3e |
| *Tpo* | 0.15 | 2.15E-03 | Lipid (Hormone) | Fig S3e |
| *Smpdl3a* | 3.91 | 1.49E-03 | Lipid (Sphingolipid) | Fig S3e |
| *Sgpp2* | 2.05 | 2.53E-03 | Lipid (Sphingolipid) | Fig S3e |
| *Sgms1* | 1.58 | 3.15E-02 | Lipid (Sphingolipid) | Fig S3e |
| *Smpd1* | 1.42 | 1.51E-03 | Lipid (Sphingolipid) | Fig S3e |
| *Ugcg* | 1.40 | 6.24E-04 | Lipid (Sphingolipid) | Fig S3e |
| *Smpd2* | 0.58 | 7.51E-03 | Lipid (Sphingolipid) | Fig S3e |
| *Acer1* | 0.09 | 6.34E-04 | Lipid (Sphingosine) | Fig S3e |
| *Hsd17b2* | 2.24 | 1.45E-02 | Lipid (Steroid) | Fig S3e |
| *Cyp1b1* | 1.86 | 3.87E-03 | Lipid (Steroid) | Fig S3e |
| *Soat1* | 1.85 | 6.24E-03 | Lipid (Steroid) | Fig S3e |
| *Lipa* | 1.42 | 9.60E-04 | Lipid (Steroid) | Fig S3e |
| *Srd5a3* | 0.70 | 7.14E-03 | Lipid (Steroid) | Fig S3e |
| *Cyp2u1* | 0.70 | 5.88E-03 | Lipid (Steroid) | Fig S3e |
| *Hsd17b1* | 0.64 | 3.95E-02 | Lipid (Steroid) | Fig S3e |
| *Hsd3b2* | 0.52 | 9.40E-03 | Lipid (Steroid) | Fig S3e |
| *Klb* | 0.33 | 2.07E-02 | Lipid (Steroid) | Fig S3e |
| *Pter* | 1.97 | 3.96E-03 | Lipid | Fig S3e |
| *Lias* | 0.76 | 2.44E-02 | Lipid | Fig S3e |
| *Lipt1* | 0.67 | 4.51E-02 | Lipid | Fig S3e |
| *Cel* | 0.10 | 2.32E-02 | Lipid (Cofactor) | Fig S3e |
| *Abcb4* | 4.63 | 6.80E-07 | ABC Transporter |  |
| *Abcb1b* | 2.86 | 6.00E-03 | ABC Transporter |  |
| *Abcc4* | 1.91 | 2.85E-03 | ABC Transporter |  |
| *Abca6* | 1.46 | 3.67E-02 | ABC Transporter |  |
| *Abcg1* | 1.34 | 1.26E-03 | ABC Transporter |  |
| *Abcb7* | 1.34 | 2.51E-02 | ABC Transporter |  |
| *Abcd3* | 1.31 | 9.59E-03 | ABC Transporter |  |
| *Abcg4* | 0.76 | 1.22E-02 | ABC Transporter |  |
| *Abcc1* | 0.73 | 3.09E-02 | ABC Transporter |  |
| *Abcd4* | 0.66 | 3.95E-03 | ABC Transporter |  |
| *Abca4* | 0.32 | 1.53E-06 | ABC Transporter |  |
| *Abca13* | 0.17 | 1.64E-03 | ABC Transporter |  |
| *Abcg5* | 0.09 | 3.42E-02 | ABC Transporter |  |
| *Pank1* | 2.29 | 1.35E-03 | CoA |  |
| *Acot11* | 1.44 | 3.80E-02 | CoA |  |
| *Acot6* | 1.39 | 1.16E-02 | CoA |  |
| *Acot8* | 0.69 | 2.56E-02 | CoA |  |
| *Pank4* | 0.45 | 6.35E-04 | CoA |  |
| *Acot4* | 0.45 | 2.59E-02 | CoA |  |
| *Acot12* | 0.11 | 7.91E-03 | CoA |  |
| *Hlcs* | 1.31 | 1.83E-02 | Cofactor(molybdenum) |  |
| *Ephx1* | 2.76 | 1.62E-03 | Detox |  |
| *Dhdh* | 1.41 | 1.92E-03 | Detox |  |
| *Ephx2* | 1.38 | 2.37E-02 | Detox |  |
| *Pon2* | 1.33 | 4.59E-02 | Detox |  |
| *Ces2f* | 0.36 | 3.65E-02 | Detox |  |
| *Epx* | 0.12 | 1.40E-06 | Detox |  |
| *Ndufs8* | 0.77 | 2.58E-02 | ETC (Complex I) |  |
| *Ndufb5* | 0.76 | 1.79E-02 | ETC (Complex I) |  |
| *Ndufb8* | 0.76 | 2.89E-02 | ETC (Complex I) |  |
| *Ndufv1* | 0.76 | 3.45E-02 | ETC (Complex I) |  |
| *Ndufb9* | 0.75 | 2.75E-02 | ETC (Complex I) |  |
| *Uqcrq* | 0.75 | 4.12E-02 | ETC (Complex III) |  |
| *Ndufa5* | 0.74 | 3.04E-02 | ETC (Complex I) |  |
| *Ndufa2* | 0.74 | 4.30E-02 | ETC (Complex I) |  |
| *Ndufa3* | 0.74 | 3.66E-02 | ETC (Complex I) |  |
| *Ndufa6* | 0.72 | 2.80E-02 | ETC (Complex I) |  |
| *Ndufs7* | 0.72 | 3.75E-02 | ETC (Complex I) |  |
| *Cox5b* | 0.71 | 3.08E-02 | ETC (Complex IV) |  |
| *Cox7a2l* | 0.71 | 1.29E-02 | ETC (Complex IV) |  |
| *Ndufa10* | 0.70 | 1.66E-03 | ETC (Complex I) |  |
| *Cox19* | 0.70 | 1.52E-02 | ETC (Complex IV) |  |
| *Ndufs3* | 0.69 | 1.74E-02 | ETC (Complex I) |  |
| *Ndufb4* | 0.66 | 2.31E-02 | ETC (Complex I) |  |
| *Ndufaf2* | 0.65 | 3.37E-02 | ETC (Complex I) |  |
| *Uqcrh* | 0.59 | 7.19E-03 | ETC (Complex III) |  |
| *Ndufa4* | 0.57 | 1.01E-02 | ETC (Complex I) |  |
| *Ndufs2* | 0.54 | 1.54E-02 | ETC (Complex I) |  |
| *Cox7a1* | 0.50 | 5.45E-03 | ETC (Complex IV) |  |
| *St3gal3* | 4.59 | 3.38E-04 | Glycan |  |
| *B4galnt3* | 4.38 | 1.90E-02 | Glycan |  |
| *Galnt13* | 4.14 | 5.16E-04 | Glycan |  |
| *Mfng* | 4.02 | 4.28E-03 | Glycan |  |
| *St3gal5* | 3.44 | 5.76E-04 | Glycan |  |
| *Galns* | 2.85 | 5.51E-04 | Glycan (Degradation) |  |
| *Chst2* | 2.78 | 9.18E-03 | Glycan (Sulfate) |  |
| *B4galt5* | 2.37 | 5.85E-03 | Glycan |  |
| *B4galnt2* | 2.20 | 6.24E-03 | Glycan |  |
| *Sulf2* | 2.20 | 2.50E-02 | Glycan (Sulfate) |  |
| *B4galt6* | 2.05 | 3.02E-02 | Glycan |  |
| *Aga* | 2.01 | 3.26E-03 | Glycan (Degradation) |  |
| *Chst12* | 1.92 | 1.44E-03 | Glycan (Sulfate) |  |
| *Pomt2* | 1.91 | 2.25E-02 | Glycan |  |
| *Dolpp1* | 1.90 | 1.91E-04 | Glycan |  |
| *Mgat5* | 1.78 | 3.23E-03 | Glycan |  |
| *Man1a2* | 1.78 | 3.95E-02 | Glycan (Degradation) |  |
| *Man2b2* | 1.74 | 1.66E-03 | Glycan (Degradation) |  |
| *Pigg* | 1.71 | 7.08E-04 | Glycan (Anchor) |  |
| *Mgat4b* | 1.68 | 2.85E-02 | Glycan |  |
| *Chst11* | 1.66 | 2.64E-02 | Glycan (Sulfate) |  |
| *Neu3* | 1.64 | 7.71E-03 | Glycan (Degradation) |  |
| *Hexb* | 1.64 | 4.32E-02 | Glycan (Degradation) |  |
| *B3gnt2* | 1.63 | 4.06E-02 | Glycan |  |
| *St8sia2* | 1.62 | 1.28E-03 | Glycan |  |
| *Manea* | 1.62 | 3.53E-03 | Glycan (Degradation) |  |
| *Man1c1* | 1.62 | 8.85E-03 | Glycan (Degradation) |  |
| *Sulf1* | 1.60 | 4.98E-02 | Glycan (Sulfate) |  |
| *B3galt5* | 1.60 | 1.03E-02 | Glycan |  |
| *Mgat3* | 1.56 | 4.56E-03 | Glycan |  |
| *Chpf* | 1.53 | 3.74E-03 | Glycan (Sulfate) |  |
| *Galnt1* | 1.53 | 3.13E-02 | Glycan |  |
| *B3gat3* | 1.49 | 3.09E-04 | Glycan |  |
| *Galnt14* | 1.49 | 1.54E-02 | Glycan |  |
| *Large1* | 1.49 | 6.84E-03 | Glycan |  |
| *Xylt1* | 1.48 | 2.65E-03 | Glycan |  |
| *Alg2* | 1.47 | 3.17E-05 | Glycan |  |
| *Man2b1* | 1.46 | 9.73E-03 | Glycan (Degradation) |  |
| *St3gal2* | 1.46 | 1.29E-04 | Glycan |  |
| *Gns* | 1.45 | 1.36E-03 | Glycan (Degradation) |  |
| *Mgat5b* | 1.45 | 1.35E-02 | Glycan |  |
| *St8sia3* | 1.45 | 1.82E-02 | Glycan |  |
| *Fut9* | 1.42 | 2.87E-02 | Glycan |  |
| *Hyal2* | 1.42 | 2.55E-02 | Glycan (Degradation) |  |
| *Ids* | 1.37 | 1.24E-02 | Glycan (Degradation) |  |
| *B4galt4* | 1.36 | 1.92E-03 | Glycan |  |
| *Ganab* | 1.32 | 3.27E-04 | Glycan |  |
| *B4galt7* | 0.76 | 1.89E-03 | Glycan |  |
| *Pigb* | 0.76 | 2.84E-03 | Glycan (Anchor) |  |
| *Alg12* | 0.74 | 2.28E-02 | Glycan |  |
| *Dpm2* | 0.72 | 1.93E-02 | Glycan (Anchor) |  |
| *Alg6* | 0.72 | 4.54E-02 | Glycan |  |
| *Pomt1* | 0.71 | 1.22E-02 | Glycan |  |
| *Csgalnact2* | 0.70 | 1.80E-02 | Glycan (Sulfate) |  |
| *B4galt2* | 0.69 | 1.95E-03 | Glycan |  |
| *Chsy1* | 0.68 | 1.37E-02 | Glycan (Sulfate) |  |
| *B4galt3* | 0.66 | 2.07E-03 | Glycan |  |
| *Man2a2* | 0.64 | 2.41E-02 | Glycan (Degradation) |  |
| *Alg8* | 0.63 | 3.58E-02 | Glycan |  |
| *B3galnt2* | 0.61 | 2.48E-02 | Glycan |  |
| *Pigc* | 0.59 | 1.35E-02 | Glycan (Anchor) |  |
| *Chpf2* | 0.57 | 2.39E-02 | Glycan (Sulfate) |  |
| *Piga* | 0.57 | 4.66E-03 | Glycan (Anchor) |  |
| *B3gnt8* | 0.55 | 4.40E-04 | Glycan |  |
| *Edem2* | 0.54 | 2.78E-02 | Glycan (Degradation) |  |
| *Pigx* | 0.53 | 2.55E-03 | Glycan (Anchor) |  |
| *Pigm* | 0.53 | 9.45E-04 | Glycan (Anchor) |  |
| *Pigh* | 0.48 | 7.25E-04 | Glycan (Anchor) |  |
| *Pigz* | 0.47 | 4.50E-03 | Glycan (Anchor) |  |
| *Hexdc* | 0.46 | 1.10E-04 | Glycan (Degradation) |  |
| *St6galnac5* | 0.46 | 2.72E-04 | Glycan |  |
| *Galnt10* | 0.46 | 1.12E-04 | Glycan |  |
| *Galnt4* | 0.44 | 1.19E-03 | Glycan |  |
| *Glt8d1* | 0.43 | 1.00E-03 | Glycan |  |
| *Man2c1* | 0.42 | 8.61E-04 | Glycan (Degradation) |  |
| *Galnt7* | 0.37 | 3.83E-03 | Glycan |  |
| *Glb1l* | 0.37 | 5.91E-03 | Glycan (Degradation) |  |
| *Ctbs* | 0.34 | 5.25E-03 | Glycan |  |
| *St6galnac2* | 0.34 | 2.89E-04 | Glycan |  |
| *B3gnt6* | 0.27 | 2.60E-02 | Glycan |  |
| *St8sia1* | 0.23 | 5.73E-04 | Glycan |  |
| *St3gal1* | 0.17 | 8.69E-05 | Glycan |  |
| *B4galt1* | 0.14 | 5.55E-07 | Glycan |  |
| *Gcnt3* | 0.12 | 1.55E-02 | Glycan |  |
| *Chst3* | 0.07 | 2.62E-04 | Glycan (Sulfate) |  |
| *Glb1l2* | 0.05 | 3.49E-06 | Glycan (Degradation) |  |
| *Glb1l3* | 2.73E-03 | 2.82E-02 | Glycan (Degradation) |  |
| *Glo1* | 2.52 | 1.91E-04 | Glyoxylate |  |
| *Hs6st3* | 4.08 | 1.61E-03 | Heparin Sulfate |  |
| *Ndst1* | 1.86 | 6.80E-03 | Heparin Sulfate |  |
| *Extl2* | 1.75 | 8.00E-05 | Heparin Sulfate |  |
| *Extl3* | 1.67 | 3.04E-03 | Heparin Sulfate |  |
| *Hs3st1* | 1.65 | 2.16E-02 | Heparin Sulfate |  |
| *Hs2st1* | 1.59 | 1.64E-03 | Heparin Sulfate |  |
| *Hs3st5* | 1.34 | 3.28E-02 | Heparin Sulfate |  |
| *Hs6st1* | 1.31 | 8.78E-03 | Heparin Sulfate |  |
| *Hs3st3b1* | 0.63 | 2.51E-02 | Heparin Sulfate |  |
| *Ndst2* | 0.55 | 1.10E-03 | Heparin Sulfate |  |
| *Ext1* | 0.30 | 5.32E-04 | Heparin Sulfate |  |
| *Chrnb4* | 18.82 | 2.42E-02 | Ion Transport |  |
| *Cngb3* | 14.36 | 1.97E-04 | Ion Transport |  |
| *Cnga3* | 13.92 | 5.55E-05 | Ion Transport |  |
| *Clca1* | 13.75 | 1.16E-03 | Ion Transport |  |
| *Kcng1* | 7.43 | 4.42E-02 | Ion Transport |  |
| *Slc24a4* | 5.81 | 3.57E-04 | Ion Transport |  |
| *Kcng3* | 5.70 | 1.33E-02 | Ion Transport |  |
| *Grik3* | 4.93 | 3.09E-02 | Ion Transport |  |
| *Slc24a2* | 4.51 | 4.58E-04 | Ion Transport |  |
| *Slc8a3* | 4.26 | 3.25E-03 | Ion Transport |  |
| *Clcnka* | 4.08 | 4.57E-02 | Ion Transport |  |
| *Ttyh2* | 4.03 | 3.16E-05 | Ion Transport |  |
| *Glra1* | 4.02 | 1.79E-02 | Ion Transport |  |
| *Kcnh5* | 3.49 | 2.01E-02 | Ion Transport |  |
| *Clic4* | 3.19 | 2.22E-04 | Ion Transport |  |
| *Kcnh6* | 2.97 | 1.63E-04 | Ion Transport |  |
| *Orai1* | 2.94 | 1.68E-03 | Ion Transport |  |
| *P2rx5* | 2.80 | 1.27E-02 | Ion Transport |  |
| *Kcne2* | 2.75 | 8.49E-03 | Ion Transport |  |
| *Cacna1h* | 2.71 | 9.25E-06 | Ion Transport |  |
| *Slc8a1* | 2.57 | 1.15E-02 | Ion Transport |  |
| *Kcnab1* | 2.55 | 1.35E-03 | Ion Transport |  |
| *Gabra2* | 2.50 | 4.57E-04 | Ion Transport |  |
| *Trpm8* | 2.50 | 2.50E-02 | Ion Transport |  |
| *Kcnd3* | 2.36 | 2.57E-04 | Ion Transport |  |
| *Kcnj6* | 2.34 | 4.20E-02 | Ion Transport |  |
| *Kcng4* | 2.33 | 1.55E-03 | Ion Transport |  |
| *Kctd12* | 2.30 | 1.22E-03 | Ion Transport |  |
| *Kcnt2* | 2.24 | 8.76E-03 | Ion Transport |  |
| *Kcna1* | 2.20 | 2.29E-02 | Ion Transport |  |
| *Atp2b3* | 2.11 | 7.31E-03 | Ion Transport |  |
| *Chrna7* | 2.10 | 8.58E-03 | Ion Transport |  |
| *Gabrb2* | 2.10 | 2.65E-02 | Ion Transport |  |
| *Gabrb3* | 2.09 | 8.92E-05 | Ion Transport |  |
| *Atp2a2* | 2.03 | 4.69E-03 | Ion Transport |  |
| *Atp1b2* | 2.02 | 5.61E-06 | Ion Transport |  |
| *Best3* | 2.01 | 2.24E-02 | Ion Transport |  |
| *Kcnj10* | 1.98 | 8.10E-03 | Ion Transport |  |
| *Slc12a4* | 1.97 | 5.11E-03 | Ion Transport |  |
| *Kcnf1* | 1.95 | 9.03E-04 | Ion Transport |  |
| *Hcn4* | 1.92 | 3.47E-03 | Ion Transport |  |
| *Slc22a23* | 1.92 | 1.28E-02 | Ion Transport |  |
| *Kcnab2* | 1.89 | 5.41E-04 | Ion Transport |  |
| *Cacng8* | 1.84 | 1.88E-02 | Ion Transport |  |
| *Kcnk12* | 1.84 | 2.37E-02 | Ion Transport |  |
| *Kcnq4* | 1.83 | 3.33E-02 | Ion Transport |  |
| *Gabra5* | 1.82 | 1.10E-03 | Ion Transport |  |
| *Atp2b2* | 1.80 | 6.88E-03 | Ion Transport |  |
| *Itpr1* | 1.78 | 1.96E-02 | Ion Transport |  |
| *Clcn3* | 1.78 | 6.93E-04 | Ion Transport |  |
| *Kcnc3* | 1.78 | 7.71E-03 | Ion Transport |  |
| *Slc12a7* | 1.76 | 2.25E-02 | Ion Transport |  |
| *Grid2* | 1.71 | 1.01E-02 | Ion Transport |  |
| *Slc39a11* | 1.68 | 2.29E-02 | Ion Transport |  |
| *Ttyh3* | 1.68 | 2.50E-03 | Ion Transport |  |
| *Kcnj12* | 1.68 | 4.02E-03 | Ion Transport |  |
| *Grik4* | 1.67 | 3.66E-02 | Ion Transport |  |
| *Heph* | 1.66 | 1.54E-04 | Ion Transport |  |
| *Grin2a* | 1.64 | 1.67E-02 | Ion Transport |  |
| *Scn5a* | 1.64 | 3.88E-02 | Ion Transport |  |
| *Slc39a14* | 1.62 | 4.21E-02 | Ion Transport |  |
| *Slc12a5* | 1.61 | 1.05E-02 | Ion Transport |  |
| *Kctd10* | 1.61 | 2.49E-03 | Ion Transport |  |
| *Kcnk3* | 1.61 | 1.84E-02 | Ion Transport |  |
| *Kcnh1* | 1.59 | 2.30E-03 | Ion Transport |  |
| *Gabrb1* | 1.56 | 1.74E-02 | Ion Transport |  |
| *Slc24a3* | 1.56 | 1.40E-02 | Ion Transport |  |
| *Slc8a2* | 1.56 | 2.94E-02 | Ion Transport |  |
| *Chrnb2* | 1.55 | 1.60E-03 | Ion Transport |  |
| *Kcna2* | 1.55 | 1.07E-02 | Ion Transport |  |
| *Nalcn* | 1.54 | 2.93E-02 | Ion Transport |  |
| *Gria1* | 1.52 | 2.52E-02 | Ion Transport |  |
| *Kctd9* | 1.50 | 3.17E-02 | Ion Transport |  |
| *Pkd2* | 1.49 | 1.88E-03 | Ion Transport |  |
| *Slc30a9* | 1.49 | 4.08E-03 | Ion Transport |  |
| *Cacng2* | 1.49 | 1.67E-02 | Ion Transport |  |
| *Kcnd2* | 1.46 | 8.30E-03 | Ion Transport |  |
| *Grid1* | 1.46 | 2.40E-02 | Ion Transport |  |
| *Slc39a7* | 1.45 | 6.52E-03 | Ion Transport |  |
| *Cacng4* | 1.45 | 3.63E-02 | Ion Transport |  |
| *Grin1* | 1.45 | 1.65E-02 | Ion Transport |  |
| *Slc39a6* | 1.45 | 4.83E-03 | Ion Transport |  |
| *Kcnj11* | 1.44 | 2.92E-02 | Ion Transport |  |
| *Kctd1* | 1.43 | 3.86E-03 | Ion Transport |  |
| *Clcn6* | 1.42 | 6.88E-03 | Ion Transport |  |
| *Atp1a3* | 1.41 | 2.60E-02 | Ion Transport |  |
| *Hcn2* | 1.40 | 2.52E-02 | Ion Transport |  |
| *Slc9a6* | 1.39 | 7.07E-04 | Ion Transport |  |
| *Slc30a1* | 1.38 | 6.49E-03 | Ion Transport |  |
| *P2rx6* | 1.37 | 2.94E-02 | Ion Transport |  |
| *Slc41a3* | 1.36 | 9.20E-03 | Ion Transport |  |
| *Nipa1* | 1.35 | 2.22E-02 | Ion Transport |  |
| *Slc39a1* | 1.35 | 1.44E-02 | Ion Transport |  |
| *Clcn7* | 1.34 | 4.20E-03 | Ion Transport |  |
| *Chrna3* | 1.33 | 1.21E-02 | Ion Transport |  |
| *Scn2b* | 1.33 | 1.01E-02 | Ion Transport |  |
| *Kcnc1* | 1.32 | 6.20E-03 | Ion Transport |  |
| *Kcnc4* | 1.31 | 1.72E-02 | Ion Transport |  |
| *Kctd15* | 1.31 | 2.45E-02 | Ion Transport |  |
| *Cacna2d4* | 0.75 | 2.60E-02 | Ion Transport |  |
| *Mfsd8* | 0.75 | 3.64E-02 | Ion Transport |  |
| *Orai2* | 0.74 | 1.55E-02 | Ion Transport |  |
| *Slc30a6* | 0.72 | 4.49E-03 | Ion Transport |  |
| *Slc39a13* | 0.72 | 8.39E-03 | Ion Transport |  |
| *Trpc6* | 0.71 | 2.71E-02 | Ion Transport |  |
| *Mcoln1* | 0.68 | 2.02E-02 | Ion Transport |  |
| *Kctd6* | 0.68 | 1.20E-02 | Ion Transport |  |
| *Pkd1* | 0.65 | 1.66E-02 | Ion Transport |  |
| *Cacna2d3* | 0.63 | 1.63E-02 | Ion Transport |  |
| *Slc12a9* | 0.62 | 1.97E-02 | Ion Transport |  |
| *Gria2* | 0.61 | 4.75E-02 | Ion Transport |  |
| *Slc30a7* | 0.61 | 2.03E-03 | Ion Transport |  |
| *Tpcn2* | 0.60 | 3.41E-02 | Ion Transport |  |
| *Kctd3* | 0.60 | 2.10E-03 | Ion Transport |  |
| *Chrna5* | 0.58 | 1.35E-02 | Ion Transport |  |
| *Grin2d* | 0.58 | 1.95E-02 | Ion Transport |  |
| *Kcnv2* | 0.57 | 5.54E-03 | Ion Transport |  |
| *Slc9a8* | 0.57 | 6.26E-04 | Ion Transport |  |
| *Mmaa* | 0.56 | 6.68E-03 | Ion Transport |  |
| *Trpc1* | 0.55 | 9.22E-03 | Ion Transport |  |
| *Cacnb2* | 0.53 | 9.09E-03 | Ion Transport |  |
| *Kcnb1* | 0.51 | 2.78E-04 | Ion Transport |  |
| *Cacna1f* | 0.51 | 4.76E-03 | Ion Transport |  |
| *Kctd18* | 0.51 | 2.35E-03 | Ion Transport |  |
| *Trpm7* | 0.51 | 2.47E-03 | Ion Transport |  |
| *Ryr3* | 0.49 | 3.78E-02 | Ion Transport |  |
| *Atp11c* | 0.49 | 1.16E-04 | Ion Transport |  |
| *Clcn1* | 0.49 | 9.87E-03 | Ion Transport |  |
| *Slc12a3* | 0.48 | 2.54E-02 | Ion Transport |  |
| *Catsper2* | 0.46 | 2.73E-03 | Ion Transport |  |
| *Slc12a8* | 0.43 | 1.67E-02 | Ion Transport |  |
| *Slc9a5* | 0.42 | 5.10E-04 | Ion Transport |  |
| *Scn4a* | 0.39 | 1.71E-02 | Ion Transport |  |
| *Slc24a5* | 0.39 | 5.05E-05 | Ion Transport |  |
| *Kcnd1* | 0.39 | 1.08E-04 | Ion Transport |  |
| *Slc31a2* | 0.38 | 3.02E-04 | Ion Transport |  |
| *Clic3* | 0.37 | 2.23E-02 | Ion Transport |  |
| *Slc3a1* | 0.33 | 1.19E-02 | Ion Transport |  |
| *Atp11b* | 0.32 | 5.78E-04 | Ion Transport |  |
| *Atp7b* | 0.27 | 8.69E-03 | Ion Transport |  |
| *Slc39a2* | 0.18 | 4.59E-03 | Ion Transport |  |
| *Atp4a* | 0.04 | 5.05E-04 | Ion Transport |  |
| *Cngb1* | 0.02 | 2.39E-04 | Ion Transport |  |
| *Slc24a1* | 0.01 | 3.05E-05 | Ion Transport |  |
| *Kcnj14* | 4.62E-03 | 5.14E-05 | Ion Transport |  |
| *Cnga1* | 2.49E-04 | 1.08E-06 | Ion Transport |  |
| *Oxct1* | 1.48 | 2.86E-02 | Ketone Bodies |  |
| *Hmgcl* | 0.65 | 1.94E-03 | Ketone Bodies |  |
| *Aacs* | 0.62 | 3.18E-02 | Ketone Bodies |  |
| *Nadk* | 1.32 | 1.23E-02 | NAD |  |
| *Nox1* | 0.62 | 2.35E-02 | NAD |  |
| *Duox2* | 0.09 | 1.64E-02 | NAD |  |
| *Bst1* | 0.02 | 4.64E-04 | NAD |  |
| *Abat* | 1.52 | 7.40E-03 | Neurotransmitter |  |
| *Ache* | 1.43 | 1.39E-02 | Neurotransmitter |  |
| *Comtd1* | 0.54 | 1.47E-02 | Neurotransmitter |  |
| *Ddc* | 0.46 | 2.38E-02 | Neurotransmitter |  |
| *Pde6c* | 9.91 | 5.16E-04 | Nucleotide (cGMP) |  |
| *Pde6h* | 4.76 | 1.56E-03 | Nucleotide (cGMP) |  |
| *Ampd2* | 4.01 | 6.89E-05 | Nucleotide (Purine) |  |
| *Adcy2* | 3.47 | 4.62E-06 | Nucleotide (cAMP) |  |
| *Oasl2* | 2.94 | 3.53E-02 | Nucleotide (Ribonuclease) |  |
| *Gucy1a2* | 2.58 | 4.22E-02 | Nucleotide (cGMP) |  |
| *Adcy9* | 2.07 | 1.35E-02 | Nucleotide (cAMP) |  |
| *Pde4d* | 2.01 | 1.11E-02 | Nucleotide (cGMP) |  |
| *Uprt* | 1.98 | 2.62E-02 | Nucleotide (Pyrimidine) |  |
| *Pde3a* | 1.90 | 6.59E-03 | Nucleotide (cGMP) |  |
| *Npr3* | 1.75 | 4.77E-02 | Nucleotide (cGMP) |  |
| *Pde11a* | 1.75 | 3.66E-02 | Nucleotide (cyclic nucleotide) |  |
| *Pde7b* | 1.68 | 5.83E-03 | Nucleotide (cyclic nucleotide) |  |
| *Dpysl3* | 1.64 | 3.22E-02 | Nucleotide (Pyrimidine) |  |
| *Uck2* | 1.63 | 3.01E-02 | Nucleotide (Pyrimidine) |  |
| *Dpysl5* | 1.63 | 2.03E-02 | Nucleotide (Pyrimidine) |  |
| *Upp2* | 1.59 | 1.86E-02 | Nucleotide (Pyrimidine) |  |
| *Nt5dc3* | 1.56 | 5.53E-03 | Nucleotide |  |
| *Dpyd* | 1.49 | 2.61E-02 | Nucleotide (Pyrimidine) |  |
| *Ampd3* | 1.44 | 4.56E-02 | Nucleotide (Purine) |  |
| *Bpnt1* | 1.39 | 4.06E-03 | Nucleotide |  |
| *Prps2* | 1.35 | 1.58E-02 | Nucleotide (Purine) |  |
| *Rrm2b* | 1.34 | 2.47E-02 | Nucleotide |  |
| *Atic* | 1.34 | 5.16E-03 | Nucleotide (Purine) |  |
| *Dpysl2* | 1.33 | 2.43E-02 | Nucleotide (Pyrimidine) |  |
| *Nme1* | 1.32 | 3.47E-02 | Nucleotide |  |
| *Adcy5* | 1.32 | 1.18E-02 | Nucleotide (cAMP) |  |
| *Pde4a* | 1.31 | 2.74E-02 | Nucleotide (cGMP) |  |
| *Guk1* | 1.31 | 3.00E-02 | Nucleotide (Purine) |  |
| *Adcy8* | 1.30 | 4.38E-02 | Nucleotide (cAMP) |  |
| *Nudt16* | 0.77 | 7.64E-03 | Nucleotide |  |
| *Tk2* | 0.71 | 4.69E-03 | Nucleotide (Pyrimidine) |  |
| *Adal* | 0.70 | 1.61E-02 | Nucleotide (Purine) |  |
| *Gmpr2* | 0.70 | 2.18E-02 | Nucleotide (Purine) |  |
| *Impdh2* | 0.69 | 8.94E-03 | Nucleotide (Purine) |  |
| *Enpp1* | 0.69 | 2.42E-02 | Nucleotide |  |
| *Pde9a* | 0.69 | 5.24E-03 | Nucleotide (cGMP) |  |
| *Pde7a* | 0.66 | 3.27E-02 | Nucleotide (cyclic nucleotide) |  |
| *Ak3* | 0.65 | 2.27E-03 | Nucleotide (Purine) |  |
| *Dhodh* | 0.64 | 5.36E-04 | Nucleotide (Pyrimidine) |  |
| *Nme3* | 0.64 | 7.68E-03 | Nucleotide |  |
| *Ak2* | 0.62 | 2.95E-03 | Nucleotide (Purine) |  |
| *Nme6* | 0.62 | 1.75E-03 | Nucleotide |  |
| *Pfas* | 0.62 | 1.21E-02 | Nucleotide (Purine) |  |
| *Dguok* | 0.59 | 3.95E-03 | Nucleotide (Purine) |  |
| *Adcy7* | 0.58 | 7.29E-04 | Nucleotide (cAMP) |  |
| *Dtymk* | 0.57 | 2.13E-03 | Nucleotide (Pyrimidine) |  |
| *Adcy6* | 0.54 | 1.00E-02 | Nucleotide (cAMP) |  |
| *Entpd4* | 0.52 | 1.56E-02 | Nucleotide |  |
| *Uckl1* | 0.50 | 6.11E-03 | Nucleotide (Pyrimidine) |  |
| *Pde4c* | 0.47 | 4.99E-02 | Nucleotide (cGMP) |  |
| *Nudt13* | 0.37 | 6.80E-03 | Nucleotide |  |
| *Nme5* | 0.37 | 1.43E-03 | Nucleotide |  |
| *Pde6g* | 0.25 | 8.19E-04 | Nucleotide (cGMP) |  |
| *Nt5e* | 0.17 | 1.16E-06 | Nucleotide |  |
| *Pde6a* | 0.07 | 8.92E-05 | Nucleotide (cGMP) |  |
| *Pde6b* | 0.02 | 3.35E-05 | Nucleotide (cGMP) |  |
| *Gucy2f* | 0.02 | 4.25E-03 | Nucleotide (cGMP) |  |
| *Alad* | 2.17 | 5.60E-04 | Porphyrin |  |
| *Cpox* | 0.74 | 4.13E-03 | Porphyrin |  |
| *Hmbs* | 0.66 | 3.20E-02 | Porphyrin |  |
| *Urod* | 0.54 | 1.98E-03 | Porphyrin |  |
| *Tap1* | 2.15 | 3.04E-02 | Proton Transport |  |
| *Tap2* | 1.97 | 1.92E-02 | Proton Transport |  |
| *Atp6v0a1* | 1.93 | 3.13E-05 | Proton Transport |  |
| *Atp9a* | 1.69 | 2.46E-03 | Proton Transport |  |
| *Atp10a* | 1.63 | 2.20E-02 | Proton Transport |  |
| *Atp13a3* | 1.58 | 1.30E-02 | Proton Transport |  |
| *Atp6v1a* | 1.45 | 2.12E-03 | Proton Transport |  |
| *Atp6v1b2* | 1.37 | 6.14E-04 | Proton Transport |  |
| *Atp6v0a2* | 1.31 | 3.61E-02 | Proton Transport |  |
| *Atp9b* | 0.77 | 4.57E-02 | Proton Transport |  |
| *Atp5l* | 0.71 | 4.86E-02 | Proton Transport |  |
| *Slc25a27* | 0.59 | 3.29E-03 | Proton Transport |  |
| *Cybb* | 8.87 | 3.27E-02 | Redox |  |
| *Frrs1* | 3.02 | 1.30E-02 | Redox |  |
| *Cyb561* | 2.13 | 2.47E-03 | Redox |  |
| *Dhrs11* | 1.53 | 2.43E-02 | Redox |  |
| *Cyba* | 1.46 | 3.52E-02 | Redox |  |
| *Enox2* | 1.46 | 4.05E-02 | Redox |  |
| *Cyb5b* | 1.31 | 7.12E-03 | Redox |  |
| *Cyb5a* | 0.72 | 3.26E-02 | Redox |  |
| *Cybrd1* | 0.64 | 2.42E-02 | Redox |  |
| *Foxred1* | 0.60 | 2.21E-03 | Redox |  |
| *Ogfod2* | 0.59 | 2.65E-03 | Redox |  |
| *Enox1* | 0.54 | 1.62E-02 | Redox |  |
| *Dhrs7* | 0.49 | 1.16E-02 | Redox |  |
| *Fdxr* | 0.39 | 2.15E-03 | Redox |  |
| *Cyb5r1* | 0.31 | 6.49E-04 | Redox |  |
| *Dhrs3* | 0.16 | 1.90E-04 | Redox |  |
| *Lalba* | 6.89 | 4.70E-03 | Sugar (Lactose) |  |
| *Impa2* | 5.15 | 7.08E-03 | Sugar (Inositol Phosphate) |  |
| *Inppl1* | 3.31 | 2.18E-04 | Sugar (Inositol Phosphate) |  |
| *H6pd* | 3.05 | 1.78E-03 | Sugar |  |
| *Inpp4a* | 1.82 | 1.20E-02 | Sugar (Inositol Phosphate) |  |
| *Itpkb* | 1.73 | 3.40E-02 | Sugar (Inositol Phosphate) |  |
| *Gfod1* | 1.69 | 7.85E-03 | Sugar |  |
| *Pik3cb* | 1.64 | 4.94E-04 | Sugar (Inositol Phosphate) |  |
| *Impad1* | 1.64 | 7.14E-04 | Sugar (Inositol Phosphate) |  |
| *Pip5k1a* | 1.57 | 1.62E-04 | Sugar (Inositol Phosphate) |  |
| *Pip5k1b* | 1.47 | 8.50E-04 | Sugar (Inositol Phosphate) |  |
| *Inpp4b* | 1.45 | 1.68E-02 | Sugar (Inositol Phosphate) |  |
| *Siae* | 1.44 | 4.57E-02 | Sugar |  |
| *Pik3r3* | 1.37 | 3.45E-02 | Sugar (Inositol Phosphate) |  |
| *Pip4k2b* | 1.35 | 9.68E-03 | Sugar (Inositol Phosphate) |  |
| *Pik3c3* | 1.33 | 3.10E-03 | Sugar (Inositol Phosphate) |  |
| *Renbp* | 1.33 | 1.56E-03 | Sugar (Aminosugar) |  |
| *Pi4k2a* | 1.32 | 1.04E-02 | Sugar (Inositol Phosphate) |  |
| *Gnpnat1* | 1.31 | 7.65E-03 | Sugar (Aminosugar) |  |
| *Sord* | 1.31 | 6.75E-04 | Sugar |  |
| *Pgls* | 0.74 | 2.22E-02 | Sugar (PPP) |  |
| *Hyi* | 0.72 | 1.57E-02 | Sugar |  |
| *Pmm2* | 0.69 | 3.98E-02 | Sugar (Mannose) |  |
| *Tsta3* | 0.68 | 2.60E-03 | Sugar |  |
| *Gnptg* | 0.66 | 9.70E-03 | Sugar (Aminosugar) |  |
| *Tgds* | 0.66 | 9.04E-03 | Sugar (Nucleotide Sugar) |  |
| *Pik3c2a* | 0.65 | 1.49E-02 | Sugar (Inositol Phosphate) |  |
| *Rbks* | 0.64 | 4.21E-02 | Sugar |  |
| *Inpp5k* | 0.62 | 2.65E-05 | Sugar (Inositol Phosphate) |  |
| *Ppa1* | 0.61 | 1.42E-02 | Sugar (PPi hydrolysis) |  |
| *Pi4k2b* | 0.60 | 4.74E-02 | Sugar (Inositol Phosphate) |  |
| *Inpp5b* | 0.57 | 6.93E-03 | Sugar (Inositol Phosphate) |  |
| *Pmm1* | 0.56 | 8.50E-06 | Sugar (Mannose) |  |
| *Rpia* | 0.56 | 9.88E-03 | Sugar (PPP) |  |
| *Khk* | 0.53 | 6.14E-04 | Sugar (Fructose) |  |
| *Pip4k2c* | 0.41 | 8.66E-05 | Sugar (Inositol Phosphate) |  |
| *Dera* | 0.38 | 9.11E-04 | Sugar (PPP) |  |
| *Amy2a5* | 0.37 | 3.54E-02 | Sugar (Amylase) |  |
| *Isyna1* | 0.35 | 3.29E-02 | Sugar (Inositol Phosphate) |  |
| *Ip6k3* | 0.33 | 4.77E-02 | Sugar (Inositol Phosphate) |  |
| *Chia1* | 0.29 | 4.44E-03 | Sugar (Aminosugar) |  |
| *Amy2a2* | 0.23 | 3.68E-02 | Sugar (Amylase) |  |
| *Amy2a3* | 0.23 | 3.68E-02 | Sugar (Amylase) |  |
| *Amy2a4* | 0.23 | 3.68E-02 | Sugar (Amylase) |  |
| *Arsb* | 1.54 | 1.41E-02 | Sulfate |  |
| *Arsg* | 0.59 | 4.63E-02 | Sulfate |  |
| *Sult1a1* | 0.38 | 2.41E-03 | Sulfate |  |
| *Coq6* | 0.74 | 3.26E-02 | Ubiquinone |  |
| *Coq9* | 0.71 | 2.76E-02 | Ubiquinone |  |
| *Coq2* | 0.68 | 5.58E-03 | Ubiquinone |  |
| *Coq10a* | 0.65 | 1.06E-03 | Ubiquinone |  |
| *Pdss1* | 0.41 | 7.88E-06 | Ubiquinone |  |
| *Folh1* | 1.84 | 3.78E-02 | Vitamin (Folate) |  |
| *Akr1b10* | 1.47 | 4.75E-03 | Vitamine (Retinoic acid, Alcohol) |  |
| *Pdxk* | 1.44 | 1.84E-02 | Vitamin (B6) |  |
| *Rdh11* | 0.74 | 1.82E-02 | Vitamin (A) |  |
| *Phospho2* | 0.67 | 1.48E-02 | Vitamin (B6) |  |
| *Adh6b* | 0.66 | 4.37E-02 | Vitamine (Retinoic acid, Alcohol) |  |
| *Rdh8* | 0.63 | 2.73E-02 | Vitamin (A) |  |
| *Dhfr* | 0.57 | 1.51E-02 | Vitamin (Folate) |  |
| *Aldh8a1* | 0.36 | 1.54E-02 | Vitamine (Retinoic acid, Alcohol) |  |
| *Rdh12* | 0.18 | 4.34E-05 | Vitamin (A) |  |

**Table S5.** **Donor information for human retina and RPE/choroid explants.**

| **Donor ID** | **Eye (s)** | **Gender** | **Age** | **Post-mortem Delay** | **Known eye diseases** |
| --- | --- | --- | --- | --- | --- |
| 190800 | both | F | 66y | 19h | No |
| 190820 | both | M | 83y | 28h | No |
| 190850 | both | F | 77y | 24h | No |
| 190853 | both | F | 71y | 28h | No |

**Table S6.** **Medium formulation for retinal organoid culture.**

| **Components** | **Concentration (mg/L)** | **mM** | **Category** | **Source** |
| --- | --- | --- | --- | --- |
| Glycine | 31.88 | 0.43 | Amino Acids | DMEM, high glucose, GlutaMAX™ + DMEM/F-12, HEPES + NEAA |
| L-Alanine | 11.12 | 0.12 | Amino Acids | DMEM/F-12, HEPES + NEAA |
| L-Alanyl-Glutamine (GlutaMAX) | 431.00 | 1.99 | Amino Acids | DMEM, high glucose, GlutaMAX™ |
| L-Arginine hydrochloride | 115.75 | 0.55 | Amino Acids | DMEM, high glucose, GlutaMAX™ + DMEM/F-12, HEPES |
| L-Asparagine | 16.50 | 0.13 | Amino Acids | DMEM/F-12, HEPES + NEAA |
| L-Aspartic acid | 16.63 | 0.13 | Amino Acids | DMEM/F-12, HEPES + NEAA |
| L-Cysteine hydrochloride-H2O | 8.78 | 0.05 | Amino Acids | DMEM/F-12, HEPES |
| L-Cystine 2HCl | 47.15 | 0.15 | Amino Acids | DMEM, high glucose, GlutaMAX™ + DMEM/F-12, HEPES |
| L-Glutamic Acid | 18.38 | 0.13 | Amino Acids | DMEM/F-12, HEPES + NEAA |
| L-Glutamine | 182.50 | 1.25 | Amino Acids | DMEM/F-12, HEPES |
| L-Histidine hydrochloride-H2O | 36.74 | 0.17 | Amino Acids | DMEM, high glucose, GlutaMAX™ + DMEM/F-12, HEPES |
| L-Isoleucine | 79.74 | 0.61 | Amino Acids | DMEM, high glucose, GlutaMAX™ + DMEM/F-12, HEPES |
| L-Leucine | 82.03 | 0.63 | Amino Acids | DMEM, high glucose, GlutaMAX™ + DMEM/F-12, HEPES |
| L-Lysine hydrochloride | 118.63 | 0.65 | Amino Acids | DMEM, high glucose, GlutaMAX™ + DMEM/F-12, HEPES |
| L-Methionine | 23.62 | 0.16 | Amino Acids | DMEM, high glucose, GlutaMAX™ + DMEM/F-12, HEPES |
| L-Phenylalanine | 50.74 | 0.31 | Amino Acids | DMEM, high glucose, GlutaMAX™ + DMEM/F-12, HEPES |
| L-Proline | 20.13 | 0.18 | Amino Acids | DMEM/F-12, HEPES + NEAA |
| L-Serine | 44.63 | 0.43 | Amino Acids | DMEM, high glucose, GlutaMAX™ + DMEM/F-12, HEPES + NEAA |
| L-Threonine | 74.23 | 0.62 | Amino Acids | DMEM, high glucose, GlutaMAX™ + DMEM/F-12, HEPES |
| L-Tryptophan | 12.51 | 0.06 | Amino Acids | DMEM, high glucose, GlutaMAX™ + DMEM/F-12, HEPES |
| L-Tyrosine disodium salt dihydrate | 79.90 | 0.31 | Amino Acids | DMEM, high glucose, GlutaMAX™ + DMEM/F-12, HEPES |
| L-Valine | 73.43 | 0.63 | Amino Acids | DMEM, high glucose, GlutaMAX™ + DMEM/F-12, HEPES |
| Biotin | 1.75E-03 | 7.17E-06 | Vitamins | DMEM/F-12, HEPES |
| Choline chloride | 6.49 | 0.05 | Vitamins | DMEM, high glucose, GlutaMAX™ + DMEM/F-12, HEPES |
| D-Calcium pantothenate | 3.12 | 0.01 | Vitamins | DMEM, high glucose, GlutaMAX™ + DMEM/F-12, HEPES |
| Folic Acid | 3.33 | 0.01 | Vitamins | DMEM, high glucose, GlutaMAX™ + DMEM/F-12, HEPES |
| i-Inositol | 9.90 | 0.06 | Vitamins | DMEM, high glucose, GlutaMAX™ + DMEM/F-12, HEPES |
| Niacinamide | 3.01 | 0.02 | Vitamins | DMEM, high glucose, GlutaMAX™ + DMEM/F-12, HEPES |
| Pyridoxal hydrochloride | 2.00 | 0.01 | Vitamins | DMEM, high glucose, GlutaMAX™ |
| Pyridoxine hydrochloride | 1.00 | 4.85E-03 | Vitamins | DMEM/F-12, HEPES |
| Riboflavin | 0.31 | 8.23E-04 | Vitamins | DMEM, high glucose, GlutaMAX™ + DMEM/F-12, HEPES |
| Thiamine hydrochloride | 3.09 | 0.01 | Vitamins | DMEM, high glucose, GlutaMAX™ + DMEM/F-12, HEPES |
| Vitamin B12 | 0.34 | 2.51E-04 | Vitamins | DMEM/F-12, HEPES |
| Calcium Chloride (CaCl2) (anhyd.) | 158.30 | 1.43 | Inorganic Salts | DMEM, high glucose, GlutaMAX™ + DMEM/F-12, HEPES |
| Cupric sulfate (CuSO4-5H2O) | 6.50E-04 | 2.60E-06 | Inorganic Salts | DMEM/F-12, HEPES |
| Ferric Nitrate (Fe(NO3)3"9H2O) | 0.08 | 1.86E-04 | Inorganic Salts | DMEM, high glucose, GlutaMAX™ + DMEM/F-12, HEPES |
| Ferrous Sulfate (FeSO4 7H2O) | 0.21 | 7.50E-04 | Inorganic Salts | DMEM/F-12, HEPES |
| Magnesium Chloride (anhydrous) | 14.32 | 0.15 | Inorganic Salts | DMEM/F-12, HEPES |
| Magnesium Sulfate (MgSO4) (anhyd.) | 73.26 | 0.61 | Inorganic Salts | DMEM, high glucose, GlutaMAX™ + DMEM/F-12, HEPES |
| Potassium Chloride (KCl) | 355.90 | 4.75 | Inorganic Salts | DMEM, high glucose, GlutaMAX™ + DMEM/F-12, HEPES |
| Sodium Bicarbonate (NaHCO3) | 2450.00 | 29.17 | Inorganic Salts | DMEM, high glucose, GlutaMAX™ + DMEM/F-12, HEPES |
| Sodium Chloride (NaCl) | 6697.75 | 115.48 | Inorganic Salts | DMEM, high glucose, GlutaMAX™ + DMEM/F-12, HEPES |
| Sodium Phosphate dibasic (Na2HPO4) anhydrous | 35.51 | 0.25 | Inorganic Salts | DMEM/F-12, HEPES |
| Sodium Phosphate monobasic (NaH2PO4-H2O) | 93.75 | 0.68 | Inorganic Salts | DMEM, high glucose, GlutaMAX™ + DMEM/F-12, HEPES |
| Zinc sulfate (ZnSO4-7H2O) | 0.22 | 7.50E-04 | Inorganic Salts | DMEM/F-12, HEPES |
| D-Glucose (Dextrose) | 3825.50 | 21.25 | Other Components | DMEM, high glucose, GlutaMAX™ + DMEM/F-12, HEPES |
| HEPES | 1787.25 | 7.51 | Other Components | DMEM/F-12, HEPES |
| Hypoxanthine Na | 1.20 | 0.01 | Other Components | DMEM/F-12, HEPES |
| Linoleic Acid | 0.02 | 7.50E-05 | Other Components | DMEM/F-12, HEPES |
| Lipoic Acid | 0.05 | 2.55E-04 | Other Components | DMEM/F-12, HEPES |
| Phenol Red | 11.55 | 0.03 | Other Components | DMEM, high glucose, GlutaMAX™ + DMEM/F-12, HEPES |
| Putrescine 2HCl | 0.04 | 2.52E-04 | Other Components | DMEM/F-12, HEPES |
| Sodium Pyruvate | 27.50 | 0.25 | Other Components | DMEM/F-12, HEPES |
| Thymidine | 0.18 | 7.54E-04 | Other Components | DMEM/F-12, HEPES |
| Taurine | 12.52 | 0.10 |  | Supplement |
| B-27™ Supplement (50X), serum free | 1X |  |  | Supplement |
| Fetal Bovine Serum | 8% (V/V) |  |  | Supplement |

**Table S7.** **Medium formulation for mouse retinal explant culture.**

| **Components** | **Concentration (mg/L)** | **mM** | **Category** | **Source** |
| --- | --- | --- | --- | --- |
| Glycine | 26.25 | 0.35 | Amino Acids | DMEM/F-12, HEPES + NEAA |
| L-Alanine | 13.35 | 0.15 | Amino Acids | DMEM/F-12, HEPES + NEAA |
| L-Arginine hydrochloride | 147.50 | 0.70 | Amino Acids | DMEM/F-12, HEPES |
| L-Asparagine | 19.80 | 0.15 | Amino Acids | DMEM/F-12, HEPES + NEAA |
| L-Aspartic acid | 19.95 | 0.15 | Amino Acids | DMEM/F-12, HEPES + NEAA |
| L-Cysteine hydrochloride-H2O | 17.56 | 0.10 | Amino Acids | DMEM/F-12, HEPES |
| L-Cystine 2HCl | 31.29 | 0.10 | Amino Acids | DMEM/F-12, HEPES |
| L-Glutamic Acid | 22.05 | 0.15 | Amino Acids | DMEM/F-12, HEPES + NEAA |
| L-Glutamine | 365.00 | 2.50 | Amino Acids | DMEM/F-12, HEPES |
| L-Histidine hydrochloride-H2O | 31.48 | 0.15 | Amino Acids | DMEM/F-12, HEPES |
| L-Isoleucine | 54.47 | 0.42 | Amino Acids | DMEM/F-12, HEPES |
| L-Leucine | 59.05 | 0.45 | Amino Acids | DMEM/F-12, HEPES |
| L-Lysine hydrochloride | 91.25 | 0.50 | Amino Acids | DMEM/F-12, HEPES |
| L-Methionine | 17.24 | 0.12 | Amino Acids | DMEM/F-12, HEPES |
| L-Phenylalanine | 35.48 | 0.22 | Amino Acids | DMEM/F-12, HEPES |
| L-Proline | 28.75 | 0.25 | Amino Acids | DMEM/F-12, HEPES + NEAA |
| L-Serine | 36.75 | 0.35 | Amino Acids | DMEM/F-12, HEPES + NEAA |
| L-Threonine | 53.45 | 0.45 | Amino Acids | DMEM/F-12, HEPES |
| L-Tryptophan | 9.02 | 0.04 | Amino Acids | DMEM/F-12, HEPES |
| L-Tyrosine disodium salt dihydrate | 55.79 | 0.21 | Amino Acids | DMEM/F-12, HEPES |
| L-Valine | 52.85 | 0.45 | Amino Acids | DMEM/F-12, HEPES |
| Biotin | 3.50E-03 | 1.43E-05 | Vitamins | DMEM/F-12, HEPES |
| Choline chloride | 8.98 | 0.06 | Vitamins | DMEM/F-12, HEPES |
| D-Calcium pantothenate | 2.24 | 4.70E-03 | Vitamins | DMEM/F-12, HEPES |
| Folic Acid | 2.65 | 0.01 | Vitamins | DMEM/F-12, HEPES |
| Niacinamide | 2.02 | 0.02 | Vitamins | DMEM/F-12, HEPES |
| Pyridoxine hydrochloride | 2.00 | 0.01 | Vitamins | DMEM/F-12, HEPES |
| Riboflavin | 0.22 | 5.82E-04 | Vitamins | DMEM/F-12, HEPES |
| Thiamine hydrochloride | 2.17 | 0.01 | Vitamins | DMEM/F-12, HEPES |
| Vitamin B12 | 0.68 | 5.02E-04 | Vitamins | DMEM/F-12, HEPES |
| i-Inositol | 12.60 | 0.07 | Vitamins | DMEM/F-12, HEPES |
| Calcium Chloride (CaCl2) (anhyd.) | 116.60 | 1.05 | Inorganic Salts | DMEM/F-12, HEPES |
| Cupric sulfate (CuSO4-5H2O) | 1.30E-03 | 5.20E-06 | Inorganic Salts | DMEM/F-12, HEPES |
| Ferric Nitrate (Fe(NO3)3"9H2O) | 0.05 | 1.24E-04 | Inorganic Salts | DMEM/F-12, HEPES |
| Ferrous Sulfate (FeSO4 7H2O) | 0.42 | 1.50E-03 | Inorganic Salts | DMEM/F-12, HEPES |
| Magnesium Chloride (anhydrous) | 28.64 | 0.30 | Inorganic Salts | DMEM/F-12, HEPES |
| Magnesium Sulfate (MgSO4) (anhyd.) | 48.84 | 0.41 | Inorganic Salts | DMEM/F-12, HEPES |
| Potassium Chloride (KCl) | 311.80 | 4.16 | Inorganic Salts | DMEM/F-12, HEPES |
| Sodium Bicarbonate (NaHCO3) | 1200.00 | 14.29 | Inorganic Salts | DMEM/F-12, HEPES |
| Sodium Chloride (NaCl) | 6995.50 | 120.61 | Inorganic Salts | DMEM/F-12, HEPES |
| Sodium Phosphate dibasic (Na2HPO4) anhydrous | 71.02 | 0.50 | Inorganic Salts | DMEM/F-12, HEPES |
| Sodium Phosphate monobasic (NaH2PO4-H2O) | 62.50 | 0.45 | Inorganic Salts | DMEM/F-12, HEPES |
| Zinc sulfate (ZnSO4-7H2O) | 0.43 | 1.50E-03 | Inorganic Salts | DMEM/F-12, HEPES |
| D-Glucose (Dextrose) | 3151.00 | 17.51 | Other Components | DMEM/F-12, HEPES |
| HEPES | 3574.50 | 15.02 | Other Components | DMEM/F-12, HEPES |
| Hypoxanthine Na | 2.39 | 0.02 | Other Components | DMEM/F-12, HEPES |
| Linoleic Acid | 0.04 | 1.50E-04 | Other Components | DMEM/F-12, HEPES |
| Lipoic Acid | 0.11 | 5.10E-04 | Other Components | DMEM/F-12, HEPES |
| Phenol Red | 8.10 | 0.02 | Other Components | DMEM/F-12, HEPES |
| Putrescine 2HCl | 0.08 | 5.03E-04 | Other Components | DMEM/F-12, HEPES |
| Sodium Pyruvate | 55.00 | 0.50 | Other Components | DMEM/F-12, HEPES |
| Thymidine | 0.37 | 1.51E-03 | Other Components | DMEM/F-12, HEPES |
| Taurine | 12.52 | 0.10 |  | Supplement |
| B-27™ Supplement (50X), serum free | 1X |  |  | Supplement |
| Bovine Serum Albumin solution | 1% (V/V) |  |  | Supplement |

**Table S8.** **Medium formulation for human retinal explant culture.**

| **Components** | **Concentration (mg/L)** | **mM** | **Category** | **Source** |
| --- | --- | --- | --- | --- |
| Glycine | 26.25 | 0.35 | Amino Acids | Advanced DMEM/F-12 + NEAA |
| L-Alanine | 13.35 | 0.15 | Amino Acids | Advanced DMEM/F-12 + NEAA |
| L-Arginine hydrochloride | 147.50 | 0.70 | Amino Acids | Advanced DMEM/F-12 |
| L-Asparagine-H2O | 7.50 | 0.05 | Amino Acids | Advanced DMEM/F-12 |
| L-Asparagine | 19.80 | 0.15 | Amino Acids | Advanced DMEM/F-12 + NEAA |
| L-Aspartic acid | 19.95 | 0.15 | Amino Acids | Advanced DMEM/F-12 + NEAA |
| L-Cysteine hydrochloride-H2O | 17.56 | 0.10 | Amino Acids | Advanced DMEM/F-12 |
| L-Cystine 2HCl | 31.29 | 0.10 | Amino Acids | Advanced DMEM/F-12 |
| L-Glutamic Acid | 22.05 | 0.15 | Amino Acids | Advanced DMEM/F-12 + NEAA |
| L-Histidine hydrochloride-H2O | 31.48 | 0.15 | Amino Acids | Advanced DMEM/F-12 |
| L-Isoleucine | 54.47 | 0.42 | Amino Acids | Advanced DMEM/F-12 |
| L-Leucine | 59.05 | 0.45 | Amino Acids | Advanced DMEM/F-12 |
| L-Lysine hydrochloride | 91.25 | 0.50 | Amino Acids | Advanced DMEM/F-12 |
| L-Methionine | 17.24 | 0.12 | Amino Acids | Advanced DMEM/F-12 |
| L-Phenylalanine | 35.48 | 0.22 | Amino Acids | Advanced DMEM/F-12 |
| L-Proline | 28.75 | 0.25 | Amino Acids | Advanced DMEM/F-12 + NEAA |
| L-Serine | 36.75 | 0.35 | Amino Acids | Advanced DMEM/F-12 + NEAA |
| L-Threonine | 53.45 | 0.45 | Amino Acids | Advanced DMEM/F-12 |
| L-Tryptophan | 9.02 | 0.04 | Amino Acids | Advanced DMEM/F-12 |
| L-Tyrosine disodium salt dihydrate | 55.79 | 0.21 | Amino Acids | Advanced DMEM/F-12 |
| L-Valine | 52.85 | 0.45 | Amino Acids | Advanced DMEM/F-12 |
| Ascorbic Acid phosphate | 2.50 | 0.01 | Vitamins | Advanced DMEM/F-12 |
| Biotin | 3.50E-03 | 1.43E-05 | Vitamins | Advanced DMEM/F-12 |
| Choline chloride | 8.98 | 0.06 | Vitamins | Advanced DMEM/F-12 |
| D-Calcium pantothenate | 2.24 | 4.70E-03 | Vitamins | Advanced DMEM/F-12 |
| Folic Acid | 2.65 | 0.01 | Vitamins | Advanced DMEM/F-12 |
| Niacinamide | 2.02 | 0.02 | Vitamins | Advanced DMEM/F-12 |
| Pyridoxine hydrochloride | 2.00 | 0.01 | Vitamins | Advanced DMEM/F-12 |
| Riboflavin | 0.22 | 5.82E-04 | Vitamins | Advanced DMEM/F-12 |
| Thiamine hydrochloride | 2.17 | 0.01 | Vitamins | Advanced DMEM/F-12 |
| Vitamin B12 | 0.68 | 5.02E-04 | Vitamins | Advanced DMEM/F-12 |
| i-Inositol | 12.60 | 0.07 | Vitamins | Advanced DMEM/F-12 |
| Calcium Chloride (CaCl2) (anhyd.) | 116.60 | 1.05 | Inorganic Salts | Advanced DMEM/F-12 |
| Cupric sulfate (CuSO4-5H2O) | 1.30E-03 | 5.20E-06 | Inorganic Salts | Advanced DMEM/F-12 |
| Ferric Nitrate (Fe(NO3)3"9H2O) | 0.05 | 1.24E-04 | Inorganic Salts | Advanced DMEM/F-12 |
| Ferric sulfate (FeSO4-7H2O) | 0.42 | 1.50E-03 | Inorganic Salts | Advanced DMEM/F-12 |
| Magnesium Chloride (anhydrous) | 28.64 | 0.30 | Inorganic Salts | Advanced DMEM/F-12 |
| Magnesium Sulfate (MgSO4) (anhyd.) | 48.84 | 0.41 | Inorganic Salts | Advanced DMEM/F-12 |
| Potassium Chloride (KCl) | 311.80 | 4.16 | Inorganic Salts | Advanced DMEM/F-12 |
| Sodium Bicarbonate (NaHCO3) | 2438.00 | 29.02 | Inorganic Salts | Advanced DMEM/F-12 |
| Sodium Chloride (NaCl) | 6995.50 | 120.61 | Inorganic Salts | Advanced DMEM/F-12 |
| Sodium Phosphate dibasic (Na2HPO4) anhydrous | 71.02 | 0.50 | Inorganic Salts | Advanced DMEM/F-12 |
| Sodium Phosphate monobasic (NaH2PO4-H2O) | 62.50 | 0.45 | Inorganic Salts | Advanced DMEM/F-12 |
| Zinc sulfate (ZnSO4-7H2O) | 0.86 | 3.00E-03 | Inorganic Salts | Advanced DMEM/F-12 |
| AlbuMAX® II | 400.00 | Infinity | Proteins | Advanced DMEM/F-12 |
| Human Transferrin (Holo) | 7.50 | Infinity | Proteins | Advanced DMEM/F-12 |
| Insulin Recombinant Full Chain | 10.00 | Infinity | Proteins | Advanced DMEM/F-12 |
| Glutathione, monosodium | 1.00 | 3.26E-03 | Reducing Agents | Advanced DMEM/F-12 |
| Ammonium Metavanadate | 3.00E-04 | 2.56E-06 | Trace Elements | Advanced DMEM/F-12 |
| Manganous Chloride | 5.00E-05 | 2.53E-07 | Trace Elements | Advanced DMEM/F-12 |
| Sodium Selenite | 0.01 | 2.89E-05 | Trace Elements | Advanced DMEM/F-12 |
| D-Glucose (Dextrose) | 3151.00 | 17.51 | Other Components | Advanced DMEM/F-12 |
| Ethanolamine | 1.90 | 0.02 | Other Components | Advanced DMEM/F-12 |
| Hypoxanthine Na | 2.39 | 0.02 | Other Components | Advanced DMEM/F-12 |
| Linoleic Acid | 0.04 | 1.50E-04 | Other Components | Advanced DMEM/F-12 |
| Lipoic Acid | 0.11 | 5.10E-04 | Other Components | Advanced DMEM/F-12 |
| Phenol Red | 8.10 | 0.02 | Other Components | Advanced DMEM/F-12 |
| Putrescine 2HCl | 0.08 | 5.03E-04 | Other Components | Advanced DMEM/F-12 |
| Sodium Pyruvate | 110.00 | 1.00 | Other Components | Advanced DMEM/F-12 |
| Thymidine | 0.37 | 1.51E-03 | Other Components | Advanced DMEM/F-12 |
| Taurine | 12.52 | 0.10 |  | Supplement |
| B-27™ Supplement (50X), serum free | 1X |  |  | Supplement |
| Bovine Serum Albumin solution | 1% (V/V) |  |  | Supplement |
| CTS™ (Cell Therapy Systems) N-2 Supplement | 1X |  |  | Supplement |
| L-Glutamine | 292.28 | 2.00 |  | Supplement |

**Table S9. Differentially expressed metabolic genes in macular and peripheral retinas.**

| **Gene symbol** | **FC** | **P value** | **Classification (Sub-category/substrate)** | **In figures** |
| --- | --- | --- | --- | --- |
| SV2C | 2.63 | 1.53E-02 | Small Molecule Transport (Sugar) | Fig 6b |
| SLC2A6 | 2.44 | 1.12E-03 | Small Molecule Transport (Glucose) | Fig 6b |
| SLC2A13 | 1.97 | 3.08E-03 | Small Molecule Transport (Myo-Inoisitol) | Fig 6b |
| SLC45A1 | 1.81 | 1.83E-02 | Small Molecule Transport (Glucose) | Fig 6b |
| SLC35F3 | 1.80 | 4.45E-02 | Small Molecule Transport (Nucleotide sugar) | Fig 6b |
| SLC35C1 | 1.51 | 1.04E-02 | Small Molecule Transport (GDP-Fructose) | Fig 6b |
| SV2A | 1.47 | 3.68E-03 | Small Molecule Transport (Sugar) | Fig 6b |
| SLC35D1 | 0.71 | 1.36E-04 | Small Molecule Transport (UDP-N-Acetyl-Glucosomine) | Fig 6b |
| SLC2A11 | 0.71 | 1.93E-02 | Small Molecule Transport (Glucose, Fructose) | Fig 6b |
| SLC2A10 | 0.69 | 3.23E-02 | Small Molecule Transport (Glucose) | Fig 6b |
| SLC35F2 | 0.68 | 7.92E-03 | Small Molecule Transport (Nucleotide sugar) | Fig 6b |
| SLC35F4 | 0.67 | 1.50E-02 | Small Molecule Transport (Nucleotide sugar) | Fig 6b |
| SLC35D2 | 0.67 | 2.65E-02 | Small Molecule Transport (Nucleotide sugar) | Fig 6b |
| SLC17A6 | 3.68 | 1.65E-03 | Small Molecule Transport (Glutamate) | Fig 6b |
| SLC1A1 | 1.40 | 3.60E-03 | Small Molecule Transport (Glutamate, Asparate) | Fig 6b |
| SLC25A18 | 1.38 | 4.84E-02 | Small Molecule Transport (Glutamate) | Fig 6b |
| SLC25A12 | 1.36 | 1.71E-03 | Small Molecule Transport (Glutamate, Aspartate) | Fig 6b |
| SLC38A3 | 0.75 | 1.70E-02 | Small Molecule Transport (His, Gln, Asn, Ala, Glutamate) | Fig 6b |
| SLC1A6 | 0.75 | 4.85E-02 | Small Molecule Transport (Glutamate, Asparate) | Fig 6b |
| SLC25A29 | 0.69 | 8.43E-03 | Small Molecule Transport (Acylcarnitine) | Fig 6b |
| SLC27A1 | 0.70 | 2.71E-03 | Small Molecule Transport (Fatty acid) | Fig 6b |
| SLC27A2 | 0.65 | 1.22E-02 | Small Molecule Transport (Fatty acid) | Fig 6b |
| SLC27A3 | 0.65 | 2.55E-02 | Small Molecule Transport (Fatty acid) | Fig 6b |
| SLC16A3 | 2.21 | 7.41E-03 | Small Molecule Transport (Lactate/Pyruvate) | Fig 6b |
| SLC16A7 | 1.85 | 6.31E-03 | Small Molecule Transport (Lactate/Pyruvate) | Fig 6b |
| SLC25A1 | 1.52 | 7.42E-04 | Small Molecule Transport (Citrate, Malate) | Fig 6b |
| SLC13A5 | 0.73 | 2.71E-02 | Small Molecule Transport (Citrate, Pyruvate) | Fig 6b |
| SLC25A23 | 0.76 | 9.61E-04 | Small Molecule Transport (Mitochondrial Phosphate) | Fig 6b |
| SLC34A1 | 0.66 | 3.72E-03 | Small Molecule Transport (Mitochondrial Phosphate) | Fig 6b |
| SLC34A3 | 0.36 | 2.27E-02 | Small Molecule Transport (Mitochondrial Phosphate) | Fig 6b |
| AQP11 | 0.73 | 2.35E-02 | Small Molecule Transport (Water, Glycerol) | Fig 6b |
| AQP3 | 0.58 | 3.61E-04 | Small Molecule Transport (Water) | Fig 6b |
| AQP9 | 0.45 | 2.08E-02 | Small Molecule Transport (Water, Glycerol) | Fig 6b |
| SLCO2A1 | 2.55 | 2.66E-04 | Small Molecule Transport (eicosanoids, prostaglandins) | Fig 6b |
| SLCO3A1 | 2.44 | 1.53E-03 | Small Molecule Transport (vasopressin, prostaglandins, thyroid hormones) | Fig 6b |
| SLC10A5 | 1.81 | 3.77E-02 | Small Molecule Transport (Bile acid) | Fig 6b |
| STARD13 | 1.58 | 2.17E-03 | Small Molecule Transport (Lipid) | Fig 6b |
| STARD4 | 1.50 | 6.04E-03 | Small Molecule Transport (Cholesterol) | Fig 6b |
| SLC10A3 | 1.31 | 6.31E-03 | Small Molecule Transport (Bile acid) | Fig 6b |
| SLCO4A1 | 0.59 | 1.38E-03 | Small Molecule Transport (bile acids, prostaglandins, steroid conjugates, thyroid hormone) | Fig 6b |
| SLC22A11 | 3.66 | 2.54E-02 | Small Molecule Transport (Uric acid ) | Fig 6b |
| SLC6A17 | 2.87 | 1.53E-03 | Small Molecule Transport (Pro, Cys, Ala) | Fig 6b |
| SLC26A8 | 2.44 | 4.23E-03 | Small Molecule Transport (Biocarbonate) | Fig 6b |
| SLC25A43 | 2.26 | 8.34E-03 | Small Molecule Transport (Nucleotide) | Fig 6b |
| RHCG | 2.18 | 3.75E-02 | Small Molecule Transport (Ammonium) | Fig 6b |
| SLC22A7 | 2.02 | 2.37E-02 | Small Molecule Transport (Anion) | Fig 6b |
| SLC5A8 | 2.04 | 1.67E-02 | Small Molecule Transport (iodide) | Fig 6b |
| SLC4A3 | 1.89 | 2.45E-03 | Small Molecule Transport (Anion) | Fig 6b |
| SLC22A18 | 1.75 | 4.94E-02 | Small Molecule Transport (Organic Cation) | Fig 6b |
| SLC6A15 | 1.63 | 5.35E-03 | Small Molecule Transport (Met, BCAA) | Fig 6b |
| SLC7A1 | 1.52 | 4.68E-03 | Small Molecule Transport (Ar, Lys, Orn, His) | Fig 6b |
| SLC19A3 | 1.45 | 2.44E-03 | Small Molecule Transport (Thiamine) | Fig 6b |
| SLC7A14 | 1.35 | 2.61E-02 | Small Molecule Transport (Amino acid) | Fig 6b |
| SLC4A4 | 1.35 | 1.58E-02 | Small Molecule Transport (Biocarbonate) | Fig 6b |
| SLC25A46 | 1.33 | 1.03E-02 | Small Molecule Transport (mitochondrial transport) | Fig 6b |
| SLC15A2 | 0.71 | 1.53E-02 | Small Molecule Transport (peptide) | Fig 6b |
| SLC25A35 | 0.71 | 6.06E-03 | Small Molecule Transport (Di-Tri-carboxylic acid) | Fig 6b |
| SLC26A11 | 0.68 | 1.34E-03 | Small Molecule Transport (Sulfate) | Fig 6b |
| SLC6A9 | 0.62 | 1.07E-03 | Small Molecule Transport (Glycine, Sarcosine) | Fig 6b |
| SLC25A45 | 0.59 | 4.01E-02 | Small Molecule Transport (Amino acid) | Fig 6b |
| SLC6A11 | 0.59 | 3.93E-02 | Small Molecule Transport (GABA, β-Alanine) | Fig 6b |
| SLC15A3 | 0.54 | 1.17E-02 | Small Molecule Transport (peptide) | Fig 6b |
| SLC44A5 | 0.53 | 1.03E-02 | Small Molecule Transport (Choline) | Fig 6b |
| FOLR1 | 0.48 | 1.08E-02 | Small Molecule Transport (Folate) | Fig 6b |
| SLC18A1 | 0.10 | 2.87E-02 | Small Molecule Transport (Monoamine) | Fig 6b |
| HEXA | 0.76 | 8.99E-03 | Glycan (Degradation) | Fig 6c |
| MAN2C1 | 0.72 | 5.79E-03 | Glycan (Degradation) | Fig 6c |
| GLB1L | 0.71 | 4.58E-02 | Glycan (Degradation) | Fig 6c |
| GUSB | 0.68 | 2.54E-02 | Glycan (Degradation) | Fig 6c |
| GLB1L2 | 0.67 | 3.46E-02 | Glycan (Degradation) | Fig 6c |
| IDUA | 0.65 | 1.16E-02 | Glycan (Degradation) | Fig 6c |
| HYAL1 | 0.56 | 2.98E-02 | Glycan (Degradation) | Fig 6c |
| CHST8 | 1.98 | 2.79E-02 | Glycan (Sulfate) | Fig 6c |
| SULF2 | 1.67 | 2.52E-03 | Glycan (Sulfate) | Fig 6c |
| CHST14 | 0.67 | 3.63E-02 | Glycan (Sulfate) | Fig 6c |
| GAL3ST3 | 0.66 | 3.07E-04 | Glycan (Sulfate) | Fig 6c |
| SGSH | 0.63 | 5.56E-05 | Glycan (Sulfate) | Fig 6c |
| CHST3 | 0.61 | 5.92E-03 | Glycan (Sulfate) | Fig 6c |
| ABO | 4.48 | 8.89E-04 | Glycan | Fig 6c |
| GCNT4 | 3.57 | 9.41E-04 | Glycan | Fig 6c |
| B3GNT8 | 3.26 | 8.37E-03 | Glycan | Fig 6c |
| ST6GAL1 | 3.26 | 5.76E-04 | Glycan | Fig 6c |
| FUT1 | 2.49 | 1.14E-03 | Glycan | Fig 6c |
| ST8SIA4 | 2.41 | 1.02E-03 | Glycan | Fig 6c |
| B4GALNT1 | 2.35 | 2.28E-03 | Glycan | Fig 6c |
| GALNT9 | 2.19 | 1.21E-03 | Glycan | Fig 6c |
| B3GNT4 | 2.16 | 1.84E-02 | Glycan | Fig 6c |
| B4GALT5 | 2.04 | 2.67E-03 | Glycan | Fig 6c |
| ST3GAL6 | 1.98 | 4.99E-04 | Glycan | Fig 6c |
| ST8SIA2 | 1.86 | 3.93E-02 | Glycan | Fig 6c |
| ST8SIA1 | 1.77 | 4.17E-03 | Glycan | Fig 6c |
| MGAT5 | 1.68 | 4.57E-03 | Glycan | Fig 6c |
| GLT1D1 | 1.68 | 2.52E-03 | Glycan | Fig 6c |
| WBSCR17 | 1.64 | 9.36E-03 | Glycan | Fig 6c |
| FUT9 | 1.59 | 9.79E-03 | Glycan | Fig 6c |
| ST6GAL2 | 1.47 | 4.07E-02 | Glycan | Fig 6c |
| MGAT3 | 1.46 | 3.03E-02 | Glycan | Fig 6c |
| GBGT1 | 1.47 | 1.70E-02 | Glycan | Fig 6c |
| MGAT4C | 1.46 | 4.53E-02 | Glycan | Fig 6c |
| ST8SIA3 | 1.39 | 7.10E-03 | Glycan | Fig 6c |
| B4GALT6 | 1.38 | 6.31E-03 | Glycan | Fig 6c |
| GLT25D2 | 1.34 | 1.93E-02 | Glycan | Fig 6c |
| DHDDS | 0.77 | 2.55E-02 | Glycan | Fig 6c |
| GALNT4 | 0.76 | 7.24E-03 | Glycan | Fig 6c |
| GALNT7 | 0.76 | 7.59E-03 | Glycan | Fig 6c |
| B4GALT1 | 0.74 | 2.05E-02 | Glycan | Fig 6c |
| FUT4 | 0.70 | 1.45E-02 | Glycan | Fig 6c |
| CTBS | 0.70 | 2.85E-02 | Glycan | Fig 6c |
| POFUT2 | 0.70 | 8.54E-03 | Glycan | Fig 6c |
| ST6GALNAC5 | 0.68 | 7.64E-04 | Glycan | Fig 6c |
| ST3GAL1 | 0.67 | 2.12E-02 | Glycan | Fig 6c |
| GALNT10 | 0.60 | 2.73E-02 | Glycan | Fig 6c |
| ST6GALNAC2 | 0.59 | 1.77E-04 | Glycan | Fig 6c |
| GYLTL1B | 0.53 | 1.23E-02 | Glycan | Fig 6c |
| FUT7 | 0.30 | 1.63E-02 | Glycan | Fig 6c |
| PYGM | 2.94 | 1.69E-04 | Sugar | Fig 6d |
| PYGB | 1.57 | 1.22E-03 | Sugar | Fig 6d |
| GBE1 | 1.49 | 2.48E-03 | Sugar | Fig 6d |
| UGP2 | 1.42 | 2.04E-05 | Sugar | Fig 6d |
| GYS1 | 0.77 | 4.76E-02 | Sugar | Fig 6d |
| AGL | 0.75 | 3.86E-02 | Sugar | Fig 6d |
| GYG1 | 0.75 | 3.71E-02 | Sugar | Fig 6d |
| PC | 1.59 | 2.97E-04 | TCA Cycle | Fig 6d |
| LDHD | 1.37 | 4.80E-02 | Glycolysis | Fig 6d |
| LDHB | 1.34 | 1.54E-04 | Glycolysis | Fig 6d |
| ME2 | 0.75 | 1.14E-02 | TCA Cycle | Fig 6d |
| GPT2 | 0.71 | 2.76E-03 | Amino Acid (Pyruvate) | Fig 6d |
| GOT1 | 1.70 | 1.92E-05 | TCA Cycle | Fig 6d |
| MDH1 | 1.55 | 4.98E-06 | TCA Cycle | Fig 6d |
| ACLY | 1.48 | 1.81E-05 | TCA Cycle | Fig 6d |
| OGDH | 1.32 | 3.68E-03 | TCA Cycle | Fig 6d |
| ACO1 | 1.31 | 2.75E-04 | TCA Cycle | Fig 6d |
| IDH2 | 0.74 | 1.35E-03 | TCA Cycle | Fig 6d |
| PGAM4 | 2.40 | 3.47E-03 | Glycolysis | Fig 6d |
| PGM2L1 | 1.54 | 2.58E-02 | Glycolysis | Fig 6d |
| GPI | 1.43 | 8.97E-03 | Glycolysis | Fig 6d |
| PGK1 | 1.33 | 4.93E-02 | Glycolysis | Fig 6d |
| PGAM2 | 0.73 | 2.35E-02 | Glycolysis | Fig 6d |
| PGM5 | 0.72 | 9.31E-04 | Glycolysis | Fig 6d |
| PFKFB2 | 0.65 | 6.13E-03 | Glycolysis | Fig 6d |
| GLYAT | 4.15 | 2.05E-02 | Amino Acid (Gly) | Fig 6e |
| GLYATL2 | 2.47 | 8.19E-05 | Amino Acid (Gly) | Fig 6e |
| GLYATL1 | 2.23 | 1.83E-04 | Amino Acid (Gly) | Fig 6e |
| PAM | 1.41 | 1.52E-02 | Amino Acid (Gly) | Fig 6e |
| PSAT | 0.69 | 1.15E-02 | Amino Acid (Ser) | Fig 6e |
| CHDH | 0.66 | 1.83E-02 | Amino Acid (Gly) | Fig 6e |
| PHGDH | 0.60 | 1.12E-02 | Amino Acid (Ser) | Fig 6e |
| CPS1 | 2.88 | 1.07E-03 | Amino Acid (Urea) | Fig 6e |
| NAGS | 2.36 | 8.48E-03 | Amino Acid (Arg) | Fig 6e |
| DDAH1 | 1.85 | 8.04E-04 | Amino Acid (Arg, Citn) | Fig 6e |
| DDAH2 | 0.59 | 4.08E-03 | Amino Acid (Arg, Citn) | Fig 6e |
| NOS1 | 0.55 | 1.34E-02 | Amino Acid (Arg) | Fig 6e |
| AOX1 | 0.44 | 3.06E-03 | Amino Acid (Arg) | Fig 6e |
| GAD2 | 0.74 | 4.28E-02 | Amino Acid (Glu) | Fig 6e |
| GPT2 | 0.71 | 2.76E-03 | Amino Acid (Glu, Ala) | Fig 6e |
| GLUL | 0.68 | 3.16E-02 | Amino Acid (Glu) | Fig 6e |
| HAGHL | 1.73 | 4.90E-02 | Amino Acid (GSH) | Fig 6e |
| MGST2 | 1.52 | 2.40E-02 | Amino Acid (GSH) | Fig 6e |
| MGST3 | 1.51 | 6.03E-04 | Amino Acid (GSH) | Fig 6e |
| GSTO1 | 1.44 | 1.48E-03 | Amino Acid (GSH) | Fig 6e |
| GPX3 | 0.43 | 4.52E-03 | Amino Acid (GSH) | Fig 6e |
| GPX8 | 0.43 | 4.09E-03 | Amino Acid (GSH) | Fig 6e |
| GPX5 | 0.06 | 3.14E-02 | Amino Acid (GSH) | Fig 6e |
| NAT1 | 2.46 | 3.50E-04 | Amino Acid (Acetylation) | Fig 6e |
| NAT8L | 1.97 | 4.70E-02 | Amino Acid (Acetylation) | Fig 6e |
| ALDH1L2 | 1.87 | 9.25E-03 | Amino Acid (One carbon) | Fig 6e |
| BBOX1 | 1.83 | 2.91E-02 | Amino Acid (Cofactor) | Fig 6e |
| BCAT1 | 1.60 | 6.26E-04 | Amino Acid (BCAAs) | Fig 6e |
| ACYP2 | 1.58 | 1.19E-06 | Amino Acid (Pyr) | Fig 6e |
| PCBD2 | 0.77 | 1.60E-02 | Amino Acid (BH4) | Fig 6e |
| ADI1 | 0.76 | 8.76E-03 | Amino Acid (Met) | Fig 6e |
| GATM | 0.75 | 1.93E-02 | Amino Acid (Cp) | Fig 6e |
| AGXT2L2 | 0.73 | 4.10E-03 | Amino Acid (Ala) | Fig 6e |
| AADAT | 0.73 | 3.34E-02 | Amino Acid (Kyneurenine, Glu) | Fig 6e |
| PIPOX | 0.72 | 1.05E-03 | Amino Acid (Lys) | Fig 6e |
| MAOB | 0.62 | 2.42E-02 | Amino Acid (Neurotransmiter) | Fig 6e |
| DDO | 0.52 | 5.53E-03 | Amino Acid (Asp) | Fig 6e |
| PRODH | 0.48 | 1.93E-02 | Amino Acid (Pro) | Fig 6e |
| HDC | 0.46 | 3.36E-02 | Amino Acid (His) | Fig 6e |
| IDO1 | 0.41 | 1.73E-02 | Amino Acid (Try, NAD) | Fig 6e |
| PNLIPRP3 | 4.84 | 2.96E-02 | Lipid (Fatty Acid) | Fig 6f |
| LIPJ | 2.87 | 3.52E-02 | Lipid (Fatty Acid) | Fig 6f |
| SCD | 2.92 | 2.13E-05 | Lipid (Fatty Acid) | Fig 6f |
| CYP4A11 | 2.60 | 1.95E-02 | Lipid (Fatty Acid) | Fig 6f |
| FAR2 | 2.55 | 5.74E-04 | Lipid (Fatty Acid) | Fig 6f |
| ACAT2 | 2.32 | 5.60E-04 | Lipid (Fatty Acid) | Fig 6f |
| ELOVL6 | 2.03 | 3.51E-05 | Lipid (Fatty Acid) | Fig 6f |
| CYP4X1 | 1.88 | 2.32E-04 | Lipid (Fatty Acid) | Fig 6f |
| FASN | 1.87 | 2.50E-03 | Lipid (Fatty Acid) | Fig 6f |
| ACSBG1 | 1.78 | 2.21E-03 | Lipid (Fatty Acid) | Fig 6f |
| ACACA | 1.61 | 3.14E-05 | Lipid (Fatty Acid) | Fig 6f |
| ABHD6 | 1.57 | 1.54E-03 | Lipid (Fatty Acid) | Fig 6f |
| MGLL | 1.54 | 3.32E-02 | Lipid (Fatty Acid) | Fig 6f |
| DDHD2 | 1.51 | 1.16E-03 | Lipid (Fatty Acid) | Fig 6f |
| ACSM4 | 1.49 | 4.64E-02 | Lipid (Fatty Acid) | Fig 6f |
| LPL | 1.41 | 3.31E-02 | Lipid (Fatty Acid) | Fig 6f |
| ACSS2 | 1.35 | 5.02E-03 | Lipid (Fatty Acid) | Fig 6f |
| ACBD4 | 0.77 | 2.49E-03 | Lipid (Fatty Acid) | Fig 6f |
| MECR | 0.77 | 4.82E-02 | Lipid (Fatty Acid) | Fig 6f |
| CPT1A | 0.76 | 1.08E-02 | Lipid (Fatty Acid) | Fig 6f |
| ELOVL1 | 0.75 | 3.56E-03 | Lipid (Fatty Acid) | Fig 6f |
| ACADS | 0.71 | 1.45E-02 | Lipid (Fatty Acid) | Fig 6f |
| AMACR | 0.70 | 1.13E-02 | Lipid (Fatty acid) | Fig 6f |
| ABHD1 | 0.69 | 1.63E-02 | Lipid (Fatty Acid) | Fig 6f |
| ACAA2 | 0.69 | 1.61E-02 | Lipid (Fatty Acid) | Fig 6f |
| ECHDC3 | 0.63 | 4.48E-02 | Lipid (Fatty Acid) | Fig 6f |
| ABHD14B | 0.61 | 2.37E-03 | Lipid (Fatty Acid) | Fig 6f |
| ACOX2 | 0.61 | 1.47E-02 | Lipid (Fatty Acid) | Fig 6f |
| HMGCR | 1.70 | 2.57E-04 | Lipid (Mevalonate) | Fig 6f |
| IDI1 | 1.50 | 6.89E-04 | Lipid (Mevalonate) | Fig 6f |
| HMGCS1 | 1.43 | 5.03E-03 | Lipid (Mevalonate) | Fig 6f |
| DHCR24 | 2.57 | 1.49E-04 | Lipid (Cholesterol) | Fig 6f |
| LSS | 2.47 | 9.45E-05 | Lipid (Cholesterol) | Fig 6f |
| SQLE | 1.63 | 5.82E-04 | Lipid (Cholesterol) | Fig 6f |
| FDPS | 1.51 | 1.42E-03 | Lipid (Cholesterol) | Fig 6f |
| DHCR7 | 1.49 | 7.39E-03 | Lipid (Cholesterol) | Fig 6f |
| SC5DL | 1.48 | 1.05E-03 | Lipid (Cholesterol) | Fig 6f |
| FDFT1 | 1.36 | 1.37E-02 | Lipid (Cholesterol) | Fig 6f |
| CYP51A1 | 1.30 | 1.37E-03 | Lipid (Cholesterol) | Fig 6f |
| OXCT2 | 1.98 | 1.59E-03 | Ketone Body (Ketone Body) | Fig 6f |
| HMGCLL1 | 1.64 | 1.20E-03 | Ketone Body (Ketone Body) | Fig 6f |
| AACS | 1.56 | 2.12E-03 | Ketone Body (Ketone Body) | Fig 6f |
| QPRT | 2.53 | 4.96E-04 | NAD (NAD) | Fig 6f |
| NMNAT2 | 2.34 | 8.18E-04 | NAD (NAD) | Fig 6f |
| NQO2 | 1.63 | 3.34E-03 | NAD (NAD) | Fig 6f |
| ABCC6 | 2.15 | 1.59E-02 | ABC Transporter |  |
| ABCB4 | 2.04 | 2.92E-02 | ABC Transporter |  |
| ABCA3 | 1.41 | 7.81E-03 | ABC Transporter |  |
| ABCG4 | 0.72 | 1.00E-02 | ABC Transporter |  |
| ABCG5 | 0.66 | 1.52E-02 | ABC Transporter |  |
| ABCG1 | 0.63 | 2.15E-02 | ABC Transporter |  |
| ABCA1 | 0.45 | 9.69E-03 | ABC Transporter |  |
| ACOT7 | 2.09 | 4.10E-04 | CoA |  |
| ARSF | 4.97 | 2.70E-03 | Cofactor (Calcium) |  |
| ALDH1B1 | 1.72 | 4.00E-03 | Detox |  |
| FMO4 | 1.61 | 4.09E-03 | Detox |  |
| ALDH7A1 | 0.76 | 1.14E-02 | Detox |  |
| EPHX2 | 0.67 | 3.91E-02 | Detox |  |
| SDHA | 1.47 | 5.99E-06 | ETC (Complex II) |  |
| COX6A2 | 1.85 | 4.63E-04 | ETC (Complex IV) |  |
| GLOD5 | 0.51 | 1.17E-02 | Glyoxylate |  |
| HS6ST3 | 2.61 | 9.44E-04 | Heparin Sulfate |  |
| HS3ST2 | 1.82 | 3.79E-03 | Heparin Sulfate |  |
| HS3ST1 | 1.71 | 3.15E-02 | Heparin Sulfate |  |
| HS3ST3B1 | 0.72 | 1.41E-02 | Heparin Sulfate |  |
| CLIC3 | 12.88 | 2.53E-02 | Ion Transport |  |
| KCNE1 | 6.04 | 9.58E-03 | Ion Transport |  |
| CATSPERB | 5.01 | 1.32E-02 | Ion Transport |  |
| P2RX2 | 3.98 | 3.56E-02 | Ion Transport |  |
| KCNC2 | 3.84 | 1.46E-03 | Ion Transport |  |
| CLCA4 | 3.72 | 3.06E-02 | Ion Transport |  |
| TRPM2 | 3.47 | 3.60E-04 | Ion Transport |  |
| CHRNA6 | 3.23 | 7.56E-05 | Ion Transport |  |
| CHRNB3 | 3.07 | 2.03E-04 | Ion Transport |  |
| TRPV2 | 2.69 | 5.50E-04 | Ion Transport |  |
| SCN1A | 2.63 | 3.05E-04 | Ion Transport |  |
| SCN4B | 2.56 | 4.68E-03 | Ion Transport |  |
| KCNN2 | 2.50 | 1.04E-04 | Ion Transport |  |
| KCTD17 | 2.49 | 1.36E-03 | Ion Transport |  |
| SLN | 2.46 | 4.73E-02 | Ion Transport |  |
| P2RX6 | 2.41 | 2.35E-05 | Ion Transport |  |
| TRPA1 | 2.37 | 3.30E-02 | Ion Transport |  |
| KCNA2 | 2.36 | 1.85E-03 | Ion Transport |  |
| ATP1B1 | 2.32 | 3.15E-04 | Ion Transport |  |
| SLC39A8 | 2.28 | 2.03E-04 | Ion Transport |  |
| SCN2B | 2.22 | 9.62E-04 | Ion Transport |  |
| GRIN2A | 2.12 | 2.03E-02 | Ion Transport |  |
| KCTD19 | 2.12 | 8.31E-04 | Ion Transport |  |
| ATP2B3 | 2.11 | 1.01E-03 | Ion Transport |  |
| GABRA4 | 2.04 | 2.18E-02 | Ion Transport |  |
| GRIN1 | 2.02 | 1.45E-02 | Ion Transport |  |
| KCNG1 | 1.97 | 2.92E-02 | Ion Transport |  |
| SCN8A | 1.94 | 5.10E-04 | Ion Transport |  |
| KCNC1 | 1.90 | 4.92E-03 | Ion Transport |  |
| GRIN2B | 1.89 | 5.78E-03 | Ion Transport |  |
| HCN2 | 1.88 | 2.30E-03 | Ion Transport |  |
| GLRA1 | 1.87 | 1.96E-03 | Ion Transport |  |
| KCNQ2 | 1.83 | 3.23E-02 | Ion Transport |  |
| ATP2B2 | 1.81 | 2.14E-03 | Ion Transport |  |
| KCNK4 | 1.78 | 1.10E-02 | Ion Transport |  |
| CACNG2 | 1.78 | 1.17E-02 | Ion Transport |  |
| KCNQ5 | 1.73 | 2.73E-02 | Ion Transport |  |
| KCNJ16 | 1.71 | 6.72E-03 | Ion Transport |  |
| KCNJ11 | 1.70 | 6.30E-03 | Ion Transport |  |
| KCNT1 | 1.69 | 3.62E-02 | Ion Transport |  |
| GLRB | 1.68 | 5.40E-04 | Ion Transport |  |
| KCNJ4 | 1.67 | 2.11E-02 | Ion Transport |  |
| KCNN1 | 1.67 | 2.43E-02 | Ion Transport |  |
| RYR2 | 1.66 | 7.72E-04 | Ion Transport |  |
| SLC9A1 | 1.66 | 4.67E-03 | Ion Transport |  |
| KCTD21 | 1.64 | 5.91E-03 | Ion Transport |  |
| KCNJ3 | 1.62 | 4.20E-02 | Ion Transport |  |
| KCNJ8 | 1.62 | 3.31E-02 | Ion Transport |  |
| TRPC3 | 1.60 | 1.29E-02 | Ion Transport |  |
| KCNE2 | 1.60 | 1.63E-02 | Ion Transport |  |
| KCNQ3 | 1.59 | 2.46E-02 | Ion Transport |  |
| VDAC1 | 1.59 | 5.50E-05 | Ion Transport |  |
| KCNJ12 | 1.58 | 2.90E-02 | Ion Transport |  |
| GABRB2 | 1.53 | 4.54E-03 | Ion Transport |  |
| CLCN6 | 1.52 | 3.14E-04 | Ion Transport |  |
| SCN2A | 1.51 | 2.97E-04 | Ion Transport |  |
| KCNH7 | 1.51 | 5.54E-03 | Ion Transport |  |
| CACNA1B | 1.51 | 1.26E-02 | Ion Transport |  |
| KCNAB2 | 1.50 | 3.66E-02 | Ion Transport |  |
| KCNN4 | 1.48 | 1.13E-03 | Ion Transport |  |
| SLC24A3 | 1.47 | 1.69E-02 | Ion Transport |  |
| SLC24A2 | 1.46 | 1.25E-02 | Ion Transport |  |
| SLC39A11 | 1.46 | 2.45E-02 | Ion Transport |  |
| TRPC7 | 1.44 | 4.25E-03 | Ion Transport |  |
| CACNB3 | 1.44 | 7.99E-03 | Ion Transport |  |
| GLRA2 | 1.43 | 4.25E-02 | Ion Transport |  |
| KCNG4 | 1.42 | 2.33E-02 | Ion Transport |  |
| SLC41A3 | 1.42 | 3.42E-03 | Ion Transport |  |
| CACNA2D3 | 1.41 | 5.60E-03 | Ion Transport |  |
| VDAC3 | 1.39 | 1.26E-03 | Ion Transport |  |
| GABRG2 | 1.38 | 6.67E-03 | Ion Transport |  |
| CHRNB2 | 1.38 | 1.92E-02 | Ion Transport |  |
| KCTD20 | 1.38 | 7.11E-04 | Ion Transport |  |
| GABRR3 | 1.37 | 4.88E-02 | Ion Transport |  |
| KCND3 | 1.36 | 1.28E-02 | Ion Transport |  |
| KCNAB3 | 1.36 | 4.01E-03 | Ion Transport |  |
| KCNK3 | 1.36 | 4.65E-03 | Ion Transport |  |
| KCNJ2 | 1.35 | 1.40E-02 | Ion Transport |  |
| SLC39A10 | 1.34 | 2.48E-02 | Ion Transport |  |
| GABRA1 | 1.32 | 2.42E-02 | Ion Transport |  |
| SLC9A6 | 1.32 | 3.10E-04 | Ion Transport |  |
| GRIA3 | 1.32 | 3.77E-02 | Ion Transport |  |
| CATSPER3 | 0.77 | 1.60E-02 | Ion Transport |  |
| RYR1 | 0.76 | 4.79E-02 | Ion Transport |  |
| ATP1B2 | 0.75 | 1.10E-03 | Ion Transport |  |
| HEPH | 0.74 | 4.54E-02 | Ion Transport |  |
| CACNA1D | 0.74 | 3.31E-02 | Ion Transport |  |
| CLIC4 | 0.73 | 2.10E-02 | Ion Transport |  |
| ITPR2 | 0.72 | 3.53E-02 | Ion Transport |  |
| CNGA1 | 0.71 | 4.88E-02 | Ion Transport |  |
| TTYH2 | 0.70 | 1.80E-02 | Ion Transport |  |
| GRIA1 | 0.69 | 2.74E-02 | Ion Transport |  |
| GRIN3A | 0.68 | 4.01E-02 | Ion Transport |  |
| TTYH1 | 0.66 | 6.47E-04 | Ion Transport |  |
| KCTD15 | 0.65 | 5.61E-03 | Ion Transport |  |
| CACNA1F | 0.64 | 2.17E-02 | Ion Transport |  |
| SCN5A | 0.63 | 1.33E-02 | Ion Transport |  |
| SLC13A4 | 0.61 | 4.92E-02 | Ion Transport |  |
| GABRB1 | 0.61 | 3.55E-02 | Ion Transport |  |
| GABRG3 | 0.60 | 1.72E-02 | Ion Transport |  |
| KCNE3 | 0.59 | 2.59E-02 | Ion Transport |  |
| KCNK1 | 0.57 | 4.26E-03 | Ion Transport |  |
| ATP2A1 | 0.55 | 4.31E-03 | Ion Transport |  |
| GABRA3 | 0.55 | 1.71E-02 | Ion Transport |  |
| KCNQ4 | 0.53 | 1.59E-02 | Ion Transport |  |
| BEST3 | 0.53 | 1.83E-02 | Ion Transport |  |
| CLIC5 | 0.52 | 8.70E-03 | Ion Transport |  |
| KCNE1L | 0.50 | 3.04E-02 | Ion Transport |  |
| KCNH8 | 0.48 | 2.14E-03 | Ion Transport |  |
| CLCNKA | 0.46 | 1.70E-02 | Ion Transport |  |
| SLC39A5 | 0.46 | 3.56E-04 | Ion Transport |  |
| TF | 0.44 | 2.30E-02 | Ion Transport |  |
| CHRNB4 | 0.42 | 3.41E-03 | Ion Transport |  |
| ATP4B | 0.32 | 1.66E-02 | Ion Transport |  |
| KCNJ13 | 0.31 | 1.10E-03 | Ion Transport |  |
| SLC30A2 | 0.29 | 1.33E-02 | Ion Transport |  |
| CACNG6 | 0.24 | 1.31E-02 | Ion Transport |  |
| BAAT | 1.66 | 2.55E-02 | Lipid (Bile Acid) |  |
| CYP27B1 | 0.46 | 4.66E-02 | Lipid (Cofactor) |  |
| HPGD | 1.52 | 2.99E-02 | Lipid (Hormone) |  |
| DIO2 | 1.46 | 3.39E-02 | Lipid (Hormone) |  |
| MMEL1 | 1.34 | 2.70E-02 | Lipid (Hormone) |  |
| PLA2G3 | 5.47 | 2.21E-03 | Lipid (Membrane Lipid) |  |
| PHOSPHO1 | 3.88 | 3.88E-03 | Lipid (Membrane Lipid) |  |
| AGPAT9 | 3.43 | 3.21E-05 | Lipid (Membrane Lipid) |  |
| PLCB1 | 2.44 | 9.37E-05 | Lipid (Membrane Lipid) |  |
| PLA2G4A | 2.43 | 1.55E-03 | Lipid (Membrane Lipid) |  |
| DAGLA | 1.85 | 1.59E-02 | Lipid (Membrane Lipid) |  |
| LPCAT4 | 1.82 | 3.74E-03 | Lipid (Membrane Lipid) |  |
| ARSE | 1.81 | 2.46E-02 | Lipid (Membrane Lipid) |  |
| DGKG | 1.74 | 2.33E-02 | Lipid (Membrane Lipid) |  |
| CDS2 | 1.61 | 1.47E-03 | Lipid (Membrane Lipid) |  |
| PTDSS1 | 1.43 | 1.34E-03 | Lipid (Membrane Lipid) |  |
| DGKA | 1.42 | 2.41E-02 | Lipid (Membrane Lipid) |  |
| PCYT2 | 1.41 | 1.57E-03 | Lipid (Membrane Lipid) |  |
| PLA2G15 | 1.37 | 2.74E-03 | Lipid (Membrane Lipid) |  |
| PLD3 | 1.34 | 3.66E-02 | Lipid (Membrane Lipid) |  |
| DGKD | 0.77 | 2.49E-02 | Lipid (Membrane Lipid) |  |
| PPAP2B | 0.75 | 2.21E-02 | Lipid (Membrane Lipid) |  |
| PLCH1 | 0.75 | 1.51E-02 | Lipid (Membrane Lipid) |  |
| PLCG2 | 0.69 | 6.59E-03 | Lipid (Membrane Lipid) |  |
| PLCXD3 | 0.66 | 4.14E-02 | Lipid (Membrane Lipid) |  |
| PLA2G4C | 0.65 | 2.74E-02 | Lipid (Membrane Lipid) |  |
| GPD1 | 0.63 | 2.71E-02 | Lipid (Membrane Lipid) |  |
| MBOAT1 | 0.60 | 2.42E-03 | Lipid (Membrane Lipid) |  |
| PPAPDC1A | 0.25 | 3.89E-02 | Lipid (Membrane Lipid) |  |
| SGMS2 | 1.85 | 1.31E-03 | Lipid (Sphingolipid) |  |
| UGT8 | 1.75 | 5.81E-03 | Lipid (Sphingolipid) |  |
| CERKL | 0.65 | 1.89E-03 | Lipid (Sphingolipid) |  |
| SMPD2 | 0.61 | 3.65E-03 | Lipid (Sphingolipid) |  |
| SPHK1 | 0.43 | 3.01E-03 | Lipid (Sphingolipid) |  |
| ACER3 | 1.42 | 5.41E-03 | Lipid (Sphingosine) |  |
| HSD11B1 | 6.75 | 2.88E-02 | Lipid (Steroid) |  |
| CYP27C1 | 3.25 | 2.23E-02 | Lipid (Steroid) |  |
| CYP2S1 | 3.14 | 3.47E-03 | Lipid (Steroid) |  |
| HSD17B6 | 2.03 | 6.84E-04 | Lipid (Steroid) |  |
| CYP3A5 | 1.92 | 6.91E-03 | Lipid (Steroid) |  |
| HSD17B13 | 1.65 | 4.52E-02 | Lipid (Steroid) |  |
| HSD17B11 | 1.44 | 4.96E-02 | Lipid (Steroid) |  |
| HSD17B3 | 0.67 | 2.65E-02 | Lipid (Steroid) |  |
| CYP39A1 | 0.65 | 3.20E-02 | Lipid (Steroid) |  |
| CYP19A1 | 0.22 | 4.89E-02 | Lipid (Steroid) |  |
| HSD17B2 | 0.12 | 1.16E-02 | Lipid (Steroid) |  |
| ACHE | 1.81 | 1.12E-02 | Neurotransmitter |  |
| DDC | 0.67 | 1.18E-02 | Neurotransmitter |  |
| NAALADL1 | 0.46 | 2.46E-03 | Neurotransmitter |  |
| CA2 | 0.57 | 1.88E-03 | Nitrogen Metabolism |  |
| CA13 | 0.45 | 2.35E-02 | Nitrogen Metabolism |  |
| ENTPD3 | 3.00 | 7.56E-04 | Nucleotide |  |
| ENPP1 | 2.43 | 8.66E-04 | Nucleotide |  |
| NUDT10 | 1.92 | 1.24E-03 | Nucleotide |  |
| NT5M | 1.86 | 2.40E-02 | Nucleotide |  |
| NUDT8 | 1.78 | 1.50E-02 | Nucleotide |  |
| NUDT11 | 1.49 | 4.12E-02 | Nucleotide |  |
| NUDT18 | 1.40 | 9.00E-03 | Nucleotide |  |
| NUDT1 | 1.31 | 5.26E-03 | Nucleotide |  |
| NT5E | 0.65 | 4.29E-02 | Nucleotide |  |
| ENTPD2 | 0.35 | 9.24E-03 | Nucleotide |  |
| ADCY3 | 3.14 | 9.53E-05 | Nucleotide (cAMP) |  |
| ADCY8 | 1.94 | 2.63E-04 | Nucleotide (cAMP) |  |
| NPR3 | 3.21 | 6.47E-04 | Nucleotide (cGMP) |  |
| PDE4A | 1.68 | 1.65E-02 | Nucleotide (cGMP) |  |
| PDE6A | 0.70 | 4.85E-02 | Nucleotide (cGMP) |  |
| GUCY2F | 0.62 | 2.81E-02 | Nucleotide (cGMP) |  |
| GUCY2C | 0.24 | 2.64E-02 | Nucleotide (cGMP) |  |
| PDE7A | 0.74 | 3.23E-02 | Nucleotide (cyclic nucleotide) |  |
| PDE8A | 0.68 | 1.94E-02 | Nucleotide (cyclic nucleotide) |  |
| PDE1B | 0.68 | 4.59E-03 | Nucleotide (cyclic nucleotide) |  |
| PDE8B | 0.63 | 1.05E-02 | Nucleotide (cyclic nucleotide) |  |
| ADAD2 | 1.87 | 3.47E-02 | Nucleotide (Purine) |  |
| ADSS | 1.39 | 6.48E-04 | Nucleotide (Purine) |  |
| AMPD3 | 1.35 | 2.97E-03 | Nucleotide (Purine) |  |
| PRPS2 | 0.72 | 4.48E-02 | Nucleotide (Purine) |  |
| DPYSL2 | 1.75 | 2.24E-04 | Nucleotide (Pyrimidine) |  |
| DPYSL4 | 1.39 | 1.19E-02 | Nucleotide (Pyrimidine) |  |
| CMPK | 1.37 | 3.97E-04 | Nucleotide (Pyrimidine) |  |
| OAS1 | 0.54 | 2.70E-02 | Nucleotide (Ribonuclease) |  |
| BLVRB | 2.12 | 8.87E-05 | Porphyrin |  |
| HMOX1 | 1.79 | 1.73E-02 | Porphyrin |  |
| HMOX2 | 1.38 | 1.32E-04 | Porphyrin |  |
| PPOX | 0.76 | 7.69E-03 | Porphyrin |  |
| ATP13A2 | 2.09 | 2.30E-03 | Proton Transport |  |
| ATP10A | 1.79 | 2.61E-02 | Proton Transport |  |
| ATP8B2 | 1.39 | 2.78E-02 | Proton Transport |  |
| ATP6V1E1 | 1.39 | 1.29E-04 | Proton Transport |  |
| ATP6V1B1 | 0.61 | 6.72E-03 | Proton Transport |  |
| MICAL2 | 3.06 | 3.73E-05 | Redox |  |
| GLRX | 2.25 | 2.84E-04 | Redox |  |
| CYB561 | 1.64 | 1.11E-03 | Redox |  |
| ENOX1 | 1.61 | 1.16E-02 | Redox |  |
| CYB5R3 | 1.49 | 5.10E-03 | Redox |  |
| PRDX5 | 1.42 | 1.44E-04 | Redox |  |
| CYB5R4 | 1.32 | 7.68E-04 | Redox |  |
| OGFOD2 | 0.74 | 2.13E-02 | Redox |  |
| CYBASC3 | 0.71 | 1.54E-02 | Redox |  |
| DHRS3 | 0.61 | 3.67E-02 | Redox |  |
| CP | 0.46 | 1.13E-02 | Redox |  |
| SIAE | 1.82 | 7.81E-05 | Sugar |  |
| ABHD10 | 1.37 | 2.67E-04 | Sugar |  |
| CRYL1 | 0.69 | 1.21E-02 | Sugar |  |
| NANS | 2.09 | 4.46E-05 | Sugar (Aminosugar) |  |
| CMAS | 1.38 | 1.03E-04 | Sugar (Aminosugar) |  |
| KHK | 1.69 | 3.41E-03 | Sugar (Fructose) |  |
| ENOSF1 | 0.60 | 3.78E-02 | Sugar (Fructose, mannose) |  |
| GALK1 | 1.60 | 2.09E-02 | Sugar (Galactose) |  |
| IP6K3 | 2.53 | 7.33E-03 | Sugar (Inositol Phosphate) |  |
| PIK3CD | 2.41 | 6.04E-03 | Sugar (Inositol Phosphate) |  |
| INPP5J | 2.07 | 1.59E-03 | Sugar (Inositol Phosphate) |  |
| INPP4B | 1.69 | 2.38E-03 | Sugar (Inositol Phosphate) |  |
| INPP5A | 1.52 | 1.07E-02 | Sugar (Inositol Phosphate) |  |
| INPP4A | 1.35 | 2.98E-02 | Sugar (Inositol Phosphate) |  |
| PI4KA | 1.35 | 3.19E-03 | Sugar (Inositol Phosphate) |  |
| INPPL1 | 0.76 | 1.30E-02 | Sugar (Inositol Phosphate) |  |
| PIP5K1B | 0.59 | 4.47E-02 | Sugar (Inositol Phosphate) |  |
| ISYNA1 | 0.39 | 1.79E-03 | Sugar (Inositol Phosphate) |  |
| GMDS | 1.31 | 3.52E-02 | Sugar (Mannose) |  |
| UXS1 | 1.45 | 1.30E-04 | Sugar (Nucleotide Sugar) |  |
| TKT | 1.30 | 1.46E-03 | Sugar (PPP) |  |
| PGLS | 0.76 | 4.19E-02 | Sugar (PPP) |  |
| XYLB | 0.72 | 1.88E-02 | Sugar (Xyulose) |  |
| SULT2B1 | 4.70 | 4.24E-03 | Sulfate |  |
| SULT4A1 | 2.06 | 8.04E-04 | Sulfate |  |
| COQ4 | 0.72 | 2.48E-03 | Ubiquinone |  |
| SQRDL | 0.62 | 4.43E-02 | Ubiquinone |  |
| SDR16C5 | 2.19 | 1.24E-03 | Vitamin (A) |  |
| RDH12 | 0.70 | 2.42E-02 | Vitamin (A) |  |
| RDH10 | 0.44 | 7.40E-03 | Vitamin (A) |  |
| CYP26B1 | 0.11 | 3.77E-02 | Vitamin (A) |  |
| MTRR | 0.75 | 2.82E-02 | Vitamin (B12) |  |
| PDXP | 1.44 | 1.48E-02 | Vitamin (B6) |  |
| PDXK | 1.42 | 3.23E-02 | Vitamin (B6) |  |
| BTD | 0.77 | 3.91E-02 | Vitamin (Biotin) |  |
| MTHFD2 | 1.76 | 4.22E-03 | Vitamin (Folate) |  |
| GGH | 1.62 | 7.12E-04 | Vitamin (Folate) |  |
| ALDH1A2 | 0.71 | 1.29E-02 | Vitamine (Retinoic acid, Alcohol) |  |

**Table S10.** **Comparing the changed metabolic genes in the retinas between WT vs NRL KO and macula vs periphery.**

| **Gene symbol** | **FC** | **P value** | **Gene symbol** | **FC** | **P value** | **Classification (Sub-category/substrate)** |
| --- | --- | --- | --- | --- | --- | --- |
| *Abcb4* | 4.63 | 6.80E-07 | ABCB4 | 2.04 | 2.92E-02 | ABC Transporter |
| *Abcg1* | 1.34 | 1.26E-03 | ABCG1 | 0.63 | 2.15E-02 | ABC Transporter |
| *Abcg4* | 0.76 | 1.22E-02 | ABCG4 | 0.72 | 1.00E-02 | ABC Transporter |
| *Abcg5* | 0.09 | 3.42E-02 | ABCG5 | 0.66 | 1.52E-02 | ABC Transporter |
| *Nos1* | 1.73 | 2.40E-02 | NOS1 | 0.55 | 1.34E-02 | Amino Acid (Arginine) |
| *Ddah1* | 1.53 | 7.66E-03 | DDAH1 | 1.85 | 8.04E-04 | Amino Acid (Arginine, Citrulline) |
| *Bcat1* | 2.32 | 1.25E-04 | BCAT1 | 1.60 | 6.26E-04 | Amino Acid (BCAAs) |
| *Gad2* | 1.50 | 1.08E-02 | GAD2 | 0.74 | 4.28E-02 | Amino Acid (Glutamate) |
| *Hdc* | 1.23E-04 | 1.55E-02 | HDC | 0.46 | 3.36E-02 | Amino Acid (histidine) |
| *Pipox* | 0.47 | 6.91E-03 | PIPOX | 0.72 | 1.05E-03 | Amino Acid (Lysine) |
| *Adi1* | 1.32 | 2.22E-02 | ADI1 | 0.76 | 8.76E-03 | Amino Acid (Methionine) |
| *Ephx2* | 1.38 | 2.37E-02 | EPHX2 | 0.67 | 3.91E-02 | Detox |
| *B4galt5* | 2.37 | 5.85E-03 | B4GALT5 | 2.04 | 2.67E-03 | Glycan |
| *B4galt6* | 2.05 | 3.02E-02 | B4GALT6 | 1.38 | 6.31E-03 | Glycan |
| *Fut9* | 1.42 | 2.87E-02 | FUT9 | 1.59 | 9.79E-03 | Glycan |
| *Mgat3* | 1.56 | 4.56E-03 | MGAT3 | 1.46 | 3.03E-02 | Glycan |
| *Mgat5* | 1.78 | 3.23E-03 | MGAT5 | 1.68 | 4.57E-03 | Glycan |
| *St8sia2* | 1.62 | 1.28E-03 | ST8SIA2 | 1.86 | 3.93E-02 | Glycan |
| *St8sia3* | 1.45 | 1.82E-02 | ST8SIA3 | 1.39 | 7.10E-03 | Glycan |
| *B3gnt8* | 0.55 | 4.40E-04 | B3GNT8 | 3.26 | 8.37E-03 | Glycan |
| *St8sia1* | 0.23 | 5.73E-04 | ST8SIA1 | 1.77 | 4.17E-03 | Glycan |
| *B4galt1* | 0.14 | 5.55E-07 | B4GALT1 | 0.74 | 2.05E-02 | Glycan |
| *Ctbs* | 0.34 | 5.25E-03 | CTBS | 0.70 | 2.85E-02 | Glycan |
| *Galnt10* | 0.46 | 1.12E-04 | GALNT10 | 0.60 | 2.73E-02 | Glycan |
| *Galnt4* | 0.44 | 1.19E-03 | GALNT4 | 0.76 | 7.24E-03 | Glycan |
| *Galnt7* | 0.37 | 3.83E-03 | GALNT7 | 0.76 | 7.59E-03 | Glycan |
| *St3gal1* | 0.17 | 8.69E-05 | ST3GAL1 | 0.67 | 2.12E-02 | Glycan |
| *St6galnac2* | 0.34 | 2.89E-04 | ST6GALNAC2 | 0.59 | 1.77E-04 | Glycan |
| *St6galnac5* | 0.46 | 2.72E-04 | ST6GALNAC5 | 0.68 | 7.64E-04 | Glycan |
| *Glb1l* | 0.37 | 5.91E-03 | GLB1L | 0.71 | 4.58E-02 | Glycan (Degradation) |
| *Glb1l2* | 0.05 | 3.49E-06 | GLB1L2 | 0.67 | 3.46E-02 | Glycan (Degradation) |
| *Man2c1* | 0.42 | 8.61E-04 | MAN2C1 | 0.72 | 5.79E-03 | Glycan (Degradation) |
| *Sulf2* | 2.20 | 2.50E-02 | SULF2 | 1.67 | 2.52E-03 | Glycan (Sulfate) |
| *Chst3* | 0.07 | 2.62E-04 | CHST3 | 0.61 | 5.92E-03 | Glycan (Sulfate) |
| *Pgm2l1* | 1.55 | 1.03E-02 | PGM2L1 | 1.54 | 2.58E-02 | Glycolysis |
| *Pgk1* | 0.67 | 8.90E-03 | PGK1 | 1.33 | 4.93E-02 | Glycolysis |
| *Pfkfb2* | 0.25 | 1.21E-02 | PFKFB2 | 0.65 | 6.13E-03 | Glycolysis |
| *Hs3st1* | 1.65 | 2.16E-02 | HS3ST1 | 1.71 | 3.15E-02 | Heparin Sulfate |
| *Hs6st3* | 4.08 | 1.61E-03 | HS6ST3 | 2.61 | 9.44E-04 | Heparin Sulfate |
| *Hs3st3b1* | 0.63 | 2.51E-02 | HS3ST3B1 | 0.72 | 1.41E-02 | Heparin Sulfate |
| *Atp2b2* | 1.80 | 6.88E-03 | ATP2B2 | 1.81 | 2.14E-03 | Ion Transport |
| *Atp2b3* | 2.11 | 7.31E-03 | ATP2B3 | 2.11 | 1.01E-03 | Ion Transport |
| *Cacng2* | 1.49 | 1.67E-02 | CACNG2 | 1.78 | 1.17E-02 | Ion Transport |
| *Chrnb2* | 1.55 | 1.60E-03 | CHRNB2 | 1.38 | 1.92E-02 | Ion Transport |
| *Clcn6* | 1.42 | 6.88E-03 | CLCN6 | 1.52 | 3.14E-04 | Ion Transport |
| *Gabrb2* | 2.10 | 2.65E-02 | GABRB2 | 1.53 | 4.54E-03 | Ion Transport |
| *Glra1* | 4.02 | 1.79E-02 | GLRA1 | 1.87 | 1.96E-03 | Ion Transport |
| *Grin1* | 1.45 | 1.65E-02 | GRIN1 | 2.02 | 1.45E-02 | Ion Transport |
| *Grin2a* | 1.64 | 1.67E-02 | GRIN2A | 2.12 | 2.03E-02 | Ion Transport |
| *Hcn2* | 1.40 | 2.52E-02 | HCN2 | 1.88 | 2.30E-03 | Ion Transport |
| *Kcna2* | 1.55 | 1.07E-02 | KCNA2 | 2.36 | 1.85E-03 | Ion Transport |
| *Kcnab2* | 1.89 | 5.41E-04 | KCNAB2 | 1.50 | 3.66E-02 | Ion Transport |
| *Kcnc1* | 1.32 | 6.20E-03 | KCNC1 | 1.90 | 4.92E-03 | Ion Transport |
| *Kcnd3* | 2.36 | 2.57E-04 | KCND3 | 1.36 | 1.28E-02 | Ion Transport |
| *Kcne2* | 2.75 | 8.49E-03 | KCNE2 | 1.60 | 1.63E-02 | Ion Transport |
| *Kcng1* | 7.43 | 4.42E-02 | KCNG1 | 1.97 | 2.92E-02 | Ion Transport |
| *Kcng4* | 2.33 | 1.55E-03 | KCNG4 | 1.42 | 2.33E-02 | Ion Transport |
| *Kcnj11* | 1.44 | 2.92E-02 | KCNJ11 | 1.70 | 6.30E-03 | Ion Transport |
| *Kcnj12* | 1.68 | 4.02E-03 | KCNJ12 | 1.58 | 2.90E-02 | Ion Transport |
| *Kcnk3* | 1.61 | 1.84E-02 | KCNK3 | 1.36 | 4.65E-03 | Ion Transport |
| *P2rx6* | 1.37 | 2.94E-02 | P2RX6 | 2.41 | 2.35E-05 | Ion Transport |
| *Scn2b* | 1.33 | 1.01E-02 | SCN2B | 2.22 | 9.62E-04 | Ion Transport |
| *Slc24a2* | 4.51 | 4.58E-04 | SLC24A2 | 1.46 | 1.25E-02 | Ion Transport |
| *Slc24a3* | 1.56 | 1.40E-02 | SLC24A3 | 1.47 | 1.69E-02 | Ion Transport |
| *Slc39a11* | 1.68 | 2.29E-02 | SLC39A11 | 1.46 | 2.45E-02 | Ion Transport |
| *Slc41a3* | 1.36 | 9.20E-03 | SLC41A3 | 1.42 | 3.42E-03 | Ion Transport |
| *Slc9a6* | 1.39 | 7.07E-04 | SLC9A6 | 1.32 | 3.10E-04 | Ion Transport |
| *Atp1b2* | 2.02 | 5.61E-06 | ATP1B2 | 0.75 | 1.10E-03 | Ion Transport |
| *Best3* | 2.01 | 2.24E-02 | BEST3 | 0.53 | 1.83E-02 | Ion Transport |
| *Chrnb4* | 18.82 | 2.42E-02 | CHRNB4 | 0.42 | 3.41E-03 | Ion Transport |
| *Clcnka* | 4.08 | 4.57E-02 | CLCNKA | 0.46 | 1.70E-02 | Ion Transport |
| *Clic4* | 3.19 | 2.22E-04 | CLIC4 | 0.73 | 2.10E-02 | Ion Transport |
| *Gabrb1* | 1.56 | 1.74E-02 | GABRB1 | 0.61 | 3.55E-02 | Ion Transport |
| *Gria1* | 1.52 | 2.52E-02 | GRIA1 | 0.69 | 2.74E-02 | Ion Transport |
| *Heph* | 1.66 | 1.54E-04 | HEPH | 0.74 | 4.54E-02 | Ion Transport |
| *Kcnq4* | 1.83 | 3.33E-02 | KCNQ4 | 0.53 | 1.59E-02 | Ion Transport |
| *Kctd15* | 1.31 | 2.45E-02 | KCTD15 | 0.65 | 5.61E-03 | Ion Transport |
| *Scn5a* | 1.64 | 3.88E-02 | SCN5A | 0.63 | 1.33E-02 | Ion Transport |
| *Ttyh2* | 4.03 | 3.16E-05 | TTYH2 | 0.70 | 1.80E-02 | Ion Transport |
| *Cacna2d3* | 0.63 | 1.63E-02 | CACNA2D3 | 1.41 | 5.60E-03 | Ion Transport |
| *Clic3* | 0.37 | 2.23E-02 | CLIC3 | 12.88 | 2.53E-02 | Ion Transport |
| *Cacna1f* | 0.51 | 4.76E-03 | CACNA1F | 0.64 | 2.17E-02 | Ion Transport |
| *Cnga1* | 2.49E-04 | 1.08E-06 | CNGA1 | 0.71 | 4.88E-02 | Ion Transport |
| *Aacs* | 0.62 | 3.18E-02 | AACS | 1.56 | 2.12E-03 | Ketone Bodies |
| *Baat* | 0.20 | 2.67E-02 | BAAT | 1.66 | 2.55E-02 | Lipid (Bile Acid) |
| *Dhcr7* | 0.69 | 1.29E-02 | DHCR7 | 1.49 | 7.39E-03 | Lipid (Cholesterol) |
| *Fdft1* | 0.74 | 1.18E-03 | FDFT1 | 1.36 | 1.37E-02 | Lipid (Cholesterol) |
| *Fdps* | 0.61 | 1.43E-03 | FDPS | 1.51 | 1.42E-03 | Lipid (Cholesterol) |
| *Acaca* | 1.53 | 2.82E-02 | ACACA | 1.61 | 3.14E-05 | Lipid (Fatty Acid) |
| *Elovl6* | 2.61 | 7.20E-03 | ELOVL6 | 2.03 | 3.51E-05 | Lipid (Fatty Acid) |
| *Acads* | 2.57 | 4.11E-03 | ACADS | 0.71 | 1.45E-02 | Lipid (Fatty Acid) |
| *Acat2* | 0.65 | 1.48E-03 | ACAT2 | 2.32 | 5.60E-04 | Lipid (Fatty Acid) |
| *Acss2* | 0.75 | 1.89E-02 | ACSS2 | 1.35 | 5.02E-03 | Lipid (Fatty Acid) |
| *Far2* | 0.68 | 2.92E-02 | FAR2 | 2.55 | 5.74E-04 | Lipid (Fatty Acid) |
| *Acox2* | 0.25 | 3.87E-02 | ACOX2 | 0.61 | 1.47E-02 | Lipid (Fatty Acid) |
| *Amacr* | 0.49 | 8.77E-04 | AMACR | 0.70 | 1.13E-02 | Lipid (Fatty acid) |
| *Elovl1* | 0.43 | 1.90E-04 | ELOVL1 | 0.75 | 3.56E-03 | Lipid (Fatty Acid) |
| *Mecr* | 0.26 | 2.22E-05 | MECR | 0.77 | 4.82E-02 | Lipid (Fatty Acid) |
| *Dio2* | 1.45 | 3.75E-02 | DIO2 | 1.46 | 3.39E-02 | Lipid (Hormone) |
| *Hpgd* | 0.72 | 1.47E-02 | HPGD | 1.52 | 2.99E-02 | Lipid (Hormone) |
| *Cds2* | 1.83 | 4.68E-03 | CDS2 | 1.61 | 1.47E-03 | Lipid (Membrane Lipid) |
| *Dagla* | 1.64 | 7.62E-03 | DAGLA | 1.85 | 1.59E-02 | Lipid (Membrane Lipid) |
| *Dgkg* | 1.40 | 2.26E-02 | DGKG | 1.74 | 2.33E-02 | Lipid (Membrane Lipid) |
| *Phospho1* | 2.37 | 1.11E-02 | PHOSPHO1 | 3.88 | 3.88E-03 | Lipid (Membrane Lipid) |
| *Pla2g15* | 1.41 | 3.07E-03 | PLA2G15 | 1.37 | 2.74E-03 | Lipid (Membrane Lipid) |
| *Plcb1* | 1.57 | 1.96E-02 | PLCB1 | 2.44 | 9.37E-05 | Lipid (Membrane Lipid) |
| *Dgkd* | 1.33 | 4.25E-02 | DGKD | 0.77 | 2.49E-02 | Lipid (Membrane Lipid) |
| *Gpd1* | 1.45 | 8.21E-03 | GPD1 | 0.63 | 2.71E-02 | Lipid (Membrane Lipid) |
| *Mboat1* | 1.35 | 3.09E-02 | MBOAT1 | 0.60 | 2.42E-03 | Lipid (Membrane Lipid) |
| *Plcg2* | 2.03 | 3.66E-02 | PLCG2 | 0.69 | 6.59E-03 | Lipid (Membrane Lipid) |
| *Plcxd3* | 2.77 | 2.27E-02 | PLCXD3 | 0.66 | 4.14E-02 | Lipid (Membrane Lipid) |
| *Dgka* | 0.61 | 3.91E-04 | DGKA | 1.42 | 2.41E-02 | Lipid (Membrane Lipid) |
| *Pcyt2* | 0.58 | 8.23E-03 | PCYT2 | 1.41 | 1.57E-03 | Lipid (Membrane Lipid) |
| *Pla2g3* | 0.26 | 4.11E-05 | PLA2G3 | 5.47 | 2.21E-03 | Lipid (Membrane Lipid) |
| *Idi1* | 0.57 | 5.60E-03 | IDI1 | 1.50 | 6.89E-04 | Lipid (Mevalonate) |
| *Smpd2* | 0.58 | 7.51E-03 | SMPD2 | 0.61 | 3.65E-03 | Lipid (Sphingolipid) |
| *Hsd17b2* | 2.24 | 1.45E-02 | HSD17B2 | 0.12 | 1.16E-02 | Lipid (Steroid) |
| *Ache* | 1.43 | 1.39E-02 | ACHE | 1.81 | 1.12E-02 | Neurotransmitter |
| *Ddc* | 0.46 | 2.38E-02 | DDC | 0.67 | 1.18E-02 | Neurotransmitter |
| *Enpp1* | 0.69 | 2.42E-02 | ENPP1 | 2.43 | 8.66E-04 | Nucleotide |
| *Nt5e* | 0.17 | 1.16E-06 | NT5E | 0.65 | 4.29E-02 | Nucleotide |
| *Adcy8* | 1.30 | 4.38E-02 | ADCY8 | 1.94 | 2.63E-04 | Nucleotide (cAMP) |
| *Npr3* | 1.75 | 4.77E-02 | NPR3 | 3.21 | 6.47E-04 | Nucleotide (cGMP) |
| *Pde4a* | 1.31 | 2.74E-02 | PDE4A | 1.68 | 1.65E-02 | Nucleotide (cGMP) |
| *Gucy2f* | 0.02 | 4.25E-03 | GUCY2F | 0.62 | 2.81E-02 | Nucleotide (cGMP) |
| *Pde6a* | 0.07 | 8.92E-05 | PDE6A | 0.70 | 4.85E-02 | Nucleotide (cGMP) |
| *Pde7a* | 0.66 | 3.27E-02 | PDE7A | 0.74 | 3.23E-02 | Nucleotide (cyclic nucleotide) |
| *Ampd3* | 1.44 | 4.56E-02 | AMPD3 | 1.35 | 2.97E-03 | Nucleotide (Purine) |
| *Prps2* | 1.35 | 1.58E-02 | PRPS2 | 0.72 | 4.48E-02 | Nucleotide (Purine) |
| *Dpysl2* | 1.33 | 2.43E-02 | DPYSL2 | 1.75 | 2.24E-04 | Nucleotide (Pyrimidine) |
| *Atp10a* | 1.63 | 2.20E-02 | ATP10A | 1.79 | 2.61E-02 | Proton Transport |
| *Cyb561* | 2.13 | 2.47E-03 | CYB561 | 1.64 | 1.11E-03 | Redox |
| *Enox1* | 0.54 | 1.62E-02 | ENOX1 | 1.61 | 1.16E-02 | Redox |
| *Dhrs3* | 0.16 | 1.90E-04 | DHRS3 | 0.61 | 3.67E-02 | Redox |
| *Ogfod2* | 0.59 | 2.65E-03 | OGFOD2 | 0.74 | 2.13E-02 | Redox |
| *Slc25a35* | 0.47 | 1.54E-03 | SLC25A35 | 0.71 | 6.06E-03 | Small Molecule Transport ( Di-Tri-carboxylic acid) |
| *Slc25a29* | 0.36 | 1.90E-05 | SLC25A29 | 0.69 | 8.43E-03 | Small Molecule Transport (Acylcarnitine) |
| *Slc7a14* | 1.63 | 1.06E-02 | SLC7A14 | 1.35 | 2.61E-02 | Small Molecule Transport (Amino acid) |
| *Slc25a45* | 1.78 | 4.62E-03 | SLC25A45 | 0.59 | 4.01E-02 | Small Molecule Transport (Amino acid) |
| *Slc4a3* | 2.02 | 1.01E-03 | SLC4A3 | 1.89 | 2.45E-03 | Small Molecule Transport (Anion) |
| *Slco4a1* | 0.19 | 2.86E-04 | SLCO4A1 | 0.59 | 1.38E-03 | Small Molecule Transport (bile acids,prostaglandins,steroid conjugates,thyroid hormone) |
| *Slc4a4* | 1.61 | 3.58E-02 | SLC4A4 | 1.35 | 1.58E-02 | Small Molecule Transport (Biocarbonate) |
| *Slc27a1* | 0.64 | 5.71E-04 | SLC27A1 | 0.70 | 2.71E-03 | Small Molecule Transport (Fatty acid) |
| *Slc27a3* | 0.64 | 4.82E-02 | SLC27A3 | 0.65 | 2.55E-02 | Small Molecule Transport (Fatty acid) |
| *Slc6a11* | 1.95 | 1.77E-03 | SLC6A11 | 0.59 | 3.93E-02 | Small Molecule Transport (GABA, β-Alanine) |
| *Slc25a18* | 0.46 | 1.72E-02 | SLC25A18 | 1.38 | 4.84E-02 | Small Molecule Transport (Glutamate) |
| *Slc1a1* | 1.56 | 9.18E-03 | SLC1A1 | 1.40 | 3.60E-03 | Small Molecule Transport (Glutamate, Asparate) |
| *Slc6a9* | 1.48 | 1.44E-02 | SLC6A9 | 0.62 | 1.07E-03 | Small Molecule Transport (Glycine, Sarcosine) |
| *Slc5a8* | 0.38 | 4.04E-02 | SLC5A8 | 2.04 | 1.67E-02 | Small Molecule Transport (iodide) |
| *Slc16a3* | 0.57 | 2.34E-03 | SLC16A3 | 2.21 | 7.41E-03 | Small Molecule Transport (Lactate/Pyruvate) |
| *Slc2a13* | 1.57 | 5.80E-03 | SLC2A13 | 1.97 | 3.08E-03 | Small Molecule Transport (Myo-Inoisitol) |
| *Slc35f3* | 1.52 | 4.15E-02 | SLC35F3 | 1.80 | 4.45E-02 | Small Molecule Transport (Nucleotide sugar) |
| *Slc35f4* | 0.53 | 5.72E-03 | SLC35F4 | 0.67 | 1.50E-02 | Small Molecule Transport (Nucleotide sugar) |
| *Slc25a43* | 0.44 | 1.53E-02 | SLC25A43 | 2.26 | 8.34E-03 | Small Molecule Transport (Nucleotide) |
| *Slc15a3* | 2.09 | 1.87E-02 | SLC15A3 | 0.54 | 1.17E-02 | Small Molecule Transport (peptide) |
| *Slc6a17* | 1.35 | 1.26E-02 | SLC6A17 | 2.87 | 1.53E-03 | Small Molecule Transport (Pro, Cys, Ala) |
| *Sv2a* | 1.75 | 1.12E-03 | SV2A | 1.47 | 3.68E-03 | Small Molecule Transport (Sugar) |
| *Sv2c* | 1.52 | 1.08E-02 | SV2C | 2.63 | 1.53E-02 | Small Molecule Transport (Sugar) |
| *Slc26a11* | 0.33 | 2.92E-05 | SLC26A11 | 0.68 | 1.34E-03 | Small Molecule Transport (Sulfate) |
| *Slc35d1* | 0.71 | 1.46E-02 | SLC35D1 | 0.71 | 1.36E-04 | Small Molecule Transport (UDP-N-Acetyl-Glucosomine) |
| *Siae* | 1.44 | 4.57E-02 | SIAE | 1.82 | 7.81E-05 | Sugar |
| *Khk* | 0.53 | 6.14E-04 | KHK | 1.69 | 3.41E-03 | Sugar (Fructose) |
| *Pygm* | 5.45 | 3.80E-05 | PYGM | 2.94 | 1.69E-04 | Sugar (Glycogen) |
| *Inpp4a* | 1.82 | 1.20E-02 | INPP4A | 1.35 | 2.98E-02 | Sugar (Inositol Phosphate) |
| *Inpp4b* | 1.45 | 1.68E-02 | INPP4B | 1.69 | 2.38E-03 | Sugar (Inositol Phosphate) |
| *Inppl1* | 3.31 | 2.18E-04 | INPPL1 | 0.76 | 1.30E-02 | Sugar (Inositol Phosphate) |
| *Pip5k1b* | 1.47 | 8.50E-04 | PIP5K1B | 0.59 | 4.47E-02 | Sugar (Inositol Phosphate) |
| *Ip6k3* | 0.33 | 4.77E-02 | IP6K3 | 2.53 | 7.33E-03 | Sugar (Inositol Phosphate) |
| *Isyna1* | 0.35 | 3.29E-02 | ISYNA1 | 0.39 | 1.79E-03 | Sugar (Inositol Phosphate) |
| *Pgls* | 0.74 | 2.22E-02 | PGLS | 0.76 | 4.19E-02 | Sugar (PPP) |
| *Acly* | 1.46 | 3.87E-03 | ACLY | 1.48 | 1.81E-05 | TCA Cycle |
| *Got1* | 1.33 | 2.35E-03 | GOT1 | 1.70 | 1.92E-05 | TCA Cycle |
| *Ogdh* | 1.64 | 1.23E-03 | OGDH | 1.32 | 3.68E-03 | TCA Cycle |
| *Pcx* | 1.59 | 8.21E-03 | PC | 1.59 | 2.97E-04 | TCA Cycle |
| *Aco1* | 0.76 | 2.41E-02 | ACO1 | 1.31 | 2.75E-04 | TCA Cycle |
| *Rdh12* | 0.18 | 4.34E-05 | RDH12 | 0.70 | 2.42E-02 | Vitamin (A) |
| *Pdxk* | 1.44 | 1.84E-02 | PDXK | 1.42 | 3.23E-02 | Vitamin (B6) |

**Table S11. Differentially expressed metabolic genes in macular and peripheral RPE/choroid.**

| **Gene symbol** | **FC** | **P value** | **Classification (Sub-category/substrate)** | **In figures** |
| --- | --- | --- | --- | --- |
| GAPDH | 0.76 | 2.30E-02 | Glycolysis | Fig 9b |
| ALDOA | 0.75 | 1.66E-02 | Glycolysis | Fig 9b |
| ADPGK | 0.74 | 4.79E-02 | Glycolysis | Fig 9b |
| PFKL | 0.66 | 3.29E-02 | Glycolysis | Fig 9b |
| LDHD | 0.62 | 3.88E-02 | Glycolysis | Fig 9b |
| GOT1 | 0.67 | 2.71E-02 | TCA Cycle | Fig 9b |
| GOT2 | 0.46 | 4.08E-02 | TCA Cycle | Fig 9b |
| COX6C | 0.76 | 6.16E-03 | ETC (Complex IV) | Fig 9b |
| ACN9 | 0.74 | 1.74E-02 | ETC (Complex II) | Fig 9b |
| NDUFB2 | 0.74 | 1.57E-02 | ETC (Complex I) | Fig 9b |
| STARD8 | 1.38 | 4.34E-02 | Small Molecule Transport (Lipid) | Fig 9c |
| SLC27A3 | 1.36 | 6.54E-03 | Small Molecule Transport (Fatty acid) | Fig 9c |
| SLC25A20 | 1.31 | 3.74E-02 | Small Molecule Transport (Carnitine) | Fig 9c |
| SLC5A9 | 2.01 | 3.27E-02 | Small Molecule Transport (Glucose) | Fig 9c |
| SLC2A4 | 1.57 | 9.21E-03 | Small Molecule Transport (Glucose, Glucosamine) | Fig 9c |
| SLC4A2 | 0.63 | 4.74E-02 | Small Molecule Transport (Biocarbonate) | Fig 9c |
| SLC4A10 | 0.21 | 2.75E-02 | Small Molecule Transport (Biocarbonate) | Fig 9c |
| AQP1 | 2.48 | 3.70E-02 | Small Molecule Transport (Water, CO2, Ammonium) | Fig 9c |
| AQP7 | 2.20 | 2.27E-02 | Small Molecule Transport (Water) | Fig 9c |
| SLCO5A1 | 2.56 | 3.18E-02 | Small Molecule Transport (peptide) | Fig 9c |
| SLC25A24 | 1.36 | 4.36E-02 | Small Molecule Transport (Mitochondrial Phosphate) | Fig 9c |
| SLC19A2 | 0.76 | 3.02E-02 | Small Molecule Transport (Thiamine) | Fig 9c |
| SLC5A6 | 0.72 | 1.42E-02 | Small Molecule Transport (Vitamine) | Fig 9c |
| SLC29A1 | 0.67 | 2.34E-02 | Small Molecule Transport (inosine) | Fig 9c |
| SLC7A1 | 0.67 | 3.27E-02 | Small Molecule Transport (Ar, Lys, Orn, His) | Fig 9c |
| SLC7A5 | 0.67 | 1.07E-02 | Small Molecule Transport (Unkown) | Fig 9c |
| SLC16A11 | 0.57 | 3.45E-02 | Small Molecule Transport (Pyryuvate) | Fig 9c |
| SLC25A10 | 0.49 | 2.97E-02 | Small Molecule Transport (Malate, Succinate) | Fig 9c |
| SLC6A12 | 0.33 | 1.91E-02 | Small Molecule Transport (GABA, Betaine) | Fig 9c |
| ABCA10 | 1.62 | 1.08E-02 | ABC Transporter | Fig 9d |
| ABCA5 | 1.62 | 1.08E-02 | ABC Transporter | Fig 9d |
| ABCA6 | 1.62 | 1.08E-02 | ABC Transporter | Fig 9d |
| ABCA8 | 1.62 | 1.08E-02 | ABC Transporter | Fig 9d |
| ABCA9 | 1.62 | 1.08E-02 | ABC Transporter | Fig 9d |
| CYP4A22 | 3.39 | 1.15E-02 | Fatty Acid | Fig 9d |
| CYP4Z1 | 3.13 | 3.62E-03 | Fatty Acid | Fig 9d |
| ACSM4 | 1.94 | 4.88E-02 | Fatty Acid | Fig 9d |
| PNPLA5 | 0.12 | 3.44E-02 | Fatty Acid | Fig 9d |
| DIO1 | 2.09 | 3.89E-02 | Hormone | Fig 9d |
| MMEL1 | 1.41 | 4.06E-02 | Hormone | Fig 9d |
| MOGAT2 | 2.78 | 4.31E-02 | Membrane Lipid | Fig 9d |
| DGKG | 2.05 | 9.69E-04 | Membrane Lipid | Fig 9d |
| DGKI | 1.64 | 1.31E-02 | Membrane Lipid | Fig 9d |
| DGKH | 1.51 | 1.45E-03 | Membrane Lipid | Fig 9d |
| PLCG2 | 1.44 | 2.50E-02 | Membrane Lipid | Fig 9d |
| LPIN3 | 1.40 | 1.55E-02 | Membrane Lipid | Fig 9d |
| PLD2 | 1.32 | 2.12E-02 | Membrane Lipid | Fig 9d |
| PPAPDC1B | 0.62 | 1.78E-02 | Membrane Lipid | Fig 9d |
| LPCAT2 | 0.52 | 3.64E-02 | Membrane Lipid | Fig 9d |
| CYP3A43 | 3.61 | 3.56E-02 | Steroid | Fig 9d |
| HSD17B6 | 1.93 | 3.66E-02 | Steroid | Fig 9d |
| CYP46A1 | 0.62 | 4.34E-02 | Steroid | Fig 9d |
| LIPE | 0.59 | 4.19E-02 | Steroid | Fig 9d |
| ASRGL1 | 0.54 | 4.43E-02 | Amino Acid (Asparagine) |  |
| AOC2 | 1.44 | 6.08E-03 | Amino acid (Dopamine, histamine) |  |
| AOC3 | 1.44 | 6.08E-03 | Amino acid (Dopamine, histamine) |  |
| GSTP1 | 0.74 | 4.98E-02 | Amino Acid (Glutathione) |  |
| HAGHL | 0.70 | 3.59E-02 | Amino Acid (Glutathione) |  |
| GPX4 | 0.61 | 2.88E-02 | Amino Acid (Glutathione) |  |
| GCAT | 0.65 | 5.17E-03 | Amino Acid (Glycine) |  |
| AADAT | 0.54 | 4.04E-02 | Amino Acid (Kyneurenine, glutamate) |  |
| MSRB3 | 1.34 | 2.26E-02 | Amino Acid (Methionine) |  |
| PNMT | 1.89 | 2.00E-02 | Amino Acid (Neurotransmitter) |  |
| PYCR1 | 0.58 | 3.91E-02 | Amino Acid (Proline) |  |
| PSAT | 0.51 | 2.74E-02 | Amino Acid (Serine) |  |
| TDO2 | 2.89 | 2.01E-02 | Amino Acid (Tryptophan) |  |
| INMT | 2.48 | 3.70E-02 | Amino Acid (Tryptophan) |  |
| ACOT6 | 0.40 | 3.97E-02 | CoA |  |
| BCHE | 2.18 | 1.15E-02 | Detox |  |
| AKR7L | 0.26 | 3.96E-02 | Detox |  |
| GBA3 | 4.10 | 2.22E-02 | Glycan |  |
| B3GALT1 | 1.99 | 1.98E-02 | Glycan |  |
| ST8SIA6 | 1.71 | 3.10E-02 | Glycan |  |
| ST6GALNAC3 | 1.65 | 7.83E-03 | Glycan |  |
| B4GALNT1 | 1.56 | 4.50E-02 | Glycan |  |
| DHDDS | 0.69 | 1.78E-02 | Glycan |  |
| MAN1A1 | 1.52 | 2.00E-02 | Glycan (Degradation) |  |
| HS3ST2 | 1.92 | 4.02E-02 | Heparin Sulfate |  |
| NDST3 | 0.23 | 4.62E-02 | Heparin Sulfate |  |
| HTR3C | 13.62 | 3.77E-02 | Ion Transport |  |
| KCNS1 | 4.51 | 1.63E-02 | Ion Transport |  |
| GRIN2A | 2.81 | 3.68E-02 | Ion Transport |  |
| GABRA4 | 2.73 | 3.69E-02 | Ion Transport |  |
| CLCA2 | 2.51 | 2.05E-02 | Ion Transport |  |
| GRIN2C | 2.31 | 5.70E-03 | Ion Transport |  |
| KCNK3 | 2.25 | 3.20E-02 | Ion Transport |  |
| GRIK2 | 1.77 | 1.81E-02 | Ion Transport |  |
| KCNMB1 | 1.58 | 2.92E-02 | Ion Transport |  |
| ATP2B4 | 1.57 | 4.06E-02 | Ion Transport |  |
| GRID1 | 1.49 | 3.68E-02 | Ion Transport |  |
| CACNB3 | 1.47 | 1.26E-03 | Ion Transport |  |
| KCTD1 | 1.45 | 4.85E-02 | Ion Transport |  |
| SLC3A2 | 0.66 | 2.01E-02 | Ion Transport |  |
| KCND3 | 0.63 | 2.36E-02 | Ion Transport |  |
| KCTD8 | 0.54 | 1.75E-02 | Ion Transport |  |
| KCNN2 | 0.45 | 4.87E-02 | Ion Transport |  |
| SLC12A1 | 0.35 | 4.14E-02 | Ion Transport |  |
| KCNK4 | 0.30 | 3.00E-03 | Ion Transport |  |
| GRIA2 | 0.26 | 4.90E-02 | Ion Transport |  |
| AACS | 0.75 | 3.14E-03 | Ketone Bodies |  |
| BDH1 | 0.58 | 1.10E-02 | Ketone Bodies |  |
| NMNAT2 | 2.70 | 1.17E-02 | NAD |  |
| NOX4 | 1.60 | 1.11E-02 | NAD |  |
| CA3 | 0.41 | 4.39E-02 | Nitrogen Metabolism |  |
| ENTPD1 | 1.40 | 6.32E-04 | Nucleotide |  |
| NT5DC3 | 0.76 | 4.54E-02 | Nucleotide |  |
| NT5M | 0.56 | 3.71E-02 | Nucleotide |  |
| ADCY10 | 2.43 | 3.92E-02 | Nucleotide (cAMP) |  |
| ADCY5 | 1.54 | 3.23E-02 | Nucleotide (cAMP) |  |
| PDE4C | 2.43 | 4.03E-02 | Nucleotide (cGMP) |  |
| PDE2A | 2.12 | 2.04E-03 | Nucleotide (cGMP) |  |
| GUCY1B3 | 1.38 | 2.55E-02 | Nucleotide (cGMP) |  |
| ATP6V1B1 | 0.62 | 4.32E-02 | Proton Transport |  |
| ATP6V0E2 | 0.38 | 3.43E-02 | Proton Transport |  |
| DHRS3 | 0.71 | 4.00E-02 | Redox |  |
| DHRS11 | 0.69 | 2.37E-02 | Redox |  |
| GFOD1 | 1.45 | 3.79E-02 | Sugar |  |
| IP6K3 | 1.68 | 2.38E-02 | Sugar (Inositol Phosphate) |  |
| INPP4A | 1.37 | 2.64E-03 | Sugar (Inositol Phosphate) |  |
| ITPKA | 0.49 | 1.59E-02 | Sugar (Inositol Phosphate) |  |
| PPA1 | 0.74 | 1.06E-02 | Sugar (PPi hydrolysis) |  |
| DCXR | 0.76 | 3.60E-02 | Sugar (Xyulose) |  |
| SULT1C4 | 1.38 | 4.38E-02 | Sulfate |  |
| ARSI | 0.73 | 2.92E-02 | Sulfate |  |
| PHOSPHO2 | 1.48 | 4.80E-04 | Vitamin (B6) |  |
| AKR7A2 | 0.70 | 1.22E-02 | Vitamine (Retinoic acid, Alcohol) |  |
| AKR7A3 | 0.26 | 3.96E-02 | Vitamine (Retinoic acid, Alcohol) |  |

**Table S12.** **Differentially expressed metabolic genes in macula between retinas and RPE/choroid.**

| **Gene symbol** | **FC** | **P value** | **Classification (Sub-category/substrate)** |
| --- | --- | --- | --- |
| ABCB11 | 513.83 | 1.49E-02 | ABC Transporter |
| ABCG4 | 85.89 | 1.93E-05 | ABC Transporter |
| ABCB4 | 42.52 | 1.97E-03 | ABC Transporter |
| ABCC8 | 16.89 | 1.38E-03 | ABC Transporter |
| ABCD2 | 13.22 | 7.65E-05 | ABC Transporter |
| ABCB9 | 8.42 | 4.00E-04 | ABC Transporter |
| ABCA3 | 7.60 | 1.22E-04 | ABC Transporter |
| ABCA7 | 5.13 | 8.53E-04 | ABC Transporter |
| ABCA4 | 4.75 | 4.83E-03 | ABC Transporter |
| ABCG1 | 3.22 | 3.96E-03 | ABC Transporter |
| ABCG5 | 2.27 | 3.83E-02 | ABC Transporter |
| ABCB6 | 1.93 | 1.30E-03 | ABC Transporter |
| ABCB8 | 1.91 | 4.81E-04 | ABC Transporter |
| ABCF3 | 1.39 | 7.92E-05 | ABC Transporter |
| ABCC9 | 0.75 | 3.59E-02 | ABC Transporter |
| ABCC1 | 0.46 | 5.82E-03 | ABC Transporter |
| ABCD3 | 0.41 | 9.14E-03 | ABC Transporter |
| ABCD1 | 0.27 | 1.20E-02 | ABC Transporter |
| ABCG2 | 0.25 | 8.31E-03 | ABC Transporter |
| ABCA10 | 0.25 | 4.85E-04 | ABC Transporter |
| ABCA5 | 0.25 | 4.85E-04 | ABC Transporter |
| ABCA6 | 0.25 | 4.85E-04 | ABC Transporter |
| ABCA8 | 0.25 | 4.85E-04 | ABC Transporter |
| ABCA9 | 0.25 | 4.85E-04 | ABC Transporter |
| ABCA1 | 0.15 | 3.79E-03 | ABC Transporter |
| ABCB1 | 0.12 | 3.04E-03 | ABC Transporter |
| ABCC11 | 0.02 | 1.15E-02 | ABC Transporter |
| ABCB5 | 1.06E-04 | 8.12E-04 | ABC Transporter |
| SMS | 1.60 | 2.24E-03 | Amino Acid (Polyamine) |
| NAT8L | 14.40 | 1.03E-02 | Amino Acid (Acetylation) |
| NAT14 | 2.45 | 1.17E-03 | Amino Acid (Acetylation) |
| NAT8 | 2.44 | 3.91E-02 | Amino Acid (Acetylation) |
| DAO | 145.85 | 9.27E-03 | Amino Acid (Alanine) |
| ADC | 1.98 | 2.48E-05 | Amino Acid (Arginine) |
| DDAH1 | 5.40 | 2.40E-04 | Amino Acid (Arginine, Citrulline) |
| OTC | 6.02 | 1.60E-02 | Amino Acid (Arginine, Urea) |
| ARG2 | 2.58 | 1.41E-03 | Amino Acid (Arginine, Urea) |
| ASNS | 3.12 | 1.92E-04 | Amino Acid (Asparagine) |
| ASRGL1 | 1.56 | 1.13E-02 | Amino Acid (Asparagine) |
| ASNSD1 | 1.31 | 2.03E-02 | Amino Acid (Asparagine) |
| AUH | 1.86 | 4.32E-05 | Amino Acid (BCAAs) |
| BCKDHA | 1.62 | 2.64E-03 | Amino Acid (BCAAs) |
| QDPR | 2.57 | 1.06E-02 | Amino Acid (BH4) |
| PTS | 1.87 | 8.71E-03 | Amino Acid (BH4) |
| PCBD2 | 1.73 | 4.85E-03 | Amino Acid (BH4) |
| BBOX1 | 2.51 | 8.75E-03 | Amino Acid (Cofactor) |
| MOCS3 | 1.57 | 2.86E-03 | Amino Acid (Cofactor) |
| CKM | 228.58 | 8.38E-03 | Amino Acid (Creatine) |
| CKMT1B | 113.92 | 1.59E-04 | Amino Acid (Creatine) |
| CKMT1A | 101.31 | 8.47E-05 | Amino Acid (Creatine) |
| CKB | 26.71 | 1.01E-06 | Amino Acid (Creatine) |
| GAMT | 2.47 | 1.78E-03 | Amino Acid (Creatine) |
| CRYM | 17.43 | 1.00E-02 | Amino Acid (Cystathionine) |
| PCYOX1L | 2.94 | 1.99E-04 | Amino Acid (Cysteine) |
| NFS1 | 1.92 | 1.54E-05 | Amino Acid (Cysteine) |
| ADO | 1.89 | 4.85E-03 | Amino Acid (Cysteine) |
| ENOPH1 | 1.68 | 2.11E-03 | Amino Acid (Cysteine, Methione) |
| GAD2 | 821.30 | 1.39E-05 | Amino Acid (Glutamate) |
| GAD1 | 127.65 | 3.26E-06 | Amino Acid (Glutamate) |
| GLS2 | 7.51 | 8.14E-05 | Amino Acid (Glutamate) |
| GLS | 3.04 | 9.73E-06 | Amino Acid (Glutamate) |
| ALDH5A1 | 2.48 | 1.12E-03 | Amino Acid (Glutamate) |
| GPT2 | 3.02 | 7.02E-05 | Amino Acid (Glutamate, Alaline) |
| GPT | 2.61 | 2.32E-02 | Amino Acid (Glutamate, Alaline) |
| HAGHL | 2.96 | 2.17E-02 | Amino Acid (Glutathione) |
| HAGH | 1.60 | 3.83E-03 | Amino Acid (Glutathione) |
| MGST3 | 1.54 | 6.87E-03 | Amino Acid (Glutathione) |
| GGT7 | 2.66 | 4.55E-04 | Amino Acid (Glutathione, Taurine, Hypotaurine) |
| GLYATL1 | 287.59 | 1.79E-05 | Amino Acid (Glycine) |
| GLYAT | 23.80 | 9.02E-03 | Amino Acid (Glycine) |
| GLYATL2 | 19.72 | 1.48E-05 | Amino Acid (Glycine) |
| FTCD | 10.43 | 3.95E-02 | Amino Acid (histidine) |
| DPEP1 | 24.64 | 6.16E-03 | Amino Acid (Hydrolysis) |
| ACY1 | 1.72 | 2.00E-04 | Amino Acid (Hydrolysis) |
| PIPOX | 9.84 | 7.26E-05 | Amino Acid (Lysine) |
| AASDHPPT | 1.65 | 1.98E-04 | Amino Acid (Lysine) |
| AASDH | 1.44 | 3.75E-02 | Amino Acid (Lysine) |
| AMD1 | 1.46 | 4.58E-03 | Amino Acid (Methionion, Polyamine) |
| TPH2 | 23.50 | 3.28E-02 | Amino Acid (Neurotransmitter) |
| TH | 3.99 | 1.42E-02 | Amino Acid (Neurotransmitter) |
| TPH1 | 3.17 | 4.21E-03 | Amino Acid (Neurotransmitter) |
| SRR | 1.64 | 4.44E-02 | Amino Acid (Neurotransmitter) |
| PRODH2 | 56.07 | 4.53E-02 | Amino Acid (Proline) |
| ACYP2 | 2.09 | 8.79E-05 | Amino Acid (Pyruvate) |
| ACYP1 | 1.67 | 9.43E-03 | Amino Acid (Pyruvate) |
| THNSL1 | 2.33 | 2.81E-05 | Amino Acid (Threonine) |
| PAH | 7.95 | 5.28E-03 | Amino Acid (Tyrosine) |
| TAT | 2.92 | 9.61E-03 | Amino Acid (Tyrosine) |
| SAT1 | 0.10 | 6.92E-03 | Amino Acid (Polyamine) |
| NAT1 | 0.45 | 3.50E-02 | Amino Acid (Acetylation) |
| AGPS | 0.68 | 3.14E-03 | Amino Acid (Alanine) |
| AGXT2L2 | 0.29 | 1.12E-04 | Amino Acid (Alanine) |
| NOS3 | 0.09 | 7.35E-03 | Amino Acid (Arginine) |
| ASL | 0.62 | 3.03E-04 | Amino Acid (Arginine ,Urea) |
| ASS1 | 0.12 | 9.47E-04 | Amino Acid (Arginine, Urea) |
| NIT2 | 0.50 | 2.86E-04 | Amino Acid (Asparagine) |
| ASPG | 0.10 | 4.89E-02 | Amino Acid (Asparagine) |
| ASPA | 0.05 | 2.65E-05 | Amino Acid (Aspartate) |
| BCKDHB | 0.72 | 4.38E-02 | Amino Acid (BCAAs) |
| HIBADH | 0.70 | 2.29E-02 | Amino Acid (BCAAs) |
| DBT | 0.59 | 1.36E-02 | Amino Acid (BCAAs) |
| IVD | 0.51 | 1.40E-05 | Amino Acid (BCAAs) |
| MCCC2 | 0.50 | 2.58E-03 | Amino Acid (BCAAs) |
| BCAT2 | 0.42 | 8.82E-04 | Amino Acid (BCAAs) |
| SPR | 0.37 | 2.12E-03 | Amino Acid (BH4) |
| GCH1 | 0.05 | 7.15E-03 | Amino Acid (BH4) |
| MOCS1 | 0.18 | 1.36E-04 | Amino Acid (Cofactor) |
| CCBL2 | 0.62 | 4.23E-03 | Amino Acid (Cysteine) |
| CSAD | 0.60 | 2.69E-02 | Amino Acid (Cysteine) |
| CBS | 0.56 | 4.43E-02 | Amino Acid (Cysteine) |
| CMBL | 0.23 | 1.17E-02 | Amino Acid (Cysteine) |
| CDO1 | 0.14 | 3.11E-02 | Amino Acid (Cysteine) |
| AOC2 | 0.07 | 7.01E-05 | Amino acid (Dopamine, histamine) |
| AOC3 | 0.07 | 7.01E-05 | Amino acid (Dopamine, histamine) |
| ALDH9A1 | 0.45 | 2.65E-02 | Amino acid (GABA) |
| ALDH18A1 | 0.55 | 4.62E-02 | Amino Acid (Glutamate) |
| ADHFE1 | 0.33 | 1.60E-02 | Amino Acid (Glutamate) |
| GLUD2 | 0.20 | 8.40E-03 | Amino Acid (Glutamate) |
| GPX4 | 0.56 | 3.03E-03 | Amino Acid (Glutathione) |
| GSTP1 | 0.49 | 2.34E-03 | Amino Acid (Glutathione) |
| GSTM3 | 0.48 | 4.95E-02 | Amino Acid (Glutathione) |
| GSTO1 | 0.34 | 9.09E-04 | Amino Acid (Glutathione) |
| GSTM2 | 0.33 | 4.10E-02 | Amino Acid (Glutathione) |
| GSTK1 | 0.19 | 1.20E-04 | Amino Acid (Glutathione) |
| MGST2 | 0.16 | 1.71E-04 | Amino Acid (Glutathione) |
| GPX7 | 0.12 | 7.25E-05 | Amino Acid (Glutathione) |
| GPX1 | 0.06 | 1.32E-03 | Amino Acid (Glutathione) |
| GPX8 | 0.02 | 5.36E-04 | Amino Acid (Glutathione) |
| MGST1 | 0.02 | 3.94E-02 | Amino Acid (Glutathione) |
| GSTA3 | 2.33E-03 | 1.43E-04 | Amino Acid (Glutathione) |
| GGT1 | 0.37 | 1.77E-03 | Amino Acid (Glutathione, Taurine, Hypotaurine) |
| GGT5 | 0.01 | 2.55E-03 | Amino Acid (Glutathione, Taurine, Hypotaurine) |
| SARDH | 0.43 | 2.64E-02 | Amino Acid (Glycine) |
| GCSH | 0.38 | 1.13E-02 | Amino Acid (Glycine) |
| DMGDH | 0.19 | 1.17E-02 | Amino Acid (Glycine) |
| BHMT | 0.12 | 1.06E-03 | Amino Acid (Glycine) |
| BHMT2 | 0.03 | 1.36E-04 | Amino Acid (Glycine) |
| AMDHD2 | 0.55 | 4.92E-03 | Amino Acid (histidine) |
| HNMT | 0.07 | 1.02E-02 | Amino Acid (histidine) |
| HAL | 0.02 | 2.53E-02 | Amino Acid (histidine) |
| APEH | 0.70 | 6.68E-04 | Amino Acid (Hydrolysis) |
| DPEP2 | 0.07 | 1.87E-02 | Amino Acid (Hydrolysis) |
| CNDP2 | 0.66 | 1.65E-02 | Amino Acid (Hydroylysis) |
| AASS | 0.20 | 2.61E-05 | Amino Acid (Lysine) |
| GCDH | 0.51 | 1.02E-04 | Amino Acid (Lysine, Tryptophan) |
| GLTP | 0.33 | 4.43E-05 | Amino Acid (Membrane lipid) |
| MTR | 0.64 | 1.34E-02 | Amino Acid (Methionine) |
| ADI1 | 0.29 | 5.43E-04 | Amino Acid (Methionine) |
| MSRA | 0.28 | 9.91E-05 | Amino Acid (Methionine) |
| MSRB3 | 0.27 | 5.84E-06 | Amino Acid (Methionine) |
| MTAP | 0.41 | 6.22E-05 | Amino Acid (Methionion, Polyamine) |
| MAOA | 0.43 | 9.08E-03 | Amino Acid (Neurotransmiter) |
| PNMT | 0.26 | 2.94E-02 | Amino Acid (Neurotransmitter) |
| ALDH1L1 | 0.24 | 2.64E-02 | Amino Acid (One carbon) |
| ALDH1L2 | 0.22 | 4.52E-02 | Amino Acid (One carbon) |
| PLOD1 | 0.54 | 4.90E-02 | Amino Acid (Proline) |
| P4HA1 | 0.44 | 1.60E-02 | Amino Acid (Proline) |
| LEPREL4 | 0.38 | 1.39E-03 | Amino Acid (Proline) |
| P4HB | 0.38 | 1.66E-02 | Amino Acid (Proline) |
| PRODH | 0.12 | 3.60E-02 | Amino Acid (Proline) |
| P4HA3 | 0.01 | 2.61E-02 | Amino Acid (Proline) |
| PHGDH | 0.19 | 4.29E-03 | Amino Acid (Serine) |
| PSAT | 0.16 | 2.27E-02 | Amino Acid (Serine) |
| THNSL2 | 0.58 | 8.27E-03 | Amino Acid (Threonine) |
| DCT | 0.01 | 3.34E-03 | Amino Acid (Tryosine, Pigment) |
| TYRP1 | 0.01 | 1.06E-04 | Amino Acid (Tryosine, Pigment) |
| TYR | 1.64E-03 | 8.54E-03 | Amino Acid (Tryosine, Pigment) |
| AFMID | 0.26 | 1.69E-03 | Amino Acid (Tryptophan) |
| KMO | 0.07 | 7.32E-05 | Amino Acid (Tryptophan) |
| INMT | 0.02 | 2.64E-02 | Amino Acid (Tryptophan) |
| TDO2 | 0.01 | 6.67E-04 | Amino Acid (Tryptophan) |
| IDO2 | 0.06 | 4.39E-02 | Amino Acid (Tryptophan, NAD)) |
| HAAO | 0.04 | 2.54E-02 | Amino Acid (Tryptophan, NAD)) |
| FAHD2B | 0.60 | 1.32E-02 | Amino Acid (Tyrosine) |
| FAHD2A | 0.42 | 2.46E-03 | Amino Acid (Tyrosine) |
| FAH | 0.18 | 8.34E-04 | Amino Acid (Tyrosine) |
| HPD | 0.02 | 4.00E-02 | Amino Acid (Tyrosine) |
| ACOT7 | 10.09 | 4.10E-04 | CoA |
| PANK4 | 2.28 | 3.13E-04 | CoA |
| ACOT8 | 1.58 | 1.87E-04 | CoA |
| PPCDC | 0.72 | 5.96E-03 | CoA |
| PPCS | 0.53 | 1.01E-04 | CoA |
| ACOT9 | 0.34 | 2.80E-04 | CoA |
| ARSF | 51.48 | 7.04E-03 | Cofactor (Calcium) |
| UGT3A1 | 143.88 | 3.82E-02 | Detox |
| DHDH | 7.38 | 4.82E-03 | Detox |
| AKR7L | 2.28 | 1.06E-02 | Detox |
| BCHE | 2.27 | 2.11E-02 | Detox |
| CES2 | 1.42 | 2.07E-02 | Detox |
| TPMT | 1.37 | 1.07E-02 | Detox |
| SOD1 | 1.30 | 3.49E-02 | Detox |
| ALDH7A1 | 0.44 | 2.01E-02 | Detox |
| ALDH1B1 | 0.41 | 8.00E-04 | Detox |
| CBR1 | 0.37 | 1.08E-04 | Detox |
| PON2 | 0.23 | 1.63E-03 | Detox |
| CAT | 0.19 | 2.67E-04 | Detox |
| EPHX1 | 0.18 | 1.51E-03 | Detox |
| CBR3 | 0.18 | 1.20E-03 | Detox |
| FMO5 | 0.15 | 1.81E-03 | Detox |
| FMO4 | 0.09 | 5.77E-04 | Detox |
| PON3 | 0.09 | 2.45E-02 | Detox |
| CES1 | 0.08 | 3.16E-02 | Detox |
| SOD3 | 0.07 | 1.20E-02 | Detox |
| FMO3 | 0.01 | 7.41E-05 | Detox |
| AADACL4 | 0.01 | 1.17E-03 | Detox |
| COX10 | 2.56 | 1.74E-04 | ETC (Complex IV) |
| SDHA | 2.35 | 4.42E-05 | ETC (Complex II) |
| COX5A | 2.19 | 5.10E-05 | ETC (Complex IV) |
| UQCRH | 2.12 | 2.11E-03 | ETC (Complex III) |
| CYCS | 1.97 | 3.88E-04 | ETC (Complex III) |
| NDUFV1 | 1.95 | 4.64E-04 | ETC (Complex I) |
| COX6A1 | 1.83 | 2.63E-04 | ETC (Complex IV) |
| NDUFS1 | 1.82 | 1.11E-02 | ETC (Complex I) |
| UQCRC1 | 1.78 | 1.26E-05 | ETC (Complex III) |
| NDUFA4 | 1.77 | 3.14E-03 | ETC (Complex I) |
| COX7A2 | 1.76 | 2.67E-04 | ETC (Complex IV) |
| NDUFA6 | 1.73 | 7.53E-05 | ETC (Complex I) |
| UQCRC2 | 1.72 | 2.34E-04 | ETC (Complex III) |
| NDUFA5 | 1.71 | 4.98E-04 | ETC (Complex I) |
| NDUFV2 | 1.63 | 1.11E-03 | ETC (Complex I) |
| NDUFA7 | 1.61 | 1.72E-03 | ETC (Complex I) |
| CYC1 | 1.57 | 9.33E-03 | ETC (Complex III) |
| NDUFB9 | 1.55 | 1.50E-03 | ETC (Complex I) |
| COX11 | 1.53 | 1.84E-05 | ETC (Complex IV) |
| NDUFAB1 | 1.53 | 1.75E-03 | ETC (Complex I) |
| COX5B | 1.52 | 1.18E-03 | ETC (Complex IV) |
| NDUFS2 | 1.50 | 8.20E-03 | ETC (Complex I) |
| UQCRFS1 | 1.49 | 7.44E-04 | ETC (Complex III) |
| NDUFA9 | 1.47 | 4.37E-03 | ETC (Complex I) |
| COX7A2L | 1.43 | 1.79E-04 | ETC (Complex IV) |
| NDUFA8 | 1.40 | 4.51E-03 | ETC (Complex I) |
| NDUFA10 | 1.37 | 4.13E-02 | ETC (Complex I) |
| NDUFB3 | 1.32 | 1.78E-02 | ETC (Complex I) |
| COX17 | 1.30 | 1.26E-02 | ETC (Complex IV) |
| COX6B2 | 0.46 | 3.61E-02 | ETC (Complex IV) |
| SDHD | 0.41 | 4.59E-03 | ETC (Complex II) |
| FUT9 | 677.59 | 6.66E-06 | Glycan |
| FUT3 | 264.82 | 1.35E-02 | Glycan |
| ST6GAL2 | 232.31 | 6.78E-04 | Glycan |
| ST8SIA3 | 130.85 | 2.28E-05 | Glycan |
| GALNT13 | 100.62 | 1.88E-05 | Glycan |
| GAL3ST3 | 96.88 | 9.08E-07 | Glycan (Sulfate) |
| FUT5 | 78.99 | 3.06E-02 | Glycan |
| B3GAT1 | 50.05 | 1.06E-04 | Glycan |
| ST8SIA1 | 48.81 | 4.93E-04 | Glycan |
| ST8SIA2 | 43.75 | 2.36E-03 | Glycan |
| GLB1L2 | 38.36 | 8.75E-04 | Glycan (Degradation) |
| CHST8 | 25.24 | 6.86E-03 | Glycan (Sulfate) |
| GAL3ST1 | 17.14 | 1.70E-03 | Glycan (Sulfate) |
| MANEAL | 16.53 | 1.09E-05 | Glycan (Degradation) |
| GALNTL5 | 16.52 | 6.64E-03 | Glycan |
| MGAT4C | 16.32 | 7.49E-04 | Glycan |
| ST8SIA5 | 14.26 | 5.78E-04 | Glycan |
| B3GALT1 | 12.81 | 5.67E-05 | Glycan |
| HAS1 | 11.89 | 1.62E-02 | Glycan |
| B3GNT4 | 9.92 | 1.26E-02 | Glycan |
| GLB1L3 | 9.62 | 3.25E-02 | Glycan (Degradation) |
| B3GALT2 | 8.97 | 3.88E-05 | Glycan |
| MGAT3 | 8.48 | 8.35E-04 | Glycan |
| B3GAT2 | 8.45 | 9.57E-05 | Glycan |
| GLT25D2 | 7.89 | 5.39E-05 | Glycan |
| B4GALNT1 | 7.72 | 3.60E-03 | Glycan |
| B4GALT6 | 7.65 | 4.83E-05 | Glycan |
| CHST5 | 7.45 | 4.39E-05 | Glycan (Sulfate) |
| GPLD1 | 6.24 | 1.25E-05 | Glycan (Anchor) |
| CHST2 | 5.09 | 3.68E-04 | Glycan (Sulfate) |
| GLT1D1 | 4.71 | 1.34E-03 | Glycan |
| GAL3ST2 | 3.93 | 2.99E-02 | Glycan (Sulfate) |
| WBSCR17 | 3.80 | 4.90E-03 | Glycan |
| GBA2 | 3.43 | 3.63E-05 | Glycan |
| PIGZ | 3.43 | 7.27E-03 | Glycan (Anchor) |
| B3GALNT1 | 3.32 | 7.73E-05 | Glycan |
| GALNT9 | 3.30 | 8.68E-03 | Glycan |
| GALNT8 | 3.21 | 3.96E-03 | Glycan |
| B4GALNT4 | 3.17 | 1.26E-03 | Glycan |
| MAN2A2 | 2.94 | 9.82E-04 | Glycan (Degradation) |
| MGAT5 | 2.79 | 2.71E-03 | Glycan |
| GBA3 | 2.76 | 5.52E-03 | Glycan |
| ALG10B | 2.76 | 6.20E-05 | Glycan |
| MGAT5B | 2.75 | 4.50E-02 | Glycan |
| LARGE | 2.67 | 2.14E-05 | Glycan |
| B3GNT1 | 2.59 | 3.22E-04 | Glycan |
| ST3GAL3 | 2.33 | 7.66E-06 | Glycan |
| ALG10 | 2.24 | 1.71E-03 | Glycan |
| IDS | 2.10 | 6.37E-03 | Glycan (Degradation) |
| POMT2 | 2.07 | 9.83E-04 | Glycan |
| GALNTL6 | 1.93 | 3.57E-02 | Glycan |
| B4GALT2 | 1.83 | 2.92E-03 | Glycan |
| DHDDS | 1.77 | 3.21E-03 | Glycan |
| XYLT1 | 1.73 | 1.40E-03 | Glycan |
| ST6GALNAC5 | 1.70 | 2.20E-02 | Glycan |
| PIGM | 1.52 | 5.64E-03 | Glycan (Anchor) |
| POMT1 | 1.50 | 1.34E-02 | Glycan |
| B3GALTL | 1.49 | 4.47E-02 | Glycan |
| PIGS | 1.44 | 2.58E-03 | Glycan (Anchor) |
| FN3K | 1.40 | 2.89E-02 | Glycan |
| FN3KRP | 1.38 | 4.00E-03 | Glycan |
| PIGX | 1.34 | 3.41E-03 | Glycan (Anchor) |
| CHST10 | 1.33 | 5.43E-03 | Glycan (Sulfate) |
| GLB1 | 0.74 | 2.36E-02 | Glycan (Degradation) |
| MANBAL | 0.71 | 2.24E-03 | Glycan (Degradation) |
| GTDC1 | 0.70 | 2.11E-02 | Glycan |
| PIGG | 0.69 | 3.93E-02 | Glycan (Anchor) |
| PIGA | 0.69 | 2.96E-02 | Glycan (Anchor) |
| PIGB | 0.67 | 1.82E-03 | Glycan (Anchor) |
| MGAT4B | 0.65 | 4.58E-02 | Glycan |
| NEU3 | 0.65 | 3.94E-02 | Glycan (Degradation) |
| PIGV | 0.64 | 1.42E-02 | Glycan (Anchor) |
| MGAT2 | 0.64 | 2.85E-02 | Glycan |
| GLT8D1 | 0.64 | 1.80E-03 | Glycan |
| NEU1 | 0.62 | 3.34E-02 | Glycan (Degradation) |
| LYG1 | 0.61 | 1.66E-02 | Glycan (Degradation) |
| ALG14 | 0.61 | 2.17E-02 | Glycan |
| ALG13 | 0.60 | 4.99E-03 | Glycan |
| B3GNT2 | 0.59 | 1.66E-03 | Glycan |
| ALG6 | 0.58 | 1.03E-02 | Glycan |
| ST3GAL2 | 0.57 | 4.80E-02 | Glycan |
| GBA | 0.56 | 2.94E-04 | Glycan |
| PIGT | 0.56 | 1.46E-03 | Glycan (Anchor) |
| ALG2 | 0.55 | 1.21E-02 | Glycan |
| B4GALT3 | 0.55 | 2.44E-03 | Glycan |
| MGAT1 | 0.53 | 5.28E-03 | Glycan |
| GALNT1 | 0.53 | 2.66E-02 | Glycan |
| FUT8 | 0.52 | 1.05E-02 | Glycan |
| MAN2B2 | 0.51 | 7.50E-03 | Glycan (Degradation) |
| ALG12 | 0.51 | 1.39E-02 | Glycan |
| MANBA | 0.51 | 1.14E-03 | Glycan (Degradation) |
| EDEM2 | 0.50 | 4.78E-03 | Glycan (Degradation) |
| MGAT4A | 0.50 | 8.68E-03 | Glycan |
| FUT1 | 0.49 | 1.50E-02 | Glycan |
| GLA | 0.48 | 3.05E-03 | Glycan (Degradation) |
| ALG8 | 0.46 | 1.03E-02 | Glycan |
| NAGPA | 0.46 | 1.07E-02 | Glycan |
| SULF2 | 0.45 | 2.24E-03 | Glycan (Sulfate) |
| ST6GAL1 | 0.44 | 1.16E-02 | Glycan |
| IDUA | 0.44 | 1.23E-02 | Glycan (Degradation) |
| CSGALNACT1 | 0.43 | 4.26E-03 | Glycan (Sulfate) |
| GUSB | 0.43 | 1.28E-02 | Glycan (Degradation) |
| B4GALT4 | 0.43 | 2.47E-03 | Glycan |
| ALG5 | 0.42 | 2.19E-04 | Glycan |
| NAGA | 0.41 | 1.16E-04 | Glycan |
| DAD1 | 0.36 | 7.95E-03 | Glycan |
| HEXB | 0.32 | 1.94E-07 | Glycan (Degradation) |
| POFUT1 | 0.32 | 1.52E-04 | Glycan |
| GCNT2 | 0.31 | 5.97E-05 | Glycan |
| GLT25D1 | 0.31 | 2.22E-03 | Glycan |
| B3GNT9 | 0.30 | 1.04E-02 | Glycan |
| CSGALNACT2 | 0.30 | 3.04E-03 | Glycan (Sulfate) |
| GNS | 0.29 | 1.18E-03 | Glycan (Degradation) |
| AGA | 0.29 | 3.80E-02 | Glycan (Degradation) |
| SULF1 | 0.29 | 1.04E-02 | Glycan (Sulfate) |
| HEXA | 0.29 | 2.17E-05 | Glycan (Degradation) |
| ST3GAL6 | 0.27 | 3.04E-03 | Glycan |
| SGSH | 0.27 | 1.75E-03 | Glycan (Sulfate) |
| CHSY1 | 0.27 | 4.78E-03 | Glycan (Sulfate) |
| GCNT4 | 0.26 | 4.90E-03 | Glycan |
| B3GNT5 | 0.25 | 3.94E-04 | Glycan |
| NAGLU | 0.24 | 6.07E-05 | Glycan (Degradation) |
| B4GALT1 | 0.24 | 1.02E-02 | Glycan |
| GLB1L | 0.23 | 2.98E-04 | Glycan (Degradation) |
| ST6GALNAC3 | 0.21 | 2.10E-04 | Glycan |
| FUCA1 | 0.19 | 4.03E-03 | Glycan (Degradation) |
| CTBS | 0.19 | 5.37E-04 | Glycan |
| GALNT10 | 0.18 | 4.62E-02 | Glycan |
| CHST3 | 0.18 | 1.07E-04 | Glycan (Sulfate) |
| FUT7 | 0.18 | 3.03E-02 | Glycan |
| MAN2B1 | 0.17 | 1.67E-04 | Glycan (Degradation) |
| ST6GALNAC4 | 0.17 | 3.98E-02 | Glycan |
| MAN1A1 | 0.16 | 1.38E-04 | Glycan (Degradation) |
| FUCA2 | 0.15 | 8.77E-05 | Glycan (Degradation) |
| HYAL1 | 0.14 | 5.90E-04 | Glycan (Degradation) |
| ST3GAL4 | 0.13 | 5.25E-04 | Glycan |
| MFNG | 0.13 | 9.92E-04 | Glycan |
| GAL3ST4 | 0.12 | 9.33E-04 | Glycan (Sulfate) |
| CHST14 | 0.12 | 1.03E-03 | Glycan (Sulfate) |
| CHST11 | 0.09 | 1.16E-06 | Glycan (Sulfate) |
| HYAL2 | 0.09 | 2.54E-03 | Glycan (Degradation) |
| CHSY3 | 0.08 | 6.31E-04 | Glycan (Sulfate) |
| GBGT1 | 0.08 | 3.43E-05 | Glycan |
| MAN2A1 | 0.08 | 1.33E-02 | Glycan (Degradation) |
| GCNT1 | 0.07 | 3.32E-04 | Glycan |
| GALNT6 | 0.06 | 9.04E-06 | Glycan |
| LYZ | 0.05 | 5.36E-03 | Glycan (Degradation) |
| ST8SIA6 | 0.05 | 4.45E-03 | Glycan |
| CHST13 | 0.04 | 2.47E-03 | Glycan (Sulfate) |
| A4GALT | 0.03 | 1.36E-02 | Glycan |
| GLT8D2 | 0.02 | 9.41E-04 | Glycan |
| GALNT5 | 0.02 | 2.21E-02 | Glycan |
| HKDC1 | 147.25 | 2.49E-03 | Glycolysis |
| ENO2 | 37.36 | 3.81E-07 | Glycolysis |
| PFKP | 14.84 | 6.32E-04 | Glycolysis |
| HK2 | 14.33 | 4.76E-03 | Glycolysis |
| ALDOC | 14.10 | 4.11E-07 | Glycolysis |
| PGAM4 | 5.02 | 2.28E-03 | Glycolysis |
| GPI | 4.54 | 5.24E-04 | Glycolysis |
| PFKFB4 | 3.71 | 4.36E-03 | Glycolysis |
| PGK1 | 3.64 | 5.27E-04 | Glycolysis |
| GCK | 3.45 | 2.34E-04 | Glycolysis |
| TPI1 | 3.42 | 2.34E-04 | Glycolysis |
| PGAM1 | 3.42 | 2.26E-05 | Glycolysis |
| GAPDH | 3.32 | 1.55E-04 | Glycolysis |
| PFKM | 3.28 | 8.66E-07 | Glycolysis |
| PGAM2 | 2.74 | 4.36E-03 | Glycolysis |
| HK1 | 2.60 | 3.34E-05 | Glycolysis |
| LDHA | 2.60 | 2.92E-03 | Glycolysis |
| PFKFB2 | 2.37 | 2.17E-02 | Glycolysis |
| PGM2 | 2.28 | 2.16E-03 | Glycolysis |
| ENO3 | 2.27 | 4.42E-03 | Glycolysis |
| ENO1 | 2.08 | 7.46E-03 | Glycolysis |
| PKLR | 1.98 | 4.75E-02 | Glycolysis |
| ALDOA | 1.86 | 3.72E-03 | Glycolysis |
| PGM2L1 | 1.86 | 2.38E-02 | Glycolysis |
| PGM1 | 1.50 | 1.69E-02 | Glycolysis |
| DLAT | 1.43 | 6.00E-03 | Glycolysis |
| ADPGK | 0.34 | 3.94E-04 | Glycolysis |
| PGM5 | 0.32 | 9.49E-03 | Glycolysis |
| PFKFB3 | 0.21 | 7.95E-03 | Glycolysis |
| HK3 | 0.01 | 2.43E-02 | Glycolysis |
| GAPDHS | 0.01 | 1.94E-03 | Glycolysis |
| GLOD5 | 451.86 | 3.52E-04 | Glyoxylate |
| GLO1 | 1.35 | 7.50E-03 | Glyoxylate |
| HS3ST4 | 75.68 | 1.11E-04 | Heparin Sulfate |
| NDST3 | 26.31 | 3.55E-06 | Heparin Sulfate |
| HS6ST3 | 23.43 | 5.64E-04 | Heparin Sulfate |
| HS3ST3B1 | 7.81 | 6.32E-04 | Heparin Sulfate |
| HS6ST2 | 7.45 | 6.59E-04 | Heparin Sulfate |
| HS3ST3A1 | 4.50 | 2.68E-02 | Heparin Sulfate |
| HS6ST1 | 3.97 | 2.41E-05 | Heparin Sulfate |
| HS3ST1 | 2.61 | 9.66E-03 | Heparin Sulfate |
| NDST1 | 1.80 | 4.67E-03 | Heparin Sulfate |
| HGSNAT | 0.71 | 9.86E-03 | Heparin Sulfate |
| EXT1 | 0.23 | 3.88E-04 | Heparin Sulfate |
| HPSE2 | 0.19 | 3.24E-02 | Heparin Sulfate |
| HPSE | 0.18 | 3.63E-02 | Heparin Sulfate |
| CACNG3 | 7448.60 | 9.06E-05 | Ion Transport |
| TRPC7 | 3359.70 | 8.65E-06 | Ion Transport |
| CHRNB3 | 2205.30 | 1.37E-04 | Ion Transport |
| KCNC2 | 1579.80 | 4.64E-03 | Ion Transport |
| CACNG7 | 1129.00 | 2.32E-03 | Ion Transport |
| SCN1A | 1107.00 | 1.99E-04 | Ion Transport |
| CACNG5 | 904.39 | 4.11E-06 | Ion Transport |
| GRIN1 | 870.67 | 2.40E-03 | Ion Transport |
| GLRA1 | 786.97 | 3.15E-05 | Ion Transport |
| KCNC1 | 735.40 | 8.88E-04 | Ion Transport |
| GABRA1 | 707.15 | 1.81E-05 | Ion Transport |
| KCNJ9 | 510.61 | 9.48E-04 | Ion Transport |
| CHRNA2 | 475.48 | 8.02E-04 | Ion Transport |
| GABRR3 | 469.74 | 7.11E-04 | Ion Transport |
| GLRA2 | 452.21 | 9.34E-05 | Ion Transport |
| KCNB2 | 396.82 | 5.85E-05 | Ion Transport |
| CHRNA6 | 337.45 | 1.04E-04 | Ion Transport |
| KCNK9 | 317.52 | 9.53E-03 | Ion Transport |
| CNGA1 | 316.50 | 1.17E-03 | Ion Transport |
| CNGB1 | 309.42 | 1.83E-03 | Ion Transport |
| SLC24A2 | 275.62 | 7.19E-05 | Ion Transport |
| CACNG2 | 265.35 | 1.34E-03 | Ion Transport |
| ATP1A4 | 255.10 | 6.94E-03 | Ion Transport |
| KCNV2 | 254.26 | 1.14E-04 | Ion Transport |
| KCNH6 | 242.33 | 1.54E-06 | Ion Transport |
| GRIK1 | 234.98 | 7.95E-06 | Ion Transport |
| HCN2 | 227.30 | 3.25E-04 | Ion Transport |
| KCNG4 | 222.95 | 2.60E-04 | Ion Transport |
| TF | 217.31 | 1.37E-03 | Ion Transport |
| GABRG2 | 204.12 | 2.80E-05 | Ion Transport |
| GABRR1 | 164.11 | 1.03E-04 | Ion Transport |
| KCNJ10 | 160.53 | 4.70E-04 | Ion Transport |
| CHRNB2 | 159.18 | 2.48E-06 | Ion Transport |
| SLN | 148.05 | 3.83E-03 | Ion Transport |
| CNGA3 | 143.14 | 3.05E-05 | Ion Transport |
| KCNH7 | 143.06 | 2.22E-05 | Ion Transport |
| KCNA1 | 131.31 | 6.46E-03 | Ion Transport |
| KCNQ2 | 126.27 | 4.63E-03 | Ion Transport |
| KCNJ6 | 116.26 | 8.17E-07 | Ion Transport |
| GABRA6 | 111.46 | 6.87E-03 | Ion Transport |
| GABRA3 | 110.66 | 1.78E-08 | Ion Transport |
| HCN1 | 101.07 | 1.92E-05 | Ion Transport |
| CACNA1G | 86.55 | 1.72E-06 | Ion Transport |
| KCNJ4 | 84.27 | 1.69E-03 | Ion Transport |
| CACNA1B | 72.14 | 2.82E-04 | Ion Transport |
| GRIA4 | 71.03 | 8.66E-06 | Ion Transport |
| CACNA1E | 70.48 | 5.24E-06 | Ion Transport |
| CHRND | 69.38 | 1.59E-02 | Ion Transport |
| SCN8A | 67.82 | 2.61E-05 | Ion Transport |
| HCN4 | 60.68 | 2.38E-03 | Ion Transport |
| KCND2 | 60.32 | 1.33E-04 | Ion Transport |
| TRPC5 | 57.31 | 4.10E-02 | Ion Transport |
| GRIA1 | 53.80 | 4.56E-04 | Ion Transport |
| KCNB1 | 53.44 | 8.83E-04 | Ion Transport |
| PKD2L1 | 52.97 | 1.45E-04 | Ion Transport |
| SCN3B | 52.58 | 1.50E-06 | Ion Transport |
| CACNG8 | 51.23 | 1.01E-06 | Ion Transport |
| KCNJ16 | 45.89 | 3.54E-05 | Ion Transport |
| SLC12A5 | 45.18 | 2.05E-07 | Ion Transport |
| GABRR2 | 45.05 | 4.11E-07 | Ion Transport |
| SLC24A4 | 42.24 | 4.90E-06 | Ion Transport |
| CACNA1F | 40.97 | 6.16E-04 | Ion Transport |
| SLC39A2 | 40.05 | 5.75E-03 | Ion Transport |
| CACNA2D4 | 40.03 | 4.57E-03 | Ion Transport |
| KCNA2 | 39.88 | 1.04E-03 | Ion Transport |
| KCNH2 | 39.79 | 2.04E-04 | Ion Transport |
| GLRA3 | 38.86 | 2.84E-04 | Ion Transport |
| GABRA5 | 38.42 | 8.74E-04 | Ion Transport |
| KCNA4 | 37.79 | 3.17E-04 | Ion Transport |
| GRIN2B | 35.95 | 2.11E-04 | Ion Transport |
| SLC24A1 | 35.49 | 9.28E-04 | Ion Transport |
| CACNG4 | 33.91 | 1.35E-05 | Ion Transport |
| CACNA1I | 33.79 | 3.66E-05 | Ion Transport |
| ATP1A3 | 33.71 | 3.56E-08 | Ion Transport |
| GABRG1 | 32.08 | 2.42E-04 | Ion Transport |
| ATP2B3 | 31.95 | 3.60E-04 | Ion Transport |
| SCNN1G | 31.72 | 1.73E-02 | Ion Transport |
| CHRNA4 | 31.07 | 1.62E-03 | Ion Transport |
| KCNF1 | 30.90 | 3.01E-04 | Ion Transport |
| SLC39A5 | 30.67 | 2.64E-03 | Ion Transport |
| GABRA2 | 30.23 | 6.75E-04 | Ion Transport |
| SCN4B | 30.23 | 2.66E-03 | Ion Transport |
| KCNV1 | 28.18 | 3.53E-02 | Ion Transport |
| SLC8A2 | 27.50 | 1.76E-04 | Ion Transport |
| SCN2A | 26.77 | 8.45E-06 | Ion Transport |
| GRIA2 | 26.25 | 4.54E-05 | Ion Transport |
| TRPV6 | 24.98 | 2.52E-05 | Ion Transport |
| GRIA3 | 24.76 | 4.76E-06 | Ion Transport |
| SLC8A3 | 24.76 | 1.30E-05 | Ion Transport |
| GABRG3 | 24.40 | 4.07E-05 | Ion Transport |
| KCNJ11 | 23.48 | 4.99E-04 | Ion Transport |
| GLRA4 | 22.99 | 6.61E-03 | Ion Transport |
| GABRB3 | 22.74 | 8.01E-06 | Ion Transport |
| CACNA1A | 21.38 | 1.02E-04 | Ion Transport |
| SLC13A4 | 20.42 | 7.63E-04 | Ion Transport |
| GRIK4 | 19.08 | 2.95E-06 | Ion Transport |
| GABRB2 | 17.56 | 6.25E-05 | Ion Transport |
| TTYH1 | 17.28 | 6.73E-05 | Ion Transport |
| CACNG6 | 17.03 | 4.41E-02 | Ion Transport |
| GRIN2C | 15.87 | 3.41E-03 | Ion Transport |
| GABRD | 15.55 | 3.48E-04 | Ion Transport |
| SLC9A2 | 15.39 | 3.20E-04 | Ion Transport |
| GRIN2A | 15.20 | 5.25E-03 | Ion Transport |
| RYR2 | 15.19 | 4.26E-05 | Ion Transport |
| BEST3 | 14.67 | 2.29E-04 | Ion Transport |
| KCTD8 | 14.62 | 1.20E-09 | Ion Transport |
| SCN2B | 14.42 | 8.12E-04 | Ion Transport |
| KCNK3 | 14.01 | 3.09E-05 | Ion Transport |
| CLCNKA | 12.95 | 2.04E-03 | Ion Transport |
| SLC9A7 | 12.83 | 5.02E-05 | Ion Transport |
| KCTD16 | 11.41 | 1.23E-04 | Ion Transport |
| HTR3E | 9.67 | 2.45E-02 | Ion Transport |
| KCND3 | 9.53 | 2.51E-06 | Ion Transport |
| P2RX6 | 9.47 | 1.93E-04 | Ion Transport |
| CNGA4 | 9.30 | 4.16E-04 | Ion Transport |
| KCNJ3 | 8.83 | 4.63E-03 | Ion Transport |
| TRPC3 | 7.73 | 4.62E-04 | Ion Transport |
| KCNN1 | 7.42 | 2.94E-03 | Ion Transport |
| SLC8A1 | 7.25 | 8.04E-05 | Ion Transport |
| KCTD19 | 7.11 | 4.22E-05 | Ion Transport |
| ATP2A1 | 7.10 | 1.82E-05 | Ion Transport |
| TRPM8 | 7.06 | 5.74E-03 | Ion Transport |
| KCNC3 | 7.04 | 5.63E-04 | Ion Transport |
| GRIN2D | 6.68 | 4.93E-03 | Ion Transport |
| ATP2B1 | 6.65 | 8.64E-05 | Ion Transport |
| KCNQ3 | 6.57 | 1.24E-03 | Ion Transport |
| CATSPER2 | 6.52 | 7.40E-03 | Ion Transport |
| ATP1B2 | 6.33 | 3.47E-05 | Ion Transport |
| CACNB4 | 5.95 | 9.71E-04 | Ion Transport |
| KCNN2 | 5.94 | 2.08E-04 | Ion Transport |
| CACNB2 | 5.80 | 2.62E-04 | Ion Transport |
| SLC13A1 | 5.68 | 2.56E-03 | Ion Transport |
| KCNK4 | 5.51 | 4.88E-03 | Ion Transport |
| GRID2 | 5.47 | 2.07E-03 | Ion Transport |
| SCN5A | 5.44 | 8.30E-03 | Ion Transport |
| KCNA5 | 5.41 | 8.92E-05 | Ion Transport |
| KCNJ2 | 5.23 | 7.33E-05 | Ion Transport |
| KCTD7 | 5.08 | 3.48E-04 | Ion Transport |
| VDAC1 | 5.04 | 3.83E-05 | Ion Transport |
| SLC12A6 | 4.92 | 4.34E-05 | Ion Transport |
| CACNA2D2 | 4.91 | 1.34E-05 | Ion Transport |
| CACNB1 | 4.78 | 4.48E-04 | Ion Transport |
| GLRB | 4.69 | 2.60E-04 | Ion Transport |
| GRIK5 | 4.49 | 4.83E-06 | Ion Transport |
| P2RX3 | 4.20 | 5.13E-03 | Ion Transport |
| GRID1 | 4.13 | 3.83E-05 | Ion Transport |
| KCNJ12 | 4.02 | 6.78E-03 | Ion Transport |
| KCNK10 | 3.93 | 3.55E-02 | Ion Transport |
| ATP2B2 | 3.78 | 6.91E-03 | Ion Transport |
| SCN3A | 3.67 | 5.01E-03 | Ion Transport |
| HCN3 | 3.61 | 1.09E-03 | Ion Transport |
| CLCN4 | 3.55 | 1.60E-03 | Ion Transport |
| PKD1 | 3.27 | 9.56E-03 | Ion Transport |
| CLCN6 | 3.24 | 3.78E-05 | Ion Transport |
| KCNA6 | 3.21 | 3.96E-03 | Ion Transport |
| KCNMA1 | 3.03 | 1.51E-03 | Ion Transport |
| CHRNE | 3.01 | 3.81E-02 | Ion Transport |
| SLC24A3 | 2.95 | 2.94E-02 | Ion Transport |
| KCNC4 | 2.93 | 1.08E-03 | Ion Transport |
| ATP1B1 | 2.88 | 5.69E-03 | Ion Transport |
| GRIK3 | 2.88 | 2.99E-03 | Ion Transport |
| KCNT1 | 2.85 | 1.14E-02 | Ion Transport |
| SLC9A6 | 2.84 | 2.92E-07 | Ion Transport |
| TRPC6 | 2.81 | 3.58E-02 | Ion Transport |
| GABRB1 | 2.76 | 6.63E-03 | Ion Transport |
| KCNN4 | 2.74 | 1.98E-03 | Ion Transport |
| TRPC1 | 2.72 | 1.03E-03 | Ion Transport |
| SLC9A4 | 2.64 | 1.66E-02 | Ion Transport |
| KCTD17 | 2.62 | 2.34E-02 | Ion Transport |
| GRIN3A | 2.55 | 4.81E-02 | Ion Transport |
| ATP7B | 2.54 | 3.33E-06 | Ion Transport |
| SCNN1D | 2.52 | 3.52E-03 | Ion Transport |
| TRPM2 | 2.51 | 2.26E-02 | Ion Transport |
| CACNA2D3 | 2.49 | 2.21E-02 | Ion Transport |
| ITPR3 | 2.34 | 9.49E-04 | Ion Transport |
| KCNAB2 | 2.24 | 1.07E-02 | Ion Transport |
| VDAC3 | 2.23 | 3.60E-04 | Ion Transport |
| GRIK2 | 2.22 | 1.42E-03 | Ion Transport |
| CLCN3 | 2.20 | 2.80E-03 | Ion Transport |
| CACNA2D1 | 2.11 | 1.87E-03 | Ion Transport |
| SLC30A9 | 1.98 | 7.39E-04 | Ion Transport |
| KCTD13 | 1.95 | 6.76E-03 | Ion Transport |
| ATP11A | 1.92 | 3.64E-03 | Ion Transport |
| RYR1 | 1.83 | 1.53E-02 | Ion Transport |
| CACNB3 | 1.76 | 9.09E-03 | Ion Transport |
| CATSPER3 | 1.73 | 4.85E-03 | Ion Transport |
| SLC39A10 | 1.67 | 5.42E-03 | Ion Transport |
| VDAC2 | 1.64 | 5.61E-03 | Ion Transport |
| ATP11B | 1.59 | 1.39E-02 | Ion Transport |
| KCNAB3 | 1.57 | 1.01E-02 | Ion Transport |
| KCTD2 | 1.46 | 1.78E-02 | Ion Transport |
| MFSD5 | 1.41 | 3.89E-02 | Ion Transport |
| MFSD8 | 1.34 | 2.53E-02 | Ion Transport |
| KCTD20 | 0.76 | 3.82E-03 | Ion Transport |
| SLC30A6 | 0.69 | 1.10E-02 | Ion Transport |
| SLC41A2 | 0.64 | 4.85E-03 | Ion Transport |
| TMCO3 | 0.59 | 1.22E-02 | Ion Transport |
| TPCN1 | 0.59 | 1.21E-03 | Ion Transport |
| KCTD11 | 0.58 | 2.62E-03 | Ion Transport |
| ATP7A | 0.56 | 2.80E-03 | Ion Transport |
| SLC39A7 | 0.56 | 1.72E-03 | Ion Transport |
| SLC31A1 | 0.51 | 5.70E-03 | Ion Transport |
| KCTD18 | 0.48 | 3.28E-03 | Ion Transport |
| HEPH | 0.47 | 1.53E-02 | Ion Transport |
| KCNE3 | 0.45 | 5.04E-03 | Ion Transport |
| SLC12A9 | 0.44 | 1.34E-02 | Ion Transport |
| SLC40A1 | 0.43 | 3.70E-02 | Ion Transport |
| KCNMB4 | 0.39 | 2.28E-02 | Ion Transport |
| SLC30A7 | 0.38 | 1.37E-03 | Ion Transport |
| KCNE1 | 0.37 | 3.57E-02 | Ion Transport |
| KCNQ1 | 0.35 | 2.05E-02 | Ion Transport |
| KCTD10 | 0.35 | 9.65E-06 | Ion Transport |
| TCN2 | 0.35 | 3.84E-03 | Ion Transport |
| SLC39A11 | 0.34 | 6.40E-03 | Ion Transport |
| TRPM7 | 0.33 | 3.89E-02 | Ion Transport |
| P2RX4 | 0.33 | 2.03E-03 | Ion Transport |
| TPCN2 | 0.32 | 1.59E-03 | Ion Transport |
| TRPM4 | 0.31 | 5.26E-04 | Ion Transport |
| PKD2 | 0.31 | 4.62E-03 | Ion Transport |
| SLC31A2 | 0.29 | 1.01E-02 | Ion Transport |
| KCNN3 | 0.26 | 6.55E-03 | Ion Transport |
| TRPC4 | 0.25 | 7.84E-03 | Ion Transport |
| ATP11C | 0.25 | 8.23E-04 | Ion Transport |
| SLC3A2 | 0.24 | 5.09E-04 | Ion Transport |
| ATP1B3 | 0.24 | 4.17E-02 | Ion Transport |
| SCN11A | 0.22 | 3.41E-02 | Ion Transport |
| SCN9A | 0.20 | 1.08E-02 | Ion Transport |
| KCNJ8 | 0.19 | 2.40E-02 | Ion Transport |
| SLC39A1 | 0.19 | 1.30E-02 | Ion Transport |
| KCNQ5 | 0.19 | 3.69E-03 | Ion Transport |
| ARSD | 0.18 | 8.16E-04 | Ion Transport |
| KCNK7 | 0.18 | 2.54E-02 | Ion Transport |
| SLC9A9 | 0.18 | 1.91E-03 | Ion Transport |
| SLC12A4 | 0.18 | 9.16E-05 | Ion Transport |
| ORAI1 | 0.18 | 6.92E-03 | Ion Transport |
| CLIC4 | 0.17 | 7.52E-03 | Ion Transport |
| KCNK15 | 0.15 | 1.54E-02 | Ion Transport |
| MAGT1 | 0.14 | 5.27E-04 | Ion Transport |
| SLC12A1 | 0.13 | 2.41E-02 | Ion Transport |
| SLC12A3 | 0.13 | 4.21E-02 | Ion Transport |
| SLC24A6 | 0.13 | 2.41E-03 | Ion Transport |
| P2RX5 | 0.12 | 3.34E-03 | Ion Transport |
| P2RX7 | 0.11 | 4.09E-03 | Ion Transport |
| CFTR | 0.11 | 3.37E-02 | Ion Transport |
| CLIC2 | 0.11 | 7.46E-04 | Ion Transport |
| KCNS3 | 0.10 | 4.58E-02 | Ion Transport |
| ITPR1 | 0.10 | 6.03E-04 | Ion Transport |
| KCNK5 | 0.09 | 7.13E-05 | Ion Transport |
| P2RX1 | 0.07 | 1.55E-02 | Ion Transport |
| KCNJ15 | 0.07 | 2.40E-02 | Ion Transport |
| KCTD12 | 0.06 | 1.13E-04 | Ion Transport |
| SLC30A3 | 0.06 | 5.63E-03 | Ion Transport |
| CLIC1 | 0.06 | 6.56E-04 | Ion Transport |
| CLCA2 | 0.04 | 1.16E-02 | Ion Transport |
| ATP2A3 | 0.03 | 1.60E-04 | Ion Transport |
| CLIC3 | 0.03 | 1.39E-02 | Ion Transport |
| TRPA1 | 0.03 | 1.15E-04 | Ion Transport |
| MCOLN3 | 0.03 | 1.57E-02 | Ion Transport |
| KCNT2 | 0.02 | 1.25E-02 | Ion Transport |
| TRPV4 | 0.02 | 4.88E-02 | Ion Transport |
| KCNE4 | 0.02 | 3.67E-03 | Ion Transport |
| SLC24A5 | 0.02 | 1.60E-02 | Ion Transport |
| KCNK6 | 0.02 | 3.08E-05 | Ion Transport |
| KCNJ13 | 0.02 | 5.56E-03 | Ion Transport |
| GABRP | 0.02 | 7.22E-03 | Ion Transport |
| OCA2 | 0.01 | 1.44E-02 | Ion Transport |
| SCN4A | 0.01 | 2.77E-02 | Ion Transport |
| SCNN1A | 0.01 | 3.94E-03 | Ion Transport |
| KCNMB1 | 0.01 | 6.26E-04 | Ion Transport |
| SLC30A2 | 0.01 | 3.69E-02 | Ion Transport |
| KCNK17 | 3.98E-04 | 1.42E-02 | Ion Transport |
| AACS | 3.93 | 7.07E-04 | Ketone Bodies |
| BDH1 | 3.52 | 4.28E-07 | Ketone Bodies |
| HMGCLL1 | 3.51 | 4.50E-04 | Ketone Bodies |
| OXCT1 | 1.81 | 8.49E-03 | Ketone Bodies |
| BDH2 | 0.16 | 5.15E-04 | Ketone Bodies |
| PLA2G4C | 408.71 | 1.39E-03 | Lipid (Membrane Lipid) |
| AANAT | 211.42 | 9.19E-06 | Lipid (Hormone) |
| PLA2G12B | 205.65 | 1.06E-03 | Lipid (Membrane Lipid) |
| PLA2G4D | 109.58 | 4.46E-04 | Lipid (Membrane Lipid) |
| TPO | 105.68 | 1.70E-02 | Lipid (Hormone) |
| MTTP | 90.93 | 1.49E-03 | Lipid (Fatty Acid) |
| PLCH2 | 63.51 | 4.71E-07 | Lipid (Membrane Lipid) |
| ELOVL4 | 61.52 | 1.69E-04 | Lipid (Fatty Acid) |
| PLA2G4E | 44.08 | 4.30E-02 | Lipid (Membrane Lipid) |
| GDPD2 | 40.13 | 3.69E-03 | Lipid (Membrane Lipid) |
| LPPR4 | 27.09 | 8.21E-06 | Lipid (Membrane Lipid) |
| CYP1A1 | 26.20 | 6.90E-06 | Lipid (Steroid) |
| ALOX12B | 25.97 | 2.02E-07 | Lipid (Hormone) |
| AKR1C4 | 23.93 | 4.39E-02 | Lipid (Steroid) |
| ACBD7 | 23.87 | 4.34E-03 | Lipid (Fatty Acid) |
| SGPP2 | 22.67 | 6.88E-04 | Lipid (Sphingolipid) |
| CDS1 | 17.80 | 1.49E-08 | Lipid (Membrane Lipid) |
| ABHD12B | 17.04 | 1.48E-07 | Lipid (Fatty Acid) |
| FAAH2 | 14.69 | 7.50E-05 | Lipid (Fatty Acid) |
| BAAT | 10.75 | 1.37E-03 | Lipid (Bile Acid) |
| ACSL6 | 10.08 | 1.43E-05 | Lipid (Fatty Acid) |
| SMPD3 | 8.73 | 5.78E-07 | Lipid (Sphingolipid) |
| PHOSPHO1 | 8.61 | 1.74E-02 | Lipid (Membrane Lipid) |
| DGKI | 8.49 | 3.02E-06 | Lipid (Membrane Lipid) |
| ASAH2 | 8.20 | 1.26E-05 | Lipid (Sphingosine) |
| PLCD4 | 7.92 | 8.99E-04 | Lipid (Membrane Lipid) |
| GDPD4 | 7.15 | 2.24E-02 | Lipid (Membrane Lipid) |
| SCD5 | 6.67 | 1.32E-04 | Lipid (Fatty Acid) |
| ALOXE3 | 6.65 | 6.60E-03 | Lipid (Hormone) |
| CERKL | 6.64 | 1.47E-03 | Lipid (Sphingolipid) |
| DGKB | 6.52 | 1.74E-03 | Lipid (Membrane Lipid) |
| GDPD1 | 6.44 | 1.45E-04 | Lipid (Membrane Lipid) |
| ELOVL6 | 6.28 | 2.65E-05 | Lipid (Fatty Acid) |
| GPD1 | 6.18 | 3.35E-03 | Lipid (Membrane Lipid) |
| PLCB1 | 6.08 | 2.75E-04 | Lipid (Membrane Lipid) |
| CYP2E1 | 5.72 | 1.36E-03 | Lipid (Steroid) |
| LPCAT1 | 5.70 | 3.96E-05 | Lipid (Membrane Lipid) |
| UGT8 | 5.51 | 2.47E-03 | Lipid (Sphingolipid) |
| SRD5A2 | 5.45 | 3.31E-02 | Lipid (Steroid) |
| PPAP2C | 5.37 | 1.81E-03 | Lipid (Membrane Lipid) |
| KLB | 5.19 | 3.70E-02 | Lipid (Steroid) |
| ACSM4 | 5.01 | 3.39E-04 | Lipid (Fatty Acid) |
| PPAPDC3 | 4.64 | 3.20E-03 | Lipid (Membrane Lipid) |
| ACSBG2 | 4.62 | 1.73E-03 | Lipid (Fatty Acid) |
| ASAH2B | 4.46 | 1.71E-04 | Lipid (Sphingosine) |
| DGKD | 4.36 | 1.18E-04 | Lipid (Membrane Lipid) |
| GDPD5 | 4.24 | 6.23E-04 | Lipid (Membrane Lipid) |
| SRD5A1 | 4.21 | 7.87E-03 | Lipid (Steroid) |
| AGPAT3 | 4.00 | 4.24E-06 | Lipid (Membrane Lipid) |
| HSD11B1L | 3.98 | 1.88E-03 | Lipid (Steroid) |
| AGPAT9 | 3.98 | 1.11E-03 | Lipid (Membrane Lipid) |
| DAGLA | 3.93 | 3.94E-03 | Lipid (Membrane Lipid) |
| CDS2 | 3.91 | 1.23E-04 | Lipid (Membrane Lipid) |
| DGKE | 3.90 | 2.89E-04 | Lipid (Membrane Lipid) |
| LPCAT4 | 3.63 | 1.47E-03 | Lipid (Membrane Lipid) |
| PCYT1B | 3.60 | 3.78E-03 | Lipid (Membrane Lipid) |
| CYP2J2 | 3.40 | 2.45E-03 | Lipid (Steroid) |
| HSDL1 | 3.31 | 6.17E-04 | Lipid (Steroid) |
| ABHD8 | 3.23 | 6.83E-03 | Lipid (Fatty Acid) |
| CYP2D6 | 3.23 | 1.74E-02 | Lipid (Steroid) |
| CROT | 3.02 | 2.57E-03 | Lipid (Membrane Lipid) |
| PLCH1 | 2.99 | 1.33E-02 | Lipid (Membrane Lipid) |
| LIPI | 2.98 | 4.84E-02 | Lipid (Fatty Acid) |
| AMACR | 2.93 | 1.07E-04 | Lipid (Fatty acid) |
| CPT1C | 2.76 | 1.16E-04 | Lipid (Fatty Acid) |
| LSS | 2.61 | 3.15E-03 | Lipid (Cholesterol) |
| FAR2 | 2.49 | 9.11E-03 | Lipid (Fatty Acid) |
| PLCD3 | 2.48 | 3.50E-04 | Lipid (Membrane Lipid) |
| CYP17A1 | 2.43 | 1.64E-03 | Lipid (Steroid) |
| HMGCR | 2.29 | 1.27E-03 | Lipid (Mevalonate) |
| CYP46A1 | 2.04 | 7.93E-03 | Lipid (Steroid) |
| LYPLA1 | 2.03 | 3.26E-04 | Lipid (Membrane Lipid) |
| MECR | 1.92 | 5.21E-03 | Lipid (Fatty Acid) |
| SC5DL | 1.90 | 1.91E-04 | Lipid (Cholesterol) |
| GK | 1.83 | 1.80E-02 | Lipid (Membrane Lipid) |
| LNPEP | 1.82 | 9.32E-03 | Lipid (Hormone) |
| DDHD1 | 1.78 | 6.60E-05 | Lipid (Membrane Lipid) |
| ACSL3 | 1.75 | 1.43E-03 | Lipid (Fatty Acid) |
| ABHD14A | 1.72 | 2.00E-04 | Lipid (Fatty Acid) |
| THEM4 | 1.59 | 4.15E-04 | Lipid (Fatty Acid) |
| LIAS | 1.56 | 3.53E-02 | Lipid |
| AGPAT5 | 1.55 | 7.01E-03 | Lipid (Membrane Lipid) |
| DAK | 1.55 | 4.69E-04 | Lipid (Cofactor) |
| LPGAT1 | 1.53 | 2.93E-02 | Lipid (Membrane Lipid) |
| ABHD6 | 1.49 | 3.42E-02 | Lipid (Fatty Acid) |
| ETNK1 | 1.47 | 7.41E-03 | Lipid (Membrane Lipid) |
| EBP | 1.46 | 1.19E-02 | Lipid (Cholesterol) |
| SDR39U1 | 1.45 | 2.29E-03 | Lipid (Fatty Acid) |
| CERK | 1.45 | 3.25E-03 | Lipid (Sphingolipid) |
| MCAT | 1.44 | 1.73E-02 | Lipid (Fatty Acid) |
| TM7SF2 | 1.40 | 1.80E-03 | Lipid (Cholesterol) |
| GGPS1 | 1.39 | 8.86E-03 | Lipid (Steroid) |
| PLA2G6 | 1.38 | 2.07E-02 | Lipid (Membrane Lipid) |
| ACBD4 | 1.38 | 2.86E-02 | Lipid (Fatty Acid) |
| ETFB | 0.74 | 2.70E-02 | Lipid (Fatty Acid) |
| SPTLC3 | 0.72 | 1.93E-02 | Lipid (Sphingolipid) |
| ILVBL | 0.70 | 2.32E-03 | Lipid (Fatty Acid) |
| GNPAT | 0.69 | 1.77E-03 | Lipid (Membrane Lipid) |
| PEMT | 0.69 | 4.21E-02 | Lipid (Membrane Lipid) |
| FADS3 | 0.69 | 2.80E-02 | Lipid (Fatty Acid) |
| HSD3B7 | 0.68 | 3.14E-02 | Lipid (Bile Acid) |
| ACBD6 | 0.66 | 1.37E-02 | Lipid (Fatty Acid) |
| PCYT1A | 0.65 | 6.53E-03 | Lipid (Membrane Lipid) |
| ACER3 | 0.64 | 7.78E-03 | Lipid (Sphingosine) |
| ETFDH | 0.64 | 5.77E-03 | Lipid (Fatty Acid) |
| PGS1 | 0.63 | 1.28E-02 | Lipid (Membrane Lipid) |
| SOAT1 | 0.61 | 1.87E-04 | Lipid (Steroid) |
| CRAT | 0.61 | 1.79E-02 | Lipid (Cofactor) |
| GALC | 0.59 | 1.81E-02 | Lipid (Sphingolipid) |
| SCP2 | 0.59 | 2.11E-06 | Lipid (Fatty Acid) |
| DGKZ | 0.59 | 1.81E-02 | Lipid (Membrane Lipid) |
| CUBN | 0.59 | 2.59E-02 | Lipid (Cofactor) |
| ACAA2 | 0.58 | 5.36E-03 | Lipid (Fatty Acid) |
| DIO2 | 0.58 | 1.61E-02 | Lipid (Hormone) |
| ASAH1 | 0.58 | 1.60E-02 | Lipid (Sphingosine) |
| LTA4H | 0.57 | 1.38E-04 | Lipid (Hormone) |
| MCEE | 0.56 | 8.08E-03 | Lipid (Propanoate) |
| ETFA | 0.56 | 1.27E-04 | Lipid (Fatty Acid) |
| KDSR | 0.54 | 3.23E-03 | Lipid (Sphingosine) |
| ELOVL1 | 0.54 | 1.27E-03 | Lipid (Fatty Acid) |
| PTGR1 | 0.53 | 4.24E-04 | Lipid (Hormone) |
| ACAA1 | 0.52 | 8.13E-04 | Lipid (Fatty Acid) |
| CEPT1 | 0.51 | 2.53E-03 | Lipid (Membrane Lipid) |
| ACAD11 | 0.51 | 3.17E-03 | Lipid (Fatty Acid) |
| HADH | 0.50 | 1.88E-03 | Lipid (Fatty Acid) |
| CYP2U1 | 0.50 | 9.76E-03 | Lipid (Steroid) |
| SPTLC1 | 0.50 | 6.76E-06 | Lipid (Sphingolipid) |
| CPT1A | 0.48 | 6.96E-04 | Lipid (Fatty Acid) |
| HADHB | 0.48 | 1.33E-03 | Lipid (Fatty Acid) |
| ABHD4 | 0.48 | 1.66E-05 | Lipid (Fatty Acid) |
| HSD17B4 | 0.48 | 8.88E-04 | Lipid (Steroid) |
| ECHDC1 | 0.47 | 7.65E-03 | Lipid (Fatty Acid) |
| PLCD1 | 0.46 | 2.34E-03 | Lipid (Membrane Lipid) |
| EBPL | 0.44 | 4.46E-03 | Lipid (Cholesterol) |
| ABHD14B | 0.43 | 8.69E-04 | Lipid (Fatty Acid) |
| DEGS1 | 0.43 | 3.14E-04 | Lipid (Sphingolipid) |
| HSD17B14 | 0.42 | 8.82E-04 | Lipid (Steroid) |
| ETNK2 | 0.41 | 2.93E-03 | Lipid (Membrane Lipid) |
| SMPD1 | 0.41 | 7.16E-06 | Lipid (Sphingolipid) |
| PNPLA2 | 0.40 | 1.04E-02 | Lipid (Fatty Acid) |
| ACADL | 0.40 | 3.80E-04 | Lipid (Fatty Acid) |
| CYP7B1 | 0.38 | 3.26E-03 | Lipid (Bile Acid) |
| CYP20A1 | 0.38 | 5.66E-04 | Lipid (Hormone) |
| PLD2 | 0.37 | 1.04E-03 | Lipid (Membrane Lipid) |
| ECH1 | 0.37 | 2.39E-03 | Lipid (Fatty Acid) |
| HSD17B6 | 0.36 | 2.71E-02 | Lipid (Steroid) |
| PLCG1 | 0.36 | 1.86E-04 | Lipid (Membrane Lipid) |
| HADHA | 0.35 | 5.08E-06 | Lipid (Fatty Acid) |
| HSDL2 | 0.35 | 3.02E-02 | Lipid (Steroid) |
| ACADVL | 0.34 | 9.84E-05 | Lipid (Fatty Acid) |
| PLCB3 | 0.33 | 4.22E-04 | Lipid (Membrane Lipid) |
| DECR1 | 0.32 | 1.22E-03 | Lipid (Fatty Acid) |
| LBR | 0.32 | 5.13E-03 | Lipid (Cholesterol) |
| DEGS2 | 0.32 | 3.68E-02 | Lipid (Sphingolipid) |
| SMPDL3A | 0.32 | 1.51E-03 | Lipid (Sphingolipid) |
| ACSL1 | 0.30 | 3.10E-02 | Lipid (Fatty Acid) |
| HSD17B11 | 0.30 | 6.65E-03 | Lipid (Steroid) |
| PLCE1 | 0.30 | 3.36E-02 | Lipid (Membrane Lipid) |
| ACSS1 | 0.29 | 1.62E-03 | Lipid (Fatty Acid) |
| HSD17B10 | 0.29 | 5.73E-03 | Lipid (Steroid) |
| ELOVL5 | 0.29 | 2.93E-02 | Lipid (Fatty Acid) |
| DECR2 | 0.29 | 2.02E-05 | Lipid (Fatty Acid) |
| PLA2G16 | 0.29 | 4.72E-03 | Lipid (Membrane Lipid) |
| DGKA | 0.28 | 3.70E-04 | Lipid (Membrane Lipid) |
| LPCAT3 | 0.26 | 1.15E-02 | Lipid (Membrane Lipid) |
| ACSS3 | 0.23 | 1.97E-02 | Lipid (Fatty Acid) |
| CYP27A1 | 0.22 | 3.94E-04 | Lipid (Bile Acid) |
| LIPA | 0.22 | 3.14E-04 | Lipid (Steroid) |
| PTGS1 | 0.22 | 1.10E-02 | Lipid (Hormone) |
| DGAT2 | 0.21 | 3.45E-03 | Lipid (Fatty Acid) |
| PPAPDC1B | 0.21 | 7.79E-03 | Lipid (Membrane Lipid) |
| ABHD3 | 0.21 | 1.13E-02 | Lipid (Fatty Acid) |
| ACACB | 0.20 | 1.32E-03 | Lipid (Fatty Acid) |
| CHPT1 | 0.20 | 3.07E-02 | Lipid (Membrane Lipid) |
| LCAT | 0.18 | 9.16E-05 | Lipid (Cholesterol) |
| ACE2 | 0.17 | 1.34E-02 | Lipid (Hormone) |
| AGPAT2 | 0.17 | 2.44E-03 | Lipid (Membrane Lipid) |
| CYP7A1 | 0.17 | 2.17E-03 | Lipid (Bile Acid) |
| SGMS2 | 0.16 | 1.81E-02 | Lipid (Sphingolipid) |
| CYP39A1 | 0.16 | 1.46E-03 | Lipid (Steroid) |
| LPCAT2 | 0.16 | 3.21E-04 | Lipid (Membrane Lipid) |
| ACSF2 | 0.16 | 6.17E-03 | Lipid (Fatty Acid) |
| ACSM5 | 0.15 | 7.56E-03 | Lipid (Fatty Acid) |
| MBOAT1 | 0.15 | 6.00E-04 | Lipid (Membrane Lipid) |
| PLA2G4A | 0.14 | 2.59E-03 | Lipid (Membrane Lipid) |
| ACOX2 | 0.14 | 7.51E-03 | Lipid (Fatty Acid) |
| HSD17B3 | 0.14 | 3.42E-04 | Lipid (Steroid) |
| ACADS | 0.14 | 5.28E-05 | Lipid (Fatty Acid) |
| LPIN3 | 0.13 | 3.04E-06 | Lipid (Membrane Lipid) |
| AOAH | 0.12 | 4.14E-03 | Lipid (Fatty Acid) |
| ACER1 | 0.12 | 2.76E-02 | Lipid (Sphingosine) |
| ARSJ | 0.12 | 8.46E-03 | Lipid (Membrane Lipid) |
| PPAP2B | 0.12 | 6.89E-04 | Lipid (Membrane Lipid) |
| CYP4X1 | 0.12 | 3.96E-02 | Lipid (Fatty Acid) |
| TBXAS1 | 0.11 | 3.92E-04 | Lipid (Hormone) |
| PPAP2A | 0.10 | 5.39E-05 | Lipid (Membrane Lipid) |
| ARSE | 0.09 | 6.70E-03 | Lipid (Membrane Lipid) |
| PLCB2 | 0.09 | 1.45E-03 | Lipid (Membrane Lipid) |
| ACSM3 | 0.09 | 1.35E-02 | Lipid (Fatty Acid) |
| ECHDC3 | 0.09 | 1.54E-04 | Lipid (Fatty Acid) |
| PLB1 | 0.08 | 3.00E-02 | Lipid (Membrane Lipid) |
| LPL | 0.08 | 6.05E-03 | Lipid (Fatty Acid) |
| HPGD | 0.07 | 1.20E-05 | Lipid (Hormone) |
| PRG2 | 0.07 | 2.41E-02 | Lipid (Membrane Lipid) |
| ALDH3B1 | 0.06 | 1.98E-04 | Lipid (Fatty Acid) |
| PLD1 | 0.05 | 4.85E-06 | Lipid (Membrane Lipid) |
| ALOX5AP | 0.05 | 9.06E-03 | Lipid (Hormone) |
| LIPC | 0.05 | 6.69E-03 | Lipid (Fatty Acid) |
| ALOX5 | 0.05 | 1.17E-05 | Lipid (Hormone) |
| PLCG2 | 0.05 | 4.24E-06 | Lipid (Membrane Lipid) |
| ACSL5 | 0.04 | 5.71E-04 | Lipid (Fatty Acid) |
| MME | 0.03 | 5.38E-03 | Lipid (Hormone) |
| SDR42E1 | 0.03 | 1.03E-04 | Lipid (Fatty Acid) |
| CYP4Z1 | 0.03 | 6.46E-03 | Lipid (Fatty Acid) |
| CYP4A22 | 0.03 | 2.95E-03 | Lipid (Fatty Acid) |
| PLA1A | 0.03 | 1.47E-05 | Lipid (Membrane Lipid) |
| KL | 0.02 | 3.22E-03 | Lipid (Steroid) |
| PTGIS | 0.01 | 3.26E-04 | Lipid (Hormone) |
| MOGAT2 | 0.01 | 3.88E-03 | Lipid (Membrane Lipid) |
| CYP27C1 | 0.01 | 4.32E-02 | Lipid (Steroid) |
| CYP1B1 | 0.01 | 3.20E-03 | Lipid (Steroid) |
| CYP4B1 | 0.01 | 1.33E-03 | Lipid (Fatty Acid) |
| PLA2G2A | 4.11E-04 | 3.35E-02 | Lipid (Membrane Lipid) |
| NOX4 | 9.82 | 1.32E-03 | NAD |
| NOX3 | 8.76 | 2.41E-02 | NAD |
| NMNAT2 | 3.23 | 6.14E-03 | NAD |
| BST1 | 0.70 | 3.82E-02 | NAD |
| NMNAT1 | 0.53 | 1.75E-02 | NAD |
| NADK | 0.46 | 8.20E-05 | NAD |
| NADSYN1 | 0.43 | 1.61E-05 | NAD |
| NQO1 | 0.37 | 7.36E-03 | NAD |
| DUOX1 | 0.14 | 2.53E-02 | NAD |
| DUOX2 | 0.08 | 9.74E-03 | NAD |
| CD38 | 0.01 | 1.71E-03 | NAD |
| DDC | 55.77 | 4.10E-04 | Neurotransmitter |
| CHAT | 15.17 | 1.16E-03 | Neurotransmitter |
| ACHE | 11.90 | 2.35E-03 | Neurotransmitter |
| ABAT | 3.58 | 8.47E-04 | Neurotransmitter |
| COMTD1 | 3.20 | 5.40E-03 | Neurotransmitter |
| NAALADL2 | 0.53 | 1.01E-02 | Neurotransmitter |
| COMT | 0.46 | 3.64E-05 | Neurotransmitter |
| DBH | 0.17 | 1.82E-02 | Neurotransmitter |
| NAALADL1 | 0.16 | 3.06E-03 | Neurotransmitter |
| CA10 | 613.90 | 8.74E-06 | Nitrogen Metabolism |
| CA11 | 9.16 | 6.87E-04 | Nitrogen Metabolism |
| CA5A | 8.39 | 4.57E-02 | Nitrogen Metabolism |
| CA2 | 6.93 | 9.96E-04 | Nitrogen Metabolism |
| CA3 | 2.58 | 2.13E-02 | Nitrogen Metabolism |
| CA13 | 1.80 | 4.87E-02 | Nitrogen Metabolism |
| CA12 | 0.02 | 3.83E-02 | Nitrogen Metabolism |
| CA4 | 9.44E-04 | 2.31E-03 | Nitrogen Metabolism |
| PDE6A | 393.76 | 5.03E-04 | Nucleotide (cGMP) |
| DPYSL5 | 323.65 | 1.54E-06 | Nucleotide (Pyrimidine) |
| GUCY2F | 309.73 | 6.42E-04 | Nucleotide (cGMP) |
| PDE6H | 303.46 | 1.16E-05 | Nucleotide (cGMP) |
| ADCY8 | 297.69 | 4.93E-05 | Nucleotide (cAMP) |
| GUCY2D | 52.39 | 1.01E-04 | Nucleotide (cGMP) |
| PDE6C | 47.73 | 9.49E-05 | Nucleotide (cGMP) |
| PDE6B | 30.71 | 2.33E-03 | Nucleotide (cGMP) |
| PDE11A | 29.69 | 1.07E-04 | Nucleotide (cyclic nucleotide) |
| ADCY1 | 25.74 | 2.07E-07 | Nucleotide (cAMP) |
| NT5C1A | 21.36 | 1.18E-02 | Nucleotide |
| DPYSL4 | 17.74 | 2.97E-04 | Nucleotide (Pyrimidine) |
| NT5E | 12.33 | 3.06E-03 | Nucleotide |
| A1CF | 11.70 | 4.48E-02 | Nucleotide |
| IMPDH1 | 9.88 | 1.47E-04 | Nucleotide (Purine) |
| GUCY1A2 | 7.00 | 4.82E-04 | Nucleotide (cGMP) |
| GDA | 6.96 | 8.53E-03 | Nucleotide (Purine) |
| NT5M | 6.46 | 5.64E-03 | Nucleotide |
| UPP2 | 5.84 | 1.63E-03 | Nucleotide (Pyrimidine) |
| ENTPD3 | 4.35 | 7.22E-03 | Nucleotide |
| HPRT1 | 4.13 | 4.73E-05 | Nucleotide (Purine) |
| AK7 | 3.67 | 4.74E-03 | Nucleotide (Purine) |
| GUK1 | 3.44 | 7.53E-06 | Nucleotide (Purine) |
| NUDT12 | 3.33 | 1.45E-03 | Nucleotide |
| NUDT10 | 3.32 | 2.16E-03 | Nucleotide |
| PDE4D | 2.71 | 1.24E-02 | Nucleotide (cGMP) |
| PDE4A | 2.63 | 7.65E-03 | Nucleotide (cGMP) |
| TYMS | 2.61 | 2.31E-03 | Nucleotide (Pyrimidine) |
| NUDT19 | 2.61 | 2.17E-03 | Nucleotide |
| DPYSL2 | 2.60 | 2.43E-03 | Nucleotide (Pyrimidine) |
| UPB1 | 2.52 | 1.09E-02 | Nucleotide (Pyrimidine) |
| PDE6D | 2.51 | 1.10E-03 | Nucleotide (cGMP) |
| GUCY1B3 | 2.43 | 7.69E-05 | Nucleotide (cGMP) |
| NUDT11 | 2.35 | 3.28E-02 | Nucleotide |
| NT5C2 | 2.30 | 2.96E-05 | Nucleotide |
| ADAL | 2.29 | 1.04E-03 | Nucleotide (Purine) |
| AMPD3 | 2.18 | 1.65E-04 | Nucleotide (Purine) |
| DPYSL3 | 2.18 | 9.49E-03 | Nucleotide (Pyrimidine) |
| ENTPD4 | 2.18 | 8.19E-04 | Nucleotide |
| NME5 | 2.14 | 2.75E-02 | Nucleotide |
| AK5 | 2.00 | 1.81E-03 | Nucleotide (Purine) |
| PDE1A | 1.93 | 4.67E-02 | Nucleotide (cyclic nucleotide) |
| PRPSAP2 | 1.87 | 6.39E-05 | Nucleotide (Purine) |
| NME7 | 1.72 | 5.78E-05 | Nucleotide |
| DCK | 1.65 | 3.64E-03 | Nucleotide |
| AMPD2 | 1.58 | 2.07E-02 | Nucleotide (Purine) |
| ADK | 1.47 | 3.70E-03 | Nucleotide (Purine) |
| RRM1 | 1.45 | 2.51E-03 | Nucleotide |
| RRM2B | 1.41 | 1.43E-02 | Nucleotide |
| NUDT16L1 | 1.36 | 7.54E-03 | Nucleotide |
| UMPS | 0.76 | 4.95E-02 | Nucleotide (Pyrimidine) |
| NUDT9 | 0.71 | 1.45E-03 | Nucleotide |
| DUT | 0.68 | 1.25E-02 | Nucleotide (Pyrimidine) |
| CANT1 | 0.66 | 2.37E-02 | Nucleotide |
| NME6 | 0.64 | 2.83E-02 | Nucleotide |
| ADCY6 | 0.62 | 5.94E-03 | Nucleotide (cAMP) |
| FHIT | 0.61 | 1.82E-02 | Nucleotide (Purine) |
| CMPK | 0.60 | 2.28E-03 | Nucleotide (Pyrimidine) |
| GMPS | 0.60 | 1.06E-02 | Nucleotide (Purine) |
| ATIC | 0.58 | 5.92E-04 | Nucleotide (Purine) |
| NT5DC1 | 0.56 | 4.50E-03 | Nucleotide |
| CTPS | 0.55 | 3.34E-02 | Nucleotide (Pyrimidine) |
| ENTPD1 | 0.55 | 1.73E-04 | Nucleotide |
| NUDT5 | 0.55 | 2.11E-04 | Nucleotide |
| ADSSL1 | 0.55 | 3.73E-02 | Nucleotide (Purine) |
| NUDT6 | 0.53 | 1.13E-02 | Nucleotide |
| TK2 | 0.52 | 3.69E-04 | Nucleotide (Pyrimidine) |
| IMPDH2 | 0.51 | 1.52E-03 | Nucleotide (Purine) |
| NUDT3 | 0.47 | 5.93E-03 | Nucleotide |
| ADCY9 | 0.46 | 4.29E-02 | Nucleotide (cAMP) |
| NT5DC2 | 0.46 | 5.78E-03 | Nucleotide |
| CECR1 | 0.40 | 2.80E-02 | Nucleotide (Purine) |
| DCTD | 0.35 | 7.09E-06 | Nucleotide (Pyrimidine) |
| APRT | 0.32 | 1.33E-02 | Nucleotide (Purine) |
| NME4 | 0.29 | 2.02E-05 | Nucleotide |
| ADCY7 | 0.25 | 5.93E-04 | Nucleotide (cAMP) |
| PDE1C | 0.24 | 1.64E-02 | Nucleotide (cGMP) |
| PDE4C | 0.15 | 3.76E-02 | Nucleotide (cGMP) |
| AK3 | 0.15 | 1.24E-05 | Nucleotide (Purine) |
| PDE3B | 0.14 | 5.20E-04 | Nucleotide (cGMP) |
| PDE1B | 0.13 | 7.85E-03 | Nucleotide (cyclic nucleotide) |
| ADA | 0.13 | 5.74E-04 | Nucleotide (Purine) |
| OAS3 | 0.11 | 7.09E-04 | Nucleotide (Ribonuclease) |
| PDE2A | 0.11 | 1.36E-03 | Nucleotide (cGMP) |
| PRPS2 | 0.10 | 4.29E-06 | Nucleotide (Purine) |
| TK1 | 0.10 | 5.52E-03 | Nucleotide (Pyrimidine) |
| OAS1 | 0.09 | 1.43E-02 | Nucleotide (Ribonuclease) |
| CDA | 0.09 | 7.80E-03 | Nucleotide (Pyrimidine) |
| RRM2 | 0.07 | 1.24E-02 | Nucleotide |
| DPYD | 0.07 | 1.78E-02 | Nucleotide (Pyrimidine) |
| GMPR | 0.07 | 4.80E-03 | Nucleotide (Purine) |
| NPR1 | 0.04 | 4.71E-02 | Nucleotide (cGMP) |
| OAS2 | 0.04 | 3.36E-03 | Nucleotide (Ribonuclease) |
| GUCY2C | 0.04 | 1.31E-03 | Nucleotide (cGMP) |
| ADCY4 | 0.04 | 4.02E-04 | Nucleotide (cAMP) |
| OASL | 0.03 | 1.05E-02 | Nucleotide (Ribonuclease) |
| ADAD2 | 0.02 | 3.99E-02 | Nucleotide (Purine) |
| ENPP6 | 0.01 | 4.00E-02 | Nucleotide |
| UROS | 1.98 | 2.52E-04 | Porphyrin |
| HMOX2 | 1.51 | 1.16E-03 | Porphyrin |
| PPOX | 1.36 | 4.24E-03 | Porphyrin |
| UROD | 0.69 | 6.10E-05 | Porphyrin |
| BLVRA | 0.61 | 7.73E-03 | Porphyrin |
| FXN | 0.45 | 1.55E-03 | Porphyrin |
| BLVRB | 0.43 | 9.93E-04 | Porphyrin |
| HMOX1 | 0.07 | 3.61E-03 | Porphyrin |
| ATP6V0D2 | 247.66 | 1.46E-04 | Proton Transport |
| ATP8A2 | 30.56 | 1.74E-06 | Proton Transport |
| ATP13A2 | 10.33 | 9.80E-04 | Proton Transport |
| ATP6V1B1 | 8.69 | 1.11E-04 | Proton Transport |
| ATP13A5 | 7.22 | 3.30E-02 | Proton Transport |
| ATP9A | 4.28 | 1.69E-03 | Proton Transport |
| ATP6V1B2 | 3.34 | 1.63E-06 | Proton Transport |
| ATP6V0A1 | 3.06 | 9.88E-06 | Proton Transport |
| ATP6V0E2 | 2.85 | 4.71E-03 | Proton Transport |
| ATP6V1E2 | 2.04 | 8.16E-04 | Proton Transport |
| ATP5SL | 2.00 | 8.52E-05 | Proton Transport |
| ATP5B | 1.99 | 2.10E-05 | Proton Transport |
| ATP5A1 | 1.96 | 3.92E-05 | Proton Transport |
| ATP6V1A | 1.96 | 9.06E-03 | Proton Transport |
| ATP6V1H | 1.69 | 9.65E-04 | Proton Transport |
| ATP6V1C1 | 1.63 | 3.41E-03 | Proton Transport |
| ATP6V0B | 1.61 | 5.14E-03 | Proton Transport |
| ATP6V1D | 1.58 | 2.86E-05 | Proton Transport |
| ATP6V1G2 | 1.50 | 1.73E-02 | Proton Transport |
| ATP6V0D1 | 1.42 | 3.30E-03 | Proton Transport |
| ATP5C1 | 1.37 | 6.81E-03 | Proton Transport |
| ATP6V1F | 1.34 | 1.99E-02 | Proton Transport |
| ATP6V1G1 | 0.73 | 6.20E-04 | Proton Transport |
| ATP13A3 | 0.59 | 3.48E-02 | Proton Transport |
| ATP6V0A2 | 0.59 | 2.71E-02 | Proton Transport |
| ATP5E | 0.56 | 1.06E-03 | Proton Transport |
| ATP5F1 | 0.50 | 1.84E-04 | Proton Transport |
| ATP10D | 0.41 | 5.24E-04 | Proton Transport |
| TAP1 | 0.32 | 5.48E-03 | Proton Transport |
| ATP8B2 | 0.21 | 4.92E-04 | Proton Transport |
| HVCN1 | 0.18 | 9.52E-03 | Proton Transport |
| UCP2 | 0.16 | 3.88E-04 | Proton Transport |
| ATP10A | 0.15 | 1.13E-02 | Proton Transport |
| ATP6V0A4 | 0.11 | 3.03E-03 | Proton Transport |
| ATP6V0E1 | 0.10 | 1.63E-05 | Proton Transport |
| TCIRG1 | 0.10 | 2.99E-02 | Proton Transport |
| ATP8B1 | 0.05 | 3.78E-03 | Proton Transport |
| ATP8B4 | 0.03 | 5.76E-03 | Proton Transport |
| DHRS11 | 5.81 | 1.41E-05 | Redox |
| PRDX2 | 2.69 | 1.11E-05 | Redox |
| CYB561 | 2.38 | 3.42E-03 | Redox |
| MICAL2 | 2.02 | 8.77E-03 | Redox |
| CYB5R4 | 2.01 | 3.87E-04 | Redox |
| CYB5D1 | 1.92 | 2.17E-03 | Redox |
| FOXRED2 | 1.73 | 4.58E-02 | Redox |
| ZADH2 | 1.64 | 5.62E-04 | Redox |
| FOXRED1 | 1.62 | 2.46E-03 | Redox |
| PYROXD1 | 1.57 | 2.79E-02 | Redox |
| DHRS7 | 1.54 | 2.28E-02 | Redox |
| PRDX5 | 1.53 | 9.74E-03 | Redox |
| ISCA1 | 1.47 | 7.18E-04 | Redox |
| CYB5R1 | 0.75 | 3.64E-02 | Redox |
| CYB5D2 | 0.72 | 3.06E-02 | Redox |
| DHRS7B | 0.61 | 3.33E-03 | Redox |
| PRDX3 | 0.58 | 1.43E-03 | Redox |
| FDX1 | 0.53 | 7.48E-03 | Redox |
| PRDX1 | 0.52 | 6.95E-04 | Redox |
| CYB5R3 | 0.52 | 1.76E-02 | Redox |
| NMRAL1 | 0.47 | 2.23E-03 | Redox |
| PRDX4 | 0.46 | 4.19E-03 | Redox |
| ENOX2 | 0.40 | 7.89E-04 | Redox |
| TXNRD3 | 0.38 | 5.49E-05 | Redox |
| CYBASC3 | 0.35 | 2.18E-04 | Redox |
| PRDX6 | 0.33 | 2.24E-02 | Redox |
| POR | 0.33 | 1.99E-03 | Redox |
| TXNRD1 | 0.32 | 5.47E-04 | Redox |
| DHRS4 | 0.31 | 8.11E-03 | Redox |
| DHRS4L2 | 0.31 | 8.11E-03 | Redox |
| CP | 0.24 | 4.40E-02 | Redox |
| CYBB | 0.20 | 1.18E-02 | Redox |
| FRRS1 | 0.14 | 1.64E-04 | Redox |
| CYB5A | 0.13 | 1.57E-05 | Redox |
| CYBRD1 | 0.11 | 1.01E-03 | Redox |
| DHRS3 | 0.05 | 2.69E-05 | Redox |
| CYBA | 0.02 | 1.69E-02 | Redox |
| SLC17A6 | 1731.20 | 3.85E-03 | Small Molecule Transport (Glutamate, Asparate, Lactate) |
| SLC32A1 | 781.05 | 7.53E-04 | Small Molecule Transport (GABA, Gly) |
| SLC17A7 | 336.47 | 6.76E-05 | Small Molecule Transport (Glutamate, Asparate, Lactate) |
| SLC6A11 | 326.30 | 1.45E-04 | Small Molecule Transport (GABA, β-Alanine) |
| SLC5A8 | 285.71 | 3.64E-03 | Small Molecule Transport (iodide) |
| SLC13A2 | 194.07 | 1.51E-03 | Small Molecule Transport (Citrate, Succinate) |
| SLC22A7 | 128.85 | 2.72E-03 | Small Molecule Transport (Anion) |
| SLC1A6 | 128.74 | 1.20E-04 | Small Molecule Transport (Glutamate) |
| SV2B | 121.78 | 3.91E-05 | Small Molecule Transport (Sugar) |
| SLC1A2 | 118.18 | 4.40E-04 | Small Molecule Transport (Glutamate, Asparate) |
| AQP4 | 109.55 | 1.58E-05 | Small Molecule Transport (Water) |
| SLC34A1 | 87.33 | 6.49E-04 | Small Molecule Transport (Mitochondrial Phosphate) |
| SV2C | 85.69 | 6.89E-03 | Small Molecule Transport (Sugar) |
| SVOP | 73.79 | 1.47E-09 | Small Molecule Transport |
| RHCG | 63.78 | 1.05E-02 | Small Molecule Transport (Ammonium) |
| SLC35F4 | 50.54 | 7.22E-04 | Small Molecule Transport (Nucleotide sugar) |
| SLC27A6 | 46.12 | 1.43E-04 | Small Molecule Transport (Fatty acid) |
| SLC6A1 | 41.92 | 1.00E-05 | Small Molecule Transport (GABA) |
| SLC38A1 | 40.16 | 9.09E-06 | Small Molecule Transport (Gln, Ala, Ser, Asn) |
| SLC22A14 | 35.61 | 3.58E-02 | Small Molecule Transport (Organic Cation) |
| SLC7A14 | 27.53 | 5.21E-05 | Small Molecule Transport (Amino acid) |
| SV2A | 20.00 | 4.12E-05 | Small Molecule Transport (Sugar) |
| SLC5A4 | 18.00 | 1.97E-02 | Small Molecule Transport (Glucose) |
| SLC17A8 | 17.60 | 1.07E-05 | Small Molecule Transport (Glutamate, Asparate, Lactate) |
| SLC4A10 | 17.30 | 2.59E-05 | Small Molecule Transport (Biocarbonate) |
| SLC13A5 | 17.08 | 1.20E-04 | Small Molecule Transport (Citrate, Pyruvate) |
| SLC35F3 | 16.26 | 6.21E-03 | Small Molecule Transport (Nucleotide sugar) |
| SLC34A2 | 15.83 | 6.10E-03 | Small Molecule Transport (Mitochondrial Phosphate) |
| MFSD4 | 14.61 | 1.19E-06 | Small Molecule Transport (Glucose) |
| SLC26A3 | 14.59 | 5.12E-04 | Small Molecule Transport (Biocarbonate) |
| SLC44A5 | 12.45 | 8.45E-04 | Small Molecule Transport (Choline) |
| SLC6A5 | 12.36 | 2.44E-03 | Small Molecule Transport (Glycine) |
| SLC4A3 | 12.21 | 4.96E-04 | Small Molecule Transport (Anion) |
| SLC47A2 | 10.94 | 2.47E-03 | Small Molecule Transport (Thiamine) |
| SLC4A8 | 10.23 | 9.30E-05 | Small Molecule Transport (Biocarbonate) |
| SLC6A17 | 10.00 | 3.83E-03 | Small Molecule Transport (Pro, Cys, Ala) |
| AQP5 | 9.83 | 6.08E-03 | Small Molecule Transport (Water) |
| SLC25A22 | 7.84 | 1.06E-04 | Small Molecule Transport (Glutamate) |
| SLC38A3 | 7.83 | 2.22E-04 | Small Molecule Transport (His, Gln, Asn, Ala, Glutamate) |
| SLC1A3 | 7.30 | 7.69E-04 | Small Molecule Transport (Glutamate, Asparate) |
| SLC1A7 | 7.23 | 8.15E-05 | Small Molecule Transport (Glutamate) |
| SLC38A4 | 6.89 | 4.07E-02 | Small Molecule Transport (His, Ar, Alan, Asn) |
| SLC35F1 | 6.73 | 6.34E-05 | Small Molecule Transport (Nucleotide sugar) |
| SLC4A7 | 6.53 | 3.94E-03 | Small Molecule Transport (Biocarbonate) |
| SLC6A7 | 6.48 | 9.15E-04 | Small Molecule Transport (Proline) |
| SLC22A20 | 6.19 | 4.39E-04 | Small Molecule Transport (Organic Cation) |
| SLC15A2 | 5.92 | 4.89E-04 | Small Molecule Transport (peptide) |
| STAR | 5.63 | 5.25E-05 | Small Molecule Transport (Cholesterol) |
| SLC35D3 | 5.39 | 9.65E-03 | Small Molecule Transport (Nucleotide sugar) |
| SLCO5A1 | 5.29 | 6.32E-04 | Small Molecule Transport (peptide) |
| SLC45A1 | 5.28 | 4.07E-03 | Small Molecule Transport (Glucose) |
| SLC6A6 | 5.13 | 1.00E-02 | Small Molecule Transport (GABA,  β-alanine,taurine) |
| SLC10A5 | 4.31 | 5.61E-03 | Small Molecule Transport (Bile acid) |
| SLC16A6 | 4.16 | 3.80E-03 | Small Molecule Transport (MCT activity) |
| SLC4A11 | 4.15 | 1.93E-04 | Small Molecule Transport (Biocarbonate) |
| SLC2A11 | 4.13 | 1.38E-03 | Small Molecule Transport (Glucose, Fructose) |
| SLC26A8 | 3.56 | 1.89E-02 | Small Molecule Transport (Biocarbonate) |
| SLC2A6 | 3.44 | 9.19E-03 | Small Molecule Transport (Glucose) |
| SLC7A11 | 3.43 | 7.63E-03 | Small Molecule Transport (Cystine, Glutamate) |
| SLC7A1 | 3.43 | 1.06E-03 | Small Molecule Transport (Ar, Lys, Orn, His) |
| SLC5A2 | 3.39 | 4.62E-02 | Small Molecule Transport (Glucose) |
| SLC29A2 | 3.32 | 1.19E-03 | Small Molecule Transport (inosine) |
| SLC25A41 | 3.20 | 7.43E-03 | Small Molecule Transport (Nucleotide) |
| SLC23A2 | 3.19 | 5.77E-04 | Small Molecule Transport (Vitamine C) |
| STARD9 | 3.09 | 2.29E-02 | Small Molecule Transport (Lipid) |
| SLC25A12 | 2.94 | 1.07E-04 | Small Molecule Transport (Glutamate, Aspartate) |
| SLC22A15 | 2.87 | 6.11E-05 | Small Molecule Transport (Organic Cation) |
| SLC29A1 | 2.65 | 1.51E-03 | Small Molecule Transport (inosine) |
| SLC36A4 | 2.61 | 1.61E-03 | Small Molecule Transport (Proline) |
| SFXN1 | 2.53 | 2.26E-04 | Small Molecule Transport (Serine) |
| SLC25A25 | 2.51 | 4.16E-04 | Small Molecule Transport (Mitochondrial Phosphate) |
| SLC35B4 | 2.44 | 2.65E-04 | Small Molecule Transport (UDP-N-Acetyl-Glucosomine) |
| SLC25A4 | 2.39 | 2.26E-03 | Small Molecule Transport (Nucleotide) |
| SFXN3 | 2.33 | 5.73E-05 | Small Molecule Transport (Serine) |
| SLC27A4 | 2.25 | 5.17E-03 | Small Molecule Transport (Fatty acid) |
| SLC35F2 | 2.23 | 4.30E-02 | Small Molecule Transport (Nucleotide sugar) |
| SLC6A15 | 2.21 | 1.37E-02 | Small Molecule Transport (Met, BCAA) |
| SLC25A40 | 2.20 | 6.73E-04 | Small Molecule Transport (Amino acid) |
| SLC35E2 | 2.18 | 1.90E-03 | Small Molecule Transport (Nucleotide sugar) |
| SLC18A2 | 2.17 | 2.50E-04 | Small Molecule Transport (Monoamine) |
| SLC25A33 | 2.08 | 2.31E-03 | Small Molecule Transport (Nucleotide) |
| SLC37A4 | 2.02 | 6.93E-04 | Small Molecule Transport (G6P) |
| SLC26A5 | 2.01 | 6.15E-03 | Small Molecule Transport (Biocarbonate) |
| SLC25A14 | 2.00 | 2.59E-05 | Small Molecule Transport (MCT activity) |
| SLC46A1 | 1.92 | 7.38E-03 | Small Molecule Transport (Folate) |
| FLVCR1 | 1.88 | 4.94E-03 | Small Molecule Transport (Hem) |
| SLC6A8 | 1.87 | 1.04E-02 | Small Molecule Transport (Creatine) |
| AQP11 | 1.87 | 3.76E-02 | Small Molecule Transport (Water, Glycerol) |
| SLC25A44 | 1.80 | 7.68E-05 | Small Molecule Transport (Amino acid) |
| SLC25A18 | 1.74 | 2.08E-02 | Small Molecule Transport (Glutamate) |
| SLC25A46 | 1.70 | 4.55E-03 | Small Molecule Transport (mitochondrial transport) |
| SLC25A23 | 1.67 | 6.05E-03 | Small Molecule Transport (Mitochondrial Phosphate) |
| SLC17A5 | 1.65 | 1.52E-02 | Small Molecule Transport (Glutamate, Asparate, Lactate) |
| SLC4A1AP | 1.58 | 3.14E-02 | Small Molecule Transport (Anion) |
| SLC1A4 | 1.56 | 1.48E-02 | Small Molecule Transport (Alanine, Cysteine, Serine) |
| SLC27A5 | 1.54 | 1.44E-02 | Small Molecule Transport (Fatty acid) |
| MFSD11 | 1.52 | 5.86E-03 | Small Molecule Transport |
| SLC35D1 | 1.46 | 4.10E-05 | Small Molecule Transport (UDP-N-Acetyl-Glucosomine) |
| SLC25A5 | 1.42 | 3.64E-02 | Small Molecule Transport (Nucleotide) |
| SLC15A4 | 1.40 | 4.54E-02 | Small Molecule Transport (peptide) |
| SLC35E3 | 1.31 | 3.65E-02 | Small Molecule Transport (Nucleotide sugar) |
| SLC37A1 | 1.30 | 4.93E-02 | Small Molecule Transport (G6P, G3P) |
| SLC10A3 | 0.76 | 3.87E-02 | Small Molecule Transport (Bile acid) |
| SFXN4 | 0.69 | 4.00E-03 | Small Molecule Transport |
| SLC25A32 | 0.69 | 3.39E-02 | Small Molecule Transport (Nucleotide) |
| SLC7A6 | 0.68 | 2.31E-02 | Small Molecule Transport (Amino acid) |
| SLC35F5 | 0.65 | 9.25E-03 | Small Molecule Transport (Nucleotide sugar) |
| SLC25A35 | 0.63 | 3.85E-03 | Small Molecule Transport ( Di-Tri-carboxylic acid) |
| NPC1 | 0.60 | 5.67E-03 | Small Molecule Transport (Cholesterol) |
| SLC22A5 | 0.60 | 3.69E-02 | Small Molecule Transport (Carnitine) |
| SLC35A1 | 0.58 | 5.21E-04 | Small Molecule Transport (Nucleotide sugar) |
| SLC25A39 | 0.57 | 2.73E-03 | Small Molecule Transport (Amino acid) |
| SLC5A6 | 0.55 | 5.53E-03 | Small Molecule Transport (Vitamine) |
| SLC7A5 | 0.53 | 1.06E-02 | Small Molecule Transport (Amino acid) |
| STARD3 | 0.51 | 2.61E-03 | Small Molecule Transport (Cholesterol) |
| SLC47A1 | 0.51 | 6.35E-03 | Small Molecule Transport (Thiamine) |
| SLC33A1 | 0.50 | 4.90E-02 | Small Molecule Transport (Acetyl CoA) |
| SLC16A2 | 0.50 | 6.77E-03 | Small Molecule Transport (T4) |
| SLC25A1 | 0.49 | 3.91E-03 | Small Molecule Transport (Citrate, Malate) |
| SLC27A1 | 0.49 | 3.99E-05 | Small Molecule Transport (Fatty acid) |
| SLC35B3 | 0.48 | 2.39E-04 | Small Molecule Transport (Nucleotide sugar) |
| SLC19A2 | 0.47 | 1.87E-03 | Small Molecule Transport (Thiamine) |
| SLC10A4 | 0.46 | 4.92E-02 | Small Molecule Transport (Bile acid) |
| SLC44A1 | 0.46 | 3.60E-03 | Small Molecule Transport (Choline) |
| SLC36A1 | 0.45 | 2.52E-02 | Small Molecule Transport (Glycine) |
| SLC45A3 | 0.45 | 9.21E-04 | Small Molecule Transport (Sugar) |
| SLC35A2 | 0.44 | 3.40E-02 | Small Molecule Transport (Nucleotide sugar) |
| SLC23A3 | 0.44 | 3.45E-02 | Small Molecule Transport (Citrate, Malate) |
| CTNS | 0.42 | 4.67E-03 | Small Molecule Transport (Cystine) |
| SLC25A20 | 0.40 | 5.20E-03 | Small Molecule Transport (Carnitine) |
| SLC15A3 | 0.40 | 1.50E-02 | Small Molecule Transport (peptide) |
| PCTP | 0.37 | 9.45E-04 | Small Molecule Transport (PC) |
| SLC20A2 | 0.37 | 1.68E-02 | Small Molecule Transport (Phosphate) |
| SLC13A3 | 0.34 | 3.40E-02 | Small Molecule Transport (Citrate, Succinate) |
| SLC29A3 | 0.33 | 1.94E-03 | Small Molecule Transport (inosine) |
| SLC7A8 | 0.31 | 4.04E-04 | Small Molecule Transport (Amino acid) |
| SLC44A2 | 0.30 | 3.89E-04 | Small Molecule Transport (Choline) |
| SLC2A4 | 0.29 | 1.96E-05 | Small Molecule Transport (Glucose, Glucosamine) |
| MFSD1 | 0.29 | 6.87E-04 | Small Molecule Transport |
| STARD13 | 0.27 | 3.04E-05 | Small Molecule Transport (Lipid) |
| SLC5A9 | 0.23 | 1.91E-02 | Small Molecule Transport (Glucose) |
| MFSD7 | 0.22 | 2.84E-03 | Small Molecule Transport |
| SLC25A45 | 0.22 | 1.52E-02 | Small Molecule Transport (Amino acid) |
| SLC4A2 | 0.21 | 2.57E-02 | Small Molecule Transport (Biocarbonate) |
| SLC44A3 | 0.20 | 1.31E-03 | Small Molecule Transport (Choline) |
| SLC43A1 | 0.20 | 1.32E-04 | Small Molecule Transport (BCAA) |
| SLC44A4 | 0.17 | 3.06E-02 | Small Molecule Transport (Choline) |
| SLC25A43 | 0.17 | 6.43E-04 | Small Molecule Transport (Nucleotide) |
| SLC37A2 | 0.14 | 1.60E-03 | Small Molecule Transport (G6P) |
| SLC27A3 | 0.14 | 7.12E-05 | Small Molecule Transport (Fatty acid) |
| SLC35D2 | 0.14 | 3.42E-04 | Small Molecule Transport (Nucleotide sugar) |
| SLC7A7 | 0.10 | 5.28E-03 | Small Molecule Transport (Amino acid) |
| SLC25A30 | 0.10 | 4.95E-04 | Small Molecule Transport (mitochondrial transport) |
| SLC6A12 | 0.09 | 2.99E-03 | Small Molecule Transport (GABA, Betaine |
| SLCO2B1 | 0.08 | 1.09E-03 | Small Molecule Transport (T4) |
| SLC2A10 | 0.08 | 1.79E-03 | Small Molecule Transport (Glucose) |
| SLC22A2 | 0.07 | 2.78E-02 | Small Molecule Transport (dopamine  histamine,PGE2) |
| SLC16A4 | 0.04 | 6.19E-03 | Small Molecule Transport (MCT activity) |
| SLC16A5 | 0.04 | 1.17E-03 | Small Molecule Transport (MCT activity) |
| STARD8 | 0.04 | 1.85E-06 | Small Molecule Transport (Lipid) |
| SLC26A4 | 0.03 | 5.75E-03 | Small Molecule Transport (Biocarbonate) |
| SLC22A3 | 0.03 | 6.90E-05 | Small Molecule Transport (Monoamine) |
| SLC16A8 | 0.03 | 1.10E-02 | Small Molecule Transport (Lactate) |
| SLC43A3 | 0.03 | 1.26E-02 | Small Molecule Transport (Amino acid) |
| AQP7 | 0.02 | 4.49E-03 | Small Molecule Transport (Water) |
| SLC45A2 | 0.02 | 1.83E-02 | Small Molecule Transport (Sugar) |
| AQP1 | 0.02 | 2.64E-02 | Small Molecule Transport( Ammonium) |
| FLVCR2 | 0.02 | 7.11E-04 | Small Molecule Transport (Hem) |
| SLC14A1 | 0.01 | 2.34E-02 | Small Molecule Transport (Urea) |
| STRA6 | 0.01 | 4.14E-02 | Small Molecule Transport (Vitamine A) |
| SLC26A7 | 0.01 | 2.69E-02 | Small Molecule Transport (Biocarbonate) |
| SLC14A2 | 0.01 | 4.13E-03 | Small Molecule Transport (Urea) |
| SLCO2A1 | 1.83E-03 | 3.63E-04 | Small Molecule Transport (eicosanoids,prostaglandins) |
| FOLR2 | 1.63E-03 | 6.13E-03 | Small Molecule Transport (Folate) |
| FOLR3 | 1.63E-03 | 6.13E-03 | Small Molecule Transport (Folate) |
| SLC5A10 | 5.73E-04 | 2.06E-02 | Small Molecule Transport (Glucose) |
| G6PC2 | 468.51 | 8.50E-03 | Sugar (gluconeogenesi) |
| MIOX | 26.49 | 1.71E-03 | Sugar (Inositol Phosphate) |
| INPP5J | 12.80 | 1.49E-03 | Sugar (Inositol Phosphate) |
| PIP5KL1 | 5.91 | 1.41E-06 | Sugar (Inositol Phosphate) |
| PIK3R4 | 5.38 | 2.64E-04 | Sugar (Inositol Phosphate) |
| KHK | 4.72 | 1.90E-03 | Sugar (Fructose) |
| AMY2A | 4.68 | 6.54E-03 | Sugar (Amylase) |
| INPP5F | 4.48 | 1.24E-05 | Sugar (Inositol Phosphate) |
| ITPKA | 4.28 | 3.03E-02 | Sugar (Inositol Phosphate) |
| TKTL1 | 4.25 | 1.39E-03 | Sugar (PPP) |
| INPP4A | 3.01 | 1.07E-03 | Sugar (Inositol Phosphate) |
| PGAP1 | 2.70 | 3.51E-03 | Sugar (Inositol Phosphate) |
| PYGB | 2.62 | 4.89E-04 | Sugar (Glycogen) |
| SYNJ1 | 2.54 | 3.38E-05 | Sugar (Inositol Phosphate) |
| ABHD10 | 2.54 | 9.32E-05 | Sugar |
| PPA1 | 2.52 | 3.71E-04 | Sugar (PPi hydrolysis) |
| MPI | 2.10 | 9.18E-04 | Sugar (Mannose) |
| AGK | 2.02 | 4.44E-06 | Sugar (PPP) |
| INPP5E | 1.95 | 1.68E-03 | Sugar (Inositol Phosphate) |
| PIP5K1C | 1.81 | 2.52E-04 | Sugar (Inositol Phosphate) |
| IP6K1 | 1.79 | 5.27E-04 | Sugar (Inositol Phosphate) |
| PIP4K2B | 1.78 | 1.18E-04 | Sugar (Inositol Phosphate) |
| IP6K2 | 1.72 | 7.17E-04 | Sugar (Inositol Phosphate) |
| AGL | 1.63 | 2.57E-02 | Sugar (Glycogen) |
| PI4KA | 1.59 | 1.70E-02 | Sugar (Inositol Phosphate) |
| CMAS | 1.55 | 1.38E-04 | Sugar (Aminosugar) |
| IPMK | 1.51 | 9.12E-03 | Sugar (Inositol Phosphate) |
| PIK3CB | 1.47 | 3.33E-02 | Sugar (Inositol Phosphate) |
| PGM3 | 1.34 | 4.93E-02 | Sugar (Aminosugar) |
| DERA | 1.32 | 2.05E-02 | Sugar (PPP) |
| DCXR | 0.75 | 4.85E-02 | Sugar (Xyulose) |
| FUK | 0.74 | 4.82E-02 | Sugar (Fructose) |
| GNPTG | 0.73 | 3.98E-04 | Sugar (Aminosugar) |
| GRHPR | 0.68 | 6.76E-04 | Sugar |
| UXS1 | 0.67 | 5.49E-03 | Sugar (Nucleotide Sugar) |
| INPPL1 | 0.65 | 3.95E-03 | Sugar (Inositol Phosphate) |
| SORD | 0.60 | 1.76E-03 | Sugar |
| INPP5B | 0.60 | 1.61E-02 | Sugar (Inositol Phosphate) |
| GAA | 0.58 | 3.86E-02 | Sugar (Glycogen) |
| PTEN | 0.56 | 1.51E-02 | Sugar (Inositol Phosphate) |
| GYG1 | 0.54 | 2.60E-05 | Sugar (Glycogen) |
| GALK2 | 0.53 | 1.72E-03 | Sugar (Galactose) |
| PIP4K2A | 0.52 | 1.29E-05 | Sugar (Inositol Phosphate) |
| INPP4B | 0.52 | 1.47E-02 | Sugar (Inositol Phosphate) |
| PIK3C2A | 0.50 | 5.92E-04 | Sugar (Inositol Phosphate) |
| TGDS | 0.47 | 8.81E-05 | Sugar (Nucleotide Sugar) |
| RPE | 0.47 | 1.97E-03 | Sugar (PPP) |
| TALDO1 | 0.46 | 1.68E-03 | Sugar (PPP) |
| GNPDA1 | 0.44 | 1.04E-03 | Sugar (Aminosugar) |
| HYI | 0.42 | 1.23E-02 | Sugar |
| GNPTAB | 0.41 | 6.61E-06 | Sugar (Aminosugar) |
| PGLS | 0.40 | 2.03E-03 | Sugar (PPP) |
| PCK2 | 0.36 | 5.55E-04 | Sugar (gluconeogenesi) |
| SYNJ2 | 0.35 | 1.88E-02 | Sugar (Inositol Phosphate) |
| UAP1L1 | 0.35 | 4.03E-04 | Sugar (Aminosugar) |
| RPIA | 0.34 | 4.23E-03 | Sugar (PPP) |
| UGDH | 0.32 | 2.36E-03 | Sugar (Nucleotide Sugar) |
| PIK3CD | 0.32 | 5.45E-04 | Sugar (Inositol Phosphate) |
| PMM2 | 0.31 | 2.76E-03 | Sugar (Mannose) |
| RENBP | 0.30 | 4.41E-03 | Sugar (Aminosugar) |
| CRYL1 | 0.25 | 5.26E-06 | Sugar |
| GFPT2 | 0.23 | 1.36E-03 | Sugar (Aminosugar) |
| NPL | 0.20 | 3.75E-04 | Sugar (Aminosugar) |
| IMPA2 | 0.16 | 3.09E-04 | Sugar (Inositol Phosphate) |
| GALM | 0.13 | 4.66E-04 | Sugar (Galactose) |
| MGAM | 0.11 | 1.58E-02 | Sugar (Mannose) |
| GMDS | 0.10 | 9.27E-03 | Sugar (Mannose) |
| FBP1 | 0.10 | 3.29E-03 | Sugar (gluconeogenesi) |
| INPP5D | 0.08 | 2.14E-05 | Sugar (Inositol Phosphate) |
| GYG2 | 0.07 | 1.84E-02 | Sugar (Glycogen) |
| PIK3R6 | 0.06 | 2.41E-03 | Sugar (Inositol Phosphate) |
| PIK3CG | 0.04 | 6.51E-05 | Sugar (Inositol Phosphate) |
| ISYNA1 | 0.03 | 7.12E-03 | Sugar (Inositol Phosphate) |
| ITPKB | 0.02 | 4.51E-04 | Sugar (Inositol Phosphate) |
| PIK3R5 | 0.02 | 7.97E-04 | Sugar (Inositol Phosphate) |
| SULT4A1 | 10.40 | 7.95E-04 | Sulfate |
| SRXN1 | 1.43 | 4.02E-04 | Sulfate |
| ARSA | 0.61 | 2.44E-03 | Sulfate |
| DSEL | 0.54 | 3.58E-03 | Sulfate |
| ARSB | 0.45 | 1.84E-02 | Sulfate |
| SULT1A1 | 0.36 | 1.84E-02 | Sulfate |
| SULT1C4 | 0.32 | 7.62E-03 | Sulfate |
| ARSI | 0.24 | 1.36E-04 | Sulfate |
| MPST | 0.23 | 1.21E-06 | Sulfate |
| TST | 0.20 | 1.27E-03 | Sulfate |
| SULT1A2 | 0.13 | 2.76E-02 | Sulfate |
| DSE | 0.07 | 9.03E-03 | Sulfate |
| PAPSS2 | 0.05 | 1.69E-02 | Sulfate |
| SULT1B1 | 0.03 | 1.43E-02 | Sulfate |
| SULT1C2 | 3.52E-03 | 2.40E-03 | Sulfate |
| GOT1 | 10.47 | 5.09E-06 | TCA Cycle |
| OGDHL | 7.11 | 9.75E-06 | TCA Cycle |
| MDH1B | 4.84 | 7.03E-03 | TCA Cycle |
| MDH1 | 3.46 | 7.82E-06 | TCA Cycle |
| L2HGDH | 2.58 | 1.72E-04 | TCA Cycle |
| PDHX | 2.45 | 9.73E-05 | TCA Cycle |
| CS | 2.08 | 5.37E-07 | TCA Cycle |
| DLD | 2.04 | 1.78E-04 | TCA Cycle |
| ACO2 | 1.96 | 5.84E-04 | TCA Cycle |
| SUCLA2 | 1.89 | 1.70E-05 | TCA Cycle |
| IDH3A | 1.89 | 1.14E-04 | TCA Cycle |
| PDHA1 | 1.80 | 9.91E-04 | TCA Cycle |
| ME2 | 1.74 | 4.11E-03 | TCA Cycle |
| FH | 1.71 | 1.62E-04 | TCA Cycle |
| ME1 | 1.59 | 3.14E-03 | TCA Cycle |
| IDH3B | 1.39 | 6.94E-03 | TCA Cycle |
| ACLY | 1.38 | 4.40E-02 | TCA Cycle |
| IDH3G | 1.31 | 1.47E-02 | TCA Cycle |
| PDHB | 0.76 | 3.39E-02 | TCA Cycle |
| SUCLG1 | 0.67 | 1.21E-02 | TCA Cycle |
| ME3 | 0.66 | 9.88E-03 | TCA Cycle |
| D2HGDH | 0.61 | 3.46E-02 | TCA Cycle |
| ACO1 | 0.34 | 1.77E-02 | TCA Cycle |
| IDH1 | 0.29 | 6.73E-03 | TCA Cycle |
| IDH2 | 0.21 | 1.95E-04 | TCA Cycle |
| SUCLG2 | 0.12 | 5.11E-05 | TCA Cycle |
| PDSS1 | 0.65 | 2.71E-02 | Ubiquinone |
| SQRDL | 0.05 | 4.84E-03 | Ubiquinone |
| RDH8 | 375.32 | 1.07E-03 | Vitamin (A) |
| RDH12 | 83.30 | 5.03E-04 | Vitamin (A) |
| SDR16C5 | 50.94 | 7.05E-05 | Vitamin (A) |
| PDXP | 6.58 | 8.60E-04 | Vitamin (B6) |
| PDXK | 3.06 | 5.81E-03 | Vitamin (B6) |
| GPHN | 2.39 | 8.13E-05 | Vitamin (Folate) |
| AKR7A3 | 2.28 | 1.06E-02 | Vitamine (Retinoic acid, Alcohol) |
| AKR1C2 | 2.14 | 9.65E-03 | Vitamine (Retinoic acid, Alcohol) |
| RDH16 | 2.10 | 6.97E-03 | Vitamin (A) |
| BCO2 | 1.87 | 8.71E-03 | Vitamin (A) |
| GGH | 1.84 | 1.27E-02 | Vitamin (Folate) |
| MTHFR | 1.79 | 2.25E-04 | Vitamin (Folate) |
| RFK | 1.77 | 8.51E-04 | Vitamin (Riboflavin) |
| VKORC1L1 | 1.60 | 3.38E-02 | Vitamin (K) |
| ALDH8A1 | 1.44 | 1.32E-02 | Vitamine (Retinoic acid, Alcohol) |
| PNPO | 0.74 | 2.99E-02 | Vitamin (B6) |
| AKR1B1 | 0.66 | 1.82E-03 | Vitamine (Retinoic acid, Alcohol) |
| MTHFS | 0.66 | 1.64E-02 | Vitamin (Folate) |
| MTRR | 0.59 | 1.70E-02 | Vitamin (B12) |
| DHFRL1 | 0.58 | 4.02E-02 | Vitamin (Folate) |
| MTHFSD | 0.56 | 6.25E-04 | Vitamin (Folate) |
| MMADHC | 0.54 | 1.61E-04 | Vitamin (B12) |
| AKR1A1 | 0.53 | 8.83E-05 | Vitamine (Retinoic acid, Alcohol) |
| MTHFD1 | 0.50 | 4.41E-04 | Vitamin (Folate) |
| RETSAT | 0.49 | 7.82E-03 | Vitamin (A) |
| ADH5 | 0.48 | 3.00E-05 | Vitamine (Retinoic acid, Alcohol) |
| MTHFD1L | 0.41 | 1.40E-02 | Vitamin (Folate) |
| FPGS | 0.35 | 2.47E-03 | Vitamin (Folate) |
| SHMT1 | 0.30 | 1.82E-03 | Vitamin (Folate) |
| TPK1 | 0.28 | 2.14E-03 | Vitamin (Thiamine) |
| MTHFD2 | 0.07 | 2.20E-02 | Vitamin (Folate) |
| ALDH1A2 | 0.05 | 7.39E-03 | Vitamine (Retinoic acid, Alcohol) |
| CYP26B1 | 0.02 | 2.74E-02 | Vitamin (A) |
| ALDH1A3 | 0.02 | 2.57E-02 | Vitamine (Retinoic acidl, Aldehyde) |
| ALDH1A1 | 0.01 | 2.79E-02 | Vitamine (Retinoic acid, Alcohol) |
| AKR1B10 | 3.06E-03 | 2.58E-03 | Vitamine (Retinoic acid, Alcohol) |
| ADH1B | 2.93E-03 | 4.85E-02 | Vitamine (Retinoic acid, Alcohol) |

**Table S13. Differentially expressed metabolic genes in periphery between retinas and RPE/choroid.**

| **Gene symbol** | **FC** | **P value** | **Classification (Sub-category/substrate)** |
| --- | --- | --- | --- |
| ABCB11 | 204.77 | 2.02E-05 | ABC Transporter |
| ABCG4 | 81.67 | 3.14E-11 | ABC Transporter |
| ABCC8 | 22.49 | 2.02E-05 | ABC Transporter |
| ABCD2 | 19.69 | 3.59E-08 | ABC Transporter |
| ABCB4 | 11.75 | 1.03E-03 | ABC Transporter |
| ABCA12 | 9.89 | 6.29E-04 | ABC Transporter |
| ABCA7 | 8.23 | 1.62E-06 | ABC Transporter |
| ABCB9 | 6.04 | 1.08E-07 | ABC Transporter |
| ABCG1 | 4.62 | 2.42E-07 | ABC Transporter |
| ABCA3 | 4.57 | 1.50E-09 | ABC Transporter |
| ABCA4 | 3.64 | 2.73E-06 | ABC Transporter |
| ABCG5 | 3.14 | 2.54E-07 | ABC Transporter |
| ABCB8 | 2.09 | 1.34E-09 | ABC Transporter |
| ABCB6 | 1.87 | 8.17E-08 | ABC Transporter |
| ABCC4 | 1.81 | 2.81E-02 | ABC Transporter |
| ABCF3 | 1.58 | 6.57E-08 | ABC Transporter |
| ABCF1 | 1.54 | 6.35E-04 | ABC Transporter |
| ABCB10 | 1.48 | 6.50E-06 | ABC Transporter |
| ABCF2 | 1.37 | 1.14E-03 | ABC Transporter |
| ABCC9 | 0.71 | 3.58E-03 | ABC Transporter |
| ABCC5 | 0.66 | 8.63E-03 | ABC Transporter |
| ABCD3 | 0.50 | 2.84E-05 | ABC Transporter |
| ABCA10 | 0.47 | 6.22E-04 | ABC Transporter |
| ABCA5 | 0.47 | 6.22E-04 | ABC Transporter |
| ABCA6 | 0.47 | 6.22E-04 | ABC Transporter |
| ABCA8 | 0.47 | 6.22E-04 | ABC Transporter |
| ABCA9 | 0.47 | 6.22E-04 | ABC Transporter |
| ABCC1 | 0.46 | 2.04E-06 | ABC Transporter |
| ABCC6 | 0.33 | 7.32E-03 | ABC Transporter |
| ABCA1 | 0.31 | 5.03E-07 | ABC Transporter |
| ABCD1 | 0.24 | 5.18E-04 | ABC Transporter |
| ABCG2 | 0.18 | 1.70E-07 | ABC Transporter |
| ABCC12 | 0.18 | 4.03E-02 | ABC Transporter |
| ABCB1 | 0.14 | 2.79E-05 | ABC Transporter |
| ABCC11 | 0.03 | 1.92E-03 | ABC Transporter |
| ABCB5 | 7.45E-04 | 1.76E-06 | ABC Transporter |
| GAD2 | 221.10 | 1.82E-09 | Amino Acid (Glutamate) |
| DAO | 117.17 | 1.00E-05 | Amino Acid (Alanine) |
| CKMT1A | 47.22 | 4.29E-13 | Amino Acid (Creatine) |
| CKMT1B | 40.11 | 4.10E-08 | Amino Acid (Creatine) |
| CRYM | 37.79 | 5.12E-06 | Amino Acid (Cystathionine) |
| DPEP1 | 30.45 | 1.91E-06 | Amino Acid (Hydrolysis) |
| GLYATL1 | 22.55 | 1.40E-06 | Amino Acid (Glycine) |
| GAD1 | 21.47 | 1.34E-10 | Amino Acid (Glutamate) |
| PAH | 16.23 | 1.22E-04 | Amino Acid (Tyrosine) |
| CKB | 15.66 | 2.51E-12 | Amino Acid (Creatine) |
| GADL1 | 13.88 | 9.17E-04 | Amino Acid (Glutamate) |
| GLYATL2 | 12.70 | 8.30E-06 | Amino Acid (Glycine) |
| GPX5 | 11.62 | 3.03E-03 | Amino Acid (Glutathione) |
| PIPOX | 10.84 | 1.14E-12 | Amino Acid (Lysine) |
| CKM | 10.35 | 9.57E-05 | Amino Acid (Creatine) |
| TPH2 | 9.23 | 3.49E-02 | Amino Acid (Neurotransmitter) |
| PRODH2 | 7.64 | 5.88E-04 | Amino Acid (Proline) |
| TH | 6.21 | 1.44E-03 | Amino Acid (Neurotransmitter) |
| NAT8 | 5.81 | 1.47E-05 | Amino Acid (Acetylation) |
| GPT | 5.53 | 2.39E-06 | Amino Acid (Glutamate, Alaline) |
| OTC | 5.51 | 9.85E-03 | Amino Acid (Arginine, Urea) |
| FTCD | 5.37 | 3.26E-03 | Amino Acid (histidine) |
| NAT8L | 5.13 | 6.08E-04 | Amino Acid (Acetylation) |
| GLS2 | 4.23 | 5.80E-09 | Amino Acid (Glutamate) |
| GPT2 | 3.82 | 1.49E-09 | Amino Acid (Glutamate, Alaline) |
| TAT | 3.41 | 1.94E-03 | Amino Acid (Tyrosine) |
| GGT7 | 3.34 | 2.12E-07 | Amino Acid (Glutathione, Taurine, Hypotaurine) |
| PCYOX1L | 3.23 | 9.85E-08 | Amino Acid (Cysteine) |
| GLS | 3.14 | 1.16E-10 | Amino Acid (Glutamate) |
| DDAH1 | 3.01 | 3.46E-06 | Amino Acid (Arginine, Citrulline) |
| TPH1 | 2.98 | 1.40E-04 | Amino Acid (Neurotransmitter) |
| ASNS | 2.84 | 7.42E-09 | Amino Acid (Asparagine) |
| THNSL1 | 2.51 | 4.15E-11 | Amino Acid (Threonine) |
| GPX2 | 2.50 | 7.66E-03 | Amino Acid (Glutathione) |
| NAT14 | 2.50 | 2.01E-06 | Amino Acid (Acetylation) |
| ADC | 2.36 | 2.28E-09 | Amino Acid (Arginine) |
| GAMT | 2.36 | 8.94E-07 | Amino Acid (Creatine) |
| ALDH5A1 | 2.15 | 3.52E-07 | Amino Acid (Glutamate) |
| QDPR | 2.08 | 3.29E-04 | Amino Acid (BH4) |
| GPX3 | 2.02 | 5.71E-03 | Amino Acid (Glutathione) |
| PCBD2 | 1.98 | 2.07E-06 | Amino Acid (BH4) |
| PTS | 1.98 | 7.55E-05 | Amino Acid (BH4) |
| GLUL | 1.87 | 1.17E-04 | Amino Acid (Glutamate) |
| ADO | 1.82 | 5.38E-05 | Amino Acid (Cysteine) |
| ARG2 | 1.81 | 1.38E-04 | Amino Acid (Arginine, Urea) |
| AASDHPPT | 1.74 | 1.86E-09 | Amino Acid (Lysine) |
| MOCS3 | 1.72 | 1.71E-04 | Amino Acid (Cofactor) |
| AASDH | 1.69 | 8.27E-06 | Amino Acid (Lysine) |
| ACY1 | 1.66 | 3.37E-04 | Amino Acid (Hydrolysis) |
| ENOPH1 | 1.65 | 8.83E-06 | Amino Acid (Cysteine, Methione) |
| AUH | 1.63 | 1.69E-07 | Amino Acid (BCAAs) |
| NFS1 | 1.63 | 1.30E-07 | Amino Acid (Cysteine) |
| ASNSD1 | 1.62 | 3.18E-08 | Amino Acid (Asparagine) |
| SRR | 1.59 | 6.36E-03 | Amino Acid (Neurotransmitter) |
| BCKDHA | 1.48 | 3.16E-04 | Amino Acid (BCAAs) |
| ACYP1 | 1.43 | 3.71E-02 | Amino Acid (Pyruvate) |
| HAGH | 1.42 | 2.61E-04 | Amino Acid (Glutathione) |
| PYCRL | 1.36 | 3.72E-02 | Amino Acid (Proline) |
| PYCR2 | 1.35 | 1.21E-02 | Amino Acid (Proline) |
| GSS | 1.35 | 1.92E-04 | Amino Acid (Glutathione) |
| SMS | 1.33 | 9.69E-04 | Amino Acid (Polyamine) |
| AMD1 | 1.32 | 1.36E-03 | Amino Acid (Methionion, Polyamine) |
| SEPHS2 | 0.75 | 4.29E-05 | Amino Acid (Slenocysteine) |
| THNSL2 | 0.72 | 1.47E-02 | Amino Acid (Threonine) |
| BCKDHB | 0.68 | 1.02E-04 | Amino Acid (BCAAs) |
| GSTM4 | 0.66 | 4.22E-03 | Amino Acid (Glutathione) |
| ASL | 0.66 | 5.55E-08 | Amino Acid (Arginine, Urea) |
| AMDHD2 | 0.65 | 8.55E-04 | Amino Acid (histidine) |
| GCDH | 0.63 | 1.38E-04 | Amino Acid (Lysine, Tryptophan) |
| CCBL2 | 0.62 | 6.11E-06 | Amino Acid (Cysteine) |
| CNDP2 | 0.60 | 3.07E-06 | Amino Acid (Hydroylysis) |
| APEH | 0.60 | 7.02E-07 | Amino Acid (Hydrolysis) |
| PEPD | 0.59 | 6.80E-03 | Amino Acid (Proline) |
| P4HA2 | 0.59 | 2.64E-06 | Amino Acid (Proline) |
| ALDH18A1 | 0.59 | 3.22E-03 | Amino Acid (Glutamate) |
| IVD | 0.58 | 5.07E-10 | Amino Acid (BCAAs) |
| GCLC | 0.58 | 6.89E-04 | Amino Acid (Glutathione) |
| PYCR1 | 0.58 | 1.81E-03 | Amino Acid (Proline) |
| NAT6 | 0.57 | 4.96E-03 | Amino Acid (Acetylation) |
| CPS1 | 0.57 | 7.92E-03 | Amino Acid (Urea) |
| BCAT1 | 0.56 | 2.51E-02 | Amino Acid (BCAAs) |
| NIT2 | 0.55 | 1.61E-08 | Amino Acid (Asparagine) |
| GSTM3 | 0.54 | 2.28E-03 | Amino Acid (Glutathione) |
| PLOD1 | 0.54 | 9.41E-03 | Amino Acid (Proline) |
| GCLM | 0.53 | 4.59E-04 | Amino Acid (Glutathione) |
| NAGS | 0.53 | 1.34E-02 | Amino Acid (Arginine) |
| HIBADH | 0.53 | 2.99E-07 | Amino Acid (BCAAs) |
| SARDH | 0.53 | 1.38E-04 | Amino Acid (Glycine) |
| CBS | 0.51 | 4.19E-04 | Amino Acid (Cysteine) |
| SDSL | 0.50 | 2.71E-02 | Amino Acid (Serine) |
| ALDH6A1 | 0.49 | 1.28E-04 | Amino Acid (BCAAs) |
| P4HA1 | 0.49 | 2.91E-04 | Amino Acid (Proline) |
| MSRB3 | 0.48 | 4.54E-05 | Amino Acid (Methionine) |
| MTAP | 0.47 | 9.83E-08 | Amino Acid (Methionion, Polyamine) |
| CHDH | 0.47 | 3.29E-03 | Amino Acid (Glycine) |
| MCCC2 | 0.47 | 1.41E-07 | Amino Acid (BCAAs) |
| FAHD2B | 0.46 | 1.97E-09 | Amino Acid (Tyrosine) |
| GSTZ1 | 0.45 | 1.89E-03 | Amino Acid (Tryrosine) |
| P4HB | 0.45 | 1.70E-04 | Amino Acid (Proline) |
| GSTP1 | 0.44 | 1.57E-06 | Amino Acid (Glutathione) |
| SRM | 0.44 | 5.76E-03 | Amino Acid (Polyamine) |
| AGXT2L2 | 0.44 | 1.89E-06 | Amino Acid (Alanine) |
| GSTM2 | 0.44 | 1.10E-03 | Amino Acid (Glutathione) |
| LEPREL4 | 0.43 | 1.85E-09 | Amino Acid (Proline) |
| ADHFE1 | 0.43 | 1.72E-03 | Amino Acid (Glutamate) |
| GCAT | 0.43 | 1.17E-06 | Amino Acid (Glycine) |
| GGT1 | 0.40 | 6.30E-06 | Amino Acid (Glutathione, Taurine, Hypotaurine) |
| GPX4 | 0.40 | 2.50E-05 | Amino Acid (Glutathione) |
| GLTP | 0.38 | 8.60E-10 | Amino Acid (Membrane lipid) |
| GSTT1 | 0.38 | 8.02E-03 | Amino Acid (Glutathione) |
| ADI1 | 0.37 | 5.23E-08 | Amino Acid (Methionine) |
| ALDH4A1 | 0.35 | 1.51E-05 | Amino Acid (Proline) |
| ALDH9A1 | 0.35 | 2.81E-04 | Amino acid (GABA) |
| CTH | 0.35 | 6.77E-04 | Amino Acid (Cysteine) |
| BCAT2 | 0.34 | 3.62E-07 | Amino Acid (BCAAs) |
| FAHD2A | 0.32 | 9.85E-06 | Amino Acid (Tyrosine) |
| ALDH1L1 | 0.32 | 4.35E-04 | Amino Acid (One carbon) |
| SPR | 0.29 | 3.41E-06 | Amino Acid (BH4) |
| AASS | 0.28 | 1.94E-09 | Amino Acid (Lysine) |
| FAH | 0.28 | 1.54E-07 | Amino Acid (Tyrosine) |
| GCSH | 0.28 | 2.26E-05 | Amino Acid (Glycine) |
| ASS1 | 0.28 | 1.24E-06 | Amino Acid (Arginine, Urea) |
| MSRA | 0.26 | 1.48E-08 | Amino Acid (Methionine) |
| DMGDH | 0.26 | 3.24E-04 | Amino Acid (Glycine) |
| AFMID | 0.26 | 1.62E-10 | Amino Acid (Tryptophan) |
| GSTO1 | 0.24 | 5.60E-08 | Amino Acid (Glutathione) |
| PHGDH | 0.24 | 5.46E-06 | Amino Acid (Serine) |
| NOS2 | 0.22 | 1.13E-05 | Amino Acid (Arginine) |
| CMBL | 0.22 | 5.06E-07 | Amino Acid (Cysteine) |
| GLUD2 | 0.21 | 3.56E-07 | Amino Acid (Glutamate) |
| PNMT | 0.21 | 3.05E-04 | Amino Acid (Neurotransmitter) |
| NAT1 | 0.20 | 4.62E-06 | Amino Acid (Acetylation) |
| MOCS1 | 0.19 | 1.99E-07 | Amino Acid (Cofactor) |
| GSTK1 | 0.19 | 1.25E-09 | Amino Acid (Glutathione) |
| BHMT | 0.18 | 1.93E-04 | Amino Acid (Glycine) |
| GPX7 | 0.18 | 5.02E-09 | Amino Acid (Glutathione) |
| AOC2 | 0.17 | 2.66E-08 | Amino acid (Dopamine, histamine) |
| AOC3 | 0.17 | 2.66E-08 | Amino acid (Dopamine, histamine) |
| ALDH1L2 | 0.17 | 4.84E-03 | Amino Acid (One carbon) |
| SMOX | 0.16 | 7.39E-04 | Amino Acid (Polyamine) |
| HNMT | 0.14 | 2.38E-06 | Amino Acid (histidine) |
| PRODH | 0.14 | 1.30E-04 | Amino Acid (Proline) |
| SAT1 | 0.13 | 2.07E-03 | Amino Acid (Polyamine) |
| PSAT | 0.12 | 1.57E-06 | Amino Acid (Serine) |
| MGST2 | 0.12 | 1.14E-10 | Amino Acid (Glutathione) |
| CKMT2 | 0.11 | 5.80E-04 | Amino Acid (Creatine) |
| CDO1 | 0.11 | 8.77E-06 | Amino Acid (Cysteine) |
| NAT2 | 0.10 | 2.31E-03 | Amino Acid (Acetylation) |
| ASPG | 0.10 | 1.42E-04 | Amino Acid (Asparagine) |
| GLDC | 0.10 | 4.49E-04 | Amino Acid (Glycine) |
| MAT1A | 0.08 | 5.17E-04 | Amino Acid (Methionine) |
| NOS3 | 0.08 | 3.91E-07 | Amino Acid (Arginine) |
| DPEP2 | 0.08 | 2.76E-04 | Amino Acid (Hydrolysis) |
| GSTA2 | 0.08 | 9.42E-03 | Amino Acid (Glutathione) |
| ARG1 | 0.08 | 3.64E-03 | Amino Acid (Arginine, Urea) |
| KMO | 0.08 | 1.87E-05 | Amino Acid (Tryptophan) |
| ASPA | 0.07 | 4.83E-10 | Amino Acid (Aspartate) |
| GPX8 | 0.07 | 8.84E-06 | Amino Acid (Glutathione) |
| GPX1 | 0.07 | 1.83E-08 | Amino Acid (Glutathione) |
| KYNU | 0.07 | 8.55E-03 | Amino Acid (Tryptophan) |
| HAAO | 0.06 | 4.23E-06 | Amino Acid (Tryptophan, NAD)) |
| INMT | 0.06 | 7.28E-04 | Amino Acid (Tryptophan) |
| HAL | 0.06 | 2.58E-05 | Amino Acid (histidine) |
| HPDL | 0.06 | 7.80E-05 | Amino Acid (Tyrosine) |
| ACY3 | 0.06 | 6.76E-04 | Amino Acid (Aspartate) |
| DPEP3 | 0.05 | 3.33E-02 | Amino Acid (Hydrolysis) |
| BHMT2 | 0.04 | 1.02E-06 | Amino Acid (Glycine) |
| GCH1 | 0.04 | 6.92E-11 | Amino Acid (BH4) |
| MGST1 | 0.03 | 3.53E-03 | Amino Acid (Glutathione) |
| HGD | 0.02 | 3.64E-04 | Amino Acid (Tyrosine) |
| P4HA3 | 0.02 | 2.95E-02 | Amino Acid (Proline) |
| HPD | 0.01 | 3.21E-04 | Amino Acid (Tyrosine) |
| TDO2 | 0.01 | 3.85E-02 | Amino Acid (Tryptophan) |
| GGT5 | 0.01 | 8.18E-04 | Amino Acid (Glutathione, Taurine, Hypotaurine) |
| DCT | 0.01 | 5.46E-04 | Amino Acid (Tryosine, Pigment) |
| CNDP1 | 0.01 | 4.13E-04 | Amino Acid (Hydroylysis) |
| GSTA1 | 0.01 | 7.18E-04 | Amino Acid (Glutathione) |
| TYRP1 | 4.65E-03 | 1.38E-12 | Amino Acid (Tryosine, Pigment) |
| TYR | 2.64E-03 | 1.14E-03 | Amino Acid (Tryosine, Pigment) |
| GSTA3 | 2.54E-03 | 2.61E-04 | Amino Acid (Glutathione) |
| GGTLC2 | 2.49E-04 | 1.69E-02 | Amino Acid (Glutathione, Taurine, Hypotaurine) |
| ACOT7 | 5.27 | 4.41E-08 | CoA |
| PANK4 | 2.91 | 3.63E-09 | CoA |
| ACOT8 | 1.42 | 3.91E-04 | CoA |
| ACOT11 | 0.72 | 2.50E-02 | CoA |
| PANK2 | 0.71 | 2.91E-05 | CoA |
| PPCS | 0.60 | 3.03E-06 | CoA |
| ACOT9 | 0.42 | 2.61E-09 | CoA |
| ACOT6 | 0.32 | 3.42E-03 | CoA |
| HLCS | 0.55 | 2.69E-03 | Cofactor(molybdenum) |
| UGT3A1 | 86.11 | 1.46E-04 | Detox |
| EPX | 9.57 | 2.94E-03 | Detox |
| BCHE | 8.45 | 6.11E-05 | Detox |
| DHDH | 6.55 | 5.55E-06 | Detox |
| TPMT | 1.72 | 1.54E-08 | Detox |
| CES2 | 1.70 | 1.36E-05 | Detox |
| ALDH2 | 0.54 | 7.72E-04 | Detox (Ethanoll) |
| ALDH7A1 | 0.44 | 2.36E-04 | Detox |
| ALDH1B1 | 0.28 | 1.73E-03 | Detox |
| CBR1 | 0.28 | 3.07E-09 | Detox |
| PON2 | 0.25 | 1.18E-07 | Detox |
| EPHX1 | 0.23 | 6.37E-07 | Detox |
| CAT | 0.19 | 4.62E-09 | Detox |
| FMO5 | 0.17 | 6.44E-06 | Detox |
| CES1 | 0.16 | 1.27E-04 | Detox |
| CBR3 | 0.13 | 2.95E-09 | Detox |
| FMO4 | 0.08 | 4.01E-05 | Detox |
| SOD3 | 0.07 | 3.49E-04 | Detox |
| PON3 | 0.07 | 5.57E-04 | Detox |
| FMO2 | 0.04 | 2.66E-03 | Detox |
| UGT3A2 | 0.03 | 7.87E-03 | Detox |
| AADACL4 | 0.02 | 7.67E-03 | Detox |
| FMO3 | 0.01 | 2.83E-06 | Detox |
| FMO1 | 1.90E-03 | 2.04E-02 | Detox |
| NDUFV1 | 2.08 | 1.80E-08 | ETC (Complex I) |
| COX10 | 2.07 | 2.55E-07 | ETC (Complex IV) |
| UQCRH | 1.93 | 4.68E-06 | ETC (Complex III) |
| SDHA | 1.67 | 5.75E-09 | ETC (Complex II) |
| UQCRC2 | 1.66 | 3.50E-07 | ETC (Complex III) |
| NDUFA4 | 1.64 | 9.36E-07 | ETC (Complex I) |
| COX11 | 1.61 | 1.00E-07 | ETC (Complex IV) |
| COX19 | 1.55 | 6.51E-06 | ETC (Complex IV) |
| NDUFA5 | 1.51 | 1.07E-05 | ETC (Complex I) |
| COX6A1 | 1.50 | 1.87E-05 | ETC (Complex IV) |
| COX5A | 1.50 | 1.86E-06 | ETC (Complex IV) |
| UQCRC1 | 1.45 | 2.26E-06 | ETC (Complex III) |
| NDUFA6 | 1.43 | 2.55E-06 | ETC (Complex I) |
| NDUFA9 | 1.41 | 1.56E-05 | ETC (Complex I) |
| NDUFS1 | 1.41 | 3.01E-03 | ETC (Complex I) |
| NDUFV2 | 1.36 | 4.98E-05 | ETC (Complex I) |
| NDUFS2 | 1.36 | 4.91E-03 | ETC (Complex I) |
| NDUFA7 | 1.33 | 3.81E-04 | ETC (Complex I) |
| CYC1 | 1.33 | 1.78E-03 | ETC (Complex III) |
| NDUFA10 | 1.31 | 2.14E-02 | ETC (Complex I) |
| CYCS | 1.31 | 8.26E-03 | ETC (Complex III) |
| COX7A2 | 1.31 | 6.16E-04 | ETC (Complex IV) |
| NDUFB5 | 0.72 | 2.56E-02 | ETC (Complex I) |
| NDUFA13 | 0.70 | 3.26E-04 | ETC (Complex I) |
| ACN9 | 0.69 | 1.48E-04 | ETC (Complex II) |
| NDUFA3 | 0.67 | 3.24E-03 | ETC (Complex I) |
| NDUFA11 | 0.67 | 3.49E-02 | ETC (Complex I) |
| COX7C | 0.59 | 2.63E-07 | ETC (Complex IV) |
| COX7A1 | 0.58 | 3.67E-04 | ETC (Complex IV) |
| SDHD | 0.49 | 7.01E-05 | ETC (Complex II) |
| COX6B2 | 0.45 | 1.85E-04 | ETC (Complex IV) |
| COX15 | 0.38 | 3.16E-02 | ETC (Complex IV) |
| COX4I2 | 0.32 | 3.66E-02 | ETC (Complex IV) |
| FUT9 | 130.14 | 5.57E-07 | Glycan |
| FUT3 | 114.59 | 5.26E-05 | Glycan |
| FUT6 | 109.07 | 1.10E-02 | Glycan |
| LYZL4 | 89.53 | 4.02E-03 | Glycan (Degradation) |
| GALNT13 | 68.25 | 7.36E-11 | Glycan |
| ST8SIA3 | 55.45 | 8.11E-11 | Glycan |
| GAL3ST3 | 53.05 | 6.46E-13 | Glycan (Sulfate) |
| ST6GAL2 | 51.01 | 2.28E-08 | Glycan |
| GLB1L2 | 36.55 | 1.19E-08 | Glycan (Degradation) |
| ST8SIA1 | 34.17 | 8.67E-09 | Glycan |
| B3GAT1 | 31.49 | 5.53E-11 | Glycan |
| B3GALT1 | 24.86 | 4.77E-08 | Glycan |
| CHST8 | 20.98 | 1.39E-05 | Glycan (Sulfate) |
| ST8SIA2 | 16.75 | 1.77E-04 | Glycan |
| FUT5 | 16.72 | 2.13E-05 | Glycan |
| NEU4 | 15.10 | 3.72E-03 | Glycan (Degradation) |
| MANEAL | 14.16 | 5.65E-11 | Glycan (Degradation) |
| ST8SIA5 | 13.94 | 9.78E-09 | Glycan |
| GBA3 | 13.51 | 6.54E-04 | Glycan |
| GAL3ST1 | 12.84 | 3.35E-05 | Glycan (Sulfate) |
| B3GALT2 | 11.24 | 4.52E-10 | Glycan |
| MGAT4C | 10.66 | 4.87E-07 | Glycan |
| GALNTL5 | 10.64 | 1.07E-02 | Glycan |
| MGAT3 | 7.80 | 3.02E-08 | Glycan |
| WBSCR17 | 6.93 | 1.87E-06 | Glycan |
| CHST5 | 6.85 | 1.25E-07 | Glycan (Sulfate) |
| B4GALT6 | 6.58 | 2.47E-10 | Glycan |
| B3GAT2 | 5.89 | 1.40E-08 | Glycan |
| CHST2 | 5.69 | 1.52E-07 | Glycan (Sulfate) |
| B4GALNT1 | 5.37 | 1.98E-06 | Glycan |
| GPLD1 | 4.67 | 1.85E-10 | Glycan (Anchor) |
| GLT25D2 | 4.66 | 4.23E-08 | Glycan |
| A4GNT | 4.64 | 2.21E-02 | Glycan |
| GALNT8 | 4.64 | 3.61E-08 | Glycan |
| GLB1L3 | 4.57 | 8.35E-04 | Glycan (Degradation) |
| GAL3ST2 | 4.07 | 2.12E-03 | Glycan (Sulfate) |
| B3GALNT1 | 3.73 | 1.21E-08 | Glycan |
| B3GNT4 | 3.68 | 2.66E-04 | Glycan |
| GBA2 | 3.60 | 1.68E-08 | Glycan |
| MAN2A2 | 3.59 | 1.35E-06 | Glycan (Degradation) |
| GALNT9 | 3.42 | 5.79E-05 | Glycan |
| ALG10B | 3.41 | 1.26E-10 | Glycan |
| LARGE | 3.34 | 8.27E-10 | Glycan |
| CHST9 | 3.21 | 8.43E-04 | Glycan (Sulfate) |
| CHST4 | 3.20 | 2.87E-02 | Glycan (Sulfate) |
| ST6GALNAC5 | 2.96 | 1.06E-09 | Glycan |
| MGAT5B | 2.77 | 1.15E-03 | Glycan |
| B4GALNT4 | 2.59 | 6.21E-07 | Glycan |
| ALG10 | 2.55 | 1.03E-05 | Glycan |
| B3GNT1 | 2.52 | 1.21E-09 | Glycan |
| GLT1D1 | 2.44 | 3.14E-05 | Glycan |
| IDS | 2.34 | 2.79E-07 | Glycan (Degradation) |
| ST3GAL3 | 2.30 | 2.53E-08 | Glycan |
| PIGZ | 2.28 | 1.11E-05 | Glycan (Anchor) |
| PIGM | 2.24 | 1.43E-06 | Glycan (Anchor) |
| POMT2 | 2.14 | 2.89E-07 | Glycan |
| B3GALTL | 2.12 | 5.53E-05 | Glycan |
| MGAT5 | 2.05 | 1.02E-04 | Glycan |
| XYLT1 | 1.95 | 3.23E-06 | Glycan |
| B3GALT5 | 1.95 | 1.24E-02 | Glycan |
| CHST7 | 1.92 | 5.87E-06 | Glycan (Sulfate) |
| FUT4 | 1.74 | 5.88E-05 | Glycan |
| FUT2 | 1.72 | 3.32E-03 | Glycan |
| DHDDS | 1.65 | 5.62E-05 | Glycan |
| POMT1 | 1.65 | 7.79E-06 | Glycan |
| B4GALT2 | 1.64 | 2.60E-06 | Glycan |
| HEXDC | 1.49 | 5.30E-03 | Glycan (Degradation) |
| HAS3 | 1.49 | 2.10E-03 | Glycan |
| PIGS | 1.48 | 9.26E-06 | Glycan (Anchor) |
| POFUT2 | 1.44 | 2.85E-04 | Glycan |
| GALNT7 | 1.42 | 3.76E-04 | Glycan |
| FN3KRP | 1.38 | 8.62E-05 | Glycan |
| GALNT4 | 1.37 | 4.51E-04 | Glycan |
| CHST10 | 1.36 | 1.12E-04 | Glycan (Sulfate) |
| POMGNT1 | 1.33 | 6.85E-06 | Glycan |
| ALG9 | 1.32 | 3.81E-04 | Glycan |
| FUT10 | 1.31 | 4.35E-02 | Glycan |
| PIGX | 1.30 | 6.62E-05 | Glycan (Anchor) |
| MGAT2 | 0.77 | 1.51E-03 | Glycan |
| GLT8D1 | 0.75 | 1.61E-03 | Glycan |
| LYG1 | 0.72 | 2.63E-04 | Glycan (Degradation) |
| GLB1 | 0.72 | 4.36E-05 | Glycan (Degradation) |
| ALG6 | 0.71 | 1.51E-03 | Glycan |
| MGAT4B | 0.71 | 1.15E-02 | Glycan |
| UST | 0.70 | 6.82E-03 | Glycan (Sulfate) |
| DOLK | 0.69 | 1.41E-04 | Glycan |
| NEU3 | 0.68 | 8.87E-05 | Glycan (Degradation) |
| ALG14 | 0.68 | 2.13E-03 | Glycan |
| IDUA | 0.65 | 1.50E-02 | Glycan (Degradation) |
| B4GALT3 | 0.65 | 1.28E-03 | Glycan |
| DPAGT1 | 0.65 | 1.53E-02 | Glycan |
| PIGV | 0.64 | 1.08E-03 | Glycan (Anchor) |
| EDEM2 | 0.64 | 7.03E-04 | Glycan (Degradation) |
| GTDC1 | 0.64 | 7.05E-06 | Glycan |
| B4GALT4 | 0.62 | 7.25E-05 | Glycan |
| PIGP | 0.62 | 1.19E-04 | Glycan (Anchor) |
| ALG12 | 0.60 | 9.85E-04 | Glycan |
| EDEM1 | 0.59 | 3.20E-02 | Glycan (Degradation) |
| CHST12 | 0.59 | 7.49E-03 | Glycan (Sulfate) |
| MGAT1 | 0.58 | 3.63E-04 | Glycan |
| ST3GAL5 | 0.58 | 9.90E-03 | Glycan |
| DDOST | 0.58 | 1.09E-02 | Glycan |
| GUSB | 0.57 | 1.89E-03 | Glycan (Degradation) |
| HYAL3 | 0.57 | 4.96E-03 | Glycan (Degradation) |
| B3GNT8 | 0.57 | 2.39E-02 | Glycan |
| ALG2 | 0.55 | 2.34E-07 | Glycan |
| NAGA | 0.54 | 3.07E-06 | Glycan |
| B3GALNT2 | 0.54 | 5.04E-05 | Glycan |
| MANBAL | 0.54 | 1.88E-08 | Glycan (Degradation) |
| B3GNT2 | 0.52 | 3.01E-07 | Glycan |
| MANBA | 0.51 | 6.39E-06 | Glycan (Degradation) |
| POFUT1 | 0.51 | 3.50E-04 | Glycan |
| PIGT | 0.50 | 7.97E-07 | Glycan (Anchor) |
| NEU1 | 0.50 | 1.99E-06 | Glycan (Degradation) |
| GLA | 0.50 | 5.83E-06 | Glycan (Degradation) |
| GALNT1 | 0.49 | 2.03E-04 | Glycan |
| GBA | 0.49 | 1.53E-07 | Glycan |
| MGAT4A | 0.49 | 4.32E-06 | Glycan |
| FUT8 | 0.48 | 7.57E-05 | Glycan |
| ALG8 | 0.48 | 3.53E-04 | Glycan |
| SGSH | 0.47 | 6.45E-06 | Glycan (Sulfate) |
| ALG5 | 0.47 | 4.38E-08 | Glycan |
| ST3GAL2 | 0.47 | 6.02E-06 | Glycan |
| NAGPA | 0.46 | 3.36E-07 | Glycan |
| GCNT2 | 0.46 | 3.66E-05 | Glycan |
| SULF1 | 0.46 | 1.29E-06 | Glycan (Sulfate) |
| CHSY1 | 0.44 | 8.24E-04 | Glycan (Sulfate) |
| MAN2B2 | 0.43 | 3.09E-06 | Glycan (Degradation) |
| CSGALNACT2 | 0.43 | 1.89E-04 | Glycan (Sulfate) |
| MAN1C1 | 0.42 | 3.09E-05 | Glycan (Degradation) |
| CSGALNACT1 | 0.41 | 1.27E-03 | Glycan (Sulfate) |
| ST6GALNAC3 | 0.40 | 2.09E-04 | Glycan |
| DAD1 | 0.40 | 2.67E-05 | Glycan |
| FUT7 | 0.40 | 9.75E-03 | Glycan |
| GALNT11 | 0.40 | 4.41E-03 | Glycan |
| CHST3 | 0.38 | 2.15E-03 | Glycan (Sulfate) |
| HEXA | 0.37 | 3.04E-13 | Glycan (Degradation) |
| GLB1L | 0.36 | 2.96E-07 | Glycan (Degradation) |
| GLT25D1 | 0.35 | 4.38E-05 | Glycan |
| HEXB | 0.34 | 4.55E-12 | Glycan (Degradation) |
| AGA | 0.33 | 1.29E-03 | Glycan (Degradation) |
| SULF2 | 0.32 | 4.74E-06 | Glycan (Sulfate) |
| B3GNT5 | 0.32 | 1.37E-07 | Glycan |
| ST6GALNAC2 | 0.30 | 5.19E-04 | Glycan |
| B4GALT5 | 0.30 | 2.84E-03 | Glycan |
| ST8SIA4 | 0.29 | 5.72E-08 | Glycan |
| B3GNT9 | 0.28 | 1.21E-04 | Glycan |
| CTBS | 0.27 | 5.03E-08 | Glycan |
| GALNT12 | 0.27 | 1.32E-06 | Glycan |
| B4GALT1 | 0.27 | 1.32E-09 | Glycan |
| FUT1 | 0.27 | 3.02E-05 | Glycan |
| MAN1A1 | 0.25 | 1.60E-05 | Glycan (Degradation) |
| MAN2B1 | 0.24 | 1.22E-07 | Glycan (Degradation) |
| ST6GALNAC4 | 0.23 | 1.32E-03 | Glycan |
| HYAL1 | 0.23 | 4.01E-05 | Glycan (Degradation) |
| CHSY3 | 0.22 | 7.35E-03 | Glycan (Sulfate) |
| GNS | 0.21 | 7.45E-10 | Glycan (Degradation) |
| CHST14 | 0.20 | 4.50E-06 | Glycan (Sulfate) |
| GALNT10 | 0.20 | 5.37E-05 | Glycan |
| ABO | 0.19 | 5.92E-04 | Glycan |
| FUCA1 | 0.18 | 8.13E-06 | Glycan (Degradation) |
| GAL3ST4 | 0.17 | 1.09E-04 | Glycan (Sulfate) |
| NAGLU | 0.17 | 4.60E-10 | Glycan (Degradation) |
| ST3GAL6 | 0.17 | 8.56E-09 | Glycan |
| ST6GAL1 | 0.16 | 1.10E-08 | Glycan |
| MFNG | 0.15 | 3.96E-06 | Glycan |
| ST3GAL4 | 0.13 | 1.40E-09 | Glycan |
| GCNT1 | 0.12 | 4.54E-05 | Glycan |
| CHST11 | 0.12 | 1.94E-11 | Glycan (Sulfate) |
| GCNT4 | 0.12 | 1.22E-04 | Glycan |
| HYAL2 | 0.11 | 4.54E-09 | Glycan (Degradation) |
| CHST13 | 0.10 | 1.75E-05 | Glycan (Sulfate) |
| FUCA2 | 0.10 | 4.82E-12 | Glycan (Degradation) |
| LYZ | 0.09 | 4.03E-05 | Glycan (Degradation) |
| MAN2A1 | 0.08 | 6.18E-06 | Glycan (Degradation) |
| GBGT1 | 0.07 | 2.31E-08 | Glycan |
| ST8SIA6 | 0.07 | 6.11E-07 | Glycan |
| GALNT6 | 0.06 | 5.32E-06 | Glycan |
| A4GALT | 0.05 | 2.76E-03 | Glycan |
| GLT8D2 | 0.04 | 9.16E-08 | Glycan |
| HAS2 | 0.01 | 3.92E-03 | Glycan |
| HKDC1 | 100.07 | 1.39E-05 | Glycolysis |
| HK2 | 24.32 | 7.78E-05 | Glycolysis |
| ENO2 | 24.30 | 2.12E-11 | Glycolysis |
| PFKP | 12.69 | 2.99E-05 | Glycolysis |
| ALDOC | 6.98 | 4.44E-10 | Glycolysis |
| ALDOB | 4.60 | 1.69E-03 | Glycolysis |
| PGAM2 | 3.72 | 9.68E-08 | Glycolysis |
| PFKFB4 | 3.61 | 1.15E-04 | Glycolysis |
| PGM2 | 3.18 | 5.56E-08 | Glycolysis |
| PFKM | 3.01 | 2.15E-15 | Glycolysis |
| GPI | 2.95 | 1.40E-08 | Glycolysis |
| HK1 | 2.94 | 9.80E-10 | Glycolysis |
| PGK1 | 2.89 | 2.47E-06 | Glycolysis |
| TPI1 | 2.71 | 5.16E-07 | Glycolysis |
| PGAM1 | 2.69 | 8.10E-09 | Glycolysis |
| GCK | 2.69 | 1.39E-05 | Glycolysis |
| LDHA | 2.48 | 4.26E-04 | Glycolysis |
| PFKFB2 | 2.41 | 2.02E-06 | Glycolysis |
| ENO3 | 2.38 | 1.62E-04 | Glycolysis |
| ENO1 | 2.29 | 4.97E-04 | Glycolysis |
| GAPDH | 2.22 | 7.42E-06 | Glycolysis |
| PKLR | 2.08 | 5.73E-03 | Glycolysis |
| PGM2L1 | 1.91 | 1.64E-02 | Glycolysis |
| PGM1 | 1.43 | 1.46E-04 | Glycolysis |
| DLAT | 1.31 | 3.92E-04 | Glycolysis |
| BPGM | 1.30 | 1.91E-04 | Glycolysis |
| PFKL | 0.76 | 1.54E-02 | Glycolysis |
| LDHB | 0.69 | 1.94E-04 | Glycolysis |
| PGM5 | 0.42 | 1.29E-04 | Glycolysis |
| LDHD | 0.30 | 6.42E-06 | Glycolysis |
| ADPGK | 0.25 | 1.89E-08 | Glycolysis |
| PFKFB3 | 0.19 | 1.31E-07 | Glycolysis |
| HK3 | 0.02 | 7.18E-05 | Glycolysis |
| GAPDHS | 0.01 | 1.01E-05 | Glycolysis |
| GLOD5 | 28.55 | 3.24E-07 | Glyoxylate |
| HS3ST4 | 50.55 | 6.27E-06 | Heparin Sulfate |
| HS3ST3B1 | 10.79 | 7.00E-11 | Heparin Sulfate |
| HS6ST2 | 8.09 | 1.40E-06 | Heparin Sulfate |
| HS6ST3 | 7.00 | 1.30E-04 | Heparin Sulfate |
| NDST3 | 6.24 | 5.38E-10 | Heparin Sulfate |
| HS6ST1 | 5.38 | 1.71E-10 | Heparin Sulfate |
| HS3ST3A1 | 4.06 | 7.33E-04 | Heparin Sulfate |
| NDST1 | 2.52 | 2.66E-06 | Heparin Sulfate |
| HS2ST1 | 1.52 | 4.50E-05 | Heparin Sulfate |
| EXTL1 | 1.45 | 4.56E-02 | Heparin Sulfate |
| GLCE | 1.39 | 1.61E-02 | Heparin Sulfate |
| EXT1 | 0.34 | 6.20E-05 | Heparin Sulfate |
| HPSE | 0.30 | 1.79E-02 | Heparin Sulfate |
| NDST4 | 0.10 | 4.98E-03 | Heparin Sulfate |
| HPSE2 | 0.09 | 1.47E-02 | Heparin Sulfate |
| CHRNB3 | 318.90 | 1.03E-04 | Ion Transport |
| SLN | 221.68 | 1.86E-02 | Ion Transport |
| KCNJ9 | 209.60 | 1.32E-08 | Ion Transport |
| GLRA1 | 201.92 | 9.27E-07 | Ion Transport |
| CACNG5 | 196.02 | 9.26E-11 | Ion Transport |
| GRIN1 | 187.90 | 8.93E-06 | Ion Transport |
| CACNG7 | 176.18 | 4.86E-08 | Ion Transport |
| CNGA3 | 163.01 | 7.33E-13 | Ion Transport |
| ATP1A4 | 161.18 | 4.70E-04 | Ion Transport |
| KCNC2 | 158.41 | 1.81E-04 | Ion Transport |
| KCNB2 | 158.24 | 8.62E-08 | Ion Transport |
| CHRNA2 | 156.13 | 2.15E-11 | Ion Transport |
| GLRA2 | 155.98 | 1.74E-07 | Ion Transport |
| SCN1A | 155.49 | 5.80E-06 | Ion Transport |
| GABRA1 | 154.67 | 4.39E-10 | Ion Transport |
| KCNC1 | 149.68 | 4.31E-08 | Ion Transport |
| KCNG4 | 144.35 | 3.98E-10 | Ion Transport |
| CHRNA6 | 140.30 | 4.67E-05 | Ion Transport |
| CACNG2 | 139.40 | 4.14E-08 | Ion Transport |
| CNGB1 | 138.52 | 6.83E-06 | Ion Transport |
| CNGA1 | 135.87 | 1.15E-09 | Ion Transport |
| TF | 130.12 | 6.07E-06 | Ion Transport |
| CACNG3 | 123.72 | 1.55E-12 | Ion Transport |
| SLC39A2 | 123.32 | 2.09E-03 | Ion Transport |
| TRPC7 | 123.30 | 5.04E-10 | Ion Transport |
| GABRR1 | 113.55 | 1.84E-12 | Ion Transport |
| SLC24A2 | 113.11 | 3.70E-09 | Ion Transport |
| GABRR3 | 108.82 | 2.00E-11 | Ion Transport |
| KCNV2 | 106.19 | 8.31E-11 | Ion Transport |
| GRIA1 | 104.52 | 1.46E-09 | Ion Transport |
| HCN2 | 99.51 | 2.77E-08 | Ion Transport |
| GABRA3 | 96.40 | 3.49E-07 | Ion Transport |
| GRIA4 | 93.36 | 4.93E-11 | Ion Transport |
| KCNJ10 | 89.12 | 1.35E-10 | Ion Transport |
| CHRND | 87.05 | 2.73E-03 | Ion Transport |
| KCNH6 | 86.92 | 2.75E-08 | Ion Transport |
| KCNK9 | 86.13 | 8.85E-06 | Ion Transport |
| GABRA5 | 84.05 | 4.00E-09 | Ion Transport |
| KCNJ6 | 82.86 | 6.88E-09 | Ion Transport |
| CHRNB2 | 76.93 | 1.24E-08 | Ion Transport |
| PKD2L1 | 73.84 | 3.46E-05 | Ion Transport |
| KCNB1 | 73.34 | 3.19E-07 | Ion Transport |
| GABRG1 | 72.16 | 7.89E-08 | Ion Transport |
| SLC39A5 | 69.61 | 5.08E-11 | Ion Transport |
| GABRG2 | 68.88 | 2.34E-11 | Ion Transport |
| HCN1 | 66.41 | 1.42E-10 | Ion Transport |
| GRIN2B | 64.69 | 5.28E-06 | Ion Transport |
| CACNA1E | 60.47 | 9.98E-08 | Ion Transport |
| CACNG8 | 59.09 | 1.68E-09 | Ion Transport |
| CACNA1G | 59.03 | 3.21E-10 | Ion Transport |
| KCNJ4 | 57.60 | 2.67E-08 | Ion Transport |
| GRIK1 | 57.20 | 2.53E-11 | Ion Transport |
| KCNA1 | 56.78 | 5.03E-04 | Ion Transport |
| KCND2 | 53.96 | 5.56E-08 | Ion Transport |
| GABRR2 | 50.69 | 3.37E-11 | Ion Transport |
| KCNQ2 | 50.42 | 1.61E-06 | Ion Transport |
| CACNA2D4 | 46.86 | 2.94E-06 | Ion Transport |
| SCN3B | 45.20 | 4.23E-10 | Ion Transport |
| HCN4 | 45.15 | 8.90E-07 | Ion Transport |
| KCNA4 | 42.76 | 3.72E-08 | Ion Transport |
| CACNA1I | 41.35 | 5.49E-09 | Ion Transport |
| SLC8A2 | 40.75 | 1.22E-11 | Ion Transport |
| SLC24A1 | 39.31 | 5.84E-10 | Ion Transport |
| SCN2A | 37.76 | 3.78E-12 | Ion Transport |
| SLC12A5 | 37.57 | 1.07E-11 | Ion Transport |
| SLC24A4 | 37.22 | 4.11E-11 | Ion Transport |
| KCNH7 | 37.04 | 7.20E-09 | Ion Transport |
| GLRA3 | 36.80 | 1.84E-07 | Ion Transport |
| CACNG4 | 34.70 | 3.98E-08 | Ion Transport |
| GABRG3 | 31.76 | 8.78E-08 | Ion Transport |
| CACNA1F | 30.90 | 1.15E-08 | Ion Transport |
| TRPC5 | 30.07 | 1.00E-03 | Ion Transport |
| CACNA1B | 29.81 | 1.33E-09 | Ion Transport |
| CLCNKA | 29.63 | 2.54E-06 | Ion Transport |
| KCNH2 | 29.31 | 8.86E-09 | Ion Transport |
| GRIA3 | 28.62 | 1.38E-08 | Ion Transport |
| GABRA6 | 28.50 | 3.11E-06 | Ion Transport |
| GRID2 | 27.68 | 2.31E-09 | Ion Transport |
| SLC8A3 | 26.90 | 2.33E-15 | Ion Transport |
| CACNA1A | 26.89 | 1.44E-07 | Ion Transport |
| SCN8A | 25.40 | 1.72E-07 | Ion Transport |
| GABRB3 | 25.36 | 1.67E-12 | Ion Transport |
| GRIN2C | 24.86 | 4.75E-07 | Ion Transport |
| KCNK3 | 24.01 | 2.03E-12 | Ion Transport |
| GABRA2 | 23.40 | 7.40E-09 | Ion Transport |
| GRIN2A | 20.88 | 1.13E-04 | Ion Transport |
| KCNF1 | 20.48 | 6.51E-09 | Ion Transport |
| CACNA1S | 19.79 | 2.59E-03 | Ion Transport |
| KCNA2 | 19.38 | 4.30E-06 | Ion Transport |
| KCNJ11 | 18.96 | 2.25E-08 | Ion Transport |
| KCTD16 | 18.76 | 2.01E-08 | Ion Transport |
| TTYH1 | 17.05 | 9.20E-13 | Ion Transport |
| TRPV6 | 16.62 | 2.08E-11 | Ion Transport |
| BEST3 | 15.92 | 9.71E-07 | Ion Transport |
| SLC9A2 | 15.42 | 1.12E-06 | Ion Transport |
| KCNJ16 | 15.38 | 3.40E-06 | Ion Transport |
| KCNH8 | 15.35 | 1.48E-08 | Ion Transport |
| ATP2A1 | 14.88 | 1.88E-08 | Ion Transport |
| GLRA4 | 14.37 | 9.30E-05 | Ion Transport |
| ATP2B3 | 14.33 | 2.91E-07 | Ion Transport |
| CACNG6 | 14.11 | 1.31E-04 | Ion Transport |
| GABRB2 | 13.17 | 5.97E-09 | Ion Transport |
| SCN5A | 12.63 | 1.51E-09 | Ion Transport |
| GRIK4 | 12.59 | 1.81E-06 | Ion Transport |
| CHRNA4 | 12.27 | 1.81E-07 | Ion Transport |
| KCNJ3 | 12.02 | 7.32E-07 | Ion Transport |
| ATP1A3 | 11.75 | 8.66E-11 | Ion Transport |
| KCNV1 | 11.67 | 4.09E-05 | Ion Transport |
| GRIN3A | 10.82 | 4.60E-08 | Ion Transport |
| SCN2B | 10.71 | 3.67E-07 | Ion Transport |
| GABRD | 10.31 | 1.05E-07 | Ion Transport |
| KCNC3 | 9.54 | 6.86E-09 | Ion Transport |
| RYR2 | 9.25 | 3.79E-09 | Ion Transport |
| CATSPER2 | 9.13 | 2.95E-06 | Ion Transport |
| KCNQ3 | 8.87 | 3.74E-06 | Ion Transport |
| SCN4B | 8.83 | 2.82E-04 | Ion Transport |
| ATP2B1 | 8.66 | 4.04E-12 | Ion Transport |
| TRPM8 | 8.57 | 5.27E-06 | Ion Transport |
| SLC8A1 | 8.57 | 2.67E-09 | Ion Transport |
| SLC9A7 | 8.24 | 2.06E-08 | Ion Transport |
| GRIN2D | 7.82 | 4.25E-07 | Ion Transport |
| SCNN1G | 7.44 | 1.37E-02 | Ion Transport |
| KCTD8 | 7.26 | 4.38E-11 | Ion Transport |
| CNGA4 | 6.67 | 2.53E-08 | Ion Transport |
| TRPC3 | 6.54 | 1.83E-06 | Ion Transport |
| ATP1B2 | 6.47 | 2.12E-12 | Ion Transport |
| SLC12A6 | 6.13 | 5.66E-09 | Ion Transport |
| KCTD7 | 5.99 | 2.73E-08 | Ion Transport |
| GRIA2 | 5.76 | 9.59E-09 | Ion Transport |
| KCNC4 | 5.34 | 8.20E-11 | Ion Transport |
| SLC13A4 | 5.16 | 2.08E-04 | Ion Transport |
| ATP1A2 | 5.05 | 3.20E-03 | Ion Transport |
| GRID1 | 5.02 | 1.72E-08 | Ion Transport |
| CACNA2D2 | 4.92 | 2.26E-10 | Ion Transport |
| GRIK2 | 4.73 | 7.60E-07 | Ion Transport |
| GRIK5 | 4.69 | 3.09E-11 | Ion Transport |
| GRIN3B | 4.65 | 1.92E-04 | Ion Transport |
| KCNA6 | 4.64 | 3.61E-08 | Ion Transport |
| GABRB1 | 4.58 | 1.17E-05 | Ion Transport |
| KCND3 | 4.55 | 6.10E-08 | Ion Transport |
| P2RX3 | 4.40 | 9.40E-05 | Ion Transport |
| PKD1 | 4.40 | 5.36E-07 | Ion Transport |
| P2RX6 | 4.35 | 8.40E-09 | Ion Transport |
| TRPM5 | 4.24 | 3.28E-02 | Ion Transport |
| CACNB1 | 4.19 | 8.25E-07 | Ion Transport |
| KCTD19 | 4.14 | 1.56E-04 | Ion Transport |
| KCNS1 | 4.07 | 9.12E-04 | Ion Transport |
| CACNB2 | 3.98 | 2.80E-07 | Ion Transport |
| SLC9A4 | 3.97 | 1.80E-05 | Ion Transport |
| KCNS2 | 3.97 | 5.89E-03 | Ion Transport |
| KCNK1 | 3.95 | 8.46E-06 | Ion Transport |
| CLCNKB | 3.88 | 1.62E-04 | Ion Transport |
| ATP7B | 3.81 | 1.36E-12 | Ion Transport |
| KCNA5 | 3.77 | 1.04E-06 | Ion Transport |
| ATP4B | 3.70 | 9.03E-04 | Ion Transport |
| KCNJ12 | 3.67 | 3.68E-06 | Ion Transport |
| KCNN1 | 3.59 | 7.43E-06 | Ion Transport |
| CHRNE | 3.44 | 1.80E-03 | Ion Transport |
| ATP4A | 3.40 | 3.05E-02 | Ion Transport |
| HCN3 | 3.39 | 2.90E-07 | Ion Transport |
| KCNMA1 | 3.20 | 6.25E-06 | Ion Transport |
| TRPC6 | 3.16 | 1.32E-05 | Ion Transport |
| CACNB4 | 3.08 | 3.61E-05 | Ion Transport |
| KCNJ2 | 2.98 | 4.22E-06 | Ion Transport |
| TRPC1 | 2.98 | 2.30E-07 | Ion Transport |
| GRIK3 | 2.96 | 1.66E-05 | Ion Transport |
| SLC24A3 | 2.91 | 3.44E-04 | Ion Transport |
| KCNT1 | 2.87 | 2.07E-03 | Ion Transport |
| SCN3A | 2.86 | 3.57E-04 | Ion Transport |
| VDAC1 | 2.66 | 1.18E-11 | Ion Transport |
| KCNAB1 | 2.62 | 3.78E-05 | Ion Transport |
| SCNN1D | 2.60 | 1.61E-07 | Ion Transport |
| GLRB | 2.54 | 3.99E-06 | Ion Transport |
| CLCN6 | 2.52 | 8.05E-09 | Ion Transport |
| CLCN3 | 2.45 | 1.19E-07 | Ion Transport |
| ATP11B | 2.43 | 1.45E-07 | Ion Transport |
| CHRNA10 | 2.36 | 1.69E-03 | Ion Transport |
| ORAI2 | 2.31 | 1.56E-04 | Ion Transport |
| RYR1 | 2.29 | 4.31E-06 | Ion Transport |
| KCTD13 | 2.26 | 8.82E-08 | Ion Transport |
| SLC9A6 | 2.25 | 1.20E-09 | Ion Transport |
| KCNG3 | 2.23 | 1.82E-02 | Ion Transport |
| KCNK10 | 2.20 | 3.48E-02 | Ion Transport |
| CACNA2D1 | 2.17 | 1.98E-05 | Ion Transport |
| CLCN4 | 2.16 | 5.76E-06 | Ion Transport |
| KCNE1L | 2.14 | 1.88E-02 | Ion Transport |
| KCNA3 | 2.07 | 2.96E-05 | Ion Transport |
| SLC30A9 | 2.05 | 8.19E-09 | Ion Transport |
| KCNN4 | 2.03 | 1.12E-04 | Ion Transport |
| ITPR3 | 2.01 | 5.75E-06 | Ion Transport |
| CACNA1H | 2.00 | 1.76E-04 | Ion Transport |
| CATSPER3 | 1.98 | 2.07E-06 | Ion Transport |
| ATP11A | 1.93 | 3.09E-06 | Ion Transport |
| CACNB3 | 1.87 | 3.65E-06 | Ion Transport |
| MFSD8 | 1.82 | 2.46E-07 | Ion Transport |
| KCNAB2 | 1.67 | 1.55E-03 | Ion Transport |
| KCTD15 | 1.62 | 2.66E-03 | Ion Transport |
| SLC39A10 | 1.62 | 1.31E-04 | Ion Transport |
| VDAC3 | 1.57 | 1.14E-05 | Ion Transport |
| ATP2B4 | 1.56 | 1.02E-02 | Ion Transport |
| VDAC2 | 1.48 | 6.99E-04 | Ion Transport |
| KCNMB3 | 1.45 | 2.40E-02 | Ion Transport |
| MFSD5 | 1.43 | 1.00E-02 | Ion Transport |
| ITPR2 | 1.42 | 2.34E-03 | Ion Transport |
| SLC41A1 | 1.32 | 4.15E-06 | Ion Transport |
| SLC39A6 | 0.77 | 3.61E-02 | Ion Transport |
| SLC30A6 | 0.77 | 3.09E-03 | Ion Transport |
| SLC39A9 | 0.74 | 2.49E-06 | Ion Transport |
| TPCN1 | 0.73 | 3.32E-03 | Ion Transport |
| KCTD20 | 0.70 | 2.63E-03 | Ion Transport |
| KCTD18 | 0.65 | 2.06E-05 | Ion Transport |
| TRPM2 | 0.63 | 2.21E-02 | Ion Transport |
| TMCO3 | 0.61 | 1.51E-05 | Ion Transport |
| SLC39A7 | 0.59 | 1.80E-06 | Ion Transport |
| NALCN | 0.57 | 2.35E-02 | Ion Transport |
| SLC31A1 | 0.56 | 3.11E-08 | Ion Transport |
| KCTD11 | 0.54 | 1.62E-07 | Ion Transport |
| SLC12A9 | 0.53 | 1.57E-03 | Ion Transport |
| TCN2 | 0.52 | 3.59E-04 | Ion Transport |
| SLC30A7 | 0.51 | 5.98E-06 | Ion Transport |
| PKD2 | 0.50 | 1.85E-04 | Ion Transport |
| SLC30A1 | 0.48 | 1.81E-03 | Ion Transport |
| CLCN5 | 0.47 | 9.79E-05 | Ion Transport |
| TPCN2 | 0.47 | 4.15E-03 | Ion Transport |
| CNGB3 | 0.46 | 3.17E-03 | Ion Transport |
| SCN9A | 0.45 | 4.69E-04 | Ion Transport |
| SLC41A2 | 0.44 | 4.23E-05 | Ion Transport |
| P2RX4 | 0.40 | 1.06E-07 | Ion Transport |
| SLC39A8 | 0.39 | 2.29E-05 | Ion Transport |
| KCNK13 | 0.39 | 6.42E-03 | Ion Transport |
| CLCN2 | 0.38 | 9.45E-04 | Ion Transport |
| KCNJ1 | 0.38 | 1.98E-02 | Ion Transport |
| KCTD10 | 0.37 | 2.16E-09 | Ion Transport |
| TRPC4 | 0.37 | 1.74E-03 | Ion Transport |
| KCNH4 | 0.36 | 2.25E-02 | Ion Transport |
| CLIC4 | 0.36 | 9.25E-03 | Ion Transport |
| CATSPER1 | 0.32 | 1.57E-04 | Ion Transport |
| CHRNA5 | 0.32 | 3.74E-03 | Ion Transport |
| SLC31A2 | 0.31 | 5.43E-08 | Ion Transport |
| SLC39A11 | 0.30 | 5.60E-05 | Ion Transport |
| SLC39A4 | 0.29 | 1.55E-02 | Ion Transport |
| SLC12A4 | 0.29 | 2.80E-08 | Ion Transport |
| CHRNA7 | 0.28 | 1.40E-03 | Ion Transport |
| TRPV2 | 0.28 | 7.70E-04 | Ion Transport |
| TRPM7 | 0.26 | 1.01E-05 | Ion Transport |
| TRPM4 | 0.24 | 1.20E-06 | Ion Transport |
| ATP1B3 | 0.24 | 1.18E-03 | Ion Transport |
| KCNQ1 | 0.24 | 1.35E-06 | Ion Transport |
| ATP11C | 0.24 | 1.96E-08 | Ion Transport |
| CHRNA3 | 0.23 | 4.27E-03 | Ion Transport |
| KCNK7 | 0.22 | 2.00E-03 | Ion Transport |
| SLC39A1 | 0.22 | 3.59E-04 | Ion Transport |
| ARSD | 0.20 | 4.11E-10 | Ion Transport |
| MAGT1 | 0.20 | 3.73E-06 | Ion Transport |
| TRPM1 | 0.19 | 4.44E-03 | Ion Transport |
| TRPM6 | 0.19 | 2.04E-03 | Ion Transport |
| SLC24A6 | 0.19 | 1.00E-05 | Ion Transport |
| ORAI1 | 0.19 | 4.32E-06 | Ion Transport |
| CFTR | 0.19 | 3.77E-03 | Ion Transport |
| SCN11A | 0.18 | 1.59E-02 | Ion Transport |
| SLC3A2 | 0.18 | 1.47E-08 | Ion Transport |
| KCNQ4 | 0.17 | 9.23E-05 | Ion Transport |
| KCNK15 | 0.17 | 8.26E-03 | Ion Transport |
| SLC9A9 | 0.17 | 5.30E-06 | Ion Transport |
| P2RX5 | 0.16 | 2.38E-04 | Ion Transport |
| KCNJ15 | 0.14 | 6.09E-05 | Ion Transport |
| KCNJ8 | 0.14 | 3.91E-05 | Ion Transport |
| P2RX7 | 0.14 | 8.83E-07 | Ion Transport |
| KCNQ5 | 0.14 | 2.51E-05 | Ion Transport |
| SLC30A3 | 0.13 | 1.20E-02 | Ion Transport |
| SLC30A10 | 0.12 | 3.57E-03 | Ion Transport |
| P2RX1 | 0.12 | 3.77E-03 | Ion Transport |
| CLIC2 | 0.11 | 1.48E-06 | Ion Transport |
| ITPR1 | 0.11 | 4.87E-08 | Ion Transport |
| KCNK5 | 0.11 | 6.79E-07 | Ion Transport |
| CLIC1 | 0.10 | 4.77E-07 | Ion Transport |
| GABRE | 0.10 | 3.19E-02 | Ion Transport |
| GABRP | 0.10 | 4.70E-04 | Ion Transport |
| KCNE1 | 0.09 | 7.04E-04 | Ion Transport |
| KCTD12 | 0.08 | 4.52E-06 | Ion Transport |
| CATSPERB | 0.07 | 1.48E-03 | Ion Transport |
| SLC39A12 | 0.06 | 8.76E-05 | Ion Transport |
| CLCA2 | 0.06 | 1.51E-03 | Ion Transport |
| KCNE4 | 0.06 | 9.76E-03 | Ion Transport |
| ATP2A3 | 0.06 | 1.56E-10 | Ion Transport |
| MCOLN3 | 0.05 | 1.22E-09 | Ion Transport |
| KCNJ13 | 0.05 | 1.75E-07 | Ion Transport |
| KCNS3 | 0.05 | 1.79E-05 | Ion Transport |
| SLC12A1 | 0.04 | 1.06E-04 | Ion Transport |
| SLC12A8 | 0.04 | 3.32E-04 | Ion Transport |
| KCNK6 | 0.03 | 1.35E-07 | Ion Transport |
| KCNT2 | 0.03 | 2.06E-04 | Ion Transport |
| TRPA1 | 0.03 | 4.97E-04 | Ion Transport |
| MCOLN2 | 0.03 | 3.09E-03 | Ion Transport |
| TRPM3 | 0.03 | 2.20E-04 | Ion Transport |
| SCN4A | 0.02 | 5.36E-04 | Ion Transport |
| SLC30A2 | 0.02 | 3.53E-04 | Ion Transport |
| SLC24A5 | 0.02 | 2.89E-08 | Ion Transport |
| OCA2 | 0.02 | 1.84E-08 | Ion Transport |
| TCN1 | 0.02 | 1.55E-07 | Ion Transport |
| HTR3A | 0.02 | 3.09E-02 | Ion Transport |
| SCNN1A | 0.01 | 3.44E-03 | Ion Transport |
| TRPV4 | 0.01 | 5.70E-05 | Ion Transport |
| BEST1 | 0.01 | 1.09E-04 | Ion Transport |
| KCNMB1 | 0.01 | 1.39E-06 | Ion Transport |
| KCTD4 | 0.01 | 4.77E-05 | Ion Transport |
| CLIC6 | 0.01 | 3.73E-06 | Ion Transport |
| CLIC3 | 4.27E-03 | 3.67E-06 | Ion Transport |
| KCNK17 | 1.44E-03 | 4.32E-05 | Ion Transport |
| BDH1 | 2.39 | 7.29E-09 | Ketone Bodies |
| HMGCLL1 | 2.29 | 3.20E-05 | Ketone Bodies |
| AACS | 1.97 | 3.78E-08 | Ketone Bodies |
| OXCT1 | 1.67 | 1.59E-04 | Ketone Bodies |
| HMGCL | 0.60 | 2.04E-04 | Ketone Bodies |
| BDH2 | 0.15 | 8.58E-07 | Ketone Bodies |
| HMGCS2 | 0.01 | 4.98E-03 | Ketone Bodies |
| PLA2G4C | 140.04 | 8.08E-09 | Lipid (Membrane Lipid) |
| AANAT | 119.72 | 4.92E-10 | Lipid (Hormone) |
| PLA2G4D | 105.35 | 1.90E-04 | Lipid (Membrane Lipid) |
| MTTP | 74.10 | 3.64E-05 | Lipid (Fatty Acid) |
| PLCH2 | 61.46 | 3.29E-10 | Lipid (Membrane Lipid) |
| HSD17B2 | 59.43 | 2.02E-04 | Lipid (Steroid) |
| ELOVL4 | 51.34 | 4.01E-11 | Lipid (Fatty Acid) |
| PLA2G12B | 46.37 | 3.39E-04 | Lipid (Membrane Lipid) |
| PLA2G4E | 35.06 | 1.10E-03 | Lipid (Membrane Lipid) |
| ALOX12B | 34.20 | 1.64E-09 | Lipid (Hormone) |
| ACBD7 | 32.65 | 5.46E-06 | Lipid (Fatty Acid) |
| LPPR4 | 23.84 | 2.02E-11 | Lipid (Membrane Lipid) |
| CYP27B1 | 20.39 | 1.82E-05 | Lipid (Cofactor) |
| CYP2W1 | 19.24 | 6.85E-03 | Lipid (Steroid) |
| AKR1C4 | 19.16 | 2.58E-03 | Lipid (Steroid) |
| SGPP2 | 16.42 | 8.49E-08 | Lipid (Sphingolipid) |
| ABHD12B | 15.31 | 2.34E-10 | Lipid (Fatty Acid) |
| BAAT | 13.44 | 3.04E-06 | Lipid (Bile Acid) |
| DGKI | 13.02 | 5.91E-09 | Lipid (Membrane Lipid) |
| CDS1 | 12.80 | 5.29E-11 | Lipid (Membrane Lipid) |
| DGKB | 12.45 | 3.39E-07 | Lipid (Membrane Lipid) |
| CERKL | 12.34 | 1.18E-12 | Lipid (Sphingolipid) |
| GDPD2 | 12.06 | 1.17E-07 | Lipid (Membrane Lipid) |
| TPO | 11.58 | 2.32E-05 | Lipid (Hormone) |
| PLCD4 | 11.55 | 9.16E-07 | Lipid (Membrane Lipid) |
| ALOXE3 | 10.72 | 4.36E-04 | Lipid (Hormone) |
| ASAH2 | 10.58 | 2.23E-08 | Lipid (Sphingosine) |
| CYP2A7 | 10.57 | 8.77E-03 | Lipid (Steroid) |
| CYP2E1 | 10.35 | 6.93E-07 | Lipid (Steroid) |
| ACSL6 | 9.56 | 3.99E-10 | Lipid (Fatty Acid) |
| PNLIPRP2 | 9.36 | 1.01E-03 | Lipid (Fatty Acid) |
| SMPD3 | 9.26 | 5.78E-08 | Lipid (Sphingolipid) |
| FAAH2 | 9.17 | 2.43E-09 | Lipid (Fatty Acid) |
| SRD5A2 | 9.06 | 1.38E-03 | Lipid (Steroid) |
| CYP2C18 | 8.37 | 1.43E-02 | Lipid (Steroid) |
| SCD5 | 7.98 | 2.35E-09 | Lipid (Fatty Acid) |
| LPCAT1 | 7.87 | 2.29E-09 | Lipid (Membrane Lipid) |
| CYP1A1 | 7.84 | 3.21E-05 | Lipid (Steroid) |
| CYP24A1 | 7.76 | 3.59E-03 | Lipid (Cofactor) |
| ACSM4 | 6.80 | 1.77E-05 | Lipid (Fatty Acid) |
| CYP2D6 | 6.75 | 6.63E-06 | Lipid (Steroid) |
| PPAP2C | 6.22 | 1.10E-03 | Lipid (Membrane Lipid) |
| DGKD | 6.17 | 5.46E-10 | Lipid (Membrane Lipid) |
| GDPD4 | 5.90 | 1.91E-04 | Lipid (Membrane Lipid) |
| GPD1 | 5.78 | 3.92E-07 | Lipid (Membrane Lipid) |
| CYP2J2 | 5.66 | 2.16E-05 | Lipid (Steroid) |
| ACSBG2 | 5.35 | 6.02E-06 | Lipid (Fatty Acid) |
| DGKE | 5.20 | 5.11E-09 | Lipid (Membrane Lipid) |
| PPAPDC3 | 4.90 | 1.91E-06 | Lipid (Membrane Lipid) |
| ASAH2B | 4.71 | 9.83E-10 | Lipid (Sphingosine) |
| GDPD5 | 4.63 | 1.18E-08 | Lipid (Membrane Lipid) |
| AMACR | 4.14 | 1.16E-08 | Lipid (Fatty acid) |
| SRD5A1 | 4.03 | 1.82E-05 | Lipid (Steroid) |
| GDPD1 | 3.77 | 1.58E-09 | Lipid (Membrane Lipid) |
| CROT | 3.59 | 1.27E-07 | Lipid (Membrane Lipid) |
| PHOSPHO1 | 3.49 | 7.02E-03 | Lipid (Membrane Lipid) |
| PLCD3 | 3.48 | 4.60E-08 | Lipid (Membrane Lipid) |
| ELOVL6 | 3.23 | 3.45E-07 | Lipid (Fatty Acid) |
| HSD11B1L | 3.19 | 7.34E-06 | Lipid (Steroid) |
| CDS2 | 3.16 | 2.11E-07 | Lipid (Membrane Lipid) |
| LIPI | 3.13 | 7.22E-05 | Lipid (Fatty Acid) |
| HSDL1 | 3.13 | 4.11E-08 | Lipid (Steroid) |
| PLCB1 | 3.10 | 1.90E-05 | Lipid (Membrane Lipid) |
| AGPAT3 | 3.09 | 1.16E-08 | Lipid (Membrane Lipid) |
| KLB | 3.08 | 2.65E-03 | Lipid (Steroid) |
| PLCH1 | 3.02 | 2.34E-06 | Lipid (Membrane Lipid) |
| PCYT1B | 2.83 | 1.01E-05 | Lipid (Membrane Lipid) |
| ABHD8 | 2.72 | 1.85E-05 | Lipid (Fatty Acid) |
| UGT8 | 2.65 | 2.64E-04 | Lipid (Sphingolipid) |
| DDHD1 | 2.53 | 1.19E-11 | Lipid (Membrane Lipid) |
| ACOXL | 2.50 | 2.45E-03 | Lipid (Fatty Acid) |
| CEL | 2.46 | 1.33E-03 | Lipid (Cofactor) |
| LPCAT4 | 2.44 | 4.42E-05 | Lipid (Membrane Lipid) |
| PLD4 | 2.42 | 2.42E-02 | Lipid (Membrane Lipid) |
| CPT1C | 2.40 | 1.52E-08 | Lipid (Fatty Acid) |
| MECR | 2.38 | 3.03E-07 | Lipid (Fatty Acid) |
| PTER | 2.31 | 1.32E-06 | Lipid |
| LYPLA1 | 2.28 | 1.76E-09 | Lipid (Membrane Lipid) |
| CYP17A1 | 2.20 | 1.11E-03 | Lipid (Steroid) |
| GK | 2.11 | 6.99E-07 | Lipid (Membrane Lipid) |
| ACBD4 | 1.99 | 9.78E-09 | Lipid (Fatty Acid) |
| DAGLA | 1.99 | 4.76E-03 | Lipid (Membrane Lipid) |
| LNPEP | 1.99 | 2.14E-04 | Lipid (Hormone) |
| THEM4 | 1.97 | 8.52E-08 | Lipid (Fatty Acid) |
| CYP2A6 | 1.74 | 5.48E-03 | Lipid (Steroid) |
| ACER2 | 1.73 | 1.44E-03 | Lipid (Sphingosine) |
| ACSL3 | 1.68 | 3.96E-06 | Lipid (Fatty Acid) |
| CERK | 1.68 | 1.96E-06 | Lipid (Sphingolipid) |
| DGKG | 1.66 | 4.05E-02 | Lipid (Membrane Lipid) |
| ABHD14A | 1.66 | 3.37E-04 | Lipid (Fatty Acid) |
| SDR39U1 | 1.66 | 1.26E-06 | Lipid (Fatty Acid) |
| ELOVL2 | 1.58 | 2.06E-03 | Lipid (Fatty Acid) |
| MCAT | 1.57 | 4.83E-05 | Lipid (Fatty Acid) |
| DAK | 1.56 | 1.48E-04 | Lipid (Cofactor) |
| MMEL1 | 1.55 | 8.68E-05 | Lipid (Hormone) |
| GGPS1 | 1.54 | 7.65E-06 | Lipid (Steroid) |
| AGPAT5 | 1.54 | 1.65E-05 | Lipid (Membrane Lipid) |
| ETNK1 | 1.53 | 5.24E-07 | Lipid (Membrane Lipid) |
| SPTLC2 | 1.52 | 4.63E-03 | Lipid (Sphingolipid) |
| DGAT1 | 1.51 | 1.43E-04 | Lipid (Fatty Acid) |
| PLD6 | 1.51 | 1.39E-04 | Lipid (Membrane Lipid) |
| PNPLA8 | 1.49 | 1.91E-03 | Lipid (Fatty Acid) |
| LPGAT1 | 1.46 | 2.62E-04 | Lipid (Membrane Lipid) |
| LIAS | 1.44 | 3.59E-03 | Lipid |
| MBOAT2 | 1.44 | 4.09E-03 | Lipid (Membrane Lipid) |
| AGPAT4 | 1.41 | 1.18E-02 | Lipid (Membrane Lipid) |
| SGPL1 | 1.40 | 2.35E-04 | Lipid (Sphingolipid) |
| IDI2 | 1.38 | 6.60E-03 | Lipid (Mevalonate) |
| PLCB4 | 1.38 | 1.41E-02 | Lipid (Membrane Lipid) |
| SC5DL | 1.35 | 5.52E-03 | Lipid (Cholesterol) |
| TM7SF2 | 1.33 | 8.12E-05 | Lipid (Cholesterol) |
| ALDH3A2 | 1.32 | 2.34E-03 | Lipid (Fatty Acid) |
| SCCPDH | 1.32 | 3.17E-05 | Lipid (Membrane lipd) |
| ECHS1 | 0.77 | 3.26E-03 | Lipid (Fatty Acid) |
| PCYT1A | 0.77 | 4.87E-03 | Lipid (Membrane Lipid) |
| LPIN1 | 0.77 | 9.77E-03 | Lipid (Membrane Lipid) |
| CPT1A | 0.76 | 1.73E-04 | Lipid (Fatty Acid) |
| ALOX12 | 0.76 | 1.48E-02 | Lipid (Hormone) |
| IAH1 | 0.74 | 4.22E-06 | Lipid |
| ETFDH | 0.74 | 5.52E-03 | Lipid (Fatty Acid) |
| SRD5A3 | 0.73 | 3.31E-02 | Lipid (Steroid) |
| GNPAT | 0.73 | 4.96E-04 | Lipid (Membrane Lipid) |
| ILVBL | 0.72 | 3.82E-05 | Lipid (Fatty Acid) |
| ABHD14B | 0.69 | 1.41E-05 | Lipid (Fatty Acid) |
| ACAA2 | 0.69 | 3.92E-03 | Lipid (Fatty Acid) |
| ABHD1 | 0.68 | 2.98E-02 | Lipid (Fatty Acid) |
| PCYT2 | 0.68 | 5.27E-04 | Lipid (Membrane Lipid) |
| PTGR1 | 0.67 | 2.60E-04 | Lipid (Hormone) |
| PTDSS1 | 0.67 | 3.09E-03 | Lipid (Membrane Lipid) |
| ACACA | 0.67 | 9.63E-03 | Lipid (Fatty Acid) |
| GALC | 0.67 | 1.16E-03 | Lipid (Sphingolipid) |
| PEMT | 0.66 | 3.96E-03 | Lipid (Membrane Lipid) |
| ELOVL1 | 0.65 | 1.21E-07 | Lipid (Fatty Acid) |
| PMVK | 0.65 | 1.38E-02 | Lipid (Mevalonate) |
| KDSR | 0.65 | 6.47E-04 | Lipid (Sphingosine) |
| FDPS | 0.65 | 1.46E-04 | Lipid (Cholesterol) |
| ABHD5 | 0.64 | 2.81E-02 | Lipid (Fatty Acid) |
| HSD17B12 | 0.64 | 8.30E-05 | Lipid (Steroid) |
| HSD17B7 | 0.64 | 3.88E-03 | Lipid (Cholesterol) |
| ETFB | 0.64 | 1.23E-03 | Lipid (Fatty Acid) |
| DGKZ | 0.63 | 2.37E-03 | Lipid (Membrane Lipid) |
| EBPL | 0.62 | 1.31E-06 | Lipid (Cholesterol) |
| ACBD6 | 0.62 | 8.32E-05 | Lipid (Fatty Acid) |
| PLD2 | 0.61 | 2.28E-04 | Lipid (Membrane Lipid) |
| SOAT1 | 0.60 | 2.26E-10 | Lipid (Steroid) |
| CYP11A1 | 0.60 | 3.61E-02 | Lipid (Steroid) |
| DHCR24 | 0.60 | 5.00E-03 | Lipid (Cholesterol) |
| CYP2U1 | 0.59 | 2.66E-06 | Lipid (Steroid) |
| MCEE | 0.58 | 1.36E-04 | Lipid (Propanoate) |
| CEPT1 | 0.58 | 6.00E-09 | Lipid (Membrane Lipid) |
| SCP2 | 0.58 | 1.08E-08 | Lipid (Fatty Acid) |
| FASN | 0.57 | 3.90E-03 | Lipid (Fatty Acid) |
| HADHB | 0.57 | 2.53E-06 | Lipid (Fatty Acid) |
| CYP20A1 | 0.57 | 6.56E-04 | Lipid (Hormone) |
| ACER3 | 0.56 | 6.63E-05 | Lipid (Sphingosine) |
| CYP51A1 | 0.55 | 5.24E-03 | Lipid (Cholesterol) |
| PLCD1 | 0.55 | 1.66E-04 | Lipid (Membrane Lipid) |
| SPTLC1 | 0.54 | 9.78E-13 | Lipid (Sphingolipid) |
| GPD1L | 0.54 | 1.11E-03 | Lipid (Membrane Lipid) |
| CRAT | 0.54 | 1.39E-09 | Lipid (Cofactor) |
| ABHD4 | 0.53 | 4.30E-07 | Lipid (Fatty Acid) |
| ETFA | 0.53 | 5.35E-08 | Lipid (Fatty Acid) |
| ETNK2 | 0.53 | 1.92E-05 | Lipid (Membrane Lipid) |
| ACAT2 | 0.52 | 1.65E-02 | Lipid (Fatty Acid) |
| PCCB | 0.52 | 4.05E-03 | Lipid (Propanoate) |
| CYP4V2 | 0.51 | 7.52E-03 | Lipid (Fatty Acid) |
| LBR | 0.51 | 1.66E-03 | Lipid (Cholesterol) |
| PLCG1 | 0.51 | 1.24E-07 | Lipid (Membrane Lipid) |
| HADH | 0.51 | 2.09E-06 | Lipid (Fatty Acid) |
| SQLE | 0.50 | 3.85E-04 | Lipid (Cholesterol) |
| CPT2 | 0.49 | 7.63E-04 | Lipid (Fatty Acid) |
| IDI1 | 0.49 | 2.34E-03 | Lipid (Mevalonate) |
| HSD17B4 | 0.49 | 7.72E-09 | Lipid (Steroid) |
| SGMS1 | 0.48 | 3.32E-05 | Lipid (Sphingolipid) |
| ECHDC1 | 0.48 | 2.50E-06 | Lipid (Fatty Acid) |
| ASAH1 | 0.47 | 2.06E-06 | Lipid (Sphingosine) |
| DIO2 | 0.47 | 2.42E-06 | Lipid (Hormone) |
| ACAA1 | 0.47 | 1.41E-07 | Lipid (Fatty Acid) |
| LYPLAL1 | 0.45 | 4.13E-03 | Lipid (Membrane Lipid) |
| DEGS2 | 0.44 | 2.65E-04 | Lipid (Sphingolipid) |
| CYP4F12 | 0.44 | 4.39E-02 | Lipid (Fatty Acid) |
| PNPLA2 | 0.43 | 4.60E-04 | Lipid (Fatty Acid) |
| CYP4F2 | 0.43 | 4.15E-02 | Lipid (Fatty Acid) |
| ACADL | 0.42 | 1.37E-07 | Lipid (Fatty Acid) |
| PLCB3 | 0.40 | 2.28E-04 | Lipid (Membrane Lipid) |
| DEGS1 | 0.40 | 3.05E-08 | Lipid (Sphingolipid) |
| LTC4S | 0.39 | 7.51E-03 | Lipid (Hormone) |
| ACAT1 | 0.39 | 2.27E-04 | Lipid (Fatty Acid) |
| ACADVL | 0.38 | 5.43E-06 | Lipid (Fatty Acid) |
| PLCE1 | 0.37 | 5.19E-05 | Lipid (Membrane Lipid) |
| SMPD1 | 0.36 | 7.58E-10 | Lipid (Sphingolipid) |
| FADS1 | 0.36 | 4.25E-05 | Lipid (Fatty Acid) |
| DECR2 | 0.36 | 1.39E-08 | Lipid (Fatty Acid) |
| HSD17B6 | 0.36 | 6.77E-03 | Lipid (Steroid) |
| CYP7B1 | 0.36 | 2.00E-08 | Lipid (Bile Acid) |
| CYP2S1 | 0.35 | 8.69E-03 | Lipid (Steroid) |
| HSD17B14 | 0.34 | 3.62E-07 | Lipid (Steroid) |
| MBOAT1 | 0.34 | 8.13E-05 | Lipid (Membrane Lipid) |
| ELOVL3 | 0.34 | 2.00E-02 | Lipid (Fatty Acid) |
| ACSS3 | 0.34 | 1.84E-04 | Lipid (Fatty Acid) |
| ARSJ | 0.34 | 1.31E-02 | Lipid (Membrane Lipid) |
| CLYBL | 0.34 | 1.74E-03 | Lipid (Fatty Acid) |
| PLCXD3 | 0.33 | 5.38E-05 | Lipid (Membrane Lipid) |
| DECR1 | 0.33 | 2.30E-09 | Lipid (Fatty Acid) |
| HADHA | 0.32 | 1.48E-08 | Lipid (Fatty Acid) |
| HSDL2 | 0.32 | 4.05E-05 | Lipid (Steroid) |
| CYP39A1 | 0.32 | 5.95E-08 | Lipid (Steroid) |
| ECH1 | 0.32 | 8.11E-08 | Lipid (Fatty Acid) |
| LIPE | 0.32 | 1.86E-05 | Lipid (Steroid) |
| PNLIPRP3 | 0.30 | 3.07E-02 | Lipid (Fatty Acid) |
| HSD17B10 | 0.30 | 3.95E-06 | Lipid (Steroid) |
| FDFT1 | 0.29 | 9.90E-05 | Lipid (Cholesterol) |
| LPCAT3 | 0.29 | 9.03E-06 | Lipid (Membrane Lipid) |
| LCAT | 0.29 | 2.80E-08 | Lipid (Cholesterol) |
| HSD17B11 | 0.28 | 1.49E-05 | Lipid (Steroid) |
| ACSS1 | 0.27 | 1.03E-06 | Lipid (Fatty Acid) |
| LIPA | 0.27 | 3.52E-07 | Lipid (Steroid) |
| ABHD2 | 0.27 | 1.38E-05 | Lipid (Fatty Acid) |
| PLA2G16 | 0.26 | 9.53E-08 | Lipid (Membrane Lipid) |
| AOAH | 0.26 | 5.42E-04 | Lipid (Fatty Acid) |
| ACSM5 | 0.26 | 1.24E-05 | Lipid (Fatty Acid) |
| LIPN | 0.26 | 2.74E-02 | Lipid (Fatty Acid) |
| DGAT2 | 0.26 | 1.54E-06 | Lipid (Fatty Acid) |
| PLCL1 | 0.26 | 7.71E-04 | Lipid (Membrane Lipid) |
| ACACB | 0.25 | 3.15E-07 | Lipid (Fatty Acid) |
| CYP27A1 | 0.25 | 8.52E-09 | Lipid (Bile Acid) |
| HSD17B3 | 0.25 | 1.62E-09 | Lipid (Steroid) |
| SMPDL3A | 0.24 | 8.85E-06 | Lipid (Sphingolipid) |
| ACSBG1 | 0.24 | 1.25E-02 | Lipid (Fatty Acid) |
| CHPT1 | 0.23 | 8.17E-04 | Lipid (Membrane Lipid) |
| PTGS1 | 0.23 | 1.95E-03 | Lipid (Hormone) |
| ACE2 | 0.23 | 6.49E-04 | Lipid (Hormone) |
| ELOVL5 | 0.23 | 3.06E-08 | Lipid (Fatty Acid) |
| LPIN3 | 0.23 | 9.13E-07 | Lipid (Membrane Lipid) |
| DGKA | 0.23 | 3.68E-08 | Lipid (Membrane Lipid) |
| ACSF2 | 0.22 | 2.46E-08 | Lipid (Fatty Acid) |
| FADS2 | 0.22 | 1.36E-04 | Lipid (Fatty Acid) |
| ACSL1 | 0.22 | 6.57E-08 | Lipid (Fatty Acid) |
| ACER1 | 0.21 | 4.90E-04 | Lipid (Sphingosine) |
| ACOX2 | 0.21 | 3.77E-05 | Lipid (Fatty Acid) |
| ACADS | 0.18 | 5.82E-08 | Lipid (Fatty Acid) |
| DHCR7 | 0.18 | 5.65E-05 | Lipid (Cholesterol) |
| PNPLA3 | 0.17 | 2.33E-04 | Lipid (Fatty Acid) |
| PLA2G5 | 0.16 | 3.10E-04 | Lipid (Membrane Lipid) |
| PPAPDC1B | 0.15 | 8.59E-09 | Lipid (Membrane Lipid) |
| TBXAS1 | 0.14 | 6.40E-05 | Lipid (Hormone) |
| CYP1A2 | 0.14 | 4.39E-02 | Lipid (Steroid) |
| ABHD3 | 0.14 | 8.26E-06 | Lipid (Fatty Acid) |
| CYP4X1 | 0.14 | 2.68E-02 | Lipid (Fatty Acid) |
| AGPAT2 | 0.14 | 1.38E-05 | Lipid (Membrane Lipid) |
| PPAP2B | 0.13 | 2.33E-06 | Lipid (Membrane Lipid) |
| GPAM | 0.13 | 7.54E-05 | Lipid (Membrane Lipid) |
| PLD1 | 0.13 | 5.43E-10 | Lipid (Membrane Lipid) |
| ECHDC3 | 0.12 | 1.03E-09 | Lipid (Fatty Acid) |
| SCD | 0.12 | 2.46E-05 | Lipid (Fatty Acid) |
| PLCB2 | 0.12 | 1.60E-04 | Lipid (Membrane Lipid) |
| LIPG | 0.12 | 4.95E-05 | Lipid (Fatty Acid) |
| LPL | 0.11 | 5.18E-03 | Lipid (Fatty Acid) |
| PLCG2 | 0.11 | 1.31E-06 | Lipid (Membrane Lipid) |
| CYP4F22 | 0.11 | 1.05E-02 | Lipid (Fatty Acid) |
| PPAP2A | 0.11 | 5.84E-15 | Lipid (Membrane Lipid) |
| ALDH3B1 | 0.11 | 1.87E-06 | Lipid (Fatty Acid) |
| CYP4Z1 | 0.09 | 1.35E-03 | Lipid (Fatty Acid) |
| LPCAT2 | 0.08 | 9.94E-06 | Lipid (Membrane Lipid) |
| PLA2G4A | 0.08 | 1.02E-04 | Lipid (Membrane Lipid) |
| PTGDS | 0.08 | 4.25E-04 | Lipid (Hormone) |
| CYP7A1 | 0.08 | 4.55E-03 | Lipid (Bile Acid) |
| PLB1 | 0.08 | 5.78E-03 | Lipid (Membrane Lipid) |
| MME | 0.08 | 9.71E-05 | Lipid (Hormone) |
| SGMS2 | 0.07 | 7.91E-09 | Lipid (Sphingolipid) |
| LIPC | 0.07 | 6.91E-04 | Lipid (Fatty Acid) |
| ARSE | 0.07 | 2.29E-05 | Lipid (Membrane Lipid) |
| EHHADH | 0.07 | 7.19E-05 | Lipid (Fatty Acid) |
| ACSM3 | 0.07 | 1.05E-03 | Lipid (Fatty Acid) |
| PRG2 | 0.06 | 4.51E-03 | Lipid (Membrane Lipid) |
| SOAT2 | 0.06 | 1.14E-03 | Lipid (Steroid) |
| ALOX5AP | 0.06 | 4.53E-04 | Lipid (Hormone) |
| ALOX5 | 0.05 | 7.50E-07 | Lipid (Hormone) |
| HPGD | 0.05 | 1.09E-06 | Lipid (Hormone) |
| CYP4F3 | 0.04 | 4.66E-03 | Lipid (Fatty Acid) |
| PTGES | 0.04 | 2.19E-02 | Lipid (Hormone) |
| PLA2G7 | 0.03 | 2.84E-04 | Lipid (Membrane Lipid) |
| KL | 0.03 | 2.28E-09 | Lipid (Steroid) |
| PNPLA5 | 0.03 | 1.31E-03 | Lipid (Fatty Acid) |
| LIPH | 0.03 | 1.42E-04 | Lipid (Fatty Acid) |
| SDR42E1 | 0.03 | 4.35E-04 | Lipid (Fatty Acid) |
| OLAH | 0.02 | 1.98E-02 | Lipid (Fatty Acid) |
| PLA1A | 0.02 | 1.02E-09 | Lipid (Membrane Lipid) |
| ACSL5 | 0.02 | 4.88E-09 | Lipid (Fatty Acid) |
| PTGIS | 0.02 | 8.84E-03 | Lipid (Hormone) |
| CYP4B1 | 0.01 | 4.59E-05 | Lipid (Fatty Acid) |
| PLD5 | 0.01 | 1.18E-04 | Lipid (Membrane Lipid) |
| CYP1B1 | 0.01 | 5.31E-06 | Lipid (Steroid) |
| CYP27C1 | 0.01 | 5.58E-07 | Lipid (Steroid) |
| HSD11B1 | 0.01 | 3.05E-02 | Lipid (Steroid) |
| PLA2G2A | 7.03E-04 | 4.37E-03 | Lipid (Membrane Lipid) |
| NOX4 | 20.50 | 6.38E-10 | NAD |
| NMNAT2 | 3.85 | 3.42E-05 | NAD |
| BST1 | 0.74 | 9.92E-05 | NAD |
| NADK | 0.61 | 4.57E-06 | NAD |
| NMNAT1 | 0.57 | 2.40E-05 | NAD |
| QPRT | 0.50 | 2.68E-03 | NAD |
| NADSYN1 | 0.45 | 1.63E-08 | NAD |
| NQO2 | 0.42 | 1.76E-05 | NAD |
| NQO1 | 0.34 | 1.32E-04 | NAD |
| DUOX2 | 0.18 | 2.01E-03 | NAD |
| DUOX1 | 0.17 | 4.87E-06 | NAD |
| NAMPT | 0.13 | 1.16E-02 | NAD |
| NNMT | 0.01 | 2.79E-03 | NAD |
| CD38 | 0.01 | 1.06E-04 | NAD |
| DDC | 32.63 | 4.14E-10 | Neurotransmitter |
| CHAT | 8.39 | 8.45E-06 | Neurotransmitter |
| ACHE | 7.69 | 4.12E-07 | Neurotransmitter |
| ABAT | 3.89 | 2.86E-07 | Neurotransmitter |
| COMTD1 | 2.04 | 6.29E-04 | Neurotransmitter |
| COMT | 0.49 | 6.16E-06 | Neurotransmitter |
| NAALADL1 | 0.42 | 6.55E-07 | Neurotransmitter |
| DBH | 0.21 | 4.67E-02 | Neurotransmitter |
| CA10 | 128.28 | 3.28E-17 | Nitrogen Metabolism |
| CA5A | 28.26 | 1.65E-03 | Nitrogen Metabolism |
| CA2 | 11.59 | 7.84E-10 | Nitrogen Metabolism |
| CA9 | 5.80 | 3.81E-02 | Nitrogen Metabolism |
| CA11 | 4.74 | 1.56E-07 | Nitrogen Metabolism |
| CA13 | 3.81 | 2.01E-04 | Nitrogen Metabolism |
| CA7 | 3.09 | 5.36E-03 | Nitrogen Metabolism |
| CA14 | 0.25 | 3.28E-03 | Nitrogen Metabolism |
| CA12 | 0.05 | 4.73E-02 | Nitrogen Metabolism |
| CA4 | 1.48E-03 | 7.20E-07 | Nitrogen Metabolism |
| GUCY2F | 152.38 | 4.74E-08 | Nucleotide (cGMP) |
| PDE6A | 142.32 | 6.82E-09 | Nucleotide (cGMP) |
| PDE6H | 135.30 | 2.75E-15 | Nucleotide (cGMP) |
| DPYSL5 | 97.59 | 8.91E-15 | Nucleotide (Pyrimidine) |
| ADCY8 | 78.33 | 4.16E-09 | Nucleotide (cAMP) |
| ADCY1 | 50.91 | 2.39E-12 | Nucleotide (cAMP) |
| PDE6C | 45.95 | 2.25E-11 | Nucleotide (cGMP) |
| GUCY2D | 42.08 | 1.20E-07 | Nucleotide (cGMP) |
| PDE11A | 28.39 | 6.57E-08 | Nucleotide (cyclic nucleotide) |
| PDE6B | 21.67 | 5.16E-07 | Nucleotide (cGMP) |
| NT5E | 17.90 | 3.76E-08 | Nucleotide |
| IMPDH1 | 15.51 | 3.74E-09 | Nucleotide (Purine) |
| DPYSL4 | 14.04 | 1.81E-11 | Nucleotide (Pyrimidine) |
| GUCY1A2 | 8.69 | 4.28E-07 | Nucleotide (cGMP) |
| A1CF | 7.85 | 1.16E-05 | Nucleotide |
| NT5C1A | 6.48 | 3.35E-06 | Nucleotide |
| AK7 | 5.57 | 4.41E-08 | Nucleotide (Purine) |
| HPRT1 | 4.71 | 3.57E-08 | Nucleotide (Purine) |
| PDE4D | 4.44 | 2.51E-05 | Nucleotide (cGMP) |
| UPP2 | 4.33 | 7.87E-04 | Nucleotide (Pyrimidine) |
| NUDT12 | 4.12 | 3.35E-08 | Nucleotide |
| GUK1 | 3.71 | 2.19E-10 | Nucleotide (Purine) |
| TYMS | 3.68 | 4.02E-06 | Nucleotide (Pyrimidine) |
| NT5C2 | 3.30 | 5.10E-13 | Nucleotide |
| DPYSL3 | 3.08 | 8.68E-08 | Nucleotide (Pyrimidine) |
| PDE6D | 2.87 | 4.61E-08 | Nucleotide (cGMP) |
| GUCY1B3 | 2.86 | 6.47E-11 | Nucleotide (cGMP) |
| XDH | 2.65 | 1.50E-02 | Nucleotide (Purine) |
| NUDT11 | 2.62 | 1.16E-04 | Nucleotide |
| ADCY2 | 2.59 | 1.57E-03 | Nucleotide (cAMP) |
| PDE1A | 2.47 | 1.27E-03 | Nucleotide (cyclic nucleotide) |
| ENTPD2 | 2.47 | 3.01E-03 | Nucleotide |
| ENTPD4 | 2.42 | 1.73E-07 | Nucleotide |
| UPB1 | 2.42 | 6.48E-04 | Nucleotide (Pyrimidine) |
| NUDT19 | 2.37 | 1.04E-07 | Nucleotide |
| PRPSAP2 | 2.22 | 2.98E-10 | Nucleotide (Purine) |
| DPYSL2 | 2.20 | 2.95E-06 | Nucleotide (Pyrimidine) |
| NME5 | 2.15 | 4.50E-04 | Nucleotide |
| AMPD3 | 2.11 | 5.21E-06 | Nucleotide (Purine) |
| NT5M | 2.05 | 4.18E-03 | Nucleotide |
| ADCY5 | 2.04 | 9.82E-05 | Nucleotide (cAMP) |
| ADAL | 2.04 | 1.15E-05 | Nucleotide (Purine) |
| PDE7A | 2.02 | 2.38E-05 | Nucleotide (cyclic nucleotide) |
| PDE4A | 1.96 | 7.39E-04 | Nucleotide (cGMP) |
| PDE8B | 1.89 | 8.73E-05 | Nucleotide (cyclic nucleotide) |
| DCK | 1.85 | 1.51E-06 | Nucleotide |
| RRM1 | 1.84 | 1.30E-07 | Nucleotide |
| NT5C3 | 1.77 | 6.58E-08 | Nucleotide |
| NUDT4 | 1.75 | 1.37E-05 | Nucleotide |
| AK5 | 1.73 | 1.44E-03 | Nucleotide (Purine) |
| PFAS | 1.72 | 3.62E-04 | Nucleotide (Purine) |
| NUDT10 | 1.69 | 5.55E-03 | Nucleotide |
| NUDT17 | 1.50 | 8.08E-03 | Nucleotide |
| PDE12 | 1.46 | 2.61E-04 | Nucleotide (cyclic nucleotide) |
| RRM2B | 1.45 | 2.20E-05 | Nucleotide |
| PDE8A | 1.40 | 1.38E-03 | Nucleotide (cyclic nucleotide) |
| AMPD2 | 1.37 | 2.17E-05 | Nucleotide (Purine) |
| NUDT16L1 | 1.37 | 9.13E-04 | Nucleotide |
| ENTPD6 | 1.33 | 2.47E-02 | Nucleotide |
| ENTPD1 | 0.77 | 9.01E-03 | Nucleotide |
| CANT1 | 0.76 | 3.84E-02 | Nucleotide |
| NME6 | 0.74 | 7.31E-03 | Nucleotide |
| ADCY6 | 0.72 | 5.34E-05 | Nucleotide (cAMP) |
| DUT | 0.71 | 2.53E-06 | Nucleotide (Pyrimidine) |
| GMPS | 0.71 | 3.55E-04 | Nucleotide (Purine) |
| GMPR2 | 0.68 | 1.09E-04 | Nucleotide (Purine) |
| NT5DC3 | 0.68 | 4.76E-03 | Nucleotide |
| TK2 | 0.67 | 2.05E-05 | Nucleotide (Pyrimidine) |
| ATIC | 0.62 | 6.44E-08 | Nucleotide (Purine) |
| IMPDH2 | 0.62 | 4.54E-04 | Nucleotide (Purine) |
| NUDT5 | 0.60 | 2.60E-05 | Nucleotide |
| ENPP5 | 0.58 | 3.06E-02 | Nucleotide |
| CTPS | 0.57 | 5.17E-03 | Nucleotide (Pyrimidine) |
| NUDT6 | 0.55 | 3.62E-06 | Nucleotide |
| NUDT7 | 0.55 | 1.84E-03 | Nucleotide |
| CECR1 | 0.52 | 9.43E-03 | Nucleotide (Purine) |
| CMPK | 0.50 | 2.74E-08 | Nucleotide (Pyrimidine) |
| NT5DC1 | 0.49 | 1.33E-06 | Nucleotide |
| NT5DC2 | 0.47 | 4.78E-07 | Nucleotide |
| NUDT18 | 0.46 | 2.08E-06 | Nucleotide |
| ENPP4 | 0.46 | 5.25E-05 | Nucleotide |
| NUDT3 | 0.46 | 2.43E-07 | Nucleotide |
| PDE4C | 0.42 | 2.74E-02 | Nucleotide (cGMP) |
| PDE5A | 0.39 | 1.40E-02 | Nucleotide (cGMP) |
| NME4 | 0.36 | 1.39E-08 | Nucleotide |
| DCTD | 0.35 | 1.87E-12 | Nucleotide (Pyrimidine) |
| PDE1C | 0.34 | 8.46E-05 | Nucleotide (cGMP) |
| ADCY9 | 0.34 | 6.20E-09 | Nucleotide (cAMP) |
| ADCY3 | 0.31 | 5.72E-03 | Nucleotide (cAMP) |
| PDE1B | 0.31 | 1.03E-02 | Nucleotide (cyclic nucleotide) |
| APRT | 0.30 | 5.43E-05 | Nucleotide (Purine) |
| PDE2A | 0.26 | 2.92E-05 | Nucleotide (cGMP) |
| OAS3 | 0.26 | 6.07E-04 | Nucleotide (Ribonuclease) |
| ADCY7 | 0.25 | 1.35E-05 | Nucleotide (cAMP) |
| OAS1 | 0.23 | 7.80E-04 | Nucleotide (Ribonuclease) |
| AK3 | 0.20 | 8.01E-10 | Nucleotide (Purine) |
| ADA | 0.18 | 2.43E-06 | Nucleotide (Purine) |
| PDE3B | 0.18 | 4.47E-05 | Nucleotide (cGMP) |
| GUCY2C | 0.16 | 1.81E-04 | Nucleotide (cGMP) |
| DPYD | 0.14 | 2.97E-04 | Nucleotide (Pyrimidine) |
| PRPS2 | 0.13 | 4.28E-16 | Nucleotide (Purine) |
| ENPP1 | 0.11 | 1.54E-05 | Nucleotide |
| TK1 | 0.09 | 5.22E-04 | Nucleotide (Pyrimidine) |
| NPR1 | 0.09 | 1.85E-03 | Nucleotide (cGMP) |
| RRM2 | 0.08 | 2.01E-03 | Nucleotide |
| CDA | 0.08 | 6.27E-06 | Nucleotide (Pyrimidine) |
| OASL | 0.06 | 2.13E-04 | Nucleotide (Ribonuclease) |
| FHIT | 0.05 | 9.39E-03 | Nucleotide (Purine) |
| OAS2 | 0.05 | 3.31E-05 | Nucleotide (Ribonuclease) |
| GMPR | 0.04 | 2.01E-06 | Nucleotide (Purine) |
| ADCY4 | 0.04 | 2.24E-06 | Nucleotide (cAMP) |
| AMPD1 | 0.03 | 3.03E-02 | Nucleotide (Purine) |
| ENPP2 | 0.01 | 7.51E-05 | Nucleotide |
| ADAD2 | 0.01 | 2.14E-05 | Nucleotide (Purine) |
| ENPP6 | 4.80E-03 | 5.94E-03 | Nucleotide |
| UROS | 1.83 | 8.15E-07 | Porphyrin |
| PPOX | 1.51 | 2.75E-05 | Porphyrin |
| ALAS1 | 0.74 | 5.55E-04 | Porphyrin |
| FECH | 0.68 | 3.90E-03 | Porphyrin |
| UROD | 0.66 | 1.35E-06 | Porphyrin |
| BLVRA | 0.61 | 4.20E-07 | Porphyrin |
| ALAD | 0.58 | 1.22E-04 | Porphyrin |
| FXN | 0.41 | 5.86E-08 | Porphyrin |
| BLVRB | 0.22 | 6.61E-12 | Porphyrin |
| HMOX1 | 0.03 | 6.43E-05 | Porphyrin |
| ALAS2 | 0.02 | 1.99E-02 | Porphyrin |
| ATP6V0D2 | 99.85 | 4.60E-07 | Proton Transport |
| ATP8A2 | 23.99 | 7.86E-09 | Proton Transport |
| ATP6V1B1 | 9.22 | 1.80E-08 | Proton Transport |
| ATP13A2 | 5.11 | 4.40E-06 | Proton Transport |
| ATP9A | 3.69 | 1.05E-06 | Proton Transport |
| ATP6V1B2 | 3.25 | 6.56E-11 | Proton Transport |
| ATP6V0A1 | 2.82 | 1.41E-10 | Proton Transport |
| ATP6V1E2 | 2.35 | 5.09E-06 | Proton Transport |
| ATP13A5 | 2.27 | 2.95E-02 | Proton Transport |
| SLC25A27 | 2.03 | 1.26E-04 | Proton Transport |
| ATP5SL | 1.88 | 5.75E-08 | Proton Transport |
| ATP5A1 | 1.77 | 3.09E-07 | Proton Transport |
| ATP6V1G2 | 1.59 | 4.17E-06 | Proton Transport |
| ATP5B | 1.59 | 1.76E-07 | Proton Transport |
| ATP6V1C1 | 1.46 | 1.45E-05 | Proton Transport |
| ATP5S | 1.43 | 1.17E-04 | Proton Transport |
| ATP6V1A | 1.35 | 5.67E-03 | Proton Transport |
| ATP8A1 | 1.34 | 3.30E-02 | Proton Transport |
| ATP6V1D | 1.31 | 1.25E-05 | Proton Transport |
| ATP6V0A2 | 0.76 | 2.65E-03 | Proton Transport |
| ATP5G2 | 0.76 | 1.21E-04 | Proton Transport |
| ATP10D | 0.61 | 5.12E-04 | Proton Transport |
| ATP13A3 | 0.58 | 5.20E-04 | Proton Transport |
| ATP5F1 | 0.52 | 3.73E-08 | Proton Transport |
| ATP5E | 0.47 | 1.20E-09 | Proton Transport |
| TAP1 | 0.38 | 8.01E-06 | Proton Transport |
| ATP13A4 | 0.35 | 7.74E-03 | Proton Transport |
| ATP6V0A4 | 0.35 | 8.25E-03 | Proton Transport |
| ATP8B2 | 0.22 | 2.71E-04 | Proton Transport |
| UCP2 | 0.16 | 2.81E-05 | Proton Transport |
| HVCN1 | 0.16 | 1.38E-05 | Proton Transport |
| ATP6V0E1 | 0.13 | 1.21E-13 | Proton Transport |
| ATP10A | 0.10 | 1.46E-04 | Proton Transport |
| TCIRG1 | 0.10 | 1.79E-03 | Proton Transport |
| ATP8B1 | 0.06 | 7.56E-06 | Proton Transport |
| ATP8B4 | 0.05 | 1.16E-04 | Proton Transport |
| ATP10B | 0.01 | 1.59E-03 | Proton Transport |
| GRXCR1 | 16.27 | 4.60E-02 | Redox |
| DHRS2 | 14.54 | 7.49E-03 | Redox |
| DHRS11 | 5.26 | 2.61E-07 | Redox |
| CYB5R2 | 4.41 | 5.06E-03 | Redox |
| CYB5D1 | 2.90 | 5.31E-08 | Redox |
| PRDX2 | 2.26 | 3.26E-10 | Redox |
| DHRS7 | 2.05 | 1.43E-06 | Redox |
| MICAL1 | 1.84 | 1.71E-03 | Redox |
| ZADH2 | 1.84 | 3.62E-09 | Redox |
| CYB5R4 | 1.84 | 8.55E-08 | Redox |
| NDOR1 | 1.72 | 2.85E-04 | Redox |
| PYROXD1 | 1.66 | 3.37E-05 | Redox |
| FOXRED1 | 1.51 | 2.75E-04 | Redox |
| CYB561 | 1.43 | 2.91E-03 | Redox |
| CRYZL1 | 1.41 | 2.51E-03 | Redox |
| FDX1L | 0.74 | 1.73E-03 | Redox |
| CYB5D2 | 0.73 | 6.66E-03 | Redox |
| FDXR | 0.73 | 3.49E-02 | Redox |
| OXA1L | 0.72 | 1.60E-04 | Redox |
| DHTKD1 | 0.72 | 3.65E-04 | Redox |
| CYB5R1 | 0.69 | 4.08E-04 | Redox |
| GLRX5 | 0.66 | 5.67E-06 | Redox |
| PRDX6 | 0.57 | 1.22E-02 | Redox |
| DHRS9 | 0.57 | 4.40E-02 | Redox |
| FDX1 | 0.56 | 5.18E-07 | Redox |
| PRDX4 | 0.53 | 1.53E-05 | Redox |
| NMRAL1 | 0.50 | 2.99E-07 | Redox |
| DHRS7B | 0.49 | 1.07E-05 | Redox |
| CYBASC3 | 0.48 | 5.89E-04 | Redox |
| PRDX3 | 0.48 | 9.21E-11 | Redox |
| PRDX1 | 0.48 | 5.10E-08 | Redox |
| CYB5R3 | 0.44 | 1.02E-05 | Redox |
| POR | 0.44 | 1.80E-03 | Redox |
| TXNRD3 | 0.41 | 1.19E-07 | Redox |
| DHRS4 | 0.40 | 4.11E-05 | Redox |
| DHRS4L2 | 0.40 | 4.11E-05 | Redox |
| ENOX2 | 0.37 | 6.47E-09 | Redox |
| TXNRD1 | 0.26 | 7.03E-09 | Redox |
| CRYZ | 0.24 | 1.87E-04 | Redox |
| CYBB | 0.22 | 2.35E-03 | Redox |
| CYBRD1 | 0.20 | 1.47E-07 | Redox |
| FRRS1 | 0.19 | 6.20E-07 | Redox |
| CYB5A | 0.12 | 2.83E-09 | Redox |
| DHRS3 | 0.07 | 4.13E-09 | Redox |
| CYBA | 0.04 | 2.43E-04 | Redox |
| SLC17A6 | 365.67 | 4.37E-04 | Small Molecule Transport (Glutamate, Asparate, Lactate) |
| SLC32A1 | 181.09 | 1.45E-07 | Small Molecule Transport (GABA, Gly) |
| SLC17A7 | 140.52 | 5.82E-12 | Small Molecule Transport (Glutamate, Asparate, Lactate) |
| SLC6A11 | 131.50 | 4.39E-07 | Small Molecule Transport (GABA, β-Alanine) |
| SLC47A2 | 97.84 | 1.65E-02 | Small Molecule Transport (Thiamine) |
| AQP4 | 90.18 | 3.60E-11 | Small Molecule Transport (Water) |
| SLC5A8 | 86.18 | 7.00E-06 | Small Molecule Transport (iodide) |
| SLC1A2 | 84.03 | 1.09E-08 | Small Molecule Transport (Glutamate, Asparate) |
| SLC1A6 | 81.47 | 3.44E-10 | Small Molecule Transport (Glutamate) |
| RHCG | 76.24 | 3.08E-04 | Small Molecule Transport (Ammonium) |
| SLC17A8 | 56.11 | 1.56E-08 | Small Molecule Transport (Glutamate, Asparate, Lactate) |
| SLC34A1 | 49.90 | 3.83E-12 | Small Molecule Transport (Mitochondrial Phosphate) |
| SLC6A1 | 48.51 | 5.67E-10 | Small Molecule Transport (GABA) |
| SLC13A2 | 47.05 | 5.00E-05 | Small Molecule Transport (Citrate, Succinate) |
| SV2B | 43.44 | 7.42E-12 | Small Molecule Transport (Sugar) |
| SLC38A1 | 39.69 | 2.71E-12 | Small Molecule Transport (Gln, Ala, Ser, Asn) |
| SLC13A5 | 38.32 | 4.32E-10 | Small Molecule Transport (Citrate, Pyruvate) |
| SLC35F4 | 25.71 | 6.49E-10 | Small Molecule Transport (Nucleotide sugar) |
| SLC26A3 | 23.20 | 8.34E-08 | Small Molecule Transport (Biocarbonate) |
| SV2C | 22.75 | 1.37E-03 | Small Molecule Transport (Sugar) |
| SLC22A7 | 22.16 | 1.61E-04 | Small Molecule Transport (Anion) |
| SLC7A14 | 20.56 | 2.53E-09 | Small Molecule Transport (Amino acid) |
| SVOP | 19.69 | 4.07E-14 | Small Molecule Transport |
| SLC44A5 | 19.24 | 1.23E-07 | Small Molecule Transport (Choline) |
| SV2A | 15.32 | 3.55E-10 | Small Molecule Transport (Sugar) |
| SLCO5A1 | 15.29 | 5.65E-06 | Small Molecule Transport (peptide) |
| SLC6A7 | 13.61 | 9.89E-08 | Small Molecule Transport (Proline) |
| SLC35F1 | 13.41 | 1.18E-11 | Small Molecule Transport (Nucleotide sugar) |
| SLC27A6 | 13.39 | 4.21E-04 | Small Molecule Transport (Fatty acid) |
| SLC34A2 | 12.34 | 2.60E-04 | Small Molecule Transport (Mitochondrial Phosphate) |
| SLC4A7 | 11.37 | 6.87E-06 | Small Molecule Transport (Biocarbonate) |
| AQP5 | 11.04 | 3.42E-05 | Small Molecule Transport (Water) |
| SLC1A3 | 10.54 | 2.94E-06 | Small Molecule Transport (Glutamate, Asparate) |
| SLC4A3 | 9.47 | 2.74E-07 | Small Molecule Transport (Anion) |
| SLC5A4 | 9.41 | 6.48E-04 | Small Molecule Transport (Glucose) |
| SLC15A2 | 9.07 | 6.58E-10 | Small Molecule Transport (peptide) |
| SLC35F3 | 8.87 | 7.69E-05 | Small Molecule Transport (Nucleotide sugar) |
| SLC1A7 | 8.43 | 2.47E-10 | Small Molecule Transport (Glutamate) |
| SLC6A6 | 7.86 | 7.58E-06 | Small Molecule Transport (GABA,  β-alanine, taurine) |
| SLC6A19 | 7.85 | 1.97E-04 | Small Molecule Transport (BCAA) |
| SLC25A22 | 7.60 | 1.42E-09 | Small Molecule Transport (Glutamate) |
| MFSD4 | 7.32 | 1.06E-08 | Small Molecule Transport (Glucose) |
| SLC35D3 | 7.05 | 1.49E-04 | Small Molecule Transport (Nucleotide sugar) |
| SLC38A3 | 6.55 | 3.72E-10 | Small Molecule Transport (His, Gln, Asn, Ala, Glutamate) |
| SLC34A3 | 6.49 | 1.49E-04 | Small Molecule Transport (Mitochondrial Phosphate) |
| SLC18A1 | 6.32 | 2.50E-03 | Small Molecule Transport (Monoamine) |
| SLC22A14 | 5.85 | 1.74E-05 | Small Molecule Transport (Organic Cation) |
| SLC6A5 | 5.61 | 1.17E-02 | Small Molecule Transport (Glycine) |
| STAR | 5.51 | 5.87E-05 | Small Molecule Transport (Cholesterol) |
| SLC2A11 | 5.21 | 7.94E-09 | Small Molecule Transport (Glucose, Fructose) |
| SLC22A20 | 5.15 | 2.92E-04 | Small Molecule Transport (Organic Cation) |
| SLC29A2 | 4.97 | 2.98E-09 | Small Molecule Transport (inosine) |
| SLC5A12 | 4.75 | 7.52E-04 | Small Molecule Transport (Nicotinic acid, Lactate, Pyruvate) |
| SLC4A11 | 4.60 | 2.00E-04 | Small Molecule Transport (Biocarbonate) |
| SLC4A8 | 4.45 | 1.96E-08 | Small Molecule Transport (Biocarbonate) |
| SLC35F2 | 4.37 | 1.21E-08 | Small Molecule Transport (Nucleotide sugar) |
| SLC6A17 | 4.29 | 1.93E-04 | Small Molecule Transport (Pro, Cys, Ala) |
| SLC4A10 | 4.17 | 5.53E-08 | Small Molecule Transport (Biocarbonate) |
| STARD9 | 3.86 | 1.22E-04 | Small Molecule Transport (Lipid) |
| SLC5A7 | 3.81 | 1.31E-03 | Small Molecule Transport (Choline) |
| SLC22A15 | 3.51 | 6.71E-06 | Small Molecule Transport (Organic Cation) |
| SLC16A6 | 3.40 | 7.73E-06 | Small Molecule Transport (MCT activity) |
| SLC45A1 | 3.31 | 1.22E-04 | Small Molecule Transport (Glucose) |
| SLC25A25 | 3.21 | 2.30E-08 | Small Molecule Transport (Mitochondrial Phosphate) |
| SLC26A5 | 3.19 | 6.19E-04 | Small Molecule Transport (Biocarbonate) |
| SLC35E2 | 3.05 | 2.14E-06 | Small Molecule Transport (Nucleotide sugar) |
| SLC10A5 | 2.93 | 3.87E-03 | Small Molecule Transport (Bile acid) |
| SLCO4A1 | 2.87 | 2.24E-07 | Small Molecule Transport (bile acids, prostaglandins, steroid conjugates, thyroid hormone) |
| SLC36A4 | 2.68 | 1.46E-05 | Small Molecule Transport (Proline) |
| SLC7A3 | 2.66 | 2.52E-02 | Small Molecule Transport (Arg, Lys, Orn) |
| SLC26A2 | 2.65 | 7.84E-05 | Small Molecule Transport (Sulfate) |
| SLC38A4 | 2.65 | 2.92E-03 | Small Molecule Transport (His, Ar, Alan, Asn) |
| SLC38A5 | 2.56 | 9.66E-03 | Small Molecule Transport (Asn, Se, His, Gln) |
| FLVCR1 | 2.48 | 3.45E-08 | Small Molecule Transport (Hem) |
| SLC25A12 | 2.43 | 7.38E-09 | Small Molecule Transport (Glutamate, Aspartate) |
| SLC23A1 | 2.38 | 6.43E-03 | Small Molecule Transport (Vitamine C) |
| SLC25A14 | 2.34 | 2.86E-09 | Small Molecule Transport (MCT activity) |
| MFSD11 | 2.31 | 4.52E-10 | Small Molecule Transport |
| SLC35D1 | 2.24 | 4.50E-10 | Small Molecule Transport (UDP-N-Acetyl-Glucosomine) |
| SLC6A9 | 2.22 | 4.40E-05 | Small Molecule Transport (Glycine, Sarcosine) |
| SLC35B4 | 2.21 | 2.72E-08 | Small Molecule Transport (UDP-N-Acetyl-Glucosomine) |
| SFXN3 | 2.13 | 1.34E-09 | Small Molecule Transport (Serine) |
| SLC25A41 | 2.10 | 6.65E-05 | Small Molecule Transport (Nucleotide) |
| SLC25A40 | 2.08 | 1.77E-07 | Small Molecule Transport (Amino acid) |
| AQP3 | 2.01 | 1.07E-04 | Small Molecule Transport (Water) |
| SLC6A8 | 1.93 | 2.51E-04 | Small Molecule Transport (Creatine) |
| SLC26A8 | 1.92 | 2.82E-02 | Small Molecule Transport (Biocarbonate) |
| SLC27A4 | 1.91 | 1.36E-05 | Small Molecule Transport (Fatty acid) |
| SLC25A33 | 1.87 | 3.21E-06 | Small Molecule Transport (Nucleotide) |
| SLC37A4 | 1.85 | 1.14E-05 | Small Molecule Transport (G6P) |
| SLC26A6 | 1.85 | 1.85E-03 | Small Molecule Transport (Biocarbonate) |
| SLC25A23 | 1.83 | 4.65E-07 | Small Molecule Transport (Mitochondrial Phosphate) |
| SLC7A11 | 1.82 | 1.93E-02 | Small Molecule Transport (Cystine, Glutamate) |
| SLC46A1 | 1.78 | 1.05E-06 | Small Molecule Transport (Folate) |
| SLC20A1 | 1.75 | 3.29E-05 | Small Molecule Transport (Phosphate) |
| SLC29A1 | 1.74 | 3.25E-04 | Small Molecule Transport (inosine) |
| SLC37A1 | 1.73 | 1.40E-03 | Small Molecule Transport (G6P, G3P) |
| SLC15A4 | 1.69 | 9.88E-03 | Small Molecule Transport (peptide) |
| SFXN1 | 1.69 | 4.44E-08 | Small Molecule Transport (Serine) |
| SLC17A5 | 1.66 | 2.99E-05 | Small Molecule Transport (Glutamate, Asparate, Lactate) |
| SLC4A1AP | 1.66 | 8.13E-05 | Small Molecule Transport (Anion) |
| SLC7A1 | 1.57 | 2.14E-04 | Small Molecule Transport (Ar, Lys, Orn, His) |
| SLC18A2 | 1.56 | 2.45E-03 | Small Molecule Transport (Monoamine) |
| SLC25A24 | 1.55 | 3.18E-04 | Small Molecule Transport (Mitochondrial Phosphate) |
| SLC1A4 | 1.55 | 5.39E-06 | Small Molecule Transport (Alanine, Cysteine, Serine) |
| SLC25A44 | 1.54 | 1.36E-07 | Small Molecule Transport (Amino acid) |
| SLC2A3 | 1.54 | 1.50E-02 | Small Molecule Transport (Glucose) |
| SLC35C2 | 1.52 | 4.85E-06 | Small Molecule Transport (Nucleotide sugar) |
| SLC25A46 | 1.51 | 5.25E-05 | Small Molecule Transport (mitochondrial transport) |
| SLC26A11 | 1.50 | 1.09E-04 | Small Molecule Transport (Sulfate) |
| AQP11 | 1.48 | 3.13E-02 | Small Molecule Transport (Water, Glycerol) |
| SLC35E3 | 1.48 | 1.24E-04 | Small Molecule Transport (Nucleotide sugar) |
| DIRC2 | 1.47 | 1.17E-05 | Small Molecule Transport |
| SLC25A29 | 1.40 | 4.74E-03 | Small Molecule Transport (Acylcarnitine) |
| SFXN2 | 1.35 | 1.80E-02 | Small Molecule Transport (Serine) |
| MFSD9 | 1.35 | 6.81E-03 | Small Molecule Transport |
| SLC25A11 | 0.76 | 7.05E-03 | Small Molecule Transport (Malate, aKG) |
| SLC35B1 | 0.75 | 1.59E-02 | Small Molecule Transport (Nucleotide sugar) |
| SLC16A2 | 0.75 | 7.92E-03 | Small Molecule Transport (T4) |
| SLC35A1 | 0.74 | 3.69E-05 | Small Molecule Transport (Nucleotide sugar) |
| SLC2A8 | 0.72 | 9.78E-03 | Small Molecule Transport (Glucose) |
| SLC25A13 | 0.71 | 1.30E-02 | Small Molecule Transport (Glutamate, Aspartate) |
| SLC22A5 | 0.70 | 2.20E-02 | Small Molecule Transport (Carnitine) |
| SLC44A1 | 0.70 | 2.25E-02 | Small Molecule Transport (Choline) |
| SLC10A3 | 0.67 | 4.73E-05 | Small Molecule Transport (Bile acid) |
| TMEM104 | 0.67 | 1.94E-05 | Small Molecule Transport (Amino Acid) |
| SLC27A1 | 0.65 | 5.70E-06 | Small Molecule Transport (Fatty acid) |
| SLC4A4 | 0.62 | 3.17E-02 | Small Molecule Transport (Biocarbonate) |
| SLC46A3 | 0.61 | 1.46E-02 | Small Molecule Transport (Folate) |
| SLC25A39 | 0.61 | 2.85E-06 | Small Molecule Transport (Amino acid) |
| SLC16A1 | 0.60 | 2.53E-02 | Small Molecule Transport (Lactate/Pyruvate/3-HB) |
| NPC1 | 0.59 | 1.27E-06 | Small Molecule Transport (Cholesterol) |
| SLC38A2 | 0.58 | 3.49E-02 | Small Molecule Transport (Met, Gln, Ala, Ser, Asn) |
| SFXN4 | 0.58 | 1.12E-05 | Small Molecule Transport |
| SLC7A5 | 0.52 | 9.16E-05 | Small Molecule Transport (Amino acid) |
| CTNS | 0.52 | 7.53E-05 | Small Molecule Transport (Cystine) |
| SLC33A1 | 0.51 | 5.38E-04 | Small Molecule Transport (Acetyl CoA) |
| SLC25A45 | 0.51 | 1.22E-02 | Small Molecule Transport (Amino acid) |
| SLC35B3 | 0.50 | 1.77E-07 | Small Molecule Transport (Nucleotide sugar) |
| SLCO3A1 | 0.50 | 3.81E-05 | Small Molecule Transport (vasopressin, prostaglandins, thyroid hormones) |
| SLC25A10 | 0.50 | 2.44E-03 | Small Molecule Transport (Malate, Succinate) |
| SLC16A11 | 0.49 | 5.51E-03 | Small Molecule Transport (Pyryuvate) |
| SLC35C1 | 0.48 | 5.01E-04 | Small Molecule Transport (GDP-Fructose) |
| PCTP | 0.46 | 6.03E-06 | Small Molecule Transport (PC) |
| MFSD10 | 0.45 | 1.34E-02 | Small Molecule Transport (Anion) |
| SLC10A4 | 0.45 | 6.64E-04 | Small Molecule Transport (Bile acid) |
| SLC25A20 | 0.43 | 4.27E-08 | Small Molecule Transport (Carnitine) |
| MFSD7 | 0.40 | 4.59E-05 | Small Molecule Transport |
| SLC19A2 | 0.40 | 1.47E-08 | Small Molecule Transport (Thiamine) |
| SLC36A1 | 0.39 | 1.88E-06 | Small Molecule Transport (Glycine) |
| SLC45A3 | 0.39 | 3.29E-02 | Small Molecule Transport (Sugar) |
| SLC22A18AS | 0.38 | 2.26E-02 | Small Molecule Transport (Organic Cation) |
| SLC44A2 | 0.38 | 1.46E-08 | Small Molecule Transport (Choline) |
| SLC35A2 | 0.38 | 2.58E-04 | Small Molecule Transport (Nucleotide sugar) |
| SLC20A2 | 0.37 | 2.26E-08 | Small Molecule Transport (Phosphate) |
| SLC23A3 | 0.35 | 3.70E-02 | Small Molecule Transport (Citrate, Malate) |
| SLC29A3 | 0.35 | 1.04E-04 | Small Molecule Transport (inosine) |
| SLC25A1 | 0.35 | 6.60E-12 | Small Molecule Transport (Citrate, Malate) |
| MFSD1 | 0.35 | 2.42E-05 | Small Molecule Transport |
| SLC22A18 | 0.34 | 3.60E-04 | Small Molecule Transport (Organic Cation) |
| SLC5A6 | 0.34 | 5.17E-09 | Small Molecule Transport (Vitamine) |
| SLC7A2 | 0.34 | 2.16E-06 | Small Molecule Transport (Arg, Lys, Orn, His) |
| SLC27A3 | 0.31 | 2.69E-08 | Small Molecule Transport (Fatty acid) |
| SLC43A1 | 0.28 | 2.59E-09 | Small Molecule Transport (BCAA) |
| SLC13A3 | 0.28 | 5.52E-05 | Small Molecule Transport (Citrate, Succinate) |
| SLC19A1 | 0.27 | 2.64E-06 | Small Molecule Transport (Thiamine, Folate) |
| SLC7A8 | 0.27 | 1.63E-08 | Small Molecule Transport (Amino acid) |
| SLC27A2 | 0.26 | 7.57E-03 | Small Molecule Transport (Fatty acid) |
| SLC35D2 | 0.25 | 1.62E-09 | Small Molecule Transport (Nucleotide sugar) |
| SLC44A3 | 0.25 | 8.23E-06 | Small Molecule Transport (Choline) |
| STARD13 | 0.23 | 6.16E-07 | Small Molecule Transport (Lipid) |
| SLC4A5 | 0.18 | 1.34E-03 | Small Molecule Transport (Biocarbonate) |
| SLC37A2 | 0.18 | 1.03E-03 | Small Molecule Transport (G6P) |
| SLC7A7 | 0.16 | 5.69E-05 | Small Molecule Transport (Amino acid) |
| SLC44A4 | 0.15 | 1.12E-04 | Small Molecule Transport (Choline) |
| SLC2A10 | 0.15 | 8.83E-05 | Small Molecule Transport (Glucose) |
| SLC16A10 | 0.15 | 3.14E-02 | Small Molecule Transport (tryptophan, phenylalanine, levodopa, tyrosine) |
| AQP7 | 0.14 | 1.40E-03 | Small Molecule Transport (Water) |
| SLC29A4 | 0.14 | 3.37E-04 | Small Molecule Transport (Dopamine) |
| SLC4A2 | 0.14 | 5.11E-08 | Small Molecule Transport (Biocarbonate) |
| SLC22A11 | 0.14 | 1.23E-03 | Small Molecule Transport (Uric acid |
| SLC25A30 | 0.14 | 4.03E-09 | Small Molecule Transport (mitochondrial transport) |
| SLC7A10 | 0.09 | 1.64E-04 | Small Molecule Transport (Amino acid) |
| SLC25A43 | 0.09 | 5.12E-08 | Small Molecule Transport (Nucleotide) |
| SLC4A1 | 0.09 | 3.18E-02 | Small Molecule Transport (Anion) |
| SLC22A3 | 0.09 | 4.12E-05 | Small Molecule Transport (Monoamine) |
| SLC16A3 | 0.09 | 1.43E-04 | Small Molecule Transport (Lactate/Pyruvate) |
| SLCO2B1 | 0.07 | 1.46E-04 | Small Molecule Transport (T4) |
| SLCO4C1 | 0.07 | 1.02E-05 | Small Molecule Transport (cAMP, steroid conjugates, thyroid hormones) |
| SLC28A3 | 0.07 | 2.00E-03 | Small Molecule Transport (inosine) |
| FOLR1 | 0.07 | 9.53E-04 | Small Molecule Transport (Folate) |
| AQP1 | 0.06 | 7.28E-04 | Small Molecule Transport (Ammonium) |
| SLC16A5 | 0.06 | 8.85E-08 | Small Molecule Transport (MCT activity) |
| SLC2A7 | 0.06 | 7.44E-03 | Small Molecule Transport (Glucose, Fructose) |
| SLC6A12 | 0.06 | 3.86E-05 | Small Molecule Transport (GABA, Betaine |
| STARD8 | 0.05 | 8.14E-07 | Small Molecule Transport (Lipid) |
| AQP9 | 0.05 | 4.70E-03 | Small Molecule Transport (Water, Glycerol) |
| SLC1A5 | 0.04 | 2.96E-03 | Small Molecule Transport (Alanine, Cysteine, Serine) |
| SLC43A3 | 0.04 | 1.80E-04 | Small Molecule Transport (Amino acid) |
| SLCO1C1 | 0.04 | 3.48E-05 | Small Molecule Transport (Thyroid) |
| SLC16A4 | 0.04 | 3.68E-08 | Small Molecule Transport (MCT activity) |
| SLC2A12 | 0.03 | 3.38E-05 | Small Molecule Transport (Glucose) |
| SLC16A14 | 0.02 | 3.27E-04 | Small Molecule Transport (MCT activity) |
| RHAG | 0.02 | 2.49E-02 | Small Molecule Transport (Ammonium) |
| SLC26A4 | 0.02 | 1.64E-10 | Small Molecule Transport (Biocarbonate) |
| SLC22A6 | 0.02 | 3.56E-03 | Small Molecule Transport (Anion) |
| FLVCR2 | 0.01 | 3.63E-07 | Small Molecule Transport (Hem) |
| SLCO1B3 | 0.01 | 4.97E-02 | Small Molecule Transport (Bile acid) |
| SLC5A5 | 0.01 | 1.08E-04 | Small Molecule Transport (iodide) |
| SLC45A2 | 0.01 | 2.56E-05 | Small Molecule Transport (Sugar) |
| SLC26A7 | 0.01 | 8.52E-06 | Small Molecule Transport (Biocarbonate) |
| SLC6A13 | 0.01 | 1.08E-04 | Small Molecule Transport (GABA, β-Alanine) |
| STRA6 | 0.01 | 4.36E-06 | Small Molecule Transport (Vitamine A) |
| SLC16A8 | 0.01 | 2.37E-04 | Small Molecule Transport (Lactate) |
| SLCO1A2 | 0.01 | 4.88E-04 | Small Molecule Transport (Bile acid) |
| SLC22A8 | 0.01 | 3.77E-05 | Small Molecule Transport (Urate) |
| SLC6A20 | 0.01 | 2.25E-05 | Small Molecule Transport (Proline) |
| SLC14A1 | 0.01 | 2.23E-03 | Small Molecule Transport (Urea) |
| SLC14A2 | 0.01 | 3.93E-07 | Small Molecule Transport (Urea) |
| SLC16A12 | 0.01 | 1.58E-04 | Small Molecule Transport (Creatine) |
| SLC5A10 | 3.92E-03 | 1.51E-05 | Small Molecule Transport (Glucose) |
| FOLR2 | 2.66E-03 | 1.47E-04 | Small Molecule Transport (Folate) |
| FOLR3 | 2.66E-03 | 1.47E-04 | Small Molecule Transport (Folate) |
| SLCO2A1 | 1.10E-03 | 8.12E-06 | Small Molecule Transport (eicosanoids, prostaglandins) |
| G6PC2 | 173.34 | 1.97E-06 | Sugar (gluconeogenesi) |
| TKTL2 | 12.75 | 1.25E-05 | Sugar (PPP) |
| PIK3R4 | 7.36 | 1.16E-07 | Sugar (Inositol Phosphate) |
| AMY2A | 6.17 | 2.73E-02 | Sugar (Amylase) |
| PIP5KL1 | 5.05 | 4.47E-07 | Sugar (Inositol Phosphate) |
| INPP5J | 4.45 | 5.62E-08 | Sugar (Inositol Phosphate) |
| INPP5F | 3.87 | 3.39E-12 | Sugar (Inositol Phosphate) |
| TKTL1 | 3.63 | 1.42E-05 | Sugar (PPP) |
| PGAP1 | 3.49 | 3.36E-06 | Sugar (Inositol Phosphate) |
| INPP4A | 3.16 | 4.40E-08 | Sugar (Inositol Phosphate) |
| KHK | 3.11 | 7.46E-09 | Sugar (Fructose) |
| AGK | 2.52 | 2.81E-10 | Sugar (PPP) |
| SYNJ1 | 2.32 | 1.71E-07 | Sugar (Inositol Phosphate) |
| AGL | 2.30 | 3.79E-07 | Sugar (Glycogen) |
| INPP5E | 2.29 | 4.63E-06 | Sugar (Inositol Phosphate) |
| IP6K2 | 2.16 | 3.57E-07 | Sugar (Inositol Phosphate) |
| PIP4K2B | 2.11 | 9.44E-14 | Sugar (Inositol Phosphate) |
| IP6K1 | 2.09 | 5.59E-07 | Sugar (Inositol Phosphate) |
| PPA1 | 2.08 | 3.03E-08 | Sugar (PPi hydrolysis) |
| MPI | 1.91 | 4.97E-07 | Sugar (Mannose) |
| DERA | 1.89 | 3.35E-05 | Sugar (PPP) |
| XYLB | 1.87 | 1.03E-03 | Sugar (Xyulose) |
| PIP5K1C | 1.83 | 6.98E-10 | Sugar (Inositol Phosphate) |
| ITPKA | 1.81 | 2.17E-03 | Sugar (Inositol Phosphate) |
| PYGB | 1.79 | 2.94E-05 | Sugar (Glycogen) |
| PGM3 | 1.71 | 1.22E-05 | Sugar (Aminosugar) |
| PIK3R1 | 1.71 | 3.86E-04 | Sugar (Inositol Phosphate) |
| IPMK | 1.63 | 2.33E-04 | Sugar (Inositol Phosphate) |
| ABHD10 | 1.61 | 2.03E-06 | Sugar |
| GFOD2 | 1.61 | 9.47E-03 | Sugar |
| MINPP1 | 1.58 | 6.73E-06 | Sugar (Inositol Phosphate) |
| PI4KB | 1.50 | 8.11E-09 | Sugar (Inositol Phosphate) |
| PIK3CA | 1.48 | 3.81E-03 | Sugar (Inositol Phosphate) |
| PIK3CB | 1.46 | 1.76E-05 | Sugar (Inositol Phosphate) |
| NAGK | 1.39 | 3.01E-05 | Sugar (Aminosugar) |
| PGD | 0.76 | 2.17E-02 | Sugar (PPP) |
| GAA | 0.75 | 3.56E-02 | Sugar (Glycogen) |
| GNPTG | 0.75 | 1.57E-06 | Sugar (Aminosugar) |
| PTEN | 0.71 | 1.27E-02 | Sugar (Inositol Phosphate) |
| TSTA3 | 0.69 | 1.03E-02 | Sugar |
| GALK2 | 0.67 | 5.02E-03 | Sugar (Galactose) |
| G6PC3 | 0.65 | 7.20E-03 | Sugar (gluconeogenesi) |
| RPE | 0.63 | 1.48E-03 | Sugar (PPP) |
| GRHPR | 0.63 | 7.35E-09 | Sugar |
| PIK3C2A | 0.57 | 3.95E-08 | Sugar (Inositol Phosphate) |
| SORD | 0.56 | 1.64E-06 | Sugar |
| UXS1 | 0.56 | 3.09E-08 | Sugar (Nucleotide Sugar) |
| TGDS | 0.54 | 2.08E-08 | Sugar (Nucleotide Sugar) |
| PGLS | 0.54 | 2.09E-05 | Sugar (PPP) |
| DCXR | 0.53 | 1.04E-05 | Sugar (Xyulose) |
| PIP4K2A | 0.53 | 3.62E-04 | Sugar (Inositol Phosphate) |
| TKT | 0.52 | 3.09E-06 | Sugar (PPP) |
| HYI | 0.50 | 1.82E-04 | Sugar |
| GNPTAB | 0.47 | 2.61E-06 | Sugar (Aminosugar) |
| PCK2 | 0.46 | 1.49E-06 | Sugar (gluconeogenesi) |
| TALDO1 | 0.46 | 1.10E-06 | Sugar (PPP) |
| RPIA | 0.44 | 3.45E-04 | Sugar (PPP) |
| GMPPB | 0.44 | 2.17E-02 | Sugar (Mannose) |
| GNPDA1 | 0.44 | 1.41E-07 | Sugar (Aminosugar) |
| GALK1 | 0.42 | 6.76E-04 | Sugar (Galactose) |
| UAP1L1 | 0.41 | 7.65E-06 | Sugar (Aminosugar) |
| GBE1 | 0.41 | 1.02E-07 | Sugar (Glycogen) |
| INPP4B | 0.39 | 9.77E-06 | Sugar (Inositol Phosphate) |
| IP6K3 | 0.38 | 4.30E-04 | Sugar (Inositol Phosphate) |
| UGDH | 0.37 | 5.72E-06 | Sugar (Nucleotide Sugar) |
| CRYL1 | 0.35 | 4.48E-08 | Sugar |
| SYNJ2 | 0.35 | 3.45E-04 | Sugar (Inositol Phosphate) |
| GFPT2 | 0.35 | 6.43E-04 | Sugar (Aminosugar) |
| SIAE | 0.33 | 6.14E-05 | Sugar |
| PYGM | 0.32 | 2.05E-04 | Sugar (Glycogen) |
| H6PD | 0.30 | 3.30E-03 | Sugar |
| NPL | 0.29 | 4.19E-05 | Sugar (Aminosugar) |
| PMM2 | 0.28 | 8.72E-06 | Sugar (Mannose) |
| RENBP | 0.25 | 4.26E-08 | Sugar (Aminosugar) |
| NANS | 0.22 | 3.78E-03 | Sugar (Aminosugar) |
| GALM | 0.21 | 3.53E-11 | Sugar (Galactose) |
| IMPA2 | 0.19 | 6.02E-07 | Sugar (Inositol Phosphate) |
| PIK3CD | 0.16 | 3.24E-09 | Sugar (Inositol Phosphate) |
| ISYNA1 | 0.13 | 1.89E-07 | Sugar (Inositol Phosphate) |
| MGAM | 0.11 | 3.23E-03 | Sugar (Mannose) |
| FBP1 | 0.11 | 1.07E-10 | Sugar (gluconeogenesi) |
| INPP5D | 0.10 | 6.32E-07 | Sugar (Inositol Phosphate) |
| GYG2 | 0.10 | 1.63E-03 | Sugar (Glycogen) |
| GMDS | 0.10 | 2.31E-07 | Sugar (Mannose) |
| PIK3R6 | 0.06 | 3.43E-04 | Sugar (Inositol Phosphate) |
| PIK3CG | 0.05 | 2.22E-05 | Sugar (Inositol Phosphate) |
| INPP5K | 0.05 | 1.94E-05 | Sugar (Inositol Phosphate) |
| PIK3R5 | 0.03 | 2.73E-06 | Sugar (Inositol Phosphate) |
| ITPKB | 0.03 | 9.92E-12 | Sugar (Inositol Phosphate) |
| FBP2 | 0.01 | 1.71E-02 | Sugar (gluconeogenesi) |
| SULT2A1 | 23.25 | 1.38E-02 | Sulfate |
| SULT4A1 | 2.02 | 1.11E-03 | Sulfate |
| ARSG | 1.48 | 4.81E-03 | Sulfate |
| ARSA | 0.77 | 2.71E-03 | Sulfate |
| DSEL | 0.66 | 1.63E-02 | Sulfate |
| ARSB | 0.60 | 2.19E-02 | Sulfate |
| SULT1C4 | 0.59 | 1.03E-05 | Sulfate |
| ARSI | 0.28 | 9.21E-08 | Sulfate |
| SULT1A1 | 0.25 | 1.62E-05 | Sulfate |
| TST | 0.17 | 8.79E-07 | Sulfate |
| MPST | 0.16 | 1.88E-10 | Sulfate |
| DSE | 0.11 | 1.31E-03 | Sulfate |
| PAPSS2 | 0.07 | 1.34E-04 | Sulfate |
| SULT1A2 | 0.06 | 2.66E-05 | Sulfate |
| SULT1B1 | 0.04 | 4.38E-04 | Sulfate |
| SULT1E1 | 0.01 | 3.06E-02 | Sulfate |
| SULT1C2 | 0.01 | 4.58E-06 | Sulfate |
| GOT1 | 4.32 | 1.42E-10 | TCA Cycle |
| MDH1B | 4.03 | 2.56E-06 | TCA Cycle |
| L2HGDH | 2.82 | 1.69E-10 | TCA Cycle |
| ME2 | 2.56 | 3.06E-08 | TCA Cycle |
| OGDHL | 2.50 | 3.33E-06 | TCA Cycle |
| PDHX | 2.20 | 2.43E-10 | TCA Cycle |
| PDHA1 | 2.14 | 1.81E-08 | TCA Cycle |
| IDH3A | 1.97 | 1.91E-07 | TCA Cycle |
| CS | 1.96 | 3.02E-10 | TCA Cycle |
| MDH1 | 1.91 | 1.25E-08 | TCA Cycle |
| ME1 | 1.81 | 2.68E-06 | TCA Cycle |
| DLD | 1.79 | 3.32E-07 | TCA Cycle |
| SUCLA2 | 1.57 | 9.89E-10 | TCA Cycle |
| ACO2 | 1.55 | 7.74E-06 | TCA Cycle |
| FH | 1.51 | 1.54E-08 | TCA Cycle |
| IDH3B | 1.35 | 6.17E-05 | TCA Cycle |
| ME3 | 0.67 | 3.82E-04 | TCA Cycle |
| PDHB | 0.60 | 1.33E-05 | TCA Cycle |
| SUCLG1 | 0.56 | 4.85E-06 | TCA Cycle |
| GOT2 | 0.55 | 9.43E-03 | TCA Cycle |
| PC | 0.54 | 3.12E-04 | TCA Cycle |
| IDH1 | 0.32 | 4.83E-06 | TCA Cycle |
| ACO1 | 0.28 | 3.22E-06 | TCA Cycle |
| IDH2 | 0.22 | 6.43E-08 | TCA Cycle |
| SUCLG2 | 0.20 | 1.49E-10 | TCA Cycle |
| COQ3 | 1.81 | 3.82E-03 | Ubiquinone |
| COQ7 | 1.52 | 2.69E-05 | Ubiquinone |
| PDSS1 | 0.60 | 5.79E-04 | Ubiquinone |
| SQRDL | 0.09 | 2.84E-05 | Ubiquinone |
| RDH8 | 109.28 | 1.14E-07 | Vitamin (A) |
| RDH12 | 64.02 | 4.63E-10 | Vitamin (A) |
| SDR16C5 | 29.24 | 1.65E-05 | Vitamin (A) |
| PDXP | 4.45 | 1.16E-10 | Vitamin (B6) |
| RDH16 | 3.02 | 3.74E-03 | Vitamin (A) |
| GPHN | 2.46 | 5.46E-10 | Vitamin (Folate) |
| PDXK | 2.17 | 3.95E-07 | Vitamin (B6) |
| MTHFR | 2.00 | 2.97E-05 | Vitamin (Folate) |
| BCO2 | 1.98 | 7.55E-05 | Vitamin (A) |
| VKORC1L1 | 1.81 | 2.09E-04 | Vitamin (K) |
| MTHFD2L | 1.67 | 4.01E-06 | Vitamin (Folate) |
| PHOSPHO2 | 1.63 | 1.03E-06 | Vitamin (B6) |
| DHFR | 1.61 | 2.62E-02 | Vitamin (Folate) |
| ALDH8A1 | 1.57 | 1.76E-05 | Vitamine (Retinoic acid, Alcohol) |
| BTD | 1.56 | 6.79E-04 | Vitamin (Biotin) |
| RFK | 1.38 | 2.28E-06 | Vitamin (Riboflavin) |
| MTHFSD | 0.66 | 9.58E-06 | Vitamin (Folate) |
| PNPO | 0.65 | 3.83E-06 | Vitamin (B6) |
| AKR7A2 | 0.64 | 4.46E-04 | Vitamine (Retinoic acid, Alcohol) |
| MMADHC | 0.57 | 8.99E-10 | Vitamin (B12) |
| AKR1B1 | 0.57 | 4.94E-08 | Vitamine (Retinoic acid, Alcohol) |
| MTHFD1L | 0.48 | 1.03E-02 | Vitamin (Folate) |
| MTHFD1 | 0.48 | 5.26E-08 | Vitamin (Folate) |
| AKR1A1 | 0.47 | 7.07E-09 | Vitamine (Retinoic acid, Alcohol) |
| FPGS | 0.44 | 5.31E-05 | Vitamin (Folate) |
| ADH5 | 0.44 | 1.59E-08 | Vitamine (Retinoic acid, Alcohol) |
| RETSAT | 0.43 | 4.37E-06 | Vitamin (A) |
| SHMT1 | 0.36 | 8.36E-08 | Vitamin (Folate) |
| TPK1 | 0.28 | 2.39E-09 | Vitamin (Thiamine) |
| CYP26B1 | 0.24 | 1.43E-02 | Vitamin (A) |
| RDH11 | 0.22 | 8.28E-04 | Vitamin (A) |
| RDH10 | 0.20 | 2.27E-05 | Vitamin (A) |
| ALDH1A2 | 0.14 | 5.30E-05 | Vitamine (Retinoic acid, Alcohol) |
| MTHFD2 | 0.07 | 2.71E-03 | Vitamin (Folate) |
| ADH4 | 0.05 | 4.75E-02 | Vitamine (Retinoic acid, Alcohol) |
| ADH1A | 0.02 | 1.00E-02 | Vitamine (Retinoic acid, Alcohol) |
| ADH1C | 0.02 | 3.73E-02 | Vitamine (Retinoic acid, Alcohol) |
| ALDH1A1 | 0.01 | 1.49E-06 | Vitamine (Retinoic acid, Alcohol) |
| LRAT | 0.01 | 9.22E-06 | Vitamin (A) |
| BCMO1 | 0.01 | 1.42E-05 | Vitamin (A) |
| ALDH1A3 | 0.01 | 3.06E-03 | Vitamine (Retinoic acidl, Aldehyde) |
| RPE65 | 0.01 | 6.00E-05 | Vitamin (A) |
| ADH1B | 1.65E-03 | 3.42E-03 | Vitamine (Retinoic acid, Alcohol) |
| AKR1B10 | 9.14E-04 | 2.10E-05 | Vitamine (Retinoic acid, Alcohol) |

**Table S14. Differentially expressed major metabolic genes between retinas and RPE/choroid.**

| **Gene symbol** | **Classification (Sub-category/substrate)** | **In Figures** |
| --- | --- | --- |
| SRM | Amino Acid (Polyamine) | Fig 5b |
| SMOX | Amino Acid (Polyamine) | Fig 5b |
| SAT1 | Amino Acid (Polyamine) | Fig 5b |
| NAT8L | Amino Acid (Acetylation) | Fig 5b |
| ADC | Amino Acid (Arginine) | Fig 5b |
| DDAH1 | Amino Acid (Arginine, Citrulline) | Fig 5b |
| OTC | Amino Acid (Arginine, Urea) | Fig 5b |
| ARG2 | Amino Acid (Arginine, Urea) | Fig 5b |
| ASL | Amino Acid (Arginine, Urea) | Fig 5b |
| ASS1 | Amino Acid (Arginine, Urea) | Fig 5b |
| ARG1 | Amino Acid (Arginine, Urea) | Fig 5b |
| ASNS | Amino Acid (Asparagine) | Fig 5b |
| ASPA | Amino Acid (Aspartate) | Fig 5b |
| MOCS1 | Amino Acid (Cofactor) | Fig 5b |
| CKMT1B | Amino Acid (Creatine) | Fig 5b |
| CKM | Amino Acid (Creatine) | Fig 5b |
| CKB | Amino Acid (Creatine) | Fig 5b |
| GAMT | Amino Acid (Creatine) | Fig 5b |
| CCBL2 | Amino Acid (Cysteine) | Fig 5b |
| CBS | Amino Acid (Cysteine) | Fig 5b |
| ADO | Amino Acid (Cysteine) | Fig 5b |
| CDO1 | Amino Acid (Cysteine) | Fig 5b |
| GAD2 | Amino Acid (Glutamate) | Fig 5b |
| GAD1 | Amino Acid (Glutamate) | Fig 5b |
| GLS2 | Amino Acid (Glutamate) | Fig 5b |
| GLS | Amino Acid (Glutamate) | Fig 5b |
| ALDH5A1 | Amino Acid (Glutamate) | Fig 5b |
| ALDH18A1 | Amino Acid (Glutamate) | Fig 5b |
| GPT | Amino Acid (Glutamate, Alaline) | Fig 5b |
| GPT2 | Amino Acid (Glutamate, Alaline) | Fig 5b |
| GPX6 | Amino Acid (Glutathione) | Fig 5b |
| GPX5 | Amino Acid (Glutathione) | Fig 5b |
| GPX2 | Amino Acid (Glutathione) | Fig 5b |
| GPX3 | Amino Acid (Glutathione) | Fig 5b |
| GPX4 | Amino Acid (Glutathione) | Fig 5b |
| GPX7 | Amino Acid (Glutathione) | Fig 5b |
| GPX1 | Amino Acid (Glutathione) | Fig 5b |
| GPX8 | Amino Acid (Glutathione) | Fig 5b |
| BHMT2 | Amino Acid (Glycine) | Fig 5b |
| GLYATL1 | Amino Acid (Glycine) | Fig 5b |
| GLYATL2 | Amino Acid (Glycine) | Fig 5b |
| GLYAT | Amino Acid (Glycine) | Fig 5b |
| GCSH | Amino Acid (Glycine) | Fig 5b |
| DMGDH | Amino Acid (Glycine) | Fig 5b |
| MTR | Amino Acid (Methionine) | Fig 5b |
| ALDH1L1 | Amino Acid (One carbon) | Fig 5b |
| ALDH1L2 | Amino Acid (One carbon) | Fig 5b |
| PRODH2 | Amino Acid (Proline) | Fig 5b |
| PYCR1 | Amino Acid (Proline) | Fig 5b |
| P4HA2 | Amino Acid (Proline) | Fig 5b |
| PEPD | Amino Acid (Proline) | Fig 5b |
| P4HA1 | Amino Acid (Proline) | Fig 5b |
| P4HB | Amino Acid (Proline) | Fig 5b |
| ALDH4A1 | Amino Acid (Proline) | Fig 5b |
| PRODH | Amino Acid (Proline) | Fig 5b |
| P4HA3 | Amino Acid (Proline) | Fig 5b |
| PHGDH | Amino Acid (Serine) | Fig 5b |
| PSAT | Amino Acid (Serine) | Fig 5b |
| KMO | Amino Acid (Tryptophan) | Fig 5b |
| KYNU | Amino Acid (Tryptophan) | Fig 5b |
| TDO2 | Amino Acid (Tryptophan) | Fig 5b |
| INMT | Amino Acid (Tryptophan) | Fig 5b |
| HAAO | Amino Acid (Tryptophan, NAD) | Fig 5b |
| HKDC1 | Glycolysis | Fig 5b |
| ENO2 | Glycolysis | Fig 5b |
| HK2 | Glycolysis | Fig 5b |
| PFKP | Glycolysis | Fig 5b |
| ALDOC | Glycolysis | Fig 5b |
| PFKFB4 | Glycolysis | Fig 5b |
| GPI | Glycolysis | Fig 5b |
| PGAM2 | Glycolysis | Fig 5b |
| PGK1 | Glycolysis | Fig 5b |
| PFKM | Glycolysis | Fig 5b |
| TPI1 | Glycolysis | Fig 5b |
| PGAM1 | Glycolysis | Fig 5b |
| PGM2 | Glycolysis | Fig 5b |
| HK1 | Glycolysis | Fig 5b |
| LDHA | Glycolysis | Fig 5b |
| GAPDH | Glycolysis | Fig 5b |
| PFKFB2 | Glycolysis | Fig 5b |
| ENO3 | Glycolysis | Fig 5b |
| ENO1 | Glycolysis | Fig 5b |
| PGM2L1 | Glycolysis | Fig 5b |
| PGAM4 | Glycolysis | Fig 5b |
| ALDOA | Glycolysis | Fig 5b |
| PGM5 | Glycolysis | Fig 5b |
| ADPGK | Glycolysis | Fig 5b |
| PFKFB3 | Glycolysis | Fig 5b |
| HK3 | Glycolysis | Fig 5b |
| GAPDHS | Glycolysis | Fig 5b |
| GLOD5 | Glyoxylate | Fig 5b |
| SLC12A8 | Ion Transport | Fig 5b |
| SLC3A2 | Ion Transport | Fig 5b |
| SPTLC1 | Lipid (Sphingolipid) | Fig 5b |
| NMNAT2 | NAD | Fig 5b |
| NADK | NAD | Fig 5b |
| NMNAT1 | NAD | Fig 5b |
| NADSYN1 | NAD | Fig 5b |
| NAMPT | NAD | Fig 5b |
| NNMT | NAD | Fig 5b |
| CD38 | NAD | Fig 5b |
| PRPS2 | Nucleotide (Purine) | Fig 5b |
| SLC1A4 | Small Molecule Transport (Alanine, Cysteine, Serine) | Fig 5b |
| SLC7A14 | Small Molecule Transport (Amino acid) | Fig 5b |
| SLC7A1 | Small Molecule Transport (Ar, Lys, Orn, His) | Fig 5b |
| SLC7A3 | Small Molecule Transport (Arg, Lys, Orn) | Fig 5b |
| SLC7A2 | Small Molecule Transport (Arg, Lys, Orn, His) | Fig 5b |
| SLC25A1 | Small Molecule Transport (Citrate, Malate) | Fig 5b |
| SLC13A5 | Small Molecule Transport (Citrate, Pyruvate) | Fig 5b |
| SLC6A8 | Small Molecule Transport (Creatine) | Fig 5b |
| SLC7A11 | Small Molecule Transport (Cystine, Glutamate) | Fig 5b |
| SLC37A4 | Small Molecule Transport (G6P) | Fig 5b |
| SLC6A6 | Small Molecule Transport (GABA,  β-alanine, taurine) | Fig 5b |
| SLC6A12 | Small Molecule Transport (GABA, Betaine | Fig 5b |
| SLC45A1 | Small Molecule Transport (Glucose) | Fig 5b |
| SLC2A6 | Small Molecule Transport (Glucose) | Fig 5b |
| SLC5A9 | Small Molecule Transport (Glucose) | Fig 5b |
| SLC2A10 | Small Molecule Transport (Glucose) | Fig 5b |
| SLC2A11 | Small Molecule Transport (Glucose, Fructose) | Fig 5b |
| SLC2A4 | Small Molecule Transport (Glucose, Glucosamine) | Fig 5b |
| SLC17A6 | Small Molecule Transport (Glutamate) | Fig 5b |
| SLC17A7 | Small Molecule Transport (Glutamate) | Fig 5b |
| SLC17A8 | Small Molecule Transport (Glutamate) | Fig 5b |
| SLC25A22 | Small Molecule Transport (Glutamate) | Fig 5b |
| SLC1A2 | Small Molecule Transport (Glutamate, Asparate) | Fig 5b |
| SLC1A3 | Small Molecule Transport (Glutamate, Asparate) | Fig 5b |
| SLC1A7 | Small Molecule Transport (Glutamate, Asparate) | Fig 5b |
| SLC25A12 | Small Molecule Transport (Glutamate, Aspartate) | Fig 5b |
| SLC6A5 | Small Molecule Transport (Glycine) | Fig 5b |
| SLC6A9 | Small Molecule Transport (Glycine, Sarcosine) | Fig 5b |
| SLC16A8 | Small Molecule Transport (Lactate) | Fig 5b |
| SLC16A6 | Small Molecule Transport (MCT activity) | Fig 5b |
| SLC16A13 | Small Molecule Transport (MCT activity) | Fig 5b |
| SLC16A5 | Small Molecule Transport (MCT activity) | Fig 5b |
| SLC16A4 | Small Molecule Transport (MCT activity) | Fig 5b |
| SLC25A30 | Small Molecule Transport (mitochondrial transport) | Fig 5b |
| SLC6A20 | Small Molecule Transport (Proline) | Fig 5b |
| SLC16A11 | Small Molecule Transport (Pyryuvate) | Fig 5b |
| SFXN3 | Small Molecule Transport (Serine) | Fig 5b |
| SFXN1 | Small Molecule Transport (Serine) | Fig 5b |
| SLC2A9 | Small Molecule Transport (Urate) | Fig 5b |
| G6PC2 | Sugar (gluconeogenesi) | Fig 5b |
| PCK2 | Sugar (gluconeogenesi) | Fig 5b |
| FBP1 | Sugar (gluconeogenesi) | Fig 5b |
| AGL | Sugar (Glycogen) | Fig 5b |
| PYGL | Sugar (Glycogen) | Fig 5b |
| GYG1 | Sugar (Glycogen) | Fig 5b |
| GAA | Sugar (Glycogen) | Fig 5b |
| GYG2 | Sugar (Glycogen) | Fig 5b |
| UGDH | Sugar (Nucleotide Sugar) | Fig 5b |
| TKTL1 | Sugar (PPP) | Fig 5b |
| RPE | Sugar (PPP) | Fig 5b |
| PGLS | Sugar (PPP) | Fig 5b |
| TALDO1 | Sugar (PPP) | Fig 5b |
| RPIA | Sugar (PPP) | Fig 5b |
| GOT1 | TCA Cycle | Fig 5b |
| MDH1B | TCA Cycle | Fig 5b |
| MDH1 | TCA Cycle | Fig 5b |
| PC | TCA Cycle | Fig 5b |
| PDHX | TCA Cycle | Fig 5b |
| PDHA1 | TCA Cycle | Fig 5b |
| OGDHL | TCA Cycle | Fig 5b |
| L2HGDH | TCA Cycle | Fig 5b |
| ME2 | TCA Cycle | Fig 5b |
| CS | TCA Cycle | Fig 5b |
| IDH3A | TCA Cycle | Fig 5b |
| ME1 | TCA Cycle | Fig 5b |
| SUCLA2 | TCA Cycle | Fig 5b |
| ACO2 | TCA Cycle | Fig 5b |
| FH | TCA Cycle | Fig 5b |
| D2HGDH | TCA Cycle | Fig 5b |
| ME3 | TCA Cycle | Fig 5b |
| SUCLG1 | TCA Cycle | Fig 5b |
| IDH1 | TCA Cycle | Fig 5b |
| ACO1 | TCA Cycle | Fig 5b |
| IDH2 | TCA Cycle | Fig 5b |
| SUCLG2 | TCA Cycle | Fig 5b |
| MTRR | Vitamin (B12) | Fig 5b |
| SHMT1 | Vitamin (Folate) | Fig 5b |
| MTHFR | Vitamin (Folate) | Fig 5b |
| MTHFS | Vitamin (Folate) | Fig 5b |
| MTHFD1 | Vitamin (Folate) | Fig 5b |
| MTHFD1L | Vitamin (Folate) | Fig 5b |
| MTHFD2 | Vitamin (Folate) | Fig 5b |

**Table S15. Key reagent and resources.**

| **Reagent or Media Components** | **Source** | **Identifier** |
| --- | --- | --- |
| DMEM, high glucose, GlutaMAX™ Supplement ^[1]^ | Thermo Fisher | 10566016 |
| DMEM/F-12, HEPES ^[1], [2]^ | Thermo Fisher | 11330032 |
| B-27™ Supplement (50X), serum free ^[1], [2], [3]^ | Thermo Fisher | 17504044 |
| MEM Non-Essential Amino Acids Solution (100X) ^[1], [2], [3], [4]^ | Thermo Fisher | 11140050 |
| Fetal Bovine Serum ^[1], [3], [4]^ | Atlanta Biologicals | S11550 |
| GlutaMAX™ Supplement ^[1]^ | Thermo Fisher | 35050061 |
| Bovine Serum Albumin solution ^[2]^ | Sigma-Aldrich | A1595-50ML |
| Taurine ^[2], [3], [4]^ | Sigma-Aldrich | T0625-10G |
| Advanced DMEM/F-12 ^[3]^ | Thermo Fisher | 12634010 |
| CTS™ (Cell Therapy Systems) N-2 Supplement ^[3]^ | Thermo Fisher | A1370701 |
| L-Glutamine ^[3]^ | Thermo Fisher | 25030081 |
| Minimum Essential Medium Eagle ^[4]^ | Sigma-Aldrich | M4526 |
| N1 Medium Supplement (100×) ^[4]^ | Sigma-Aldrich | N6530-5ML |
| Hydrocortisone ^[4]^ | Sigma-Aldrich | H0888-1G |
| 3, 3', 5-Triiodo-L-Thyronine ^[4]^ | Sigma-Aldrich | T6397-100MG |
| HBSS, no calcium, no magnesium | Thermo Fisher | 14170112 |
| Corning® Transwell® polyester membrane cell culture inserts | Sigma-Aldrich | CLS3470 |
| CO_2_ Independent Medium | Thermo Fisher | 18045088 |
| Neurobasal™-A Medium | Thermo Fisher | 10888022 |
| Biopsy Punch | Kai Medical | BP-50F, 5.0 mm |
| Corning® Transwell® polycarbonate membrane cell culture inserts | Sigma-Aldrich | CLS3421 |
| **Mass Spectrometry (GC MS)** | | |
| Water, Optima™ LC/MS Grade | Fisher Chemical | 166415 |
| Methanol | Fisher Chemical | 164905 |
| Methoxyamine hydrochloride | Sigma-Aldrich | 226904 |
| Pyridine | Sigma-Aldrich | 270970 |
| N-tert-Butyldimethylsilyl-N-methyltrifluoroacetamide | Sigma-Aldrich | 394882 |
| DB-5ms GC Column, 30 m, 0.25 mm, 0.25 um | Agilent Technologies | 1225532 |
| Gel Pump | Savant | GP110 |
| Speed vac Plus | Savant | SC110A |
| Centrifuge | Eppendorf | 5424 |
| Acquity UPLC BEH Amide 1.7 μm Vanguard pre-column 2.1 x 5mm column | Waters | 186004799 |
| Acquity UPLC BEH Amide 1.7 μm 2.1 x 50mm column | Waters | 186004800 |
| Acetonitrile, Optima™ LC/MS Grade | Fisher Chemical | 75-05-8 |
| Ammonium Acetate | Sigma-Aldrich | 431311 |
| Ammonium hydroxide 28 - 30% in water ACS | VWR | AC42330-5000 |
| **Animals** | | |
| C57BL/6J | Jackson Lab | 000664 |
| Nrl knockout mice | Anand Swaroop Lab |  |

[1] Retinal organoid culture medium

[2] Mouse retinal explant culture medium

[3] Human retinal explant culture medium

[4] Human RPE explant culture medium

**Table S16. The medium formulation for human RPE/choroid culture.**

| **Components** | **Concentration (mg/L)** | **mM** | **Category** | **Source** |
| --- | --- | --- | --- | --- |
| Glycine | 57.5 | 0.77 | Amino Acids | MEM alpha + NEAA |
| L-Alanine | 33.9 | 0.38 | Amino Acids | MEM alpha + NEAA |
| L-Arginine hydrochloride | 105.0 | 0.50 | Amino Acids | MEM alpha |
| L-Asparagine-H2O | 63.2 | 0.43 | Amino Acids | MEM alpha + NEAA |
| L-Aspartic acid | 43.3 | 0.33 | Amino Acids | MEM alpha + NEAA |
| L-Cysteine hydrochloride-H2O | 100.0 | 0.57 | Amino Acids | MEM alpha |
| L-Cystine 2HCl | 31.0 | 0.10 | Amino Acids | MEM alpha |
| L-Glutamic Acid | 89.8 | 0.61 | Amino Acids | MEM alpha + NEAA |
| L-Glutamine | 292.0 | 2.00 | Amino Acids | MEM alpha |
| L-Histidine hydrochloride-H2O | 42.0 | 0.20 | Amino Acids | MEM alpha |
| L-Isoleucine | 52.4 | 0.40 | Amino Acids | MEM alpha |
| L-Leucine | 52.0 | 0.40 | Amino Acids | MEM alpha |
| L-Lysine hydrochloride | 73.0 | 0.40 | Amino Acids | MEM alpha |
| L-Methionine | 15.0 | 0.10 | Amino Acids | MEM alpha |
| L-Phenylalanine | 32.0 | 0.19 | Amino Acids | MEM alpha |
| L-Proline | 51.5 | 0.45 | Amino Acids | MEM alpha + NEAA |
| L-Serine | 35.5 | 0.34 | Amino Acids | MEM alpha + NEAA |
| L-Threonine | 48.0 | 0.40 | Amino Acids | MEM alpha |
| L-Tryptophan | 10.0 | 0.05 | Amino Acids | MEM alpha |
| L-Tyrosine disodium salt | 52.0 | 0.20 | Amino Acids | MEM alpha |
| L-Valine | 46.0 | 0.39 | Amino Acids | MEM alpha |
| Ascorbic Acid | 50.0 | 0.28 | Vitamins | MEM alpha |
| Biotin | 1.0E-01 | 4.00E-04 | Vitamins | MEM alpha |
| Choline chloride | 1.0 | 7.10E-03 | Vitamins | MEM alpha |
| D-Calcium pantothenate | 1.0 | 2.10E-03 | Vitamins | MEM alpha |
| Folic Acid | 1.0 | 2.30E-03 | Vitamins | MEM alpha |
| Niacinamide | 1.0 | 8.20E-03 | Vitamins | MEM alpha |
| Pyridoxal hydrochloride | 1.0 | 4.90E-03 | Vitamins | MEM alpha |
| Riboflavin | 1.0E-01 | 3.00E-04 | Vitamins | MEM alpha |
| Thiamine hydrochloride | 1.0 | 3.00E-03 | Vitamins | MEM alpha |
| Vitamin B12 | 1.4 | 1.00E-03 | Vitamins | MEM alpha |
| i-Inositol | 2.0 | 1.11E-02 | Vitamins | MEM alpha |
| Calcium Chloride (CaCl2) (anhyd.) | 200.0 | 1.80 | Inorganic Salts | MEM alpha |
| Magnesium Sulfate (MgSO4) (anhyd.) | 97.7 | 0.81 | Inorganic Salts | MEM alpha |
| Potassium Chloride (KCl) | 400.0 | 5.33 | Inorganic Salts | MEM alpha |
| Sodium Bicarbonate (NaHCO3) | 2200.0 | 26.19 | Inorganic Salts | MEM alpha |
| Sodium Chloride (NaCl) | 6800.0 | 117.24 | Inorganic Salts | MEM alpha |
| Sodium Phosphate monobasic (NaH2PO4-H2O) | 140.0 | 1.01 | Inorganic Salts | MEM alpha |
| D-Glucose (Dextrose) | 1000.0 | 5.56 | Other components | MEM alpha |
| Lipoic Acid | 0.2 | 1.00E-03 | Other components | MEM alpha |
| Phenol Red | 10.0 | 0.03 | Other components | MEM alpha |
| Sodium Pyruvate | 110.0 | 1.00 | Other components | MEM alpha |
| Recombinant human insulin | 5.0 |  | Other components | N1 Supplement |
| human transferrin | 5.0 |  | Other components | N1 Supplement |
| Sodium selenite | 5.0E-03 |  | Other components | N1 Supplement |
| Putrescine | 16.0 |  | Other components | N1 Supplement |
| Progesterone | 7.3E-03 |  | Other components | N1 Supplement |
| Taurine | 250.0 |  |  | Supplement |
| Hydrocortisone | 2.0E-02 |  |  | Supplement |
| triiodo-thyronine | 1.3E-05 |  |  | Supplement |
| Fetal Bovine Serum | 1% (V/V) |  |  | Supplement |
